# A description of the genus *Denitromonas* nom. rev.: *Denitromonas iodatirespirans* sp. nov. a novel iodate-reducing bacterium and two novel perchlorate-reducing bacteria *Denitromonas halophila* and *Denitromonas ohlonensis* isolated from San Francisco Bay intertidal mudflats

**DOI:** 10.1101/2022.10.10.511631

**Authors:** Victor M. Reyes-Umana, John D. Coates

**Author notes:** **Identifiers:** *Denitromonas iodatirespirans* IR12 (genome): GCA_018524365.1 (16S sequence): MW380749.3 *Denitromonas halophila* SFB-3 (genome): GCA_007625155.1 (16S sequence): KP137428.1 *Denitromonas ohlonensis* SFB-1 (genome): GCA_007625205.1 (16S sequence): KP137426.2 *Denitromonas ohlonensis* SFB-2 (genome): GCA_007625125.1 (16S sequence): KP137427.1. **Corresponding author**, 241 Koshland Hall, University of California, Berkeley Ca 94720.

## Abstract

The genus *Denitromonas* is currently a non-validated taxon that has been identified in several recent publications as members of microbial communities arising from marine environments. Very little is known about the biology of *Denitromonas* spp., and no pure cultures are presently found in any culture collections. The current epitaph of *Denitromonas* was given to the organism under the assumption that all members of this genus are denitrifying bacteria. This study performs phenotypic and genomic analyses on three new *Denitromonas* spp. isolated from tidal mudflats in the San Francisco Bay. We demonstrate that *Denitromonas* spp. are indeed all facultative denitrifying bacteria that utilize a variety of carbon sources such as acetate, lactate, and succinate. In addition, individual strains also use the esoteric electron acceptors perchlorate, chlorate, and iodate. Both 16S and Rps/Rpl phylogenetic analyses place *Denitromonas* spp. as a deep branching clade in the family *Zoogloeaceae*, separate from either *Thauera* spp., *Azoarcus* spp., or *Aromatoleum* spp. Genome sequencing reveals a G+C content ranging from 63.72% to 66.54%, and genome sizes range between 4.39-5.18 Mb. Genes for salt tolerance and denitrification are distinguishing features that separate *Denitromonas* spp. from the closely related *Azoarcus* and *Aromatoleum* genera.

## Introduction

The family *Zoogloeaceae* includes an ecologically and metabolically diverse group of organisms involved in nitrogen cycling and consists of numerous diazotrophs and denitrifiers with different ecological niches(1). Among the several genera in the family, there exists a poorly defined boundary between genera *Azoarcus*(2), *Thauera*, the recently proposed genus *Aromatoleum*(3), and the proposed genus *Denitromonas*. *Azoarcus* commonly possess denitrification genes and persist in the environment as either endophytes or free-living microorganisms. Endophytic *Azoarcus* spp. such as strain BH72 exhibit exclusively endophytic lifestyles and have only been observed as part of the rhizosphere(4). These *Azoarcus* are commonly studied to understand endophytic nitrogen fixation and the evolution and regulation of their nitrogenase genes(5, 6). Free-living *Azoarcus* spp. are often found in BTEX (benzene, toluene, ethylbenzene, and xylene) contaminated water systems where they can degrade these toxic hydrocarbons anaerobically under denitrifying conditions(7, 8). The combination of different lifestyles, genome identity, and ecological niches has led some researchers to propose that some free-living *Azoarcus* be reclassified as *Aromatoleum*(3), *Cognatazoarcus*, or *Pseudazoarcus* (9).

Unlike *Azoarcus* or *Aromatoleum, Denitromonas* has only been described as constituents of marine-derived microbial communities. All characterized *Denitromonas* spp. are free living microorganisms isolated from saline environments, making salt tolerance yet another distinguishing feature between *Aromatoleum, Azoarcus*, and *Denitromonas* (Table S1). However, various descriptions of *Denitromonas* are limited to a 16S rRNA gene on NCBI. The lack of complete genomes and publicly available isolates makes distinguishing the ecological and phylogenetic patterns between *Azoarcus*, *Aromatoleum*, and *Denitromonas* challenging. S. Orla-Jensen first proposed the genus in 1909 because denitrification was a unique characteristic limited to this genus at the time(10, 11). A review of the literature since Jensen’s proposal identifies three separate studies characterizing five distinct *Denitromonas* isolates that focus primarily on their respiratory metabolisms. Yip and Gu characterize the halophilic isolate *Denitromonas indolicum* MPKc and its ability to degrade 2-methylindole; however, the strain is not deposited in any culture collections and information about its genome is missing(12). Of the four *Denitromonas spp*. with sequenced genomes, Carlstrom *et al*. briefly describe three isolates of *Denitromonas halophila* from an anaerobic marine microbial community that reduces perchlorate to chloride and couples its reduction to growth(13). Barnum *et al*. further describe a parasitic symbiosis between several chlorate respiring bacteria and *Denitromonas sp*. SFB-1, based on the competitive utilization of the chlorate intermediate formed during anaerobic perchlorate reduction by SFB-1 for growth(14). Similarly, Reyes-Umana *et al*. describe an isolate of *Denitromonas* capable of using iodate as a terminal electron acceptor and coupling its reduction to growth(15).

While several studies have identified organisms belonging to the genus *Denitromonas*, the genus itself has not validly been characterized, and thus has no nomenclature standing on the List of Prokaryotic names with Standing in Nomenclature (LSPN)(16, 17). This study seeks to validly publish the taxon *Denitromonas* nom. rev. and holistically demonstrate a clear distinction between the four sequenced *Denitromonas* isolates, the recently discovered *Nitrogeniibacter* spp., and other *Zoogloeaceae* through 16S ribosomal gene phylogeny, ribosomal protein phylogeny, average nucleotide identity, and biochemical assays. Additionally, while Orla-Jensen raised issues with the epitaph *Denitromonas* due to the ubiquity of denitrification among bacteria, we chose to maintain the original epitaph for consistency and historical recognition(10). We also demonstrate that *Denitromonas* spp. are free living, metabolically versatile, possess genes associated with salt tolerance, but do not have an absolute requirement for NaCl. These defining features of the genus likely explain why isolates arise from marine environments, and suggest that *Denitromonas* forms a new, possibly under-sampled, genus belonging to the family *Zoogloeaceae* that inhabits a distinct niche from *Aromatoleum* and *Azoarcus*.

## Methods

### Media, chemicals, and culture conditions

*Denitromonas* spp. were isolated from San Francisco Bay intertidal flat sediment as previously described(13). All isolates were confirmed by single colony selection on agar plates with subsequent sequencing of the 16S ribosomal gene (16S rRNA gene: see below). Isolates were grown routinely both aerobically or anaerobically at 30°C using either a defined marine media, or Reasoner’s 2A (R2A) agar or medium (HiMedia, USA). The defined marine media contains the following per liter: 30.8g NaCl, 1.0g NH_4_Cl, 0.77g KCl, 0.1g KH_2_PO_4_, 0.20g MgSO_4_·7H_2_O, 0.02g CaCl_2_·2 H_2_O, 7.16g HEPES, along with vitamin and mineral mixes as described in Carlstrom *et al*. (13). The anaerobic medium was boiled and dispensed under an atmosphere of 100% N_2_ and sealed with a butyl rubber stopper. After autoclaving, each liter of media was aseptically amended with 34.24mL 0.4M CaCl_2_ and 26.07mL 2M MgCl_2_·6H_2_O from sterile stock solutions. All chemicals used in this study used sodium salts and were purchased through Sigma Aldrich (Sigma Aldrich, USA). Thermo Scientific™ GENESYS™ 20 spectrophotometer was used to measure growth via optical density at 600 nm (OD_600_). The Analytical Profile Index (API) 20 NE system was used to determine enzymatic activities following the manufacturer’s instructions (BioMérieux, France). Carbon utilization profiles, pH and salinity optima were determined by monitoring OD_600_ increase using the TECAN Sunrise™ 96-well microplate reader. Temperature range was determined by growth of isolates on R2A agar plates at 4°C, 14°C, 20°C, 25°C, 30°C, and 37°C.

### Genome sequencing and assembly

Original methods for genome sequencing and assembly for *D. halophila* and *D. ohlonensis* can be found in Barnum *et al*. (14), while the methods for the *D. iodatirespirans* genome sequencing and assembly can be found in Reyes-Umana *et al*. (15). In brief: For *D. halophila* and *D. ohlonensis*, DNA library preparation and sequencing were performed by either the Adam Arkin Laboratory or the Vincent J. Coates Genomics Sequencing Laboratory at the California Institute of Quantitative Biosciences (QB3, Berkeley, CA) using either an Illumina MiSeq V2 (150PE or 250PE) and an Illumina Hiseq4000 (100PE), respectively. Paired-end reads from each sample were trimmed using Sickle v. 1.33 with default parameters(18), error-corrected using SGA v. 0.10.15(19), and assembled using MEGAHIT v. 1.1.2 with the parameters --no-mercy and --min-count 3(20). Library preparation and sequencing for *D. iodatirespirans* was done on an Illumina HiSeq4000 (150PE) by the Vincent J. Coates Genomics Sequencing Laboratory at the California Institute of Quantitative Biosciences (QB3, Berkeley, CA). Reads were trimmed using sickle 1.33 and genome was assembled using SPAdes 3.9.(18, 21). All draft genome sequences were deposited onto the NCBI database.

### Taxonomic assessment

Taxonomy assignments were made using (i) 16S rRNA gene sequence; (ii) alignment of the ribosomal marker proteins (Rps and Rpl)(22); (iii) multigene alignments using the Genome Taxonomy Database (GTDB)(23); and (iv) alignment fraction analysis of average nucleotide identity (ANI) (24). Primers 27F and 1492R were used to amplify the 16S rRNA gene in all *Denitromonas* isolates by PCR. Resulting products were sequenced by Sanger sequencing, and sequences were deposited on NCBI. 16S rRNA gene sequences belonging to genera in the families *Zoogloeaceae*, some belonging to *Rhodocyclaceae*, and *Denitromonas* (N=60) were downloaded from NCBI in the FASTA (.fna) file format and aligned using MUSCLE 3.8(25). An approximately maximum-likelihood phylogenetic tree was generated using FastTree, specifying 10,000 resamples and using the standard settings for all other options(26). The resultant tree was visualized in a ladderized format using the *ete3* toolkit(27).

Whole genomes belonging to the families *Zoogloeaceae* and *Rhodocyclaceae*, and the genus *Denitromonas* (N=60), were downloaded from NCBI as protein FASTA files (.faa) and were run using the following select modules from the phylogenomics workflow described by Graham *et al*.(28) (https://github.com/edgraham/PhylogenomicsWorkflow). All .faa files were concatenated into a single FASTA file, and the script *identifyHMM* was used at standard settings without the *--performProdigal* flag to identify the set of Ribosomal proteins (Rps) and Ribosomal Protein Large subunit (Rpl) markers from Hug *et al*. (29). The *identifyHMM* script produced individual lists of hits for each Rps or Rpl marker, and hits for each marker were then aligned using MUSCLE 3.8 with the *-maxiters 16* flag included. Alignments were trimmed using trimAl v1.3(30) with the *-automated 1* flag included, and the *concat* script that comes packaged with BinSanity v.0.5.3 was run with the minimum number of sequences to be concatenated set at 8. The phylogenetic tree was built using FastTree at standard settings while also including the *-gamma* and *-lg* flags. The tree was visualized using the *ete3* toolkit. In parallel, the Genome Taxonomy Database Toolkit (GTDB-Tk) was also used to search for the taxonomy of *Denitromonas*.

### Average nucleotide identity (ANI) and alignment fraction (AF) analysis

Whole genomes belonging to the family *Zoogloeaceae* and the genus *Denitromonas* were downloaded from NCBI (N=50) as nucleotide FASTA files (.fna) and analyzed with FastANI using the *“many to many”* option(31) (https://github.com/ParBLiSS/FastANI). The output file was then transformed into a pivot table using pandas 1.2, and setting the index, columns, and values to the query, reference, and ANI, respectively. The index and column were sorted using the tree order from the RpS/RpL phylogenetic analysis and displayed as a heatmap using the seaborn 0.11 package. Alignment fractions were calculated from the output by dividing the count of bidirectional mappings by the total number of mappings for any one pair. Genus boundaries were determined using the methods outlined by Barco *et al*., which uses alignment frequency and average nucleotide identity(24). Briefly, a non-parametric Wilcoxon test was used to determine the boundaries for the *Denitromonas* genus (p ≤ 0.01) by comparing type species belonging to *Denitromonas* (N=4) against non-type species (N=46). *Denitromonas iodatirespirans* was used as the reference strain, and strains exceeding both AF and ANI boundaries were classified as *Denitromonas*.

### Protein subfamily and Denitromonas core genome analysis

Protein subfamilies were identified using the *subfamilies.py* script from Méheust *et al*.(32) on the concatenated protein FASTA file (.faa) for the genomes belonging to the families *Zoogloeaceae* and *Rhodocyclaceae*, and the genus *Denitromonas*. This script uses MMSeqs2 to perform an *all vs. all* search using parameters set at an e-value: 0.001, sensitivity: 7.5, and cover: 0.5. A sequence similarity network was built based on the pairwise similarities, and the *greedy set-cover* algorithm from MMseqs2 was performed to define protein subclusters. The resultant clusters were exported as a tab-separated file listing all identified proteins and their associated subfamilies. Feature tables belonging to each respective genome were downloaded from NCBI as feature_table.txt files to associate locus tag, genome ID, and gene product information with the MMseqs2 output using pandas 1.2. The resultant dataframe was converted to a presence and absence matrix with values of either 1 (present) or 0 (absent) using the *groupby* function, transposed, and genomes were then sorted using the phylogenetic tree order from the RpS/RpL analysis. Clusters were subsequently ordered by the presence frequency in descending order, starting with conserved protein subfamilies found in all genomes on the left. A global *all vs all* Jaccard similarity score was calculated for each genome in the dataset using the *jaccard_score* function on scikitlearn 0.24 and setting the method to “micro”. Results of the presence/absence matrix and the Jaccard similarity index were displayed using seaborn 0.11. Protein subfamilies unique to all *Denitromonas* were identified by taking the inverse intersection of the *non-Denitromonas* subfamily dataframe and *Denitromonas* subfamily dataframe. A similar analysis was repeated for the subfamily dataframe containing *Nitrogeniibacter, Cognatazoarcus*, and *Denitromonas*. A similar analysis of the “shell” proteins (defined as present in all but one *Denitromonas* spp.) used a similar method. The Kyoto Encyclopedia of Genes and Genomes (KEGG) was used to annotate the *Denitromonas* genomes using the *BlastKOALA* tool, and denitrification genes were identified by inputting the resulting K-numbers table into the KEGG Mapper *Reconstruct* tool.

### Microscopy

A Meiji MT4200L compound microscope (Meiji Techno Co., Japan) set with a 10x ocular lens was used to determine cell size under the 100x objective. Ocular lens was set with a scale bar and was calibrated to a 45μM nylon fiber to determine size of bars in the eyepiece. Individual cells were measured, and size was reported as range for vegetative cells. Motility was determined by observing twitching or movement under the 100x objective.

## Results and Discussion

### Cultured *Denitromonas spp*. forms a phylogenetically distinct clade

A 16S rRNA gene phylogenetic analysis of *Zoogloeaceae* is shown in Figure 1A. To allow for a valid comparison, several members belonging to the closely related family *Rhodocyclaceae* are provided as an outgroup. Our analyses combined with the taxonomic reassignments by Huang *et al*. show that all genera in *Zoogloeaceae*, including *Denitromonas* spp. form distinct monophyletic clades(9). *Denitromonas* is most closely related to the recently described *Nitrogeniibacter mangrovi*., with 94.80% 16S rRNA gene sequence similarity between the type strains *D. iodatirespirans* and *N. mangrovi*. There is good bootstrap support (≥99%) separating the majority of *Denitromonas* spp. (except for the uncultured *Denitromonas* sp. clone DMSN28) from *Nitrogeniibacter* spp. (Figure 1A), suggesting that the four isolates described herein belong to a distinct monophyletic group within the family *Zoogloeaceae*. Furthermore, all *Denitromonas* spp. show greater than 97.07% intraspecies sequence identity in their 16S rRNA gene; thus, the branch containing *D. halophila*, *D. ohlonensis*, and *D. iodatirespirans* belongs to a monophyletic clade containing other *Denitromonas* spp. exclusively. *Denitromonas* spp. are more distantly related to the recently reclassified *Pseudazoarcus* spp. and *Pseudothauera* spp. along with the more clearly defined *Thauera* spp. and *Aromatoleum* spp. clades supporting prior observations that each of these genera form phylogenetically distinct monophyletic clades(3, 9).

**Figure 1:**
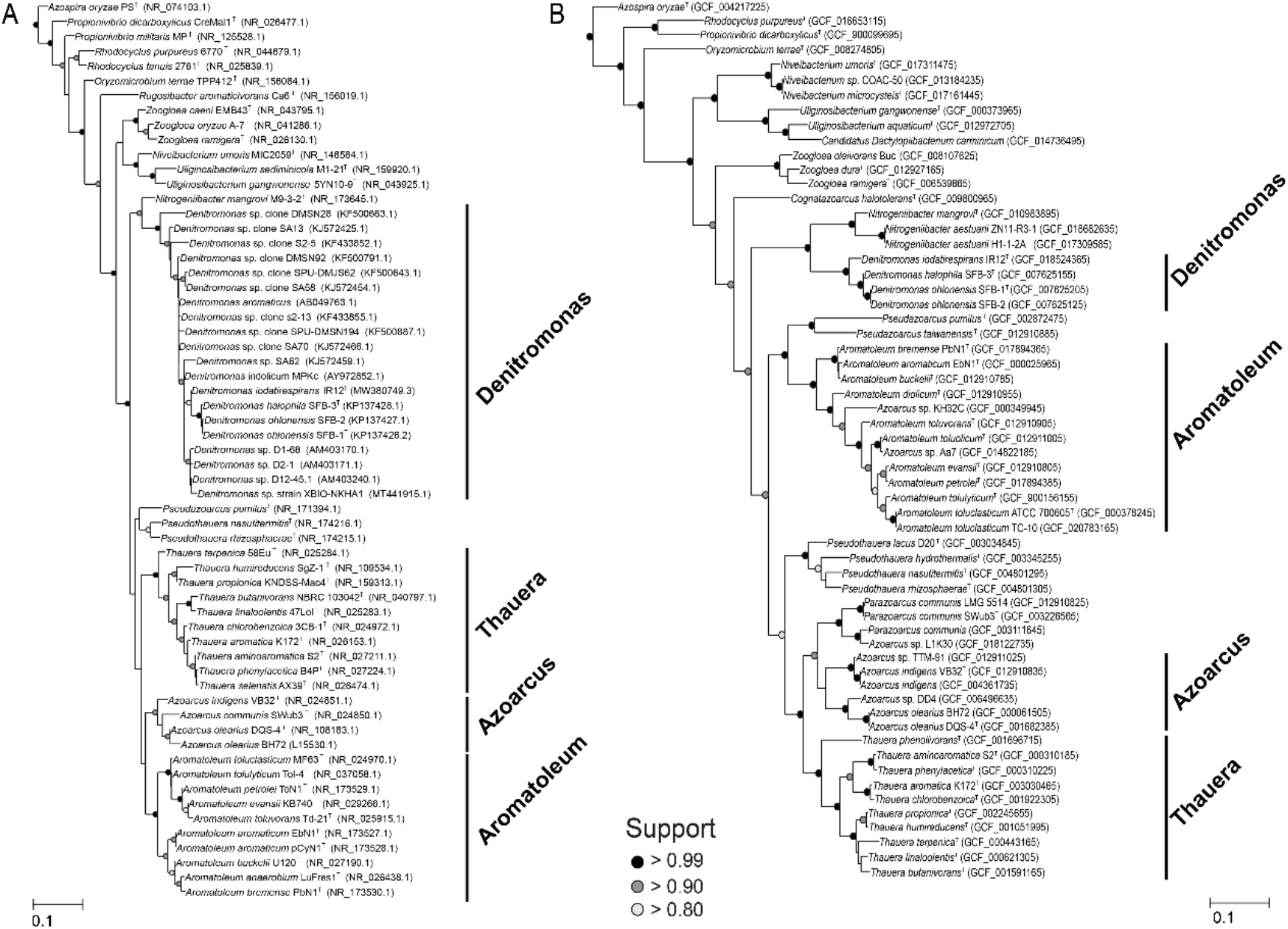
16S and ribosomal protein phylogeny. A) The 16S phylogeny and B) Ribosomal protein phylogeny of several *Zoogloeaceae* is summarized. Support values are signified by the color at each bifurcation.

While 16S rRNA gene phylogeny provides an overview of the diversity of *Zoogloeaceae*, current standards of prokaryotic classification recommend using the entire genome to determine reliable taxonomic assignment(33). Currently, whole genomes for the members belonging to the *Zoogloeaceae* family: *Azoarcus*, *Aromatoleum*, *Denitromonas*, *Dactylopiibacterium*, *Nitrogeniibacter*, *Pseudazoarcus*, *Pseudothauera, Parazoarcus, Thauera*, and *Uliginosibacterium* are available on NCBI. Figure 1B shows the concatenated ribosomal protein phylogeny of the different species within these genera. Using the *Rhodocyclaceae* bacteria *Azospira oryzae, Rhodocyclus* spp., *Oryzomicrobium terrae*, and *Niveibacterium* spp. as an outgroup, *Denitromonas spp*. form a distinct, monophyletic, deep branching clade that is separate from either *Azoarcus, Aromatoleum*, or *Nitrogeniibacter*. There is good bootstrap support (≥99%) for the distinction between *Denitromonas* spp. and *Aromatoleum* spp. consistent with distinct evolutionary histories for these two genera. Additionally, all *Zoogloeaceae* show less than 82.00% average nucleotide identity (ANI) to any *Denitromonas* spp. (Figure 2A, Table S6). While these observations suggest that *Denitromonas* spp. form a separate genus, the determination of a genus is fluid and varies among different bacterial families and orders. Recently, Barco *et al*. proposed a statistical test to discern the differences in the alignment fraction (AF) and average nucleotide identity between type and non-type strains of a genus to estimate a cutoff for a genus(24). Using this method, we calculated the genus cutoff for *Denitromonas* spp. to be at an ANI of 82.33% (p < 0.01) and an AF of 0.55 (p < 0.01). All fully sequenced *Denitromonas* currently fall above this threshold, suggesting that all belong to the same genus (Figure 2B). Crucially, the closely related *Nitrogeniibacter mangrovi* falls below the calculated cutoff with an ANI of 82.00% to the most closely related *Denitromonas* sp. (*D. iodatirespirans*) and an AF of 0.58, suggesting that *N. mangrovi* is correctly classified as a *Nitrogeniibacter* sp. Within the *Denitromonas* spp. clade, *D. ohlonensis* SFB-1 and *D. ohlonensis* SFB-2 show 99.99% ANI, suggesting that these are different strains of the same species. *D. halophila* shows 94% ANI with either *D. ohlonensis* isolates, suggesting that *D. halophila* is a different species. *D. iodatirespirans* shows the lowest ANI (85%) to other *Denitromonas* spp., further suggesting that the species diversity of the genus is likely under-sampled. Although the family *Zoogloeaceae* is a validly published taxon(34), a recent proposal to reduce polyphyly reclassifies the class *Betaproteobacteria* as an order of *Gammaproteobacteria* thus changing the higher taxonomic ranks of *Denitromonas, Azoarcus*, and *Aromatoleum*(35). The taxonomic ranks proposed by the Genome Taxonomy Database version 207 (http://gtdb.ecogenomic.org/) classifies the family *Zoogloeaceae* into the familiy *Rhodocyclaceae*, which belongs to the order *Burkholderiales* in the class *Gammaproteobacteria*. Within the family *Rhodocyclaceae*, *Denitromonas* is proposed as a distinct genus, further supporting our observations. Together, these data suggest that *Denitromonas* forms a distinct genus with a different evolutionary history from other members of the same family.

**Figure 2:**
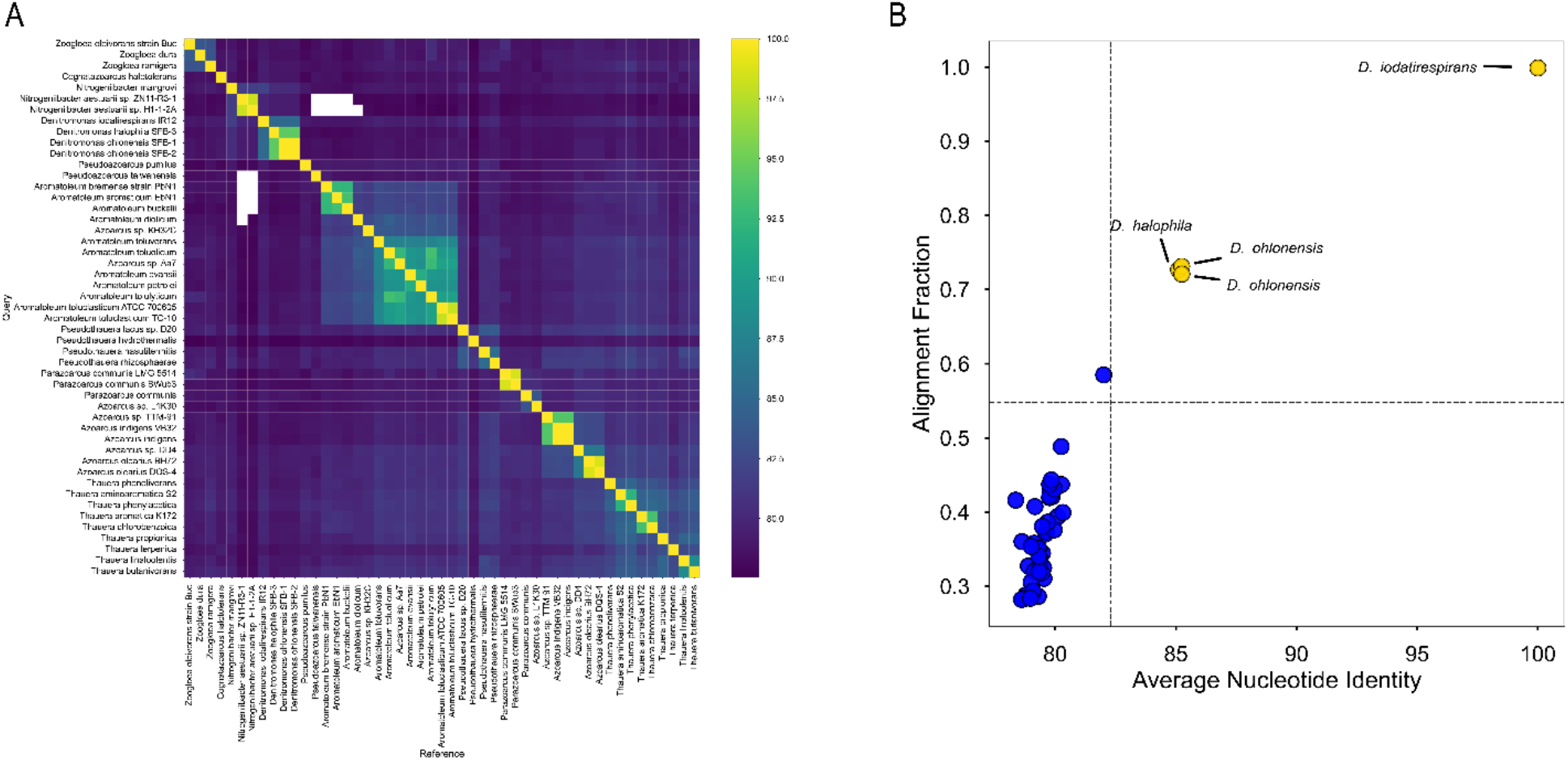
Average nucleotide identity and average nucleotide identity plotted against alignment fraction. A) The pairwise average nucleotide identity for organisms in the family *Zoogloeaceae*. Average nucleotide identity represented as a percentage. Squares in white denote an ANI below 78%. Outgroups are excluded due to an ANI below 78% B) The alignment fraction plotted against the average nucleotide identity is shown above. *Denitromonas iodatirespirans* is used as the reference strain. Yellow data points demonstrate the isolates that are confidently (Wilcoxon p ≤ 0.05) classified as *Denitromonas* whereas points in blue demonstrate related organisms that are significantly different from the type species. Significance is demonstrated by the dotted black lines where only strains in the top right quadrant are considered to belong the same genus.

### *Denitromonas spp*. possess protein subfamilies suited for marine environments

To further delineate possible functional differences between *Denitromonas spp*. and other *Zoogloeaceae*, an analysis of the proteome of *Zoogloeaceae* was performed. Additional genomes belonging to the family *Rhodocyclaceae* were included to serve as an outgroup for comparison. Proteins were clustered into 10,627 different subfamilies using MMSeqs2 and displayed as a presence-absence matrix (Table S2, Figure S1). The matrix was then scored by calculating the Jaccard similarity index on pairwise comparisons for each genome (Figure 3, Table S7). Figure 3 shows several distinct groupings within *Zoogloeaceae* in line with patterns observed in Figure 1 and Figure 2. *Pseudothauera* spp., *Parazoarcus* spp., *Azoarcus* spp., and *Thauera* spp. consistently show Jaccard scores ≥ 0.70 intergenera, suggesting that these genera share a common evolutionary history and lifestyle. Likewise, *Denitromonas* spp., *Nitrogeniibacter* spp., and *Cognatazoarcus halotolerans* show Jaccard scores ≥ 0.70 within these genera. We observe a difference in protein subfamily similarity between species related to *A. diolicum* and those belonging to the clade with *A. aromaticum*, indicating a possible difference in lifestyle within *Aromatoleum* spp. as previously reported(3). Likewise, *Denitromonas* spp. shows a different set of proteins that are internally similar within the genus but distinct from other genera. *Denitromonas* clearly forms a core cluster with similarity scores of 0.825 or above between all members of the genus (Figure 3). Similarity scores greater than 0.825 are in line with the similarity score between other genera in *Zoogloeaceae*, which all form similar clusters that track closely with taxonomy and ANI (Figure 1B, Figure 2A). *Cognatazoarcus halotolerans* shows relatively close Jaccard scores between 0.74 to 0.77 to *Denitromonas* spp., which is in line with the ribosomal protein phylogeny described earlier (Figure 1B). Similarly, *Nitrogeniibacter* spp. have Jaccard score ranges between 0.76 and 0.81, with the highest pairwise similarity of 0.81 between *Nitrogeniibacter mangrovi* and *Denitromonas ohlonensis* SFB-1. To understand the functional differences between *Denitromonas* spp. and other *Zoogloeaceae*, we analyzed the panproteome of the family *Zoogloeaceae* for protein subfamilies (i) present only in *Denitromonas*; (ii) present only in *Denitromonas, Nitrogeniibacter* spp., and *Cognatazoarcus*; and (iii) missing from *Denitromonas* (Tables S3, S4, and S5 respectively). Out of the 10,627 protein subfamilies, there are 132 subfamilies unique to *Denitromonas* and absent from all other *Zoogloeaceae* and *Rhodocyclaceae*, which accounts for 1.24% of the analyzed panproteome. Of these 132 protein subfamilies, 74/132 subfamilies (56%) are annotated as hypothetical proteins, and 28/132 subfamilies (21%) are present in all *Denitromonas* spp..

**Figure 3.**
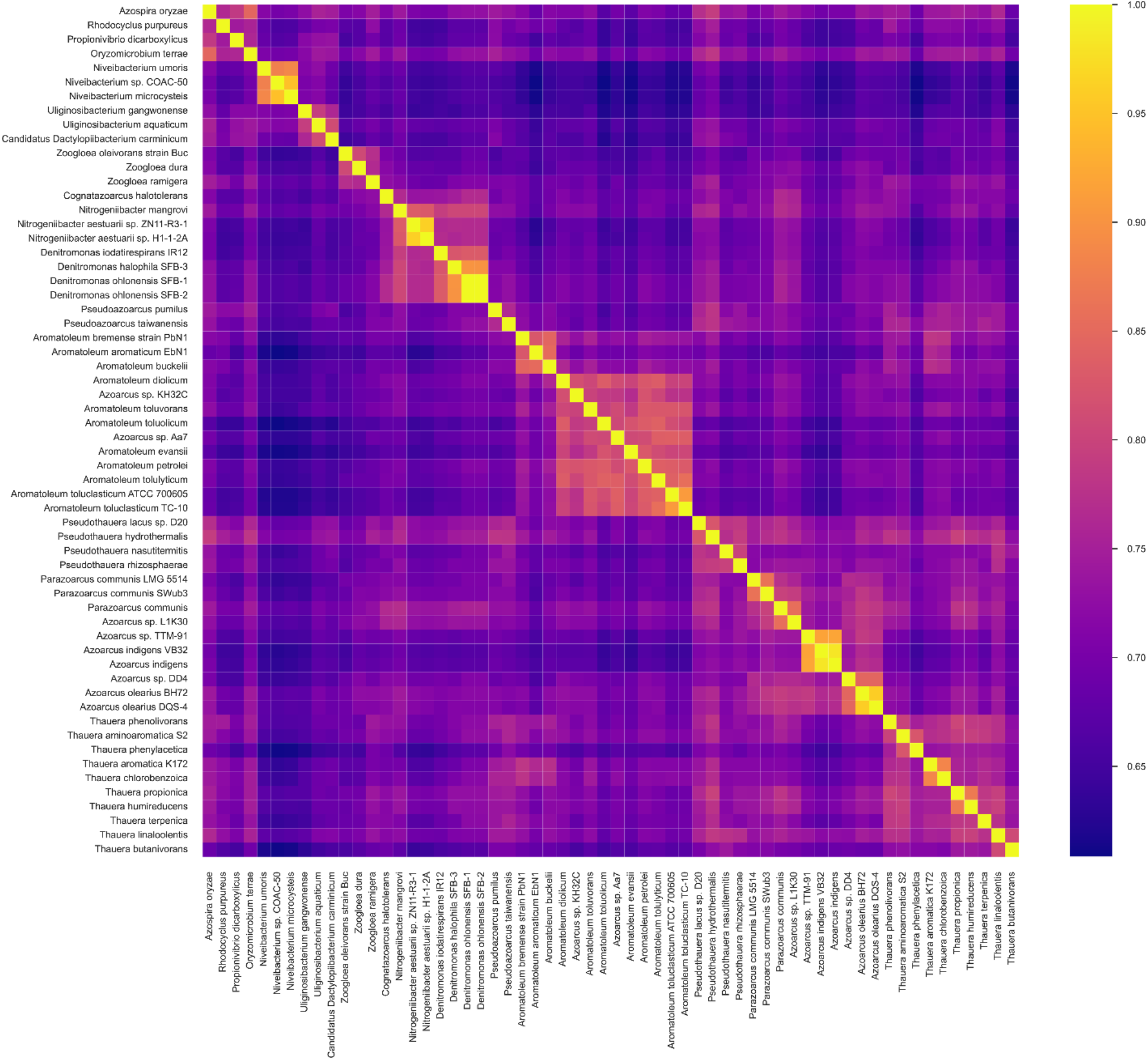
Quantification of the protein subfamily clustering using the Jaccard similarity index. Protein clusters for each genome were represented as a presence and absence matrix (Figure S1) and a pairwise comparison of each genome was performed using the Jaccard similarity index. Scores of 1.0 represents an identical set of protein subfamilies between any two given genomes.

*Denitromonas* spp. possess several protein subfamilies involved in metal homeostasis that are not found in other *Zoogloeaceae*. Two subfamilies found in all *Denitromonas* spp. include a FeoA ferrous iron transporter-associated protein and ChrE, which is involved in processing of chromium-glutathione complexes. FeoA proteins are cytoplasmically expressed and are essential in ferrous iron transport broadly; however, their exact function in iron transport in *Denitromonas* remains unknown(36) but may be related to an excess ferrous iron detoxification mechanism for nitrate reducing species as previously outlined by Carlson *et al*(37). ChrE is a rhodanese type enzyme that is alternatively annotated as a sulfotransferase. While its function has not been clearly defined, other enzymes of similar function show activity against seleno-glutathione complexes and are involved in selenate resistance. ChrE is believed to perform a similar function since chromate ions can also interact with glutathione, suggesting that ChrE may be involved in chromium resistance by cleaving chromium-glutathione complexes(38). Similarly, not much is known about the metal ABC transporter permeases found in *Denitromonas iodatirespirans*. Proteins belonging to this cluster are inconsistently annotated as *znuB* on NCBI, suggesting that this protein may be involved in zinc, manganese, or divalent metal metabolism. ZnuB homologs in other bacteria have been noted for their involvement in copper resistance or iron transport(39). Several proteins common to close relatives of *Denitromonas* are notably absent from *Aromatoleum* spp. and some *Azoarcus* spp. For instance, calcium proton antiporters are common in organisms like *Denitromonas* spp. and *Parazoarcus* spp. These proteins have been characterized in *E. coli* and *Azotobacter vinelandii*, where they have been shown to regulate calcium transport into and out of the cell and generate a proton motive force(40, 41). While the function of calcium in bacteria remains enigmatic, Ca^2+^/2H^+^ antiporters have been found in numerous bacteria from saline and alkaline environments, where they have been associated with salt tolerance(42, 43). Since *Azoarcus* and *Aromatoleum* are not exclusively found in marine environments, proteins belonging to these clusters likely demonstrate that *Denitromonas* spp. have adapted to mildly alkaline marine environments where they compete for the paucity of divalent metals(44, 45).

Organisms reclassified as *Aromatoleum* have been exclusively identified in freshwater systems(46), whereas *Denitromonas* spp. and *Azoarcus* spp. (e.g., *A. indigens* and *A. olearius* BH72) are consistently identified in marine environments or material sourced from marine environments (see table 1). Several genes are present in *Denitromonas* and *Azoarcus* but absent in *Aromatoleum* include the *nqr* operon (*nqrACFM*), the LEA type 2 proteins, and class GN sortases. The *nqr* operon is involved in maintaining a sodium motive force in numerous marine microorganisms and enables respiration in high salinity/high pH ecosystems(47, 48). *Denitromonas* spp. and *Cognatazoarcus halotolerans* have LEA type 2 proteins which play an essential role in abiotic stress caused by drought, cold, or salinity. The LEA type 2 proteins specifically contain a domain known as the water stress and hypersensitive response (WHy) domain which protects against dehydration(49). Lastly, both *Denitromonas* spp. and *Azoarcus* spp. possess the poorly described class GN sortase enzymes. Sortase enzymes are widespread in gram-positive bacteria and enable cell wall protein sorting; however, their role in gram-negative organisms remains unknown, and their distribution is limited to halotolerant proteobacteria(50).

**Table 1:** *Metabolic profiles of Denitromonas* spp.. + dictates ability to utilize substrate, - dictates an inability to utilize substrate. Only conditions with at least a single discrepancy are shown.

|  | *Denitromonas*<br>*iodatirespirans* IR-12 | *Denitromonas*<br>*ohlonensis* SFB-1 | *Denitromonas*<br>*halophila* SFB-3 |
| --- | --- | --- | --- |
|  | **Physical Characteristics** |  |  |
| Temperature (Range) | 14°C - 37°C | 14°C - 30°C | 14°C - 30°C |
| Salinity (Range) | 0-5% | 0-3% | 0-5% |
| Length (µM) | 1.5-2.0 | 1.0-1.5 | 1.5-2.0 |
| Genome size (Mb) | 5.18 | 4.42 | 4.39 |
| Mol% G+C | 66.54% | 64.05% | 63.72% |
|  | **Enzymatic Activity (72 Hours)** |  |  |
| Arginine Dihydrolase | + | - | + |
|  | **Carbon Assimilation (48 Hours)** |  |  |
| Yeast Extract (1%) | + | + | - |
|  | **Terminal Electron Acceptor Usage** |  |  |
| Oxygen | + | + | + |
| Nitrate | + | + | + |
| Perchlorate | - | + | + |
| Chlorate | - | + | + |
| Iodate | + | - | - |

*Denitromonas* spp., *Cognatazoarcus* spp, *Nitrogeniibacter* spp., and *Azoarcus* spp. have origins in saline or brackish environments(51), whereas *Aromatoleum* arise from a relatively new clade of organisms found in fresh contaminated water. Aside from *Aromatoleum*, organisms in the family *Zoogloeaceae* form a diverse clade of organisms that have adapted to survive in both saline and freshwater systems, while *Denitromonas* forms a clade of organisms that are predominantly found in brackish or saline environments. Interestingly, while some *Nitrogeniibacter* spp. have the absolute requirement for NaCl, *Denitromonas* spp. are able to grow in NaCl-free media such as R2A, suggesting a different lifestyle between these two closely related genera(51). Intertidal mudflats in estuarine environments like the San Francisco Bay are exposed to consistent changes in salinity and oxygen(52). The presence of genes associated with salt tolerance and metal transport, along with the ability to use numerous terminal electron acceptors, suggests that *Denitromonas* spp. have adapted to an environment with transient changes in salinity and nutrients. The metabolic diversity of *Denitromonas*, along with the dynamic environment they inhabit, suggests that this genus adopts a generalist, free-living lifestyle in brackish marine environments.

### Description of *Denitromonas* nom. rev

*Denitromonas* nom. rev. (De.ni.tro.mo’nas) L. prep. *de*, from, of; N.L. masc. n. *nitras* (gen. *nitratis*) nitrate; N.L. pref. *nitro-*, pertaining to nitrate; L. fem. n. *monas* a unit; N.L. fem. n. *Denitromonas* unit descending from nitrogen, which refers to the ability of all members from the genus to obtain energy through dissimilatory nitrate reduction to N_2_ gas.

Cells are rod-shaped, motile, mesophilic, and heterotrophic. All *Denitromonas* spp. are facultatively anaerobic chemoorganotrophs between 1.0-2.0 μM long by 0.5-1.0 μM wide. Cells grow in a wide range of salinity but optimally in brackish or saline water between 1% and 3%. Cells grow over a pH range of 6.8 to 8.2 with a growth optimum at pH 7.2. *Denitromonas* spp. grow over a wide temperature range (Table 1), but are routinely cultured at 30°C. The G+C content ranges from 63.72% to 66.54%, and genome sizes range between 4.39-5.18 Mb. *Denitromonas* spp. are metabolically versatile and utilize a variety of carbon sources including acetate, lactate, succinate, butyrate, fumarate, pyruvate, and propionate. However, *Denitromonas* spp. characterized to date are unable to utilize malate, formate, glycerol, or glucose as growth substrates. *Denitromonas* spp. do not ferment glucose, hydrolyze gelatin, or show beta-galactosidase activity; however, all strains are urease positive and can hydrolyze esculin. An analysis of the genome shows all the genes involved in denitrification and assimilatory nitrate reduction (NarGHI, NapAB, NirK, NorBC, and NosZ), and nitrogen fixation (nifDHK); thus, all *Denitromonas* are denitrifying bacteria that utilize nitrate as a terminal electron acceptor and produce N_2_ as an end product.

The type species for *Denitromonas* spp. is *Denitromonas iodatirespirans*.

### Description of *Denitromonas iodatirespirans* sp. nov

*Denitromonas iodatirespirans* (io.da.ti.res.pi’rans) N.L. masc. n. *iodas* (gen. *iodatis*), iodate; L. pres. part. *respirans*, breathing; N.L. part. adj. *iodatirespirans*, iodate-breathing, referencing the ability to use iodate as a terminal electron acceptor for growth.

*Denitromonas iodatirespirans* is a facultatively aerobic organoheterotroph isolated from an intertidal mudflat in the San Francisco Bay near Berkeley, CA. It grows by oxidizing lactate or acetate with concomitant reduction of oxygen (O_2_), nitrate (NO_3_^-^), or iodate (IO_3_^-^) in a defined marine medium at 3% salinity at a pH of 7.2. Alternatively, *D. iodatirespirans* can grow on R2A in either liquid or solid agar plates and can use a variety of carbon sources aerobically. On solid R2A agar, it forms small smooth reddish white colonies with a minute portion of crimson red and ash grey that deepen in color to a peach blossom red after 48-72 hours. Growth occurs between 14-37°C but is consistently grown at 30°C.

The type strain is *Denitromonas iodatirespirans* IR-12^T^ (=ATCC TSD-242^T^, = DSM 113304^T^), which has a 5,181,847 bp (average coverage 64.2x) genome with 4697 CDS, a G+C content of 66.54%, 57 tRNAs, one tmRNA, and one CRISPR array with associated Cas2-3, Cas 5d, and a Cas4-Cas1 fusion. It has a single plasmid 81,584 bp long whose function remains unclear. The 16S rRNA gene sequence (accession number: MW380749.3) and the complete genome has been deposited in GenBank (GCA_018524365.1), currently consisting of 202 contigs.

### Description of *Denitromonas halophila* sp. nov

*Denitromonas halophila* (ha.lo’phi.la.) Gr. masc. n. *hals* (gen. *halos*), salt; N.L. masc. adj.*philus* (from Gr. masc. adj. *philos*), loving; N.L. fem. adj. *halophila* salt-loving, referring to the organism’s ability to grow in saline environments

The description of *Denitromonas halophila* SFB-3 expands on observations from Carlstrom *et al*. and was amended to include details regarding its genome. *D. halophila* is a facultatively aerobic organoheterotroph isolated from an intertidal mudflat in the San Francisco Bay near Berkeley, CA(13). It grows by completely oxidizing lactate or acetate with concomitant reduction of oxygen (O_2_), nitrate (NO_3_^-^), perchlorate (ClO_4_^-^), or chlorate (ClO_3_^-^) in a defined marine medium at 3% salinity and a pH of 7.2. It forms small, round, rough, white colonies. Unlike other *Denitromonas* spp., *D. halophila* is unable to use 1% yeast extract for growth. Colony colors deepen to an orange-colored white colonies with a very small portion of tile red, king’s yellow, and ash grey on R2A agar after 48-72 hours.

The type strain is *Denitromonas halophila* SFB-3^T^ (=ATCC TSD-270^T^, =DSM 113305^T^) which has a 4,393,467 bp (average coverage 151.3x) genome with 4154 CDS, with 4,105 coding for proteins, 49 tRNAs, and no CRISPR array or plasmid. Its G+C content is 63.72%. The 16S rRNA gene sequence (accession number: KP137428.1) and complete genome has been deposited in GenBank (accession number: GCA_007625155.1), consisting of 34 contigs.

### Description of *Denitromonas ohlonensis* nom. nov

*Denitromonas ohlonensis* (oh.lo.nen’.sis) N. *ohlone* as it pertains to the occupied Ohlone land in contemporary Berkeley, California; L. nom. *ensis* belonging to; NL nom. *ohlonensis* belonging to the Ohlone land.

The description of *Denitromonas ohlonensis* expands from Barnum *et al*. and was amended to include details regarding its genome and the newly proposed nomenclature(14). *Denitromonas ohlonensis* SFB-1 is a facultatively aerobic organoheterotroph isolated from an intertidal mudflat in the San Francisco Bay near Berkeley, CA. It grows by oxidizing lactate or acetate with concomitant reduction of oxygen (O_2_), nitrate (NO_3_^-^), or perchlorate (ClO_4_^-^) in a defined marine medium at 3% salinity at a pH of 7.2. Alternatively, it grows in a defined marine medium at a salinity of 1% NaCl, pH of 7.2, and temperature of 30°C. On R2A it forms small smooth white colonies after 48-72 hours. The lack of arginine dihydrolase activity differentiates *D. ohlonensis* from other *Denitromonas* spp.

The species has two strains, the type strain is *Denitromonas ohlonensis* is SFB-1^T^ (=ATCC TSD-271^T^, =DSM 113310^T^). *Denitromonas ohlonensis* SFB-1 and SFB-2 was originally provided the epitaph of *Denitromonas halophilus* SFB-1 but the ANI between both species falls below the cutoff of 95%(53), resulting in homonymy; thus, reclassification is merited. *Denitromonas ohlonensis* SFB-2 is another strain belonging to this species. The genome of *D. ohlonensis* is 4,416,963 bp (average coverage of 129.6x) with 4,121 CDS with 4,111 coding for proteins. Its G+C content is 64.05% and has no CRISPR array or plasmid DNA. The 16S rRNA gene sequence (accession number: KP137426.2) and complete genome for SFB-1 has been deposited in GenBank (accession number: GCA_007625205.1), consisting of 40 contigs. Similarly, the 16S rRNA gene sequence (accession number: KP137427.1) and complete genome (accession number: GCA_007625125.1) has been deposited on GenBank.

## Author Contributions

Formal analysis, investigation, and visualizations were performed by VRU. Supervision was performed by JDC. Both VRU and JDC were involved in the conceptualization and writing of the manuscript.

## Conflicts of interest

The authors declare no conflicts of interest

## Funding information

Funding for research on iodate in the Coates lab was provided to VRU through the NSF GRFP Base Award: DGE1752814.

## Acknowledgements

The authors would like to thank Irania Alarcon for support with the microscopy work.

## Notes

### Competing Interest Statement

The authors have declared no competing interest.

