## Supplemental Data for "A description of the genus *Denitromonas* nom. rev.: *Denitromonas iodatirespirans* sp. nov. a novel iodate-reducing bacterium and two novel perchlorate-reducing bacteria *Denitromonas halophila* and *Denitromonas ohlonensis* isolated from San Francisco Bay intertidal mudflats"

**Supplemental Material**

Figure S1: Presence and absence of conserved proteins
Table S1: List of Denitromonas spp. 16S sequences
Table S2: Protein subfamilies of all genomes searched
Table S3: Protein subfamilies unique to Denitromonas spp.
Table S4: Protein subfamilies unique to close relatives of Denitromonas spp.
Table S5: Protein subfamilies missing from Denitromonas spp. and close relatives
Figure S6: Average nucleotide identities presented as percent pairwise identity
Figure S7: Jaccard similarity index scores

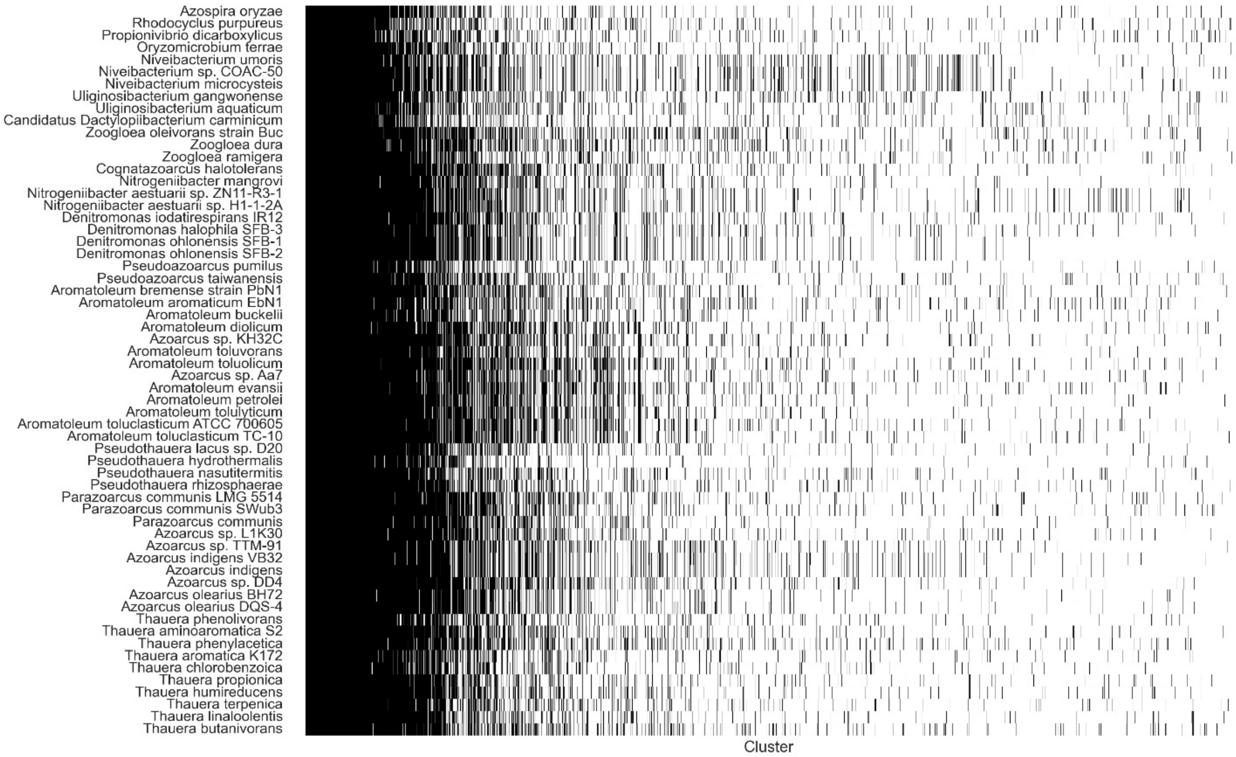


Figure S1: *Presence and absence of conserved proteins.* Proteins from all genomes were clustered using the MMSeqs2 algorithm and displayed as a presence and absence matrix. Black denotes presence, white denotes absence for any particular gene cluster. Clusters are ordered with core genes represented by the black squares on the left.

#### Table S1: List of Denitromonas spp. 16S sequences

| Name | Location | Associated Metabolism | Accession |
|---|---|---|---|
| *Denitromonas sp.* S2.5 | Kalo Vig (sediment) | Nitrate-reducing Fe (II)oxidizer | LN896468.1 |
| *Denitromonas sp. D2-1* | marine aquaculture | Nitrate-reducing | AM403171.1 |
| *Denitromonas sp. D12-45.1* | marine aquaculture | Nitrate-reducing | AM403240.1 |
| *Denitromonas sp. XBIO-NKHA1* | oil brine | n.d. | MT441915.1 |
| *Denitromonas sp. D1-68* | marine aquaculture | Nitrate-reducing | AM403170.1 |
| *Denitromonas sp. 37* | Wastewater treatment | n.d | DQ104209.1 |
| *Denitromonas indolicum strain MPKc 16S* | Inner Deep Bay (Hong Kong, Sediment) | 2-methylindole oxidation(11) | AY972852.1 |
| *Denitromonas aromaticus* | Activated sludge | n.d. | AB049763.1 |
| *Uncultured Denitromonas sp. clone strain ARE21* | Air Cathode Microbial Fuel Cell Reactor | n.d. | MN880253.1 |
| *Uncultured Denitromonas sp. isolate endo43* | Date palm endophyte | n.d. | KU579242.1 |
| *Uncultured Denitromonas sp. clone SA13* | Sludge | n.d. | KJ572425.1 |
| *Uncultured Denitromonas sp. clone SA58* | Sludge | n.d. | KJ572454.1 |
| *Uncultured Denitromonas sp. clone SA62* | Sludge | n.d. | KJ572459.1 |
| *Uncultured Denitromonas sp. clone SA70* | Sludge | n.d. | KJ572466.1 |
| *Uncultured Denitromonas sp.* | Microbial Fuel Cell Having Carbon | n.d. | KJ411935.1 |
| *Uncultured Denitromonas sp. clone SPU:DMJS40* | activated sludge | n.d. | KF500624 |
| *Uncultured Denitromonas sp. clone SPU:DMJS62* | activated sludge | n.d. | KF500643.1 |
| *Uncultured Denitromonas sp. clone SPU:DMJS65* | activated sludge | n.d. | KF500645.1 |
| *Uncultured Denitromonas sp. clone DMSN28* | activated sludge | n.d. | KF500663.1 |
| *Uncultured Denitromonas sp. clone SPU:DMSN194* | activated sludge | n.d. | KF500887.1 |
| *Uncultured Denitromonas sp. clone F5OHPNU07HZJ1B* | Gypsum-Treated Oil Sands Tailings Pond | n.d. | HQ067056.1 |
| *Uncultured Denitromonas sp. clone F5OHPNU07IF648* | Gypsum-Treated Oil Sands Tailings Pond | n.d. | HQ095246.1 |
| *Uncultured Denitromonas sp. clone F5OHPNU07H41ZX* | Gypsum-Treated Oil Sands Tailings Pond | n.d. | HQ095979.1 |
| *Uncultured Denitromonas sp. isolate DGGE gel band 3alfa* | Fish canning effluent | n.d. | HQ184392.1 |
| *Uncultured Denitromonas sp. isolate DGGE gel band 3beta 16S* | Fish canning effluent | n.d. | HQ184393.1 |

Table S2: Protein subfamilies of all genomes searched

| Cluster | Gene Count | Mode Gene Product Name |
|---|---|---|
| **00926** | 2750 | ABC transporter ATP-binding protein [Azoarcus communis] |
| **02985** | 1281 | SDR family oxidoreductase [Aromatoleum toluclasticum] |
| **02686** | 1206 | EAL domain-containing protein [Denitromonas halophilus] |
| **00755** | 1065 | sigma-54 dependent transcriptional regulator [Aromatoleum toluclasticum] |
| **05429** | 1061 | AMP-binding protein [Aromatoleum toluclasticum] |
| **03716** | 930 | ATP-binding protein [Nitrogeniibacter aestuarii] |
| **05818** | 873 | response regulator transcription factor [Azoarcus communis] |
| **01051** | 858 | enoyl-CoA hydratase-related protein [Aromatoleum toluclasticum] |
| **02507** | 795 | LysR family transcriptional regulator [Denitromonas halophilus] |
| **05502** | 777 | methyl-accepting chemotaxis protein [Azoarcus communis] |
| **00159** | 770 | acyl-CoA dehydrogenase family protein [Aromatoleum toluclasticum] |
| **07769** | 644 | aldehyde dehydrogenase family protein [Aromatoleum toluclasticum] |
| **02857** | 636 | response regulator transcription factor [Aromatoleum toluclasticum] |
| **04706** | 613 | efflux RND transporter permease subunit [Azoarcus communis] |
| **09257** | 599 | ABC transporter substrate-binding protein [Aromatoleum toluclasticum] |
| **09296** | 556 | LysR family transcriptional regulator [Denitromonas halophilus] |
| **01194** | 539 | PAS domain S-box protein [Denitromonas halophilus] |
| **01691** | 530 | PDR/VanB family oxidoreductase [Azoarcus indigens] |
| **04210** | 520 | molybdopterin-dependent oxidoreductase [Aromatoleum toluclasticum] |
| **00915** | 512 | TonB-dependent receptor [Thauera linaloolentis] |
| **00315** | 501 | acetyl-CoA C-acetyltransferase [Aromatoleum toluclasticum] |
| **03097** | 481 | TRAP transporter large permease [Azoarcus communis] |
| **09902** | 481 | ATP-binding cassette domain-containing protein [Azoarcus communis] |
| **04598** | 474 | ABC transporter ATP-binding protein/permease [Azoarcus indigens] |
| **07072** | 454 | LysR family transcriptional regulator [Denitromonas halophilus] |
| **07538** | 452 | branched-chain amino acid ABC transporter permease [Azoarcus indigens] |
| **02416** | 448 | DMT family transporter [Azoarcus communis] |
| **05102** | 446 | ATP-binding protein [Aromatoleum petrolei] |
| **06374** | 421 | PAS domain S-box protein [Nitrogeniibacter aestuarii] |
| **09125** | 395 | TRAP transporter substrate-binding protein [Propionivibrio dicarboxylicus] |
| **07864** | 391 | GDP-L-fucose synthase [Azoarcus olearius] |
| **03254** | 361 | branched-chain amino acid ABC transporter permease [Azoarcus communis] |
| **07670** | 359 | response regulator [Denitromonas halophilus] |
| **09991** | 358 | methyltransferase domain-containing protein [Denitromonas halophilus] |
| **00007** | 355 | heavy metal translocating P-type ATPase [Azoarcus communis] |
| **05442** | 348 | CBS domain-containing protein [Denitromonas halophilus] |
| **07943** | 345 | efflux RND transporter periplasmic adaptor subunit [Denitromonas halophilus] |
| **02421** | 344 | aspartate aminotransferase family protein [Azoarcus communis] |
| **09632** | 328 | DEAD/DEAH box helicase [Azoarcus communis] |
| **10504** | 325 | PLP-dependent aminotransferase family protein [Azoarcus indigens] |
| **04039** | 320 | alpha/beta hydrolase [Azoarcus communis] |
| **02593** | 316 | diguanylate cyclase [Aromatoleum toluclasticum] |
| **02800** | 312 | CoA transferase [Aromatoleum toluclasticum] |
| **09018** | 309 | ABC transporter permease [Azoarcus sp. L1K30] |
| **07099** | 304 | histidinol-phosphate transaminase [Azoarcus communis] |
| **00422** | 302 | ABC transporter ATP-binding protein [Aromatoleum toluclasticum] |
| **03704** | 301 | efflux transporter outer membrane subunit [Azoarcus indigens] |
| **05859** | 296 | glycosyltransferase [Nitrogeniibacter aestuarii] |
| **01713** | 294 | GspE/PulE family protein [Azoarcus communis] |
| **06722** | 293 | rhodanese-like domain-containing protein [Denitromonas halophilus] |
| **06154** | 290 | sensor domain-containing diguanylate cyclase [Azoarcus communis] |
| **07853** | 288 | ABC transporter permease [Azoarcus communis] |
| **00268** | 285 | maleylacetoacetate isomerase [Azoarcus communis] |
| **06609** | 281 | GntR family transcriptional regulator [Azoarcus communis] |
| **00096** | 277 | HDOD domain-containing protein [Azoarcus communis] |
| **07547** | 274 | efflux RND transporter periplasmic adaptor subunit [Denitromonas halophilus] |
| **05272** | 272 | electron transport complex subunit E [Azoarcus communis] |
| **04402** | 272 | nucleotidyltransferase family protein [Azoarcus communis] |
| **04113** | 262 | DUF1329 domain-containing protein [Aromatoleum toluclasticum] |
| **05716** | 262 | cation acetate symporter [Denitromonas halophilus] |
| **04013** | 258 | multidrug effflux MFS transporter [Azoarcus communis] |
| **03797** | 254 | biopolymer transporter ExbD [Azoarcus indigens] |
| **04173** | 253 | FAD-dependent oxidoreductase [Aromatoleum toluclasticum] |
| **03920** | 251 | cystathionine gamma-synthase family protein [Denitromonas halophilus] |
| **08247** | 251 | MULTISPECIES: YfhL family 4Fe-4S dicluster ferredoxin [Aromatoleum] |
| **05928** | 247 | Lrp/AsnC family transcriptional regulator [Denitromonas halophilus] |
| **01203** | 247 | GTP 3',8-cyclase MoaA [Denitromonas halophilus] |
| **00335** | 245 | YCF48-related protein [Aromatoleum toluclasticum] |
| **03178** | 245 | sigma-70 family RNA polymerase sigma factor [Thauera linaloolentis] |
| **02764** | 242 | peroxiredoxin [Denitromonas halophilus] |
| **02313** | 242 | cytochrome c oxidase accessory protein CcoG [Azoarcus communis] |
| **10508** | 242 | LysR family transcriptional regulator [Denitromonas halophilus] |
| **03807** | 242 | sulfoacetaldehyde acetyltransferase, partial [Denitromonas halophilus] |
| **02871** | 242 | HAD hydrolase-like protein [Azoarcus communis] |
| **04901** | 241 | 2-isopropylmalate synthase [Azoarcus communis] |
| **10260** | 234 | pseudouridine synthase [Azoarcus communis] |
| **03413** | 231 | MMPL family transporter [Aromatoleum toluclasticum] |
| **05915** | 231 | DegQ family serine endoprotease [Azoarcus indigens] |
| **08055** | 230 | cysteine desulfurase [Aromatoleum evansii] |
| **06620** | 228 | response regulator [Azoarcus communis] |
| **07031** | 228 | RluA family pseudouridine synthase [Azoarcus communis] |
| **03107** | 223 | OmpA family protein [Azoarcus communis] |
| **06650** | 218 | U32 family peptidase [Aromatoleum toluclasticum] |
| **09585** | 218 | TlpA family protein disulfide reductase [Aromatoleum toluclasticum] |
| **07827** | 217 | M48 family metalloprotease [Azoarcus communis] |
| **05663** | 217 | D-amino acid dehydrogenase [Denitromonas halophilus] |
| **03632** | 216 | bacteriohemerythrin [Azoarcus olearius] |
| **07143** | 215 | AMP-binding protein [Azoarcus indigens] |
| **06199** | 214 | LysE family translocator [Denitromonas halophilus] |
| **02652** | 210 | MULTISPECIES: integration host factor subunit alpha [Azoarcus] |
| **02668** | 209 | F0F1 ATP synthase subunit beta [Denitromonas halophilus] |
| **08886** | 208 | 4Fe-4S dicluster domain-containing protein [Aromatoleum toluclasticum] |

| | | |
|---|---|---|
| **03051** | 207 | monovalent cation/H+ antiporter subunit D family protein [Denitromonas halophilus] |
| **05400** | 207 | response regulator [Azoarcus communis] |
| **01425** | 205 | MBL fold metallo-hydrolase [Azoarcus communis] |
| **08008** | 204 | type II secretion system F family protein [Azoarcus communis] |
| **09197** | 202 | dihydrolipoyl dehydrogenase [Aromatoleum toluclasticum] |
| **02362** | 201 | PilT/PilU family type 4a pilus ATPase [Aromatoleum toluclasticum] |
| **03478** | 200 | MFS transporter [Aromatoleum toluclasticum] |
| **05523** | 200 | class I poly(R)-hydroxyalkanoic acid synthase [Azoarcus communis] |
| **05638** | 197 | MULTISPECIES: rod shape-determining protein [Aromatoleum] |
| **01760** | 197 | NAD(P)-dependent alcohol dehydrogenase [Nitrogeniibacter aestuarii] |
| **03326** | 197 | ParA family protein [Denitromonas halophilus] |
| **05150** | 196 | oxidoreductase [Denitromonas halophilus] |
| **07473** | 194 | FecR domain-containing protein [Thauera linaloolentis] |
| **02226** | 193 | ABC transporter permease [Denitromonas halophilus] |
| **08388** | 193 | DUF1302 domain-containing protein [Aromatoleum toluclasticum] |
| **00909** | 189 | 2-oxo acid dehydrogenase subunit E2 [Aromatoleum toluclasticum] |
| **02748** | 188 | porin [Aromatoleum toluclasticum] |
| **06821** | 186 | amino acid ABC transporter permease [Candidatus Dactylopiibacterium carminicum] |
| **01233** | 186 | universal stress protein [Aromatoleum toluclasticum] |
| **04111** | 186 | NADH-quinone oxidoreductase subunit M [Aromatoleum toluclasticum] |
| **07367** | 185 | NAD(P)-dependent oxidoreductase [Denitromonas halophilus] |
| **07578** | 185 | 3-hydroxybutyryl-CoA dehydrogenase [Aromatoleum toluclasticum] |
| **06778** | 184 | acetyl-CoA carboxylase biotin carboxylase subunit [Azoarcus communis] |
| **04777** | 183 | type I methionyl aminopeptidase [Azoarcus communis] |
| **05448** | 183 | protein phosphatase 2C domain-containing protein [Azoarcus olearius] |
| **00011** | 182 | FAD-binding protein [Azoarcus communis] |
| **00189** | 182 | AraC family transcriptional regulator [Aromatoleum toluclasticum] |
| **08471** | 182 | RNA polymerase sigma factor FliA [Azoarcus communis] |
| **07850** | 181 | glycosyltransferase [Azoarcus olearius] |
| **02866** | 180 | MerR family transcriptional regulator [Azoarcus olearius] |
| **09893** | 180 | isoleucine--tRNA ligase [Azoarcus communis] |
| **09074** | 177 | c-type cytochrome [Azoarcus communis] |
| **02908** | 177 | FAD-binding protein [Azoarcus indigens] |
| **01540** | 176 | TRAP transporter substrate-binding protein [Azoarcus indigens] |
| **00332** | 175 | LysR family transcriptional regulator [Azoarcus indigens] |
| **01443** | 175 | FAD-binding protein [Aromatoleum toluolicum] |
| **05269** | 172 | cation:proton antiporter [Azoarcus communis] |
| **04698** | 171 | TRAP transporter small permease [Azoarcus communis] |
| **04638** | 170 | chemotaxis response regulator protein-glutamate methylesterase [Azoarcus sp. KH32C] |
| **07833** | 170 | glycosyltransferase family 9 protein [Azoarcus indigens] |
| **10528** | 170 | 2-oxo-3-hexenedioate decarboxylase [Zoogloea dura] |
| **01782** | 169 | FAD-dependent oxidoreductase [Aromatoleum toluvorans] |
| **09726** | 169 | indolepyruvate ferredoxin oxidoreductase family protein [Denitromonas halophilus] |
| **07758** | 169 | MULTISPECIES: iron-sulfur cluster insertion protein ErpA [Thauera] |
| **03019** | 168 | EAL domain-containing protein [Aromatoleum toluclasticum] |
| **02062** | 168 | SulP family inorganic anion transporter [Denitromonas halophilus] |
| **05697** | 168 | MULTISPECIES: cold-shock protein [Thauera] |
| **07573** | 168 | transporter substrate-binding domain-containing protein [Azoarcus communis] |
| **08583** | 168 | UvrD-helicase domain-containing protein [Azoarcus communis] |
| **00871** | 167 | HIT family protein [Azoarcus communis] |
| **10496** | 167 | amidase [Azoarcus communis] |
| **01435** | 167 | electron transfer flavoprotein subunit beta/FixA family protein [Aromatoleum toluolicum] |
| **00073** | 167 | inositol monophosphatase [Azoarcus communis] |
| **07845** | 166 | elongation factor G [Denitromonas halophilus] |
| **04576** | 165 | sulfate ABC transporter substrate-binding protein [Azoarcus olearius] |
| **04969** | 165 | ATP-dependent Clp protease ATP-binding subunit ClpA [Azoarcus communis] |
| **02658** | 164 | Crp/Fnr family transcriptional regulator [Azoarcus communis] |
| **09034** | 163 | IclR family transcriptional regulator [Azoarcus communis] |
| **09598** | 161 | chemotaxis protein CheA [Azoarcus communis] |
| **02049** | 161 | nucleotide sugar dehydrogenase [Azoarcus communis] |
| **04353** | 160 | c-type cytochrome [Denitromonas halophilus] |
| **01450** | 159 | NAD(P)H-dependent oxidoreductase subunit E [Azoarcus communis] |
| **08997** | 159 | response regulator transcription factor [Azoarcus indigens] |
| **02852** | 159 | asparagine synthase (glutamine-hydrolyzing) [Nitrogeniibacter aestuarii] |
| **07706** | 158 | DMT family transporter [Azoarcus communis] |
| **00013** | 157 | ABC transporter substrate-binding protein [Denitromonas halophilus] |
| **01664** | 157 | cation diffusion facilitator family transporter [Azoarcus indigens] |
| **01470** | 156 | pirin family protein [Azoarcus communis] |
| **08737** | 156 | MotA/TolQ/ExbB proton channel family protein [Azoarcus indigens] |
| **06265** | 155 | MULTISPECIES: class 1 integron integrase IntI1 [Bacteria] |
| **09869** | 154 | GlxA family transcriptional regulator [Aromatoleum toluclasticum] |
| **07647** | 153 | SDR family oxidoreductase [Aromatoleum toluclasticum] |
| **00297** | 153 | nitronate monooxygenase [Aromatoleum toluclasticum] |
| **03863** | 151 | formyltetrahydrofolate deformylase [Azoarcus communis] |
| **07669** | 150 | ABC transporter permease [Aromatoleum toluclasticum] |
| **09673** | 150 | beta-ketoacyl-[acyl-carrier-protein] synthase family protein [Azoarcus indigens] |
| **08953** | 150 | enoyl-CoA hydratase/isomerase family protein [Denitromonas halophilus] |
| **09242** | 149 | sensor histidine kinase [Denitromonas halophilus] |
| **04177** | 149 | FtsX-like permease family protein [Azoarcus indigens] |
| **10318** | 148 | PEP-CTERM system TPR-repeat protein PrsT [Azoarcus communis] |
| **10197** | 147 | PLP-dependent cysteine synthase family protein [Denitromonas halophilus] |
| **00547** | 147 | fumarylacetoacetate hydrolase family protein [Aromatoleum toluclasticum] |
| **00675** | 146 | 8-oxoguanine deaminase [Azoarcus communis] |
| **04932** | 146 | serine/threonine protein kinase [Azoarcus communis] |
| **03070** | 145 | MULTISPECIES: P-II family nitrogen regulator [Aromatoleum] |
| **07160** | 144 | phosphoglucosamine mutase [Azoarcus communis] |
| **09267** | 143 | aromatic ring-hydroxylating dioxygenase subunit alpha [Azoarcus communis] |
| **05129** | 142 | TetR/AcrR family transcriptional regulator [Azoarcus communis] |
| **06976** | 142 | MFS transporter [Denitromonas halophilus] |
| **08533** | 142 | Rrf2 family transcriptional regulator [Denitromonas halophilus] |
| **07940** | 142 | cytochrome c [Denitromonas halophilus] |
| **10264** | 142 | helix-turn-helix domain-containing protein [Denitromonas halophilus] |
| **05089** | 142 | xanthine dehydrogenase family protein molybdopterin-binding subunit [Azoarcus indigens] |
| **01386** | 141 | MULTISPECIES: thioredoxin TrxA [Azoarcus] |

| | | |
|---|---|---|
| **06500** | 140 | methanol/ethanol family PQQ-dependent dehydrogenase [Denitromonas halophilus] |
| **01196** | 139 | thiolase family protein [Aromatoleum toluclasticum] |
| **04896** | 138 | aminodeoxychorismate synthase component I [Aromatoleum toluclasticum] |
| **02803** | 138 | hemerythrin domain-containing protein [Aromatoleum toluclasticum] |
| **06047** | 138 | TetR/AcrR family transcriptional regulator [Denitromonas halophilus] |
| **07911** | 137 | 3-hydroxyacyl-CoA dehydrogenase [Aromatoleum toluclasticum] |
| **05768** | 137 | ABC transporter permease [Denitromonas halophilus] |
| **01372** | 136 | sigma-70 family RNA polymerase sigma factor [Denitromonas halophilus] |
| **06498** | 136 | YceI family protein [Azoarcus communis] |
| **02154** | 136 | gamma carbonic anhydrase family protein [Denitromonas halophilus] |
| **03898** | 136 | cbb3-type cytochrome c oxidase subunit I [Denitromonas halophilus] |
| **08967** | 135 | D-2-hydroxyacid dehydrogenase [Denitromonas halophilus] |
| **08036** | 135 | RnfABCDGE type electron transport complex subunit D [Aromatoleum toluclasticum] |
| **05769** | 135 | MULTISPECIES: translation initiation factor IF-1 [Thauera] |
| **02150** | 134 | type I glyceraldehyde-3-phosphate dehydrogenase [Azoarcus communis] |
| **10285** | 133 | acyl-CoA thioesterase [Azoarcus indigens] |
| **00374** | 133 | ABC transporter substrate-binding protein [Azoarcus communis] |
| **05621** | 131 | DnaJ domain-containing protein [Denitromonas halophilus] |
| **00296** | 130 | aldo/keto reductase [Azoarcus indigens] |
| **07046** | 129 | HD-GYP domain-containing protein [Azoarcus communis] |
| **04320** | 129 | tRNA dihydrouridine synthase DusB [Azoarcus communis] |
| **03100** | 129 | uroporphyrinogen-III C-methyltransferase [Azoarcus communis] |
| **09441** | 129 | FadR family transcriptional regulator [Propionivibrio dicarboxylicus] |
| **05925** | 128 | patatin-like phospholipase family protein [Denitromonas halophilus] |
| **07316** | 128 | HAD-IB family hydrolase [Azoarcus communis] |
| **04703** | 128 | MBL fold metallo-hydrolase [Azoarcus communis] |
| **04184** | 128 | TonB-dependent receptor [Azoarcus indigens] |
| **07211** | 128 | branched-chain amino acid transaminase [Azoarcus communis] |
| **09351** | 127 | 6-hydroxycyclohex-1-ene-1-carbonyl-CoA dehydrogenase [Aromatoleum buckelii] |
| **00260** | 127 | MBL fold metallo-hydrolase [Azoarcus communis] |
| **08376** | 127 | FKBP-type peptidyl-prolyl cis-trans isomerase [Azoarcus communis] |
| **06998** | 127 | MULTISPECIES: TolC family protein [Rhodocyclales] |
| **07624** | 127 | selenide, water dikinase SelD [Azoarcus communis] |
| **02693** | 127 | diaminopimelate decarboxylase [Azoarcus communis] |
| **05054** | 126 | methylmalonyl-CoA mutase family protein [Aromatoleum toluclasticum] |
| **00797** | 126 | 3-isopropylmalate dehydratase large subunit [Azoarcus communis] |
| **03500** | 126 | TatD family hydrolase [Azoarcus communis] |
| **03660** | 126 | acyl-CoA carboxylase subunit beta [Azoarcus communis] |
| **06974** | 126 | 3-isopropylmalate dehydrogenase [Azoarcus communis] |
| **10036** | 125 | MULTISPECIES: flagellar basal-body rod protein FlgF [Thauera] |
| **05021** | 124 | alkyl hydroperoxide reductase subunit F [Azoarcus communis] |
| **00082** | 124 | diaminopropionate ammonia-lyase [Azoarcus communis] |
| **02159** | 124 | HlyD family type I secretion periplasmic adaptor subunit [Azoarcus communis] |
| **08475** | 124 | ATP-dependent DNA helicase RecG [Azoarcus communis] |
| **07037** | 123 | polyprenyl synthetase family protein [Denitromonas halophilus] |
| **01048** | 123 | protein-L-isoaspartate O-methyltransferase [Azoarcus communis] |
| **08731** | 123 | cytochrome c oxidase subunit 3 family protein [Denitromonas halophilus] |
| **01559** | 123 | ribose-phosphate pyrophosphokinase [Azoarcus communis] |
| **09292** | 122 | metalloprotease PmbA [Azoarcus communis] |
| **01460** | 122 | MULTISPECIES: acetylglutamate kinase [Azoarcus] |
| **03751** | 122 | glutathione S-transferase family protein [Denitromonas halophilus] |
| **08153** | 122 | response regulator transcription factor [Aromatoleum toluclasticum] |
| **07329** | 122 | MULTISPECIES: peptidylprolyl isomerase [Thauera] |
| **04415** | 122 | TolC family outer membrane protein [Azoarcus communis] |
| **04409** | 122 | Hsp20/alpha crystallin family protein [Aromatoleum toluclasticum] |
| **03176** | 121 | integrase arm-type DNA-binding domain-containing protein [Denitromonas halophilus] |
| **10366** | 121 | glycosyltransferase [Aromatoleum toluclasticum] |
| **06820** | 121 | oxygen-independent coproporphyrinogen III oxidase [Azoarcus communis] |
| **04862** | 121 | chemotaxis protein [Azoarcus communis] |
| **08701** | 121 | FtsH protease activity modulator HflK [Azoarcus communis] |
| **09880** | 121 | aspartate--tRNA ligase [Azoarcus communis] |
| **04230** | 121 | threonylcarbamoyl-AMP synthase [Aromatoleum toluclasticum] |
| **01329** | 121 | putative lipid II flippase FtsW [Azoarcus communis] |
| **00833** | 120 | PhoH family protein [Azoarcus communis] |
| **02335** | 120 | aspartate carbamoyltransferase catalytic subunit [Azoarcus communis] |
| **02082** | 120 | 3-deoxy-7-phosphoheptulonate synthase AroG [Azoarcus communis] |
| **08400** | 120 | DNA topoisomerase (ATP-hydrolyzing) subunit B [Azoarcus communis] |
| **02690** | 120 | SCO family protein [Aromatoleum toluclasticum] |
| **08359** | 120 | DNA gyrase subunit A [Azoarcus communis] |
| **00493** | 120 | presqualene diphosphate synthase HpnD [Azoarcus communis] |
| **01262** | 120 | FAD-dependent monooxygenase [Azoarcus communis] |
| **04164** | 120 | RNA polymerase-binding protein DksA [Azoarcus communis] |
| **06531** | 120 | MULTISPECIES: lysophospholipid acyltransferase family protein [Thauera] |
| **08389** | 120 | 16S rRNA (cytosine(967)-C(5))-methyltransferase RsmB [Azoarcus communis] |
| **09599** | 120 | ATP phosphoribosyltransferase regulatory subunit [Azoarcus communis] |
| **01063** | 120 | exodeoxyribonuclease III [Azoarcus communis] |
| **01637** | 120 | penicillin-binding protein 2 [Azoarcus communis] |
| **00241** | 120 | Maf family nucleotide pyrophosphatase [Azoarcus communis] |
| **00162** | 120 | D-alanyl-D-alanine carboxypeptidase [Azoarcus communis] |
| **09863** | 120 | signal recognition particle protein [Azoarcus communis] |
| **06591** | 120 | GTPase ObgE [Azoarcus communis] |
| **07766** | 120 | glutamate--tRNA ligase [Azoarcus communis] |
| **00094** | 119 | 30S ribosomal protein S12 methylthiotransferase RimO [Azoarcus communis] |
| **08872** | 119 | C40 family peptidase [Azoarcus communis] |
| **02148** | 119 | sulfite exporter TauE/SafE family protein [Azoarcus communis] |
| **00893** | 119 | peptide chain release factor 1 [Azoarcus communis] |
| **08558** | 119 | cell division protein FtsA [Azoarcus communis] |
| **01485** | 119 | MULTISPECIES: hypothetical protein [Niveibacterium] |
| **01387** | 119 | bifunctional (p)ppGpp synthetase/guanosine-3',5'-bis(diphosphate) 3'-pyrophosphohydrolase [Azoarcus communis] |
| **03544** | 119 | 23S rRNA (guanosine(2251)-2'-O)-methyltransferase RlmB [Azoarcus communis] |
| **00618** | 118 | MarR family transcriptional regulator [Aromatoleum toluclasticum] |
| **07364** | 118 | NnrS family protein [Denitromonas halophilus] |
| **07946** | 118 | UDP-N-acetylmuramate--L-alanine ligase [Azoarcus communis] |
| **06476** | 118 | flagellar motor stator protein MotA [Azoarcus communis] |

| | | |
|---|---|---|
| 05831 | 117 | formate dehydrogenase subunit gamma [Azoarcus indigens] |
| 10262 | 117 | SAM-dependent methyltransferase [Denitromonas halophilus] |
| 03743 | 117 | UbiD family decarboxylase [Aromatoleum toluclasticum] |
| 10531 | 117 | MULTISPECIES: BolA/IbaG family iron-sulfur metabolism protein [Thauera] |
| 03853 | 117 | efflux RND transporter periplasmic adaptor subunit [Aromatoleum toluclasticum] |
| 01198 | 117 | autotransporter assembly complex protein TamA [Denitromonas halophilus] |
| 05017 | 116 | MULTISPECIES: rubredoxin [Azoarcus] |
| 09300 | 116 | ATP-binding protein [Azoarcus olearius] |
| 01474 | 116 | histone deacetylase family protein [Azoarcus indigens] |
| 00010 | 116 | molybdopterin-synthase adenylyltransferase MoeB [Aromatoleum toluclasticum] |
| 02406 | 116 | UDP-N-acetylmuramoyl-L-alanine--D-glutamate ligase [Azoarcus communis] |
| 05532 | 116 | acetate--CoA ligase family protein [Aromatoleum toluclasticum] |
| 01538 | 115 | NADP-dependent malic enzyme [Azoarcus communis] |
| 06158 | 115 | CobD/CbiB family protein [Azoarcus communis] |
| 05163 | 115 | DNA repair protein RadC [Azoarcus communis] |
| 03211 | 115 | NAD(P)/FAD-dependent oxidoreductase [Azoarcus communis] |
| 09647 | 115 | HDOD domain-containing protein [Azoarcus communis] |
| 02846 | 115 | ABC transporter permease [Azoarcus communis] |
| 07056 | 115 | 16S rRNA (cytidine(1402)-2'-O)-methyltransferase [Azoarcus communis] |
| 03849 | 114 | transcription elongation factor GreA [Azoarcus communis] |
| 00698 | 114 | ABC transporter permease [Azoarcus communis] |
| 10302 | 114 | chemotaxis protein CheR [Azoarcus communis] |
| 05364 | 114 | class II aldolase/adducin family protein [Azoarcus indigens] |
| 02996 | 114 | RnfABCDGE type electron transport complex subunit B [Aromatoleum toluclasticum] |
| 03530 | 114 | phasin family protein [Thauera phenolivorans] |
| 04078 | 114 | MFS transporter [Azoarcus indigens] |
| 02307 | 113 | pyridoxal phosphate-dependent aminotransferase [Azoarcus communis] |
| 06347 | 113 | ammonium transporter [Azoarcus communis] |
| 07675 | 113 | phosphate ABC transporter permease PstA [Aromatoleum toluclasticum] |
| 04486 | 113 | UDP-2,3-diacylglucosamine diphosphatase [Azoarcus communis] |
| 08424 | 113 | MULTISPECIES: NADH-quinone oxidoreductase subunit B [Rhodocyclales] |
| 02245 | 113 | electron transport complex subunit RsxC [Aromatoleum toluclasticum] |
| 04163 | 113 | AarF/UbiB family protein [Aromatoleum toluclasticum] |
| 07889 | 113 | MFS transporter [Azoarcus communis] |
| 00514 | 113 | acyl-CoA thioesterase [Azoarcus communis] |
| 00925 | 112 | glutamate 5-kinase [Azoarcus communis] |
| 08669 | 112 | aspartate ammonia-lyase [Denitromonas halophilus] |
| 08295 | 112 | MULTISPECIES: glutathione S-transferase N-terminal domain-containing protein [Azoarcus] |
| 04624 | 112 | 4Fe-4S binding protein [Denitromonas halophilus] |
| 03154 | 112 | N-acetylmuramoyl-L-alanine amidase [Azoarcus communis] |
| 08307 | 111 | DegT/DnrJ/EryC1/StrS family aminotransferase [Azoarcus communis] |
| 01574 | 111 | fused MFS/spermidine synthase [Azoarcus indigens] |
| 04032 | 111 | branched-chain amino acid ABC transporter permease [Azoarcus indigens] |
| 04772 | 111 | cytochrome b/b6 domain-containing protein [Azoarcus olearius] |
| 00729 | 111 | RnfH family protein [Aromatoleum toluclasticum] |
| 07105 | 111 | DNA polymerase III subunit alpha [Azoarcus communis] |
| 02509 | 111 | ABC transporter substrate-binding protein [Azoarcus indigens] |
| 08168 | 111 | ABC transporter substrate-binding protein [Azoarcus indigens] |
| 02721 | 111 | mechanosensitive ion channel [Nitrogeniibacter aestuarii] |
| 00002 | 111 | 23S rRNA (adenine(2503)-C(2))-methyltransferase RlmN [Azoarcus communis] |
| 00875 | 110 | D-glycero-beta-D-manno-heptose 1-phosphate adenylyltransferase [Azoarcus communis] |
| 02144 | 110 | NAD(P)H-dependent oxidoreductase subunit E [Denitromonas halophilus] |
| 02465 | 110 | elongation factor G [Azoarcus communis] |
| 02084 | 110 | class I SAM-dependent methyltransferase [Denitromonas halophilus] |
| 08775 | 110 | 23S rRNA (uracil(1939)-C(5))-methyltransferase RlmD [Azoarcus communis] |
| 03024 | 110 | molybdopterin molybdotransferase MoeA [Azoarcus communis] |
| 05931 | 110 | c-type cytochrome [Azoarcus communis] |
| 03990 | 110 | (2Fe-2S)-binding protein [Aromatoleum toluclasticum] |
| 07147 | 110 | electron transport complex subunit RsxG [Azoarcus communis] |
| 02792 | 109 | helix-turn-helix domain-containing protein [Denitromonas halophilus] |
| 07891 | 109 | serine O-acetyltransferase [Azoarcus communis] |
| 01475 | 109 | methyl-accepting chemotaxis protein [Zoogloea dura] |
| 09705 | 109 | EAL domain-containing protein [Azoarcus communis] |
| 06035 | 109 | succinate dehydrogenase iron-sulfur subunit [Denitromonas halophilus] |
| 02096 | 109 | GTP-binding protein [Aromatoleum toluclasticum] |
| 03922 | 108 | peptide deformylase [Azoarcus communis] |
| 08853 | 108 | hypothetical protein [Aromatoleum toluclasticum] |
| 06852 | 108 | circularly permuted type 2 ATP-grasp protein [Azoarcus communis] |
| 08859 | 108 | type I restriction-modification system subunit M [Denitromonas halophilus] |
| 07688 | 108 | tripartite tricarboxylate transporter substrate binding protein [Propionivibrio dicarboxylicus] |
| 08364 | 108 | phospholipase D-like domain-containing protein [Nitrogeniibacter aestuarii] |
| 07009 | 108 | FAD-dependent oxidoreductase [Aromatoleum toluclasticum] |
| 10512 | 108 | nickel-dependent hydrogenase large subunit [Azoarcus communis] |
| 02312 | 108 | MULTISPECIES: non-heme iron oxygenase ferredoxin subunit [Rhodocyclales] |
| 09818 | 108 | AEC family transporter [Denitromonas halophilus] |
| 04128 | 107 | AAA family ATPase [Denitromonas halophilus] |
| 09620 | 107 | Glu/Leu/Phe/Val dehydrogenase [Azoarcus communis] |
| 01826 | 107 | AAA family ATPase [Denitromonas halophilus] |
| 05977 | 107 | serine hydroxymethyltransferase [Aromatoleum toluclasticum] |
| 08588 | 107 | tryptophan synthase subunit beta [Azoarcus communis] |
| 06885 | 106 | GntR family transcriptional regulator [Azoarcus communis] |
| 03746 | 106 | type II secretion system GspH family protein [Aromatoleum toluclasticum] |
| 06539 | 106 | iron ABC transporter permease [Azoarcus nasutitermitis] |
| 10101 | 106 | lytic murein transglycosylase B [Azoarcus communis] |
| 04637 | 106 | glycosyltransferase family 4 protein [Denitromonas halophilus] |
| 04310 | 106 | chain length determinant protein EpsF [Azoarcus communis] |
| 00619 | 105 | type VI secretion system tip protein VgrG [Azoarcus communis] |
| 06416 | 105 | NAD(P) transhydrogenase subunit alpha [Azoarcus communis] |
| 01363 | 105 | 2-hydroxychromene-2-carboxylate isomerase [Denitromonas halophilus] |
| 03889 | 105 | hemolysin family protein [Azoarcus communis] |
| 10022 | 105 | NAD-dependent deacylase [Azoarcus communis] |
| 00984 | 105 | cytochrome b [Denitromonas halophilus] |
| 02864 | 104 | flavin reductase family protein [Azoarcus communis] |
| 05635 | 104 | SDR family oxidoreductase [Aromatoleum toluclasticum] |

| | | |
|---|---|---|
| 02423 | 103 | outer membrane lipoprotein carrier protein LolA [Azoarcus communis] |
| 04109 | 103 | carboxymuconolactone decarboxylase family protein [Aromatoleum toluclasticum] |
| 08693 | 103 | putative toxin-antitoxin system toxin component, PIN family [Aromatoleum aromaticum] |
| 02614 | 103 | NAD(P)H-dependent oxidoreductase [Azoarcus indigens] |
| 04155 | 103 | LemA family protein [Azoarcus communis] |
| 03325 | 103 | TorF family putative porin [Aromatoleum toluclasticum] |
| 01852 | 103 | glycosyltransferase [Azoarcus indigens] |
| 03644 | 103 | YkgJ family cysteine cluster protein [Azoarcus communis] |
| 03401 | 103 | H-NS histone family protein [Azoarcus indigens] |
| 05011 | 103 | GMC family oxidoreductase [Azoarcus communis] |
| 05953 | 102 | lytic transglycosylase domain-containing protein [Denitromonas halophilus] |
| 03482 | 102 | CoA ester lyase [Denitromonas halophilus] |
| 08463 | 102 | chemotaxis protein CheW [Azoarcus indigens] |
| 00016 | 101 | alpha/beta hydrolase [Nitrogeniibacter aestuarii] |
| 00863 | 101 | general secretion pathway protein GspD [Denitromonas halophilus] |
| 01504 | 101 | MULTISPECIES: 3-hydroxyacyl-ACP dehydratase FabZ [Azoarcus] |
| 09706 | 101 | AEC family transporter [Denitromonas halophilus] |
| 04108 | 101 | M20 family metallopeptidase [Aromatoleum toluclasticum] |
| 04937 | 100 | ABC transporter substrate-binding protein [Azoarcus communis] |
| 08012 | 100 | PBP1A family penicillin-binding protein [Denitromonas halophilus] |
| 08756 | 100 | TonB-dependent receptor [Azoarcus communis] |
| 01378 | 100 | 3'-5' exonuclease [Azoarcus communis] |
| 06266 | 100 | NapC/NirT family cytochrome c [Denitromonas halophilus] |
| 00041 | 100 | (Fe-S)-binding protein [Aromatoleum toluclasticum] |
| 09696 | 99 | DUF4212 domain-containing protein [Denitromonas halophilus] |
| 04297 | 99 | helix-turn-helix transcriptional regulator [Azoarcus indigens] |
| 00854 | 99 | glycosyltransferase family 2 protein [Azoarcus communis] |
| 03503 | 99 | ubiquinol-cytochrome c reductase iron-sulfur subunit [Azoarcus communis] |
| 03103 | 99 | PaaI family thioesterase [Denitromonas halophilus] |
| 04875 | 99 | formate dehydrogenase accessory sulfurtransferase FdhD [Aromatoleum toluclasticum] |
| 09702 | 99 | electron transfer flavoprotein-ubiquinone oxidoreductase [Azoarcus communis] |
| 02124 | 98 | AsnC family transcriptional regulator [Denitromonas halophilus] |
| 06838 | 97 | alpha/beta hydrolase [Azoarcus communis] |
| 01151 | 97 | DUF2189 domain-containing protein [Azoarcus communis] |
| 00913 | 97 | TIGR03013 family PEP-CTERM/XrtA system glycosyltransferase [Azoarcus communis] |
| 00404 | 97 | aminodeoxychorismate/anthranilate synthase component II [Azoarcus communis] |
| 03713 | 97 | FHA domain-containing protein [Azoarcus communis] |
| 00696 | 96 | PHB depolymerase family esterase [Aromatoleum buckelii] |
| 08566 | 96 | YbhB/YbcL family Raf kinase inhibitor-like protein [Aromatoleum toluclasticum] |
| 01722 | 96 | cytochrome C oxidase subunit IV family protein [Denitromonas halophilus] |
| 08042 | 96 | flavodoxin family protein [Denitromonas halophilus] |
| 10097 | 96 | amino acid ABC transporter permease [Azoarcus olearius] |
| 09974 | 96 | pyruvate kinase [Aromatoleum toluclasticum] |
| 00181 | 96 | chaperonin GroEL [Azoarcus communis] |
| 01107 | 95 | glutathione peroxidase [Azoarcus communis] |
| 07346 | 94 | bacterioferritin [Azoarcus indigens] |
| 02004 | 94 | hydrogenase nickel incorporation protein HypB [Azoarcus communis] |
| 09409 | 94 | slipin family protein [Nitrogeniibacter aestuarii] |
| 02594 | 94 | PAS domain S-box protein [Azoarcus indigens] |
| 06257 | 94 | 2-oxoacid:ferredoxin oxidoreductase subunit beta [Aromatoleum toluclasticum] |
| 05505 | 93 | MFS transporter [Aromatoleum toluclasticum] |
| 10489 | 93 | MCE family protein [Aromatoleum toluclasticum] |
| 09329 | 93 | sulfate adenylyltransferase subunit CysN [Denitromonas halophilus] |
| 05558 | 93 | polysaccharide export protein EpsE [Azoarcus communis] |
| 09796 | 92 | cache domain-containing protein [Aromatoleum toluclasticum] |
| 01812 | 92 | MULTISPECIES: type I glutamate--ammonia ligase [Thauera] |
| 02878 | 92 | 5-(carboxyamino)imidazole ribonucleotide synthase [Azoarcus communis] |
| 05262 | 92 | MULTISPECIES: 2Fe-2S iron-sulfur cluster binding domain-containing protein [Azoarcus] |
| 04611 | 92 | dihydroxy-acid dehydratase [Propionivibrio dicarboxylicus] |
| 05349 | 92 | sulfate adenylyltransferase subunit CysD [Denitromonas halophilus] |
| 01373 | 92 | TRAP transporter permease [Denitromonas halophilus] |
| 03083 | 92 | dinitrogenase iron-molybdenum cofactor biosynthesis protein [Azoarcus communis] |
| 03324 | 91 | dihydroneopterin aldolase [Azoarcus communis] |
| 09304 | 91 | TRAP transporter small permease subunit [Azoarcus communis] |
| 09355 | 91 | DedA family protein [Nitrogeniibacter aestuarii] |
| 06825 | 91 | NarK/NasA family nitrate transporter [Aromatoleum toluclasticum] |
| 06953 | 91 | peptide-methionine (S)-S-oxide reductase MsrA [Denitromonas halophilus] |
| 10212 | 91 | CNNM domain-containing protein [Nitrogeniibacter aestuarii] |
| 08893 | 91 | SPOR domain-containing protein [Azoarcus communis] |
| 04285 | 90 | molybdopterin-guanine dinucleotide biosynthesis protein B [Azoarcus communis] |
| 00532 | 90 | nitrous oxide reductase accessory protein NosL [Azoarcus communis] |
| 06056 | 90 | MULTISPECIES: nitrogenase molybdenum-iron protein alpha chain [Azoarcus] |
| 05682 | 90 | MULTISPECIES: helix-turn-helix domain-containing protein [Proteobacteria] |
| 06796 | 90 | DNA topoisomerase III [Azoarcus communis] |
| 04303 | 89 | Fic family protein [Aromatoleum bremense] |
| 04314 | 89 | ferric reductase-like transmembrane domain-containing protein [Azoarcus communis] |
| 04273 | 89 | transketolase [Azoarcus communis] |
| 07076 | 89 | FAD-dependent oxidoreductase [Aromatoleum toluclasticum] |
| 09635 | 89 | FKBP-type peptidyl-prolyl cis-trans isomerase [Denitromonas halophilus] |
| 08790 | 89 | DUF485 domain-containing protein [Aromatoleum toluclasticum] |
| 08802 | 89 | adenosylcobalamin-dependent ribonucleoside-diphosphate reductase [Azoarcus communis] |
| 06631 | 89 | 30S ribosomal protein S4 [Denitromonas halophilus] |
| 03227 | 89 | cytochrome-c peroxidase [Azoarcus communis] |
| 09091 | 89 | DUF294 nucleotidyltransferase-like domain-containing protein [Azoarcus communis] |
| 07005 | 88 | patatin-like phospholipase family protein [Zoogloea oleivorans] |
| 02528 | 88 | carboxylating nicotinate-nucleotide diphosphorylase [Aromatoleum toluclasticum] |
| 03170 | 88 | hydantoinase/oxoprolinase family protein [Aromatoleum toluclasticum] |
| 03346 | 88 | cytochrome c [Azoarcus communis] |
| 08553 | 87 | asparaginase [Denitromonas halophilus] |
| 07080 | 87 | thioredoxin family protein [Denitromonas halophilus] |
| 02632 | 87 | ankyrin repeat domain-containing protein [Aromatoleum toluclasticum] |
| 00583 | 87 | nitrogenase iron-molybdenum cofactor biosynthesis protein NifN [Azoarcus communis] |
| 06375 | 87 | ABC transporter substrate-binding protein [Azoarcus communis] |
| 07299 | 87 | alpha-amylase family glycosyl hydrolase [Niveibacterium microcysteis] |

| | | |
|---|---|---|
| 07216 | 87 | class I SAM-dependent methyltransferase [Azoarcus communis] |
| 02999 | 87 | thiazole synthase [Denitromonas halophilus] |
| 09217 | 86 | RidA family protein [Azoarcus communis] |
| 05794 | 86 | GTP cyclohydrolase FolE2 [Azoarcus communis] |
| 01511 | 86 | inorganic phosphate transporter [Denitromonas halophilus] |
| 03652 | 86 | MULTISPECIES: TIGR00730 family Rossman fold protein [Aromatoleum] |
| 01623 | 86 | type II secretion system GspH family protein [Aromatoleum toluclasticum] |
| 02141 | 86 | TSUP family transporter [Denitromonas halophilus] |
| 03387 | 86 | sulfur carrier protein ThiS [Denitromonas halophilus] |
| 06817 | 86 | MULTISPECIES: bifunctional pyr operon transcriptional regulator/uracil phosphoribosyltransferase PyrR [Azoarcus] |
| 10203 | 85 | HlyD family efflux transporter periplasmic adaptor subunit [Azoarcus communis] |
| 09452 | 85 | cytochrome c oxidase subunit I [Aromatoleum toluclasticum] |
| 04093 | 85 | NADH-quinone oxidoreductase subunit D [Azoarcus communis] |
| 03542 | 85 | GTPase HflX [Azoarcus communis] |
| 01874 | 85 | GGDEF domain-containing protein [Azoarcus communis] |
| 08697 | 85 | heme-binding protein [Aromatoleum toluclasticum] |
| 08896 | 85 | shikimate dehydrogenase [Aromatoleum diolicum] |
| 04917 | 85 | dihydrodipicolinate synthase family protein [Propionivibrio dicarboxylicus] |
| 05903 | 85 | MaoC family dehydratase [Aromatoleum toluclasticum] |
| 03619 | 85 | TrkH family potassium uptake protein [Denitromonas halophilus] |
| 00124 | 85 | AAA family ATPase [Denitromonas halophilus] |
| 01070 | 85 | succinate dehydrogenase, cytochrome b556 subunit [Aromatoleum toluclasticum] |
| 09950 | 85 | dihydroorotase [Azoarcus communis] |
| 06748 | 85 | NADH-quinone oxidoreductase subunit NuoH [Aromatoleum toluclasticum] |
| 04132 | 84 | diacylglycerol kinase [Denitromonas halophilus] |
| 04488 | 84 | TAXI family TRAP transporter solute-binding subunit [Azoarcus communis] |
| 03737 | 84 | aspartate/tyrosine/aromatic aminotransferase [Aromatoleum toluclasticum] |
| 01843 | 84 | ATP-dependent zinc metalloprotease FtsH [Denitromonas halophilus] |
| 00960 | 84 | outer membrane protein transport protein [Zoogloea dura] |
| 00995 | 84 | UbiX family flavin prenyltransferase [Aromatoleum toluclasticum] |
| 00409 | 84 | polysaccharide deacetylase family protein [Denitromonas halophilus] |
| 03836 | 84 | fructose-bisphosphate aldolase class II [Azoarcus communis] |
| 07204 | 84 | MBL fold metallo-hydrolase [Azoarcus communis] |
| 07239 | 84 | succinate dehydrogenase, hydrophobic membrane anchor protein [Aromatoleum toluclasticum] |
| 02877 | 84 | isochorismatase family protein [Aromatoleum toluclasticum] |
| 05943 | 83 | carbohydrate kinase family protein [Azoarcus communis] |
| 02149 | 83 | hypoxanthine-guanine phosphoribosyltransferase [Azoarcus communis] |
| 03647 | 83 | aspartate/glutamate racemase family protein [Azoarcus communis] |
| 05342 | 83 | MULTISPECIES: IS21-like element helper ATPase IstB [Thauera] |
| 09122 | 83 | DsbC family protein [Aromatoleum aromaticum] |
| 08844 | 83 | alpha-hydroxy-acid oxidizing protein [Azoarcus communis] |
| 04780 | 82 | acetaldehyde dehydrogenase (acetylating) [Aromatoleum toluclasticum] |
| 03463 | 82 | alpha/beta fold hydrolase [Azoarcus communis] |
| 02112 | 82 | ABC transporter substrate-binding protein [Aromatoleum toluclasticum] |
| 03501 | 82 | MULTISPECIES: co-chaperone GroES [Rhodocyclales] |
| 03028 | 82 | DUF502 domain-containing protein [Azoarcus communis] |
| 05836 | 82 | AraC family transcriptional regulator [Azoarcus olearius] |
| 06504 | 82 | malonic semialdehyde reductase [Azoarcus indigens] |
| 07869 | 81 | (Fe-S)-binding protein [Azoarcus communis] |
| 08237 | 81 | DUF3473 domain-containing protein [Azoarcus communis] |
| 02774 | 81 | arylesterase [Azoarcus communis] |
| 05763 | 81 | bifunctional methylenetetrahydrofolate dehydrogenase/methenyltetrahydrofolate cyclohydrolase FolD [Azoarcus communis] |
| 04723 | 81 | hypothetical protein [Azoarcus communis] |
| 08283 | 81 | ribonuclease E activity regulator RraA [Azoarcus communis] |
| 10400 | 81 | MULTISPECIES: copper-binding protein [Zoogloeaceae] |
| 07894 | 81 | cyclopropane-fatty-acyl-phospholipid synthase family protein [Aromatoleum evansii] |
| 00036 | 80 | hypothetical protein [Denitromonas halophilus] |
| 02604 | 80 | DUF2062 domain-containing protein [Denitromonas halophilus] |
| 07525 | 80 | dicarboxylate/amino acid:cation symporter [Nitrogeniibacter aestuarii] |
| 02801 | 80 | site-specific integrase [Azoarcus indigens] |
| 06518 | 80 | ABC transporter permease [Azoarcus communis] |
| 10617 | 80 | MULTISPECIES: Crp/Fnr family transcriptional regulator [Niveibacterium] |
| 01529 | 80 | extracellular solute-binding protein [Nitrogeniibacter aestuarii] |
| 05353 | 79 | glycine zipper 2TM domain-containing protein [Denitromonas halophilus] |
| 10373 | 79 | sulfurtransferase [Azoarcus communis] |
| 06893 | 79 | acetate/propionate family kinase [Azoarcus communis] |
| 06982 | 79 | alpha-ketoacid dehydrogenase subunit beta [Aromatoleum toluclasticum] |
| 03186 | 79 | sigma 54-interacting transcriptional regulator [Aromatoleum toluclasticum] |
| 10411 | 79 | phosphopyruvate hydratase [Denitromonas halophilus] |
| 06403 | 79 | YeeE/YedE family protein [Azoarcus communis] |
| 08572 | 79 | nucleoside deaminase [Azoarcus communis] |
| 03248 | 79 | dTDP-4-dehydrorhamnose 3,5-epimerase [Azoarcus indigens] |
| 07416 | 79 | MULTISPECIES: sulfonamide-resistant dihydropteroate synthase Sul1 [Bacteria] |
| 06678 | 79 | thiamine pyrophosphate-dependent dehydrogenase E1 component subunit alpha [Azoarcus communis] |
| 02455 | 78 | 2-oxoacid:acceptor oxidoreductase subunit alpha [Aromatoleum toluclasticum] |
| 04547 | 78 | AAA family ATPase [Azoarcus communis] |
| 07988 | 78 | iron ABC transporter permease [Azoarcus indigens] |
| 01689 | 78 | DUF411 domain-containing protein [Denitromonas halophilus] |
| 06516 | 78 | lipoyl synthase [Aromatoleum toluclasticum] |
| 01824 | 78 | MULTISPECIES: elongation factor Tu [Azoarcus] |
| 09093 | 77 | LytTR family DNA-binding domain-containing protein [Niveibacterium umoris] |
| 08874 | 77 | DNA photolyase family protein [Nitrogeniibacter aestuarii] |
| 07788 | 77 | catechol 2,3-dioxygenase [Azoarcus indigens] |
| 02403 | 77 | TonB-dependent receptor [Azoarcus communis] |
| 09886 | 77 | phosphoglycerate kinase [Denitromonas halophilus] |
| 04237 | 77 | MULTISPECIES: NADH-quinone oxidoreductase subunit NuoE [Thauera] |
| 01180 | 77 | class 1 fructose-bisphosphatase [Azoarcus communis] |
| 02450 | 77 | S41 family peptidase [Azoarcus communis] |
| 08453 | 76 | MaoC family dehydratase [Denitromonas halophilus] |
| 09890 | 76 | lactoylglutathione lyase [Azoarcus communis] |
| 02104 | 76 | 7-carboxy-7-deazaguanine synthase QueE [Azoarcus communis] |
| 06673 | 76 | peptidylprolyl isomerase [Azoarcus indigens] |
| 08601 | 76 | phosphoribulokinase [Azoarcus communis] |
| 01625 | 76 | MULTISPECIES: mercury resistance system periplasmic binding protein MerP [Proteobacteria] |

| | | |
|---|---|---|
| 01967 | 76 | FAD-dependent oxidoreductase [Aromatoleum toluclasticum] |
| 09117 | 76 | tripartite tricarboxylate transporter permease [Propionivibrio dicarboxylicus] |
| 08526 | 75 | 2-dehydropantoate 2-reductase [Azoarcus indigens] |
| 07682 | 75 | recombination-associated protein RdgC [Azoarcus communis] |
| 00353 | 75 | transglutaminase family protein [Denitromonas halophilus] |
| 08146 | 75 | MULTISPECIES: bifunctional demethylmenaquinone methyltransferase/2-methoxy-6-polyprenyl-1,4-benzoquinol methylase UbiE [Aromatoleum] |
| 03884 | 75 | metal ABC transporter permease [Azoarcus olearius] |
| 07261 | 75 | chemotaxis protein CheD [Azoarcus indigens] |
| 04446 | 75 | methylenetetrahydrofolate reductase [NAD(P)H] [Azoarcus communis] |
| 09516 | 74 | histidine kinase [Azoarcus communis] |
| 06078 | 74 | carbon-nitrogen hydrolase family protein [Denitromonas halophilus] |
| 00244 | 74 | dTDP-4-dehydrorhamnose reductase [Azoarcus communis] |
| 08998 | 74 | D-glycero-beta-D-manno-heptose 1,7-bisphosphate 7-phosphatase [Azoarcus communis] |
| 06785 | 74 | pyruvate dehydrogenase (acetyl-transferring), homodimeric type [Denitromonas halophilus] |
| 08162 | 74 | transcriptional repressor LexA [Azoarcus communis] |
| 00184 | 74 | methylated-DNA--[protein]-cysteine S-methyltransferase [Denitromonas halophilus] |
| 09816 | 74 | DUF2889 domain-containing protein [Aromatoleum toluclasticum] |
| 02042 | 73 | Crp/Fnr family transcriptional regulator [Azoarcus communis] |
| 07237 | 73 | mannose-1-phosphate guanylyltransferase/mannose-6-phosphate isomerase [Aromatoleum toluclasticum] |
| 01798 | 73 | multifunctional CCA addition/repair protein [Azoarcus communis] |
| 10445 | 73 | heme lyase CcmF/NrfE family subunit [Azoarcus communis] |
| 08556 | 73 | VOC family protein [Denitromonas halophilus] |
| 04123 | 73 | Y-family DNA polymerase [Denitromonas halophilus] |
| 05325 | 73 | ethanolamine utilization protein [Azoarcus communis] |
| 09108 | 73 | 2-methylcitrate synthase [Azoarcus communis] |
| 08846 | 73 | ABC transporter permease [Azoarcus communis] |
| 01436 | 73 | glycine cleavage system protein GcvH [Aromatoleum toluclasticum] |
| 06140 | 73 | ShlB/FhaC/HecB family hemolysin secretion/activation protein [Azoarcus indigens] |
| 05917 | 73 | DUF3422 domain-containing protein [Denitromonas halophilus] |
| 01459 | 73 | preprotein translocase subunit SecA [Azoarcus communis] |
| 08636 | 72 | YifB family Mg chelatase-like AAA ATPase [Azoarcus communis] |
| 04150 | 72 | MULTISPECIES: superoxide dismutase [Thauera] |
| 05653 | 72 | 39S ribosomal protein L45 [Aromatoleum buckelii] |
| 00212 | 72 | 4-oxalomesaconate tautomerase [Denitromonas halophilus] |
| 00321 | 72 | ADP-forming succinate--CoA ligase subunit beta [Azoarcus communis] |
| 00620 | 72 | ABC transporter substrate-binding protein [Azoarcus indigens] |
| 08442 | 72 | chromate transporter [Aromatoleum diolicum] |
| 09138 | 72 | copper chaperone PCu(A)C [Denitromonas halophilus] |
| 06806 | 72 | ACR3 family arsenite efflux transporter [Azoarcus communis] |
| 03972 | 72 | MULTISPECIES: integration host factor subunit beta [Rhodocyclales] |
| 01547 | 72 | F0F1 ATP synthase subunit A [Denitromonas halophilus] |
| 06082 | 72 | arsenate reductase ArsC [Azoarcus communis] |
| 02184 | 72 | MULTISPECIES: cytochrome-c oxidase, cbb3-type subunit II [Thauera] |
| 00699 | 72 | LacI family transcriptional regulator [Uliginosibacterium gangwonense] |
| 01732 | 72 | aminomethyltransferase family protein [Azoarcus communis] |
| 00014 | 72 | c-type cytochrome [Nitrogeniibacter aestuarii] |
| 01582 | 71 | acyl-CoA dehydratase activase [Aromatoleum toluclasticum] |
| 08238 | 71 | IS3 family transposase [Zoogloea oleivorans] |
| 06919 | 71 | fumarate hydratase [Azoarcus communis] |
| 07133 | 71 | heme exporter protein CcmD [Aromatoleum toluclasticum] |
| 05683 | 71 | AmmeMemoRadiSam system radical SAM enzyme [Azoarcus communis] |
| 06004 | 71 | low molecular weight phosphotyrosine protein phosphatase [Azoarcus communis] |
| 07116 | 71 | DUF4118 domain-containing protein [Azoarcus indigens] |
| 03839 | 71 | peptide-methionine (R)-S-oxide reductase MsrB [Denitromonas halophilus] |
| 00202 | 71 | hypothetical protein [Niveibacterium umoris] |
| 10396 | 71 | IS5 family transposase [Zoogloea oleivorans] |
| 09662 | 71 | carbonate dehydratase [Azoarcus communis] |
| 00989 | 71 | TOBE domain-containing protein [Azoarcus communis] |
| 00867 | 71 | ParB/RepB/Spo0J family partition protein [Aromatoleum aromaticum] |
| 06839 | 71 | DEAD/DEAH box helicase family protein [Thauera phenylacetica] |
| 04551 | 71 | 4-oxalocrotonate tautomerase family protein [Nitrogeniibacter aestuarii] |
| 05533 | 71 | aromatic/alkene/methane monooxygenase hydroxylase/oxygenase subunit alpha [Aromatoleum toluclasticum] |
| 09319 | 71 | ACP S-malonyltransferase [Azoarcus communis] |
| 06550 | 70 | nitrite/sulfite reductase [Azoarcus communis] |
| 07575 | 70 | NADH-quinone oxidoreductase subunit K [Azoarcus communis] |
| 08065 | 70 | DUF1840 domain-containing protein [Denitromonas halophilus] |
| 04196 | 70 | DUF3108 domain-containing protein [Aromatoleum toluclasticum] |
| 02187 | 70 | magnesium transporter [Denitromonas halophilus] |
| 07087 | 70 | endonuclease/exonuclease/phosphatase family protein [Azoarcus indigens] |
| 07884 | 70 | glycosyltransferase family 39 protein [Denitromonas halophilus] |
| 02847 | 70 | cache domain-containing protein [Azoarcus olearius] |
| 00476 | 70 | MULTISPECIES: ferric iron uptake transcriptional regulator [Rhodocyclales] |
| 03191 | 70 | beta-N-acetylhexosaminidase [Azoarcus communis] |
| 04447 | 70 | methionyl-tRNA formyltransferase [Azoarcus communis] |
| 00320 | 70 | phosphoenolpyruvate carboxylase [Denitromonas halophilus] |
| 04022 | 70 | aromatic/alkene monooxygenase hydroxylase subunit beta [Aromatoleum toluclasticum] |
| 00671 | 70 | molybdopterin-dependent oxidoreductase [Aromatoleum toluclasticum] |
| 08993 | 70 | sulfite exporter TauE/SafE family protein [Azoarcus communis] |
| 04800 | 70 | acetyl-CoA carboxylase, carboxyltransferase subunit beta [Azoarcus communis] |
| 05058 | 70 | F0F1 ATP synthase subunit gamma [Denitromonas halophilus] |
| 09513 | 70 | hypothetical protein [Niveibacterium umoris] |
| 08522 | 70 | signal peptidase I [Denitromonas halophilus] |
| 03133 | 70 | aminomethyl-transferring glycine dehydrogenase [Aromatoleum toluclasticum] |
| 06961 | 70 | MULTISPECIES: heme ABC transporter permease CcmC [Aromatoleum] |
| 02923 | 70 | DUF934 domain-containing protein [Azoarcus communis] |
| 09534 | 70 | OmpA family protein [Azoarcus communis] |
| 01902 | 70 | phosphoenolpyruvate--protein phosphotransferase [Uliginosibacterium gangwonense] |
| 01913 | 70 | phosphoheptose isomerase [Azoarcus communis] |
| 02205 | 70 | acyltransferase [Azoarcus sp. DD4] |
| 10510 | 69 | class II glutamine amidotransferase [Azoarcus olearius] |
| 03406 | 69 | SDR family oxidoreductase [Azoarcus indigens] |
| 01639 | 69 | RibD family protein [Azoarcus communis] |
| 03122 | 69 | phosphotransferase family protein [Aromatoleum toluclasticum] |
| 07603 | 69 | NAD(P)(+) transhydrogenase (Re/Si-specific) subunit beta [Azoarcus communis] |

| | | |
|---|---|---|
| 05137 | 69 | tryptophan--tRNA ligase [Azoarcus communis] |
| 10353 | 69 | methyl-accepting chemotaxis protein [Thauera aromatica] |
| 05648 | 69 | phosphoenolpyruvate synthase [Azoarcus communis] |
| 02524 | 69 | BON domain-containing protein [Azoarcus indigens] |
| 03450 | 69 | MULTISPECIES: K+/H+ antiporter subunit F [Thauera] |
| 09069 | 69 | AAA family ATPase [Aromatoleum toluclasticum] |
| 05368 | 69 | 3,4-dihydroxy-2-butanone-4-phosphate synthase [Nitrogeniibacter aestuarii] |
| 07262 | 69 | RNA polymerase factor sigma-54 [Azoarcus indigens] |
| 06969 | 69 | YggS family pyridoxal phosphate-dependent enzyme [Azoarcus communis] |
| 09540 | 69 | uroporphyrinogen decarboxylase [Azoarcus communis] |
| 09190 | 69 | UDP-N-acetylglucosamine 2-epimerase (non-hydrolyzing) [Denitromonas halophilus] |
| 02666 | 69 | AzlC family ABC transporter permease [Denitromonas halophilus] |
| 04504 | 69 | glycosyltransferase family 2 protein [Azoarcus communis] |
| 08660 | 69 | MULTISPECIES: NADH-quinone oxidoreductase subunit NuoI [Aromatoleum] |
| 10466 | 69 | AmpG family muropeptide MFS transporter [Denitromonas halophilus] |
| 06872 | 68 | nitrous oxide reductase family maturation protein NosD [Denitromonas halophilus] |
| 05640 | 68 | flagellar brake protein [Azoarcus communis] |
| 05378 | 68 | bifunctional tetrahydrofolate synthase/dihydrofolate synthase [Azoarcus communis] |
| 03833 | 68 | single-stranded DNA-binding protein [Azoarcus communis] |
| 10307 | 68 | UPF0149 family protein [Aromatoleum buckelii] |
| 03209 | 68 | hydroxysqualene dehydroxylase HpnE [Azoarcus communis] |
| 01899 | 68 | FAD:protein FMN transferase [Azoarcus olearius] |
| 10299 | 68 | hypothetical protein [Aromatoleum toluclasticum] |
| 00225 | 68 | 4Fe-4S binding protein [Nitrogeniibacter aestuarii] |
| 04354 | 68 | IS110 family transposase [Azoarcus sp. DD4] |
| 07549 | 68 | 3-phosphoserine/phosphohydroxythreonine transaminase [Denitromonas halophilus] |
| 01861 | 68 | hydroxypyruvate isomerase [Azoarcus communis] |
| 09789 | 68 | Gfo/Idh/MocA family oxidoreductase [Propionivibrio dicarboxylicus] |
| 01524 | 67 | MULTISPECIES: homoserine O-acetyltransferase [Thauera] |
| 00824 | 67 | MULTISPECIES: peptidoglycan-associated lipoprotein Pal [Azoarcus] |
| 10454 | 67 | PQQ-dependent sugar dehydrogenase [Azoarcus communis] |
| 00235 | 67 | oligosaccharide flippase family protein [Denitromonas halophilus] |
| 04502 | 67 | potassium transporter Kup [Aromatoleum toluclasticum] |
| 06291 | 67 | ketoacyl-ACP synthase III [Azoarcus communis] |
| 07109 | 67 | biotin synthase BioB [Aromatoleum toluclasticum] |
| 09098 | 67 | response regulator [Azoarcus communis] |
| 10041 | 67 | CDP-diacylglycerol--glycerol-3-phosphate 3-phosphatidyltransferase [Azoarcus communis] |
| 00359 | 67 | AI-2E family transporter [Azoarcus communis] |
| 10096 | 67 | phosphomethylpyrimidine synthase ThiC [Azoarcus communis] |
| 05715 | 67 | SLC13 family permease [Denitromonas halophilus] |
| 08621 | 67 | MULTISPECIES: Fe-S cluster assembly scaffold IscU [Rhodocyclales] |
| 06124 | 67 | group II truncated hemoglobin [Azoarcus communis] |
| 07964 | 67 | 1-acyl-sn-glycerol-3-phosphate acyltransferase [Azoarcus communis] |
| 00165 | 67 | helix-turn-helix domain-containing protein [Nitrogeniibacter aestuarii] |
| 02209 | 67 | chorismate synthase [Aromatoleum toluclasticum] |
| 00258 | 67 | CinA family protein [Azoarcus indigens] |
| 03003 | 67 | ATP-dependent Clp protease ATP-binding subunit ClpX [Azoarcus communis] |
| 05732 | 66 | flagellar biosynthetic protein FliR [Azoarcus communis] |
| 04905 | 66 | flagellar type III secretion system pore protein FliP [Azoarcus communis] |
| 05539 | 66 | cytochrome c oxidase subunit II [Aromatoleum toluclasticum] |
| 05699 | 66 | F0F1 ATP synthase subunit epsilon [Denitromonas halophilus] |
| 01563 | 66 | HD domain-containing protein [Denitromonas halophilus] |
| 06034 | 66 | MULTISPECIES: glutaredoxin 3 [Niveibacterium] |
| 05471 | 66 | threonine--tRNA ligase [Azoarcus communis] |
| 07278 | 66 | NUDIX hydrolase [Azoarcus communis] |
| 09464 | 66 | alanine racemase [Azoarcus communis] |
| 03878 | 66 | GNAT family N-acetyltransferase [Azoarcus indigens] |
| 02685 | 66 | cytochrome-c oxidase, cbb3-type subunit III [Azoarcus communis] |
| 00828 | 66 | ATP-dependent Clp endopeptidase proteolytic subunit ClpP [Aromatoleum tolulyticum] |
| 04019 | 66 | transcriptional repressor [Azoarcus communis] |
| 03582 | 65 | extracellular solute-binding protein [Azoarcus communis] |
| 04505 | 65 | replicative DNA helicase [Azoarcus indigens] |
| 04359 | 65 | gamma-glutamyltransferase family protein [Nitrogeniibacter aestuarii] |
| 07062 | 65 | hydantoinase B/oxoprolinase family protein [Aromatoleum toluclasticum] |
| 01617 | 65 | XdhC family protein [Aromatoleum toluclasticum] |
| 06975 | 65 | MULTISPECIES: guanylate kinase [Thauera] |
| 06780 | 65 | acetyl-CoA carboxylase biotin carboxyl carrier protein [Aromatoleum toluclasticum] |
| 08619 | 65 | flagellar biosynthesis protein FlhA [Azoarcus communis] |
| 04116 | 65 | 1,6-anhydro-N-acetylmuramyl-L-alanine amidase AmpD [Azoarcus communis] |
| 08831 | 65 | HupU protein [Azoarcus communis] |
| 08413 | 65 | DUF2325 domain-containing protein [Azoarcus olearius] |
| 08863 | 65 | histidine kinase [Azoarcus communis] |
| 03840 | 65 | GNAT family N-acetyltransferase [Azoarcus communis] |
| 01893 | 65 | magnesium/cobalt transporter CorA [Azoarcus communis] |
| 00021 | 65 | flagellar basal body L-ring protein FlgH [Azoarcus communis] |
| 01578 | 65 | glycerol kinase GlpK [Azoarcus communis] |
| 07627 | 65 | nitronate monooxygenase [Azoarcus communis] |
| 08503 | 65 | flagellar basal body-associated FliL family protein [Azoarcus communis] |
| 07487 | 65 | protein-disulfide reductase DsbD [Azoarcus communis] |
| 02719 | 65 | VacJ family lipoprotein [Azoarcus communis] |
| 07179 | 65 | Sec-dependent nitrous-oxide reductase [Denitromonas halophilus] |
| 02698 | 65 | ABC transporter ATP-binding protein/permease [Azoarcus indigens] |
| 09089 | 65 | type II 3-dehydroquinate dehydratase [Azoarcus communis] |
| 05113 | 65 | flagellar type III secretion system protein FlhB [Azoarcus communis] |
| 00344 | 65 | M48 family metallopeptidase [Azoarcus indigens] |
| 03643 | 65 | 3-isopropylmalate dehydratase small subunit [Azoarcus communis] |
| 04841 | 65 | MBL fold metallo-hydrolase [Aromatoleum toluclasticum] |
| 08107 | 65 | HAD family phosphatase [Azoarcus communis] |
| 06479 | 64 | PAS domain-containing sensor histidine kinase [Denitromonas halophilus] |
| 10523 | 64 | histidine phosphatase family protein [Denitromonas halophilus] |
| 01731 | 64 | cation:proton antiporter [Aromatoleum toluclasticum] |
| 02018 | 64 | bifunctional protein-serine/threonine kinase/phosphatase [Denitromonas halophilus] |
| 04236 | 64 | lipid IV(A) 3-deoxy-D-manno-octulosonic acid transferase [Azoarcus communis] |
| 02019 | 64 | 1-(5-phosphoribosyl)-5-[(5-phosphoribosylamino)methylideneamino]imidazole-4-carboxamide isomerase [Azoarcus communis] |

| | | |
|---|---|---|
| 08094 | 64 | type II toxin-antitoxin system VapC family toxin [Thauera phenylacetica] |
| 06290 | 64 | TMEM165/GDT1 family protein [Azoarcus communis] |
| 05868 | 64 | MULTISPECIES: NADH-quinone oxidoreductase subunit NuoK [Thauera] |
| 07054 | 64 | cytochrome c-550 PedF [Azoarcus indigens] |
| 01223 | 64 | MULTISPECIES: bifunctional enoyl-CoA hydratase/phosphate acetyltransferase [Azoarcus] |
| 04583 | 64 | RDD family protein [Azoarcus communis] |
| 10146 | 64 | ATP-dependent DNA helicase [Azoarcus indigens] |
| 05437 | 64 | MULTISPECIES: PTS IIA-like nitrogen regulatory protein PtsN [Azoarcus] |
| 06106 | 64 | cell division protein FtsZ [Azoarcus communis] |
| 02429 | 64 | MULTISPECIES: flagellar biosynthesis protein FliQ [Rhodocyclales] |
| 01359 | 64 | IS66 family insertion sequence element accessory protein TnpB [Aromatoleum aromaticum] |
| 08779 | 64 | FtsX-like permease family protein [Azoarcus communis] |
| 04141 | 64 | MmoB/DmpM family protein [Aromatoleum toluclasticum] |
| 05487 | 64 | AzlD domain-containing protein [Denitromonas halophilus] |
| 07751 | 64 | DUF748 domain-containing protein [Azoarcus indigens] |
| 03505 | 64 | IMP dehydrogenase [Azoarcus communis] |
| 00100 | 64 | DsbA family oxidoreductase [Denitromonas halophilus] |
| 01335 | 63 | 4-hydroxythreonine-4-phosphate dehydrogenase PdxA [Azoarcus communis] |
| 01666 | 63 | flagellar hook-basal body complex protein FliE [Azoarcus communis] |
| 00502 | 63 | NADH-quinone oxidoreductase subunit J [Aromatoleum toluclasticum] |
| 02554 | 63 | flagellar M-ring protein FliF [Azoarcus communis] |
| 03452 | 63 | UDP-N-acetylmuramoyl-tripeptide--D-alanyl-D-alanine ligase [Azoarcus communis] |
| 09581 | 63 | 6,7-dimethyl-8-ribityllumazine synthase [Nitrogeniibacter aestuarii] |
| 02130 | 63 | MULTISPECIES: arginine/lysine/ornithine decarboxylase [Azoarcus] |
| 07704 | 63 | dioxygenase [Azoarcus communis] |
| 01201 | 63 | flagellar filament capping protein FliD [Azoarcus communis] |
| 06728 | 63 | endopeptidase La [Azoarcus communis] |
| 03950 | 63 | tyrosine--tRNA ligase [Azoarcus communis] |
| 04054 | 63 | MULTISPECIES: NADH-quinone oxidoreductase subunit A [Rhodocyclales] |
| 07830 | 63 | nitrite reductase large subunit NirB [Azoarcus communis] |
| 08983 | 63 | ketol-acid reductoisomerase, partial [Denitromonas halophilus] |
| 04152 | 63 | SDR family oxidoreductase [Uliginosibacterium gangwonense] |
| 06448 | 63 | isopentenyl-diphosphate Delta-isomerase [Aromatoleum toluclasticum] |
| 02156 | 63 | signal peptidase II [Azoarcus communis] |
| 03046 | 63 | DUF883 family protein [Azoarcus communis] |
| 03640 | 63 | NAD kinase [Azoarcus communis] |
| 08969 | 63 | 7-cyano-7-deazaguanine synthase QueC [Azoarcus communis] |
| 04643 | 63 | MULTISPECIES: CDP-diacylglycerol--serine O-phosphatidyltransferase [Niveibacterium] |
| 05127 | 63 | disulfide bond formation protein B [Azoarcus communis] |
| 07431 | 63 | PEP-CTERM/exosortase system-associated acyltransferase [Azoarcus indigens] |
| 06884 | 63 | MoxR family ATPase [Denitromonas halophilus] |
| 04282 | 63 | NADH-quinone oxidoreductase subunit C [Aromatoleum toluclasticum] |
| 04562 | 63 | SAM-dependent methyltransferase [Aromatoleum toluclasticum] |
| 07226 | 63 | MULTISPECIES: transcription termination factor Rho [Azoarcus] |
| 03302 | 63 | trimeric intracellular cation channel family protein [Azoarcus communis] |
| 04697 | 63 | flagellar basal body rod protein FlgC [Azoarcus communis] |
| 04647 | 63 | MULTISPECIES: sulfurtransferase TusA family protein [Rhodocyclales] |
| 01397 | 63 | prolipoprotein diacylglyceryl transferase [Azoarcus communis] |
| 03607 | 63 | D-alanine--D-alanine ligase [Azoarcus communis] |
| 10457 | 63 | exodeoxyribonuclease VII large subunit [Azoarcus communis] |
| 01241 | 63 | flagellar export chaperone FliS [Azoarcus indigens] |
| 00037 | 63 | 6-carboxytetrahydropterin synthase QueD [Azoarcus communis] |
| 03393 | 63 | glutamate synthase large subunit [Aromatoleum toluclasticum] |
| 10532 | 63 | DUF2892 domain-containing protein [Azoarcus communis] |
| 07282 | 63 | NAD+ synthase [Azoarcus communis] |
| 08565 | 62 | DUF3460 family protein [Aromatoleum toluclasticum] |
| 05450 | 62 | apolipoprotein N-acyltransferase [Azoarcus communis] |
| 08484 | 62 | histidinol dehydrogenase [Azoarcus communis] |
| 09118 | 62 | nicotinate-nucleotide--dimethylbenzimidazole phosphoribosyltransferase [Azoarcus communis] |
| 00555 | 62 | cytochrome c-type biogenesis protein CcmH [Azoarcus communis] |
| 03829 | 62 | alanine:cation symporter family protein [Denitromonas halophilus] |
| 09294 | 62 | MULTISPECIES: flagellar basal body rod protein FlgB [Thauera] |
| 07559 | 62 | DUF3683 domain-containing protein [Denitromonas halophilus] |
| 10474 | 62 | MULTISPECIES: 30S ribosomal protein S14 [Azoarcus] |
| 08155 | 62 | M3 family metallopeptidase [Azoarcus communis] |
| 03335 | 62 | bifunctional diguanylate cyclase/phosphodiesterase [Denitromonas halophilus] |
| 03550 | 62 | thiamine phosphate synthase [Azoarcus communis] |
| 04252 | 62 | zinc-dependent peptidase [Azoarcus communis] |
| 10202 | 62 | outer membrane lipid asymmetry maintenance protein MlaD [Azoarcus communis] |
| 03831 | 62 | iron-sulfur cluster carrier protein ApbC [Azoarcus communis] |
| 00176 | 62 | imidazole glycerol phosphate synthase subunit HisF [Azoarcus communis] |
| 09044 | 62 | cysteine--tRNA ligase [Azoarcus communis] |
| 02152 | 62 | glutamine--fructose-6-phosphate transaminase (isomerizing) [Azoarcus communis] |
| 01891 | 62 | LrgB family protein [Azoarcus indigens] |
| 01838 | 62 | DNA-binding protein YbiB [Azoarcus communis] |
| 00187 | 62 | CidA/LrgA family protein [Azoarcus indigens] |
| 00046 | 62 | flagellar basal body P-ring protein FlgI [Azoarcus indigens] |
| 07466 | 62 | phosphate signaling complex protein PhoU [Azoarcus communis] |
| 02696 | 62 | MltA domain-containing protein [Denitromonas halophilus] |
| 06418 | 62 | imidazole glycerol phosphate synthase subunit HisH [Denitromonas halophilus] |
| 02021 | 62 | tryptophan synthase subunit alpha [Azoarcus communis] |
| 05841 | 62 | TIGR01841 family phasin [Denitromonas halophilus] |
| 06612 | 62 | NUDIX hydrolase [Azoarcus communis] |
| 03377 | 62 | flagellar hook assembly protein FlgD [Azoarcus communis] |
| 07246 | 62 | type II toxin-antitoxin system prevent-host-death family antitoxin [Azoarcus nasutitermitis] |
| 03914 | 62 | 4-hydroxy-tetrahydrodipicolinate reductase [Azoarcus communis] |
| 06094 | 62 | murein biosynthesis integral membrane protein MurJ [Azoarcus communis] |
| 09131 | 61 | N-acetyl-gamma-glutamyl-phosphate reductase [Azoarcus communis] |
| 02233 | 61 | aminopeptidase N [Azoarcus communis] |
| 05003 | 61 | CYTH and CHAD domain-containing protein [Azoarcus communis] |
| 05483 | 61 | Bax inhibitor-1/YccA family protein [Azoarcus communis] |
| 09722 | 61 | ribosomal protein S18-alanine N-acetyltransferase [Azoarcus communis] |
| 07342 | 61 | GNAT family N-acetyltransferase [Nitrogeniibacter aestuarii] |
| 09793 | 61 | Tol-Pal system beta propeller repeat protein TolB [Azoarcus communis] |

| | | |
|---|---|---|
| 02222 | 61 | M48 family metallopeptidase [Azoarcus communis] |
| 07507 | 61 | MULTISPECIES: 50S ribosomal protein L27 [Aromatoleum] |
| 02368 | 61 | divalent-cation tolerance protein CutA [Azoarcus communis] |
| 04721 | 61 | class I SAM-dependent methyltransferase [Azoarcus communis] |
| 07540 | 61 | thymidylate synthase [Azoarcus communis] |
| 01098 | 61 | argininosuccinate synthase [Azoarcus communis] |
| 07635 | 61 | RNA polymerase sigma factor RpoD [Azoarcus communis] |
| 07539 | 61 | S-methyl-5'-thioinosine phosphorylase [Azoarcus communis] |
| 01851 | 61 | UvrD-helicase domain-containing protein [Denitromonas halophilus] |
| 01849 | 61 | cell division protein ZapD [Azoarcus communis] |
| 07215 | 61 | undecaprenyldiphospho-muramoylpentapeptide beta-N-acetylglucosaminyltransferase [Azoarcus communis] |
| 09658 | 61 | DUF2905 domain-containing protein [Nitrogeniibacter aestuarii] |
| 01258 | 61 | 50S ribosomal protein L9 [Azoarcus communis] |
| 01825 | 61 | DNA repair protein RecN [Azoarcus communis] |
| 05124 | 61 | phenylalanine--tRNA ligase subunit beta [Azoarcus communis] |
| 04863 | 61 | bifunctional 2-polyprenyl-6-hydroxyphenol methylase/3-demethylubiquinol 3-O-methyltransferase UbiG [Azoarcus communis] |
| 04878 | 61 | dihydroorotase [Azoarcus communis] |
| 01405 | 61 | exodeoxyribonuclease VII small subunit [Azoarcus communis] |
| 04820 | 61 | MULTISPECIES: LysR family substrate-binding domain-containing protein [Rhodocyclales] |
| 01174 | 61 | MULTISPECIES: methionine biosynthesis protein MetW [Thauera] |
| 02289 | 61 | ATP-binding protein [Azoarcus communis] |
| 01653 | 61 | phosphoribosylformylglycinamidine synthase [Azoarcus communis] |
| 01771 | 61 | MULTISPECIES: 30S ribosomal protein S13 [Thauera] |
| 01145 | 61 | cell division protein ZapE [Azoarcus communis] |
| 07702 | 61 | MULTISPECIES: VOC family protein [Aromatoleum] |
| 07701 | 61 | 23S rRNA (pseudouridine(1915)-N(3))-methyltransferase RlmH [Azoarcus communis] |
| 07281 | 61 | OsmC family protein [Azoarcus communis] |
| 09982 | 61 | ComF family protein [Azoarcus communis] |
| 07275 | 61 | YebC/PmpR family DNA-binding transcriptional regulator [Azoarcus communis] |
| 07036 | 61 | MULTISPECIES: F0F1 ATP synthase subunit C [Rhodocyclales] |
| 05288 | 61 | MULTISPECIES: 30S ribosomal protein S19 [Zoogloeaceae] |
| 09827 | 61 | 1-acyl-sn-glycerol-3-phosphate acyltransferase [Azoarcus communis] |
| 05474 | 61 | porphobilinogen synthase [Azoarcus communis] |
| 09487 | 61 | tRNA epoxyqueuosine(34) reductase QueG [Azoarcus communis] |
| 02439 | 61 | phosphoadenylyl-sulfate reductase [Azoarcus communis] |
| 02031 | 61 | recombinase RecA [Azoarcus communis] |
| 09371 | 61 | segregation/condensation protein A [Azoarcus communis] |
| 03135 | 61 | phosphatidylglycerophosphatase A [Azoarcus communis] |
| 03443 | 61 | DUF2917 domain-containing protein [Denitromonas halophilus] |
| 00314 | 61 | flagellar hook-associated protein FlgK [Azoarcus indigens] |
| 06786 | 61 | 50S ribosomal protein L4 [Azoarcus communis] |
| 06045 | 61 | cytochrome d ubiquinol oxidase subunit II [Azoarcus indigens] |
| 03824 | 61 | methionine adenosyltransferase [Azoarcus communis] |
| 02636 | 61 | lytic transglycosylase domain-containing protein [Azoarcus communis] |
| 10237 | 61 | acyl carrier protein [Denitromonas halophilus] |
| 05804 | 61 | hydroxymethylglutaryl-CoA lyase [Azoarcus communis] |
| 03818 | 61 | MULTISPECIES: 50S ribosomal protein L32 [Rhodocyclales] |
| 02959 | 61 | alanine--tRNA ligase [Azoarcus communis] |
| 07757 | 61 | HPr family phosphocarrier protein [Azoarcus communis] |
| 06613 | 61 | ribonuclease HI [Azoarcus communis] |
| 08778 | 61 | flagellar motor protein MotB [Azoarcus communis] |
| 10313 | 61 | uracil-DNA glycosylase [Azoarcus communis] |
| 05782 | 61 | triose-phosphate isomerase [Azoarcus communis] |
| 08285 | 61 | undecaprenyl-diphosphate phosphatase [Azoarcus communis] |
| 03734 | 61 | MULTISPECIES: 50S ribosomal protein L34 [Rhodocyclales] |
| 06040 | 61 | MULTISPECIES: oxidative damage protection protein [Aromatoleum] |
| 02997 | 61 | MULTISPECIES: 30S ribosomal protein S9 [Rhodocyclales] |
| 08668 | 61 | flagellar transcriptional regulator FlhC [Azoarcus communis] |
| 10115 | 61 | argininosuccinate lyase [Azoarcus communis] |
| 00050 | 61 | SirB2 family protein [Azoarcus communis] |
| 00277 | 61 | molecular chaperone HtpG [Azoarcus communis] |
| 04065 | 61 | 2-C-methyl-D-erythritol 4-phosphate cytidylyltransferase [Azoarcus communis] |
| 07809 | 61 | DUF4390 domain-containing protein [Azoarcus communis] |
| 06299 | 61 | thiol:disulfide interchange protein DsbA/DsbL [Azoarcus communis] |
| 03769 | 61 | MULTISPECIES: acyl carrier protein [Rhodocyclales] |
| 04082 | 61 | MULTISPECIES: 30S ribosomal protein S8 [Aromatoleum] |
| 00813 | 61 | tRNA uridine-5-carboxymethylaminomethyl(34) synthesis enzyme MnmG [Azoarcus communis] |
| 06483 | 61 | ATP phosphoribosyltransferase [Azoarcus communis] |
| 06360 | 61 | flagellar motor switch protein FliG [Azoarcus communis] |
| 03865 | 61 | MULTISPECIES: NAD(P) transhydrogenase subunit alpha [Thauera] |
| 02622 | 61 | 3-deoxy-8-phosphooctulonate synthase [Azoarcus communis] |
| 10177 | 61 | DUF615 domain-containing protein [Azoarcus communis] |
| 08488 | 61 | 5'/3'-nucleotidase SurE [Azoarcus communis] |
| 05643 | 61 | ribulose-phosphate 3-epimerase [Azoarcus communis] |
| 00215 | 61 | MULTISPECIES: accessory factor UbiK family protein [Aromatoleum] |
| 08784 | 61 | flagellar hook-associated protein FlgL [Azoarcus communis] |
| 06745 | 61 | MULTISPECIES: polyprenyl diphosphate synthase [Thauera] |
| 10543 | 61 | M48 family metallopeptidase [Azoarcus communis] |
| 05865 | 61 | molybdopterin synthase catalytic subunit MoaE [Azoarcus communis] |
| 00407 | 61 | 3-deoxy-manno-octulosonate cytidylyltransferase [Denitromonas halophilus] |
| 00334 | 60 | phosphate acyltransferase PlsX [Azoarcus communis] |
| 09186 | 60 | Gx transporter family protein [Azoarcus communis] |
| 03963 | 60 | MULTISPECIES: 30S ribosomal protein S10 [Thauera] |
| 08639 | 60 | MULTISPECIES: DUF2788 domain-containing protein [Rhodocyclales] |
| 08642 | 60 | excinuclease ABC subunit UvrB [Azoarcus communis] |
| 05693 | 60 | ribosome silencing factor [Azoarcus communis] |
| 05120 | 60 | riboflavin synthase [Azoarcus communis] |
| 10519 | 60 | 6-phosphofructokinase [Azoarcus communis] |
| 01139 | 60 | ATP-dependent RNA helicase HrpA [Azoarcus communis] |
| 01712 | 60 | fluoride efflux transporter CrcB [Azoarcus communis] |
| 03956 | 60 | peptidylprolyl isomerase [Azoarcus communis] |
| 02922 | 60 | nucleotide exchange factor GrpE [Azoarcus communis] |
| 10258 | 60 | rod shape-determining protein MreD [Azoarcus communis] |
| 02027 | 60 | 30S ribosomal protein S1 [Azoarcus communis] |

| | | |
|---|---|---|
| 06970 | 60 | 30S ribosomal protein S5 [Azoarcus communis] |
| 07110 | 60 | lipopolysaccharide assembly protein LapB [Azoarcus communis] |
| 07407 | 60 | glutamine-hydrolyzing GMP synthase [Azoarcus communis] |
| 01158 | 60 | MULTISPECIES: 50S ribosomal protein L3 [Aromatoleum] |
| 02151 | 60 | Holliday junction branch migration protein RuvA [Azoarcus communis] |
| 03857 | 60 | 50S ribosomal protein L15 [Azoarcus communis] |
| 05121 | 60 | DUF4124 domain-containing protein [Azoarcus communis] |
| 05862 | 60 | outer membrane protein assembly factor BamD [Azoarcus communis] |
| 01915 | 60 | MULTISPECIES: 30S ribosomal protein S20 [Aromatoleum] |
| 06062 | 60 | DNA translocase FtsK [Azoarcus communis] |
| 07394 | 60 | cytochrome c1 [Azoarcus communis] |
| 03714 | 60 | MULTISPECIES: succinate--CoA ligase subunit alpha [Rhodocyclales] |
| 05268 | 60 | MULTISPECIES: 30S ribosomal protein S15 [Aromatoleum] |
| 06723 | 60 | tRNA threonylcarbamoyladenosine dehydratase [Azoarcus communis] |
| 03715 | 60 | MULTISPECIES: UMP kinase [Rhodocyclales] |
| 03985 | 60 | MULTISPECIES: 50S ribosomal protein L11 [Aromatoleum] |
| 08341 | 60 | chromosomal replication initiator protein DnaA [Azoarcus communis] |
| 03992 | 60 | STAS domain-containing protein [Azoarcus communis] |
| 02854 | 60 | Rne/Rng family ribonuclease [Azoarcus communis] |
| 05878 | 60 | cobyric acid synthase [Azoarcus communis] |
| 10536 | 60 | oxygen-dependent coproporphyrinogen oxidase [Azoarcus communis] |
| 03688 | 60 | cytochrome ubiquinol oxidase subunit I [Azoarcus indigens] |
| 07365 | 60 | cyclic pyranopterin monophosphate synthase MoaC [Azoarcus communis] |
| 04716 | 60 | MULTISPECIES: ribosome-associated translation inhibitor RaiA [Thauera] |
| 06743 | 60 | phosphatidate cytidylyltransferase [Azoarcus communis] |
| 04324 | 60 | FHA domain-containing protein [Azoarcus communis] |
| 10537 | 60 | 50S ribosomal protein L24 [Azoarcus communis] |
| 04699 | 60 | carbonic anhydrase family protein [Azoarcus communis] |
| 07363 | 60 | RdgB/HAM1 family non-canonical purine NTP pyrophosphatase [Azoarcus communis] |
| 03999 | 60 | 50S ribosomal protein L17 [Azoarcus communis] |
| 10175 | 60 | DUF2249 domain-containing protein [Azoarcus communis] |
| 08021 | 60 | membrane protein insertase YidC [Azoarcus communis] |
| 04006 | 60 | MULTISPECIES: acetolactate synthase small subunit [Aromatoleum] |
| 04007 | 60 | transcription termination factor NusA [Azoarcus communis] |
| 09388 | 60 | methionine--tRNA ligase [Azoarcus communis] |
| 03131 | 60 | adenylosuccinate synthase [Azoarcus communis] |
| 03691 | 60 | BON domain-containing protein [Azoarcus communis] |
| 08512 | 60 | NAD-dependent DNA ligase LigA [Azoarcus communis] |
| 02354 | 60 | endolytic transglycosylase MltG [Azoarcus communis] |
| 03143 | 60 | LPS-assembly protein LptD [Azoarcus communis] |
| 06726 | 60 | protein translocase subunit SecD [Azoarcus communis] |
| 00383 | 60 | MULTISPECIES: Fis family transcriptional regulator [Azoarcus] |
| 04360 | 60 | cbb3-type cytochrome oxidase assembly protein CcoS [Azoarcus communis] |
| 08454 | 60 | polynucleotide adenylyltransferase PcnB [Azoarcus communis] |
| 01548 | 60 | crossover junction endodeoxyribonuclease RuvC [Azoarcus communis] |
| 01307 | 60 | OmpH family outer membrane protein [Aromatoleum toluclasticum] |
| 02337 | 60 | CbiX/SirB N-terminal domain-containing protein [Azoarcus communis] |
| 03984 | 60 | phosphoribosylaminoimidazolesuccinocarboxamide synthase [Azoarcus communis] |
| 10349 | 60 | tRNA guanosine(34) transglycosylase Tgt [Azoarcus communis] |
| 07381 | 60 | polysaccharide biosynthesis protein [Azoarcus communis] |
| 04954 | 60 | MULTISPECIES: DUF465 domain-containing protein [Aromatoleum] |
| 06950 | 60 | ribonuclease III [Azoarcus communis] |
| 04358 | 60 | MULTISPECIES: cytochrome c maturation protein CcmE [Aromatoleum] |
| 05291 | 60 | ribosome biogenesis GTPase YlqF [Azoarcus communis] |
| 08336 | 60 | Asp-tRNA(Asn)/Glu-tRNA(Gln) amidotransferase subunit GatB [Azoarcus communis] |
| 07478 | 60 | lipoprotein insertase outer membrane protein LolB [Azoarcus communis] |
| 05873 | 60 | c-type cytochrome biogenesis protein CcmI [Azoarcus communis] |
| 02297 | 60 | MULTISPECIES: 50S ribosomal protein L22 [Rhodocyclales] |
| 09328 | 60 | alpha/beta fold hydrolase [Denitromonas halophilus] |
| 07975 | 60 | transcription antitermination factor NusB [Azoarcus communis] |
| 01167 | 60 | DNA mismatch repair endonuclease MutL [Azoarcus communis] |
| 02208 | 60 | peptidoglycan DD-metalloendopeptidase family protein [Azoarcus communis] |
| 09274 | 60 | primosomal replication protein N [Azoarcus communis] |
| 08404 | 60 | monofunctional biosynthetic peptidoglycan transglycosylase [Azoarcus communis] |
| 06667 | 60 | cytochrome c biogenesis protein CcsA [Azoarcus communis] |
| 00282 | 60 | UDP-N-acetylglucosamine 1-carboxyvinyltransferase [Azoarcus communis] |
| 06457 | 60 | lipid-A-disaccharide synthase [Azoarcus communis] |
| 07019 | 60 | DUF484 family protein [Azoarcus communis] |
| 07018 | 60 | CvpA family protein [Azoarcus communis] |
| 09278 | 60 | FixH family protein [Azoarcus communis] |
| 07139 | 60 | MULTISPECIES: 50S ribosomal protein L23 [Thauera] |
| 03888 | 60 | 50S ribosomal protein L6 [Azoarcus communis] |
| 07141 | 60 | MULTISPECIES: Trm112 family protein [Rhodocyclales] |
| 01469 | 60 | MULTISPECIES: pantetheine-phosphate adenylyltransferase [Azoarcus] |
| 03208 | 60 | DNA primase [Azoarcus communis] |
| 07053 | 60 | type II toxin-antitoxin system RatA family toxin [Azoarcus communis] |
| 01473 | 60 | outer membrane protein assembly factor BamE [Azoarcus communis] |
| 05195 | 60 | prephenate dehydrogenase/arogenate dehydrogenase family protein [Azoarcus communis] |
| 03752 | 60 | MULTISPECIES: 50S ribosomal protein L29 [Thauera] |
| 04441 | 60 | MULTISPECIES: 30S ribosomal protein S16 [Aromatoleum] |
| 08537 | 60 | tetratricopeptide repeat protein [Azoarcus communis] |
| 05806 | 60 | protein translocase subunit SecF [Azoarcus communis] |
| 07959 | 60 | peptidoglycan editing factor PgeF [Azoarcus communis] |
| 09747 | 60 | LPS export ABC transporter permease LptF [Azoarcus communis] |
| 05040 | 60 | ribosome maturation factor RimP [Azoarcus communis] |
| 05986 | 60 | transcriptional regulator NrdR [Azoarcus communis] |
| 06477 | 60 | MULTISPECIES: division/cell wall cluster transcriptional repressor MraZ [Rhodocyclales] |
| 09742 | 60 | aspartate-semialdehyde dehydrogenase [Azoarcus communis] |
| 03032 | 60 | sigma-70 family RNA polymerase sigma factor [Denitromonas halophilus] |
| 09160 | 60 | MULTISPECIES: 50S ribosomal protein L7/L12 [Aromatoleum] |
| 04483 | 60 | RIP metalloprotease RseP [Azoarcus communis] |
| 03185 | 60 | anthranilate phosphoribosyltransferase [Azoarcus communis] |
| 07038 | 60 | cell division protein FtsL [Azoarcus communis] |
| 03011 | 60 | RlmE family RNA methyltransferase [Azoarcus communis] |

| | | |
|---|---|---|
| 07953 | 60 | TerB family tellurite resistance protein [Azoarcus communis] |
| 06009 | 60 | helix-turn-helix transcriptional regulator [Azoarcus communis] |
| 05161 | 60 | ribosome maturation factor RimM [Azoarcus communis] |
| 07956 | 60 | MULTISPECIES: glycine--tRNA ligase subunit alpha [Thauera] |
| 09645 | 60 | MULTISPECIES: phosphoribosyl-ATP diphosphatase [Azoarcus] |
| 07897 | 60 | transcription-repair coupling factor [Azoarcus communis] |
| 07895 | 60 | dTMP kinase [Azoarcus communis] |
| 00492 | 60 | MULTISPECIES: dCTP deaminase [Aromatoleum] |
| 06027 | 60 | bifunctional UDP-N-acetylglucosamine diphosphorylase/glucosamine-1-phosphate N-acetyltransferase GlmU [Azoarcus communis] |
| 04433 | 60 | MULTISPECIES: 30S ribosomal protein S7 [Aromatoleum] |
| 07476 | 60 | 4-hydroxybenzoate octaprenyltransferase [Azoarcus communis] |
| 03936 | 60 | dihydrofolate reductase [Denitromonas halophilus] |
| 08435 | 60 | SsrA-binding protein SmpB [Azoarcus communis] |
| 04995 | 60 | aminoacyl-tRNA hydrolase [Azoarcus communis] |
| 09317 | 60 | lipopolysaccharide transport periplasmic protein LptA [Azoarcus communis] |
| 03244 | 60 | DUF721 domain-containing protein [Denitromonas halophilus] |
| 07867 | 60 | MULTISPECIES: hypothetical protein [Thauera] |
| 03164 | 60 | THUMP domain-containing protein [Denitromonas halophilus] |
| 05322 | 60 | MULTISPECIES: 50S ribosomal protein L35 [Rhodocyclales] |
| 01512 | 60 | MULTISPECIES: cell division protein ZapA [Azoarcus] |
| 02026 | 60 | phenylalanine--tRNA ligase subunit alpha [Azoarcus communis] |
| 00565 | 60 | carotenoid 1,2-hydratase [Azoarcus communis] |
| 04648 | 60 | peroxide stress protein YaaA [Azoarcus communis] |
| 03942 | 60 | MULTISPECIES: 30S ribosomal protein S12 [Azoarcus] |
| 09597 | 60 | ribonuclease HII [Azoarcus communis] |
| 01933 | 60 | ribosome biogenesis GTPase Der [Azoarcus communis] |
| 09590 | 60 | iron donor protein CyaY [Azoarcus communis] |
| 01930 | 60 | Fe-S cluster assembly protein IscX [Azoarcus communis] |
| 03049 | 60 | arginine--tRNA ligase [Azoarcus communis] |
| 01520 | 60 | CoA pyrophosphatase [Azoarcus communis] |
| 01954 | 60 | amidophosphoribosyltransferase [Azoarcus communis] |
| 03909 | 60 | ZipA [Azoarcus communis] |
| 06030 | 60 | DNA-directed RNA polymerase subunit alpha [Azoarcus communis] |
| 08379 | 60 | orotate phosphoribosyltransferase [Azoarcus communis] |
| 03894 | 60 | serine--tRNA ligase [Azoarcus communis] |
| 09297 | 60 | preprotein translocase subunit SecE [Azoarcus communis] |
| 00552 | 60 | deoxyguanosinetriphosphate triphosphohydrolase [Azoarcus communis] |
| 06031 | 60 | Holliday junction branch migration DNA helicase RuvB [Azoarcus communis] |
| 07875 | 60 | 30S ribosomal protein S2 [Azoarcus communis] |
| 06032 | 60 | HPr(Ser) kinase/phosphatase [Azoarcus communis] |
| 03862 | 60 | MULTISPECIES: 30S ribosomal protein S21 [Zoogloeaceae] |
| 03738 | 60 | MULTISPECIES: 50S ribosomal protein L21 [Azoarcus] |
| 01205 | 60 | outer membrane protein assembly factor BamB [Azoarcus communis] |
| 01492 | 60 | LPS assembly lipoprotein LptE [Aromatoleum toluclasticum] |
| 08377 | 60 | MULTISPECIES: STAS/SEC14 domain-containing protein [Azoarcus] |
| 08589 | 60 | Sec-independent protein translocase protein TatB [Azoarcus communis] |
| 09788 | 60 | NusG domain II-containing protein [Azoarcus communis] |
| 05000 | 60 | class I SAM-dependent methyltransferase [Denitromonas halophilus] |
| 05743 | 60 | tRNA pseudouridine(55) synthase TruB [Azoarcus communis] |
| 09100 | 60 | trigger factor [Azoarcus communis] |
| 03906 | 60 | 50S ribosomal protein L2 [Azoarcus communis] |
| 01490 | 60 | MULTISPECIES: 50S ribosomal protein L36 [Rhodocyclales] |
| 00674 | 60 | LPS export ABC transporter periplasmic protein LptC [Azoarcus communis] |
| 03864 | 60 | MULTISPECIES: 50S ribosomal protein L20 [Rhodocyclales] |
| 08990 | 60 | DUF1289 domain-containing protein [Azoarcus communis] |
| 06549 | 60 | oligoribonuclease [Azoarcus communis] |
| 05435 | 60 | 30S ribosomal protein S6 [Azoarcus communis] |
| 05552 | 60 | 3-methyl-2-oxobutanoate hydroxymethyltransferase [Azoarcus communis] |
| 09518 | 60 | ribosome recycling factor [Azoarcus communis] |
| 08213 | 60 | ribose-5-phosphate isomerase RpiA [Azoarcus communis] |
| 09949 | 60 | glucose-6-phosphate isomerase [Azoarcus communis] |
| 03441 | 60 | DNA gyrase inhibitor YacG [Azoarcus communis] |
| 07300 | 60 | biotin--[acetyl-CoA-carboxylase] ligase [Azoarcus communis] |
| 00414 | 60 | MULTISPECIES: recombination mediator RecR [Thauera] |
| 04207 | 60 | DUF2065 domain-containing protein [Azoarcus communis] |
| 09451 | 60 | adenosylhomocysteinase [Azoarcus communis] |
| 05445 | 60 | MULTISPECIES: imidazoleglycerol-phosphate dehydratase HisB [Aromatoleum] |
| 06548 | 60 | DUF2069 domain-containing protein [Azoarcus communis] |
| 09963 | 60 | septum site-determining protein MinC [Azoarcus communis] |
| 08144 | 60 | CTP synthase [Azoarcus communis] |
| 00943 | 60 | acyl-ACP--UDP-N-acetylglucosamine O-acyltransferase [Azoarcus communis] |
| 05449 | 60 | 30S ribosomal protein S3 [Denitromonas halophilus] |
| 00816 | 60 | rRNA maturation RNase YbeY [Azoarcus communis] |
| 00818 | 60 | DNA repair protein RadA [Azoarcus communis] |
| 03438 | 60 | lipoyl(octanoyl) transferase LipB [Azoarcus communis] |
| 07302 | 60 | MULTISPECIES: heme exporter protein CcmB [Azoarcus] |
| 03838 | 60 | glutamate-5-semialdehyde dehydrogenase [Azoarcus communis] |
| 08220 | 60 | recombination regulator RecX [Azoarcus communis] |
| 09927 | 60 | A/G-specific adenine glycosylase [Azoarcus communis] |
| 02044 | 60 | ribonuclease PH [Azoarcus communis] |
| 08112 | 60 | 50S ribosomal protein L19 [Azoarcus communis] |
| 08905 | 60 | 16S rRNA (uracil(1498)-N(3))-methyltransferase [Azoarcus communis] |
| 08035 | 60 | hypothetical protein [Aromatoleum toluclasticum] |
| 02683 | 60 | DUF177 domain-containing protein [Denitromonas halophilus] |
| 08477 | 60 | pantoate--beta-alanine ligase [Azoarcus communis] |
| 00790 | 60 | DUF971 domain-containing protein [Azoarcus communis] |
| 01396 | 60 | 5-(carboxyamino)imidazole ribonucleotide mutase [Azoarcus communis] |
| 09938 | 60 | MULTISPECIES: protein-export chaperone SecB [Azoarcus] |
| 00791 | 60 | YicC family protein [Azoarcus communis] |
| 07708 | 60 | pyrroline-5-carboxylate reductase [Azoarcus communis] |
| 01786 | 60 | translation elongation factor Ts [Azoarcus communis] |
| 04223 | 60 | MULTISPECIES: malate dehydrogenase [Azoarcus] |
| 05099 | 60 | 2-polyprenyl-3-methyl-6-methoxy-1,4-benzoquinone monooxygenase [Azoarcus communis] |
| 07305 | 60 | MULTISPECIES: membrane protein insertion efficiency factor YidD [Thauera] |

| | | |
|---|---|---|
| **02129** | 60 | proline--tRNA ligase [Azoarcus communis] |
| **07100** | 60 | adenylate kinase [Azoarcus communis] |
| **05091** | 60 | tRNA pseudouridine(38-40) synthase TruA [Azoarcus communis] |
| **03821** | 60 | 50S ribosomal protein L5 [Azoarcus communis] |
| **03072** | 60 | MULTISPECIES: nucleoside-diphosphate kinase [Azoarcus] |
| **00034** | 60 | MULTISPECIES: 30S ribosomal protein S17 [Aromatoleum] |
| **08145** | 60 | glycerol-3-phosphate 1-O-acyltransferase PlsY [Azoarcus communis] |
| **09224** | 60 | 6-bladed beta-propeller [Aromatoleum toluclasticum] |
| **02630** | 60 | MULTISPECIES: CcoQ/FixQ family Cbb3-type cytochrome c oxidase assembly chaperone [Thauera] |
| **06851** | 60 | phosphoribosylamine--glycine ligase [Azoarcus communis] |
| **03835** | 60 | MULTISPECIES: 50S ribosomal protein L33 [Zoogloeaceae] |
| **06523** | 60 | exopolyphosphatase [Azoarcus communis] |
| **04521** | 60 | MULTISPECIES: YbaB/EbfC family nucleoid-associated protein [Aromatoleum] |
| **06240** | 60 | bifunctional phosphopantothenoylcysteine decarboxylase/phosphopantothenate--cysteine ligase CoaBC [Azoarcus communis] |
| **09223** | 60 | DUF192 domain-containing protein [Azoarcus communis] |
| **10032** | 60 | MULTISPECIES: 30S ribosomal protein S18 [Zoogloeaceae] |
| **02081** | 60 | competence/damage-inducible protein A [Aromatoleum toluclasticum] |
| **05486** | 60 | YdiU family protein [Azoarcus communis] |
| **06527** | 60 | SCP2 sterol-binding domain-containing protein [Azoarcus communis] |
| **09486** | 60 | tRNA (cytidine(34)-2'-O)-methyltransferase [Denitromonas halophilus] |
| **10015** | 60 | 50S ribosomal protein L25/general stress protein Ctc [Azoarcus communis] |
| **03096** | 60 | RNase adapter RapZ [Azoarcus communis] |
| **06253** | 60 | MULTISPECIES: Asp-tRNA(Asn)/Glu-tRNA(Gln) amidotransferase subunit GatC [Denitromonas] |
| **10614** | 60 | ClpXP protease specificity-enhancing factor [Azoarcus communis] |
| **01360** | 60 | MULTISPECIES: 50S ribosomal protein L18 [Thauera] |
| **09471** | 60 | tRNA 2-thiouridine(34) synthase MnmA [Azoarcus communis] |
| **05458** | 60 | ferrochelatase [Azoarcus communis] |
| **04832** | 60 | RidA family protein [Denitromonas halophilus] |
| **08871** | 60 | 1-deoxy-D-xylulose-5-phosphate reductoisomerase [Azoarcus communis] |
| **08204** | 60 | 2-oxoglutarate dehydrogenase E1 component [Azoarcus communis] |
| **00826** | 60 | ribonuclease G [Azoarcus communis] |
| **04186** | 60 | MULTISPECIES: 50S ribosomal protein L13 [Rhodocyclales] |
| **00362** | 60 | YajQ family cyclic di-GMP-binding protein [Azoarcus communis] |
| **03822** | 60 | MULTISPECIES: transcription termination/antitermination protein NusG [Aromatoleum] |
| **04829** | 60 | Grx4 family monothiol glutaredoxin [Azoarcus communis] |
| **04831** | 60 | MULTISPECIES: cell division topological specificity factor MinE [Azoarcus] |
| **03823** | 60 | MULTISPECIES: 30S ribosomal protein S11 [Rhodocyclales] |
| **06520** | 60 | tRNA (adenosine(37)-N6)-threonylcarbamoyltransferase complex transferase subunit TsaD [Azoarcus communis] |
| **03541** | 60 | polyphosphate kinase 1 [Azoarcus communis] |
| **00841** | 60 | sensor histidine kinase [Azoarcus communis] |
| **01385** | 60 | DNA-directed RNA polymerase subunit beta [Azoarcus communis] |
| **04133** | 60 | MULTISPECIES: 50S ribosomal protein L31 [Aromatoleum] |
| **02573** | 60 | 16S rRNA (adenine(1518)-N(6)/adenine(1519)-N(6))-dimethyltransferase RsmA [Azoarcus communis] |
| **04180** | 60 | MULTISPECIES: 50S ribosomal protein L28 [Rhodocyclales] |
| **04834** | 60 | SurA N-terminal domain-containing protein [Azoarcus communis] |
| **04179** | 60 | flavodoxin-dependent (E)-4-hydroxy-3-methylbut-2-enyl-diphosphate synthase [Azoarcus communis] |
| **05414** | 60 | trifunctional transcriptional regulator/proline dehydrogenase/L-glutamate gamma-semialdehyde dehydrogenase [Denitromonas halophilus] |
| **04799** | 60 | Nudix family hydrolase [Azoarcus communis] |
| **05574** | 60 | quinolinate synthase NadA [Azoarcus communis] |
| **00730** | 60 | tRNA preQ1(34) S-adenosylmethionine ribosyltransferase-isomerase QueA [Azoarcus communis] |
| **09701** | 60 | MULTISPECIES: RNA chaperone Hfq [Rhodocyclales] |
| **05352** | 60 | DNA-directed RNA polymerase subunit beta' [Azoarcus communis] |
| **02762** | 60 | cytochrome bc complex cytochrome b subunit [Azoarcus communis] |
| **07664** | 60 | transglycosylase SLT domain-containing protein [Azoarcus communis] |
| **09414** | 60 | leucyl/phenylalanyl-tRNA--protein transferase [Azoarcus communis] |
| **01832** | 60 | prephenate dehydratase [Azoarcus communis] |
| **07791** | 60 | MULTISPECIES: Sec-independent protein translocase subunit TatA [Rhodocyclales] |
| **04059** | 60 | YqgE/AlgH family protein [Azoarcus communis] |
| **00728** | 60 | DNA polymerase III subunit beta [Azoarcus communis] |
| **06767** | 60 | zinc ribbon domain-containing protein [Azoarcus communis] |
| **01033** | 60 | indole-3-glycerol phosphate synthase TrpC [Azoarcus communis] |
| **03069** | 60 | RNB domain-containing ribonuclease [Azoarcus communis] |
| **04929** | 60 | MULTISPECIES: F0F1 ATP synthase subunit B [Thauera] |
| **01599** | 60 | MULTISPECIES: RNA pyrophosphohydrolase [Rhodocyclales] |
| **00733** | 60 | endonuclease III [Azoarcus communis] |
| **08252** | 60 | bifunctional [glutamate--ammonia ligase]-adenylyl-L-tyrosine phosphorylase/[glutamate--ammonia-ligase] adenylyltransferase [Azoarcus communis] |
| **06906** | 60 | SMC-Scp complex subunit ScpB [Azoarcus communis] |
| **04060** | 60 | translation initiation factor IF-2 [Azoarcus communis] |
| **10418** | 60 | DUF493 family protein [Denitromonas halophilus] |
| **04926** | 60 | MULTISPECIES: dUTP diphosphatase [Thauera] |
| **06583** | 60 | Smr/MutS family protein [Azoarcus communis] |
| **04045** | 60 | RNA methyltransferase [Azoarcus communis] |
| **08718** | 60 | tRNA (adenosine(37)-N6)-threonylcarbamoyltransferase complex ATPase subunit type 1 TsaE [Azoarcus communis] |
| **02133** | 60 | cupin domain-containing protein [Azoarcus communis] |
| **08989** | 60 | MULTISPECIES: PTS fructose transporter subunit IIA [Azoarcus] |
| **00681** | 60 | tRNA (adenosine(37)-N6)-threonylcarbamoyltransferase complex dimerization subunit type 1 TsaB [Azoarcus communis] |
| **10165** | 60 | NAD(P)H-hydrate dehydratase [Azoarcus communis] |
| **03177** | 60 | bifunctional phosphoribosylaminoimidazolecarboxamide formyltransferase/IMP cyclohydrolase [Azoarcus communis] |
| **02137** | 60 | 50S ribosomal protein L1 [Azoarcus communis] |
| **08304** | 60 | hydroxymethylbilane synthase [Azoarcus communis] |
| **07907** | 60 | hypothetical protein [Azoarcus communis] |
| **05642** | 60 | phosphoribosylformylglycinamidine cyclo-ligase [Azoarcus communis] |
| **05108** | 60 | phosphoribosylanthranilate isomerase [Azoarcus communis] |
| **01319** | 60 | adenylosuccinate lyase [Azoarcus communis] |
| **04745** | 60 | bifunctional glutamate N-acetyltransferase/amino-acid acetyltransferase ArgJ [Azoarcus communis] |
| **03645** | 60 | twin-arginine translocase subunit TatC [Azoarcus communis] |
| **04935** | 60 | SlyX family protein [Azoarcus communis] |
| **07654** | 60 | diaminopimelate epimerase [Azoarcus communis] |
| **07524** | 60 | glutamyl-tRNA reductase [Azoarcus communis] |
| **00711** | 60 | PHP domain-containing protein [Azoarcus communis] |
| **08715** | 60 | 16S rRNA (guanine(527)-N(7))-methyltransferase RsmG [Azoarcus communis] |
| **06757** | 60 | LutB/LldF family L-lactate oxidation iron-sulfur protein [Thauera phenylacetica] |
| **00715** | 60 | Fe-S protein assembly co-chaperone HscB [Azoarcus communis] |
| **08941** | 60 | single-stranded-DNA-specific exonuclease RecJ [Azoarcus communis] |

| | | |
|---|---|---|
| 00738 | 60 | ABC transporter permease [Azoarcus communis] |
| 06783 | 60 | UDP-3-O-acyl-N-acetylglucosamine deacetylase [Azoarcus communis] |
| 06353 | 60 | tol-pal system protein YbgF [Azoarcus communis] |
| 05391 | 60 | DNA mismatch repair protein MutS [Azoarcus communis] |
| 09538 | 60 | 4-(cytidine 5'-diphospho)-2-C-methyl-D-erythritol kinase [Azoarcus communis] |
| 06895 | 60 | MULTISPECIES: ATP-dependent Clp protease adapter ClpS [Rhodocyclales] |
| 07703 | 60 | YggT family protein [Azoarcus communis] |
| 09532 | 60 | outer membrane protein assembly factor BamC [Azoarcus communis] |
| 03847 | 60 | MULTISPECIES: 50S ribosomal protein L14 [Zoogloeaceae] |
| 02454 | 60 | YhbY family RNA-binding protein [Denitromonas halophilus] |
| 01797 | 60 | 1-deoxy-D-xylulose-5-phosphate synthase [Azoarcus communis] |
| 06577 | 60 | Holliday junction resolvase RuvX [Azoarcus communis] |
| 10109 | 60 | glutathione synthase [Azoarcus communis] |
| 05594 | 60 | alpha-ribazole phosphatase family protein [Azoarcus communis] |
| 05590 | 60 | bifunctional riboflavin kinase/FAD synthetase [Azoarcus communis] |
| 03403 | 60 | UDP-N-acetylmuramate dehydrogenase [Azoarcus communis] |
| 01331 | 60 | arginyltransferase [Azoarcus communis] |
| 04086 | 60 | preprotein translocase subunit SecY [Azoarcus communis] |
| 03604 | 60 | holo-ACP synthase [Azoarcus communis] |
| 01806 | 60 | DUF2863 family protein [Azoarcus communis] |
| 00104 | 60 | lytic transglycosylase domain-containing protein [Denitromonas halophilus] |
| 08246 | 60 | elongation factor P [Azoarcus communis] |
| 05104 | 60 | 3-dehydroquinate synthase [Azoarcus communis] |
| 08249 | 60 | replication-associated recombination protein A [Azoarcus communis] |
| 08936 | 60 | cell division protein FtsQ/DivIB [Azoarcus communis] |
| 05412 | 60 | bifunctional DNA-formamidopyrimidine glycosylase/DNA-(apurinic or apyrimidinic site) lyase [Azoarcus communis] |
| 10120 | 60 | tRNA (guanosine(37)-N1)-methyltransferase TrmD [Azoarcus communis] |
| 05958 | 60 | MerR family transcriptional regulator [Azoarcus communis] |
| 04760 | 60 | MULTISPECIES: 50S ribosomal protein L16 [Thauera] |
| 04768 | 60 | MULTISPECIES: 50S ribosomal protein L30 [Rhodocyclales] |
| 08248 | 60 | glycine--tRNA ligase subunit beta [Azoarcus communis] |
| 03813 | 60 | 2-C-methyl-D-erythritol 2,4-cyclodiphosphate synthase [Azoarcus communis] |
| 08930 | 60 | MotA/TolQ/ExbB proton channel family protein [Azoarcus communis] |
| 04088 | 60 | F0F1 ATP synthase subunit delta [Azoarcus communis] |
| 01809 | 60 | 5-formyltetrahydrofolate cyclo-ligase [Azoarcus communis] |
| 07321 | 60 | ribosome biogenesis GTP-binding protein YihA/YsxC [Azoarcus communis] |
| 00458 | 59 | Cys-tRNA(Pro) deacylase [Azoarcus communis] |
| 05879 | 59 | YraN family protein [Azoarcus communis] |
| 02074 | 59 | DNA polymerase III subunit chi [Azoarcus communis] |
| 07050 | 59 | type 4a pilus biogenesis protein PilO [Azoarcus communis] |
| 00459 | 59 | VanZ family protein [Azoarcus communis] |
| 03834 | 59 | GNAT family N-acetyltransferase [Azoarcus communis] |
| 01289 | 59 | heat-inducible transcriptional repressor HrcA [Azoarcus communis] |
| 07052 | 59 | Hsp33 family molecular chaperone HslO [Azoarcus communis] |
| 04497 | 59 | tRNA 2-selenouridine(34) synthase MnmH [Azoarcus communis] |
| 02162 | 59 | TIGR03862 family flavoprotein [Azoarcus communis] |
| 03795 | 59 | carbamoyl-phosphate synthase large subunit [Denitromonas halophilus] |
| 09193 | 59 | excinuclease ABC subunit UvrC [Azoarcus communis] |
| 09694 | 59 | LPS export ABC transporter permease LptG [Azoarcus communis] |
| 06555 | 59 | DNA repair protein RecO [Azoarcus communis] |
| 01388 | 59 | GTPase Era [Azoarcus communis] |
| 06608 | 59 | DNA internalization-related competence protein ComEC/Rec2 [Azoarcus communis] |
| 02120 | 59 | NUDIX hydrolase [Azoarcus communis] |
| 07526 | 59 | preprotein translocase subunit YajC [Azoarcus communis] |
| 03701 | 59 | protein phosphatase CheZ [Azoarcus communis] |
| 01437 | 59 | polyhydroxyalkanoate synthesis repressor PhaR [Azoarcus communis] |
| 04104 | 59 | DNA polymerase III subunit gamma/tau [Azoarcus communis] |
| 10062 | 59 | UDP-3-O-(3-hydroxymyristoyl)glucosamine N-acyltransferase [Azoarcus communis] |
| 06332 | 59 | homoserine dehydrogenase [Azoarcus communis] |
| 04803 | 59 | type III pantothenate kinase [Azoarcus communis] |
| 04216 | 59 | methionine synthase [Denitromonas halophilus] |
| 09930 | 59 | tetraacyldisaccharide 4'-kinase [Azoarcus communis] |
| 08114 | 59 | LuxR C-terminal-related transcriptional regulator [Aromatoleum toluclasticum] |
| 06341 | 59 | NAD(P)-dependent glycerol-3-phosphate dehydrogenase [Azoarcus communis] |
| 00988 | 59 | 3',5'-cyclic-nucleotide phosphodiesterase [Azoarcus communis] |
| 03404 | 59 | DnaA regulatory inactivator Hda [Azoarcus communis] |
| 06120 | 59 | isocitrate lyase [Azoarcus communis] |
| 00092 | 59 | DNA polymerase III subunit delta' [Azoarcus communis] |
| 08749 | 59 | YafY family transcriptional regulator [Azoarcus communis] |
| 08925 | 59 | elongation factor P maturation arginine rhamnosyltransferase EarP [Azoarcus communis] |
| 04771 | 59 | hypothetical protein [Azoarcus communis] |
| 07471 | 59 | DUF4115 domain-containing protein [Denitromonas halophilus] |
| 03627 | 59 | excinuclease ABC subunit UvrA [Azoarcus communis] |
| 04043 | 59 | phosphotransferase [Azoarcus communis] |
| 02778 | 59 | 2-amino-4-hydroxy-6-hydroxymethyldihydropteridine diphosphokinase [Azoarcus communis] |
| 06206 | 59 | RNA-binding transcriptional accessory protein [Azoarcus communis] |
| 04204 | 59 | TRAP transporter small permease [Azoarcus indigens] |
| 04890 | 59 | hypothetical protein [Aromatoleum toluclasticum] |
| 02664 | 59 | rod shape-determining protein MreC [Azoarcus communis] |
| 06287 | 59 | DNA-processing protein DprA [Azoarcus communis] |
| 03523 | 59 | MULTISPECIES: zinc-finger domain-containing protein [Thauera] |
| 03469 | 59 | YaeQ family protein [Azoarcus communis] |
| 03466 | 59 | 16S rRNA (cytosine(1402)-N(4))-methyltransferase RsmH [Azoarcus communis] |
| 04142 | 59 | YfiR family protein [Nitrogeniibacter aestuarii] |
| 05480 | 59 | 30S ribosome-binding factor RbfA [Azoarcus communis] |
| 08848 | 59 | Nif3-like dinuclear metal center hexameric protein [Azoarcus communis] |
| 09472 | 59 | 4-hydroxy-3-methylbut-2-enyl diphosphate reductase [Azoarcus communis] |
| 06835 | 59 | hypothetical protein [Denitromonas halophilus] |
| 02577 | 59 | SCP2 sterol-binding domain-containing protein [Azoarcus communis] |
| 01739 | 59 | DUF4845 domain-containing protein [Azoarcus communis] |
| 08812 | 59 | MULTISPECIES: inorganic diphosphatase [Aromatoleum] |
| 06858 | 59 | [protein-PII] uridylyltransferase [Azoarcus communis] |
| 07247 | 59 | metal-sulfur cluster assembly factor [Denitromonas halophilus] |
| 08866 | 59 | DNA polymerase III subunit delta [Azoarcus communis] |

| | | |
|---|---|---|
| **02560** | 59 | Mth938-like domain-containing protein [Azoarcus communis] |
| **07756** | 59 | flagellar assembly protein FliH [Azoarcus communis] |
| **08704** | 59 | DNA polymerase I [Azoarcus communis] |
| **09887** | 59 | pyrimidine/purine nucleoside phosphorylase [Azoarcus communis] |
| **02405** | 59 | LysM peptidoglycan-binding domain-containing protein [Azoarcus communis] |
| **05736** | 59 | pyridoxine 5'-phosphate synthase [Azoarcus communis] |
| **09347** | 59 | TIGR02099 family protein [Azoarcus communis] |
| **00688** | 59 | 2,3,4,5-tetrahydropyridine-2,6-dicarboxylate N-succinyltransferase [Azoarcus communis] |
| **06712** | 59 | single-strand selective monofunctional uracil-DNA glycosylase [Denitromonas halophilus] |
| **09075** | 59 | MFS transporter [Azoarcus communis] |
| **02296** | 59 | CopD family protein [Azoarcus communis] |
| **05252** | 59 | MULTISPECIES: molybdopterin adenylyltransferase [Thauera] |
| **03927** | 59 | YIP1 family protein [Denitromonas halophilus] |
| **10501** | 59 | TonB family protein [Denitromonas halophilus] |
| **08590** | 59 | 50S ribosomal protein L11 methyltransferase [Azoarcus communis] |
| **09061** | 59 | polyphosphate kinase 2 [Nitrogeniibacter aestuarii] |
| **06036** | 59 | polyribonucleotide nucleotidyltransferase [Azoarcus communis] |
| **02242** | 59 | dephospho-CoA kinase [Azoarcus communis] |
| **00541** | 59 | DUF4124 domain-containing protein [Azoarcus indigens] |
| **03210** | 59 | MULTISPECIES: DUF2782 domain-containing protein [Aromatoleum] |
| **09119** | 59 | symmetrical bis(5'-nucleosyl)-tetraphosphatase [Azoarcus communis] |
| **01989** | 59 | acyl-CoA thioesterase [Denitromonas halophilus] |
| **00286** | 59 | shikimate kinase [Azoarcus communis] |
| **03763** | 59 | kinase/pyrophosphorylase [Azoarcus communis] |
| **03954** | 59 | MerR family transcriptional regulator [Azoarcus communis] |
| **04382** | 59 | DUF2322 family protein [Azoarcus communis] |
| **04539** | 59 | RNA-binding S4 domain-containing protein [Azoarcus communis] |
| **07600** | 59 | transglycosylase SLT domain-containing protein [Azoarcus communis] |
| **05297** | 59 | uroporphyrinogen-III synthase [Azoarcus communis] |
| **07623** | 59 | hypothetical protein [Azoarcus communis] |
| **01144** | 59 | A24 family peptidase [Azoarcus communis] |
| **04709** | 59 | YbfB/YjiJ family MFS transporter [Azoarcus communis] |
| **09398** | 59 | tetratricopeptide repeat protein [Aromatoleum pumilum] |
| **08659** | 59 | MULTISPECIES: succinate dehydrogenase assembly factor 2 [Thauera] |
| **02872** | 59 | Fe-S cluster assembly protein SufB [Denitromonas halophilus] |
| **07646** | 59 | molybdenum cofactor guanylyltransferase MobA [Azoarcus communis] |
| **01138** | 59 | type IV pilus biogenesis/stability protein PilW [Azoarcus communis] |
| **03316** | 59 | histidine phosphatase family protein [Azoarcus communis] |
| **04695** | 59 | preprotein translocase subunit SecG [Azoarcus communis] |
| **02355** | 59 | molybdopterin converting factor subunit 1 [Azoarcus communis] |
| **01458** | 58 | PilZ domain-containing protein [Azoarcus communis] |
| **04918** | 58 | ParA family protein [Azoarcus communis] |
| **06372** | 58 | nucleotide pyrophosphohydrolase [Azoarcus communis] |
| **03532** | 58 | tRNA 2-thiocytidine(32) synthetase TtcA [Azoarcus communis] |
| **03614** | 58 | esterase [Azoarcus communis] |
| **07270** | 58 | serine/threonine protein kinase [Azoarcus communis] |
| **01643** | 58 | ribonuclease R [Azoarcus communis] |
| **08200** | 58 | STAS domain-containing protein [Azoarcus indigens] |
| **04916** | 58 | 8-amino-7-oxononanoate synthase [Azoarcus communis] |
| **08327** | 58 | SoxR reducing system RseC family protein [Azoarcus communis] |
| **06439** | 58 | chemotaxis protein CheW [Azoarcus communis] |
| **04785** | 58 | diguanylate cyclase [Azoarcus communis] |
| **07495** | 58 | hypothetical protein [Azoarcus communis] |
| **08207** | 58 | D-alanyl-D-alanine carboxypeptidase/D-alanyl-D-alanine-endopeptidase [Azoarcus communis] |
| **07291** | 58 | NAD(P)H-binding protein [Denitromonas halophilus] |
| **07295** | 58 | methyl-accepting chemotaxis protein [Azoarcus communis] |
| **01536** | 58 | acetyl-CoA carboxylase carboxyltransferase subunit alpha [Azoarcus communis] |
| **01500** | 58 | nitrogen fixation protein NifM [Azoarcus communis] |
| **02607** | 58 | hypothetical protein [Azoarcus communis] |
| **05664** | 58 | ATP synthase subunit I [Azoarcus communis] |
| **01096** | 58 | PepSY domain-containing protein [Azoarcus indigens] |
| **09168** | 58 | barstar family protein [Azoarcus communis] |
| **06730** | 58 | bifunctional phosphoserine phosphatase/homoserine phosphotransferase ThrH [Azoarcus communis] |
| **00793** | 58 | hemerythrin domain-containing protein [Azoarcus communis] |
| **02336** | 58 | acetolactate synthase small subunit [Azoarcus communis] |
| **01334** | 58 | alpha/beta hydrolase [Denitromonas halophilus] |
| **07568** | 58 | ribosome small subunit-dependent GTPase A [Azoarcus communis] |
| **02550** | 58 | bifunctional adenosylcobinamide kinase/adenosylcobinamide-phosphate guanylyltransferase [Azoarcus communis] |
| **06578** | 58 | PilN domain-containing protein [Azoarcus communis] |
| **09811** | 58 | 23S rRNA (adenine(2030)-N(6))-methyltransferase RlmJ [Azoarcus communis] |
| **06697** | 58 | ZIP family metal transporter [Azoarcus indigens] |
| **05511** | 58 | GTP cyclohydrolase I FolE [Azoarcus communis] |
| **02139** | 58 | TetR family transcriptional regulator [Denitromonas halophilus] |
| **08513** | 58 | site-2 protease family protein [Azoarcus communis] |
| **10257** | 58 | aldehyde ferredoxin oxidoreductase family protein [Azoarcus communis] |
| **09792** | 58 | PDZ domain-containing protein [Azoarcus communis] |
| **01179** | 58 | D-hexose-6-phosphate mutarotase [Azoarcus communis] |
| **03048** | 58 | ribonuclease [Azoarcus communis] |
| **10381** | 58 | tRNA lysidine(34) synthetase TilS [Azoarcus communis] |
| **03913** | 58 | folate-binding protein YgfZ [Azoarcus communis] |
| **10305** | 58 | ATP-dependent protease subunit HslV [Azoarcus communis] |
| **10168** | 58 | MULTISPECIES: cell division protein FtsB [Azoarcus] |
| **05359** | 58 | alpha-E domain-containing protein [Azoarcus communis] |
| **04311** | 58 | tRNA (adenosine(37)-N6)-dimethylallyltransferase MiaA [Azoarcus communis] |
| **00749** | 58 | (2Fe-2S)-binding protein [Nitrogeniibacter aestuarii] |
| **04057** | 58 | DUF494 domain-containing protein [Azoarcus communis] |
| **04064** | 58 | 16S rRNA (guanine(966)-N(2))-methyltransferase RsmD [Azoarcus communis] |
| **03837** | 58 | ATP-dependent protease ATPase subunit HslU [Azoarcus communis] |
| **01351** | 58 | restriction endonuclease subunit S [Denitromonas halophilus] |
| **01368** | 58 | LUD domain-containing protein [Denitromonas halophilus] |
| **06830** | 57 | peptidoglycan DD-metalloendopeptidase family protein [Azoarcus communis] |
| **08804** | 57 | HDOD domain-containing protein [Azoarcus communis] |
| **01667** | 57 | sensor domain-containing diguanylate cyclase [Propionivibrio dicarboxylicus] |
| **04443** | 57 | energy transducer TonB [Azoarcus communis] |

| | | |
|---|---|---|
| **09962** | 57 | heme biosynthesis protein HemY [Azoarcus communis] |
| **09136** | 57 | phospholipase D-like domain-containing protein [Nitrogeniibacter aestuarii] |
| **09603** | 57 | serine protease [Azoarcus olearius] |
| **04188** | 57 | MbcA/ParS/Xre antitoxin family protein [Aromatoleum evansii] |
| **01756** | 57 | bifunctional aconitate hydratase 2/2-methylisocitrate dehydratase [Azoarcus communis] |
| **03207** | 57 | MULTISPECIES: flagellar motor switch protein FliN [Thauera] |
| **00530** | 57 | bifunctional tRNA (5-methylaminomethyl-2-thiouridine)(34)-methyltransferase MnmD/FAD-dependent 5-carboxymethylaminomethyl-2-thiouridine(34) oxidoreductase MnmC [Azoarcus communis] |
| **00537** | 57 | VF530 family protein [Denitromonas halophilus] |
| **00531** | 57 | iron-containing alcohol dehydrogenase [Propionivibrio dicarboxylicus] |
| **01224** | 57 | dienelactone hydrolase family protein [Azoarcus olearius] |
| **06216** | 57 | flagellar basal body P-ring formation protein FlgA [Azoarcus communis] |
| **09960** | 57 | cytochrome c oxidase subunit 3 [Azoarcus communis] |
| **08393** | 57 | ureidoglycolate lyase [Azoarcus communis] |
| **07059** | 57 | SRPBCC family protein [Aromatoleum toluclasticum] |
| **04379** | 57 | DNA-deoxyinosine glycosylase [Azoarcus communis] |
| **00572** | 57 | hypothetical protein [Azoarcus communis] |
| **04987** | 57 | MULTISPECIES: alkylphosphonate utilization protein [Azoarcus] |
| **08928** | 57 | MULTISPECIES: hypothetical protein [Thauera] |
| **01003** | 57 | bifunctional hydroxymethylpyrimidine kinase/phosphomethylpyrimidine kinase [Denitromonas halophilus] |
| **08929** | 57 | ABC transporter substrate-binding protein [Azoarcus communis] |
| **02494** | 57 | YceH family protein [Azoarcus communis] |
| **04235** | 57 | Crp/Fnr family transcriptional regulator [Azoarcus communis] |
| **02442** | 57 | GGDEF domain-containing protein [Denitromonas halophilus] |
| **03343** | 57 | DUF853 domain-containing protein [Azoarcus communis] |
| **10576** | 57 | ArsC family reductase [Azoarcus communis] |
| **10170** | 57 | hypothetical protein [Azoarcus communis] |
| **07314** | 57 | agmatinase [Aromatoleum petrolei] |
| **02376** | 57 | GNAT family N-acetyltransferase [Aromatoleum aromaticum] |
| **00075** | 57 | MFS transporter [Azoarcus communis] |
| **02339** | 57 | general secretion pathway protein GspD [Azoarcus communis] |
| **07985** | 57 | Co2+/Mg2+ efflux protein ApaG [Azoarcus communis] |
| **03273** | 57 | DUF1841 family protein [Azoarcus communis] |
| **05274** | 57 | adenosylcobinamide-GDP ribazoletransferase [Azoarcus communis] |
| **07597** | 57 | hypothetical protein [Aromatoleum toluclasticum] |
| **02193** | 57 | TerB family tellurite resistance protein [Denitromonas halophilus] |
| **06715** | 57 | pilus assembly protein PilP [Denitromonas halophilus] |
| **06709** | 57 | EscU/YscU/HrcU family type III secretion system export apparatus switch protein [Azoarcus communis] |
| **08613** | 57 | LapA family protein [Azoarcus communis] |
| **01743** | 57 | DUF1343 domain-containing protein [Azoarcus communis] |
| **02960** | 57 | NADPH-dependent 7-cyano-7-deazaguanine reductase QueF [Denitromonas halophilus] |
| **02022** | 57 | AmmeMemoRadiSam system protein A [Azoarcus communis] |
| **09192** | 57 | flagellar export protein FliJ [Azoarcus communis] |
| **05151** | 57 | sigma-E factor negative regulatory protein [Azoarcus communis] |
| **08857** | 57 | hypothetical protein [Azoarcus communis] |
| **00452** | 57 | flagellar hook protein FlgE [Azoarcus communis] |
| **03867** | 57 | NRDE family protein [Azoarcus communis] |
| **02109** | 57 | YciI family protein [Azoarcus communis] |
| **03820** | 57 | threonine synthase [Denitromonas halophilus] |
| **05849** | 57 | hypothetical protein [Azoarcus communis] |
| **02155** | 57 | hypothetical protein [Azoarcus communis] |
| **02083** | 57 | AmmeMemoRadiSam system protein B [Azoarcus communis] |
| **03539** | 57 | septation protein A [Azoarcus communis] |
| **01389** | 57 | methylmalonyl Co-A mutase-associated GTPase MeaB [Azoarcus communis] |
| **09081** | 56 | excinuclease ABC subunit UvrA [Azoarcus communis] |
| **00587** | 56 | GspH/FimT family pseudopilin [Aromatoleum toluclasticum] |
| **08785** | 56 | Uma2 family endonuclease [Thauera phenylacetica] |
| **06054** | 56 | 23S rRNA (cytidine(2498)-2'-O)-methyltransferase RlmM [Azoarcus communis] |
| **03247** | 56 | PilC/PilY family type IV pilus protein [Zoogloea ramigera] |
| **00810** | 56 | RNA-binding S4 domain-containing protein [Denitromonas halophilus] |
| **00330** | 56 | acyl-CoA dehydrogenase [Denitromonas halophilus] |
| **05962** | 56 | DUF4922 domain-containing protein [Azoarcus communis] |
| **08851** | 56 | cob(I)yrinic acid a,c-diamide adenosyltransferase [Azoarcus communis] |
| **08885** | 56 | flagellar protein FlaG [Azoarcus communis] |
| **10497** | 56 | nitrate reductase cytochrome c-type subunit [Azoarcus olearius] |
| **02153** | 56 | DUF58 domain-containing protein [Azoarcus communis] |
| **05242** | 56 | MarR family transcriptional regulator [Azoarcus communis] |
| **09939** | 56 | tRNA uridine-5-carboxymethylaminomethyl(34) synthesis GTPase MnmE [Denitromonas halophilus] |
| **01698** | 56 | zinc ABC transporter substrate-binding protein [Azoarcus indigens] |
| **06087** | 56 | MULTISPECIES: phenol hydroxylase subunit [Azoarcus] |
| **07217** | 56 | glycerate kinase [Denitromonas halophilus] |
| **07687** | 56 | insulinase family protein [Azoarcus communis] |
| **00396** | 56 | EI24 domain-containing protein [Azoarcus communis] |
| **10567** | 56 | beta-propeller fold lactonase family protein [Nitrogeniibacter aestuarii] |
| **10616** | 56 | transporter [Azoarcus indigens] |
| **05942** | 56 | insulinase family protein [Azoarcus communis] |
| **03641** | 56 | MULTISPECIES: flagellar motor switch protein FliM [Azoarcus] |
| **07811** | 56 | flagellar biosynthesis protein FlhF [Azoarcus communis] |
| **02885** | 56 | type II toxin-antitoxin system RelE/ParE family toxin [Aromatoleum toluclasticum] |
| **09923** | 56 | NYN domain-containing protein [Azoarcus communis] |
| **01583** | 56 | phenol hydroxylase subunit P4 [Aromatoleum toluclasticum] |
| **01795** | 56 | deoxynucleoside kinase [Azoarcus communis] |
| **00789** | 56 | permease-like cell division protein FtsX [Azoarcus communis] |
| **08208** | 56 | hypothetical protein [Azoarcus communis] |
| **05424** | 56 | MucB/RseB C-terminal domain-containing protein [Azoarcus communis] |
| **10190** | 56 | S-methyl-5-thioribose-1-phosphate isomerase [Azoarcus communis] |
| **00101** | 56 | STAS domain-containing protein [Azoarcus communis] |
| **07144** | 56 | MULTISPECIES: flagellar protein FlgN [Thauera] |
| **09103** | 56 | 2Fe-2S iron-sulfur cluster binding domain-containing protein [Aromatoleum toluclasticum] |
| **08547** | 56 | signal peptide peptidase SppA [Denitromonas halophilus] |
| **09650** | 56 | uroporphyrinogen-III C-methyltransferase [Azoarcus communis] |
| **08423** | 56 | chromate efflux transporter [Azoarcus communis] |
| **00525** | 56 | type II toxin-antitoxin system VapC family toxin [Rhodocyclus purpureus] |
| **02246** | 56 | phosphoribosyl-AMP cyclohydrolase [Azoarcus communis] |
| **01481** | 56 | MULTISPECIES: thiol peroxidase [Azoarcus] |

| | | |
|---|---|---|
| **05839** | 56 | cobyrinate a,c-diamide synthase [Azoarcus communis] |
| **03739** | 56 | MULTISPECIES: cob(I)yrinic acid a,c-diamide adenosyltransferase [Thauera] |
| **04135** | 56 | glutamate--cysteine ligase [Denitromonas halophilus] |
| **09177** | 56 | flagellar biosynthetic protein FliO [Azoarcus communis] |
| **07755** | 56 | TonB C-terminal domain-containing protein [Azoarcus communis] |
| **08571** | 56 | SIMPL domain-containing protein [Azoarcus communis] |
| **07472** | 56 | phosphoenolpyruvate carboxykinase (GTP) [Azoarcus communis] |
| **02970** | 56 | DUF3426 domain-containing protein, partial [Denitromonas halophilus] |
| **03756** | 56 | succinyl-diaminopimelate desuccinylase [Azoarcus communis] |
| **00840** | 55 | TIGR00645 family protein [Azoarcus communis] |
| **08218** | 55 | DUF3579 domain-containing protein [Azoarcus communis] |
| **10192** | 55 | transaldolase [Azoarcus communis] |
| **08632** | 55 | M48 family metallopeptidase [Denitromonas halophilus] |
| **08311** | 55 | DUF3617 family protein [Denitromonas halophilus] |
| **03030** | 55 | M14-type cytosolic carboxypeptidase [Aromatoleum toluclasticum] |
| **01636** | 55 | MULTISPECIES: aromatic-ring-hydroxylating dioxygenase subunit beta [Rhodocyclales] |
| **06509** | 55 | ABC-type transport auxiliary lipoprotein family protein [Azoarcus communis] |
| **01572** | 55 | Hpt domain-containing protein [Azoarcus communis] |
| **08487** | 55 | DOMON-like domain-containing protein [Azoarcus communis] |
| **08645** | 55 | MULTISPECIES: hypothetical protein [Azoarcus] |
| **03871** | 55 | homoserine kinase [Azoarcus communis] |
| **08508** | 55 | MULTISPECIES: DUF1328 domain-containing protein [Azoarcus] |
| **09238** | 55 | site-specific tyrosine recombinase XerD [Azoarcus communis] |
| **05553** | 55 | c-type cytochrome [Denitromonas halophilus] |
| **03759** | 55 | bifunctional 3-phosphoshikimate 1-carboxyvinyltransferase/cytidylate kinase [Azoarcus communis] |
| **09584** | 55 | haloacid dehalogenase type II [Denitromonas halophilus] |
| **01737** | 55 | MULTISPECIES: cytochrome c [Thauera] |
| **05389** | 55 | acetyl-CoA hydrolase/transferase family protein [Azoarcus communis] |
| **07551** | 55 | dynamin family protein [Azoarcus communis] |
| **09637** | 55 | hypothetical protein [Azoarcus communis] |
| **01243** | 55 | MinD/ParA family protein [Denitromonas halophilus] |
| **03858** | 55 | DUF1178 family protein [Azoarcus communis] |
| **05382** | 55 | glycolate oxidase subunit GlcE [Azoarcus communis] |
| **06890** | 55 | AraC family transcriptional regulator [Azoarcus communis] |
| **02102** | 55 | glycerophosphodiester phosphodiesterase [Azoarcus communis] |
| **00229** | 55 | nitric oxide reductase [Denitromonas halophilus] |
| **09014** | 55 | ornithine cyclodeaminase [Denitromonas halophilus] |
| **02615** | 55 | nucleotidyltransferase family protein [Azoarcus communis] |
| **04300** | 55 | ribonuclease P protein component [Azoarcus communis] |
| **03943** | 55 | hypothetical protein [Azoarcus communis] |
| **09135** | 55 | YihY/virulence factor BrkB family protein [Azoarcus indigens] |
| **06184** | 55 | YaiI/YqxD family protein [Azoarcus communis] |
| **02430** | 55 | translation initiation factor IF-3 [Denitromonas halophilus] |
| **04559** | 54 | flagellar assembly peptidoglycan hydrolase FlgJ [Azoarcus communis] |
| **08830** | 54 | PACE efflux transporter [Azoarcus communis] |
| **00743** | 54 | CNP1-like family protein [Azoarcus communis] |
| **00164** | 54 | glutamine--tRNA ligase/YqeY domain fusion protein [Azoarcus communis] |
| **06674** | 54 | leucyl aminopeptidase [Denitromonas halophilus] |
| **03576** | 54 | MULTISPECIES: flagellar biosynthesis anti-sigma factor FlgM [Thauera] |
| **05913** | 54 | DmsE family decaheme c-type cytochrome [Aromatoleum toluclasticum] |
| **01580** | 54 | glutaredoxin family protein [Azoarcus communis] |
| **05423** | 54 | mandelate racemase/muconate lactonizing enzyme family protein [Azoarcus indigens] |
| **00132** | 54 | flagellar hook-length control protein FliK [Azoarcus communis] |
| **02051** | 54 | flagellin [Azoarcus communis] |
| **05369** | 54 | class I SAM-dependent methyltransferase [Azoarcus communis] |
| **02260** | 54 | host attachment protein [Azoarcus communis] |
| **06739** | 54 | MULTISPECIES: sulfoxide reductase heme-binding subunit YedZ [Azoarcus] |
| **01539** | 54 | outer membrane lipoprotein-sorting protein [Azoarcus indigens] |
| **06234** | 54 | MULTISPECIES: flagellar transcriptional regulator FlhD [Aromatoleum] |
| **10298** | 54 | DUF1857 family protein [Azoarcus communis] |
| **09303** | 54 | uracil-xanthine permease family protein [Azoarcus communis] |
| **10267** | 54 | YdcF family protein [Azoarcus communis] |
| **02635** | 54 | hypothetical protein [Azoarcus communis] |
| **05749** | 54 | ATP-dependent RecD-like DNA helicase [Aromatoleum petrolei] |
| **03056** | 54 | DUF3488 and transglutaminase-like domain-containing protein [Azoarcus communis] |
| **00040** | 54 | alpha/beta hydrolase [Denitromonas halophilus] |
| **07403** | 54 | AI-2E family transporter [Aromatoleum diolicum] |
| **08352** | 54 | OsmC domain/YcaO domain-containing protein [Azoarcus communis] |
| **00860** | 54 | VWA domain-containing protein [Azoarcus communis] |
| **10238** | 54 | DNA-3-methyladenine glycosylase I [Azoarcus indigens] |
| **09932** | 54 | MULTISPECIES: nucleoside diphosphate kinase regulator [Thauera] |
| **02482** | 54 | DUF6394 family protein [Denitromonas halophilus] |
| **09237** | 54 | translocation/assembly module TamB domain-containing protein [Denitromonas halophilus] |
| **07101** | 54 | chromosome segregation protein SMC [Azoarcus communis] |
| **05888** | 54 | peptidoglycan DD-metalloendopeptidase family protein [Azoarcus communis] |
| **10389** | 53 | LON peptidase substrate-binding domain-containing protein [Azoarcus communis] |
| **05271** | 53 | orotidine-5'-phosphate decarboxylase [Azoarcus communis] |
| **08561** | 53 | hypothetical protein [Azoarcus communis] |
| **02063** | 53 | SPOR domain-containing protein [Azoarcus communis] |
| **01601** | 53 | SOS response-associated peptidase [Thauera propionica] |
| **10534** | 53 | polyphosphate:AMP phosphotransferase [Azoarcus communis] |
| **02437** | 53 | thiosulfohydrolase SoxB [Azoarcus communis] |
| **09762** | 53 | dethiobiotin synthase [Azoarcus communis] |
| **04068** | 53 | D-aminoacyl-tRNA deacylase [Azoarcus communis] |
| **05304** | 53 | transglutaminase family protein [Azoarcus communis] |
| **08535** | 53 | hypothetical protein [Azoarcus communis] |
| **02667** | 53 | hypothetical protein [Azoarcus communis] |
| **03023** | 53 | chloride channel protein [Azoarcus communis] |
| **04997** | 53 | ADP-ribosylglycohydrolase family protein [Azoarcus communis] |
| **10111** | 53 | flagellar assembly protein FliW [Azoarcus communis] |
| **00725** | 53 | MULTISPECIES: periplasmic Cu(I)/Cu(II)-binding protein CopK [Rhodocyclales] |
| **00067** | 53 | glucokinase [Azoarcus indigens] |
| **09110** | 53 | heme o synthase [Azoarcus indigens] |
| **02599** | 53 | DUF2478 domain-containing protein [Azoarcus communis] |

| | | |
|---|---|---|
| **01395** | 53 | NAD-binding protein [Denitromonas halophilus] |
| **05485** | 53 | hemin uptake protein HemP [Azoarcus communis] |
| **07612** | 53 | hypothetical protein [Azoarcus communis] |
| **00402** | 53 | TetR/AcrR family transcriptional regulator [Aromatoleum toluclasticum] |
| **00691** | 53 | NADP-dependent isocitrate dehydrogenase [Azoarcus communis] |
| **07455** | 53 | class I SAM-dependent methyltransferase [Azoarcus communis] |
| **09277** | 53 | translesion DNA synthesis-associated protein ImuA [Azoarcus communis] |
| **06115** | 53 | alpha/beta fold hydrolase [Denitromonas halophilus] |
| **03337** | 53 | FAD-dependent oxidoreductase [Azoarcus indigens] |
| **04175** | 53 | DUF1415 domain-containing protein [Azoarcus communis] |
| **03268** | 53 | hydrolase [Azoarcus communis] |
| **05411** | 52 | SpoVR family protein [Azoarcus communis] |
| **06310** | 52 | YeaH/YhbH family protein [Azoarcus communis] |
| **02533** | 52 | YgiQ family radical SAM protein [Azoarcus communis] |
| **01710** | 52 | HPF/RaiA family ribosome-associated protein [Azoarcus communis] |
| **06321** | 52 | MULTISPECIES: DUF2892 domain-containing protein [Aromatoleum] |
| **08919** | 52 | MATE family efflux transporter [Azoarcus indigens] |
| **04215** | 52 | hypothetical protein [Azoarcus communis] |
| **04804** | 52 | MULTISPECIES: type VI secretion system contractile sheath large subunit [Niveibacterium] |
| **04859** | 52 | energy-coupling factor ABC transporter permease [Azoarcus communis] |
| **02522** | 52 | type VI secretion system baseplate subunit TssG [Azoarcus communis] |
| **09043** | 52 | bifunctional hydroxymethylpyrimidine kinase/phosphomethylpyrimidine kinase [Azoarcus communis] |
| **04071** | 52 | PrkA family serine protein kinase [Azoarcus communis] |
| **00182** | 52 | cyclase family protein [Azoarcus communis] |
| **07997** | 52 | glutamine-hydrolyzing carbamoyl-phosphate synthase small subunit [Denitromonas halophilus] |
| **10450** | 52 | 3'-5' exonuclease [Azoarcus communis] |
| **10191** | 52 | putative sulfate exporter family transporter [Aromatoleum toluclasticum] |
| **03034** | 52 | DUF3301 domain-containing protein [Azoarcus communis] |
| **05189** | 52 | hypothetical protein [Denitromonas halophilus] |
| **09614** | 52 | type VI secretion system tube protein Hcp [Azoarcus communis] |
| **05740** | 52 | polyamine aminopropyltransferase [Azoarcus communis] |
| **06103** | 52 | 3'-5' exonuclease [Denitromonas halophilus] |
| **02037** | 52 | MULTISPECIES: recombinase family protein [Proteobacteria] |
| **01411** | 52 | ferredoxin III, nif-specific [Azoarcus communis] |
| **04301** | 52 | DNA polymerase III subunit epsilon [Azoarcus communis] |
| **05103** | 52 | 4Fe-4S binding protein [Azoarcus communis] |
| **02268** | 52 | 50S ribosomal protein L10 [Denitromonas halophilus] |
| **07805** | 52 | TRAP transporter small permease [Azoarcus communis] |
| **07084** | 52 | allantoicase [Azoarcus communis] |
| **07517** | 52 | ethylbenzene dehydrogenase-related protein [Denitromonas halophilus] |
| **08932** | 52 | acylphosphatase [Denitromonas halophilus] |
| **02546** | 51 | RNA 2',3'-cyclic phosphodiesterase [Denitromonas halophilus] |
| **01185** | 51 | glycerol-3-phosphate dehydrogenase/oxidase [Azoarcus communis] |
| **02987** | 51 | xanthine dehydrogenase small subunit [Azoarcus communis] |
| **07581** | 51 | universal stress protein [Aromatoleum toluclasticum] |
| **01745** | 51 | GNAT family N-acetyltransferase [Azoarcus communis] |
| **10440** | 51 | nuclear transport factor 2 family protein [Azoarcus communis] |
| **07541** | 51 | type VI secretion system baseplate subunit TssF [Azoarcus communis] |
| **09500** | 51 | FAD-binding oxidoreductase [Denitromonas halophilus] |
| **05548** | 51 | type VI secretion system membrane subunit TssM [Azoarcus communis] |
| **10028** | 51 | flagellin FliC [Denitromonas halophilus] |
| **03064** | 51 | MULTISPECIES: chaperone NapD [Thauera] |
| **00862** | 51 | MULTISPECIES: hydrogenase maturation nickel metallochaperone HypA [Azoarcus] |
| **10609** | 51 | hypothetical protein [Azoarcus communis] |
| **08828** | 51 | glutamine amidotransferase [Denitromonas halophilus] |
| **01336** | 51 | septum site-determining protein MinD [Denitromonas halophilus] |
| **01657** | 51 | 3-carboxyethylcatechol 2,3-dioxygenase [Azoarcus communis] |
| **07572** | 51 | hypothetical protein [Azoarcus communis] |
| **06814** | 51 | MFS transporter [Azoarcus indigens] |
| **06155** | 51 | hypothetical protein [Azoarcus communis] |
| **06399** | 51 | MFS transporter [Azoarcus indigens] |
| **01565** | 51 | ATP-binding domain-containing protein [Aromatoleum toluclasticum] |
| **07599** | 51 | uracil phosphoribosyltransferase [Azoarcus communis] |
| **02382** | 51 | septal ring lytic transglycosylase RlpA family protein [Azoarcus communis] |
| **05645** | 51 | LysM peptidoglycan-binding domain-containing protein [Azoarcus communis] |
| **05580** | 51 | type II secretion system secretin GspD [Azoarcus indigens] |
| **02295** | 51 | ABC transporter permease [Azoarcus communis] |
| **03547** | 51 | MtrB/PioB family decaheme-associated outer membrane protein [Aromatoleum toluclasticum] |
| **03722** | 51 | DUF3079 domain-containing protein [Denitromonas halophilus] |
| **00892** | 50 | thiosulfate oxidation carrier protein SoxY [Azoarcus communis] |
| **10007** | 50 | TPM domain-containing protein [Azoarcus communis] |
| **01735** | 50 | putative selenate ABC transporter substrate-binding protein [Denitromonas halophilus] |
| **10388** | 50 | DUF167 family protein [Denitromonas halophilus] |
| **05142** | 50 | c-type cytochrome [Aromatoleum toluclasticum] |
| **03677** | 50 | HigA family addiction module antitoxin [Nitrogeniibacter aestuarii] |
| **05565** | 50 | HypC/HybG/HupF family hydrogenase formation chaperone [Azoarcus communis] |
| **08944** | 50 | ProQ/FINO family protein [Azoarcus communis] |
| **04033** | 50 | 5,6-dimethylbenzimidazole synthase [Azoarcus communis] |
| **07733** | 50 | COX15/CtaA family protein [Azoarcus communis] |
| **07063** | 50 | MULTISPECIES: phasin family protein [Thauera] |
| **01556** | 50 | ACP phosphodiesterase [Azoarcus communis] |
| **08733** | 50 | MgtC/SapB family protein [Azoarcus communis] |
| **10016** | 50 | universal stress protein [Aromatoleum toluclasticum] |
| **06816** | 50 | helix-turn-helix transcriptional regulator [Azoarcus indigens] |
| **07947** | 50 | malonyl-CoA decarboxylase [Azoarcus communis] |
| **01553** | 50 | catalase/peroxidase HPI [Azoarcus communis] |
| **10516** | 50 | SURF1 family protein [Azoarcus communis] |
| **09804** | 50 | transporter [Aromatoleum aromaticum] |
| **01507** | 50 | hydrogenase expression/formation protein HypE [Azoarcus communis] |
| **00210** | 50 | Lrp/AsnC family transcriptional regulator [Denitromonas halophilus] |
| **09907** | 50 | cytochrome c oxidase assembly protein [Azoarcus communis] |
| **08134** | 50 | type VI secretion system baseplate subunit TssK [Azoarcus communis] |
| **05462** | 50 | heptose kinase [Azoarcus communis] |
| **06534** | 50 | type VI secretion system protein TssA [Azoarcus communis] |

| | | |
|---|---|---|
| 06065 | 50 | hypothetical protein [Azoarcus communis] |
| 03753 | 50 | MULTISPECIES: DNA-directed RNA polymerase subunit omega [Azoarcus] |
| 05815 | 50 | hypothetical protein [Aromatoleum toluclasticum] |
| 05016 | 50 | carboxymuconolactone decarboxylase family protein [Denitromonas halophilus] |
| 02502 | 49 | DUF2249 domain-containing protein [Denitromonas halophilus] |
| 02189 | 49 | thiosulfate oxidation carrier complex protein SoxZ [Azoarcus communis] |
| 02694 | 49 | DUF2802 domain-containing protein [Azoarcus communis] |
| 03809 | 49 | MaoC family dehydratase N-terminal domain-containing protein [Azoarcus indigens] |
| 03907 | 49 | hypothetical protein [Azoarcus communis] |
| 04584 | 49 | DNA polymerase Y family protein [Azoarcus communis] |
| 07273 | 49 | VanZ family protein [Aromatoleum aromaticum] |
| 02781 | 49 | integrase family protein [Aromatoleum bremense] |
| 08370 | 49 | CopD family protein [Azoarcus communis] |
| 02566 | 49 | fatty acid desaturase [Denitromonas halophilus] |
| 03733 | 49 | long-chain N-acyl amino acid synthase [Azoarcus communis] |
| 04362 | 49 | IS21 family transposase [Aromatoleum aromaticum] |
| 02251 | 49 | hypothetical protein [Azoarcus communis] |
| 02068 | 49 | GlsB/YeaQ/YmgE family stress response membrane protein [Azoarcus communis] |
| 09307 | 49 | Trk system potassium transporter TrkA [Denitromonas halophilus] |
| 04828 | 49 | hypothetical protein [Azoarcus communis] |
| 05384 | 49 | CoA transferase subunit A [Aromatoleum diolicum] |
| 09995 | 49 | flagellar protein FliT [Azoarcus communis] |
| 00219 | 49 | NAD(P)H-dependent oxidoreductase [Aromatoleum buckelii] |
| 05936 | 49 | extracellular solute-binding protein [Aromatoleum toluclasticum] |
| 06544 | 49 | DMT family protein [Azoarcus communis] |
| 07207 | 49 | YihY family inner membrane protein [Denitromonas halophilus] |
| 01381 | 49 | phosphonate ABC transporter, permease protein PhnE [Azoarcus nasutitermitis] |
| 01346 | 49 | arsenate reductase (glutaredoxin) [Azoarcus indigens] |
| 05170 | 49 | hypothetical protein [Denitromonas halophilus] |
| 00586 | 49 | type VI secretion system baseplate subunit TssE [Azoarcus communis] |
| 05306 | 49 | TIGR03087 family PEP-CTERM/XrtA system glycosyltransferase [Azoarcus communis] |
| 05355 | 49 | alkaline phosphatase family protein [Azoarcus communis] |
| 05370 | 49 | D-glycero-beta-D-manno-heptose-7-phosphate kinase [Azoarcus communis] |
| 09895 | 49 | Crp/Fnr family transcriptional regulator [Aromatoleum diolicum] |
| 06853 | 49 | alpha/beta fold hydrolase [Denitromonas halophilus] |
| 06707 | 49 | hypothetical protein [Azoarcus communis] |
| 01909 | 48 | transglutaminase-like cysteine peptidase [Azoarcus communis] |
| 04172 | 48 | c-type cytochrome [Aromatoleum toluclasticum] |
| 00475 | 48 | hypothetical protein [Denitromonas halophilus] |
| 09990 | 48 | glycosyltransferase family 39 protein [Azoarcus communis] |
| 02118 | 48 | hypothetical protein [Azoarcus communis] |
| 01291 | 48 | LysR family transcriptional regulator [Azoarcus communis] |
| 03168 | 48 | TPM domain-containing protein [Azoarcus communis] |
| 06987 | 48 | 2,3-diphosphoglycerate-dependent phosphoglycerate mutase [Denitromonas halophilus] |
| 02489 | 48 | NfeD family protein [Azoarcus communis] |
| 06984 | 48 | formylglycine-generating enzyme family protein [Denitromonas halophilus] |
| 07584 | 48 | phage holin family protein [Azoarcus communis] |
| 02290 | 48 | MULTISPECIES: YjfB family protein [Azoarcus] |
| 03372 | 48 | FemAB family PEP-CTERM system-associated protein [Azoarcus communis] |
| 05250 | 48 | MULTISPECIES: quaternary ammonium compound efflux SMR transporter QacE delta 1 [Bacteria] |
| 02661 | 48 | radical SAM protein [Azoarcus communis] |
| 00821 | 48 | polymer-forming cytoskeletal protein [Azoarcus communis] |
| 09577 | 48 | DUF2946 family protein [Azoarcus communis] |
| 03035 | 48 | hydrogenase formation protein HypD [Azoarcus communis] |
| 06129 | 48 | type VI secretion system contractile sheath small subunit [Azoarcus communis] |
| 09264 | 48 | MULTISPECIES: phage tail protein [Aromatoleum] |
| 09657 | 48 | glycosyltransferase family 4 protein [Denitromonas halophilus] |
| 08489 | 48 | class I SAM-dependent methyltransferase [Azoarcus communis] |
| 08355 | 48 | PAS domain S-box protein [Nitrogeniibacter aestuarii] |
| 08709 | 48 | thioesterase family protein [Azoarcus communis] |
| 03331 | 48 | hypothetical protein [Azoarcus communis] |
| 03149 | 48 | MarC family protein [Denitromonas halophilus] |
| 01403 | 48 | hypothetical protein [Azoarcus communis] |
| 04080 | 48 | serine/threonine protein kinase [Azoarcus communis] |
| 06371 | 48 | chorismate lyase [Denitromonas halophilus] |
| 03724 | 48 | extracellular solute-binding protein [Azoarcus communis] |
| 04171 | 48 | BCCT family transporter [Nitrogeniibacter aestuarii] |
| 04372 | 47 | phenylacetate-CoA oxygenase subunit PaaJ [Azoarcus communis] |
| 09045 | 47 | DUF4936 family protein [Azoarcus communis] |
| 06239 | 47 | CoA-binding protein [Azoarcus communis] |
| 07445 | 47 | exodeoxyribonuclease V subunit gamma [Azoarcus communis] |
| 04391 | 47 | 1,2-phenylacetyl-CoA epoxidase subunit A [Azoarcus communis] |
| 05015 | 47 | MULTISPECIES: 1,2-phenylacetyl-CoA epoxidase subunit B [Rhodocyclales] |
| 02271 | 47 | TIGR03016 family PEP-CTERM system-associated outer membrane protein [Azoarcus communis] |
| 01608 | 47 | MULTISPECIES: nitrogenase iron protein [Niveibacterium] |
| 03065 | 47 | XrtA-associated tyrosine autokinase [Azoarcus communis] |
| 10279 | 47 | putative O-glycosylation ligase, exosortase A system-associated [Azoarcus communis] |
| 06212 | 47 | Na+/H+ antiporter subunit G [Azoarcus communis] |
| 03792 | 47 | CpaF family protein [Aromatoleum toluclasticum] |
| 02962 | 47 | exosortase A [Azoarcus communis] |
| 07015 | 47 | hypothetical protein [Azoarcus communis] |
| 03651 | 47 | MULTISPECIES: hypothetical protein [Thauera] |
| 09150 | 47 | flavodoxin [Azoarcus indigens] |
| 09746 | 47 | HDOD domain-containing protein [Denitromonas halophilus] |
| 01284 | 47 | cytochrome P450 [Aromatoleum toluclasticum] |
| 02147 | 47 | MULTISPECIES: Spy/CpxP family protein refolding chaperone [Azoarcus] |
| 10544 | 47 | fructosamine kinase family protein [Denitromonas halophilus] |
| 05880 | 47 | DNA recombination protein RmuC [Azoarcus communis] |
| 00326 | 47 | MULTISPECIES: DUF2933 domain-containing protein [Zoogloeaceae] |
| 09478 | 47 | MULTISPECIES: phenylacetate-CoA oxygenase subunit PaaC [Azoarcus] |
| 00152 | 47 | putative Na+/H+ antiporter [Azoarcus indigens] |
| 03932 | 47 | polymer-forming cytoskeletal protein [Denitromonas halophilus] |
| 02825 | 47 | HprK-related kinase A [Azoarcus communis] |
| 09996 | 47 | calcium/sodium antiporter [Denitromonas halophilus] |

| | | |
|---|---|---|
| 02470 | 47 | Na+/H+ antiporter subunit E [Azoarcus communis] |
| 08380 | 47 | DUF1631 domain-containing protein [Aromatoleum toluclasticum] |
| 08892 | 47 | hydroxyphenylacetyl-CoA thioesterase PaaI [Azoarcus communis] |
| 08312 | 47 | MipA/OmpV family protein [Azoarcus communis] |
| 08308 | 47 | hypothetical protein [Azoarcus communis] |
| 01688 | 47 | SAM-dependent methyltransferase [Azoarcus communis] |
| 05499 | 47 | S-formylglutathione hydrolase [Azoarcus communis] |
| 04165 | 46 | hypothetical protein [Denitromonas halophilus] |
| 06013 | 46 | allantoinase PuuE [Azoarcus communis] |
| 03193 | 46 | pentapeptide repeat-containing protein [Denitromonas halophilus] |
| 02882 | 46 | nucleotidyltransferase family protein [Azoarcus communis] |
| 05681 | 46 | sugar ABC transporter permease [Uliginosibacterium gangwonense] |
| 04927 | 46 | PEP-CTERM system histidine kinase PrsK [Azoarcus communis] |
| 05200 | 46 | tetratricopeptide repeat protein [Nitrogeniibacter aestuarii] |
| 06309 | 46 | CreA family protein [Azoarcus communis] |
| 01676 | 46 | zf-HC2 domain-containing protein [Denitromonas halophilus] |
| 05162 | 46 | flavin reductase family protein [Azoarcus indigens] |
| 05788 | 46 | DUF1924 domain-containing protein [Azoarcus communis] |
| 02938 | 46 | hypothetical protein [Azoarcus indigens] |
| 04077 | 46 | (2Fe-2S)-binding protein [Denitromonas halophilus] |
| 09202 | 46 | nucleoside recognition domain-containing protein [Denitromonas halophilus] |
| 06593 | 46 | DUF839 domain-containing protein [Azoarcus communis] |
| 10358 | 46 | DUF86 domain-containing protein [Azoarcus communis] |
| 00099 | 46 | DUF924 domain-containing protein [Azoarcus communis] |
| 05354 | 46 | Hsp70 family protein [Azoarcus communis] |
| 02392 | 46 | aromatic ring-hydroxylating dioxygenase subunit alpha [Nitrogeniibacter aestuarii] |
| 05293 | 46 | diheme cytochrome c [Azoarcus communis] |
| 01060 | 46 | BMP family ABC transporter substrate-binding protein [Azoarcus communis] |
| 02364 | 46 | MULTISPECIES: TerC family protein [Niveibacterium] |
| 06442 | 46 | NAD(P)H-dependent oxidoreductase [Azoarcus communis] |
| 06138 | 46 | DUF2789 domain-containing protein [Denitromonas halophilus] |
| 05853 | 45 | hypothetical protein [Azoarcus communis] |
| 00545 | 45 | YqjK-like family protein [Azoarcus communis] |
| 04319 | 45 | NAD(P)-dependent oxidoreductase [Uliginosibacterium gangwonense] |
| 06914 | 45 | PilZ domain-containing protein [Azoarcus communis] |
| 04748 | 45 | putative addiction module antidote protein [Aromatoleum buckelii] |
| 00883 | 45 | MFS transporter [Nitrogeniibacter aestuarii] |
| 03537 | 45 | DUF3820 family protein [Azoarcus communis] |
| 08864 | 45 | MBL fold metallo-hydrolase [Azoarcus communis] |
| 09675 | 45 | archaetidylserine decarboxylase [Azoarcus communis] |
| 01686 | 45 | PstS family phosphate ABC transporter substrate-binding protein [Denitromonas halophilus] |
| 01714 | 45 | AAA family ATPase [Azoarcus communis] |
| 02672 | 45 | Cd(II)/Pb(II)-responsive transcriptional regulator [Denitromonas halophilus] |
| 05636 | 45 | U32 family peptidase [Azoarcus communis] |
| 02075 | 45 | DUF1778 domain-containing protein [Azoarcus communis] |
| 01075 | 45 | M23 family metallopeptidase [Azoarcus communis] |
| 04295 | 45 | PepSY domain-containing protein [Aromatoleum aromaticum] |
| 00878 | 45 | L,D-transpeptidase [Azoarcus communis] |
| 02895 | 45 | glycine zipper 2TM domain-containing protein [Denitromonas halophilus] |
| 03845 | 45 | DUF2970 domain-containing protein [Azoarcus communis] |
| 02531 | 45 | pyrroloquinoline quinone biosynthesis peptide chaperone PqqD [Denitromonas halophilus] |
| 04747 | 45 | DNA starvation/stationary phase protection protein [Azoarcus communis] |
| 05649 | 45 | transglycosylase domain-containing protein [Azoarcus communis] |
| 04652 | 45 | DUF2946 domain-containing protein [Azoarcus communis] |
| 03941 | 45 | transcription regulator [Azoarcus communis] |
| 04675 | 45 | RES family NAD+ phosphorylase [Aromatoleum toluclasticum] |
| 02273 | 45 | MULTISPECIES: hypothetical protein [Thauera] |
| 02867 | 45 | MULTISPECIES: hypothetical protein [Rhodocyclales] |
| 09344 | 45 | MULTISPECIES: conjugative transfer ATPase [Rhodocyclales] |
| 01926 | 45 | TetR family transcriptional regulator [Aromatoleum toluclasticum] |
| 00032 | 45 | precorrin-4 C(11)-methyltransferase [Azoarcus communis] |
| 04156 | 45 | nitrogen fixation protein NifQ [Azoarcus communis] |
| 09129 | 45 | phospholipase A [Azoarcus communis] |
| 06294 | 45 | DUF3619 family protein [Denitromonas halophilus] |
| 05706 | 44 | energy-coupling factor ABC transporter permease [Azoarcus communis] |
| 07616 | 44 | hypothetical protein [Azoarcus communis] |
| 01648 | 44 | hypothetical protein [Azoarcus communis] |
| 08832 | 44 | hypothetical protein [Azoarcus communis] |
| 01841 | 44 | fumarylacetoacetate hydrolase family protein [Azoarcus communis] |
| 08964 | 44 | ISAs1 family transposase [Zoogloea ramigera] |
| 04098 | 44 | IS66 family transposase [Thauera aromatica] |
| 07658 | 44 | benzoyl-CoA reductase, bzd-type, subunit N [Aromatoleum toluclasticum] |
| 00274 | 44 | sulfite exporter TauE/SafE family protein [Denitromonas halophilus] |
| 03521 | 44 | aspartate/glutamate racemase family protein [Azoarcus communis] |
| 03437 | 44 | membrane-bound lytic murein transglycosylase MltF [Azoarcus communis] |
| 00693 | 44 | NADPH-dependent F420 reductase [Aromatoleum toluclasticum] |
| 03804 | 44 | MULTISPECIES: carbohydrate ABC transporter permease [Niveibacterium] |
| 01110 | 44 | DUF4197 domain-containing protein [Azoarcus communis] |
| 07439 | 44 | zinc metallopeptidase [Denitromonas halophilus] |
| 00689 | 44 | quinoprotein dehydrogenase-associated SoxYZ-like carrier [Nitrogeniibacter aestuarii] |
| 00556 | 44 | DUF1289 domain-containing protein [Azoarcus communis] |
| 10159 | 44 | thioredoxin family protein [Azoarcus communis] |
| 08991 | 44 | L-serine ammonia-lyase [Aromatoleum toluclasticum] |
| 00199 | 44 | PAS domain-containing protein [Denitromonas halophilus] |
| 08438 | 44 | site-specific recombinase [Azoarcus communis] |
| 03596 | 44 | DUF2783 domain-containing protein [Azoarcus communis] |
| 02623 | 43 | hydrogenase maturation protease [Azoarcus communis] |
| 05634 | 43 | cytochrome c [Denitromonas halophilus] |
| 02773 | 43 | MULTISPECIES: nitrogenase-stabilizing/protective protein NifW [Niveibacterium] |
| 06915 | 43 | beta-ketoacyl synthase chain length factor [Azoarcus communis] |
| 07927 | 43 | AraC family transcriptional regulator [Azoarcus rhizosphaerae] |
| 07168 | 43 | urease subunit beta [Azoarcus communis] |
| 02292 | 43 | bifunctional isocitrate dehydrogenase kinase/phosphatase [Aromatoleum toluclasticum] |
| 07978 | 43 | 1-acyl-sn-glycerol-3-phosphate acyltransferase [Azoarcus communis] |

| | | |
|---|---|---|
| 09240 | 43 | acyl-CoA synthetase [Azoarcus communis] |
| 00716 | 43 | EamA family transporter RarD [Denitromonas halophilus] |
| 00576 | 43 | MFS transporter [Aromatoleum toluclasticum] |
| 06148 | 43 | virulence RhuM family protein [Azoarcus communis] |
| 00652 | 43 | O-acetyl-ADP-ribose deacetylase [Azoarcus olearius] |
| 00568 | 43 | group II intron reverse transcriptase/maturase [Thauera humireducens] |
| 05210 | 43 | NifX-associated nitrogen fixation protein [Azoarcus communis] |
| 04343 | 43 | benzoate/H(+) symporter BenE family transporter [Azoarcus communis] |
| 07672 | 43 | TRAP transporter small permease [Propionivibrio dicarboxylicus] |
| 00972 | 43 | urease accessory protein UreF [Azoarcus communis] |
| 02979 | 43 | hypothetical protein [Azoarcus communis] |
| 00621 | 43 | MULTISPECIES: HlyD family efflux transporter periplasmic adaptor subunit [Thauera] |
| 05772 | 43 | MMPL family transporter [Azoarcus communis] |
| 07852 | 43 | AMP-binding protein [Azoarcus communis] |
| 08282 | 43 | MULTISPECIES: acyl carrier protein [Niveibacterium] |
| 02308 | 43 | nuclear transport factor 2 family protein [Azoarcus communis] |
| 08801 | 43 | prenyltransferase [Azoarcus communis] |
| 02398 | 43 | TRAP transporter substrate-binding protein DctP [Denitromonas halophilus] |
| 00454 | 43 | nitrogenase cofactor biosynthesis protein NifB [Azoarcus communis] |
| 01392 | 43 | urease subunit gamma [Azoarcus communis] |
| 05343 | 43 | helix-turn-helix transcriptional regulator [Aromatoleum toluclasticum] |
| 05641 | 43 | tripartite tricarboxylate transporter TctB family protein [Propionivibrio dicarboxylicus] |
| 02458 | 43 | cytochrome c [Denitromonas halophilus] |
| 05578 | 43 | large conductance mechanosensitive channel protein MscL [Denitromonas halophilus] |
| 01823 | 43 | DNA ligase [Azoarcus communis] |
| 01671 | 43 | MULTISPECIES: dodecin family protein [Aromatoleum] |
| 04838 | 43 | precorrin-6y C5,15-methyltransferase (decarboxylating) subunit CbiE [Denitromonas halophilus] |
| 00006 | 43 | MOSC domain-containing protein [Denitromonas halophilus] |
| 01705 | 43 | SPOR domain-containing protein [Azoarcus communis] |
| 10438 | 43 | methyl-accepting chemotaxis protein [Nitrogeniibacter aestuarii] |
| 08381 | 43 | lipoprotein [Azoarcus communis] |
| 05084 | 42 | nitroreductase [Aromatoleum toluclasticum] |
| 09053 | 42 | GspH/FimT family pseudopilin [Aromatoleum pumilum] |
| 00336 | 42 | MULTISPECIES: PepSY domain-containing protein [Thauera] |
| 07163 | 42 | DedA family protein [Denitromonas halophilus] |
| 04644 | 42 | urease subunit alpha [Azoarcus communis] |
| 07839 | 42 | VOC family protein [Azoarcus communis] |
| 05240 | 42 | hypothetical protein [Azoarcus communis] |
| 10348 | 42 | ATP-binding protein [Azoarcus communis] |
| 05695 | 42 | AlpA family phage regulatory protein [Aromatoleum buckelii] |
| 06099 | 42 | phosphate ABC transporter substrate-binding protein PstS [Azoarcus olearius] |
| 09039 | 42 | endonuclease/exonuclease/phosphatase family protein [Azoarcus communis] |
| 05464 | 42 | type IV pilus modification protein PilV [Azoarcus communis] |
| 02624 | 42 | exonuclease SbcCD subunit D [Azoarcus communis] |
| 07256 | 42 | type IV pili methyl-accepting chemotaxis transducer N-terminal domain-containing protein [Nitrogeniibacter aestuarii] |
| 10611 | 42 | FAD-dependent oxidoreductase [Azoarcus communis] |
| 01120 | 42 | DMT family transporter [Aromatoleum toluclasticum] |
| 05267 | 42 | hypothetical protein [Aromatoleum toluclasticum] |
| 05904 | 42 | MULTISPECIES: type II toxin-antitoxin system HicA family toxin [Aromatoleum] |
| 04903 | 42 | urease accessory protein UreD [Azoarcus communis] |
| 09741 | 42 | PsiF family protein [Denitromonas halophilus] |
| 00617 | 42 | hypothetical protein [Azoarcus communis] |
| 03771 | 42 | FIST C-terminal domain-containing protein [Nitrogeniibacter aestuarii] |
| 07395 | 42 | carboxymuconolactone decarboxylase family protein [Aromatoleum toluclasticum] |
| 07047 | 42 | c-type cytochrome [Azoarcus sp. KH32C] |
| 03784 | 42 | MULTISPECIES: putative nitrogen fixation protein NifT [Azoarcus] |
| 08444 | 41 | RtcB family protein [Nitrogeniibacter aestuarii] |
| 06178 | 41 | MULTISPECIES: hypothetical protein [Azoarcus] |
| 03376 | 41 | ferrous iron transport protein B [Azoarcus communis] |
| 00246 | 41 | Vat family streptogramin A O-acetyltransferase [Azoarcus indigens] |
| 07162 | 41 | hypothetical protein [Azoarcus communis] |
| 09166 | 41 | 4Fe-4S dicluster domain-containing protein [Denitromonas halophilus] |
| 08516 | 41 | DUF3106 domain-containing protein [Denitromonas halophilus] |
| 04520 | 41 | heavy metal-binding domain-containing protein [Azoarcus communis] |
| 00858 | 41 | precorrin-2 C(20)-methyltransferase [Azoarcus communis] |
| 08534 | 41 | DUF1330 domain-containing protein [Denitromonas halophilus] |
| 08299 | 41 | MULTISPECIES: cupredoxin family protein [Zoogloeaceae] |
| 08643 | 41 | DNA-3-methyladenine glycosylase [Azoarcus indigens] |
| 01375 | 41 | rhodoquinone biosynthesis methyltransferase RquA [Azoarcus communis] |
| 05724 | 41 | S8 family serine peptidase [Aromatoleum evansii] |
| 03277 | 41 | amidohydrolase [Azoarcus indigens] |
| 04836 | 41 | ABC transporter permease [Azoarcus communis] |
| 03626 | 41 | MULTISPECIES: type IV conjugative transfer system coupling protein TraD [Rhodocyclales] |
| 03880 | 41 | urease accessory protein UreE [Azoarcus communis] |
| 00495 | 41 | CsbD family protein [Aromatoleum diolicum] |
| 02395 | 41 | ABC transporter substrate-binding protein [Azoarcus sp. KH32C] |
| 00180 | 41 | YkgJ family cysteine cluster protein [Azoarcus communis] |
| 05775 | 41 | NAD(P)/FAD-dependent oxidoreductase [Denitromonas halophilus] |
| 00242 | 41 | TerC family protein [Propionivibrio dicarboxylicus] |
| 07900 | 41 | L,D-transpeptidase family protein [Denitromonas halophilus] |
| 02334 | 41 | FAD-dependent oxidoreductase [Azoarcus communis] |
| 00230 | 40 | feruloyl-CoA synthase [Aromatoleum toluclasticum] |
| 05072 | 40 | helix-turn-helix transcriptional regulator [Thauera humireducens] |
| 02070 | 40 | 5-oxoprolinase subunit PxpA [Azoarcus communis] |
| 01665 | 40 | DUF2238 domain-containing protein [Nitrogeniibacter aestuarii] |
| 08229 | 40 | DJ-1/PfpI family protein [Azoarcus communis] |
| 10160 | 40 | bifunctional DNA-binding transcriptional regulator/O6-methylguanine-DNA methyltransferase Ada [Aromatoleum toluclasticum] |
| 01758 | 40 | HAD family hydrolase [Azoarcus olearius] |
| 03613 | 40 | chain length determinant protein tyrosine kinase EpsG [Azoarcus communis] |
| 06487 | 40 | Flp family type IVb pilin [Aromatoleum toluclasticum] |
| 07531 | 40 | TetR family transcriptional regulator [Denitromonas halophilus] |
| 10370 | 40 | precorrin-8X methylmutase [Azoarcus communis] |
| 00924 | 40 | DUF779 domain-containing protein [Azoarcus communis] |
| 04625 | 40 | MBOAT family protein [Azoarcus sp. KH32C] |

| | | |
|---|---|---|
| 01065 | 40 | precorrin-3B C(17)-methyltransferase [Azoarcus communis] |
| 02829 | 40 | nodulation protein NfeD [Denitromonas halophilus] |
| 00137 | 40 | IclR family transcriptional regulator [Azoarcus sp. Aa7] |
| 03777 | 40 | tetratricopeptide repeat protein [Propionivibrio dicarboxylicus] |
| 07545 | 40 | PilZ domain-containing protein [Azoarcus communis] |
| 04749 | 40 | xanthine dehydrogenase family protein subunit M [Azoarcus indigens] |
| 00087 | 40 | Lrp/AsnC family transcriptional regulator [Denitromonas halophilus] |
| 09340 | 40 | two-component sensor histidine kinase [Aromatoleum toluclasticum] |
| 02876 | 40 | MULTISPECIES: AbrB/MazE/SpoVT family DNA-binding domain-containing protein [Aromatoleum] |
| 09048 | 40 | 4-hydroxyphenylpyruvate dioxygenase [Azoarcus communis] |
| 07175 | 40 | putative DNA binding domain-containing protein [Denitromonas halophilus] |
| 00138 | 40 | hypothetical protein [Denitromonas halophilus] |
| 01554 | 40 | MoaD/ThiS family protein [Azoarcus communis] |
| 05684 | 40 | hypothetical protein [Denitromonas halophilus] |
| 00522 | 40 | DUF533 domain-containing protein [Azoarcus communis] |
| 03224 | 40 | MliC family protein [Denitromonas halophilus] |
| 06046 | 40 | biotin-dependent carboxyltransferase family protein [Azoarcus communis] |
| 07319 | 40 | calcium-binding protein [Aromatoleum toluclasticum] |
| 04232 | 40 | LTA synthase family protein [Azoarcus olearius] |
| 01071 | 40 | chalcone isomerase family protein [Aromatoleum toluclasticum] |
| 09931 | 40 | aminoacyl-tRNA hydrolase [Denitromonas halophilus] |
| 10130 | 40 | tautomerase family protein [Azoarcus indigens] |
| 00659 | 40 | hypothetical protein [Aromatoleum toluclasticum] |
| 05312 | 40 | DUF1365 domain-containing protein [Azoarcus communis] |
| 08675 | 39 | MULTISPECIES: TraI domain-containing protein [Rhodocyclales] |
| 01406 | 39 | glycosyltransferase [Azoarcus communis] |
| 10360 | 39 | hydrolase 2, exosortase A system-associated [Denitromonas halophilus] |
| 00437 | 39 | MULTISPECIES: cation transporter [Thauera] |
| 01801 | 39 | GTPase/DUF3482 domain-containing protein [Denitromonas halophilus] |
| 09834 | 39 | MULTISPECIES: conjugal transfer protein TraG N-terminal domain-containing protein [Rhodocyclales] |
| 00299 | 39 | MULTISPECIES: VOC family protein [Rhodocyclales] |
| 09837 | 39 | prepilin-type N-terminal cleavage/methylation domain-containing protein [Azoarcus communis] |
| 03698 | 39 | NAD(+)--dinitrogen-reductase ADP-D-ribosyltransferase [Azoarcus communis] |
| 02069 | 39 | TlyA family RNA methyltransferase [Azoarcus communis] |
| 05290 | 39 | MULTISPECIES: RidA family protein [Aromatoleum] |
| 09219 | 39 | NTP transferase domain-containing protein [Azoarcus communis] |
| 01321 | 39 | hypothetical protein [Azoarcus communis] |
| 06903 | 39 | response regulator transcription factor [Azoarcus communis] |
| 10619 | 39 | dihydroneopterin triphosphate diphosphatase [Denitromonas halophilus] |
| 07860 | 39 | aminotransferase class V-fold PLP-dependent enzyme [Azoarcus communis] |
| 07191 | 39 | DUF3833 domain-containing protein [Azoarcus communis] |
| 04365 | 39 | EAL domain-containing protein [Azoarcus olearius] |
| 06963 | 39 | radical SAM protein [Azoarcus communis] |
| 08361 | 39 | alpha/beta fold hydrolase [Denitromonas halophilus] |
| 00262 | 39 | hypothetical protein [Denitromonas halophilus] |
| 01723 | 39 | MULTISPECIES: isoprenylcysteine carboxylmethyltransferase family protein [Zoogloeaceae] |
| 03910 | 39 | GatB/YqeY domain-containing protein [Denitromonas halophilus] |
| 01176 | 39 | arsenite efflux transporter metallochaperone ArsD [Denitromonas halophilus] |
| 02783 | 39 | flippase-like domain-containing protein [Denitromonas halophilus] |
| 04370 | 39 | hydrolase 1, exosortase A system-associated [Denitromonas halophilus] |
| 08910 | 38 | 5-oxoprolinase subunit PxpB [Azoarcus communis] |
| 02288 | 38 | arylsulfatase [Aromatoleum toluclasticum] |
| 07941 | 38 | TonB-dependent receptor [Azoarcus indigens] |
| 01807 | 38 | DEAD/DEAH box helicase [Azoarcus communis] |
| 07834 | 38 | type II secretion system GspH family protein [Denitromonas halophilus] |
| 02757 | 38 | hypothetical protein [Azoarcus communis] |
| 09769 | 38 | cobaltochelatase subunit CobN [Azoarcus communis] |
| 01834 | 38 | HPP family protein [Azoarcus communis] |
| 09181 | 38 | IS21 family transposase [Aromatoleum aromaticum] |
| 02736 | 38 | IS630 family transposase [Aromatoleum bremense] |
| 07684 | 38 | acetone carboxylase subunit gamma [Aromatoleum toluclasticum] |
| 01377 | 38 | cyanophycin synthetase [Denitromonas halophilus] |
| 03108 | 38 | 3-hydroxybenzoate 6-monooxygenase [Azoarcus communis] |
| 00086 | 38 | DUF3149 domain-containing protein [Azoarcus communis] |
| 01376 | 38 | tetratricopeptide repeat protein [Azoarcus communis] |
| 02671 | 38 | chalcone isomerase family protein [Azoarcus communis] |
| 07077 | 38 | DUF3025 domain-containing protein [Azoarcus communis] |
| 00836 | 38 | OB-fold domain-containing protein [Aromatoleum toluclasticum] |
| 00724 | 38 | NCS1 family transporter [Aromatoleum toluclasticum] |
| 08901 | 38 | xanthine dehydrogenase accessory protein XdhC [Azoarcus communis] |
| 01940 | 38 | NnrU family protein [Aromatoleum petrolei] |
| 00564 | 38 | methyl-accepting chemotaxis protein [Azoarcus communis] |
| 04487 | 38 | DUF938 domain-containing protein [Azoarcus communis] |
| 05266 | 38 | response regulator [Azoarcus communis] |
| 00704 | 38 | hypothetical protein [Azoarcus communis] |
| 08760 | 38 | MULTISPECIES: TIGR03757 family integrating conjugative element protein [Rhodocyclales] |
| 03624 | 38 | IS1182 family transposase [Aromatoleum bremense] |
| 10107 | 38 | FTR1 family protein [Aromatoleum toluclasticum] |
| 05482 | 38 | ankyrin repeat domain-containing protein [Denitromonas halophilus] |
| 01953 | 38 | MULTISPECIES: integrating conjugative element protein [Proteobacteria] |
| 05396 | 38 | DUF3306 domain-containing protein [Azoarcus communis] |
| 01707 | 38 | hypothetical protein [Azoarcus communis] |
| 07280 | 38 | GTP cyclohydrolase II [Nitrogeniibacter aestuarii] |
| 07255 | 38 | MULTISPECIES: proteasome-type protease [Niveibacterium] |
| 01427 | 38 | molecular chaperone TorD family protein [Azoarcus communis] |
| 09648 | 38 | CZB domain-containing protein [Azoarcus communis] |
| 01569 | 37 | nitroreductase family protein [Denitromonas halophilus] |
| 00293 | 37 | DUF488 family protein [Denitromonas halophilus] |
| 10452 | 37 | benzoyl-CoA reductase, bzd-type, subunit O [Aromatoleum toluclasticum] |
| 02240 | 37 | cytochrome c3 family protein [Nitrogeniibacter aestuarii] |
| 01829 | 37 | PAS domain S-box protein [Nitrogeniibacter aestuarii] |
| 06826 | 37 | Rieske 2Fe-2S domain-containing protein [Azoarcus communis] |
| 05425 | 37 | 4a-hydroxytetrahydrobiopterin dehydratase [Azoarcus communis] |
| 01649 | 37 | 1,4-alpha-glucan branching protein GlgB [Azoarcus olearius] |

| | | |
|---|---|---|
| 04976 | 37 | EpsD family peptidyl-prolyl cis-trans isomerase [Azoarcus communis] |
| 10246 | 37 | GNAT family N-acetyltransferase [Azoarcus communis] |
| 02637 | 37 | hypothetical protein [Azoarcus communis] |
| 02085 | 37 | DUF2189 domain-containing protein [Denitromonas halophilus] |
| 08143 | 37 | type II secretion system F family protein [Azoarcus nasutitermitis] |
| 03179 | 37 | pyrroloquinoline-quinone synthase PqqC [Azoarcus communis] |
| 07069 | 37 | cupredoxin domain-containing protein [Aromatoleum toluclasticum] |
| 01751 | 37 | malate synthase G [Azoarcus communis] |
| 09016 | 37 | ABC transporter ATP-binding protein [Aromatoleum bremense] |
| 10055 | 37 | MULTISPECIES: YciI family protein [Niveibacterium] |
| 00879 | 37 | diguanylate cyclase [Azoarcus communis] |
| 07879 | 37 | EAL domain-containing protein [Denitromonas halophilus] |
| 10284 | 37 | hypothetical protein [Azoarcus communis] |
| 10464 | 37 | PAAR domain-containing protein [Azoarcus communis] |
| 02629 | 37 | MFS transporter [Azoarcus communis] |
| 09322 | 36 | hypothetical protein [Aromatoleum toluclasticum] |
| 06497 | 36 | MULTISPECIES: TIGR03751 family conjugal transfer lipoprotein [Proteobacteria] |
| 03661 | 36 | ABC transporter permease [Aromatoleum bremense] |
| 01964 | 36 | MULTISPECIES: single-stranded DNA-binding protein [Proteobacteria] |
| 07082 | 36 | rubrerythrin family protein [Aromatoleum toluclasticum] |
| 05808 | 36 | DUF4399 domain-containing protein [Denitromonas halophilus] |
| 06794 | 36 | polysulfide reductase NrfD [Aromatoleum toluclasticum] |
| 07229 | 36 | PilW family protein [Azoarcus communis] |
| 05780 | 36 | DUF47 family protein [Azoarcus communis] |
| 10090 | 36 | DUF302 domain-containing protein [Azoarcus communis] |
| 01292 | 36 | restriction endonuclease [Denitromonas halophilus] |
| 08972 | 36 | MULTISPECIES: TIGR03756 family integrating conjugative element protein [Rhodocyclales] |
| 01545 | 36 | protocatechuate 4,5-dioxygenase subunit alpha [Azoarcus communis] |
| 02451 | 36 | hypothetical protein [Azoarcus communis] |
| 00796 | 36 | form I ribulose bisphosphate carboxylase large subunit [Azoarcus communis] |
| 08914 | 36 | phage holin family protein [Denitromonas halophilus] |
| 04302 | 36 | YdbL family protein [Azoarcus communis] |
| 09088 | 36 | DUF808 domain-containing protein [Azoarcus communis] |
| 08151 | 36 | DUF2784 domain-containing protein [Azoarcus communis] |
| 10225 | 36 | organic hydroperoxide resistance protein [Azoarcus indigens] |
| 01936 | 36 | B12-binding domain-containing radical SAM protein [Azoarcus communis] |
| 00538 | 36 | DMT family transporter [Denitromonas halophilus] |
| 04998 | 36 | cytochrome bd-I oxidase subunit CydX [Azoarcus indigens] |
| 07223 | 36 | NYN domain-containing protein [Denitromonas halophilus] |
| 05245 | 36 | hypothetical protein [Azoarcus communis] |
| 01727 | 36 | twin transmembrane helix small protein [Azoarcus communis] |
| 02912 | 36 | 3-oxoacid CoA-transferase subunit B [Azoarcus sp. KH32C] |
| 04137 | 36 | tetratricopeptide repeat protein [Aromatoleum toluclasticum] |
| 07502 | 36 | LLM class flavin-dependent oxidoreductase [Azoarcus indigens] |
| 05519 | 36 | MULTISPECIES: TIGR03746 family integrating conjugative element protein [Proteobacteria] |
| 09393 | 36 | DUF364 domain-containing protein [Aromatoleum bremense] |
| 02956 | 36 | phosphatase PAP2 family protein [Azoarcus communis] |
| 07732 | 36 | right-handed parallel beta-helix repeat-containing protein [Azoarcus communis] |
| 08862 | 36 | DUF3305 domain-containing protein [Azoarcus communis] |
| 05742 | 36 | hypothetical protein [Azoarcus communis] |
| 09431 | 36 | MULTISPECIES: superoxide dismutase family protein [Niveibacterium] |
| 00911 | 36 | DUF6531 domain-containing protein, partial [Thauera butanivorans] |
| 03755 | 36 | beta-1,6-glucan synthase [Azoarcus communis] |
| 06107 | 36 | PEPxxWA-CTERM sorting domain-containing protein [Nitrogeniibacter aestuarii] |
| 00905 | 36 | amidohydrolase family protein [Aromatoleum toluclasticum] |
| 10001 | 36 | Na/Pi symporter [Azoarcus communis] |
| 03494 | 36 | carbamoyltransferase [Azoarcus indigens] |
| 09254 | 36 | GtrA family protein [Azoarcus communis] |
| 03675 | 36 | hypothetical protein [Denitromonas halophilus] |
| 02057 | 36 | hypothetical protein [Aromatoleum toluclasticum] |
| 03042 | 36 | GPW/gp25 family protein [Azoarcus communis] |
| 02642 | 36 | TetR/AcrR family transcriptional regulator [Azoarcus olearius] |
| 10550 | 36 | GAF domain-containing protein [Azoarcus indigens] |
| 00070 | 36 | MULTISPECIES: TIGR03761 family integrating conjugative element protein [Proteobacteria] |
| 04735 | 36 | MULTISPECIES: TIGR03750 family conjugal transfer protein [Rhodocyclales] |
| 09017 | 35 | L-lactate permease [Thauera phenolivorans] |
| 00865 | 35 | TIGR02281 family clan AA aspartic protease [Denitromonas halophilus] |
| 08382 | 35 | hypothetical protein [Aromatoleum toluclasticum] |
| 09402 | 35 | ABC transporter permease [Azoarcus communis] |
| 08677 | 35 | hypothetical protein [Azoarcus communis] |
| 05689 | 35 | hypothetical protein [Denitromonas halophilus] |
| 05557 | 35 | MULTISPECIES: copper oxidase [Zoogloeaceae] |
| 07847 | 35 | MULTISPECIES: TIGR03749 family integrating conjugative element protein [Rhodocyclales] |
| 08120 | 35 | transporter [Azoarcus communis] |
| 01216 | 35 | MBL fold metallo-hydrolase [Aromatoleum toluclasticum] |
| 06611 | 35 | transporter substrate-binding domain-containing protein [Niveibacterium umoris] |
| 07284 | 35 | OmpA family protein [Denitromonas halophilus] |
| 09800 | 35 | PilZ domain-containing protein [Azoarcus communis] |
| 08091 | 35 | cobalt-precorrin-5B (C(1))-methyltransferase [Denitromonas halophilus] |
| 09808 | 35 | DUF1992 domain-containing protein [Azoarcus communis] |
| 00777 | 35 | ChaN family lipoprotein [Denitromonas halophilus] |
| 05728 | 35 | CBS domain-containing protein [Denitromonas halophilus] |
| 06459 | 35 | peptidoglycan-binding protein LysM [Azoarcus communis] |
| 01257 | 35 | NAD(P)/FAD-dependent oxidoreductase [Azoarcus communis] |
| 00161 | 35 | pyridoxamine 5'-phosphate oxidase family protein [Nitrogeniibacter aestuarii] |
| 09570 | 35 | ROK family protein [Azoarcus communis] |
| 07353 | 35 | hypothetical protein [Azoarcus communis] |
| 05374 | 35 | sirohydrochlorin chelatase [Denitromonas halophilus] |
| 04214 | 35 | MULTISPECIES: type II toxin-antitoxin system PrlF family antitoxin [Rhodocyclales] |
| 05510 | 35 | hypothetical protein [Denitromonas halophilus] |
| 05708 | 35 | MetQ/NlpA family ABC transporter substrate-binding protein [Azoarcus rhizosphaerae] |
| 06648 | 35 | SIR2 family protein [Azoarcus communis] |
| 09015 | 35 | RNA polymerase sigma factor [Denitromonas halophilus] |
| 01778 | 35 | 2-oxoacid:acceptor oxidoreductase family protein [Aromatoleum toluclasticum] |

| | | |
|---|---|---|
| 08616 | 35 | 3-hydroxylacyl-ACP dehydratase [Azoarcus communis] |
| 02911 | 34 | nitroreductase family protein [Azoarcus indigens] |
| 02946 | 34 | hypothetical protein [Azoarcus communis] |
| 03960 | 34 | flagellar hook-length control protein FliK [Denitromonas halophilus] |
| 08615 | 34 | MULTISPECIES: polyamine aminopropyltransferase [Niveibacterium] |
| 05976 | 34 | MULTISPECIES: TIGR03747 family integrating conjugative element membrane protein [Proteobacteria] |
| 05211 | 34 | hypothetical protein [Azoarcus communis] |
| 05734 | 34 | hypothetical protein [Azoarcus communis] |
| 02935 | 34 | MULTISPECIES: entericidin A/B family lipoprotein [Thauera] |
| 04410 | 34 | gamma-glutamylcyclotransferase [Azoarcus communis] |
| 05307 | 34 | MULTISPECIES: TIGR03759 family integrating conjugative element protein [Proteobacteria] |
| 05178 | 34 | hypothetical protein [Aromatoleum toluclasticum] |
| 03148 | 34 | lysine 2,3-aminomutase [Denitromonas halophilus] |
| 00420 | 34 | MULTISPECIES: TIGR03745 family integrating conjugative element membrane protein [Proteobacteria] |
| 04619 | 34 | mechanosensitive ion channel family protein [Azoarcus communis] |
| 05043 | 34 | VOC family protein [Aromatoleum toluclasticum] |
| 04815 | 34 | hypothetical protein [Denitromonas halophilus] |
| 05564 | 34 | cobalamin biosynthesis protein CbiG [Denitromonas halophilus] |
| 00217 | 34 | MULTISPECIES: PilL N-terminal domain-containing protein [Proteobacteria] |
| 08908 | 34 | DUF2149 domain-containing protein [Nitrogeniibacter aestuarii] |
| 00253 | 34 | PhnD/SsuA/transferrin family substrate-binding protein [Aromatoleum buckelii] |
| 05614 | 34 | 3-keto-5-aminohexanoate cleavage protein [Aromatoleum toluclasticum] |
| 04968 | 34 | hypothetical protein [Azoarcus communis] |
| 08887 | 34 | type II toxin-antitoxin system RelE/ParE family toxin [Azoarcus communis] |
| 00358 | 34 | c-type cytochrome [Aromatoleum toluclasticum] |
| 09965 | 34 | glycine cleavage system protein R [Denitromonas halophilus] |
| 06137 | 34 | MULTISPECIES: transglycosylase SLT domain-containing protein [Proteobacteria] |
| 04181 | 34 | TonB-dependent receptor [Denitromonas halophilus] |
| 03055 | 34 | PAS domain S-box protein [Denitromonas halophilus] |
| 07103 | 34 | ABC transporter permease [Denitromonas halophilus] |
| 01773 | 34 | phosphotransferase [Azoarcus communis] |
| 07844 | 34 | ferritin-like domain-containing protein [Denitromonas halophilus] |
| 01412 | 34 | DoxX family protein [Aromatoleum toluclasticum] |
| 05023 | 34 | MULTISPECIES: DUF3275 family protein [Proteobacteria] |
| 01046 | 34 | TIGR04283 family arsenosugar biosynthesis glycosyltransferase [Aromatoleum toluclasticum] |
| 10155 | 34 | cytochrome c [Denitromonas halophilus] |
| 01534 | 34 | potassium/proton antiporter [Azoarcus communis] |
| 04266 | 34 | MULTISPECIES: AlpA family transcriptional regulator [Betaproteobacteria] |
| 08637 | 34 | hypothetical protein [Azoarcus communis] |
| 01693 | 34 | hypothetical protein [Azoarcus communis] |
| 04262 | 34 | Lin0512 family protein [Denitromonas halophilus] |
| 00294 | 34 | hypothetical protein [Azoarcus communis] |
| 06696 | 33 | PAS domain-containing protein [Azoarcus communis] |
| 09770 | 33 | Fe2+-dependent dioxygenase [Azoarcus communis] |
| 07881 | 33 | XRE family transcriptional regulator [Azoarcus communis] |
| 05141 | 33 | DUF4124 domain-containing protein [Azoarcus indigens] |
| 06357 | 33 | energy transducer TonB [Azoarcus sp. KH32C] |
| 04727 | 33 | PRC-barrel domain-containing protein [Azoarcus indigens] |
| 08867 | 33 | hypothetical protein [Denitromonas halophilus] |
| 06002 | 33 | RES family NAD+ phosphorylase [Azoarcus communis] |
| 01062 | 33 | DUF1615 domain-containing protein [Nitrogeniibacter aestuarii] |
| 07153 | 33 | MotA/TolQ/ExbB proton channel family protein [Nitrogeniibacter aestuarii] |
| 03082 | 33 | DUF2868 domain-containing protein [Azoarcus communis] |
| 05691 | 33 | hypothetical protein [Aromatoleum toluclasticum] |
| 03262 | 33 | 4Fe-4S binding protein [Aromatoleum toluclasticum] |
| 03892 | 33 | DUF2237 domain-containing protein [Azoarcus communis] |
| 03005 | 33 | hypothetical protein [Azoarcus communis] |
| 08557 | 33 | caspase family protein [Denitromonas halophilus] |
| 02496 | 33 | NAD(+) diphosphatase [Azoarcus communis] |
| 03973 | 33 | MULTISPECIES: GNAT family N-acetyltransferase [Aromatoleum] |
| 09068 | 33 | antibiotic biosynthesis monooxygenase [Denitromonas halophilus] |
| 09511 | 33 | arsenical pump-driving ATPase [Denitromonas halophilus] |
| 01220 | 33 | DUF4149 domain-containing protein [Azoarcus communis] |
| 02122 | 33 | hypothetical protein [Denitromonas halophilus] |
| 10275 | 33 | type II toxin-antitoxin system VapC family toxin [Aromatoleum aromaticum] |
| 08368 | 33 | M1 family metallopeptidase [Denitromonas halophilus] |
| 07625 | 33 | cobalamin biosynthesis protein [Denitromonas halophilus] |
| 03104 | 32 | carbamoyltransferase HypF [Azoarcus communis] |
| 10580 | 32 | hypothetical protein [Azoarcus indigens] |
| 05232 | 32 | hypothetical protein [Azoarcus communis] |
| 06801 | 32 | CHAT domain-containing protein [Denitromonas halophilus] |
| 08428 | 32 | indolepyruvate oxidoreductase subunit beta family protein [Azoarcus indigens] |
| 10093 | 32 | MULTISPECIES: integrating conjugative element protein [Rhodocyclales] |
| 08037 | 32 | hypothetical protein [Denitromonas halophilus] |
| 09798 | 32 | LysE family translocator [Azoarcus indigens] |
| 07594 | 32 | EAL domain-containing protein [Aromatoleum toluclasticum] |
| 02697 | 32 | acyl CoA:acetate/3-ketoacid CoA transferase [Azoarcus communis] |
| 06703 | 32 | LEA type 2 family protein [Denitromonas halophilus] |
| 01482 | 32 | MULTISPECIES: outer membrane beta-barrel protein [Niveibacterium] |
| 01894 | 32 | hypothetical protein [Azoarcus communis] |
| 04947 | 32 | M48 family metallopeptidase [Denitromonas halophilus] |
| 03502 | 32 | hypothetical protein [Azoarcus communis] |
| 03609 | 32 | YajD family HNH nuclease [Denitromonas halophilus] |
| 05004 | 32 | TIGR00730 family Rossman fold protein [Azoarcus indigens] |
| 07146 | 32 | cupin domain-containing protein [Aromatoleum toluclasticum] |
| 08611 | 32 | DUF3299 domain-containing protein [Azoarcus communis] |
| 04519 | 32 | adenosylmethionine decarboxylase [Denitromonas halophilus] |
| 04420 | 32 | hypothetical protein [Denitromonas halophilus] |
| 05452 | 32 | type II secretion system minor pseudopilin GspI [Azoarcus indigens] |
| 02166 | 32 | anti-sigma factor [Denitromonas halophilus] |
| 01031 | 32 | NAD-glutamate dehydrogenase [Denitromonas halophilus] |
| 01254 | 32 | type I restriction endonuclease subunit R [Denitromonas halophilus] |
| 01341 | 32 | ABC transporter substrate-binding protein [Aromatoleum toluclasticum] |
| 01077 | 32 | NUDIX domain-containing protein [Denitromonas halophilus] |

| | | |
|---|---|---|
| 04259 | 32 | MULTISPECIES: hypothetical protein [Proteobacteria] |
| 07836 | 32 | lipocalin family protein [Azoarcus communis] |
| 01094 | 32 | TIGR04282 family arsenosugar biosynthesis glycosyltransferase [Aromatoleum toluclasticum] |
| 00958 | 32 | MULTISPECIES: DUF6094 domain-containing protein [Proteobacteria] |
| 09649 | 32 | VOC family protein [Denitromonas halophilus] |
| 09959 | 32 | hypothetical protein [Aromatoleum toluclasticum] |
| 09968 | 32 | hypothetical protein [Azoarcus communis] |
| 06409 | 32 | hypothetical protein [Azoarcus communis] |
| 04208 | 32 | arsenosugar biosynthesis radical SAM protein ArsS [Aromatoleum toluclasticum] |
| 02557 | 32 | TerC family protein [Denitromonas halophilus] |
| 04573 | 32 | pyrroloquinoline quinone biosynthesis protein PqqB [Azoarcus communis] |
| 08492 | 32 | cupin domain-containing protein [Azoarcus communis] |
| 08310 | 32 | hypothetical protein [Azoarcus communis] |
| 04588 | 32 | MoaD/ThiS family protein [Denitromonas halophilus] |
| 05456 | 31 | metal-dependent hydrolase [Azoarcus communis] |
| 10000 | 31 | sel1 repeat family protein [Aromatoleum toluclasticum] |
| 05632 | 31 | LLM class flavin-dependent oxidoreductase [Azoarcus communis] |
| 10624 | 31 | 4-hydroxy-2-oxoheptanedioate aldolase [Azoarcus indigens] |
| 03966 | 31 | MULTISPECIES: YnfA family protein [Rhodocyclales] |
| 06402 | 31 | hypothetical protein [Aromatoleum diolicum] |
| 07674 | 31 | hypothetical protein [Aromatoleum toluclasticum] |
| 04769 | 31 | ABC transporter permease [Uliginosibacterium gangwonense] |
| 08824 | 31 | MULTISPECIES: DEAD/DEAH box helicase [Rhodocyclales] |
| 00773 | 31 | MULTISPECIES: STY4534 family ICE replication protein [Rhodocyclales] |
| 06924 | 31 | hypothetical protein [Aromatoleum toluclasticum] |
| 02695 | 31 | formate/nitrite transporter family protein [Denitromonas halophilus] |
| 01610 | 31 | glycine betaine/L-proline ABC transporter substrate-binding protein ProX [Denitromonas halophilus] |
| 07804 | 31 | MFS transporter [Azoarcus communis] |
| 10087 | 31 | hypothetical protein [Azoarcus communis] |
| 09539 | 31 | DUF3309 domain-containing protein [Azoarcus communis] |
| 06160 | 31 | patatin-like phospholipase family protein [Azoarcus communis] |
| 06370 | 31 | HyaD/HybD family hydrogenase maturation endopeptidase [Azoarcus communis] |
| 01113 | 31 | L,D-transpeptidase family protein [Azoarcus communis] |
| 00912 | 31 | HNH endonuclease [Azoarcus communis] |
| 05299 | 31 | hypothetical protein [Aromatoleum toluclasticum] |
| 06822 | 31 | hypothetical protein [Denitromonas halophilus] |
| 04288 | 31 | indolepyruvate oxidoreductase subunit beta [Denitromonas halophilus] |
| 00030 | 31 | 2,3-bisphosphoglycerate-independent phosphoglycerate mutase [Aromatoleum toluclasticum] |
| 10397 | 31 | type II secretion system major pseudopilin GspG [Azoarcus indigens] |
| 10304 | 31 | TetR/AcrR family transcriptional regulator [Aromatoleum toluclasticum] |
| 04530 | 31 | hypothetical protein [Denitromonas halophilus] |
| 07149 | 31 | CzcE family metal-binding protein [Aromatoleum toluclasticum] |
| 01494 | 31 | DUF429 domain-containing protein [Azoarcus communis] |
| 07035 | 31 | LLM class flavin-dependent oxidoreductase [Aromatoleum toluclasticum] |
| 05871 | 31 | DUF6129 family protein [Azoarcus communis] |
| 00338 | 31 | hypothetical protein [Aromatoleum toluclasticum] |
| 05828 | 31 | hypothetical protein [Azoarcus communis] |
| 07576 | 31 | PDDEXK nuclease domain-containing protein [Zoogloea oleivorans] |
| 07208 | 31 | FxDxF family PEP-CTERM protein [Azoarcus communis] |
| 08219 | 31 | autotransporter domain-containing protein [Azoarcus indigens] |
| 02186 | 31 | MbcA/ParS/Xre antitoxin family protein [Azoarcus communis] |
| 00084 | 31 | ribbon-helix-helix domain-containing protein [Azoarcus communis] |
| 03017 | 31 | hypothetical protein [Azoarcus communis] |
| 09571 | 31 | ferredoxin family protein [Azoarcus indigens] |
| 02160 | 31 | phage tail sheath subtilisin-like domain-containing protein [Denitromonas halophilus] |
| 01542 | 31 | EAL domain-containing response regulator [Aromatoleum toluclasticum] |
| 07918 | 31 | hypothetical protein [Denitromonas halophilus] |
| 05194 | 31 | hypothetical protein [Azoarcus communis] |
| 04451 | 31 | MarR family transcriptional regulator [Azoarcus indigens] |
| 01315 | 31 | GAF domain-containing protein [Azoarcus communis] |
| 10573 | 30 | cupin domain-containing protein [Denitromonas halophilus] |
| 00406 | 30 | FIST C-terminal domain-containing protein [Azoarcus communis] |
| 03948 | 30 | carbohydrate porin [Denitromonas halophilus] |
| 01383 | 30 | glycosyltransferase family 2 protein [Aromatoleum toluclasticum] |
| 03363 | 30 | MULTISPECIES: quinohemoprotein amine dehydrogenase subunit beta [Thauera] |
| 09936 | 30 | hypothetical protein [Denitromonas halophilus] |
| 05739 | 30 | 2-oxo-4-hydroxy-4-carboxy-5-ureidoimidazoline decarboxylase [Azoarcus communis] |
| 00206 | 30 | hypothetical protein [Uliginosibacterium aquaticum] |
| 10390 | 30 | iron transporter [Denitromonas halophilus] |
| 04561 | 30 | amidohydrolase family protein [Nitrogeniibacter aestuarii] |
| 05726 | 30 | pyridoxamine 5'-phosphate oxidase family protein [Azoarcus communis] |
| 09674 | 30 | glycosyltransferase [Denitromonas halophilus] |
| 02495 | 30 | ethanolamine ammonia-lyase subunit EutC [Azoarcus indigens] |
| 07297 | 30 | serine protease [Azoarcus communis] |
| 10137 | 30 | regulator [Nitrogeniibacter aestuarii] |
| 10213 | 30 | glycosyltransferase [Nitrogeniibacter aestuarii] |
| 05678 | 30 | HypC/HybG/HupF family hydrogenase formation chaperone [Azoarcus communis] |
| 09198 | 30 | GGDEF domain-containing protein [Denitromonas halophilus] |
| 06680 | 30 | DUF2796 domain-containing protein [Azoarcus communis] |
| 04496 | 30 | MULTISPECIES: hypothetical protein [Aromatoleum] |
| 08187 | 30 | acetyl-CoA hydrolase/transferase family protein [Aromatoleum toluclasticum] |
| 08970 | 30 | hydroxyisourate hydrolase [Azoarcus communis] |
| 01844 | 30 | cold shock and DUF1294 domain-containing protein [Azoarcus communis] |
| 02176 | 30 | MULTISPECIES: TolC family protein [Zoogloeaceae] |
| 08063 | 30 | glycosyltransferase family 39 protein [Denitromonas halophilus] |
| 09866 | 30 | quinohemoprotein amine dehydrogenase subunit alpha [Aromatoleum aromaticum] |
| 08236 | 30 | hypothetical protein [Azoarcus communis] |
| 01535 | 30 | MULTISPECIES: arginine N-succinyltransferase [Niveibacterium] |
| 04075 | 30 | cytochrome c biogenesis protein CcsA [Aromatoleum diolicum] |
| 08820 | 30 | DUF3135 domain-containing protein [Denitromonas halophilus] |
| 09496 | 30 | nucleoid-associated protein [Azoarcus communis] |
| 07761 | 30 | hypothetical protein [Denitromonas halophilus] |
| 10278 | 30 | MULTISPECIES: quinohemoprotein amine dehydrogenase subunit gamma [Rhodocyclales] |
| 06510 | 30 | hypothetical protein [Denitromonas halophilus] |

| | | |
|---|---|---|
| 02284 | 30 | MarR family transcriptional regulator [Azoarcus communis] |
| 04427 | 30 | [NiFe]-hydrogenase assembly chaperone HybE [Azoarcus communis] |
| 03348 | 30 | type II toxin-antitoxin system RelE/ParE family toxin [Aromatoleum pumilum] |
| 01837 | 29 | hypothetical protein [Aromatoleum toluclasticum] |
| 10479 | 29 | twin-arginine translocation signal domain-containing protein [Azoarcus olearius] |
| 02503 | 29 | hypothetical protein [Azoarcus communis] |
| 02279 | 29 | 6-oxocyclohex-1-ene-1-carbonyl-CoA hydratase [Aromatoleum toluclasticum] |
| 06597 | 29 | IS91 family transposase [Thauera terpenica] |
| 06098 | 29 | type IVB secretion system protein IcmH/DotU [Denitromonas halophilus] |
| 03112 | 29 | DUF2242 domain-containing protein [Aromatoleum toluclasticum] |
| 02776 | 29 | xanthine phosphoribosyltransferase [Azoarcus communis] |
| 07377 | 29 | EcsC family protein [Aromatoleum toluclasticum] |
| 06828 | 29 | ethanolamine ammonia-lyase subunit EutB [Aromatoleum evansii] |
| 09035 | 29 | copper resistance system multicopper oxidase [Denitromonas halophilus] |
| 00651 | 29 | dimethyl sulfoxide reductase anchor subunit [Aromatoleum toluclasticum] |
| 01340 | 29 | type II secretion system minor pseudopilin GspK [Azoarcus indigens] |
| 04920 | 29 | CopG family ribbon-helix-helix protein [Denitromonas halophilus] |
| 06818 | 29 | DEAD/DEAH box helicase [Aromatoleum aromaticum] |
| 02745 | 29 | transglutaminase family protein [Aromatoleum toluclasticum] |
| 03127 | 29 | adenosine deaminase [Azoarcus communis] |
| 07790 | 29 | glucosylglycerol-phosphate synthase [Azoarcus communis] |
| 00003 | 29 | 2,3-epoxybenzoyl-CoA dihydrolase [Aromatoleum toluclasticum] |
| 01976 | 29 | HupE/UreJ family protein [Denitromonas halophilus] |
| 08876 | 29 | PilX N-terminal domain-containing pilus assembly protein [Azoarcus communis] |
| 05447 | 29 | sodium-dependent bicarbonate transport family permease [Azoarcus communis] |
| 09228 | 29 | MULTISPECIES: DUF3742 family protein [Proteobacteria] |
| 08284 | 29 | hypothetical protein [Azoarcus communis] |
| 07336 | 29 | sulfite exporter TauE/SafE family protein [Azoarcus indigens] |
| 08221 | 29 | aminoacyl-histidine dipeptidase [Aromatoleum toluclasticum] |
| 01566 | 29 | porin [Denitromonas halophilus] |
| 09733 | 29 | NAD(P)/FAD-dependent oxidoreductase [Azoarcus communis] |
| 03608 | 29 | tetrathionate reductase family octaheme c-type cytochrome [Denitromonas halophilus] |
| 06567 | 29 | DUF2721 domain-containing protein [Azoarcus communis] |
| 02717 | 29 | MULTISPECIES: DUF945 domain-containing protein [Proteobacteria] |
| 01225 | 29 | DUF3862 domain-containing protein [Aromatoleum toluclasticum] |
| 03355 | 29 | DoxX family protein [Azoarcus indigens] |
| 00778 | 29 | type VI secretion system lipoprotein TssJ [Denitromonas halophilus] |
| 03013 | 29 | LysE/ArgO family amino acid transporter [Denitromonas halophilus] |
| 08691 | 29 | MULTISPECIES: hypothetical protein [Proteobacteria] |
| 05751 | 29 | DMT family transporter [Thauera propionica] |
| 07828 | 29 | ISL3 family transposase [Azoarcus sp. Aa7] |
| 05694 | 29 | transposase [Azoarcus communis] |
| 04694 | 29 | UDP-N-acetylglucosamine 4,6-dehydratase (inverting) [Denitromonas halophilus] |
| 04248 | 29 | SGNH/GDSL hydrolase family protein [Denitromonas halophilus] |
| 09335 | 28 | GNAT family N-acetyltransferase [Niveibacterium sp. COAC-50] |
| 06113 | 28 | hypothetical protein [Denitromonas halophilus] |
| 08655 | 28 | GGDEF domain-containing protein [Nitrogeniibacter aestuarii] |
| 10121 | 28 | ABC transporter substrate-binding protein [Aromatoleum toluclasticum] |
| 07678 | 28 | Asp/Glu racemase [Denitromonas halophilus] |
| 08538 | 28 | MULTISPECIES: pyrroloquinoline quinone precursor peptide PqqA [Rhodocyclales] |
| 10215 | 28 | DUF898 domain-containing protein [Denitromonas halophilus] |
| 05055 | 28 | MULTISPECIES: disulfide bond formation protein B [Thauera] |
| 09566 | 28 | EAL domain-containing protein [Azoarcus communis] |
| 05922 | 28 | glycosyltransferase [Azoarcus rhizosphaerae] |
| 04168 | 28 | nitrate reductase molybdenum cofactor assembly chaperone [Denitromonas halophilus] |
| 07457 | 28 | FAD-binding oxidoreductase [Aromatoleum aromaticum] |
| 07115 | 28 | (2Fe-2S)-binding protein [Azoarcus communis] |
| 04613 | 28 | RcnB family protein [Azoarcus communis] |
| 09734 | 28 | DUF2269 domain-containing protein [Azoarcus indigens] |
| 02572 | 28 | SUF system NifU family Fe-S cluster assembly protein [Denitromonas halophilus] |
| 00919 | 28 | MULTISPECIES: hypothetical protein [Thauera] |
| 03109 | 28 | MULTISPECIES: hypothetical protein [Proteobacteria] |
| 04061 | 28 | PAS domain-containing protein [Zoogloea oleivorans] |
| 10134 | 28 | hypothetical protein [Denitromonas halophilus] |
| 07451 | 28 | hypothetical protein [Azoarcus communis] |
| 10355 | 28 | MULTISPECIES: hypothetical protein [Thauera] |
| 03998 | 28 | glucose-6-phosphate dehydrogenase [Aromatoleum diolicum] |
| 00814 | 28 | 5-histidylcysteine sulfoxide synthase [Azoarcus communis] |
| 10198 | 28 | TonB-dependent receptor [Denitromonas halophilus] |
| 09321 | 28 | hypothetical protein [Candidatus Dactylopiibacterium carminicum] |
| 10204 | 28 | cupin domain-containing protein [Denitromonas halophilus] |
| 04743 | 28 | DUF692 domain-containing protein [Nitrogeniibacter aestuarii] |
| 02425 | 28 | OmpA family protein [Denitromonas halophilus] |
| 01859 | 28 | 2Fe-2S iron-sulfur cluster-binding protein [Aromatoleum toluvorans] |
| 04448 | 28 | respiratory nitrate reductase subunit gamma [Denitromonas halophilus] |
| 09085 | 28 | pseudaminic acid synthase [Denitromonas halophilus] |
| 01624 | 28 | glycosyltransferase family 39 protein [Aromatoleum toluclasticum] |
| 01114 | 28 | type VI secretion system-associated protein TagF [Azoarcus communis] |
| 06934 | 28 | PEP-CTERM sorting domain-containing protein [Azoarcus halotolerans] |
| 04787 | 28 | HipA domain-containing protein [Aromatoleum pumilum] |
| 05176 | 28 | sel1 repeat family protein [Denitromonas halophilus] |
| 02206 | 28 | PilW family protein [Aromatoleum toluclasticum] |
| 04423 | 28 | methyltransferase [Denitromonas halophilus] |
| 07770 | 28 | MFS transporter [Aromatoleum toluclasticum] |
| 09530 | 28 | ATP-grasp domain-containing protein [Denitromonas halophilus] |
| 08089 | 28 | WYL domain-containing protein [Aromatoleum bremense] |
| 07343 | 28 | DUF4351 domain-containing protein [Rhodocyclus purpureus] |
| 06925 | 28 | ABC transporter substrate-binding protein [Azoarcus indigens] |
| 05212 | 28 | HD domain-containing protein [Aromatoleum bremense] |
| 06085 | 28 | response regulator transcription factor [Zoogloea dura] |
| 03945 | 28 | sulfite dehydrogenase [Azoarcus communis] |
| 03929 | 28 | hypothetical protein [Denitromonas halophilus] |
| 03778 | 27 | VIT family protein [Denitromonas halophilus] |
| 08159 | 27 | nicotinate phosphoribosyltransferase [Aromatoleum toluclasticum] |

| | | |
|---|---|---|
| 07636 | 27 | alpha/beta hydrolase [Nitrogeniibacter aestuarii] |
| 01362 | 27 | YitT family protein [Denitromonas halophilus] |
| 05605 | 27 | PEP-CTERM sorting domain-containing protein [Azoarcus communis] |
| 08819 | 27 | SDR family oxidoreductase [Aromatoleum toluclasticum] |
| 03229 | 27 | NAD(+)/NADH kinase [Denitromonas halophilus] |
| 06972 | 27 | MULTISPECIES: stabilization protein [Proteobacteria] |
| 08730 | 27 | AsmA family protein [Denitromonas halophilus] |
| 03199 | 27 | YbaK/EbsC family protein [Denitromonas halophilus] |
| 10428 | 27 | DUF3365 domain-containing protein [Nitrogeniibacter aestuarii] |
| 04212 | 27 | radical SAM protein [Azoarcus communis] |
| 09670 | 27 | MULTISPECIES: helix-turn-helix domain-containing protein [Proteobacteria] |
| 02017 | 27 | MULTISPECIES: phenylacetic acid degradation operon negative regulatory protein PaaX [Thauera] |
| 02586 | 27 | prepilin-type N-terminal cleavage/methylation domain-containing protein [Azoarcus indigens] |
| 04720 | 27 | permease [Azoarcus communis] |
| 03395 | 27 | DNA cytosine methyltransferase [Aromatoleum evansii] |
| 10367 | 27 | nitrate regulatory protein [Azoarcus communis] |
| 07826 | 27 | SdiA-regulated domain-containing protein [Azoarcus communis] |
| 09011 | 27 | TIM barrel protein [Thauera humireducens] |
| 04823 | 27 | hypothetical protein [Thauera phenylacetica] |
| 04565 | 27 | S8 family serine peptidase [Aromatoleum aromaticum] |
| 00331 | 27 | MULTISPECIES: cupredoxin domain-containing protein [Azoarcus] |
| 06407 | 27 | hypothetical protein [Azoarcus communis] |
| 02553 | 27 | MULTISPECIES: helix-turn-helix transcriptional regulator [Niveibacterium] |
| 08240 | 27 | MULTISPECIES: benzoyl-CoA 2,3-epoxidase subunit BoxB [Aromatoleum] |
| 10056 | 27 | SoxR reducing system RseC family protein [Aromatoleum toluclasticum] |
| 00678 | 27 | acyloxyacyl hydrolase [Azoarcus indigens] |
| 01477 | 27 | MULTISPECIES: ribonucleotide-diphosphate reductase subunit beta [Niveibacterium] |
| 10437 | 27 | YbdK family carboxylate-amine ligase [Aromatoleum toluclasticum] |
| 09690 | 27 | iron transporter [Azoarcus rhizosphaerae] |
| 01165 | 27 | MgtC/SapB family protein [Azoarcus olearius] |
| 01604 | 27 | 4Fe-4S binding protein [Aromatoleum toluclasticum] |
| 03120 | 27 | glycoside hydrolase family 5 protein [Aromatoleum toluclasticum] |
| 05982 | 27 | MULTISPECIES: STY4528 family pathogenicity island replication protein [Rhodocyclales] |
| 08227 | 27 | hemerythrin domain-containing protein [Denitromonas halophilus] |
| 09610 | 27 | MULTISPECIES: hypothetical protein [Rhodocyclales] |
| 03654 | 27 | hypothetical protein [Azoarcus communis] |
| 00239 | 27 | hypothetical protein [Azoarcus communis] |
| 00626 | 27 | TIGR02281 family clan AA aspartic protease [Azoarcus communis] |
| 05116 | 27 | hypothetical protein [Azoarcus communis] |
| 04696 | 27 | HAMP domain-containing protein [Thauera butanivorans] |
| 09448 | 27 | DUF3047 domain-containing protein [Azoarcus communis] |
| 07932 | 27 | hypothetical protein [Denitromonas halophilus] |
| 08132 | 27 | DUF3750 domain-containing protein [Denitromonas halophilus] |
| 09353 | 26 | MULTISPECIES: nitrogen fixation protein NifZ [Azoarcus] |
| 02600 | 26 | DUF423 domain-containing protein [Aromatoleum toluclasticum] |
| 05773 | 26 | transglutaminase domain-containing protein [Azoarcus indigens] |
| 00023 | 26 | DUF2339 domain-containing protein [Azoarcus communis] |
| 07992 | 26 | (Fe-S) protein [Azoarcus communis] |
| 03052 | 26 | urate hydroxylase PuuD [Azoarcus communis] |
| 00063 | 26 | TIGR01458 family HAD-type hydrolase [Aromatoleum toluclasticum] |
| 04750 | 26 | polysaccharide pyruvyl transferase family protein [Azoarcus indigens] |
| 03507 | 26 | alpha/beta hydrolase [Denitromonas halophilus] |
| 00060 | 26 | formate dehydrogenase accessory protein FdhE [Azoarcus indigens] |
| 04893 | 26 | N-formylglutamate amidohydrolase [Nitrogeniibacter aestuarii] |
| 08593 | 26 | IS1634 family transposase [Thauera chlorobenzoica] |
| 03386 | 26 | N(2)-fixation sustaining protein CowN [Azoarcus communis] |
| 09841 | 26 | osmoprotectant NAGGN system M42 family peptidase [Denitromonas halophilus] |
| 03358 | 26 | low specificity L-threonine aldolase [Denitromonas halophilus] |
| 03161 | 26 | tyrosine-protein phosphatase [Nitrogeniibacter aestuarii] |
| 03259 | 26 | DUF4863 family protein [Aromatoleum toluclasticum] |
| 08189 | 26 | hydrogenase maturation protein [Denitromonas halophilus] |
| 05118 | 26 | BREX-2 system adenine-specific DNA-methyltransferase PglX [Azoarcus indigens] |
| 00403 | 26 | zinc ABC transporter substrate-binding protein [Denitromonas halophilus] |
| 08769 | 26 | hypothetical protein [Aromatoleum toluclasticum] |
| 01592 | 26 | hypothetical protein [Aromatoleum aromaticum] |
| 09850 | 26 | type II toxin-antitoxin system prevent-host-death family antitoxin [Azoarcus communis] |
| 02333 | 26 | hypothetical protein [Zoogloea ramigera] |
| 04509 | 26 | hypothetical protein [Azoarcus communis] |
| 03138 | 26 | DUF882 domain-containing protein [Aromatoleum toluclasticum] |
| 02493 | 26 | N-acetylglutaminylglutamine synthetase [Denitromonas halophilus] |
| 06057 | 26 | ATP-binding cassette domain-containing protein [Denitromonas halophilus] |
| 04952 | 26 | phosphatidylserine decarboxylase [Denitromonas halophilus] |
| 08542 | 26 | HlyD family efflux transporter periplasmic adaptor subunit [Azoarcus halotolerans] |
| 03750 | 26 | cytochrome C [Azoarcus communis] |
| 04887 | 26 | nitrogen fixation protein NifZ [Azoarcus communis] |
| 09851 | 26 | MULTISPECIES: DUF3158 family protein [Proteobacteria] |
| 00471 | 26 | Mut7-C ubiquitin/RNAse domain-containing protein [Denitromonas halophilus] |
| 00208 | 26 | ChbG/HpnK family deacetylase [Azoarcus communis] |
| 03772 | 26 | M15 family metallopeptidase [Denitromonas halophilus] |
| 02816 | 25 | aromatic amino acid ammonia-lyase [Denitromonas halophilus] |
| 08262 | 25 | tetratricopeptide repeat protein [Denitromonas halophilus] |
| 04688 | 25 | MULTISPECIES: RAQPRD family integrative conjugative element protein [Proteobacteria] |
| 08027 | 25 | aromatic ring-hydroxylating dioxygenase subunit alpha [Azoarcus indigens] |
| 03637 | 25 | cytochrome c [Azoarcus communis] |
| 08157 | 25 | class I SAM-dependent methyltransferase [Azoarcus indigens] |
| 07906 | 25 | MmgE/PrpD family protein [Aromatoleum toluvorans] |
| 07443 | 25 | ectoine/hydroxyectoine ABC transporter substrate-binding protein EhuB [Azoarcus olearius] |
| 08395 | 25 | class I SAM-dependent methyltransferase [Denitromonas halophilus] |
| 07010 | 25 | amino acid permease [Aromatoleum evansii] |
| 09256 | 25 | AraC family transcriptional regulator [Denitromonas halophilus] |
| 09087 | 25 | histidine phosphatase family protein [Azoarcus communis] |
| 09652 | 25 | hypothetical protein [Azoarcus halotolerans] |
| 09430 | 25 | exo-alpha-sialidase [Azoarcus sp. DD4] |
| 04885 | 25 | DNA oxidative demethylase AlkB [Aromatoleum toluclasticum] |

| | | |
|---|---|---|
| 04480 | 25 | toluene tolerance protein [Azoarcus communis] |
| 02272 | 25 | tyrosine-type recombinase/integrase [Aromatoleum aromaticum] |
| 09525 | 25 | prepilin-type N-terminal cleavage/methylation domain-containing protein [Aromatoleum toluclasticum] |
| 06393 | 25 | cytochrome c [Aromatoleum aromaticum] |
| 04484 | 25 | hypothetical protein [Azoarcus communis] |
| 10553 | 25 | AraC family transcriptional regulator [Denitromonas halophilus] |
| 00186 | 25 | malate synthase A [Azoarcus communis] |
| 01152 | 25 | YCF48-related protein [Aromatoleum bremense] |
| 07821 | 25 | sel1 repeat family protein [Nitrogeniibacter aestuarii] |
| 10008 | 25 | hypothetical protein [Aromatoleum toluclasticum] |
| 07857 | 25 | transcriptional repressor [Denitromonas halophilus] |
| 07219 | 25 | type II toxin-antitoxin system RelE/ParE family toxin [Aromatoleum evansii] |
| 10351 | 25 | MULTISPECIES: Txe/YoeB family addiction module toxin [Thauera] |
| 05858 | 25 | MULTISPECIES: bifunctional 4-hydroxy-2-oxoglutarate aldolase/2-dehydro-3-deoxy-phosphogluconate aldolase [Niveibacterium] |
| 04374 | 25 | FecR domain-containing protein [Denitromonas halophilus] |
| 07070 | 25 | UDP-glucuronosyltransferase [Azoarcus communis] |
| 02411 | 25 | DUF5710 domain-containing protein [Denitromonas halophilus] |
| 01181 | 25 | hypothetical protein [Azoarcus communis] |
| 06977 | 25 | TlpA family protein disulfide reductase [Denitromonas halophilus] |
| 08324 | 25 | hydrolase [Denitromonas halophilus] |
| 07400 | 25 | VWA domain-containing protein [Azoarcus communis] |
| 06517 | 25 | HlyD family efflux transporter periplasmic adaptor subunit [Zoogloea oleivorans] |
| 03696 | 25 | TetR/AcrR family transcriptional regulator [Thauera terpenica] |
| 09436 | 25 | MULTISPECIES: TIGR03758 family integrating conjugative element protein [Proteobacteria] |
| 04416 | 25 | rhodanese-like domain-containing protein [Aromatoleum toluclasticum] |
| 06891 | 25 | EF-hand domain-containing protein [Azoarcus communis] |
| 09559 | 25 | glycogen synthase GlgA [Azoarcus indigens] |
| 03957 | 25 | hypothetical protein [Denitromonas halophilus] |
| 03983 | 25 | MULTISPECIES: site-specific integrase [Thauera] |
| 08300 | 25 | type II toxin-antitoxin system Phd/YefM family antitoxin [Aromatoleum bremense] |
| 01215 | 25 | AAA family ATPase [Aromatoleum toluclasticum] |
| 03885 | 25 | MULTISPECIES: ParB family protein [Rhodocyclales] |
| 02617 | 25 | Na(+)-translocating NADH-quinone reductase subunit A [Azoarcus communis] |
| 07210 | 25 | YdcH family protein [Denitromonas halophilus] |
| 07026 | 25 | carbon starvation protein A [Aromatoleum toluclasticum] |
| 06506 | 25 | alanine-phosphoribitol ligase [Azoarcus olearius] |
| 09961 | 25 | phosphatase PAP2 family protein [Denitromonas halophilus] |
| 01508 | 25 | caspase family protein [Propionivibrio dicarboxylicus] |
| 01770 | 25 | DUF3334 family protein [Azoarcus communis] |
| 07893 | 25 | OsmC family protein [Denitromonas halophilus] |
| 05443 | 25 | Fe-S cluster assembly protein NifU [Azoarcus communis] |
| 06481 | 25 | MULTISPECIES: TIGR03752 family integrating conjugative element protein [Rhodocyclales] |
| 05344 | 25 | TetR/AcrR family transcriptional regulator [Denitromonas halophilus] |
| 00221 | 25 | hypothetical protein [Azoarcus olearius] |
| 01662 | 25 | CHASE2 domain-containing protein [Denitromonas halophilus] |
| 01313 | 25 | AAA family ATPase [Aromatoleum bremense] |
| 02003 | 24 | DUF1850 domain-containing protein [Azoarcus communis] |
| 10416 | 24 | transposase [Denitromonas halophilus] |
| 06995 | 24 | SLC13 family permease [Denitromonas halophilus] |
| 08050 | 24 | c-type cytochrome [Aromatoleum evansii] |
| 00356 | 24 | cysteine-rich CWC family protein [Aromatoleum aromaticum] |
| 08833 | 24 | FMN-binding glutamate synthase family protein [Azoarcus communis] |
| 09624 | 24 | MULTISPECIES: hypothetical protein [Rhodocyclales] |
| 02643 | 24 | GGDEF domain-containing protein [Azoarcus indigens] |
| 02752 | 24 | MULTISPECIES: cytochrome c [Thauera] |
| 07368 | 24 | glycogen debranching protein GlgX [Niveibacterium umoris] |
| 08391 | 24 | oxidoreductase [Azoarcus communis] |
| 09417 | 24 | MULTISPECIES: hypothetical protein [Proteobacteria] |
| 04407 | 24 | IMPACT family protein [Azoarcus indigens] |
| 05738 | 24 | ABC-type transport auxiliary lipoprotein family protein [Aromatoleum toluclasticum] |
| 05984 | 24 | hypothetical protein [Denitromonas halophilus] |
| 08356 | 24 | PEP-CTERM sorting domain-containing protein [Denitromonas halophilus] |
| 01965 | 24 | MFS transporter [Aromatoleum buckelii] |
| 02540 | 24 | type II toxin-antitoxin system RelE/ParE family toxin [Aromatoleum aromaticum] |
| 05034 | 24 | hypothetical protein [Denitromonas halophilus] |
| 09411 | 24 | SPOR domain-containing protein [Nitrogeniibacter aestuarii] |
| 03995 | 24 | U32 family peptidase [Aromatoleum aromaticum] |
| 06168 | 24 | carbohydrate porin [Uliginosibacterium gangwonense] |
| 10429 | 24 | DUF599 domain-containing protein [Aromatoleum toluclasticum] |
| 08985 | 24 | DUF3274 domain-containing protein [Azoarcus olearius] |
| 04895 | 24 | YbaN family protein [Denitromonas halophilus] |
| 00482 | 24 | Dyp-type peroxidase [Aromatoleum toluclasticum] |
| 00148 | 24 | arsenate reductase family protein [Azoarcus communis] |
| 08765 | 24 | Na(+)-translocating NADH-quinone reductase subunit C [Azoarcus communis] |
| 09143 | 24 | DUF805 domain-containing protein [Denitromonas halophilus] |
| 04938 | 24 | RNA polymerase sigma factor [Denitromonas halophilus] |
| 08018 | 24 | DUF2169 domain-containing protein [Azoarcus communis] |
| 06645 | 24 | VWA domain-containing protein [Azoarcus indigens] |
| 07771 | 24 | methyltransferase domain-containing protein [Azoarcus communis] |
| 10317 | 24 | pyruvate carboxylase [Denitromonas halophilus] |
| 07055 | 24 | bifunctional salicylyl-CoA 5-hydroxylase/oxidoreductase [Aromatoleum evansii] |
| 05357 | 24 | PA0069 family radical SAM protein [Denitromonas halophilus] |
| 02178 | 24 | MULTISPECIES: DUF2857 domain-containing protein [Proteobacteria] |
| 02157 | 24 | hypothetical protein [Azoarcus communis] |
| 07660 | 24 | DeoR/GlpR family DNA-binding transcription regulator [Uliginosibacterium gangwonense] |
| 01064 | 24 | 5-carboxymethyl-2-hydroxymuconate Delta-isomerase [Azoarcus communis] |
| 05261 | 24 | amidohydrolase family protein [Aromatoleum toluclasticum] |
| 06965 | 24 | DUF802 domain-containing protein [Azoarcus communis] |
| 04531 | 24 | SLAC1 anion channel family protein [Azoarcus communis] |
| 09763 | 24 | GNAT family N-acetyltransferase [Azoarcus communis] |
| 09396 | 24 | glycosyltransferase [Azoarcus olearius] |
| 03952 | 24 | DUF3348 domain-containing protein [Azoarcus communis] |
| 10113 | 24 | glycoside hydrolase family protein [Uliginosibacterium gangwonense] |
| 04544 | 24 | transketolase [Propionivibrio dicarboxylicus] |

| | | |
|---|---|---|
| **00328** | 24 | cytochrome P460 family protein [Denitromonas halophilus] |
| **04541** | 24 | MFS transporter [Azoarcus indigens] |
| **05074** | 24 | MULTISPECIES: thioredoxin domain-containing protein [Thauera] |
| **02805** | 24 | hypothetical protein [Propionivibrio dicarboxylicus] |
| **06856** | 24 | DUF2817 domain-containing protein [Azoarcus olearius] |
| **02035** | 24 | hypothetical protein [Denitromonas halophilus] |
| **07937** | 24 | KTSC domain-containing protein [Azoarcus indigens] |
| **08774** | 24 | PepSY domain-containing protein [Denitromonas halophilus] |
| **02689** | 24 | sugar transferase [Aromatoleum diolicum] |
| **00278** | 24 | type II toxin-antitoxin system HipA family toxin [Azoarcus communis] |
| **03844** | 24 | MULTISPECIES: hypothetical protein [Niveibacterium] |
| **02055** | 24 | DUF1778 domain-containing protein [Aromatoleum toluclasticum] |
| **05951** | 23 | amidohydrolase family protein [Denitromonas halophilus] |
| **02681** | 23 | serine protein kinase RIO [Thauera chlorobenzoica] |
| **05236** | 23 | IS5 family transposase [Aromatoleum aromaticum] |
| **10050** | 23 | 2'-5' RNA ligase family protein [Azoarcus indigens] |
| **05567** | 23 | hypothetical protein [Azoarcus communis] |
| **02942** | 23 | MULTISPECIES: hypothetical protein [Rhodocyclales] |
| **01531** | 23 | STAS domain-containing protein [Denitromonas halophilus] |
| **09588** | 23 | FUSC family protein [Aromatoleum toluclasticum] |
| **07196** | 23 | hypothetical protein [Azoarcus indigens] |
| **00754** | 23 | transporter substrate-binding domain-containing protein [Niveibacterium microcysteis] |
| **07779** | 23 | MULTISPECIES: prepilin-type N-terminal cleavage/methylation domain-containing protein [Niveibacterium] |
| **04924** | 23 | type II secretion system F family protein [Aromatoleum toluclasticum] |
| **01519** | 23 | GNAT family N-acetyltransferase [Niveibacterium umoris] |
| **07421** | 23 | DMT family transporter [Aromatoleum toluclasticum] |
| **06966** | 23 | MULTISPECIES: MarR family transcriptional regulator [Niveibacterium] |
| **05948** | 23 | nuclear transport factor 2 family protein [Denitromonas halophilus] |
| **02370** | 23 | CcdB family protein [Azoarcus communis] |
| **00327** | 23 | WecB/TagA/CpsF family glycosyltransferase [Aromatoleum toluclasticum] |
| **08214** | 23 | YdgA family protein [Denitromonas halophilus] |
| **00456** | 23 | endonuclease/exonuclease/phosphatase family protein [Denitromonas halophilus] |
| **03345** | 23 | VOC family protein [Aromatoleum toluclasticum] |
| **00981** | 23 | TIGR01212 family radical SAM protein [Aromatoleum toluclasticum] |
| **01364** | 23 | hypothetical protein [Denitromonas halophilus] |
| **06467** | 23 | CDGSH iron-sulfur domain-containing protein [Aromatoleum toluclasticum] |
| **03285** | 23 | BON domain-containing protein [Denitromonas halophilus] |
| **01088** | 23 | hypothetical protein [Aromatoleum toluclasticum] |
| **01858** | 23 | (Na+)-NQR maturation NqrM [Azoarcus communis] |
| **06926** | 23 | putative zinc-binding protein [Denitromonas halophilus] |
| **09867** | 23 | hypothetical protein [Aromatoleum aromaticum] |
| **05085** | 23 | phage baseplate assembly protein V [Aromatoleum toluclasticum] |
| **01350** | 23 | MULTISPECIES: hypothetical protein [Proteobacteria] |
| **07683** | 23 | glutaredoxin family protein [Azoarcus communis] |
| **07520** | 23 | antibiotic biosynthesis monooxygenase [Denitromonas halophilus] |
| **04298** | 23 | catalase [Azoarcus indigens] |
| **08328** | 23 | sodium-dependent transporter [Azoarcus communis] |
| **09707** | 23 | GGDEF domain-containing protein [Nitrogeniibacter aestuarii] |
| **06188** | 23 | response regulator [Aromatoleum toluclasticum] |
| **09524** | 23 | DUF1987 domain-containing protein [Azoarcus indigens] |
| **04290** | 23 | MULTISPECIES: ferritin family protein [Niveibacterium] |
| **02163** | 23 | WD40 repeat domain-containing protein [Aromatoleum evansii] |
| **09957** | 23 | DUF4442 domain-containing protein [Denitromonas halophilus] |
| **04221** | 23 | thymidine phosphorylase family protein [Azoarcus communis] |
| **04018** | 23 | flavin reductase family protein [Azoarcus indigens] |
| **02927** | 23 | DUF2894 domain-containing protein [Azoarcus communis] |
| **02761** | 23 | pilin [Uliginosibacterium gangwonense] |
| **01479** | 23 | phosphatase PAP2 family protein [Azoarcus communis] |
| **00851** | 23 | polysaccharide pyruvyl transferase family protein [Aromatoleum toluclasticum] |
| **10610** | 23 | tyrosine-protein phosphatase [Aromatoleum toluclasticum] |
| **05283** | 23 | NdvB [Thauera terpenica] |
| **02340** | 23 | Spy/CpxP family protein refolding chaperone [Denitromonas halophilus] |
| **04753** | 23 | PhnD/SsuA/transferrin family substrate-binding protein [Aromatoleum toluclasticum] |
| **04840** | 23 | DNA-binding domain-containing protein [Nitrogeniibacter aestuarii] |
| **00388** | 23 | hypothetical protein [Denitromonas halophilus] |
| **02556** | 23 | MULTISPECIES: antibiotic biosynthesis monooxygenase [Niveibacterium] |
| **09987** | 23 | monovalent cation/H(+) antiporter subunit G [Azoarcus communis] |
| **06023** | 22 | hypothetical protein [Denitromonas halophilus] |
| **00304** | 22 | bile acid:sodium symporter [Denitromonas halophilus] |
| **07831** | 22 | DNA adenine methylase, partial [Uliginosibacterium gangwonense] |
| **00322** | 22 | L-histidine N(alpha)-methyltransferase [Denitromonas halophilus] |
| **06110** | 22 | EpsI family protein [Azoarcus communis] |
| **06649** | 22 | cytochrome C [Aromatoleum evansii] |
| **03817** | 22 | sulfur oxidation c-type cytochrome SoxA [Azoarcus communis] |
| **07801** | 22 | nucleotidyltransferase domain-containing protein [Aromatoleum evansii] |
| **05874** | 22 | alpha-glucan family phosphorylase [Rhodocyclus purpureus] |
| **05652** | 22 | extracellular solute-binding protein [Uliginosibacterium gangwonense] |
| **10223** | 22 | MULTISPECIES: TetR/AcrR family transcriptional regulator [Rhodocyclales] |
| **06623** | 22 | tyrosine-type recombinase/integrase [Aromatoleum aromaticum] |
| **03088** | 22 | DUF393 domain-containing protein [Aromatoleum toluclasticum] |
| **08609** | 22 | methyl-accepting chemotaxis protein [Denitromonas halophilus] |
| **00544** | 22 | acetyltransferase/hydrolase [Aromatoleum aromaticum] |
| **03719** | 22 | EVE domain-containing protein [Denitromonas halophilus] |
| **10229** | 22 | acyl carrier protein [Aromatoleum toluclasticum] |
| **08024** | 22 | MULTISPECIES: type II secretion system protein GspM [Niveibacterium] |
| **06738** | 22 | hypothetical protein [Aromatoleum toluclasticum] |
| **03219** | 22 | helix-turn-helix transcriptional regulator [Azoarcus communis] |
| **03955** | 22 | AMP nucleosidase [Azoarcus communis] |
| **00168** | 22 | flagellar brake protein [Aromatoleum pumilum] |
| **08711** | 22 | MULTISPECIES: sensor domain-containing diguanylate cyclase [Niveibacterium] |
| **03134** | 22 | MULTISPECIES: hypothetical protein [Thauera] |
| **05748** | 22 | hypothetical protein [Aromatoleum toluclasticum] |
| **03499** | 22 | MULTISPECIES: winged helix DNA-binding protein [Azoarcus] |
| **09535** | 22 | sulfotransferase family 2 domain-containing protein [Denitromonas halophilus] |

| | | |
|---|---|---|
| 09613 | 22 | alpha-1,2-mannosidase [Azoarcus indigens] |
| 02504 | 22 | MULTISPECIES: hypothetical protein [Niveibacterium] |
| 06832 | 22 | PepSY domain-containing protein [Denitromonas halophilus] |
| 04516 | 22 | ParD-like family protein [Aromatoleum toluclasticum] |
| 02110 | 22 | type II secretion system protein N [Azoarcus indigens] |
| 10071 | 22 | lipo-like protein [Denitromonas halophilus] |
| 05246 | 22 | NAD(P)-dependent oxidoreductase [Aromatoleum toluclasticum] |
| 00931 | 22 | DUF4255 domain-containing protein [Aromatoleum toluclasticum] |
| 01299 | 22 | O-antigen ligase family protein [Azoarcus communis] |
| 07017 | 22 | ferrous iron transport protein A [Azoarcus communis] |
| 09555 | 22 | MULTISPECIES: phosphate-starvation-inducible PsiE family protein [Thauera] |
| 01210 | 22 | cytochrome c [Denitromonas halophilus] |
| 01879 | 22 | DUF2235 domain-containing protein [Aromatoleum toluclasticum] |
| 02597 | 22 | putative baseplate assembly protein [Aromatoleum toluclasticum] |
| 09564 | 22 | DUF2889 domain-containing protein [Aromatoleum toluclasticum] |
| 04422 | 22 | exosortase B [Azoarcus communis] |
| 06958 | 22 | cobalt chelatase [Denitromonas halophilus] |
| 04672 | 22 | MULTISPECIES: hypothetical protein [Rhodocyclales] |
| 01006 | 22 | nuclear transport factor 2 family protein [Aromatoleum toluclasticum] |
| 00914 | 22 | hypothetical protein [Aromatoleum diolicum] |
| 06831 | 22 | maltose/maltodextrin ABC transporter substrate-binding protein MalE [Niveibacterium umoris] |
| 08935 | 22 | hypothetical protein [Azoarcus olearius] |
| 02461 | 22 | HD domain-containing protein [Azoarcus communis] |
| 09999 | 22 | MerR family transcriptional regulator [Azoarcus indigens] |
| 06899 | 22 | hypothetical protein [Azoarcus communis] |
| 06366 | 22 | SiaB family protein kinase [Azoarcus indigens] |
| 05373 | 22 | alpha-2-macroglobulin family protein [Aromatoleum toluclasticum] |
| 08116 | 22 | nuclear transport factor 2 family protein [Aromatoleum toluclasticum] |
| 08172 | 22 | alpha/beta hydrolase [Azoarcus olearius] |
| 08980 | 22 | MULTISPECIES: Tn3 family transposase [Proteobacteria] |
| 09860 | 22 | sulfur oxidation c-type cytochrome SoxX [Azoarcus communis] |
| 05075 | 22 | phosphate acetyltransferase [Denitromonas halophilus] |
| 09654 | 22 | type IV pilin protein [Azoarcus communis] |
| 09520 | 22 | methyl-accepting chemotaxis protein [Aromatoleum toluclasticum] |
| 02580 | 22 | sulfatase-like hydrolase/transferase [Aromatoleum toluclasticum] |
| 03361 | 21 | nucleotidyltransferase domain-containing protein [Aromatoleum bremense] |
| 08852 | 21 | ATP-binding protein [Aromatoleum evansii] |
| 09187 | 21 | OsmC family protein [Denitromonas halophilus] |
| 02552 | 21 | MULTISPECIES: TniB family NTP-binding protein [Proteobacteria] |
| 03881 | 21 | MULTISPECIES: GTPase [Rhodocyclales] |
| 02099 | 21 | ATP-binding protein [Rhodocyclus purpureus] |
| 04522 | 21 | TfoX/Sxy family protein [Denitromonas halophilus] |
| 04550 | 21 | MULTISPECIES: type II toxin-antitoxin system RelE/ParE family toxin [Aromatoleum] |
| 05799 | 21 | diguanylate cyclase [Azoarcus indigens] |
| 02835 | 21 | cytochrome c3 family protein [Aromatoleum toluclasticum] |
| 05797 | 21 | type II toxin-antitoxin system CcdA family antitoxin [Azoarcus communis] |
| 04395 | 21 | GNAT family N-acetyltransferase [Azoarcus olearius] |
| 09899 | 21 | thioredoxin [Azoarcus communis] |
| 02457 | 21 | 4-vinyl reductase [Nitrogeniibacter aestuarii] |
| 02385 | 21 | AAA family ATPase [Thauera butanivorans] |
| 06720 | 21 | Flp pilus assembly protein CpaB [Aromatoleum toluclasticum] |
| 08338 | 21 | DcaP family trimeric outer membrane transporter [Azoarcus sp. KH32C] |
| 05298 | 21 | cupin domain-containing protein [Aromatoleum toluclasticum] |
| 07613 | 21 | NADH-quinone oxidoreductase subunit B family protein [Aromatoleum toluclasticum] |
| 01126 | 21 | 6-phosphogluconolactonase [Aromatoleum toluclasticum] |
| 05420 | 21 | MULTISPECIES: hypothetical protein [Azoarcus] |
| 09728 | 21 | PA2778 family cysteine peptidase [Aromatoleum toluclasticum] |
| 10526 | 21 | TonB-dependent receptor [Denitromonas halophilus] |
| 00976 | 21 | DUF445 domain-containing protein [Aromatoleum toluclasticum] |
| 04225 | 21 | sarcosine oxidase subunit delta [Azoarcus communis] |
| 02235 | 21 | GNAT family N-acetyltransferase [Uliginosibacterium gangwonense] |
| 09775 | 21 | glycosyltransferase family 4 protein [Azospira oryzae] |
| 09773 | 21 | toll/interleukin-1 receptor domain-containing protein [Denitromonas halophilus] |
| 02210 | 21 | PA2779 family protein [Aromatoleum toluclasticum] |
| 00534 | 21 | class I SAM-dependent methyltransferase [Azoarcus indigens] |
| 05067 | 21 | type II toxin-antitoxin system RelE/ParE family toxin [Aromatoleum bremense] |
| 08000 | 21 | isochorismatase family protein [Aromatoleum toluclasticum] |
| 06324 | 21 | type I-C CRISPR-associated endonuclease Cas1c [Azoarcus indigens] |
| 00375 | 21 | hydrogenase expression/formation protein [Azoarcus communis] |
| 06220 | 21 | AbrB family transcriptional regulator [Azoarcus communis] |
| 00042 | 21 | flagellar hook-length control protein FliK [Azoarcus communis] |
| 02645 | 21 | type II and III secretion system protein family protein [Aromatoleum toluclasticum] |
| 03799 | 21 | tyrosine-type recombinase/integrase [Azoarcus sp. Aa7] |
| 00815 | 21 | OprD family outer membrane porin [Denitromonas halophilus] |
| 08292 | 21 | hypothetical protein [Aromatoleum toluclasticum] |
| 00808 | 21 | MULTISPECIES: type II toxin-antitoxin system YhaV family toxin [Rhodocyclales] |
| 00323 | 21 | metallophosphoesterase [Denitromonas halophilus] |
| 04112 | 21 | histidine kinase [Aromatoleum toluclasticum] |
| 01631 | 21 | Fic family protein [Aromatoleum aromaticum] |
| 01640 | 21 | DUF3830 family protein [Denitromonas halophilus] |
| 00150 | 21 | helix-turn-helix domain-containing protein [Nitrogeniibacter aestuarii] |
| 01775 | 21 | DUF3501 family protein [Denitromonas halophilus] |
| 06465 | 21 | MULTISPECIES: site-specific integrase [Thauera] |
| 04833 | 21 | HD-GYP domain-containing protein [Aromatoleum toluclasticum] |
| 04839 | 21 | ABC transporter substrate-binding protein [Azoarcus communis] |
| 00255 | 21 | thermonuclease family protein [Denitromonas sp. IR12] |
| 01982 | 21 | 2Fe-2S iron-sulfur cluster-binding protein [Azoarcus communis] |
| 01552 | 21 | ferritin-like domain-containing protein [Aromatoleum toluclasticum] |
| 09646 | 21 | MULTISPECIES: hypothetical protein [Niveibacterium] |
| 01996 | 21 | site-specific integrase [Zoogloea oleivorans] |
| 01417 | 21 | MULTISPECIES: helix-turn-helix domain-containing protein [Proteobacteria] |
| 00057 | 21 | c-type cytochrome [Azoarcus communis] |
| 08506 | 21 | MFS transporter [Denitromonas halophilus] |
| 03642 | 21 | DotU family type VI secretion system protein [Azoarcus communis] |

| | | |
|---|---|---|
| 03371 | 20 | ATPase P [Denitromonas halophilus] |
| 07089 | 20 | MULTISPECIES: hypothetical protein [Rhodocyclales] |
| 03860 | 20 | permease [Denitromonas sp. IR12] |
| 09831 | 20 | FprA family A-type flavoprotein [Azoarcus communis] |
| 03876 | 20 | GatB/YqeY domain-containing protein [Azoarcus communis] |
| 04907 | 20 | DUF6502 family protein [Aromatoleum toluclasticum] |
| 03390 | 20 | sodium:proton antiporter NhaD [Rhodocyclus purpureus] |
| 00306 | 20 | acyl-CoA/acyl-ACP dehydrogenase [Azoarcus indigens] |
| 05277 | 20 | MULTISPECIES: DDE-type integrase/transposase/recombinase [Proteobacteria] |
| 07588 | 20 | DUF2182 domain-containing protein [Aromatoleum toluclasticum] |
| 02010 | 20 | HD-GYP domain-containing protein [Denitromonas halophilus] |
| 05308 | 20 | hypothetical protein [Azoarcus communis] |
| 04390 | 20 | hypothetical protein [Azoarcus indigens] |
| 06312 | 20 | YbdD/YjiX family protein [Aromatoleum toluclasticum] |
| 01763 | 20 | VOC family protein [Aromatoleum evansii] |
| 04818 | 20 | DUF2798 domain-containing protein [Azoarcus communis] |
| 03380 | 20 | Zn-dependent hydrolase [Azoarcus communis] |
| 01066 | 20 | helix-turn-helix domain-containing protein [Rhodocyclus purpureus] |
| 03493 | 20 | TRAP transporter small permease [Azoarcus communis] |
| 07668 | 20 | glycosyltransferase [Azoarcus communis] |
| 08142 | 20 | phage tail protein [Aromatoleum toluclasticum] |
| 03216 | 20 | DctP family TRAP transporter solute-binding subunit [Azoarcus communis] |
| 01692 | 20 | hypothetical protein [Azoarcus communis] |
| 01719 | 20 | DUF3567 family protein [Denitromonas halophilus] |
| 03498 | 20 | metallophosphoesterase [Aromatoleum toluclasticum] |
| 09533 | 20 | metallophosphoesterase [Azoarcus communis] |
| 01600 | 20 | hypothetical protein [Denitromonas halophilus] |
| 07233 | 20 | MULTISPECIES: S49 family peptidase [Niveibacterium] |
| 07890 | 20 | hydrogenase [Denitromonas halophilus] |
| 07523 | 20 | 2,4'-dihydroxyacetophenone dioxygenase family protein [Azoarcus communis] |
| 09633 | 20 | MULTISPECIES: formate hydrogenlyase [Aromatoleum] |
| 03830 | 20 | type II toxin-antitoxin system VapB family antitoxin [Aromatoleum toluclasticum] |
| 01974 | 20 | C39 family peptidase [Azoarcus communis] |
| 02123 | 20 | response regulator [Denitromonas halophilus] |
| 01352 | 20 | PsiF family protein [Azoarcus olearius] |
| 03119 | 20 | AP2 domain-containing protein [Nitrogeniibacter aestuarii] |
| 05128 | 20 | ribonuclease HI family protein [Azoarcus communis] |
| 00035 | 20 | tail fiber protein [Denitromonas halophilus] |
| 10422 | 20 | IS110 family transposase [Aromatoleum pumilum] |
| 07537 | 20 | TolC family protein [Aromatoleum toluclasticum] |
| 01910 | 20 | GAF domain-containing protein [Azoarcus communis] |
| 09689 | 20 | DUF1326 domain-containing protein [Aromatoleum toluclasticum] |
| 07199 | 20 | MULTISPECIES: DUF3365 domain-containing protein [Azoarcus] |
| 02976 | 20 | hypothetical protein [Denitromonas halophilus] |
| 05070 | 20 | ABC transporter substrate-binding protein [Azoarcus rhizosphaerae] |
| 01890 | 20 | AraC family transcriptional regulator [Candidatus Dactylopiibacterium carminicum] |
| 03471 | 20 | MULTISPECIES: P-type conjugative transfer protein TrbJ [Proteobacteria] |
| 07366 | 20 | MULTISPECIES: YeeE/YedE family protein [Niveibacterium] |
| 07569 | 20 | MULTISPECIES: hypothetical protein [Aromatoleum] |
| 06194 | 20 | YidB family protein [Denitromonas halophilus] |
| 10590 | 20 | toll/interleukin-1 receptor domain-containing protein [Azoarcus communis] |
| 09215 | 20 | TraB/GumN family protein [Denitromonas halophilus] |
| 07577 | 20 | VOC family protein [Azoarcus communis] |
| 00767 | 20 | hypothetical protein [Azoarcus communis] |
| 04450 | 20 | MULTISPECIES: RidA family protein [Proteobacteria] |
| 10034 | 20 | ROK family protein [Azoarcus sp. L1K30] |
| 05618 | 20 | SDR family NAD(P)-dependent oxidoreductase [Aromatoleum toluclasticum] |
| 06846 | 20 | glycosyltransferase [Denitromonas halophilus] |
| 05407 | 20 | phosphopantetheine-binding protein [Denitromonas halophilus] |
| 02849 | 20 | helix-turn-helix domain-containing protein [Thauera phenylacetica] |
| 03350 | 20 | creatininase family protein [Azoarcus communis] |
| 00147 | 20 | APC family permease [Nitrogeniibacter aestuarii] |
| 08692 | 20 | glycosyltransferase [Denitromonas halophilus] |
| 05630 | 20 | GbsR/MarR family transcriptional regulator [Azoarcus indigens] |
| 08771 | 20 | hypothetical protein [Azoarcus communis] |
| 05418 | 20 | DedA family protein [Azoarcus communis] |
| 10004 | 20 | acyl carrier protein [Azoarcus indigens] |
| 02527 | 20 | dihydrofolate reductase family protein [Azoarcus olearius] |
| 03693 | 20 | isoprenylcysteine carboxylmethyltransferase family protein [Azoarcus communis] |
| 06380 | 20 | acyl-CoA/acyl-ACP dehydrogenase [Azoarcus communis] |
| 02716 | 20 | MULTISPECIES: thioredoxin domain-containing protein [Rhodocyclales] |
| 09933 | 20 | alanine--glyoxylate aminotransferase family protein [Azoarcus indigens] |
| 02824 | 20 | hypothetical protein [Denitromonas halophilus] |
| 06900 | 20 | carbonic anhydrase [Nitrogeniibacter aestuarii] |
| 00928 | 20 | hypothetical protein [Aromatoleum toluclasticum] |
| 03293 | 20 | META domain-containing protein [Azoarcus communis] |
| 03195 | 19 | flagellar motor protein MotA [Denitromonas halophilus] |
| 07086 | 19 | SDR family oxidoreductase [Azoarcus indigens] |
| 04751 | 19 | 5-amino-6-(D-ribitylamino)uracil--L-tyrosine 4-hydroxyphenyl transferase CofH [Aromatoleum toluclasticum] |
| 01150 | 19 | ADP-ribosylglycohydrolase family protein [Zoogloea oleivorans] |
| 07602 | 19 | DUF3141 domain-containing protein [Azoarcus olearius] |
| 09554 | 19 | MULTISPECIES: hypothetical protein [Thauera] |
| 03264 | 19 | cryptochrome/photolyase family protein [Azoarcus communis] |
| 07032 | 19 | DUF3304 domain-containing protein [Denitromonas halophilus] |
| 07114 | 19 | dimethyl sulfoxide reductase anchor subunit [Aromatoleum toluclasticum] |
| 07361 | 19 | hypothetical protein [Azoarcus communis] |
| 06754 | 19 | LysR family transcriptional regulator [Aromatoleum toluclasticum] |
| 06282 | 19 | IS66 family insertion sequence element accessory protein TnpB [Aromatoleum bremense] |
| 06060 | 19 | hypothetical protein [Denitromonas halophilus] |
| 04827 | 19 | D-(-)-3-hydroxybutyrate oligomer hydrolase [Denitromonas halophilus] |
| 01753 | 19 | type II toxin-antitoxin system RelB/DinJ family antitoxin [Aromatoleum toluclasticum] |
| 01765 | 19 | molybdopterin-dependent oxidoreductase [Azoarcus communis] |
| 06590 | 19 | HupE/UreJ family protein [Aromatoleum toluclasticum] |
| 02280 | 19 | pyridoxamine 5'-phosphate oxidase family protein [Aromatoleum toluclasticum] |

| | | |
|---|---|---|
| **01399** | 19 | cyanase [Denitromonas halophilus] |
| **02276** | 19 | CoA-binding protein [Aromatoleum bremense] |
| **02344** | 19 | LytR C-terminal domain-containing protein [Aromatoleum bremense] |
| **02691** | 19 | HD domain-containing protein [Azoarcus halotolerans] |
| **04805** | 19 | AAA family ATPase [Aromatoleum toluclasticum] |
| **01593** | 19 | hypothetical protein [Aromatoleum evansii] |
| **09097** | 19 | ABC transporter substrate-binding protein [Azoarcus communis] |
| **04030** | 19 | 4'-phosphopantetheinyl transferase superfamily protein [Denitromonas halophilus] |
| **00449** | 19 | group III truncated hemoglobin [Azoarcus indigens] |
| **10535** | 19 | tyrosine-type recombinase/integrase [Aromatoleum evansii] |
| **00090** | 19 | DUF4347 domain-containing protein [Zoogloea oleivorans] |
| **07095** | 19 | MULTISPECIES: DUF3085 domain-containing protein [Rhodocyclales] |
| **07604** | 19 | adenylate/guanylate cyclase domain-containing protein [Denitromonas halophilus] |
| **04919** | 19 | mechanosensitive ion channel family protein [Azoarcus communis] |
| **05514** | 19 | hypothetical protein [Aromatoleum toluvorans] |
| **08254** | 19 | MULTISPECIES: lytic transglycosylase domain-containing protein [Betaproteobacteria] |
| **05647** | 19 | YMGG-like glycine zipper-containing protein [Propionivibrio dicarboxylicus] |
| **09495** | 19 | MULTISPECIES: helix-turn-helix domain-containing protein [Rhodocyclales] |
| **07848** | 19 | IclR family transcriptional regulator [Thauera aromatica] |
| **00616** | 19 | hypothetical protein [Denitromonas halophilus] |
| **02862** | 19 | MULTISPECIES: molybdopterin-binding protein [Azoarcus] |
| **04127** | 19 | hypothetical protein [Denitromonas halophilus] |
| **08965** | 19 | PRC-barrel domain-containing protein [Aromatoleum aromaticum] |
| **06445** | 19 | exo-alpha-sialidase [Azoarcus communis] |
| **04908** | 19 | FMN-binding negative transcriptional regulator [Aromatoleum toluclasticum] |
| **03130** | 19 | TSUP family transporter [Aromatoleum pumilum] |
| **06446** | 19 | marine proteobacterial sortase target protein [Denitromonas halophilus] |
| **09342** | 19 | MULTISPECIES: SIMPL domain-containing protein [Thauera] |
| **00130** | 19 | hypothetical protein [Aromatoleum toluclasticum] |
| **08777** | 19 | glycosyltransferase [Aromatoleum toluclasticum] |
| **07417** | 19 | hypothetical protein [Aromatoleum toluclasticum] |
| **09200** | 19 | conjugal transfer protein TraG [Azoarcus communis] |
| **00154** | 19 | alpha/beta hydrolase-fold protein [Thauera linaloolentis] |
| **01687** | 19 | DUF1854 domain-containing protein [Denitromonas halophilus] |
| **02810** | 19 | hypothetical protein [Aromatoleum toluclasticum] |
| **01452** | 19 | MULTISPECIES: carboxymuconolactone decarboxylase family protein [Niveibacterium] |
| **08740** | 19 | carbamoyltransferase HypF [Aromatoleum diolicum] |
| **00827** | 19 | phosphatidate cytidylyltransferase [Azoarcus olearius] |
| **09744** | 19 | GGDEF domain-containing protein [Denitromonas halophilus] |
| **06219** | 19 | adenine phosphoribosyltransferase [Denitromonas halophilus] |
| **08354** | 19 | type II toxin-antitoxin system RelE/ParE family toxin [Aromatoleum bremense] |
| **04010** | 19 | hypothetical protein [Denitromonas halophilus] |
| **09288** | 19 | alkaline phosphatase [Azoarcus olearius] |
| **06263** | 19 | MULTISPECIES: mercuric ion transporter MerT [Proteobacteria] |
| **08827** | 19 | hypothetical protein [Aromatoleum toluclasticum] |
| **06264** | 19 | 2-phospho-L-lactate guanylyltransferase [Aromatoleum toluclasticum] |
| **09703** | 18 | hypothetical protein [Denitromonas halophilus] |
| **05829** | 18 | hypothetical protein [Niveibacterium microcysteis] |
| **03276** | 18 | M23 family metallopeptidase [Denitromonas halophilus] |
| **00741** | 18 | glutathione S-transferase C-terminal domain-containing protein [Aromatoleum toluclasticum] |
| **00196** | 18 | 2,5-didehydrogluconate reductase DkgB [Azoarcus indigens] |
| **09199** | 18 | ceramidase domain-containing protein [Denitromonas halophilus] |
| **04517** | 18 | hypothetical protein [Denitromonas halophilus] |
| **06634** | 18 | acetate kinase [Azoarcus olearius] |
| **01820** | 18 | hypothetical protein [Aromatoleum evansii] |
| **09201** | 18 | hypothetical protein [Azoarcus communis] |
| **10162** | 18 | prolyl oligopeptidase family serine peptidase [Niveibacterium sp. COAC-50] |
| **02812** | 18 | MULTISPECIES: heavy-metal-associated domain-containing protein [Rhodocyclales] |
| **09427** | 18 | sodium:proton antiporter [Denitromonas halophilus] |
| **01573** | 18 | FapA family protein [Azoarcus olearius] |
| **02569** | 18 | urocanate hydratase [Aromatoleum toluclasticum] |
| **10147** | 18 | aromatic-ring-hydroxylating dioxygenase subunit beta [Azoarcus communis] |
| **00847** | 18 | MULTISPECIES: metalloregulator ArsR/SmtB family transcription factor [Azoarcus] |
| **01895** | 18 | DUF5610 domain-containing protein [Azoarcus indigens] |
| **01925** | 18 | alpha/beta hydrolase [Azoarcus communis] |
| **02647** | 18 | non-ribosomal peptide synthetase [Thauera phenylacetica] |
| **08728** | 18 | YkgJ family cysteine cluster protein [Azoarcus communis] |
| **04985** | 18 | DoxX family protein [Azoarcus olearius] |
| **09602** | 18 | tautomerase family protein [Denitromonas sp. IR12] |
| **01493** | 18 | PhnD/SsuA/transferrin family substrate-binding protein [Denitromonas halophilus] |
| **06237** | 18 | glycine zipper 2TM domain-containing protein [Denitromonas halophilus] |
| **08203** | 18 | MULTISPECIES: type II toxin-antitoxin system Phd/YefM family antitoxin [Thauera] |
| **07807** | 18 | UDP-2,4-diacetamido-2,4,6-trideoxy-beta-L-altropyranose hydrolase [Denitromonas halophilus] |
| **06765** | 18 | AAA family ATPase [Denitromonas halophilus] |
| **04146** | 18 | MULTISPECIES: EexN family lipoprotein [Proteobacteria] |
| **04607** | 18 | NAD-dependent epimerase/dehydratase family protein [Nitrogeniibacter aestuarii] |
| **07509** | 18 | DNA mismatch endonuclease Vsr [Azoarcus nasutitermitis] |
| **05497** | 18 | hypothetical protein [Azoarcus indigens] |
| **01568** | 18 | PEP-utilizing enzyme [Aromatoleum aromaticum] |
| **05031** | 18 | ATP-binding protein [Azoarcus rhizosphaerae] |
| **03522** | 18 | MarR family transcriptional regulator [Aromatoleum toluclasticum] |
| **10123** | 18 | MULTISPECIES: hypothetical protein [Zoogloeaceae] |
| **02601** | 18 | glycosyltransferase [Aromatoleum aromaticum] |
| **09241** | 18 | amino acid adenylation domain-containing protein, partial [Azoarcus indigens] |
| **01401** | 18 | MULTISPECIES: DUF1097 domain-containing protein [Azoarcus] |
| **01581** | 18 | hypothetical protein [Aromatoleum toluclasticum] |
| **06061** | 18 | N-carbamoyl-D-amino-acid hydrolase [Azoarcus communis] |
| **05069** | 18 | hypothetical protein [Azoarcus communis] |
| **04027** | 18 | filamentous hemagglutinin N-terminal domain-containing protein, partial [Azoarcus indigens] |
| **07090** | 18 | DUF1501 domain-containing protein [Nitrogeniibacter aestuarii] |
| **05122** | 18 | ferredoxin family protein [Aromatoleum toluvorans] |
| **02874** | 18 | DUF1043 family protein [Denitromonas halophilus] |
| **02967** | 18 | DUF481 domain-containing protein [Azoarcus communis] |
| **08171** | 18 | ribulose bisphosphate carboxylase small subunit [Azoarcus communis] |

| | | |
|---|---|---|
| 08584 | 18 | arsenical resistance protein ArsH [Azoarcus communis] |
| 00118 | 18 | MULTISPECIES: DUF4148 domain-containing protein [Thauera] |
| 01681 | 18 | Hsp70 family protein [Aromatoleum buckelii] |
| 04897 | 18 | 1-acyl-sn-glycerol-3-phosphate acyltransferase [Azoarcus olearius] |
| 09113 | 18 | DUF1295 domain-containing protein [Aromatoleum diolicum] |
| 09779 | 18 | DUF4126 domain-containing protein [Denitromonas halophilus] |
| 10477 | 18 | cytochrome c biogenesis protein ResB [Aromatoleum diolicum] |
| 05203 | 18 | VWA domain-containing protein [Azoarcus olearius] |
| 10083 | 18 | ABC transporter substrate-binding protein [Aromatoleum petrolei] |
| 07626 | 18 | type VI secretion system lipoprotein TssJ [Azoarcus communis] |
| 04336 | 18 | 4a-hydroxytetrahydrobiopterin dehydratase [Azoarcus olearius] |
| 08579 | 18 | 1-acyl-sn-glycerol-3-phosphate acyltransferase [Denitromonas halophilus] |
| 01207 | 18 | TonB-dependent receptor [Azoarcus communis] |
| 10566 | 18 | tetratricopeptide repeat protein [Aromatoleum petrolei] |
| 03951 | 18 | hypothetical protein [Azoarcus communis] |
| 10475 | 18 | THxN family PEP-CTERM protein [Denitromonas halophilus] |
| 02490 | 18 | ATP-binding protein [Azoarcus olearius] |
| 00998 | 18 | MULTISPECIES: P-type conjugative transfer protein TrbG [Betaproteobacteria] |
| 02733 | 18 | hypothetical protein [Aromatoleum toluclasticum] |
| 02353 | 18 | pilus assembly protein [Aromatoleum toluclasticum] |
| 06456 | 18 | xanthine/uracil/vitamin C permease [Azoarcus communis] |
| 09828 | 18 | phosphatase PAP2 family protein [Aromatoleum tolulyticum] |
| 01057 | 18 | iron export ABC transporter permease subunit FetB [Denitromonas halophilus] |
| 00146 | 18 | PepSY domain-containing protein [Azoarcus indigens] |
| 00083 | 18 | ribonucleotide reductase subunit alpha [Denitromonas halophilus] |
| 08952 | 18 | Crp/Fnr family transcriptional regulator [Azoarcus communis] |
| 05376 | 18 | hypothetical protein [Aromatoleum toluclasticum] |
| 05386 | 18 | IS256 family transposase [Aromatoleum buckelii] |
| 01007 | 18 | hypothetical protein [Aromatoleum toluclasticum] |
| 00524 | 18 | A24 family peptidase [Aromatoleum petrolei] |
| 03336 | 18 | nitrite reductase small subunit NirD [Azoarcus communis] |
| 04110 | 18 | PEP/pyruvate-binding domain-containing protein [Aromatoleum toluclasticum] |
| 07686 | 18 | hypothetical protein [Azoarcus olearius] |
| 08231 | 18 | peptidoglycan -binding protein [Denitromonas halophilus] |
| 08761 | 18 | hypothetical protein [Aromatoleum toluclasticum] |
| 02165 | 18 | metallophosphatase family protein [Azoarcus sp. KH32C] |
| 07546 | 18 | DUF58 domain-containing protein [Azoarcus communis] |
| 08574 | 18 | hypothetical protein [Aromatoleum toluclasticum] |
| 09259 | 17 | MULTISPECIES: thermostable hemolysin [Thauera] |
| 05895 | 17 | energy transducer TonB [Azoarcus olearius] |
| 10558 | 17 | hypothetical protein [Nitrogeniibacter aestuarii] |
| 09906 | 17 | RNA polymerase sigma factor [Azoarcus indigens] |
| 02756 | 17 | HD domain-containing protein [Aromatoleum toluclasticum] |
| 07475 | 17 | MCP four helix bundle domain-containing protein [Azoarcus sp. L1K30] |
| 01718 | 17 | potassium channel protein [Denitromonas halophilus] |
| 02501 | 17 | GntP family permease [Thauera butanivorans] |
| 04603 | 17 | class I SAM-dependent methyltransferase [Aromatoleum toluclasticum] |
| 09562 | 17 | MULTISPECIES: acyl-CoA thioesterase [Aromatoleum] |
| 04347 | 17 | hypothetical protein [Denitromonas halophilus] |
| 05028 | 17 | YXWGXW repeat-containing protein [Azoarcus communis] |
| 04608 | 17 | MULTISPECIES: PAS domain S-box protein [Rhodocyclales] |
| 04035 | 17 | DUF2145 domain-containing protein [Denitromonas halophilus] |
| 06451 | 17 | MULTISPECIES: rhodanese-like domain-containing protein [Niveibacterium] |
| 04049 | 17 | type II toxin-antitoxin system VapC family toxin [Azoarcus communis] |
| 01701 | 17 | ABC transporter permease [Azoarcus communis] |
| 05358 | 17 | PEP-utilizing enzyme [Aromatoleum petrolei] |
| 01946 | 17 | MULTISPECIES: pyridoxamine 5'-phosphate oxidase [Thauera] |
| 06026 | 17 | EthD family reductase [Denitromonas halophilus] |
| 08922 | 17 | AAA family ATPase [Azoarcus halotolerans] |
| 00266 | 17 | hypothetical protein [Azoarcus communis] |
| 07164 | 17 | alpha/beta hydrolase [Azoarcus communis] |
| 06928 | 17 | tetratricopeptide repeat protein [Denitromonas halophilus] |
| 03843 | 17 | hypothetical protein [Azoarcus communis] |
| 01977 | 17 | class I SAM-dependent methyltransferase [Azoarcus communis] |
| 05965 | 17 | sarcosine oxidase subunit gamma family protein [Aromatoleum petrolei] |
| 02731 | 17 | tryptophanase [Aromatoleum toluclasticum] |
| 09285 | 17 | aldo/keto reductase [Niveibacterium microcysteis] |
| 01937 | 17 | hypothetical protein [Denitromonas halophilus] |
| 08465 | 17 | MULTISPECIES: iron-containing redox enzyme family protein [Thauera] |
| 07136 | 17 | OST-HTH/LOTUS domain-containing protein [Aromatoleum toluclasticum] |
| 01935 | 17 | H-NS histone family protein [Aromatoleum aromaticum] |
| 00076 | 17 | DEAD/DEAH box helicase [Aromatoleum petrolei] |
| 08347 | 17 | DUF6285 domain-containing protein [Aromatoleum toluclasticum] |
| 02549 | 17 | AAA family ATPase [Niveibacterium microcysteis] |
| 05543 | 17 | dihydrolipoamide acyltransferase [Aromatoleum aromaticum] |
| 07957 | 17 | nitrogen fixation protein NifZ [Aromatoleum toluclasticum] |
| 07048 | 17 | GFA family protein [Denitromonas halophilus] |
| 02581 | 17 | lipoyl domain-containing protein [Azoarcus communis] |
| 06753 | 17 | transporter substrate-binding domain-containing protein [Denitromonas halophilus] |
| 02237 | 17 | TRAP transporter substrate-binding protein DctP [Uliginosibacterium gangwonense] |
| 09250 | 17 | PAS domain S-box protein [Denitromonas halophilus] |
| 08139 | 17 | DUF4142 domain-containing protein [Azoarcus olearius] |
| 00970 | 17 | ion transporter [Azoarcus communis] |
| 01774 | 17 | MULTISPECIES: replication initiator protein A [Proteobacteria] |
| 04072 | 17 | SRPBCC family protein [Nitrogeniibacter aestuarii] |
| 01338 | 17 | hypothetical protein [Nitrogeniibacter aestuarii] |
| 07025 | 17 | fumarate reductase [Azoarcus toluclasticus] |
| 09165 | 17 | cytochrome C oxidase subunit IV family protein [Denitromonas halophilus] |
| 04510 | 17 | filamentous haemagglutinin family protein [Azoarcus indigens] |
| 00203 | 17 | aldose 1-epimerase [Aromatoleum diolicum] |
| 05436 | 17 | AraC family transcriptional regulator [Aromatoleum toluclasticum] |
| 01251 | 17 | AlpA family phage regulatory protein [Azoarcus taiwanensis] |
| 02254 | 17 | isocitrate lyase/PEP mutase family protein [Azoarcus taiwanensis] |
| 00801 | 17 | phosphate ABC transporter substrate-binding protein [Aromatoleum toluclasticum] |

| | | |
|---|---|---|
| 09556 | 17 | DUF3592 domain-containing protein [Azoarcus communis] |
| 07067 | 17 | extracellular solute-binding protein [Aromatoleum toluclasticum] |
| 06883 | 17 | hypothetical protein [Aromatoleum toluclasticum] |
| 10005 | 17 | transglutaminase family protein [Azoarcus communis] |
| 03939 | 17 | efflux RND transporter periplasmic adaptor subunit [Oryzomicrobium terrae] |
| 07234 | 17 | response regulator [Aromatoleum toluclasticum] |
| 03308 | 17 | MULTISPECIES: formate C-acetyltransferase [Niveibacterium] |
| 06355 | 17 | hypothetical protein [Aromatoleum toluclasticum] |
| 07108 | 17 | hypothetical protein [Denitromonas halophilus] |
| 01157 | 17 | YbaK/EbsC family protein [Aromatoleum toluclasticum] |
| 09145 | 17 | hypothetical protein [Aromatoleum toluclasticum] |
| 00174 | 17 | cache domain-containing protein [Azoarcus sp. KH32C] |
| 02140 | 17 | TonB-dependent receptor [Azoarcus indigens] |
| 07290 | 17 | fumarate reductase subunit FrdD [Aromatoleum toluclasticum] |
| 02282 | 17 | hypothetical protein [Denitromonas halophilus] |
| 02998 | 17 | conjugal transfer protein TrbE [Azoarcus communis] |
| 03238 | 17 | HDOD domain-containing protein [Azoarcus indigens] |
| 05421 | 17 | pyridoxamine 5'-phosphate oxidase family protein [Thauera propionica] |
| 01781 | 16 | restriction endonuclease subunit S [Azoarcus taiwanensis] |
| 02714 | 16 | hypothetical protein [Azoarcus indigens] |
| 01635 | 16 | tyrosine-type recombinase/integrase [Denitromonas halophilus] |
| 05615 | 16 | hypothetical protein [Aromatoleum toluclasticum] |
| 03183 | 16 | thioredoxin fold domain-containing protein [Aromatoleum toluclasticum] |
| 10468 | 16 | AbrB/MazE/SpoVT family DNA-binding domain-containing protein [Azoarcus communis] |
| 09422 | 16 | methylamine utilization protein MauE [Azoarcus indigens] |
| 03172 | 16 | ferritin [Aromatoleum aromaticum] |
| 09493 | 16 | ATP-binding protein, partial [Zoogloea oleivorans] |
| 01822 | 16 | MULTISPECIES: hypothetical protein [Aromatoleum] |
| 08766 | 16 | BON domain-containing protein [Azoarcus indigens] |
| 01848 | 16 | methylamine utilization protein [Azoarcus communis] |
| 00782 | 16 | Crp/Fnr family transcriptional regulator [Nitrogeniibacter aestuarii] |
| 00237 | 16 | 2-phospho-L-lactate transferase [Aromatoleum toluclasticum] |
| 03236 | 16 | helix-turn-helix transcriptional regulator [Azoarcus communis] |
| 08450 | 16 | coenzyme F420-0:L-glutamate ligase [Aromatoleum toluclasticum] |
| 04741 | 16 | flagellar filament capping protein FliD [Azoarcus indigens] |
| 04740 | 16 | AmiS/UreI family transporter [Aromatoleum toluclasticum] |
| 03790 | 16 | bacterioferritin [Aromatoleum toluclasticum] |
| 04725 | 16 | DUF2946 domain-containing protein [Azoarcus indigens] |
| 03730 | 16 | amine dehydrogenase [Azoarcus communis] |
| 10003 | 16 | rhomboid family intramembrane serine protease [Aromatoleum toluclasticum] |
| 00339 | 16 | CbbX protein [Azoarcus communis] |
| 02514 | 16 | molecular chaperone TorD family protein [Aromatoleum toluclasticum] |
| 08671 | 16 | cytochrome oxidase small assembly protein [Azoarcus communis] |
| 08325 | 16 | MOSC domain-containing protein [Aromatoleum evansii] |
| 01132 | 16 | MULTISPECIES: type II toxin-antitoxin system YafQ family toxin [Bacteria] |
| 00582 | 16 | ankyrin repeat domain-containing protein [Azoarcus communis] |
| 09929 | 16 | MFS transporter [Aromatoleum evansii] |
| 03918 | 16 | hypothetical protein [Azoarcus sp. KH32C] |
| 04000 | 16 | EAL domain-containing protein [Aromatoleum toluclasticum] |
| 01208 | 16 | hypothetical protein [Denitromonas halophilus] |
| 04224 | 16 | DUF4150 domain-containing protein [Azoarcus communis] |
| 00598 | 16 | sulfurtransferase [Azoarcus communis] |
| 03903 | 16 | beta-ketoacyl-ACP synthase III [Azoarcus communis] |
| 05198 | 16 | CsgG/HfaB family protein [Azoarcus halotolerans] |
| 04445 | 16 | acyltransferase [Oryzomicrobium terrae] |
| 05750 | 16 | PEP-CTERM sorting domain-containing protein [Azoarcus olearius] |
| 06798 | 16 | DEAD/DEAH box helicase [Thauera phenylacetica] |
| 10303 | 16 | MULTISPECIES: TIGR00266 family protein [Niveibacterium] |
| 08889 | 16 | hypothetical protein [Aromatoleum aromaticum] |
| 05970 | 16 | CDP-alcohol phosphatidyltransferase family protein [Azoarcus olearius] |
| 08670 | 16 | phosphatase PAP2/dual specificity phosphatase family protein [Azoarcus olearius] |
| 06359 | 16 | PilZ domain-containing protein [Aromatoleum toluclasticum] |
| 02181 | 16 | PepSY-associated TM helix domain-containing protein [Azoarcus communis] |
| 03379 | 16 | PIN domain-containing protein [Azoarcus communis] |
| 03981 | 16 | MULTISPECIES: DNA-binding protein [Proteobacteria] |
| 09900 | 16 | hypothetical protein [Azoarcus communis] |
| 09896 | 16 | SEL1-like repeat protein [Thauera aminoaromatica] |
| 02453 | 16 | antibiotic biosynthesis monooxygenase [Azoarcus communis] |
| 02865 | 16 | GntR family transcriptional regulator [Aromatoleum toluclasticum] |
| 04270 | 16 | hypothetical protein [Denitromonas halophilus] |
| 08640 | 16 | hypothetical protein [Denitromonas halophilus] |
| 02427 | 16 | MurR/RpiR family transcriptional regulator [Azoarcus communis] |
| 08634 | 16 | DUF120 domain-containing protein [Aromatoleum buckelii] |
| 07855 | 16 | MULTISPECIES: phosphohydrolase [Betaproteobacteria] |
| 05332 | 16 | recombinase family protein [Aromatoleum bremense] |
| 07645 | 16 | CHAT domain-containing protein [Propionivibrio dicarboxylicus] |
| 08056 | 16 | VOC family protein [Azoarcus communis] |
| 04239 | 16 | amine dehydrogenase [Azoarcus communis] |
| 07637 | 16 | sulfotransferase [Aromatoleum diolicum] |
| 09919 | 16 | DUF3325 domain-containing protein [Azoarcus indigens] |
| 02914 | 16 | MULTISPECIES: PIN domain-containing protein [Thauera] |
| 09009 | 16 | response regulator [Aromatoleum toluclasticum] |
| 08683 | 16 | ATP-binding protein [Aromatoleum bremense] |
| 10486 | 16 | class I SAM-dependent methyltransferase [Aromatoleum buckelii] |
| 04209 | 16 | prepilin-type N-terminal cleavage/methylation domain-containing protein [Azoarcus indigens] |
| 00058 | 16 | MULTISPECIES: type IV toxin-antitoxin system AbiEi family antitoxin domain-containing protein [Proteobacteria] |
| 03058 | 16 | MULTISPECIES: Ni/Fe-hydrogenase cytochrome b subunit [Azospira] |
| 09234 | 16 | glucan biosynthesis protein D [Azoarcus indigens] |
| 00887 | 16 | universal stress protein [Azoarcus communis] |
| 09236 | 16 | MULTISPECIES: ABC transporter substrate-binding protein [Rhodocyclales] |
| 06314 | 16 | hypothetical protein [Azoarcus communis] |
| 03444 | 16 | sulfite exporter TauE/SafE family protein [Denitromonas halophilus] |
| 08210 | 16 | MULTISPECIES: methylamine dehydrogenase accessory protein MauD [Azoarcus] |
| 00156 | 16 | DUF2256 domain-containing protein [Azoarcus communis] |

| | | |
|---|---|---|
| 08494 | 16 | GNAT family N-acetyltransferase [Azoarcus communis] |
| 03717 | 16 | DUF2271 domain-containing protein [Azoarcus communis] |
| 06311 | 16 | TetR/AcrR family transcriptional regulator [Azoarcus indigens] |
| 06845 | 16 | MULTISPECIES: plasmid stabilization protein [Azoarcus] |
| 07460 | 16 | NTP transferase domain-containing protein [Denitromonas halophilus] |
| 10394 | 16 | DUF2721 domain-containing protein [Denitromonas halophilus] |
| 05503 | 16 | DUF799 domain-containing protein [Azoarcus indigens] |
| 00371 | 16 | Na+/H+ antiporter subunit E [Azoarcus communis] |
| 05506 | 16 | diguanylate cyclase [Azoarcus communis] |
| 00717 | 16 | hypothetical protein [Aromatoleum toluclasticum] |
| 10157 | 16 | glycerophosphodiester phosphodiesterase [Azoarcus olearius] |
| 04026 | 16 | YeeE/YedE family protein [Nitrogeniibacter aestuarii] |
| 09210 | 16 | hypothetical protein [Nitrogeniibacter aestuarii] |
| 08695 | 16 | DUF6279 family lipoprotein [Azoarcus communis] |
| 05833 | 16 | hypothetical protein [Azoarcus communis] |
| 00484 | 16 | MFS transporter [Aromatoleum toluclasticum] |
| 09208 | 16 | protein NosL [Denitromonas halophilus] |
| 00481 | 16 | PEP-CTERM sorting domain-containing protein [Aromatoleum diolicum] |
| 10336 | 16 | BrnA antitoxin family protein [Rhodocyclus purpureus] |
| 03272 | 16 | TRAP transporter small permease [Azoarcus communis] |
| 02815 | 16 | MULTISPECIES: hemolysin III family protein [Niveibacterium] |
| 06632 | 16 | VOC family protein [Azoarcus olearius] |
| 08688 | 16 | LLM class flavin-dependent oxidoreductase [Azoarcus indigens] |
| 02548 | 16 | VOC family protein [Aromatoleum toluclasticum] |
| 05465 | 16 | DUF1127 domain-containing protein [Denitromonas halophilus] |
| 04016 | 16 | DUF4810 domain-containing protein [Azoarcus indigens] |
| 00933 | 16 | hypothetical protein [Aromatoleum toluclasticum] |
| 08197 | 15 | nickel-responsive transcriptional regulator NikR [Azoarcus communis] |
| 08194 | 15 | DUF2345 domain-containing protein, partial [Denitromonas halophilus] |
| 06156 | 15 | phosphate/phosphite/phosphonate ABC transporter substrate-binding protein [Azoarcus olearius] |
| 08277 | 15 | cyclic nucleotide-binding protein [Aromatoleum toluolicum] |
| 08306 | 15 | formate dehydrogenase subunit delta [Aromatoleum pumilum] |
| 06315 | 15 | MATE family efflux transporter [Candidatus Dactylopiibacterium carminicum] |
| 03301 | 15 | ASKHA domain-containing protein [Aromatoleum toluclasticum] |
| 00172 | 15 | dipeptidase [Aromatoleum toluclasticum] |
| 06350 | 15 | type II toxin-antitoxin system RelE/ParE family toxin [Thauera aromatica] |
| 03633 | 15 | class I SAM-dependent methyltransferase [Azoarcus communis] |
| 03417 | 15 | hypothetical protein [Denitromonas halophilus] |
| 02605 | 15 | phage holin family protein [Azoarcus communis] |
| 07441 | 15 | DUF2157 domain-containing protein [Aromatoleum toluclasticum] |
| 02036 | 15 | TauD/TfdA family dioxygenase [Aromatoleum toluclasticum] |
| 07829 | 15 | DUF4198 domain-containing protein [Azoarcus communis] |
| 07512 | 15 | TonB-dependent receptor [Azoarcus communis] |
| 07093 | 15 | MULTISPECIES: ferritin-like domain-containing protein [Denitromonas] |
| 07591 | 15 | glycosyltransferase family 39 protein [Azoarcus communis] |
| 09889 | 15 | DMT family transporter [Denitromonas halophilus] |
| 07098 | 15 | paraquat-inducible protein A [Azoarcus halotolerans] |
| 09664 | 15 | MULTISPECIES: TrbC/VirB2 family protein [Proteobacteria] |
| 09663 | 15 | MULTISPECIES: M48 family metalloprotease [Betaproteobacteria] |
| 09660 | 15 | MULTISPECIES: S26 family signal peptidase [Betaproteobacteria] |
| 04585 | 15 | class I SAM-dependent methyltransferase [Denitromonas halophilus] |
| 10167 | 15 | site-specific DNA-methyltransferase [Aromatoleum bremense] |
| 07117 | 15 | AsmA family protein [Aromatoleum toluclasticum] |
| 02850 | 15 | DUF2141 domain-containing protein [Azoarcus communis] |
| 04589 | 15 | MULTISPECIES: AraC family transcriptional regulator [Niveibacterium] |
| 09253 | 15 | HD domain-containing protein [Denitromonas halophilus] |
| 02002 | 15 | bifunctional alpha/beta hydrolase/class I SAM-dependent methyltransferase [Azoarcus olearius] |
| 05525 | 15 | DEAD/DEAH box helicase [Azoarcus communis] |
| 05025 | 15 | PepSY-associated TM helix domain-containing protein [Aromatoleum toluclasticum] |
| 06967 | 15 | adenylate/guanylate cyclase domain-containing protein [Azoarcus indigens] |
| 02919 | 15 | DUF853 family protein [Denitromonas halophilus] |
| 08686 | 15 | DUF4401 domain-containing protein [Aromatoleum toluclasticum] |
| 07824 | 15 | SDR family NAD(P)-dependent oxidoreductase [Aromatoleum toluclasticum] |
| 02928 | 15 | MULTISPECIES: YiaA/YiaB family protein [Niveibacterium] |
| 09167 | 15 | IS4 family transposase [Denitromonas halophilus] |
| 09758 | 15 | general secretion pathway protein GspK [Aromatoleum buckelii] |
| 09757 | 15 | MULTISPECIES: phasin family protein [Proteobacteria] |
| 00776 | 15 | hypothetical protein [Denitromonas halophilus] |
| 04456 | 15 | cytochrome c [Denitromonas halophilus] |
| 00774 | 15 | DUF1513 domain-containing protein [Azoarcus communis] |
| 02194 | 15 | helix-turn-helix transcriptional regulator [Zoogloea dura] |
| 02712 | 15 | acyl-CoA dehydrogenase family protein [Aromatoleum toluclasticum] |
| 07042 | 15 | AI-2E family transporter [Aromatoleum toluclasticum] |
| 04091 | 15 | cupredoxin domain-containing protein [Aromatoleum diolicum] |
| 04074 | 15 | alpha/beta hydrolase [Denitromonas halophilus] |
| 00831 | 15 | filamentous hemagglutinin N-terminal domain-containing protein [Azoarcus sp. TTM-91] |
| 09169 | 15 | FAD-dependent oxidoreductase, partial [Denitromonas halophilus] |
| 00740 | 15 | DUF2282 domain-containing protein [Nitrogeniibacter aestuarii] |
| 02750 | 15 | hypothetical protein [Zoogloea ramigera] |
| 05235 | 15 | ATP-binding protein [Azoarcus communis] |
| 01288 | 15 | MULTISPECIES: chemotaxis protein CheX [Niveibacterium] |
| 07081 | 15 | DUF2857 domain-containing protein [Aromatoleum aromaticum] |
| 02787 | 15 | tetratricopeptide repeat protein [Azoarcus olearius] |
| 06980 | 15 | aromatic ring-hydroxylating dioxygenase subunit alpha [Aromatoleum toluclasticum] |
| 02790 | 15 | hypothetical protein [Azoarcus communis] |
| 04632 | 15 | ATP-binding protein [Aromatoleum toluolicum] |
| 10077 | 15 | MULTISPECIES: nucleotidyl transferase AbiEii/AbiGii toxin family protein [Proteobacteria] |
| 05453 | 15 | hypothetical protein [Azoarcus indigens] |
| 01549 | 15 | GspH/FimT family pseudopilin [Azoarcus indigens] |
| 00408 | 15 | serine acetyltransferase [Azoarcus indigens] |
| 01885 | 15 | cold shock domain-containing protein [Denitromonas halophilus] |
| 08439 | 15 | chromate resistance protein [Denitromonas halophilus] |
| 10395 | 15 | hypothetical protein [Azoarcus communis] |
| 02951 | 15 | hypothetical protein [Azoarcus indigens] |

| | | |
|---|---|---|
| 03786 | 15 | MULTISPECIES: conjugal transfer protein TrbF [Proteobacteria] |
| 07719 | 15 | phenylphosphate carboxylase subunit gamma [Aromatoleum buckelii] |
| 07225 | 15 | DUF1445 domain-containing protein [Azoarcus indigens] |
| 04198 | 15 | NF038104 family lipoprotein [Aromatoleum toluclasticum] |
| 05340 | 15 | sulfatase-like hydrolase/transferase [Aromatoleum toluclasticum] |
| 01668 | 15 | prepilin-type N-terminal cleavage/methylation domain-containing protein [Azoarcus indigens] |
| 08464 | 15 | hypothetical protein [Rhodocyclus purpureus] |
| 01634 | 15 | nucleotidyl transferase AbiEii/AbiGii toxin family protein [Thauera terpenica] |
| 05408 | 15 | c-type cytochrome [Azoarcus communis] |
| 08459 | 15 | PilN domain-containing protein [Azoarcus indigens] |
| 01106 | 15 | hypothetical protein [Aromatoleum toluclasticum] |
| 03073 | 15 | SemiSWEET transporter [Aromatoleum toluclasticum] |
| 01678 | 15 | SoxR reducing system RseC family protein [Azoarcus communis] |
| 07265 | 15 | hypothetical protein [Aromatoleum buckelii] |
| 09884 | 15 | DsrE family protein [Aromatoleum evansii] |
| 10461 | 15 | cytochrome c [Azoarcus communis] |
| 00549 | 15 | hypothetical protein [Aromatoleum evansii] |
| 05776 | 15 | DUF2726 domain-containing protein [Denitromonas halophilus] |
| 04286 | 15 | NADPH-dependent FMN reductase [Azoarcus indigens] |
| 04639 | 15 | hypothetical protein [Aromatoleum toluclasticum] |
| 10334 | 15 | amidohydrolase family protein [Azoarcus communis] |
| 09314 | 15 | DUF3999 domain-containing protein [Denitromonas halophilus] |
| 00479 | 15 | tetratricopeptide repeat protein [Azoarcus communis] |
| 06617 | 15 | secondary thiamine-phosphate synthase enzyme YjbQ [Denitromonas halophilus] |
| 10343 | 15 | GspH/FimT family pseudopilin [Denitromonas halophilus] |
| 04361 | 15 | MFS transporter [Aromatoleum toluclasticum] |
| 01677 | 15 | DUF3096 domain-containing protein [Aromatoleum toluclasticum] |
| 10368 | 15 | DUF3014 domain-containing protein [Azoarcus communis] |
| 01715 | 15 | TMAO reductase system periplasmic protein TorT [Aromatoleum toluclasticum] |
| 04306 | 14 | type II toxin-antitoxin system PemK/MazF family toxin [Azoarcus nasutitermitis] |
| 02420 | 14 | UxaA family hydrolase [Azoarcus communis] |
| 04424 | 14 | UbiA family prenyltransferase [Denitromonas halophilus] |
| 06343 | 14 | AAA domain-containing protein [Nitrogeniibacter aestuarii] |
| 00108 | 14 | ATP-binding protein [Niveibacterium sp. COAC-50] |
| 02252 | 14 | cupin domain-containing protein [Nitrogeniibacter aestuarii] |
| 08988 | 14 | biotin/lipoyl-binding protein, partial [Nitrogeniibacter aestuarii] |
| 06938 | 14 | ComEA family DNA-binding protein [Aromatoleum buckelii] |
| 03385 | 14 | DNA-binding protein [Azoarcus communis] |
| 09005 | 14 | type II toxin-antitoxin system Phd/YefM family antitoxin [Azoarcus rhizosphaerae] |
| 01164 | 14 | MBL fold metallo-hydrolase [Azoarcus indigens] |
| 03388 | 14 | esterase-like activity of phytase family protein [Azoarcus olearius] |
| 07622 | 14 | L-seryl-tRNA(Sec) selenium transferase [Azoarcus indigens] |
| 01155 | 14 | biopolymer transporter ExbD [Denitromonas halophilus] |
| 06964 | 14 | hypothetical protein [Denitromonas halophilus] |
| 01136 | 14 | hypothetical protein [Aromatoleum toluclasticum] |
| 02315 | 14 | hypothetical protein [Denitromonas halophilus] |
| 07553 | 14 | cysteine protease [Aromatoleum buckelii] |
| 03225 | 14 | hypothetical protein [Denitromonas halophilus] |
| 04546 | 14 | class I SAM-dependent methyltransferase [Aromatoleum bremense] |
| 09133 | 14 | CRISPR-associated helicase Cas3' [Azoarcus indigens] |
| 08160 | 14 | GNAT family N-acetyltransferase [Denitromonas halophilus] |
| 03472 | 14 | ABC transporter permease [Aromatoleum toluclasticum] |
| 09384 | 14 | DUF2135 domain-containing protein [Azoarcus indigens] |
| 01570 | 14 | molybdopterin-binding protein [Aromatoleum evansii] |
| 01847 | 14 | class I SAM-dependent methyltransferase [Aromatoleum toluclasticum] |
| 04762 | 14 | hypothetical protein [Denitromonas halophilus] |
| 03481 | 14 | hypothetical protein [Azoarcus indigens] |
| 08166 | 14 | MULTISPECIES: cytochrome-c oxidase [Aromatoleum] |
| 04653 | 14 | MULTISPECIES: hypothetical protein [Rhodocyclales] |
| 04902 | 14 | DUF6156 family protein [Azoarcus olearius] |
| 09521 | 14 | DUF927 domain-containing protein [Thauera aromatica] |
| 07287 | 14 | acyltransferase [Uliginosibacterium aquaticum] |
| 07249 | 14 | methyltransferase type 11 [Aromatoleum toluclasticum] |
| 04867 | 14 | carbon-nitrogen hydrolase family protein [Denitromonas halophilus] |
| 04860 | 14 | transposase [Thauera terpenica] |
| 07195 | 14 | diiron oxygenase [Azoarcus communis] |
| 07174 | 14 | 2-oxoacid:acceptor oxidoreductase subunit alpha [Aromatoleum toluclasticum] |
| 00080 | 14 | tetratricopeptide repeat protein [Aromatoleum toluclasticum] |
| 04537 | 14 | pectate lyase [Uliginosibacterium aquaticum] |
| 01274 | 14 | type VI secretion system-associated protein TagF [Azoarcus communis] |
| 06334 | 14 | PQQ-binding-like beta-propeller repeat protein [Azoarcus sp. DD4] |
| 07532 | 14 | hypothetical protein [Denitromonas halophilus] |
| 09679 | 14 | retroviral-like aspartic protease family protein [Aromatoleum toluclasticum] |
| 06329 | 14 | helix-turn-helix domain-containing protein [Azoarcus olearius] |
| 02115 | 14 | CRISPR-associated protein Cas4 [Azoarcus indigens] |
| 02008 | 14 | hypothetical protein [Azoarcus taiwanensis] |
| 09309 | 14 | tricarballylate utilization 4Fe-4S protein TcuB [Denitromonas halophilus] |
| 07470 | 14 | phage tail sheath subtilisin-like domain-containing protein [Azoarcus communis] |
| 08205 | 14 | DUF1705 domain-containing protein, partial [Thauera phenylacetica] |
| 01445 | 14 | PHP domain-containing protein [Aromatoleum toluclasticum] |
| 09628 | 14 | nuclear transport factor 2 family protein [Denitromonas halophilus] |
| 03534 | 14 | type II toxin-antitoxin system RelE/ParE family toxin [Aromatoleum toluclasticum] |
| 04633 | 14 | MULTISPECIES: polysaccharide export protein [Thauera] |
| 05363 | 14 | type II toxin-antitoxin system VapB family antitoxin [Aromatoleum bremense] |
| 08272 | 14 | hypothetical protein [Aromatoleum toluclasticum] |
| 05651 | 14 | carboxymuconolactone decarboxylase family protein [Aromatoleum toluclasticum] |
| 10116 | 14 | WbuC family cupin fold metalloprotein [Oryzomicrobium terrae] |
| 00625 | 14 | hypothetical protein [Azoarcus sp. KH32C] |
| 03974 | 14 | MULTISPECIES: cobalamin B12-binding domain-containing protein [Aromatoleum] |
| 02881 | 14 | hypothetical protein [Aromatoleum toluclasticum] |
| 07840 | 14 | TetR/AcrR family transcriptional regulator [Azoarcus communis] |
| 00631 | 14 | hypothetical protein [Aromatoleum toluclasticum] |
| 05685 | 14 | MarR family transcriptional regulator [Denitromonas halophilus] |
| 05676 | 14 | C13 family peptidase [Denitromonas halophilus] |

| | | |
|---|---|---|
| 06746 | 14 | DUF1566 domain-containing protein [Azoarcus communis] |
| 10184 | 14 | CRISPR-associated endonuclease Cas2 [Azoarcus indigens] |
| 02845 | 14 | sensor histidine kinase [Denitromonas halophilus] |
| 06400 | 14 | antibiotic biosynthesis monooxygenase [Azoarcus communis] |
| 10181 | 14 | AbrB/MazE/SpoVT family DNA-binding domain-containing protein [Aromatoleum pumilum] |
| 05385 | 14 | DUF6088 family protein [Thauera terpenica] |
| 06752 | 14 | retention module-containing protein, partial [Denitromonas halophilus] |
| 08679 | 14 | type VI secretion system Vgr family protein, partial [Denitromonas sp. IR12] |
| 04011 | 14 | TIGR02281 family clan AA aspartic protease [Azoarcus olearius] |
| 10169 | 14 | helix-turn-helix domain-containing protein [Denitromonas halophilus] |
| 00731 | 14 | peptidoglycan-binding protein [Azoarcus communis] |
| 00747 | 14 | hypothetical protein [Azoarcus communis] |
| 00613 | 14 | HEAT repeat domain-containing protein [Azoarcus communis] |
| 02906 | 14 | DUF2254 domain-containing protein [Denitromonas halophilus] |
| 10290 | 14 | type II toxin-antitoxin system HigB family toxin [Thauera aromatica] |
| 07922 | 14 | MULTISPECIES: TolC family protein [Rhodocyclales] |
| 10467 | 14 | DUF3429 domain-containing protein [Nitrogeniibacter aestuarii] |
| 06005 | 14 | type I-C CRISPR-associated protein Cas8c/Csd1 [Azoarcus indigens] |
| 03768 | 14 | RimK family alpha-L-glutamate ligase [Azoarcus olearius] |
| 10505 | 14 | DUF559 domain-containing protein [Azoarcus olearius] |
| 07934 | 14 | hypothetical protein [Azoarcus communis] |
| 03251 | 14 | PhzF family phenazine biosynthesis protein [Denitromonas halophilus] |
| 10425 | 14 | MULTISPECIES: hypothetical protein [Niveibacterium] |
| 00368 | 14 | hypothetical protein [Azoarcus indigens] |
| 03257 | 14 | hypothetical protein [Zoogloea oleivorans] |
| 03008 | 14 | BON domain-containing protein [Azoarcus olearius] |
| 10404 | 14 | MULTISPECIES: TrbI/VirB10 family protein [Proteobacteria] |
| 00382 | 14 | transposase, partial [Azoarcus halotolerans] |
| 00392 | 14 | rhomboid family intramembrane serine protease [Denitromonas halophilus] |
| 05890 | 14 | DM13 domain-containing protein [Denitromonas halophilus] |
| 03851 | 14 | ribbon-helix-helix domain-containing protein [Aromatoleum aromaticum] |
| 03861 | 14 | winged helix-turn-helix domain-containing protein [Denitromonas halophilus] |
| 08548 | 14 | hypothetical protein [Denitromonas halophilus] |
| 06644 | 14 | hypothetical protein [Azoarcus communis] |
| 03658 | 14 | TetR/AcrR family transcriptional regulator [Aromatoleum toluclasticum] |
| 09482 | 14 | hypothetical protein [Denitromonas halophilus] |
| 06793 | 14 | type I-C CRISPR-associated protein Cas7/Csd2 [Azoarcus indigens] |
| 04136 | 14 | transcriptional regulator HexR [Aromatoleum toluclasticum] |
| 05551 | 14 | hypothetical protein [Denitromonas halophilus] |
| 00999 | 14 | response regulator [Thauera propionica] |
| 05546 | 14 | MULTISPECIES: hypothetical protein [Rhodocyclales] |
| 04120 | 14 | glutamate-cysteine ligase family protein [Azoarcus olearius] |
| 04122 | 14 | MULTISPECIES: hypothetical protein [Rhodocyclales] |
| 07753 | 14 | modified peptide precursor CbpA [Aromatoleum bremense] |
| 04131 | 14 | TolC family protein [Zoogloea oleivorans] |
| 08278 | 14 | IS30 family transposase [Aromatoleum toluclasticum] |
| 06444 | 14 | CapA family protein [Aromatoleum bremense] |
| 06862 | 14 | tellurite resistance TerB family protein [Azoarcus communis] |
| 10555 | 14 | LuxR C-terminal-related transcriptional regulator [Azoarcus communis] |
| 00771 | 14 | hypothetical protein [Aromatoleum toluclasticum] |
| 10013 | 14 | DUF5938 domain-containing protein [Azoarcus sp. KH32C] |
| 02613 | 14 | MULTISPECIES: hypothetical protein [Thauera] |
| 04167 | 14 | 3-hydroxyacyl-CoA dehydrogenase/enoyl-CoA hydratase family protein [Aromatoleum evansii] |
| 06854 | 14 | ParB/RepB/Spo0J family partition protein [Azoarcus communis] |
| 06123 | 14 | lipoate--protein ligase [Azoarcus communis] |
| 06859 | 14 | hypothetical protein [Azoarcus olearius] |
| 00787 | 14 | ATP-binding protein [Aromatoleum toluclasticum] |
| 08773 | 14 | hypothetical protein [Denitromonas halophilus] |
| 04243 | 14 | cell envelope integrity protein CreD [Azoarcus communis] |
| 05404 | 14 | MULTISPECIES: hemerythrin domain-containing protein [Thauera] |
| 04106 | 14 | TMAO reductase system sensor histidine kinase/response regulator TorS [Aromatoleum toluclasticum] |
| 04101 | 14 | NCS2 family permease [Denitromonas halophilus] |
| 04915 | 13 | DUF2157 domain-containing protein [Azoarcus communis] |
| 06365 | 13 | type I-E CRISPR-associated protein Cas5/CasD [Azoarcus communis] |
| 03628 | 13 | MULTISPECIES: DUF2958 domain-containing protein [Proteobacteria] |
| 03767 | 13 | MULTISPECIES: haloacid dehalogenase type II [Azoarcus] |
| 05902 | 13 | archease [Aromatoleum bremense] |
| 04861 | 13 | MULTISPECIES: sterol desaturase family protein [Thauera] |
| 04949 | 13 | endonuclease/exonuclease/phosphatase family protein [Aromatoleum toluclasticum] |
| 07935 | 13 | hypothetical protein [Aromatoleum petrolei] |
| 06569 | 13 | anti-sigma factor [Nitrogeniibacter aestuarii] |
| 06025 | 13 | MOSC N-terminal beta barrel domain-containing protein [Nitrogeniibacter aestuarii] |
| 03785 | 13 | pyridoxamine 5'-phosphate oxidase family protein [Aromatoleum toluclasticum] |
| 06873 | 13 | hypothetical protein [Azoarcus communis] |
| 07354 | 13 | high-potential iron-sulfur protein [Aromatoleum toluclasticum] |
| 07971 | 13 | transporter substrate-binding domain-containing protein [Niveibacterium umoris] |
| 05336 | 13 | zinc ribbon domain-containing protein [Aromatoleum toluclasticum] |
| 06429 | 13 | class I SAM-dependent methyltransferase [Azoarcus olearius] |
| 07339 | 13 | hypothetical protein [Denitromonas halophilus] |
| 04894 | 13 | metal ABC transporter permease [Denitromonas halophilus] |
| 05973 | 13 | sugar kinase [Candidatus Dactylopiibacterium carminicum] |
| 05430 | 13 | DUF6351 family protein [Aromatoleum toluclasticum] |
| 05337 | 13 | ABC transporter substrate-binding protein [Niveibacterium sp. COAC-50] |
| 07423 | 13 | MULTISPECIES: DUF2914 domain-containing protein [Thauera] |
| 04764 | 13 | PEP-CTERM sorting domain-containing protein [Denitromonas halophilus] |
| 06230 | 13 | hypothetical protein [Aromatoleum toluclasticum] |
| 06952 | 13 | tetratricopeptide repeat protein [Denitromonas halophilus] |
| 03996 | 13 | DUF4125 family protein [Aromatoleum toluclasticum] |
| 06391 | 13 | alanyl-tRNA editing protein [Denitromonas halophilus] |
| 07544 | 13 | AHH domain-containing protein [Azoarcus communis] |
| 06772 | 13 | DsbA family protein [Azoarcus indigens] |
| 04459 | 13 | Rieske 2Fe-2S domain-containing protein [Azoarcus indigens] |
| 07060 | 13 | hypothetical protein [Denitromonas halophilus] |
| 07064 | 13 | type I-E CRISPR-associated protein Cas7/Cse4/CasC [Azoarcus communis] |

| | | |
|---|---|---|
| 05616 | 13 | winged helix-turn-helix domain-containing protein [Denitromonas halophilus] |
| 05639 | 13 | nucleotide 5'-monophosphate nucleosidase PpnN [Aromatoleum toluclasticum] |
| 04029 | 13 | pectinesterase family protein [Uliginosibacterium aquaticum] |
| 03554 | 13 | zinc ribbon domain-containing protein [Nitrogeniibacter aestuarii] |
| 07511 | 13 | type I-C CRISPR-associated protein Cas5c [Azoarcus indigens] |
| 07033 | 13 | RecQ family ATP-dependent DNA helicase [Aromatoleum toluclasticum] |
| 03678 | 13 | hypothetical protein [Aromatoleum toluclasticum] |
| 07491 | 13 | methyltransferase domain-containing protein [Aromatoleum toluclasticum] |
| 04649 | 13 | DUF4037 domain-containing protein [Aromatoleum toluclasticum] |
| 05561 | 13 | 3'-5' exonuclease [Azoarcus communis] |
| 07481 | 13 | DUF262 domain-containing HNH endonuclease family protein [Azoarcus indigens] |
| 05051 | 13 | IS110 family transposase [Aromatoleum toluclasticum] |
| 04432 | 13 | DUF2846 domain-containing protein [Uliginosibacterium aquaticum] |
| 07752 | 13 | MULTISPECIES: hypothetical protein [Niveibacterium] |
| 06354 | 13 | RNA-binding protein [Denitromonas halophilus] |
| 05526 | 13 | 5-methyltetrahydropteroyltriglutamate--homocysteine S-methyltransferase [Azoarcus indigens] |
| 06169 | 13 | DUF3291 domain-containing protein [Denitromonas halophilus] |
| 05007 | 13 | TetR family transcriptional regulator [Azoarcus indigens] |
| 05703 | 13 | PEP-CTERM sorting domain-containing protein [Azoarcus communis] |
| 06716 | 13 | FAD-dependent oxidoreductase [Denitromonas halophilus] |
| 06710 | 13 | diguanylate cyclase [Niveibacterium umoris] |
| 04151 | 13 | MULTISPECIES: glycosyltransferase family 2 protein [Niveibacterium] |
| 04260 | 13 | DNA recombination protein RmuC [Aromatoleum toluclasticum] |
| 06268 | 13 | hypothetical protein [Aromatoleum toluclasticum] |
| 01498 | 13 | hypothetical protein [Denitromonas halophilus] |
| 02327 | 13 | hypothetical protein [Aromatoleum toluclasticum] |
| 09605 | 13 | hypothetical protein [Aromatoleum toluclasticum] |
| 08696 | 13 | class I SAM-dependent methyltransferase [Azoarcus indigens] |
| 01931 | 13 | 1,4-dihydroxy-2-naphthoate polyprenyltransferase [Denitromonas halophilus] |
| 00467 | 13 | hypothetical protein [Azoarcus communis] |
| 10345 | 13 | DUF4224 domain-containing protein [Aromatoleum evansii] |
| 03430 | 13 | hypothetical protein [Aromatoleum toluclasticum] |
| 08653 | 13 | 50S ribosome-binding protein YggL [Denitromonas halophilus] |
| 01366 | 13 | spondin domain-containing protein [Denitromonas halophilus] |
| 02400 | 13 | hemerythrin domain-containing protein [Aromatoleum aromaticum] |
| 00399 | 13 | alkaline phosphatase D family protein [Nitrogeniibacter aestuarii] |
| 00775 | 13 | redoxin domain-containing protein [Niveibacterium sp. COAC-50] |
| 08631 | 13 | hypothetical protein [Denitromonas halophilus] |
| 03439 | 13 | polysaccharide deacetylase [Azoarcus indigens] |
| 08630 | 13 | NAD(P)-binding protein [Niveibacterium umoris] |
| 02915 | 13 | CRISPR-associated helicase Cas3' [Rhodocyclus purpureus] |
| 03250 | 13 | VWA domain-containing protein [Denitromonas halophilus] |
| 08735 | 13 | choice-of-anchor I family protein [Azoarcus indigens] |
| 10310 | 13 | DUF4123 domain-containing protein [Denitromonas halophilus] |
| 00451 | 13 | HAD-IA family hydrolase [Aromatoleum toluclasticum] |
| 08839 | 13 | phage portal protein [Thauera butanivorans] |
| 00700 | 13 | ABC transporter substrate-binding protein [Aromatoleum pumilum] |
| 08899 | 13 | YfjI family protein [Aromatoleum toluolicum] |
| 01813 | 13 | glycine zipper domain-containing protein [Azoarcus olearius] |
| 08897 | 13 | MULTISPECIES: autotransporter domain-containing protein [Niveibacterium] |
| 08882 | 13 | hypothetical protein [Denitromonas halophilus] |
| 10449 | 13 | ABC transporter substrate-binding protein [Azoarcus communis] |
| 09424 | 13 | hypothetical protein [Thauera butanivorans] |
| 08836 | 13 | GDSL-type esterase/lipase family protein [Denitromonas halophilus] |
| 08748 | 13 | FAD-dependent oxidoreductase [Azoarcus communis] |
| 02641 | 13 | isopenicillin N synthase family oxygenase [Azoarcus communis] |
| 02646 | 13 | type II toxin-antitoxin system VapC family toxin [Azoarcus communis] |
| 03411 | 13 | AAA family ATPase [Denitromonas halophilus] |
| 01605 | 13 | hypothetical protein [Aromatoleum toluclasticum] |
| 08776 | 13 | hypothetical protein [Azoarcus communis] |
| 08223 | 13 | co-chaperone YbbN [Azoarcus indigens] |
| 00519 | 13 | BREX system Lon protease-like protein BrxL [Zoogloea oleivorans] |
| 02940 | 13 | DNA/RNA nuclease SfsA [Azoarcus indigens] |
| 10612 | 13 | SRPBCC family protein [Aromatoleum toluclasticum] |
| 09756 | 13 | MULTISPECIES: CopG family transcriptional regulator [Aromatoleum] |
| 08555 | 13 | class D beta-lactamase [Nitrogeniibacter aestuarii] |
| 00786 | 13 | hypothetical protein [Aromatoleum toluclasticum] |
| 01082 | 13 | VOC family protein [Denitromonas halophilus] |
| 09226 | 13 | TolC family protein [Azoarcus olearius] |
| 00238 | 13 | transporter [Aromatoleum toluclasticum] |
| 10405 | 13 | O-antigen ligase family protein [Azoarcus communis] |
| 08461 | 13 | MULTISPECIES: acetamidase/formamidase family protein [Niveibacterium] |
| 10021 | 13 | hypothetical protein [Azoarcus communis] |
| 02094 | 13 | siderophore-interacting protein [Azoarcus indigens] |
| 10554 | 13 | RcnB family protein [Azoarcus indigens] |
| 08415 | 13 | four-helix bundle copper-binding protein [Azoarcus olearius] |
| 00250 | 13 | cytochrome C peroxidase [Aromatoleum toluclasticum] |
| 03375 | 13 | hypothetical protein [Azoarcus communis] |
| 08410 | 13 | SRPBCC family protein [Aromatoleum toluclasticum] |
| 09161 | 13 | hypothetical protein [Aromatoleum evansii] |
| 09970 | 13 | hypothetical protein [Aromatoleum toluclasticum] |
| 10242 | 13 | condensation domain-containing protein, partial [Denitromonas sp. IR12] |
| 00908 | 13 | DegV family protein [Denitromonas halophilus] |
| 03012 | 13 | PIN domain nuclease [Aromatoleum bremense] |
| 09439 | 13 | type I-E CRISPR-associated endonuclease Cas1e [Azoarcus communis] |
| 00804 | 13 | Poxvirus G6 [Azoarcus indigens] |
| 02006 | 13 | class I SAM-dependent methyltransferase [Azoarcus indigens] |
| 10082 | 13 | Crp/Fnr family transcriptional regulator [Aromatoleum toluclasticum] |
| 00216 | 13 | MULTISPECIES: mercury resistance system transport protein MerF [Proteobacteria] |
| 01134 | 13 | GTPase domain-containing protein [Aromatoleum petrolei] |
| 00611 | 13 | BREX system P-loop protein BrxC [Zoogloea oleivorans] |
| 09768 | 13 | nitrate/nitrite transporter NrtS [Aromatoleum toluclasticum] |
| 02264 | 13 | MULTISPECIES: hypothetical protein [Niveibacterium] |
| 10075 | 13 | MULTISPECIES: OB-fold domain-containing protein [Aromatoleum] |

| | | |
|---|---|---|
| 02243 | 13 | ParD-like family protein [Aromatoleum toluclasticum] |
| 01189 | 13 | hypothetical protein [Aromatoleum toluclasticum] |
| 01192 | 13 | MULTISPECIES: plasmid pRiA4b ORF-3 family protein [Rhodocyclales] |
| 06692 | 12 | MULTISPECIES: hypothetical protein [Proteobacteria] |
| 04385 | 12 | HIRAN domain-containing protein [Azospira oryzae] |
| 04238 | 12 | SGNH/GDSL hydrolase family protein [Azoarcus indigens] |
| 05785 | 12 | BrnT family toxin [Aromatoleum pumilum] |
| 00117 | 12 | MULTISPECIES: DUF2285 domain-containing protein [Proteobacteria] |
| 02305 | 12 | AAA family ATPase [Azoarcus sp. DD4] |
| 02467 | 12 | CHRD domain-containing protein [Denitromonas halophilus] |
| 07593 | 12 | helix-turn-helix domain-containing protein [Azoarcus indigens] |
| 02338 | 12 | cysteine dioxygenase [Azoarcus indigens] |
| 04463 | 12 | DUF883 family protein [Azoarcus indigens] |
| 09153 | 12 | phage tail protein [Denitromonas halophilus] |
| 02197 | 12 | hypothetical protein [Aromatoleum toluclasticum] |
| 06685 | 12 | alpha/beta hydrolase [Azoarcus indigens] |
| 04438 | 12 | MULTISPECIES: hypothetical protein [Zoogloeaceae] |
| 04321 | 12 | response regulator [Aromatoleum toluclasticum] |
| 09003 | 12 | PEP-CTERM sorting domain-containing protein [Azoarcus communis] |
| 05967 | 12 | DUF5666 domain-containing protein [Aromatoleum toluclasticum] |
| 10315 | 12 | MULTISPECIES: metal/formaldehyde-sensitive transcriptional repressor [Thauera] |
| 02966 | 12 | PEP-CTERM sorting domain-containing protein [Denitromonas halophilus] |
| 03902 | 12 | hypothetical protein [Azoarcus indigens] |
| 06224 | 12 | hypothetical protein [Aromatoleum petrolei] |
| 05866 | 12 | hypothetical protein [Aromatoleum bremense] |
| 08569 | 12 | PD-(D/E)XK nuclease family protein [Denitromonas halophilus] |
| 08530 | 12 | CRISPR-associated endonuclease Cas2 [Niveibacterium sp. COAC-50] |
| 08500 | 12 | helix-turn-helix domain-containing protein [Rhodocyclus purpureus] |
| 03091 | 12 | hypothetical protein [Aromatoleum toluclasticum] |
| 00429 | 12 | MULTISPECIES: MarR family winged helix-turn-helix transcriptional regulator [Thauera] |
| 06572 | 12 | hypothetical protein [Azoarcus olearius] |
| 03789 | 12 | MULTISPECIES: type II toxin-antitoxin system ParD family antitoxin [Thauera] |
| 03770 | 12 | DUF4157 domain-containing protein [Aromatoleum toluclasticum] |
| 06235 | 12 | hypothetical protein [Azoarcus communis] |
| 03192 | 12 | DUF1972 domain-containing protein [Denitromonas halophilus] |
| 08408 | 12 | MULTISPECIES: bifunctional acetaldehyde-CoA/alcohol dehydrogenase [Niveibacterium] |
| 06552 | 12 | type VI secretion system-associated FHA domain protein TagH [Azoarcus communis] |
| 03904 | 12 | hypothetical protein [Aromatoleum diolicum] |
| 02939 | 12 | c-type cytochrome [Azoarcus indigens] |
| 04219 | 12 | 3-dehydro-L-gulonate 2-dehydrogenase [Propionivibrio dicarboxylicus] |
| 06213 | 12 | DUF2118 domain-containing protein [Zoogloea ramigera] |
| 02545 | 12 | hypothetical protein [Aromatoleum diolicum] |
| 04169 | 12 | flavin reductase family protein [Aromatoleum toluclasticum] |
| 02634 | 12 | SRPBCC family protein [Aromatoleum toluclasticum] |
| 08811 | 12 | DNA-3-methyladenine glycosylase 2 [Denitromonas halophilus] |
| 02692 | 12 | class GN sortase [Denitromonas halophilus] |
| 06494 | 12 | YegP family protein [Aromatoleum evansii] |
| 04070 | 12 | hypothetical protein [Denitromonas halophilus] |
| 05826 | 12 | peptidoglycan-binding protein [Aromatoleum toluclasticum] |
| 06630 | 12 | hypothetical protein [Aromatoleum toluclasticum] |
| 04482 | 12 | 4-alpha-glucanotransferase [Azoarcus taiwanensis] |
| 08690 | 12 | type IV pilin protein [Aromatoleum toluclasticum] |
| 02814 | 12 | IS3 family transposase, partial [Aromatoleum toluclasticum] |
| 03993 | 12 | MULTISPECIES: amidohydrolase [Rhodocyclales] |
| 02886 | 12 | MULTISPECIES: mechanosensitive ion channel family protein [Thauera] |
| 03964 | 12 | triphosphoribosyl-dephospho-CoA synthase MdcB [Nitrogeniibacter aestuarii] |
| 06151 | 12 | GTP cyclohydrolase II [Azoarcus communis] |
| 06133 | 12 | 2-phosphosulfolactate phosphatase [Aromatoleum toluclasticum] |
| 02145 | 12 | MULTISPECIES: hypothetical protein [Rhodocyclales] |
| 00120 | 12 | type VI secretion system tip protein VgrG, partial [Azoarcus indigens] |
| 07034 | 12 | flippase [Aromatoleum petrolei] |
| 01252 | 12 | hypothetical protein [Denitromonas halophilus] |
| 05160 | 12 | hypothetical protein [Azoarcus olearius] |
| 05155 | 12 | URC4/urg3 family protein [Azoarcus communis] |
| 01277 | 12 | BLUF domain-containing protein [Nitrogeniibacter aestuarii] |
| 06458 | 12 | VanZ family protein [Azoarcus indigens] |
| 01287 | 12 | hypothetical protein [Aromatoleum toluclasticum] |
| 01318 | 12 | MULTISPECIES: hypothetical protein [Thauera] |
| 07079 | 12 | hypothetical protein [Denitromonas halophilus] |
| 01355 | 12 | MULTISPECIES: DUF2274 domain-containing protein [Betaproteobacteria] |
| 06414 | 12 | periplasmic heavy metal sensor [Denitromonas halophilus] |
| 05584 | 12 | MULTISPECIES: hypothetical protein [Rhodocyclales] |
| 07121 | 12 | MULTISPECIES: DUF2116 family Zn-ribbon domain-containing protein [Niveibacterium] |
| 07129 | 12 | MULTISPECIES: TolC family protein [Zoogloeaceae] |
| 01438 | 12 | potassium-transporting ATPase subunit KdpC [Azoarcus indigens] |
| 10108 | 12 | glycerate kinase [Azoarcus communis] |
| 01442 | 12 | CoA-binding protein [Denitromonas halophilus] |
| 01462 | 12 | LuxR family transcriptional regulator [Nitrogeniibacter aestuarii] |
| 04992 | 12 | Fic family protein [Aromatoleum toluclasticum] |
| 01227 | 12 | hypothetical protein [Azoarcus communis] |
| 01222 | 12 | DJ-1/PfpI family protein [Azoarcus indigens] |
| 05199 | 12 | MarR family winged helix-turn-helix transcriptional regulator [Denitromonas halophilus] |
| 05381 | 12 | hypothetical protein [Azoarcus halotolerans] |
| 00921 | 12 | hypothetical protein [Denitromonas halophilus] |
| 00254 | 12 | MULTISPECIES: type II toxin-antitoxin system Phd/YefM family antitoxin [Azoarcus] |
| 00964 | 12 | helix-turn-helix domain-containing protein [Aromatoleum pumilum] |
| 10485 | 12 | OB-fold domain-containing protein [Aromatoleum toluolicum] |
| 09920 | 12 | DUF1932 domain-containing protein [Nitrogeniibacter aestuarii] |
| 05504 | 12 | HD domain-containing protein [Zoogloea dura] |
| 05403 | 12 | retention module-containing protein, partial [Aromatoleum toluclasticum] |
| 06896 | 12 | DUF262 domain-containing protein [Aromatoleum tolulyticum] |
| 09888 | 12 | phage virion morphogenesis protein [Thauera phenylacetica] |
| 10465 | 12 | hypothetical protein [Denitromonas halophilus] |
| 01043 | 12 | IS4 family transposase [Aromatoleum bremense] |

| | | |
|---|---|---|
| 01050 | 12 | exopolysaccharide biosynthesis protein [Azoarcus communis] |
| 05345 | 12 | hypothetical protein [Azoarcus communis] |
| 00849 | 12 | TonB-dependent receptor [Azoarcus indigens] |
| 00843 | 12 | MULTISPECIES: relaxase/mobilization nuclease and DUF3363 domain-containing protein [Betaproteobacteria] |
| 05305 | 12 | F0F1 ATP synthase subunit C [Denitromonas halophilus] |
| 00829 | 12 | GNAT family N-acetyltransferase, partial [Azoarcus halotolerans] |
| 05547 | 12 | trimethylamine methyltransferase family protein [Aromatoleum toluvorans] |
| 01509 | 12 | F0F1 ATP synthase subunit B [Denitromonas halophilus] |
| 07173 | 12 | PEP-CTERM sorting domain-containing protein [Denitromonas halophilus] |
| 04975 | 12 | PD40 domain-containing protein [Aromatoleum evansii] |
| 02001 | 12 | DUF433 domain-containing protein [Aromatoleum aromaticum] |
| 04684 | 12 | hypothetical protein [Azoarcus olearius] |
| 07397 | 12 | HNH endonuclease [Aromatoleum toluvorans] |
| 04680 | 12 | GDYXXLXY domain-containing protein [Aromatoleum toluclasticum] |
| 00142 | 12 | magnesium chelatase [Azoarcus communis] |
| 01928 | 12 | type II toxin-antitoxin system Phd/YefM family antitoxin [Rhodocyclus purpureus] |
| 09316 | 12 | acetate uptake transporter [Nitrogeniibacter aestuarii] |
| 04634 | 12 | TRAFs-binding domain-containing protein [Azoarcus olearius] |
| 04623 | 12 | hypothetical protein [Azoarcus communis] |
| 09247 | 12 | quinoprotein dehydrogenase-associated putative ABC transporter substrate-binding protein [Nitrogeniibacter aestuarii] |
| 01870 | 12 | recombinase family protein [Aromatoleum toluclasticum] |
| 02034 | 12 | toprim domain-containing protein [Aromatoleum aromaticum] |
| 05702 | 12 | OB-fold domain-containing protein [Aromatoleum toluclasticum] |
| 02067 | 12 | DUF1499 domain-containing protein [Aromatoleum toluclasticum] |
| 04529 | 12 | DUF2818 family protein [Azoarcus nasutitermitis] |
| 04528 | 12 | NUDIX hydrolase [Aromatoleum toluclasticum] |
| 02090 | 12 | PepSY domain-containing protein [Thauera phenylacetica] |
| 02103 | 12 | type II toxin-antitoxin system VapC family toxin [Azoarcus rhizosphaerae] |
| 04508 | 12 | alkaline phosphatase D family protein [Aromatoleum toluclasticum] |
| 05670 | 12 | D-xylulose 5-phosphate/D-fructose 6-phosphate phosphoketolase [Thauera terpenica] |
| 04732 | 12 | Nif11-like leader peptide family natural product precursor [Azoarcus communis] |
| 10529 | 12 | ethylbenzene dehydrogenase-related protein [Aromatoleum aromaticum] |
| 01652 | 12 | hypothetical protein [Aromatoleum evansii] |
| 07209 | 12 | AAA family ATPase [Azoarcus communis] |
| 05607 | 12 | hypothetical protein [Denitromonas halophilus] |
| 01603 | 12 | MULTISPECIES: DUF2840 domain-containing protein [Proteobacteria] |
| 00183 | 12 | AtpZ/AtpI family protein [Denitromonas halophilus] |
| 04909 | 12 | MULTISPECIES: VOC family protein [Niveibacterium] |
| 01630 | 12 | OB-fold domain-containing protein [Aromatoleum toluclasticum] |
| 07236 | 12 | two-component system response regulator [Azoarcus indigens] |
| 10133 | 12 | hypothetical protein [Azoarcus communis] |
| 01655 | 12 | MULTISPECIES: hypothetical protein [Bacteria] |
| 09399 | 12 | L(+)-tartrate dehydratase subunit beta [Denitromonas halophilus] |
| 01663 | 12 | sodium:solute symporter [Denitromonas halophilus] |
| 04870 | 12 | hypothetical protein [Aromatoleum toluclasticum] |
| 00166 | 12 | hypothetical protein [Azoarcus communis] |
| 01734 | 12 | MULTISPECIES: lmo0937 family membrane protein [Niveibacterium] |
| 01740 | 12 | ABC transporter permease [Azoarcus communis] |
| 09463 | 12 | ABC transporter substrate-binding protein [Azoarcus communis] |
| 00670 | 12 | helix-turn-helix domain-containing protein [Aromatoleum toluolicum] |
| 07331 | 12 | cytochrome c [Nitrogeniibacter aestuarii] |
| 07967 | 12 | pectate lyase [Uliginosibacterium aquaticum] |
| 10556 | 12 | DUF3240 family protein [Azoarcus olearius] |
| 10391 | 12 | prolyl oligopeptidase family serine peptidase [Niveibacterium microcysteis] |
| 10436 | 12 | hypothetical protein [Aromatoleum aromaticum] |
| 03275 | 12 | MULTISPECIES: type I 3-dehydroquinate dehydratase [Aromatoleum] |
| 03595 | 12 | hypothetical protein [Aromatoleum toluclasticum] |
| 00347 | 12 | hypothetical protein [Azoarcus communis] |
| 03545 | 12 | MULTISPECIES: cytochrome c biogenesis protein ResB [Thauera] |
| 03477 | 12 | hypothetical protein [Aromatoleum toluolicum] |
| 00324 | 12 | phage terminase large subunit family protein [Candidatus Dactylopiibacterium carminicum] |
| 03575 | 12 | hypothetical protein [Denitromonas halophilus] |
| 03468 | 12 | acetyltransferase [Azoarcus sp. KH32C] |
| 05912 | 12 | GDP-mannose--glycolipid 4-beta-D-mannosyltransferase [Azoarcus indigens] |
| 05256 | 11 | GT4 family glycosyltransferase PelF [Denitromonas halophilus] |
| 05300 | 11 | MULTISPECIES: hypothetical protein [Aromatoleum] |
| 07570 | 11 | O-antigen ligase family protein [Aromatoleum toluclasticum] |
| 02729 | 11 | DnaJ domain-containing protein [Aromatoleum aromaticum] |
| 01419 | 11 | MULTISPECIES: DUF736 domain-containing protein [Proteobacteria] |
| 05313 | 11 | helix-turn-helix domain-containing protein [Denitromonas halophilus] |
| 01993 | 11 | DUF3262 family protein [Aromatoleum aromaticum] |
| 03978 | 11 | AAA family ATPase [Thauera lacus] |
| 08658 | 11 | MULTISPECIES: DUF3262 family protein [Aromatoleum] |
| 05529 | 11 | hypothetical protein [Azoarcus taiwanensis] |
| 01703 | 11 | hypothetical protein [Denitromonas halophilus] |
| 01696 | 11 | MULTISPECIES: VirB3 family type IV secretion system protein [Proteobacteria] |
| 04009 | 11 | MULTISPECIES: hypothetical protein [Aromatoleum] |
| 04014 | 11 | hypothetical protein [Azoarcus communis] |
| 00489 | 11 | MULTISPECIES: TIGR03752 family integrating conjugative element protein [Aromatoleum] |
| 10054 | 11 | MULTISPECIES: hypothetical protein [Niveibacterium] |
| 09523 | 11 | TnsA endonuclease N-terminal domain-containing protein [Aromatoleum toluclasticum] |
| 10584 | 11 | hypothetical protein [Aromatoleum toluclasticum] |
| 05286 | 11 | carboxypeptidase M32 [Azoarcus olearius] |
| 00163 | 11 | saccharopine dehydrogenase NADP-binding domain-containing protein [Denitromonas halophilus] |
| 01166 | 11 | MULTISPECIES: conjugal transfer protein TraF [Betaproteobacteria] |
| 05544 | 11 | DUF421 domain-containing protein [Azoarcus indigens] |
| 04047 | 11 | hypothetical protein [Aromatoleum toluclasticum] |
| 04050 | 11 | electron transfer flavoprotein subunit beta [Aromatoleum evansii] |
| 10585 | 11 | nuclear transport factor 2 family protein [Azoarcus indigens] |
| 09806 | 11 | helix-turn-helix domain-containing protein [Azoarcus nasutitermitis] |
| 09023 | 11 | MCP four helix bundle domain-containing protein [Aromatoleum toluclasticum] |
| 07459 | 11 | FAD-dependent oxidoreductase [Aromatoleum pumilum] |
| 10273 | 11 | hypothetical protein [Aromatoleum toluclasticum] |
| 05952 | 11 | TonB-dependent receptor [Azoarcus olearius] |

| | | |
|---|---|---|
| 09378 | 11 | MULTISPECIES: type IV conjugative transfer system protein TraL [Betaproteobacteria] |
| 00145 | 11 | PqiC family protein [Azoarcus halotolerans] |
| 00654 | 11 | hypothetical protein [Denitromonas halophilus] |
| 03038 | 11 | hypothetical protein [Uliginosibacterium aquaticum] |
| 00302 | 11 | mechanosensitive ion channel family protein [Azoarcus indigens] |
| 10196 | 11 | DsrE family protein [Rhodocyclus purpureus] |
| 00701 | 11 | hypothetical protein [Aromatoleum toluclasticum] |
| 08568 | 11 | Ig-like domain-containing protein [Nitrogeniibacter aestuarii] |
| 03891 | 11 | PIN domain-containing protein [Aromatoleum diolicum] |
| 02983 | 11 | cytochrome b5 domain-containing protein [Nitrogeniibacter aestuarii] |
| 01757 | 11 | iron-sulfur cluster assembly scaffold protein [Aromatoleum toluclasticum] |
| 02978 | 11 | nitrate- and nitrite sensing domain-containing protein [Azoarcus communis] |
| 02973 | 11 | SDR family oxidoreductase [Denitromonas halophilus] |
| 02972 | 11 | putative baseplate assembly protein [Denitromonas halophilus] |
| 02952 | 11 | DUF3016 domain-containing protein [Aromatoleum toluclasticum] |
| 03418 | 11 | helix-hairpin-helix domain-containing protein [Azoarcus communis] |
| 01905 | 11 | hypothetical protein [Azoarcus indigens] |
| 06441 | 11 | serine hydrolase [Azoarcus communis] |
| 05674 | 11 | diguanylate cyclase [Aromatoleum toluclasticum] |
| 05047 | 11 | glycosyl hydrolase 53 family protein [Niveibacterium umoris] |
| 08079 | 11 | MULTISPECIES: periplasmic nitrate reductase, NapE protein [Azoarcus] |
| 01420 | 11 | type II toxin-antitoxin system Phd/YefM family antitoxin [Azoarcus communis] |
| 03937 | 11 | hypothetical protein [Denitromonas halophilus] |
| 09470 | 11 | hypothetical protein [Rhodocyclus purpureus] |
| 00462 | 11 | MULTISPECIES: type-F conjugative transfer system protein TraW [Betaproteobacteria] |
| 05334 | 11 | Tad domain-containing protein [Aromatoleum tolulyticum] |
| 06921 | 11 | hypothetical protein [Aromatoleum toluclasticum] |
| 02907 | 11 | hypothetical protein [Azoarcus communis] |
| 03961 | 11 | hypothetical protein [Aromatoleum toluclasticum] |
| 03962 | 11 | DUF3379 family protein [Denitromonas halophilus] |
| 01995 | 11 | MULTISPECIES: type-F conjugative transfer system pilin assembly protein TrbC [Betaproteobacteria] |
| 00622 | 11 | DUF6491 family protein [Aromatoleum toluclasticum] |
| 07461 | 11 | toll/interleukin-1 receptor domain-containing protein [Denitromonas halophilus] |
| 04222 | 11 | potassium-transporting ATPase subunit KdpA [Azoarcus indigens] |
| 05566 | 11 | hypothetical protein [Uliginosibacterium gangwonense] |
| 02506 | 11 | hypothetical protein [Aromatoleum tolulyticum] |
| 03447 | 11 | glutathione S-transferase [Denitromonas halophilus] |
| 02079 | 11 | RHS domain-containing protein [Azoarcus communis] |
| 04536 | 11 | hypothetical protein [Aromatoleum toluclasticum] |
| 08276 | 11 | GDSL-type esterase/lipase family protein [Aromatoleum toluclasticum] |
| 07695 | 11 | hypothetical protein [Nitrogeniibacter aestuarii] |
| 02479 | 11 | Hpt domain-containing protein [Azoarcus communis] |
| 02478 | 11 | LysM domain-containing protein [Denitromonas halophilus] |
| 05709 | 11 | phage baseplate assembly protein V [Denitromonas halophilus] |
| 02473 | 11 | hypothetical protein [Aromatoleum aromaticum] |
| 04272 | 11 | DinB family protein [Denitromonas halophilus] |
| 00520 | 11 | TrkA family potassium uptake protein [Aromatoleum aromaticum] |
| 05130 | 11 | DUF2341 domain-containing protein [Denitromonas halophilus] |
| 10300 | 11 | Ku protein [Aromatoleum diolicum] |
| 09589 | 11 | general secretion pathway protein GspB [Aromatoleum toluclasticum] |
| 09716 | 11 | diguanylate cyclase [Azoarcus communis] |
| 00599 | 11 | PEP-CTERM sorting domain-containing protein [Niveibacterium sp. COAC-50] |
| 10091 | 11 | DUF2164 domain-containing protein [Azoarcus indigens] |
| 01503 | 11 | MULTISPECIES: transcriptional regulator [Proteobacteria] |
| 06463 | 11 | hypothetical protein [Aromatoleum toluclasticum] |
| 05583 | 11 | hypothetical protein [Nitrogeniibacter aestuarii] |
| 09684 | 11 | sulfotransferase [Thauera terpenica] |
| 01501 | 11 | hypothetical protein [Aromatoleum toluclasticum] |
| 02359 | 11 | MULTISPECIES: S26 family signal peptidase [Betaproteobacteria] |
| 07620 | 11 | lactoylglutathione lyase [Denitromonas halophilus] |
| 07097 | 11 | hypothetical protein [Denitromonas halophilus] |
| 08275 | 11 | acyl CoA--acetate/3-ketoacid CoA transferase subunit beta [Aromatoleum toluclasticum] |
| 09616 | 11 | acyl-protein synthetase [Aromatoleum diolicum] |
| 09759 | 11 | N-6 DNA methylase [Aromatoleum toluolicum] |
| 05177 | 11 | helix-turn-helix domain-containing protein [Denitromonas halophilus] |
| 00807 | 11 | MULTISPECIES: TraU family protein [Betaproteobacteria] |
| 05924 | 11 | DUF1223 domain-containing protein [Denitromonas halophilus] |
| 09912 | 11 | MULTISPECIES: ribbon-helix-helix protein, CopG family [Betaproteobacteria] |
| 00116 | 11 | energy transducer TonB [Denitromonas halophilus] |
| 02293 | 11 | MULTISPECIES: (2Fe-2S)-binding protein [Azoarcus] |
| 04090 | 11 | M67 family metallopeptidase [Aromatoleum toluclasticum] |
| 10228 | 11 | wax ester/triacylglycerol synthase family O-acyltransferase [Niveibacterium umoris] |
| 04094 | 11 | hypothetical protein [Aromatoleum toluclasticum] |
| 00727 | 11 | hypothetical protein [Rhodocyclus purpureus] |
| 02707 | 11 | PEP-CTERM sorting domain-containing protein [Azoarcus olearius] |
| 01193 | 11 | hypothetical protein [Aromatoleum toluclasticum] |
| 01200 | 11 | MULTISPECIES: hypothetical protein [Rhodocyclales] |
| 05218 | 11 | MULTISPECIES: OmpA family protein [Betaproteobacteria] |
| 00025 | 11 | mechanosensitive ion channel [Azoarcus indigens] |
| 01590 | 11 | MULTISPECIES: conjugal transfer protein TraH [Betaproteobacteria] |
| 06406 | 11 | malonate decarboxylase acyl carrier protein [Aromatoleum pumilum] |
| 05809 | 11 | ATP-binding cassette domain-containing protein, partial [Aromatoleum toluclasticum] |
| 09639 | 11 | DUF6516 family protein [Aromatoleum bremense] |
| 02306 | 11 | malonate decarboxylase subunit alpha [Aromatoleum pumilum] |
| 01564 | 11 | DUF3304 domain-containing protein [Aromatoleum toluclasticum] |
| 03315 | 11 | SHOCT domain-containing protein [Niveibacterium microcysteis] |
| 01230 | 11 | hypothetical protein [Denitromonas halophilus] |
| 10232 | 11 | putative baseplate assembly protein [Denitromonas halophilus] |
| 00504 | 11 | type II toxin-antitoxin system RelE/ParE family toxin [Thauera phenylacetica] |
| 00505 | 11 | VTT domain-containing protein [Azoarcus indigens] |
| 04182 | 11 | hypothetical protein [Denitromonas halophilus] |
| 07731 | 11 | energy-coupling factor transporter transmembrane protein EcfT [Azoarcus communis] |
| 01235 | 11 | GrpB family protein [Azoarcus taiwanensis] |
| 04189 | 11 | MULTISPECIES: conjugal transfer protein TraN [Betaproteobacteria] |

| | | |
|---|---|---|
| 00373 | 11 | addiction module antitoxin [Denitromonas halophilus] |
| 04195 | 11 | YceI family protein [Denitromonas halophilus] |
| 01423 | 11 | MULTISPECIES: helix-turn-helix domain-containing protein [Proteobacteria] |
| 00119 | 11 | MULTISPECIES: class A beta-lactamase [Niveibacterium] |
| 03588 | 11 | nuclear transport factor 2 family protein [Aromatoleum diolicum] |
| 02244 | 11 | carboxypeptidase-like regulatory domain-containing protein [Azoarcus communis] |
| 06397 | 11 | MULTISPECIES: P-type conjugative transfer protein TrbL [Proteobacteria] |
| 05877 | 11 | pilus assembly protein [Aromatoleum aromaticum] |
| 09971 | 11 | FAD-binding oxidoreductase [Denitromonas halophilus] |
| 05501 | 11 | glutamate-ammonia-ligase adenylyltransferase [Denitromonas halophilus] |
| 01428 | 11 | MULTISPECIES: YHS domain-containing protein [Denitromonas] |
| 04440 | 11 | PEP-CTERM sorting domain-containing protein [Denitromonas halophilus] |
| 08373 | 11 | MULTISPECIES: RAQPRD family integrative conjugative element protein [Aromatoleum] |
| 08339 | 11 | chemotaxis protein CheW [Azoarcus indigens] |
| 08468 | 11 | tellurite resistance/C4-dicarboxylate transporter family protein [Denitromonas halophilus] |
| 03212 | 11 | hypothetical protein [Azoarcus communis] |
| 00657 | 11 | O-antigen ligase family protein [Denitromonas halophilus] |
| 04756 | 11 | DUF4255 domain-containing protein [Denitromonas halophilus] |
| 08421 | 11 | NAD(P)/FAD-dependent oxidoreductase [Azoarcus indigens] |
| 04733 | 11 | phage Gp37/Gp68 family protein [Aromatoleum pumilum] |
| 10407 | 11 | MULTISPECIES: hypothetical protein [Niveibacterium] |
| 03815 | 11 | DNA ligase D [Azoarcus olearius] |
| 06029 | 11 | biliverdin-producing heme oxygenase [Azoarcus communis] |
| 04426 | 11 | aspartate kinase [Denitromonas halophilus] |
| 01817 | 11 | ATP-binding protein [Aromatoleum tolulyticum] |
| 06296 | 11 | BtrH N-terminal domain-containing protein [Azoarcus communis] |
| 00661 | 11 | amidohydrolase family protein [Thauera phenylacetica] |
| 00978 | 11 | DUF72 domain-containing protein [Azoarcus indigens] |
| 05470 | 11 | MULTISPECIES: hypothetical protein [Thauera] |
| 03852 | 11 | zinc-binding alcohol dehydrogenase [Azoarcus communis] |
| 04758 | 11 | DUF3427 domain-containing protein [Azoarcus communis] |
| 09115 | 11 | aquaporin Z [Azoarcus communis] |
| 04739 | 11 | Arc family DNA-binding protein [Azoarcus indigens] |
| 00416 | 11 | YgiT-type zinc finger protein [Aromatoleum toluclasticum] |
| 00114 | 11 | alpha-ketoglutarate-dependent dioxygenase AlkB [Azoarcus communis] |
| 05439 | 11 | ankyrin repeat domain-containing protein [Denitromonas sp. IR12] |
| 03810 | 11 | hypothetical protein [Azoarcus communis] |
| 01787 | 11 | toprim domain-containing protein [Aromatoleum toluolicum] |
| 00967 | 11 | helix-turn-helix domain-containing protein [Uliginosibacterium gangwonense] |
| 05466 | 11 | TonB-dependent receptor [Thauera linaloolentis] |
| 05398 | 11 | MULTISPECIES: hypothetical protein [Denitromonas] |
| 04586 | 10 | MULTISPECIES: hypothetical protein [Aromatoleum] |
| 05213 | 10 | PEP-CTERM sorting domain-containing protein [Aromatoleum aromaticum] |
| 08845 | 10 | hypothetical protein [Propionivibrio dicarboxylicus] |
| 04166 | 10 | ABC-type transport auxiliary lipoprotein family protein [Aromatoleum toluclasticum] |
| 05020 | 10 | hypothetical protein [Denitromonas halophilus] |
| 10231 | 10 | DUF2924 domain-containing protein [Aromatoleum bremense] |
| 04557 | 10 | VOC family protein [Azoarcus communis] |
| 10383 | 10 | polysaccharide pyruvyl transferase family protein [Azoarcus indigens] |
| 01831 | 10 | CAAX prenyl protease-related protein [Azoarcus communis] |
| 03535 | 10 | DUF4198 domain-containing protein [Azoarcus communis] |
| 00934 | 10 | glutaminase [Denitromonas halophilus] |
| 10080 | 10 | hypothetical protein [Aromatoleum aromaticum] |
| 04945 | 10 | hypothetical protein [Nitrogeniibacter aestuarii] |
| 05457 | 10 | glycosyltransferase family 2 protein [Aromatoleum toluclasticum] |
| 06245 | 10 | VOC family protein [Denitromonas halophilus] |
| 05610 | 10 | MULTISPECIES: pyridoxal phosphatase [Niveibacterium] |
| 04581 | 10 | CYTH domain-containing protein [Aromatoleum evansii] |
| 02616 | 10 | AAA family ATPase [Aromatoleum diolicum] |
| 10331 | 10 | hypothetical protein [Azoarcus sp. L1K30] |
| 07008 | 10 | PRC-barrel domain-containing protein [Aromatoleum aromaticum] |
| 07483 | 10 | azurin [Azoarcus indigens] |
| 00385 | 10 | DUF2875 family protein [Azoarcus indigens] |
| 07764 | 10 | Mov34/MPN/PAD-1 family protein [Aromatoleum toluclasticum] |
| 06815 | 10 | hypothetical protein [Aromatoleum toluclasticum] |
| 09137 | 10 | YciI family protein [Denitromonas halophilus] |
| 00424 | 10 | putative porin [Azoarcus indigens] |
| 06260 | 10 | hypothetical protein [Propionivibrio dicarboxylicus] |
| 00135 | 10 | MULTISPECIES: NAD(P)H-binding protein [Rhodocyclales] |
| 04453 | 10 | MULTISPECIES: peptidoglycan-binding protein [Thauera] |
| 05938 | 10 | histidine phosphatase family protein [Aromatoleum toluclasticum] |
| 05624 | 10 | hypothetical protein [Aromatoleum aromaticum] |
| 00487 | 10 | isochorismatase family protein [Denitromonas halophilus] |
| 07241 | 10 | hypothetical protein [Aromatoleum toluclasticum] |
| 04631 | 10 | NrdH-redoxin [Rhodocyclus purpureus] |
| 00171 | 10 | transposase family protein, partial [Azoarcus communis] |
| 04636 | 10 | MBL fold metallo-hydrolase [Azoarcus halotolerans] |
| 01682 | 10 | urea carboxylase-associated family protein [Azoarcus communis] |
| 05886 | 10 | MULTISPECIES: hypothetical protein [Zoogloeaceae] |
| 04873 | 10 | DUF1640 domain-containing protein [Aromatoleum aromaticum] |
| 06134 | 10 | MliC family protein [Aromatoleum toluclasticum] |
| 09801 | 10 | hypothetical protein [Azoarcus olearius] |
| 08228 | 10 | molecular chaperone TorD family protein [Denitromonas halophilus] |
| 06986 | 10 | hypothetical protein [Aromatoleum toluclasticum] |
| 06992 | 10 | hypothetical protein [Aromatoleum toluclasticum] |
| 06988 | 10 | adenosine-specific kinase [Aromatoleum toluclasticum] |
| 05741 | 10 | hypothetical protein [Azoarcus sp. DD4] |
| 08431 | 10 | mechanosensitive channel MscK [Thauera aromatica] |
| 09265 | 10 | helix-turn-helix domain-containing protein [Zoogloea dura] |
| 00065 | 10 | hypothetical protein [Azoarcus indigens] |
| 01434 | 10 | hypothetical protein, partial [Aromatoleum toluclasticum] |
| 06978 | 10 | hypothetical protein [Aromatoleum toluclasticum] |
| 08767 | 10 | MULTISPECIES: P-type conjugative transfer protein TrbL [Proteobacteria] |
| 06501 | 10 | hypothetical protein [Azoarcus communis] |

| | | |
|---|---|---|
| 02723 | 10 | hypothetical protein [Aromatoleum toluclasticum] |
| 06863 | 10 | hypothetical protein [Aromatoleum aromaticum] |
| 01854 | 10 | hypothetical protein [Azoarcus indigens] |
| 08150 | 10 | MULTISPECIES: benzylsuccinate synthase gamma subunit family protein [Thauera] |
| 04174 | 10 | MULTISPECIES: hypothetical protein [Thauera] |
| 06226 | 10 | MULTISPECIES: hypothetical protein [Niveibacterium] |
| 00557 | 10 | alpha/beta fold hydrolase [Azoarcus indigens] |
| 06368 | 10 | bacteriohemerythrin [Denitromonas halophilus] |
| 05934 | 10 | class I SAM-dependent methyltransferase [Aromatoleum toluclasticum] |
| 00103 | 10 | PEP-CTERM sorting domain-containing protein [Denitromonas sp. IR12] |
| 01303 | 10 | hypothetical protein [Aromatoleum petrolei] |
| 07535 | 10 | hypothetical protein [Azoarcus olearius] |
| 06272 | 10 | hypothetical protein [Azoarcus olearius] |
| 03625 | 10 | ribbon-helix-helix protein, CopG family [Azoarcus communis] |
| 06261 | 10 | CbtA family protein [Aromatoleum toluclasticum] |
| 07965 | 10 | 5'-nucleotidase [Aromatoleum toluolicum] |
| 06411 | 10 | type I-E CRISPR-associated protein Cas6/Cse3/CasE [Azoarcus communis] |
| 07661 | 10 | hypothetical protein [Aromatoleum evansii] |
| 00098 | 10 | DUF4157 domain-containing protein [Aromatoleum toluclasticum] |
| 04277 | 10 | malonate decarboxylase holo-[acyl-carrier-protein] synthase [Nitrogeniibacter aestuarii] |
| 00289 | 10 | hypothetical protein [Aromatoleum toluclasticum] |
| 00122 | 10 | SDR family oxidoreductase [Azoarcus taiwanensis] |
| 10569 | 10 | penicillin acylase family protein [Azoarcus indigens] |
| 08294 | 10 | MULTISPECIES: ester cyclase [Aromatoleum] |
| 07071 | 10 | hypothetical protein [Aromatoleum toluclasticum] |
| 09568 | 10 | hypothetical protein [Azoarcus communis] |
| 02383 | 10 | Rossmann-like and DUF2520 domain-containing protein [Azoarcus communis] |
| 05079 | 10 | DUF1320 domain-containing protein [Aromatoleum pumilum] |
| 00310 | 10 | AlpA family phage regulatory protein [Azoarcus olearius] |
| 06848 | 10 | AbrB/MazE/SpoVT family DNA-binding domain-containing protein [Aromatoleum aromaticum] |
| 01365 | 10 | hypothetical protein [Denitromonas halophilus] |
| 06795 | 10 | MULTISPECIES: type IV conjugative transfer system lipoprotein TraV [Betaproteobacteria] |
| 00897 | 10 | cupin domain-containing protein [Azoarcus indigens] |
| 08096 | 10 | WYL domain-containing protein [Zoogloea oleivorans] |
| 05036 | 10 | hypothetical protein [Aromatoleum toluclasticum] |
| 07152 | 10 | type II toxin-antitoxin system RelE/ParE family toxin [Denitromonas halophilus] |
| 05579 | 10 | glucans biosynthesis glucosyltransferase MdoH [Nitrogeniibacter aestuarii] |
| 02366 | 10 | MULTISPECIES: hypothetical protein [Aromatoleum] |
| 02369 | 10 | endonuclease/exonuclease/phosphatase family protein [Aromatoleum toluclasticum] |
| 01814 | 10 | nuclear transport factor 2 family protein [Azoarcus indigens] |
| 04494 | 10 | hypothetical protein [Aromatoleum toluclasticum] |
| 05762 | 10 | ChaN family lipoprotein [Aromatoleum aromaticum] |
| 05136 | 10 | hypothetical protein [Denitromonas halophilus] |
| 10539 | 10 | MULTISPECIES: type II toxin-antitoxin system HicB family antitoxin [Aromatoleum] |
| 08060 | 10 | DUF935 domain-containing protein [Candidatus Dactylopiibacterium carminicum] |
| 08937 | 10 | glycine cleavage system protein H [Azoarcus rhizosphaerae] |
| 09057 | 10 | hypothetical protein [Azoarcus olearius] |
| 05175 | 10 | DUF302 domain-containing protein [Aromatoleum toluclasticum] |
| 08888 | 10 | ankyrin repeat domain-containing protein [Nitrogeniibacter aestuarii] |
| 08883 | 10 | MULTISPECIES: response regulator [Thauera] |
| 05655 | 10 | MULTISPECIES: TraB/VirB10 family protein [Betaproteobacteria] |
| 04202 | 10 | IS66 family transposase zinc-finger binding domain-containing protein, partial [Azoarcus sp. Aa7] |
| 04201 | 10 | MULTISPECIES: hypothetical protein [Thauera] |
| 06659 | 10 | hypothetical protein [Aromatoleum toluclasticum] |
| 04197 | 10 | DUF2760 domain-containing protein [Aromatoleum aromaticum] |
| 00486 | 10 | hypothetical protein [Azospira oryzae] |
| 08881 | 10 | GspH/FimT family pseudopilin [Azoarcus indigens] |
| 05704 | 10 | bile acid:sodium symporter family protein [Denitromonas halophilus] |
| 01555 | 10 | hypothetical protein [Denitromonas halophilus] |
| 02314 | 10 | thioredoxin domain-containing protein [Denitromonas halophilus] |
| 04553 | 10 | ATP-dependent helicase [Aromatoleum diolicum] |
| 08890 | 10 | hypothetical protein [Aromatoleum toluclasticum] |
| 00511 | 10 | MULTISPECIES: metallophosphoesterase [Niveibacterium] |
| 06142 | 10 | DUF3175 domain-containing protein [Azoarcus indigens] |
| 08912 | 10 | zonular occludens toxin [Azoarcus indigens] |
| 09738 | 10 | GH1 family beta-glucosidase [Niveibacterium microcysteis] |
| 06426 | 10 | hypothetical protein [Azoarcus indigens] |
| 07681 | 10 | cupredoxin domain-containing protein [Aromatoleum diolicum] |
| 00735 | 10 | hypothetical protein [Azoarcus halotolerans] |
| 10266 | 10 | acyl-CoA thioesterase [Propionivibrio dicarboxylicus] |
| 03384 | 10 | hypothetical protein [Aromatoleum pumilum] |
| 07506 | 10 | monooxygenase [Azoarcus olearius] |
| 07503 | 10 | SMP-30/gluconolactonase/LRE family protein [Azoarcus olearius] |
| 05792 | 10 | transposase [Thauera propionica] |
| 05606 | 10 | mobilization protein [Denitromonas halophilus] |
| 00680 | 10 | hypothetical protein [Azoarcus indigens] |
| 09760 | 10 | CbtB-domain containing protein [Aromatoleum toluclasticum] |
| 06705 | 10 | glycosyltransferase [Aromatoleum evansii] |
| 07504 | 10 | PilW family protein [Aromatoleum toluclasticum] |
| 09826 | 10 | glycine zipper 2TM domain-containing protein [Aromatoleum toluclasticum] |
| 08147 | 10 | cytochrome B6 [Azoarcus indigens] |
| 02873 | 10 | MULTISPECIES: type-F conjugative transfer system secretin TraK [Betaproteobacteria] |
| 10046 | 10 | DUF504 domain-containing protein [Aromatoleum diolicum] |
| 08322 | 10 | rhamnogalacturonan acetylesterase [Uliginosibacterium aquaticum] |
| 03819 | 10 | hypothetical protein [Denitromonas halophilus] |
| 05919 | 10 | hypothetical protein [Aromatoleum aromaticum] |
| 02988 | 10 | hypothetical protein [Azoarcus indigens] |
| 05963 | 10 | hypothetical protein [Aromatoleum toluclasticum] |
| 06288 | 10 | hypothetical protein [Aromatoleum toluclasticum] |
| 01761 | 10 | hypothetical protein [Aromatoleum toluclasticum] |
| 09943 | 10 | hypothetical protein [Denitromonas halophilus] |
| 07996 | 10 | hypothetical protein [Azoarcus communis] |
| 06390 | 10 | hypothetical protein [Azoarcus indigens] |
| 03680 | 10 | alkaline phosphatase family protein [Thauera chlorobenzoica] |

| | | |
|---|---|---|
| 08663 | 10 | outer membrane beta-barrel protein [Nitrogeniibacter aestuarii] |
| 08564 | 10 | hypothetical protein [Azoarcus communis] |
| 03639 | 10 | Hpt domain-containing protein [Aromatoleum bremense] |
| 08666 | 10 | phasin family protein [Aromatoleum toluclasticum] |
| 03303 | 10 | M23 family metallopeptidase [Aromatoleum evansii] |
| 08563 | 10 | class I SAM-dependent methyltransferase [Aromatoleum toluclasticum] |
| 07380 | 10 | OmpA family protein [Azoarcus communis] |
| 06385 | 10 | class C beta-lactamase [Azoarcus communis] |
| 07379 | 10 | WbqC family protein [Azoarcus communis] |
| 05375 | 10 | Atrophin-1 multi-domain protein [Azoarcus indigens] |
| 02891 | 10 | DsrE family protein [Aromatoleum pumilum] |
| 00444 | 10 | anaerobic ribonucleoside-triphosphate reductase [Candidatus Dactylopiibacterium carminicum] |
| 06443 | 10 | flippase-like domain-containing protein [Azoarcus communis] |
| 03926 | 10 | adenylyl-sulfate kinase [Azoarcus indigens] |
| 07399 | 10 | hypothetical protein [Azoarcus indigens] |
| 10347 | 10 | acetyltransferase [Aromatoleum diolicum] |
| 09935 | 10 | SH3 domain-containing protein [Denitromonas halophilus] |
| 03585 | 10 | MULTISPECIES: GNAT family N-acetyltransferase [Niveibacterium] |
| 05341 | 10 | MULTISPECIES: type IV conjugative transfer system protein TraE [Betaproteobacteria] |
| 07283 | 10 | VOC family protein [Nitrogeniibacter aestuarii] |
| 07863 | 10 | hypothetical protein [Azoarcus communis] |
| 06584 | 10 | hypothetical protein [Azoarcus indigens] |
| 10153 | 10 | hypothetical protein [Azoarcus indigens] |
| 09387 | 10 | ChaB family protein [Azoarcus indigens] |
| 03587 | 10 | class I SAM-dependent methyltransferase [Denitromonas halophilus] |
| 00979 | 10 | lipocalin-like domain-containing protein [Azoarcus communis] |
| 02913 | 10 | FxDxF family PEP-CTERM protein [Denitromonas halophilus] |
| 09354 | 10 | DUF3325 domain-containing protein [Nitrogeniibacter aestuarii] |
| 05361 | 10 | carbonic anhydrase [Aromatoleum toluclasticum] |
| 07565 | 10 | glycosyltransferase family 39 protein [Denitromonas halophilus] |
| 00149 | 10 | fatty acid cis/trans isomerase [Nitrogeniibacter aestuarii] |
| 07289 | 10 | MULTISPECIES: DUF642 domain-containing protein [Niveibacterium] |
| 01780 | 10 | MULTISPECIES: benzylsuccinate synthase beta subunit family protein [Aromatoleum] |
| 06907 | 10 | helix-turn-helix transcriptional regulator [Azoarcus indigens] |
| 03811 | 10 | hypothetical protein [Denitromonas halophilus] |
| 03520 | 10 | OpgC domain-containing protein [Azoarcus communis] |
| 10364 | 10 | sulfite exporter TauE/SafE family protein [Azoarcus communis] |
| 04865 | 10 | O-antigen ligase family protein [Aromatoleum petrolei] |
| 10359 | 10 | Na+/H+ antiporter NhaA [Azoarcus indigens] |
| 08681 | 10 | SEL1-like repeat protein [Aromatoleum aromaticum] |
| 09502 | 10 | DUF2167 domain-containing protein [Azoarcus indigens] |
| 00650 | 10 | P-II family nitrogen regulator [Nitrogeniibacter aestuarii] |
| 03866 | 10 | MULTISPECIES: sodium-translocating pyrophosphatase [Niveibacterium] |
| 06624 | 10 | VWA domain-containing protein [Aromatoleum toluclasticum] |
| 06301 | 10 | DUF3427 domain-containing protein [Aromatoleum toluclasticum] |
| 04641 | 10 | hypothetical protein [Aromatoleum toluclasticum] |
| 04003 | 10 | transposase [Zoogloea oleivorans] |
| 02214 | 10 | acetylornithine deacetylase [Azoarcus communis] |
| 05538 | 10 | hypothetical protein [Aromatoleum toluclasticum] |
| 05867 | 10 | cytochrome c family protein [Aromatoleum toluclasticum] |
| 05105 | 9 | cupin domain-containing protein [Denitromonas halophilus] |
| 06452 | 9 | hypothetical protein [Denitromonas halophilus] |
| 06732 | 9 | FUSC family protein [Azoarcus indigens] |
| 02161 | 9 | hypothetical protein [Azoarcus communis] |
| 08396 | 9 | hypothetical protein [Aromatoleum evansii] |
| 09211 | 9 | phage holin family protein [Azoarcus indigens] |
| 05787 | 9 | copper resistance protein B [Denitromonas halophilus] |
| 06586 | 9 | cobalamin-dependent protein [Aromatoleum toluclasticum] |
| 00604 | 9 | hypothetical protein [Azoarcus indigens] |
| 01042 | 9 | hypothetical protein [Aromatoleum pumilum] |
| 10314 | 9 | nucleotidyltransferase [Aromatoleum bremense] |
| 08496 | 9 | FecR domain-containing protein [Azoarcus communis] |
| 03381 | 9 | gluconate 2-dehydrogenase subunit 3 family protein [Azoarcus communis] |
| 00441 | 9 | helix-turn-helix domain-containing protein [Thauera aminoaromatica] |
| 01901 | 9 | 3'-5' exoribonuclease [Niveibacterium sp. COAC-50] |
| 03742 | 9 | hypothetical protein [Aromatoleum toluclasticum] |
| 07041 | 9 | 2-oxoacid:acceptor oxidoreductase family protein [Rhodocyclus purpureus] |
| 08591 | 9 | 3-hydroxyacyl-CoA dehydrogenase NAD-binding domain-containing protein [Aromatoleum toluclasticum] |
| 07317 | 9 | MULTISPECIES: virulence factor [Aromatoleum] |
| 05906 | 9 | RNA-guided endonuclease TnpB family protein [Aromatoleum aromaticum] |
| 09170 | 9 | MULTISPECIES: integrase domain-containing protein [Betaproteobacteria] |
| 00387 | 9 | tetratricopeptide repeat protein [Denitromonas halophilus] |
| 08326 | 9 | excinuclease ATPase subunit [Denitromonas halophilus] |
| 08173 | 9 | FeoC-like transcriptional regulator [Nitrogeniibacter aestuarii] |
| 01467 | 9 | cytochrome c3 family protein, partial [Propionivibrio dicarboxylicus] |
| 06669 | 9 | hypothetical protein [Aromatoleum toluclasticum] |
| 08602 | 9 | hypothetical protein [Aromatoleum aromaticum] |
| 03692 | 9 | DNA-binding protein [Aromatoleum diolicum] |
| 05582 | 9 | dienelactone hydrolase family protein [Propionivibrio dicarboxylicus] |
| 08521 | 9 | DEAD/DEAH box helicase [Aromatoleum toluolicum] |
| 07286 | 9 | C-GCAxxG-C-C family protein [Aromatoleum diolicum] |
| 05173 | 9 | helix-turn-helix domain-containing protein [Denitromonas halophilus] |
| 04816 | 9 | DUF2730 family protein [Aromatoleum tolulyticum] |
| 07148 | 9 | KamA family radical SAM protein [Aromatoleum toluclasticum] |
| 09455 | 9 | hypothetical protein [Denitromonas halophilus] |
| 01908 | 9 | EAL domain-containing protein [Nitrogeniibacter aestuarii] |
| 02348 | 9 | hypothetical protein [Denitromonas halophilus] |
| 03362 | 9 | FlgO family outer membrane protein [Rhodocyclus purpureus] |
| 03721 | 9 | DUF2325 domain-containing protein [Azoarcus indigens] |
| 02519 | 9 | TIGR02680 family protein [Azoarcus indigens] |
| 07711 | 9 | heavy-metal-associated domain-containing protein [Aromatoleum toluclasticum] |
| 04218 | 9 | AhpC/TSA family protein [Denitromonas halophilus] |
| 01015 | 9 | hypothetical protein [Denitromonas halophilus] |
| 08543 | 9 | glycerol dehydrogenase [Azoarcus indigens] |

| | | |
|---|---|---|
| **04373** | 9 | hypothetical protein [Azoarcus communis] |
| **01748** | 9 | NAD(P)/FAD-dependent oxidoreductase [Nitrogeniibacter aestuarii] |
| **00049** | 9 | FRG domain-containing protein [Azoarcus indigens] |
| **03087** | 9 | GNAT family N-acetyltransferase [Denitromonas halophilus] |
| **00641** | 9 | ethylbenzene dehydrogenase-related protein [Aromatoleum toluclasticum] |
| **09128** | 9 | STN domain-containing protein [Azoarcus rhizosphaerae] |
| **05791** | 9 | PEP-CTERM sorting domain-containing protein [Nitrogeniibacter aestuarii] |
| **03449** | 9 | hypothetical protein [Azoarcus olearius] |
| **10311** | 9 | hypothetical protein [Aromatoleum toluclasticum] |
| **09209** | 9 | hypothetical protein [Denitromonas halophilus] |
| **02341** | 9 | PEP-CTERM sorting domain-containing protein [Nitrogeniibacter aestuarii] |
| **08934** | 9 | MULTISPECIES: methylenetetrahydrofolate reductase C-terminal domain-containing protein [Aromatoleum] |
| **03886** | 9 | mechanosensitive ion channel [Azoarcus indigens] |
| **03567** | 9 | gamma-glutamylcyclotransferase [Aromatoleum toluclasticum] |
| **09365** | 9 | DUF1804 family protein [Aromatoleum pumilum] |
| **09621** | 9 | alpha/beta hydrolase [Niveibacterium umoris] |
| **07652** | 9 | cytochrome c oxidase assembly protein [Azoarcus indigens] |
| **01514** | 9 | transporter substrate-binding domain-containing protein [Niveibacterium umoris] |
| **01298** | 9 | class I SAM-dependent methyltransferase [Azoarcus communis] |
| **02134** | 9 | hypothetical protein [Aromatoleum aromaticum] |
| **00201** | 9 | prepilin-type N-terminal cleavage/methylation domain-containing protein [Propionivibrio dicarboxylicus] |
| **05713** | 9 | AIPR family protein [Aromatoleum buckelii] |
| **10099** | 9 | MULTISPECIES: hemerythrin domain-containing protein [Niveibacterium] |
| **09688** | 9 | hypothetical protein [Aromatoleum toluclasticum] |
| **08974** | 9 | hypothetical protein [Nitrogeniibacter aestuarii] |
| **09695** | 9 | protein glxC [Azoarcus communis] |
| **09993** | 9 | flocculation-associated PEP-CTERM protein PepA [Aromatoleum aromaticum] |
| **05399** | 9 | TIGR02677 family protein [Azoarcus indigens] |
| **02407** | 9 | EAL domain-containing protein [Nitrogeniibacter aestuarii] |
| **07644** | 9 | hypothetical protein [Niveibacterium microcysteis] |
| **10455** | 9 | MULTISPECIES: hypothetical protein [Betaproteobacteria] |
| **02401** | 9 | MULTISPECIES: MAPEG family protein [Niveibacterium] |
| **01314** | 9 | bifunctional glutathionylspermidine amidase/synthase [Denitromonas halophilus] |
| **06533** | 9 | DNA-binding protein [Aromatoleum evansii] |
| **06687** | 9 | DUF475 domain-containing protein [Candidatus Dactylopiibacterium carminicum] |
| **02331** | 9 | hypothetical protein [Zoogloea oleivorans] |
| **02132** | 9 | ligase-associated DNA damage response DEXH box helicase [Azoarcus olearius] |
| **06688** | 9 | DUF971 domain-containing protein [Azoarcus communis] |
| **08217** | 9 | hypothetical protein [Aromatoleum pumilum] |
| **01793** | 9 | FHA domain-containing protein [Azoarcus indigens] |
| **01005** | 9 | transcriptional regulator BetI [Aromatoleum toluclasticum] |
| **00412** | 9 | formate dehydrogenase [Denitromonas halophilus] |
| **05759** | 9 | amino acid racemase [Denitromonas halophilus] |
| **07902** | 9 | SPOR domain-containing protein [Nitrogeniibacter aestuarii] |
| **03015** | 9 | EAL domain-containing protein [Azoarcus communis] |
| **01811** | 9 | glycosyltransferase [Aromatoleum bremense] |
| **07183** | 9 | sensor histidine kinase [Azoarcus communis] |
| **04822** | 9 | FkbM family methyltransferase [Azoarcus nasutitermitis] |
| **06474** | 9 | integrase core domain-containing protein, partial [Thauera linaloolentis] |
| **04367** | 9 | hypothetical protein [Azoarcus communis] |
| **05383** | 9 | BatD family protein [Aromatoleum diolicum] |
| **04268** | 9 | hypothetical protein [Aromatoleum bremense] |
| **00602** | 9 | DUF1911 domain-containing protein [Nitrogeniibacter aestuarii] |
| **04332** | 9 | DNA-binding transcriptional regulator [Azoarcus communis] |
| **00982** | 9 | MULTISPECIES: hypothetical protein [Aromatoleum] |
| **09905** | 9 | polyhydroxyalkanoate depolymerase [Azoarcus halotolerans] |
| **05478** | 9 | DUF4160 domain-containing protein [Aromatoleum bremense] |
| **09988** | 9 | helix-turn-helix domain-containing protein [Aromatoleum aromaticum] |
| **06841** | 9 | restriction endonuclease subunit S [Denitromonas sp. IR12] |
| **10387** | 9 | MSMEG_0568 family radical SAM protein [Azoarcus indigens] |
| **03334** | 9 | hypothetical protein [Azoarcus communis] |
| **01037** | 9 | BrnT family toxin [Azoarcus communis] |
| **02371** | 9 | iron-regulated protein [Azoarcus communis] |
| **04328** | 9 | hypothetical protein [Azoarcus indigens] |
| **09040** | 9 | poly-beta-1,6 N-acetyl-D-glucosamine synthase [Nitrogeniibacter aestuarii] |
| **05907** | 9 | hypothetical protein [Aromatoleum bremense] |
| **09672** | 9 | amino acid ABC transporter permease [Aromatoleum toluvorans] |
| **09765** | 9 | potassium channel family protein [Aromatoleum toluclasticum] |
| **01886** | 9 | alpha/beta hydrolase [Azoarcus communis] |
| **06626** | 9 | hypothetical protein [Thauera butanivorans] |
| **05957** | 9 | hypothetical protein [Azoarcus nasutitermitis] |
| **06100** | 9 | hypothetical protein [Denitromonas halophilus] |
| **00942** | 9 | MULTISPECIES: oxidase [Niveibacterium] |
| **06790** | 9 | hypothetical protein [Aromatoleum toluclasticum] |
| **08433** | 9 | DUF2029 domain-containing protein [Aromatoleum toluclasticum] |
| **09266** | 9 | hypothetical protein [Azoarcus communis] |
| **09642** | 9 | FapA family protein [Aromatoleum toluclasticum] |
| **00418** | 9 | hypothetical protein [Aromatoleum toluclasticum] |
| **10339** | 9 | LuxR family transcriptional regulator [Azoarcus indigens] |
| **10048** | 9 | GNAT family N-acetyltransferase [Denitromonas halophilus] |
| **03169** | 9 | lysine 5,6-aminomutase subunit alpha [Aromatoleum toluclasticum] |
| **02727** | 9 | hypothetical protein [Aromatoleum toluclasticum] |
| **03473** | 9 | DUF4381 domain-containing protein [Aromatoleum diolicum] |
| **00055** | 9 | putative zinc-binding metallopeptidase [Azoarcus indigens] |
| **09401** | 9 | VWA domain-containing protein [Aromatoleum aromaticum] |
| **10484** | 9 | hypothetical protein [Aromatoleum aromaticum] |
| **03793** | 9 | PAS domain S-box protein [Azoarcus olearius] |
| **06618** | 9 | phage protease [Aromatoleum pumilum] |
| **05884** | 9 | DUF4178 domain-containing protein [Azoarcus sp. Aa7] |
| **07560** | 9 | hypothetical protein [Aromatoleum bremense] |
| **01609** | 9 | four-carbon acid sugar kinase family protein [Aromatoleum diolicum] |
| **06996** | 9 | DUF11 domain-containing protein [Azoarcus communis] |
| **03102** | 9 | TIGR02679 family protein [Azoarcus indigens] |
| **04117** | 9 | HPP family protein [Azoarcus olearius] |

| | | |
|---|---|---|
| 01416 | 9 | thioesterase family protein [Aromatoleum toluclasticum] |
| 08791 | 9 | hypothetical protein [Azoarcus olearius] |
| 05993 | 9 | Mu-like prophage major head subunit gpT family protein [Aromatoleum pumilum] |
| 02700 | 9 | ferrous iron transport protein A [Nitrogeniibacter aestuarii] |
| 00690 | 9 | porin [Uliginosibacterium aquaticum] |
| 09548 | 9 | hypothetical protein [Azoarcus communis] |
| 00496 | 9 | alpha-amylase family glycosyl hydrolase [Niveibacterium microcysteis] |
| 08163 | 9 | hypothetical protein [Aromatoleum evansii] |
| 00468 | 9 | DNA-binding protein [Azoarcus sp. Aa7] |
| 07837 | 9 | TIGR02678 family protein [Azoarcus indigens] |
| 00291 | 9 | VOC family protein [Azoarcus olearius] |
| 06250 | 9 | MBL fold metallo-hydrolase [Denitromonas halophilus] |
| 04393 | 9 | RHS repeat protein [Azoarcus indigens] |
| 05814 | 9 | MaoC family dehydratase N-terminal domain-containing protein [Aromatoleum petrolei] |
| 04418 | 9 | hypothetical protein [Denitromonas halophilus] |
| 08687 | 9 | hypothetical protein [Azoarcus indigens] |
| 09306 | 9 | hypothetical protein [Aromatoleum toluclasticum] |
| 01675 | 9 | N(2)-acetyl-L-2,4-diaminobutanoate deacetylase DoeB [Denitromonas halophilus] |
| 01143 | 9 | VIT1/CCC1 transporter family protein [Azoarcus indigens] |
| 01683 | 9 | DUF1801 domain-containing protein [Aromatoleum petrolei] |
| 05629 | 9 | inorganic diphosphatase [Nitrogeniibacter aestuarii] |
| 06162 | 9 | filamentous hemagglutinin N-terminal domain-containing protein [Azoarcus indigens] |
| 06871 | 9 | MULTISPECIES: hypothetical protein [Thauera] |
| 02799 | 9 | glycosyltransferase family 4 protein [Azoarcus olearius] |
| 10059 | 9 | glycosyltransferase [Azoarcus halotolerans] |
| 00483 | 9 | AraC family transcriptional regulator [Azoarcus olearius] |
| 08700 | 9 | hypothetical protein [Azoarcus communis] |
| 07347 | 9 | hypothetical protein [Nitrogeniibacter aestuarii] |
| 01123 | 9 | N-acetylmuramoyl-L-alanine amidase [Aromatoleum evansii] |
| 02822 | 9 | hypothetical protein [Azoarcus indigens] |
| 08721 | 9 | 3-aminobutyryl-CoA ammonia lyase [Aromatoleum evansii] |
| 00474 | 9 | 6-carboxytetrahydropterin synthase [Azoarcus communis] |
| 06274 | 9 | Flp pilus assembly protein CpaB [Azoarcus nasutitermitis] |
| 03152 | 9 | type II toxin-antitoxin system RelE/ParE family toxin [Azoarcus communis] |
| 06621 | 9 | TIR domain-containing protein [Denitromonas halophilus] |
| 06823 | 9 | rhomboid family intramembrane serine protease [Denitromonas halophilus] |
| 07583 | 9 | zinc-binding dehydrogenase [Aromatoleum toluclasticum] |
| 00948 | 9 | hypothetical protein [Aromatoleum toluclasticum] |
| 03280 | 9 | hypothetical protein [Aromatoleum aromaticum] |
| 04650 | 9 | DUF4043 family protein [Azoarcus indigens] |
| 01154 | 9 | MarR family winged helix-turn-helix transcriptional regulator [Azoarcus indigens] |
| 03140 | 9 | hypothetical protein [Azoarcus communis] |
| 10219 | 9 | MSHA biogenesis protein MshK [Niveibacterium umoris] |
| 00227 | 9 | hypothetical protein [Denitromonas halophilus] |
| 03132 | 9 | glycosyltransferase [Rhodocyclus purpureus] |
| 10335 | 9 | NERD domain-containing protein [Azoarcus indigens] |
| 03801 | 9 | DUF6519 domain-containing protein [Denitromonas halophilus] |
| 00365 | 9 | hypothetical protein [Azoarcus indigens] |
| 08794 | 9 | hypothetical protein [Azoarcus communis] |
| 06105 | 9 | DUF1156 domain-containing protein [Aromatoleum bremense] |
| 01080 | 9 | helix-turn-helix domain-containing protein [Azoarcus indigens] |
| 09346 | 9 | hypothetical protein [Denitromonas halophilus] |
| 08628 | 9 | HlyD family efflux transporter periplasmic adaptor subunit [Azoarcus halotolerans] |
| 03490 | 9 | MULTISPECIES: queuosine precursor transporter [Niveibacterium] |
| 05656 | 9 | IS200/IS605 family transposase [Aromatoleum aromaticum] |
| 02592 | 9 | FAD-binding protein, partial [Thauera butanivorans] |
| 07980 | 9 | VCBS domain-containing protein, partial [Niveibacterium umoris] |
| 07126 | 9 | hypothetical protein [Aromatoleum bremense] |
| 02575 | 9 | MULTISPECIES: IS1380 family transposase [Thauera] |
| 06331 | 9 | FAD-binding oxidoreductase [Nitrogeniibacter aestuarii] |
| 03687 | 9 | MULTISPECIES: GTP-binding protein, partial [Betaproteobacteria] |
| 05190 | 9 | hypothetical protein [Niveibacterium microcysteis] |
| 00348 | 9 | class II glutamine amidotransferase [Azoarcus communis] |
| 09848 | 9 | MaoC family dehydratase N-terminal domain-containing protein [Denitromonas halophilus] |
| 09861 | 9 | helix-turn-helix domain-containing protein [Azoarcus indigens] |
| 00053 | 9 | hypothetical protein [Azoarcus indigens] |
| 02917 | 9 | RHS repeat protein, partial [Azoarcus communis] |
| 05459 | 9 | LTA synthase family protein [Azoarcus nasutitermitis] |
| 09227 | 9 | ATP/GTP-binding protein [Nitrogeniibacter aestuarii] |
| 09767 | 9 | isoaspartyl peptidase/L-asparaginase [Aromatoleum toluclasticum] |
| 04200 | 9 | exopolysaccharide biosynthesis protein [Aromatoleum aromaticum] |
| 04199 | 9 | hypothetical protein [Azoarcus olearius] |
| 10482 | 9 | alpha/beta hydrolase [Aromatoleum toluclasticum] |
| 07020 | 9 | MULTISPECIES: hypothetical protein [Niveibacterium] |
| 10442 | 9 | DUF839 domain-containing protein [Denitromonas halophilus] |
| 03758 | 9 | urate hydroxylase PuuD [Denitromonas halophilus] |
| 06652 | 9 | hypothetical protein [Aromatoleum toluclasticum] |
| 04674 | 9 | MULTISPECIES: PaaI family thioesterase [Azospira] |
| 09979 | 9 | oxalurate catabolism protein HpxZ [Azoarcus communis] |
| 07402 | 9 | sulfatase-like hydrolase/transferase [Aromatoleum bremense] |
| 01726 | 9 | TIGR02647 family protein [Denitromonas halophilus] |
| 02565 | 9 | MULTISPECIES: sugar phosphate isomerase/epimerase [Azoarcus] |
| 05492 | 9 | helix-turn-helix domain-containing protein [Denitromonas halophilus] |
| 10144 | 9 | hypothetical protein [Aromatoleum toluclasticum] |
| 08712 | 9 | hypothetical protein [Azoarcus indigens] |
| 01865 | 9 | hypothetical protein [Aromatoleum tolulyticum] |
| 04148 | 9 | phospholipase D family protein [Azoarcus olearius] |
| 01589 | 9 | acetyltransferase, CysE/LacA/LpxA/NodL family [Azoarcus sp. KH32C] |
| 04138 | 9 | hypothetical protein [Thauera terpenica] |
| 01211 | 9 | SPFH domain-containing protein [Azoarcus sp. Aa7] |
| 04575 | 9 | zf-HC2 domain-containing protein [Denitromonas halophilus] |
| 10142 | 9 | hypothetical protein [Aromatoleum evansii] |
| 07482 | 9 | nuclear transport factor 2 family protein [Denitromonas halophilus] |
| 06769 | 9 | hypothetical protein [Denitromonas halophilus] |

| | | |
|---|---|---|
| 01393 | 9 | ATP-binding protein [Azoarcus halotolerans] |
| 07104 | 9 | MBL fold metallo-hydrolase [Denitromonas halophilus] |
| 09345 | 9 | translation initiation factor Sui1 [Denitromonas halophilus] |
| 10038 | 9 | MULTISPECIES: hypothetical protein [Azospira] |
| 06197 | 9 | PEP-CTERM sorting domain-containing protein [Nitrogeniibacter aestuarii] |
| 07744 | 9 | 3,4-dihydroxyphenylacetate 2,3-dioxygenase [Azoarcus communis] |
| 05204 | 9 | DUF499 domain-containing protein [Aromatoleum bremense] |
| 10214 | 9 | hypothetical protein [Aromatoleum toluclasticum] |
| 07406 | 9 | glycoside hydrolase family 28 protein [Candidatus Dactylopiibacterium carminicum] |
| 03114 | 9 | hypothetical protein [Denitromonas halophilus] |
| 10076 | 9 | DUF2442 domain-containing protein [Aromatoleum buckelii] |
| 03597 | 9 | DUF1638 domain-containing protein [Aromatoleum toluclasticum] |
| 07134 | 9 | site-specific integrase [Thauera propionica] |
| 05612 | 9 | DUF5668 domain-containing protein [Denitromonas halophilus] |
| 09858 | 9 | VWA domain-containing protein [Aromatoleum evansii] |
| 02609 | 9 | hypothetical protein [Aromatoleum toluclasticum] |
| 09230 | 9 | FkbM family methyltransferase [Azoarcus communis] |
| 05811 | 9 | OmpA family protein [Azoarcus communis] |
| 05012 | 8 | hypothetical protein [Azoarcus rhizosphaerae] |
| 07552 | 8 | ion transporter [Denitromonas halophilus] |
| 07127 | 8 | hypothetical protein [Propionivibrio dicarboxylicus] |
| 05730 | 8 | cupin domain-containing protein [Azoarcus communis] |
| 02212 | 8 | MULTISPECIES: hypothetical protein [Niveibacterium] |
| 01447 | 8 | hypothetical protein [Aromatoleum toluclasticum] |
| 01432 | 8 | molecular chaperone TorD family protein [Aromatoleum diolicum] |
| 02217 | 8 | tripartite tricarboxylate transporter TctB family protein [Aromatoleum diolicum] |
| 07128 | 8 | CDP-alcohol phosphatidyltransferase family protein [Azoarcus communis] |
| 03631 | 8 | organoarsenical effux MFS transporter ArsJ [Azoarcus communis] |
| 05022 | 8 | hypothetical protein [Azoarcus indigens] |
| 04460 | 8 | CHAT domain-containing protein [Denitromonas halophilus] |
| 09156 | 8 | NIPSNAP family protein [Azoarcus sp. DD4] |
| 06180 | 8 | DUF1302 family protein [Azoarcus sp. L1K30] |
| 00757 | 8 | alpha-1,4-glucan--maltose-1-phosphate maltosyltransferase [Azoarcus indigens] |
| 10136 | 8 | alginate lyase family protein [Zoogloea dura] |
| 01744 | 8 | surface lipoprotein assembly modifier, partial [Thauera linaloolentis] |
| 09475 | 8 | hypothetical protein [Azoarcus indigens] |
| 07413 | 8 | FAD-dependent oxidoreductase [Azoarcus olearius] |
| 07267 | 8 | YqaJ viral recombinase family protein [Azoarcus indigens] |
| 07263 | 8 | hypothetical protein [Azoarcus indigens] |
| 06097 | 8 | DMT family transporter [Oryzomicrobium terrae] |
| 06126 | 8 | aromatic ring-hydroxylating dioxygenase subunit alpha [Thauera linaloolentis] |
| 00635 | 8 | NAD(P)-dependent oxidoreductase [Azoarcus indigens] |
| 01496 | 8 | methyltransferase domain-containing protein [Aromatoleum pumilum] |
| 06127 | 8 | hypothetical protein [Aromatoleum aromaticum] |
| 04872 | 8 | D-Ala-D-Ala carboxypeptidase family metallohydrolase [Nitrogeniibacter aestuarii] |
| 09312 | 8 | alpha/beta hydrolase [Niveibacterium sp. COAC-50] |
| 06727 | 8 | helix-turn-helix domain-containing protein [Aromatoleum tolulyticum] |
| 05628 | 8 | hypothetical protein [Azoarcus rhizosphaerae] |
| 01963 | 8 | RES family NAD+ phosphorylase [Aromatoleum toluclasticum] |
| 06093 | 8 | GNAT family N-acetyltransferase [Azoarcus communis] |
| 01746 | 8 | fumarate reductase cytochrome b subunit, partial [Thauera phenolivorans] |
| 01750 | 8 | hypothetical protein [Propionivibrio dicarboxylicus] |
| 04692 | 8 | DUF5615 family PIN-like protein [Aromatoleum aromaticum] |
| 01752 | 8 | hypothetical protein [Aromatoleum toluclasticum] |
| 01802 | 8 | hypothetical protein [Aromatoleum evansii] |
| 04770 | 8 | CHASE domain-containing protein [Denitromonas halophilus] |
| 03592 | 8 | hypothetical protein [Denitromonas halophilus] |
| 08232 | 8 | hypothetical protein [Azoarcus indigens] |
| 04790 | 8 | DegT/DnrJ/EryC1/StrS family aminotransferase [Niveibacterium microcysteis] |
| 09419 | 8 | glycoside hydrolase family 19 protein [Azoarcus taiwanensis] |
| 07344 | 8 | DEAD/DEAH box helicase [Aromatoleum aromaticum] |
| 01789 | 8 | hypothetical protein [Aromatoleum toluclasticum] |
| 05657 | 8 | hypothetical protein [Aromatoleum buckelii] |
| 01856 | 8 | glutamate--cysteine ligase [Denitromonas halophilus] |
| 09445 | 8 | M48 family metallopeptidase [Aromatoleum pumilum] |
| 10185 | 8 | hypothetical protein [Aromatoleum toluclasticum] |
| 04722 | 8 | nitrous oxide reductase accessory protein NosL [Azospira oryzae] |
| 04811 | 8 | hypothetical protein [Denitromonas halophilus] |
| 04813 | 8 | hypothetical protein [Denitromonas halophilus] |
| 05945 | 8 | MULTISPECIES: sulfide/dihydroorotate dehydrogenase-like FAD/NAD-binding protein [Niveibacterium] |
| 06763 | 8 | hypothetical protein [Aromatoleum toluclasticum] |
| 04910 | 8 | glycine zipper 2TM domain-containing protein [Uliginosibacterium gangwonense] |
| 02073 | 8 | hypothetical protein [Denitromonas halophilus] |
| 04533 | 8 | hypothetical protein [Niveibacterium umoris] |
| 04527 | 8 | alpha/beta hydrolase [Denitromonas halophilus] |
| 07190 | 8 | DNA-binding protein [Aromatoleum bremense] |
| 06781 | 8 | MlaD family protein [Azoarcus halotolerans] |
| 01530 | 8 | DUF3293 domain-containing protein [Aromatoleum pumilum] |
| 02101 | 8 | hypothetical protein [Aromatoleum evansii] |
| 07515 | 8 | beta-hydroxyacyl-ACP dehydratase [Denitromonas halophilus] |
| 04511 | 8 | DUF4124 domain-containing protein [Azoarcus indigens] |
| 07170 | 8 | type II toxin-antitoxin system MqsA family antitoxin [Aromatoleum evansii] |
| 06717 | 8 | hypothetical protein [Denitromonas halophilus] |
| 04498 | 8 | hypothetical protein [Aromatoleum toluclasticum] |
| 04490 | 8 | TnsA endonuclease N-terminal domain-containing protein [Aromatoleum evansii] |
| 03427 | 8 | T6SS amidase immunity protein Tai4 family protein [Denitromonas halophilus] |
| 01497 | 8 | heavy-metal-associated domain-containing protein [Azoarcus communis] |
| 02164 | 8 | hypothetical protein [Denitromonas halophilus] |
| 04540 | 8 | UPF0175 family protein [Azoarcus nasutitermitis] |
| 04545 | 8 | SurA N-terminal domain-containing protein [Aromatoleum toluclasticum] |
| 08348 | 8 | SRPBCC family protein [Aromatoleum toluclasticum] |
| 08066 | 8 | DUF2231 domain-containing protein [Azoarcus indigens] |
| 07462 | 8 | MULTISPECIES: hypothetical protein [Aromatoleum] |
| 06466 | 8 | hypothetical protein [Aromatoleum evansii] |

| | | |
|---|---|---|
| **09255** | 8 | SCO family protein [Aromatoleum toluclasticum] |
| **06405** | 8 | DUF2703 domain-containing protein [Azoarcus olearius] |
| **10129** | 8 | hypothetical protein [Aromatoleum buckelii] |
| **02030** | 8 | MULTISPECIES: hypothetical protein [Proteobacteria] |
| **07486** | 8 | hypothetical protein [Azoarcus indigens] |
| **02045** | 8 | hypothetical protein [Aromatoleum toluclasticum] |
| **01577** | 8 | TRAP transporter large permease subunit, partial [Propionivibrio dicarboxylicus] |
| **03424** | 8 | acyl-CoA thioesterase [Denitromonas halophilus] |
| **00126** | 8 | DUF4124 domain-containing protein [Niveibacterium microcysteis] |
| **02056** | 8 | MULTISPECIES: hypothetical protein [Thauera] |
| **02058** | 8 | MULTISPECIES: lipid A deacylase LpxR family protein [Proteobacteria] |
| **00188** | 8 | phosphate acetyltransferase [Propionivibrio dicarboxylicus] |
| **04951** | 8 | CZB domain-containing protein [Azoarcus communis] |
| **09112** | 8 | MULTISPECIES: ethanolamine utilization microcompartment protein EutM [Thauera] |
| **04774** | 8 | DUF2059 domain-containing protein [Azoarcus indigens] |
| **07561** | 8 | ArsR family transcriptional regulator [Azoarcus communis] |
| **06997** | 8 | IS5 family transposase [Aromatoleum bremense] |
| **08823** | 8 | DUF350 domain-containing protein [Azoarcus indigens] |
| **07006** | 8 | mechanosensitive ion channel [Niveibacterium microcysteis] |
| **02638** | 8 | hypothetical protein [Azoarcus indigens] |
| **01032** | 8 | MULTISPECIES: hypothetical protein [Aromatoleum] |
| **03283** | 8 | MULTISPECIES: PRTRC system protein D [Betaproteobacteria] |
| **08028** | 8 | DUF192 domain-containing protein [Thauera terpenica] |
| **03314** | 8 | TIGR02285 family protein [Uliginosibacterium aquaticum] |
| **04130** | 8 | hypothetical protein [Azoarcus indigens] |
| **02992** | 8 | hypothetical protein [Nitrogeniibacter aestuarii] |
| **03462** | 8 | BREX-1 system adenine-specific DNA-methyltransferase PglX [Aromatoleum aromaticum] |
| **06993** | 8 | pilus assembly protein TadG-related protein [Aromatoleum toluclasticum] |
| **00243** | 8 | M14 family metallopeptidase [Aromatoleum toluolicum] |
| **02680** | 8 | DUF3696 domain-containing protein [Aromatoleum bremense] |
| **03890** | 8 | ABC transporter permease [Azoarcus nasutitermitis] |
| **05230** | 8 | response regulator [Azoarcus olearius] |
| **05817** | 8 | hypothetical protein [Aromatoleum toluclasticum] |
| **05819** | 8 | hypothetical protein [Aromatoleum diolicum] |
| **06639** | 8 | hypothetical protein [Azoarcus communis] |
| **02968** | 8 | winged helix-turn-helix domain-containing protein [Denitromonas halophilus] |
| **04092** | 8 | hypothetical protein [Aromatoleum toluclasticum] |
| **01041** | 8 | WD40 repeat domain-containing protein [Thauera humireducens] |
| **06981** | 8 | HutD family protein [Aromatoleum toluclasticum] |
| **07749** | 8 | MSMEG_0569 family flavin-dependent oxidoreductase [Azoarcus indigens] |
| **10471** | 8 | EAL domain-containing protein [Azoarcus communis] |
| **08301** | 8 | type I-F CRISPR-associated helicase Cas3f [Azoarcus halotolerans] |
| **07730** | 8 | hypothetical protein [Azoarcus indigens] |
| **00389** | 8 | phospholipase D-like domain-containing protein [Denitromonas halophilus] |
| **08898** | 8 | hypothetical protein [Aromatoleum toluclasticum] |
| **04220** | 8 | DUF5655 domain-containing protein [Azoarcus communis] |
| **06285** | 8 | M10 family metallopeptidase C-terminal domain-containing protein [Aromatoleum toluclasticum] |
| **03323** | 8 | nitroreductase family deazaflavin-dependent oxidoreductase [Aromatoleum toluclasticum] |
| **00512** | 8 | addiction module protein [Azoarcus communis] |
| **05563** | 8 | hypothetical protein [Candidatus Dactylopiibacterium carminicum] |
| **05422** | 8 | MFS transporter [Aromatoleum pumilum] |
| **00985** | 8 | TerD family protein [Aromatoleum aromaticum] |
| **02558** | 8 | ATP-binding protein [Denitromonas halophilus] |
| **08858** | 8 | MULTISPECIES: hypothetical protein [Niveibacterium] |
| **02610** | 8 | glycosyltransferase [Aromatoleum pumilum] |
| **10324** | 8 | LexA family transcriptional regulator [Azoarcus rhizosphaerae] |
| **00074** | 8 | S8 family peptidase [Azoarcus sp. Aa7] |
| **09917** | 8 | helix-turn-helix domain-containing protein [Aromatoleum pumilum] |
| **05559** | 8 | DUF1993 domain-containing protein [Niveibacterium umoris] |
| **05394** | 8 | hypothetical protein [Azoarcus olearius] |
| **01012** | 8 | hypothetical protein [Azoarcus nasutitermitis] |
| **00214** | 8 | VOC family protein [Azoarcus sp. KH32C] |
| **00500** | 8 | cytosine permease [Aromatoleum toluclasticum] |
| **03045** | 8 | DNA alkylation repair protein [Azoarcus indigens] |
| **03484** | 8 | dihydroneopterin aldolase [Aromatoleum evansii] |
| **04087** | 8 | DUF1800 domain-containing protein [Niveibacterium sp. COAC-50] |
| **10276** | 8 | cytochrome D1 [Aromatoleum toluclasticum] |
| **10581** | 8 | maltose alpha-D-glucosyltransferase [Azoarcus indigens] |
| **06943** | 8 | DUF3883 domain-containing protein [Aromatoleum bremense] |
| **06453** | 8 | sel1 repeat family protein [Azoarcus communis] |
| **02780** | 8 | DUF1214 domain-containing protein [Thauera terpenica] |
| **05837** | 8 | biotin-independent malonate decarboxylase subunit gamma [Aromatoleum pumilum] |
| **01117** | 8 | efflux RND transporter permease subunit, partial [Denitromonas halophilus] |
| **10051** | 8 | helix-turn-helix domain-containing protein [Thauera butanivorans] |
| **06596** | 8 | hypothetical protein [Denitromonas halophilus] |
| **08617** | 8 | hypothetical protein [Nitrogeniibacter aestuarii] |
| **03476** | 8 | pilus assembly protein [Aromatoleum toluclasticum] |
| **07854** | 8 | type I-F CRISPR-associated endoribonuclease Cas6/Csy4 [Azoarcus nasutitermitis] |
| **06215** | 8 | hypothetical protein [Denitromonas halophilus] |
| **05333** | 8 | arsenic resistance N-acetyltransferase ArsN2 [Nitrogeniibacter aestuarii] |
| **07777** | 8 | sulfite exporter TauE/SafE family protein [Azoarcus indigens] |
| **00477** | 8 | mitochondrial fission ELM1 family protein [Thauera phenolivorans] |
| **00232** | 8 | M23 family metallopeptidase [Aromatoleum tolulyticum] |
| **06454** | 8 | hypothetical protein [Denitromonas halophilus] |
| **08633** | 8 | Hpt domain-containing protein, partial [Azoarcus sp. TTM-91] |
| **03967** | 8 | CopG family transcriptional regulator [Aromatoleum aromaticum] |
| **07849** | 8 | DUF4351 domain-containing protein [Thauera butanivorans] |
| **10342** | 8 | hypothetical protein [Zoogloea oleivorans] |
| **00838** | 8 | cupin domain-containing protein [Aromatoleum toluclasticum] |
| **10344** | 8 | DUF4105 domain-containing protein [Azoarcus indigens] |
| **08656** | 8 | hypothetical protein [Denitromonas halophilus] |
| **10423** | 8 | DNA repair ATPase [Azoarcus communis] |
| **10594** | 8 | NUDIX hydrolase [Nitrogeniibacter aestuarii] |
| **09813** | 8 | ATP-binding protein [Aromatoleum aromaticum] |

| | | |
|---|---|---|
| 04044 | 8 | L-lactate dehydrogenase [Azoarcus indigens] |
| 07778 | 8 | PspA/IM30 family protein [Azoarcus communis] |
| 01178 | 8 | MULTISPECIES: hypothetical protein [Thauera] |
| 08743 | 8 | hypothetical protein [Azoarcus indigens] |
| 06304 | 8 | DsbA family protein [Aromatoleum toluclasticum] |
| 02954 | 8 | FHA domain-containing protein [Aromatoleum aromaticum] |
| 03919 | 8 | hypothetical protein [Azoarcus halotolerans] |
| 03672 | 8 | MULTISPECIES: pilus assembly protein TadG-related protein [Thauera] |
| 07782 | 8 | patatin-like phospholipase family protein [Azoarcus indigens] |
| 02740 | 8 | ferritin [Azoarcus olearius] |
| 05516 | 8 | chaperone NapD [Azoarcus indigens] |
| 10352 | 8 | galactosyldiacylglycerol synthase [Nitrogeniibacter aestuarii] |
| 02747 | 8 | molybdenum storage protein subunit alpha [Azoarcus nasutitermitis] |
| 01162 | 8 | MULTISPECIES: type II toxin-antitoxin system Phd/YefM family antitoxin [Thauera] |
| 01160 | 8 | DNA-binding protein [Aromatoleum aromaticum] |
| 02944 | 8 | alpha/beta fold hydrolase [Aromatoleum toluclasticum] |
| 07792 | 8 | type I-E CRISPR-associated endoribonuclease Cas2e [Aromatoleum diolicum] |
| 06834 | 8 | hypothetical protein [Aromatoleum diolicum] |
| 09809 | 8 | site-specific integrase [Denitromonas halophilus] |
| 05264 | 8 | hypothetical protein [Aromatoleum toluclasticum] |
| 04048 | 8 | TfoX/Sxy family protein [Aromatoleum toluclasticum] |
| 01056 | 8 | toxic anion resistance protein, partial [Thauera phenolivorans] |
| 02488 | 8 | TIR domain-containing protein [Denitromonas halophilus] |
| 05182 | 8 | hypothetical protein [Nitrogeniibacter aestuarii] |
| 00259 | 8 | glycosyltransferase [Aromatoleum aromaticum] |
| 06551 | 8 | type II toxin-antitoxin system RelE/ParE family toxin [Thauera terpenica] |
| 07618 | 8 | hypothetical protein [Azoarcus communis] |
| 05756 | 8 | calcium:proton antiporter [Denitromonas halophilus] |
| 00763 | 8 | hypothetical protein [Aromatoleum petrolei] |
| 03773 | 8 | DUF4337 domain-containing protein [Niveibacterium umoris] |
| 03508 | 8 | cupin domain-containing protein [Azoarcus olearius] |
| 02291 | 8 | hypothetical protein [Aromatoleum toluclasticum] |
| 06293 | 8 | prepilin-type N-terminal cleavage/methylation domain-containing protein [Azoarcus nasutitermitis] |
| 00106 | 8 | DUF309 domain-containing protein [Aromatoleum toluclasticum] |
| 07629 | 8 | S1 family peptidase [Nitrogeniibacter aestuarii] |
| 05093 | 8 | MULTISPECIES: hypothetical protein [Thauera] |
| 01344 | 8 | hypothetical protein, partial [Azoarcus nasutitermitis] |
| 05760 | 8 | type I-F CRISPR-associated protein Csy3 [Azoarcus nasutitermitis] |
| 09685 | 8 | hypothetical protein [Aromatoleum diolicum] |
| 06565 | 8 | glycine zipper 2TM domain-containing protein [Azoarcus halotolerans] |
| 01342 | 8 | alpha,alpha-trehalase TreF [Aromatoleum diolicum] |
| 07944 | 8 | PepSY domain-containing protein [Azoarcus indigens] |
| 10289 | 8 | Wzz/FepE/Etk N-terminal domain-containing protein [Aromatoleum diolicum] |
| 05059 | 8 | CoA transferase [Aromatoleum toluclasticum] |
| 03787 | 8 | EthD domain-containing protein [Aromatoleum toluclasticum] |
| 09013 | 8 | radical SAM protein [Aromatoleum toluclasticum] |
| 08417 | 8 | MULTISPECIES: hypothetical protein [Thauera] |
| 07986 | 8 | rhomboid family intramembrane serine protease [Aromatoleum aromaticum] |
| 05461 | 8 | type I-F CRISPR-associated endonuclease Cas1f [Azoarcus halotolerans] |
| 09986 | 8 | MFS transporter [Azoarcus taiwanensis] |
| 02321 | 8 | MULTISPECIES: DNA-binding protein [Thauera] |
| 00553 | 8 | FdtA/QdtA family cupin domain-containing protein [Azoarcus communis] |
| 08392 | 8 | GNAT family N-acetyltransferase [Azoarcus indigens] |
| 06165 | 8 | hypothetical protein [Azoarcus indigens] |
| 08399 | 8 | metal-dependent phosphohydrolase [Denitromonas halophilus] |
| 09981 | 8 | HipA domain-containing protein [Azoarcus communis] |
| 04368 | 8 | transporter substrate-binding domain-containing protein [Azoarcus communis] |
| 03741 | 8 | hypothetical protein [Aromatoleum toluclasticum] |
| 08383 | 8 | hypothetical protein [Aromatoleum toluolicum] |
| 08403 | 8 | replication initiation factor domain-containing protein [Azoarcus indigens] |
| 05073 | 8 | conjugal transfer protein TraG N-terminal domain-containing protein [Azoarcus indigens] |
| 03349 | 8 | formaldehyde-activating enzyme [Aromatoleum toluvorans] |
| 00109 | 8 | helix-turn-helix domain-containing protein [Aromatoleum evansii] |
| 05752 | 8 | uracil-DNA glycosylase family protein [Denitromonas halophilus] |
| 06163 | 8 | alpha/beta fold hydrolase [Azoarcus taiwanensis] |
| 08414 | 8 | hypothetical protein [Azoarcus indigens] |
| 08286 | 8 | CoA transferase [Aromatoleum toluclasticum] |
| 00560 | 8 | PAS domain S-box protein [Azoarcus indigens] |
| 03745 | 8 | glycosyltransferase [Aromatoleum aromaticum] |
| 01310 | 8 | DUF4198 domain-containing protein [Azoarcus indigens] |
| 00539 | 8 | glycine zipper 2TM domain-containing protein [Nitrogeniibacter aestuarii] |
| 06077 | 8 | tandem-95 repeat protein [Denitromonas halophilus] |
| 00566 | 8 | FecR family protein [Azoarcus indigens] |
| 06346 | 8 | lipid-binding SYLF domain-containing protein [Azoarcus sp. KH32C] |
| 07567 | 8 | 3'-5' exonuclease [Azoarcus indigens] |
| 10281 | 8 | CopD family protein [Azoarcus indigens] |
| 09735 | 8 | Abi family protein [Rhodocyclus purpureus] |
| 06231 | 8 | hypothetical protein [Aromatoleum bremense] |
| 08931 | 8 | DUF4395 family protein [Azoarcus communis] |
| 07120 | 8 | SapC family protein [Azoarcus indigens] |
| 06079 | 8 | hypothetical protein [Denitromonas halophilus] |
| 09942 | 8 | helix-turn-helix transcriptional regulator [Nitrogeniibacter aestuarii] |
| 01264 | 8 | hypothetical protein [Propionivibrio dicarboxylicus] |
| 06255 | 8 | AAA family ATPase [Azoarcus sp. Aa7] |
| 03117 | 8 | 4-oxalocrotonate tautomerase [Denitromonas halophilus] |
| 05988 | 8 | DUF4351 domain-containing protein [Rhodocyclus purpureus] |
| 05434 | 8 | CBS domain-containing protein [Nitrogeniibacter aestuarii] |
| 08040 | 8 | hypothetical protein [Denitromonas halophilus] |
| 03557 | 8 | hypothetical protein [Nitrogeniibacter aestuarii] |
| 07977 | 8 | MULTISPECIES: DUF190 domain-containing protein [Niveibacterium] |
| 08479 | 8 | Wzz/FepE/Etk N-terminal domain-containing protein [Aromatoleum aromaticum] |
| 03706 | 8 | hypothetical protein [Azoarcus indigens] |
| 07930 | 8 | succinylglutamate desuccinylase/aspartoacylase family protein [Denitromonas halophilus] |
| 03269 | 8 | MaoC family dehydratase N-terminal domain-containing protein [Aromatoleum toluclasticum] |

| | | |
|---|---|---|
| 08966 | 8 | DUF5602 domain-containing protein [Aromatoleum toluclasticum] |
| 09711 | 8 | PEP-CTERM sorting domain-containing protein [Zoogloea dura] |
| 00411 | 8 | DnaJ domain-containing protein [Thauera phenylacetica] |
| 05494 | 8 | GtrA family protein [Aromatoleum toluvorans] |
| 09713 | 8 | DUF4332 domain-containing protein [Niveibacterium microcysteis] |
| 00947 | 8 | hypothetical protein [Aromatoleum toluclasticum] |
| 09717 | 8 | HEAT repeat domain-containing protein [Azoarcus indigens] |
| 07107 | 8 | XRE family transcriptional regulator [Denitromonas halophilus] |
| 09092 | 8 | flippase [Azoarcus communis] |
| 05050 | 8 | nuclear transport factor 2 family protein [Aromatoleum toluclasticum] |
| 06791 | 8 | DUF3616 domain-containing protein [Aromatoleum aromaticum] |
| 07974 | 8 | MULTISPECIES: hypothetical protein [Niveibacterium] |
| 08961 | 8 | hypothetical protein [Denitromonas halophilus] |
| 08960 | 8 | hypothetical protein [Aromatoleum bremense] |
| 06799 | 8 | hypothetical protein [Azoarcus indigens] |
| 06348 | 8 | hypothetical protein [Aromatoleum aromaticum] |
| 06482 | 8 | hypothetical protein [Nitrogeniibacter aestuarii] |
| 05339 | 7 | serine/threonine protein kinase [Aromatoleum aromaticum] |
| 09878 | 7 | TonB family protein [Propionivibrio dicarboxylicus] |
| 01061 | 7 | sodium ion-translocating decarboxylase subunit beta [Azoarcus communis] |
| 05319 | 7 | phage baseplate assembly protein V [Uliginosibacterium gangwonense] |
| 09997 | 7 | hypothetical protein [Aromatoleum toluclasticum] |
| 05488 | 7 | C45 family autoproteolytic acyltransferase/hydolase [Denitromonas halophilus] |
| 00234 | 7 | MULTISPECIES: EthD domain-containing protein [Rhodocyclales] |
| 00697 | 7 | hydrogenase maturation protease [Nitrogeniibacter aestuarii] |
| 09467 | 7 | DUF1826 domain-containing protein [Azoarcus communis] |
| 01784 | 7 | AAA family ATPase [Aromatoleum toluclasticum] |
| 06844 | 7 | cytochrome c [Nitrogeniibacter aestuarii] |
| 01013 | 7 | DUF1116 domain-containing protein [Azoarcus indigens] |
| 06843 | 7 | DUF1446 domain-containing protein [Aromatoleum toluclasticum] |
| 09457 | 7 | ABC transporter permease [Azoarcus taiwanensis] |
| 09456 | 7 | hypothetical protein [Denitromonas sp. IR12] |
| 09951 | 7 | hypothetical protein [Denitromonas halophilus] |
| 09925 | 7 | hypothetical protein [Azoarcus communis] |
| 01053 | 7 | Coenzyme F420 hydrogenase/dehydrogenase, beta subunit C-terminal domain [Denitromonas halophilus] |
| 09446 | 7 | arsenate reductase (azurin) small subunit [Aromatoleum pumilum] |
| 06935 | 7 | DUF1153 domain-containing protein [Azoarcus halotolerans] |
| 00962 | 7 | bifunctional proline dehydrogenase/L-glutamate gamma-semialdehyde dehydrogenase PutA [Azoarcus indigens] |
| 00252 | 7 | DUF1376 domain-containing protein [Aromatoleum aromaticum] |
| 00971 | 7 | DUF4332 domain-containing protein [Aromatoleum aromaticum] |
| 08184 | 7 | hypothetical protein [Thauera humireducens] |
| 01018 | 7 | antitoxin MazE family protein [Aromatoleum aromaticum] |
| 08190 | 7 | hypothetical protein [Azoarcus indigens] |
| 05380 | 7 | cytochrome oxidase small assembly protein [Aromatoleum toluclasticum] |
| 00345 | 7 | DUF2170 family protein [Azoarcus communis] |
| 09891 | 7 | MULTISPECIES: entry exclusion lipoprotein TrbK [Proteobacteria] |
| 09460 | 7 | hypothetical protein [Azoarcus indigens] |
| 05513 | 7 | PEP-CTERM sorting domain-containing protein [Denitromonas halophilus] |
| 04798 | 7 | DUF2138 family protein [Azoarcus indigens] |
| 00856 | 7 | GGDEF domain-containing protein [Azoarcus olearius] |
| 01039 | 7 | HAD hydrolase-like protein [Aromatoleum aromaticum] |
| 00936 | 7 | GAF domain-containing protein [Denitromonas halophilus] |
| 08191 | 7 | hypothetical protein [Aromatoleum toluclasticum] |
| 05362 | 7 | GNAT family N-acetyltransferase [Aromatoleum diolicum] |
| 05460 | 7 | hypothetical protein [Azoarcus indigens] |
| 00932 | 7 | M35 family metallo-endopeptidase [Rhodocyclus purpureus] |
| 06909 | 7 | type I-F CRISPR-associated protein Csy2 [Azoarcus nasutitermitis] |
| 09466 | 7 | MULTISPECIES: hypothetical protein [Proteobacteria] |
| 06929 | 7 | hypothetical protein [Denitromonas halophilus] |
| 10061 | 7 | DUF2007 domain-containing protein [Denitromonas halophilus] |
| 00231 | 7 | hypothetical protein [Aromatoleum toluclasticum] |
| 06805 | 7 | DUF2442 domain-containing protein [Zoogloea ramigera] |
| 05573 | 7 | B12-binding domain-containing radical SAM protein [Niveibacterium sp. COAC-50] |
| 07167 | 7 | low molecular weight phosphatase family protein [Azoarcus indigens] |
| 04981 | 7 | EF-hand domain-containing protein [Nitrogeniibacter aestuarii] |
| 06070 | 7 | hypothetical protein [Denitromonas halophilus] |
| 03548 | 7 | hypothetical protein [Niveibacterium microcysteis] |
| 05133 | 7 | cupin domain-containing protein [Azoarcus indigens] |
| 01285 | 7 | hypothetical protein [Denitromonas halophilus] |
| 08211 | 7 | DNA topoisomerase IB [Azoarcus indigens] |
| 01275 | 7 | DUF2853 family protein [Denitromonas halophilus] |
| 09579 | 7 | hypothetical protein [Nitrogeniibacter aestuarii] |
| 06425 | 7 | DUF2938 domain-containing protein [Denitromonas halophilus] |
| 07043 | 7 | ATPase [Nitrogeniibacter mangrovi] |
| 07068 | 7 | hypothetical protein [Aromatoleum toluclasticum] |
| 07330 | 7 | hypothetical protein [Denitromonas halophilus] |
| 05154 | 7 | MULTISPECIES: hypothetical protein [Azoarcus] |
| 03446 | 7 | type IV toxin-antitoxin system AbiEi family antitoxin domain-containing protein [Candidatus Dactylopiibacterium carminicum] |
| 09578 | 7 | right-handed parallel beta-helix repeat-containing protein [Uliginosibacterium gangwonense] |
| 06427 | 7 | hypothetical protein [Azoarcus indigens] |
| 00734 | 7 | GGDEF domain-containing protein [Azoarcus sp. KH32C] |
| 04957 | 7 | MULTISPECIES: sodium/glutamate symporter [Proteobacteria] |
| 06016 | 7 | hypothetical protein [Aromatoleum toluclasticum] |
| 10084 | 7 | hypothetical protein [Aromatoleum evansii] |
| 00207 | 7 | cephalosporin hydroxylase family protein [Aromatoleum aromaticum] |
| 03426 | 7 | DUF1304 domain-containing protein [Denitromonas halophilus] |
| 03436 | 7 | S1/P1 nuclease [Azospira oryzae] |
| 01316 | 7 | hypothetical protein [Nitrogeniibacter aestuarii] |
| 04858 | 7 | DUF4917 family protein [Azoarcus communis] |
| 01391 | 7 | DUF547 domain-containing protein [Denitromonas halophilus] |
| 06413 | 7 | hypothetical protein [Denitromonas halophilus] |
| 05039 | 7 | DUF2300 domain-containing protein [Azoarcus indigens] |
| 10417 | 7 | hypothetical protein [Denitromonas halophilus] |
| 07131 | 7 | DUF192 domain-containing protein [Azoarcus nasutitermitis] |

| | | |
|---|---|---|
| 01414 | 7 | ATP-binding cassette domain-containing protein, partial [Aromatoleum evansii] |
| 09661 | 7 | type I restriction enzyme HsdR N-terminal domain-containing protein [Denitromonas halophilus] |
| 01407 | 7 | MULTISPECIES: hypothetical protein [Betaproteobacteria] |
| 05061 | 7 | hypothetical protein [Azoarcus nasutitermitis] |
| 09640 | 7 | gamma-glutamylcyclotransferase [Aromatoleum toluclasticum] |
| 01400 | 7 | hypothetical protein [Nitrogeniibacter aestuarii] |
| 05065 | 7 | aspartate kinase [Nitrogeniibacter aestuarii] |
| 06514 | 7 | hypothetical protein [Azoarcus communis] |
| 00008 | 7 | BREX-1 system adenine-specific DNA-methyltransferase PglX [Azoarcus communis] |
| 09667 | 7 | DUF2778 domain-containing protein [Aromatoleum toluclasticum] |
| 09623 | 7 | Na(+)/H(+) antiporter subunit B [Denitromonas halophilus] |
| 06076 | 7 | GYD domain-containing protein [Denitromonas halophilus] |
| 01369 | 7 | DUF1289 domain-containing protein [Denitromonas halophilus] |
| 07145 | 7 | TonB-system energizer ExbB [Azoarcus indigens] |
| 06789 | 7 | TniQ family protein [Aromatoleum tolulyticum] |
| 03431 | 7 | type VI secretion system amidase effector protein Tae4 [Denitromonas halophilus] |
| 05081 | 7 | alpha/beta fold hydrolase [Azoarcus indigens] |
| 01353 | 7 | DUF4124 domain-containing protein [Nitrogeniibacter aestuarii] |
| 01349 | 7 | hypothetical protein [Azoarcus rhizosphaerae] |
| 07157 | 7 | glycoside hydrolase [Azoarcus communis] |
| 05184 | 7 | MULTISPECIES: phosphoadenosine phosphosulfate reductase family protein [Rhodocyclales] |
| 03457 | 7 | TRAP transporter small permease subunit [Aromatoleum toluclasticum] |
| 09772 | 7 | hypothetical protein [Denitromonas halophilus] |
| 06955 | 7 | hypothetical protein [Denitromonas halophilus] |
| 05972 | 7 | cellulase family glycosylhydrolase [Niveibacterium microcysteis] |
| 05255 | 7 | MULTISPECIES: (d)CMP kinase [Niveibacterium] |
| 04891 | 7 | DUF1501 domain-containing protein [Azoarcus olearius] |
| 09509 | 7 | glycosyltransferase family 4 protein [Azoarcus olearius] |
| 05623 | 7 | TetR/AcrR family transcriptional regulator [Denitromonas halophilus] |
| 05625 | 7 | N-acetylmuramidase domain-containing protein [Azoarcus sp. Aa7] |
| 06760 | 7 | cupin domain-containing protein [Aromatoleum petrolei] |
| 05265 | 7 | DUF1819 family protein [Aromatoleum aromaticum] |
| 01137 | 7 | DUF3445 domain-containing protein [Azoarcus communis] |
| 00170 | 7 | MULTISPECIES: hypothetical protein [Aromatoleum] |
| 05276 | 7 | bifunctional lysylphosphatidylglycerol flippase/synthetase MprF [Azoarcus indigens] |
| 01122 | 7 | adenosylcobinamide amidohydrolase [Denitromonas halophilus] |
| 05192 | 7 | hypothetical protein [Zoogloea dura] |
| 07258 | 7 | DUF3592 domain-containing protein [Nitrogeniibacter aestuarii] |
| 05537 | 7 | PQQ-binding-like beta-propeller repeat protein [Aromatoleum diolicum] |
| 00169 | 7 | universal stress protein [Aromatoleum toluclasticum] |
| 06945 | 7 | DUF1549 and DUF1553 domain-containing protein [Azoarcus olearius] |
| 00825 | 7 | hypothetical protein [Nitrogeniibacter aestuarii] |
| 04864 | 7 | NAD(P)H-dependent oxidoreductase [Azoarcus olearius] |
| 01105 | 7 | hypothetical protein [Azoarcus indigens] |
| 10626 | 7 | hypothetical protein [Aromatoleum evansii] |
| 09488 | 7 | DUF1018 domain-containing protein [Aromatoleum evansii] |
| 09842 | 7 | MULTISPECIES: hypothetical protein [Rhodocyclales] |
| 06020 | 7 | type I-F CRISPR-associated protein Csy1 [Azoarcus nasutitermitis] |
| 01651 | 7 | hypothetical protein [Azoarcus halotolerans] |
| 05251 | 7 | GIY-YIG nuclease family protein [Aromatoleum evansii] |
| 10060 | 7 | DUF1499 domain-containing protein [Denitromonas halophilus] |
| 03578 | 7 | APC family permease [Azoarcus communis] |
| 03459 | 7 | hypothetical protein [Azoarcus communis] |
| 10081 | 7 | hypothetical protein [Azoarcus olearius] |
| 01218 | 7 | hypothetical protein [Azoarcus indigens] |
| 05202 | 7 | hypothetical protein [Denitromonas halophilus] |
| 09783 | 7 | hypothetical protein [Nitrogeniibacter aestuarii] |
| 08113 | 7 | glycosyltransferase family 4 protein [Nitrogeniibacter aestuarii] |
| 05205 | 7 | flagellar protein FliT [Denitromonas halophilus] |
| 05206 | 7 | GNAT family N-acetyltransferase [Denitromonas halophilus] |
| 01584 | 7 | hypothetical protein [Azoarcus sp. KH32C] |
| 05207 | 7 | hypothetical protein [Denitromonas sp. IR12] |
| 06428 | 7 | BREX-1 system phosphatase PglZ type A [Aromatoleum aromaticum] |
| 01591 | 7 | MULTISPECIES: hypothetical protein [Aromatoleum] |
| 09553 | 7 | HlyD family secretion protein [Azoarcus indigens] |
| 01595 | 7 | PepSY domain-containing protein [Azoarcus communis] |
| 07002 | 7 | hypothetical protein [Azoarcus communis] |
| 07001 | 7 | FecR domain-containing protein [Nitrogeniibacter aestuarii] |
| 08199 | 7 | hypothetical protein [Azoarcus nasutitermitis] |
| 01607 | 7 | TIR domain-containing protein [Aromatoleum diolicum] |
| 00220 | 7 | MULTISPECIES: capsular polysaccharide biosynthesis protein CapF [Thauera] |
| 01619 | 7 | hypothetical protein [Niveibacterium microcysteis] |
| 01190 | 7 | hypothetical protein [Aromatoleum toluclasticum] |
| 03528 | 7 | hypothetical protein [Aromatoleum toluclasticum] |
| 01183 | 7 | MULTISPECIES: MoaD/ThiS family protein [Aromatoleum] |
| 00033 | 7 | DUF3369 domain-containing protein [Thauera hydrothermalis] |
| 10002 | 7 | hypothetical protein [Rhodocyclus purpureus] |
| 02902 | 7 | carboxymuconolactone decarboxylase family protein [Aromatoleum diolicum] |
| 06131 | 7 | HPr-rel-A system PqqD family peptide chaperone [Aromatoleum petrolei] |
| 09245 | 7 | hypothetical protein [Aromatoleum diolicum] |
| 06373 | 7 | MULTISPECIES: hypothetical protein [Thauera] |
| 02657 | 7 | hypothetical protein [Azoarcus communis] |
| 04338 | 7 | hypothetical protein [Azoarcus indigens] |
| 02038 | 7 | AAA family ATPase [Thauera linaloolentis] |
| 07617 | 7 | AAA family ATPase [Azoarcus communis] |
| 09239 | 7 | hypothetical protein [Aromatoleum toluclasticum] |
| 02941 | 7 | alpha/beta fold hydrolase [Azoarcus indigens] |
| 00313 | 7 | SWIM zinc finger family protein [Azoarcus sp. KH32C] |
| 02350 | 7 | ribosomal protein L7/L12 [Denitromonas halophilus] |
| 07859 | 7 | Fic family protein [Azoarcus rhizosphaerae] |
| 07493 | 7 | hypothetical protein [Niveibacterium microcysteis] |
| 08793 | 7 | heparan-alpha-glucosaminide N-acetyltransferase domain-containing protein [Azoarcus indigens] |
| 04351 | 7 | hypothetical protein [Azoarcus communis] |
| 06694 | 7 | LysM peptidoglycan-binding domain-containing protein [Azoarcus olearius] |

| | | |
|---|---|---|
| **03309** | 7 | DUF29 domain-containing protein [Aromatoleum diolicum] |
| **08067** | 7 | DUF5681 domain-containing protein [Aromatoleum aromaticum] |
| **00607** | 7 | DEAD/DEAH box helicase family protein [Aromatoleum bremense] |
| **08622** | 7 | DMT family transporter, partial [Azoarcus taiwanensis] |
| **03344** | 7 | GNAT family N-acetyltransferase, partial [Thauera phenolivorans] |
| **02332** | 7 | hypothetical protein [Zoogloea oleivorans] |
| **06317** | 7 | hypothetical protein [Azoarcus indigens] |
| **04366** | 7 | stage II sporulation protein M [Candidatus Dactylopiibacterium carminicum] |
| **02704** | 7 | IS630 family transposase, partial [Azoarcus indigens] |
| **04097** | 7 | hypothetical protein [Aromatoleum aromaticum] |
| **07870** | 7 | hypothetical protein [Zoogloea dura] |
| **00614** | 7 | DUF4157 domain-containing protein [Azoarcus sp. DD4] |
| **02024** | 7 | hypothetical protein [Niveibacterium umoris] |
| **00623** | 7 | substrate-binding domain-containing protein [Denitromonas halophilus] |
| **02394** | 7 | hypothetical protein [Azoarcus indigens] |
| **08532** | 7 | M48 family metallopeptidase [Denitromonas halophilus] |
| **08539** | 7 | DUF4040 domain-containing protein [Denitromonas halophilus] |
| **08544** | 7 | hypothetical protein [Aromatoleum toluclasticum] |
| **04157** | 7 | DUF5110 domain-containing protein [Niveibacterium sp. COAC-50] |
| **04617** | 7 | glycosyltransferase [Azoarcus olearius] |
| **09276** | 7 | hypothetical protein [Propionivibrio dicarboxylicus] |
| **03020** | 7 | glycine reductase [Azoarcus indigens] |
| **02387** | 7 | hypothetical protein [Aromatoleum toluclasticum] |
| **03877** | 7 | hypothetical protein [Azoarcus indigens] |
| **07456** | 7 | AAA family ATPase [Azoarcus nasutitermitis] |
| **08552** | 7 | flagellar basal body protein FlgE [Azoarcus indigens] |
| **07631** | 7 | OPT/YSL family transporter [Nitrogeniibacter aestuarii] |
| **06328** | 7 | helix-turn-helix transcriptional regulator [Azoarcus indigens] |
| **08567** | 7 | DUF3568 family protein [Candidatus Dactylopiibacterium carminicum] |
| **04604** | 7 | YjiH family protein [Aromatoleum evansii] |
| **00069** | 7 | FHA domain-containing protein [Azoarcus olearius] |
| **00133** | 7 | transposase [Azoarcus sp. DD4] |
| **04129** | 7 | DUF2892 domain-containing protein [Azoarcus indigens] |
| **07469** | 7 | DUF2309 domain-containing protein [Azoarcus communis] |
| **06170** | 7 | MULTISPECIES: GNAT family N-acetyltransferase [Bacteria] |
| **05812** | 7 | N-6 DNA methylase [Aromatoleum bremense] |
| **09006** | 7 | leukotoxin LktA family filamentous adhesin [Denitromonas halophilus] |
| **09251** | 7 | NUDIX domain-containing protein [Nitrogeniibacter mangrovi] |
| **10447** | 7 | hypothetical protein [Nitrogeniibacter aestuarii] |
| **05745** | 7 | BREX system serine/threonine kinase PglW [Azoarcus indigens] |
| **03958** | 7 | YfiR family protein [Nitrogeniibacter aestuarii] |
| **04038** | 7 | hypothetical protein [Aromatoleum evansii] |
| **02820** | 7 | conjugal transfer transcriptional regulator TraJ [Thauera phenolivorans] |
| **08318** | 7 | DUF3024 domain-containing protein [Aromatoleum tolulyticum] |
| **08317** | 7 | YCF48-related protein [Uliginosibacterium aquaticum] |
| **00485** | 7 | hypothetical protein [Nitrogeniibacter aestuarii] |
| **04479** | 7 | helix-turn-helix transcriptional regulator [Zoogloea oleivorans] |
| **08702** | 7 | SEC-C domain-containing protein [Aromatoleum aromaticum] |
| **00316** | 7 | AAA family ATPase [Aromatoleum bremense] |
| **09180** | 7 | hypothetical protein [Azoarcus olearius] |
| **02261** | 7 | DUF6484 domain-containing protein [Azoarcus communis] |
| **07780** | 7 | glycosyltransferase family 4 protein [Azoarcus sp. KH32C] |
| **08741** | 7 | hypothetical protein [Azoarcus nasutitermitis] |
| **02758** | 7 | MULTISPECIES: RDD family protein [Niveibacterium] |
| **04015** | 7 | hypothetical protein [Aromatoleum toluolicum] |
| **03636** | 7 | polysaccharide deacetylase family protein [Azoarcus indigens] |
| **09140** | 7 | Na+/H+ antiporter subunit E [Denitromonas halophilus] |
| **08714** | 7 | glycoside hydrolase family 3 C-terminal domain-containing protein [Uliginosibacterium gangwonense] |
| **08716** | 7 | cytochrome c biogenesis CcdA family protein [Propionivibrio dicarboxylicus] |
| **05727** | 7 | head decoration protein [Thauera butanivorans] |
| **04444** | 7 | cupin domain-containing protein [Aromatoleum diolicum] |
| **04421** | 7 | amidinotransferase [Denitromonas halophilus] |
| **09126** | 7 | DUF2399 domain-containing protein [Azoarcus rhizosphaerae] |
| **08727** | 7 | MULTISPECIES: MFS transporter [Thauera] |
| **06529** | 7 | methyltransferase domain-containing protein [Denitromonas halophilus] |
| **08722** | 7 | hypothetical protein [Aromatoleum toluclasticum] |
| **00112** | 7 | hypothetical protein [Aromatoleum aromaticum] |
| **08684** | 7 | DUF2474 domain-containing protein [Azoarcus sp. DD4] |
| **08064** | 7 | Imm71 family immunity protein [Aromatoleum tolulyticum] |
| **02843** | 7 | SLC13 family permease [Azoarcus taiwanensis] |
| **08038** | 7 | hypothetical protein [Nitrogeniibacter aestuarii] |
| **09206** | 7 | gamma-glutamylcyclotransferase [Nitrogeniibacter aestuarii] |
| **03353** | 7 | MULTISPECIES: DUF1173 domain-containing protein [Betaproteobacteria] |
| **02897** | 7 | AAA family ATPase [Aromatoleum aromaticum] |
| **07516** | 7 | multicopper oxidase domain-containing protein [Denitromonas halophilus] |
| **08648** | 7 | ABC transporter substrate-binding protein [Niveibacterium microcysteis] |
| **00376** | 7 | hypothetical protein [Azoarcus indigens] |
| **09203** | 7 | hypothetical protein [Aromatoleum toluclasticum] |
| **08757** | 7 | hypothetical protein [Aromatoleum toluclasticum] |
| **03979** | 7 | regulatory protein GemA [Azoarcus communis] |
| **06635** | 7 | PEP-CTERM sorting domain-containing protein [Azoarcus olearius] |
| **05844** | 7 | 4-oxalocrotonate tautomerase [Aromatoleum pumilum] |
| **09094** | 7 | hypothetical protein [Azoarcus communis] |
| **10579** | 7 | MULTISPECIES: hypothetical protein [Thauera] |
| **02281** | 7 | hypothetical protein [Aromatoleum evansii] |
| **07776** | 7 | AAA family ATPase [Aromatoleum aromaticum] |
| **02837** | 7 | MULTISPECIES: universal stress protein [Thauera] |
| **04001** | 7 | GTP-binding protein [Aromatoleum diolicum] |
| **02275** | 7 | DUF4160 domain-containing protein [Aromatoleum evansii] |
| **02830** | 7 | type IV pili methyl-accepting chemotaxis transducer N-terminal domain-containing protein [Thauera phenylacetica] |
| **04492** | 7 | cytochrome C oxidase subunit IV family protein [Aromatoleum toluclasticum] |
| **07527** | 7 | site-specific DNA-methyltransferase [Aromatoleum bremense] |
| **08682** | 7 | AAA family ATPase [Aromatoleum aromaticum] |
| **08267** | 7 | hypothetical protein [Denitromonas halophilus] |

| | | |
|---|---|---|
| **00628** | 7 | hypothetical protein [Denitromonas halophilus] |
| **06157** | 7 | pectate lyase [Uliginosibacterium aquaticum] |
| **10561** | 7 | hypothetical protein [Denitromonas halophilus] |
| **02466** | 7 | type II toxin-antitoxin system HicB family antitoxin [Thauera aromatica] |
| **02462** | 7 | hypothetical protein [Niveibacterium umoris] |
| **06558** | 7 | LuxR C-terminal-related transcriptional regulator [Aromatoleum toluclasticum] |
| **04705** | 7 | GNAT family N-acetyltransferase [Zoogloea oleivorans] |
| **09364** | 7 | calcium-binding protein [Azoarcus communis] |
| **06381** | 7 | 4-hydroxyphenylacetate 3-monooxygenase [Azoarcus indigens] |
| **07942** | 7 | DUF4440 domain-containing protein, partial [Azoarcus sp. TTM-91] |
| **07385** | 7 | hypothetical protein [Denitromonas halophilus] |
| **03165** | 7 | DUF2971 domain-containing protein [Azoarcus indigens] |
| **05979** | 7 | hypothetical protein [Aromatoleum toluclasticum] |
| **05893** | 7 | hypothetical protein [Denitromonas halophilus] |
| **10377** | 7 | MULTISPECIES: hypothetical protein [Thauera] |
| **07714** | 7 | iron transporter [Aromatoleum toluclasticum] |
| **10448** | 7 | SIR2 family protein [Denitromonas halophilus] |
| **07389** | 7 | SLC13 family permease, partial [Thauera phenylacetica] |
| **07392** | 7 | IS630 family transposase [Thauera phenylacetica] |
| **08083** | 7 | MULTISPECIES: ATP-binding protein [Rhodocyclales] |
| **08456** | 7 | tail protein X [Uliginosibacterium gangwonense] |
| **08982** | 7 | DUF2867 domain-containing protein [Azoarcus olearius] |
| **07673** | 7 | MTH1187 family thiamine-binding protein [Aromatoleum pumilum] |
| **06522** | 7 | patatin-like phospholipase family protein [Niveibacterium sp. COAC-50] |
| **00639** | 7 | TraI domain-containing protein [Azoarcus indigens] |
| **03814** | 7 | thiol reductant ABC exporter subunit CydD [Azoarcus indigens] |
| **01912** | 7 | fructuronate reductase [Candidatus Dactylopiibacterium carminicum] |
| **08472** | 7 | phage tail tape measure protein [Azoarcus indigens] |
| **00077** | 7 | SHOCT domain-containing protein [Nitrogeniibacter aestuarii] |
| **03774** | 7 | ATP-grasp domain-containing protein [Azoarcus olearius] |
| **01875** | 7 | DUF3300 domain-containing protein [Azoarcus indigens] |
| **09349** | 7 | DUF2778 domain-containing protein [Nitrogeniibacter aestuarii] |
| **07981** | 7 | DUF4350 domain-containing protein [Candidatus Dactylopiibacterium carminicum] |
| **06489** | 7 | helix-turn-helix domain-containing protein [Aromatoleum tolulyticum] |
| **03731** | 7 | malto-oligosyltrehalose synthase [Azoarcus olearius] |
| **02477** | 7 | hypothetical protein [Denitromonas halophilus] |
| **04766** | 7 | terminase ATPase subunit family protein [Uliginosibacterium gangwonense] |
| **03414** | 7 | hypothetical protein [Aromatoleum aromaticum] |
| **03242** | 7 | hypothetical protein [Azoarcus olearius] |
| **07334** | 7 | hypothetical protein [Denitromonas halophilus] |
| **06395** | 7 | ethanolamine ammonia-lyase reactivating factor EutA [Aromatoleum toluclasticum] |
| **01821** | 7 | phosphomethylpyrimidine synthase ThiC, partial [Denitromonas halophilus] |
| **03232** | 7 | hypothetical protein [Azoarcus indigens] |
| **08387** | 7 | toluene-4-monooxygenase system B family protein [Aromatoleum toluclasticum] |
| **03744** | 7 | DUF1778 domain-containing protein [Aromatoleum aromaticum] |
| **03271** | 7 | MMPL family transporter, partial [Aromatoleum toluclasticum] |
| **07710** | 7 | Mu transposase C-terminal domain-containing protein [Aromatoleum pumilum] |
| **08902** | 7 | class I SAM-dependent methyltransferase [Azoarcus indigens] |
| **08402** | 7 | 2-hydroxyacyl-CoA dehydratase family protein [Aromatoleum toluclasticum] |
| **02515** | 7 | Txe/YoeB family addiction module toxin [Aromatoleum aromaticum] |
| **04742** | 7 | PH domain-containing protein [Aromatoleum aromaticum] |
| **01857** | 7 | DUF47 family protein [Aromatoleum bremense] |
| **00413** | 7 | hypothetical protein [Denitromonas halophilus] |
| **00658** | 7 | D-serine ammonia-lyase [Aromatoleum toluclasticum] |
| **00151** | 7 | DUF748 domain-containing protein [Dechlorosoma suillum] |
| **09389** | 7 | DUF1788 domain-containing protein [Aromatoleum aromaticum] |
| **04726** | 7 | MULTISPECIES: hypothetical protein [Azoarcus] |
| **03410** | 7 | PIN domain-containing protein [Achromobacter xylosoxidans] |
| **00432** | 7 | Ig-like domain-containing protein [Aromatoleum toluclasticum] |
| **04264** | 7 | urea carboxylase [Azoarcus communis] |
| **07418** | 7 | integrase [Denitromonas halophilus] |
| **03317** | 7 | DUF4129 domain-containing protein [Candidatus Dactylopiibacterium carminicum] |
| **10328** | 7 | hypothetical protein [Aromatoleum toluclasticum] |
| **03079** | 7 | hypothetical protein [Denitromonas halophilus] |
| **07912** | 7 | MULTISPECIES: replication protein RepA [Proteobacteria] |
| **04629** | 7 | MULTISPECIES: menaquinone-dependent protoporphyrinogen IX dehydrogenase [Thauera] |
| **09429** | 7 | hypothetical protein [Azoarcus indigens] |
| **00632** | 7 | sensory rhodopsin transducer [Azoarcus olearius] |
| **03842** | 7 | ankyrin repeat domain-containing protein [Aromatoleum diolicum] |
| **10301** | 7 | ABC transporter substrate-binding protein [Aromatoleum aromaticum] |
| **05998** | 7 | hypothetical protein [Azoarcus sp. KH32C] |
| **00527** | 7 | hypothetical protein [Niveibacterium sp. COAC-50] |
| **02598** | 7 | murein L,D-transpeptidase catalytic domain family protein [Azoarcus communis] |
| **03703** | 7 | YebB family permuted papain-like enzyme [Candidatus Dactylopiibacterium carminicum] |
| **09331** | 7 | GspH/FimT family pseudopilin [Aromatoleum bremense] |
| **06679** | 7 | DEAD/DEAH box helicase family protein [Zoogloea oleivorans] |
| **02424** | 7 | NirD/YgiW/YdeI family stress tolerance protein [Azoarcus indigens] |
| **02417** | 7 | hypothetical protein [Niveibacterium umoris] |
| **04661** | 7 | DUF6290 family protein [Thauera terpenica] |
| **03101** | 7 | PIN domain-containing protein [Aromatoleum bremense] |
| **07414** | 7 | DUF1175 domain-containing protein [Azoarcus indigens] |
| **03407** | 7 | hypothetical protein [Azoarcus rhizosphaerae] |
| **08491** | 7 | DNA-binding protein [Aromatoleum buckelii] |
| **08976** | 7 | RNA-binding protein [Denitromonas halophilus] |
| **03832** | 7 | radical SAM protein [Nitrogeniibacter aestuarii] |
| **07433** | 7 | AAA family ATPase [Azoarcus communis] |
| **03110** | 7 | orotidine-5'-phosphate decarboxylase [Denitromonas halophilus] |
| **03057** | 7 | aldose 1-epimerase [Uliginosibacterium gangwonense] |
| **08033** | 7 | EF-hand domain-containing protein [Denitromonas halophilus] |
| **08978** | 7 | hypothetical protein [Denitromonas halophilus] |
| **03076** | 7 | M13 family metallopeptidase [Niveibacterium microcysteis] |
| **08720** | 6 | MarR family transcriptional regulator [Aromatoleum toluclasticum] |
| **04162** | 6 | glycoside hydrolase family 16 protein [Niveibacterium umoris] |
| **06430** | 6 | hypothetical protein [Azoarcus communis] |

| | | |
|---|---|---|
| 09799 | 6 | Gp49 family protein [Thauera humireducens] |
| 00071 | 6 | DUF3486 family protein [Azoarcus communis] |
| 07737 | 6 | DUF932 domain-containing protein [Azoarcus indigens] |
| 03455 | 6 | aspartate aminotransferase family protein [Denitromonas halophilus] |
| 07738 | 6 | MULTISPECIES: relaxase/mobilization nuclease domain-containing protein [Proteobacteria] |
| 02675 | 6 | MULTISPECIES: DUF87 domain-containing protein [Rhodocyclales] |
| 02742 | 6 | hypothetical protein [Azoarcus communis] |
| 08738 | 6 | carboxynorspermidine decarboxylase, partial [Thauera phenolivorans] |
| 02749 | 6 | hypothetical protein [Azoarcus olearius] |
| 07739 | 6 | DDE-type integrase/transposase/recombinase [Azoarcus communis] |
| 06971 | 6 | hypothetical protein [Aromatoleum toluclasticum] |
| 04154 | 6 | hypothetical protein [Candidatus Dactylopiibacterium carminicum] |
| 02759 | 6 | endo-1,4-beta-xylanase [Uliginosibacterium gangwonense] |
| 09780 | 6 | DUF302 domain-containing protein [Denitromonas halophilus] |
| 00783 | 6 | fatty acid desaturase [Nitrogeniibacter aestuarii] |
| 02511 | 6 | MULTISPECIES: hypothetical protein [Thauera] |
| 01156 | 6 | ATP-grasp domain-containing protein [Denitromonas halophilus] |
| 05790 | 6 | hypothetical protein [Azoarcus sp. Aa7] |
| 06063 | 6 | PelD GGDEF domain-containing protein [Denitromonas halophilus] |
| 06210 | 6 | alkane 1-monooxygenase [Aromatoleum petrolei] |
| 07745 | 6 | ferrous iron transport protein A [Azoarcus rhizosphaerae] |
| 02530 | 6 | radical SAM protein [Aromatoleum aromaticum] |
| 05172 | 6 | penicillin-binding protein activator LpoB [Denitromonas halophilus] |
| 10511 | 6 | MULTISPECIES: trehalose-phosphatase [Niveibacterium] |
| 06189 | 6 | diguanylate cyclase [Niveibacterium microcysteis] |
| 05974 | 6 | MULTISPECIES: putative motility protein [Niveibacterium] |
| 04115 | 6 | glycosyltransferase [Propionivibrio dicarboxylicus] |
| 06251 | 6 | hypothetical protein [Nitrogeniibacter aestuarii] |
| 05187 | 6 | glycosyltransferase [Azoarcus taiwanensis] |
| 06809 | 6 | type II toxin-antitoxin system HicA family toxin [Thauera phenylacetica] |
| 04193 | 6 | GNAT family N-acetyltransferase [Azoarcus olearius] |
| 10578 | 6 | PilX N-terminal domain-containing pilus assembly protein [Aromatoleum bremense] |
| 02708 | 6 | hypothetical protein [Azoarcus communis] |
| 04121 | 6 | transposase, partial [Aromatoleum tolulyticum] |
| 04096 | 6 | hypothetical protein [Azoarcus communis] |
| 08764 | 6 | contractile injection system protein, VgrG/Pvc8 family [Uliginosibacterium gangwonense] |
| 10575 | 6 | metal-sensing transcriptional repressor [Aromatoleum diolicum] |
| 05222 | 6 | MULTISPECIES: ABC transporter permease [Zoogloeaceae] |
| 02659 | 6 | glycosyltransferase family 29 protein [Azoarcus communis] |
| 00224 | 6 | cupin domain-containing protein [Nitrogeniibacter aestuarii] |
| 03529 | 6 | SPOR domain-containing protein [Nitrogeniibacter aestuarii] |
| 06323 | 6 | DUF2783 domain-containing protein [Azoarcus indigens] |
| 08752 | 6 | MULTISPECIES: molecular chaperone TorD [Thauera] |
| 06330 | 6 | TfoX/Sxy family protein [Azoarcus olearius] |
| 08810 | 6 | hypothetical protein [Azoarcus nasutitermitis] |
| 07003 | 6 | DUF4384 domain-containing protein [Propionivibrio dicarboxylicus] |
| 07713 | 6 | hypothetical protein [Aromatoleum evansii] |
| 04170 | 6 | cytochrome b/b6 domain-containing protein [Aromatoleum toluclasticum] |
| 09787 | 6 | hypothetical protein [Aromatoleum toluclasticum] |
| 04134 | 6 | nuclear transport factor 2 family protein [Aromatoleum bremense] |
| 08031 | 6 | J domain-containing protein [Azoarcus olearius] |
| 05822 | 6 | hypothetical protein [Aromatoleum diolicum] |
| 08313 | 6 | IS5 family transposase [Thauera humireducens] |
| 02620 | 6 | SHOCT domain-containing protein [Denitromonas halophilus] |
| 01805 | 6 | hypothetical protein [Niveibacterium microcysteis] |
| 02770 | 6 | hypothetical protein [Azospira oryzae] |
| 06840 | 6 | hypothetical protein [Azoarcus communis] |
| 09926 | 6 | hypothetical protein [Azoarcus indigens] |
| 00991 | 6 | hypothetical protein [Uliginosibacterium gangwonense] |
| 08504 | 6 | hypothetical protein [Azospira oryzae] |
| 10014 | 6 | MULTISPECIES: hypothetical protein [Niveibacterium] |
| 09924 | 6 | histidine phosphatase family protein [Azoarcus sp. DD4] |
| 06887 | 6 | hypothetical protein [Azoarcus communis] |
| 08188 | 6 | ParB/RepB/Spo0J family partition protein [Azospira oryzae] |
| 01002 | 6 | HORMA domain containing protein [Aromatoleum evansii] |
| 08509 | 6 | hypothetical protein [Azoarcus indigens] |
| 00047 | 6 | MmoB/DmpM family protein [Azoarcus indigens] |
| 03856 | 6 | type I secretion C-terminal target domain-containing protein [Aromatoleum toluclasticum] |
| 03053 | 6 | hypothetical protein [Uliginosibacterium gangwonense] |
| 01011 | 6 | hypothetical protein [Aromatoleum evansii] |
| 07990 | 6 | alkaline phosphatase D family protein [Aromatoleum aromaticum] |
| 09810 | 6 | hypothetical protein [Azoarcus indigens] |
| 05508 | 6 | hypothetical protein [Azoarcus indigens] |
| 03027 | 6 | N-6 DNA methylase [Aromatoleum toluvorans] |
| 08545 | 6 | PEP-CTERM sorting domain-containing protein [Denitromonas halophilus] |
| 03021 | 6 | retention module-containing protein [Azoarcus indigens] |
| 05379 | 6 | ImmA/IrrE family metallo-endopeptidase [Aromatoleum toluclasticum] |
| 06041 | 6 | DUF6152 family protein [Aromatoleum pumilum] |
| 03879 | 6 | terminase large subunit [Azoarcus indigens] |
| 03282 | 6 | DUF4254 domain-containing protein [Azoarcus olearius] |
| 06902 | 6 | PEP-CTERM sorting domain-containing protein [Azoarcus indigens] |
| 01035 | 6 | hypothetical protein [Thauera phenolivorans] |
| 03895 | 6 | type II toxin-antitoxin system RelE/ParE family toxin [Azoarcus indigens] |
| 02971 | 6 | amidohydrolase family protein [Denitromonas halophilus] |
| 06546 | 6 | nucleotidyltransferase domain-containing protein [Aromatoleum bremense] |
| 00987 | 6 | GPW/gp25 family protein [Denitromonas sp. IR12] |
| 05416 | 6 | MULTISPECIES: TAXI family TRAP transporter solute-binding subunit [Aromatoleum] |
| 05417 | 6 | hypothetical protein [Aromatoleum toluclasticum] |
| 00877 | 6 | MarR family transcriptional regulator [Thauera aromatica] |
| 03720 | 6 | hypothetical protein [Azoarcus indigens] |
| 03252 | 6 | hypothetical protein [Aromatoleum toluclasticum] |
| 07976 | 6 | exopolysaccharide Pel transporter PelG [Denitromonas halophilus] |
| 06849 | 6 | hypothetical protein [Thauera butanivorans] |
| 07968 | 6 | DUF4254 domain-containing protein [Azoarcus indigens] |

| | | |
|---|---|---|
| 00263 | 6 | ThiF family adenylyltransferase [Zoogloea oleivorans] |
| 05473 | 6 | GNAT family N-acetyltransferase [Azoarcus indigens] |
| 03749 | 6 | WD40 repeat domain-containing protein [Azoarcus indigens] |
| 06857 | 6 | dihydroorotate dehydrogenase-like protein [Azoarcus communis] |
| 07962 | 6 | DUF3592 domain-containing protein [Denitromonas halophilus] |
| 08407 | 6 | DUF134 domain-containing protein [Propionivibrio dicarboxylicus] |
| 07955 | 6 | hypothetical protein [Azoarcus sp. KH32C] |
| 03188 | 6 | zinc ribbon domain-containing protein [Azoarcus olearius] |
| 03184 | 6 | AAA family ATPase [Thauera humireducens] |
| 09972 | 6 | hypothetical protein [Aromatoleum evansii] |
| 07987 | 6 | IS1595 family transposase [Aromatoleum toluclasticum] |
| 08445 | 6 | insulinase family protein [Niveibacterium umoris] |
| 06870 | 6 | MULTISPECIES: hypothetical protein [Thauera] |
| 06298 | 6 | YhcH/YjgK/YiaL family protein [Propionivibrio dicarboxylicus] |
| 06232 | 6 | prepilin-type N-terminal cleavage/methylation domain-containing protein [Azoarcus sp. KH32C] |
| 06579 | 6 | transporter substrate-binding domain-containing protein [Niveibacterium umoris] |
| 00264 | 6 | hypothetical protein [Aromatoleum evansii] |
| 05431 | 6 | hypothetical protein [Azoarcus communis] |
| 03825 | 6 | hypothetical protein [Thauera propionica] |
| 00433 | 6 | LysR family transcriptional regulator [Candidatus Dactylopiibacterium carminicum] |
| 00435 | 6 | AAA family ATPase [Aromatoleum diolicum] |
| 03086 | 6 | homocysteine S-methyltransferase family protein [Azoarcus olearius] |
| 07999 | 6 | phage major capsid protein [Aromatoleum aromaticum] |
| 03905 | 6 | MSMEG_0572 family nitrogen starvation response protein [Azoarcus indigens] |
| 08596 | 6 | hypothetical protein [Thauera terpenica] |
| 03299 | 6 | LuxR C-terminal-related transcriptional regulator [Azoarcus indigens] |
| 03970 | 6 | hypothetical protein [Aromatoleum diolicum] |
| 00464 | 6 | type II toxin-antitoxin system PemK/MazF family toxin [Aromatoleum aromaticum] |
| 07841 | 6 | MULTISPECIES: K(+)-transporting ATPase subunit F [Azoarcus] |
| 02848 | 6 | capsular biosynthesis protein [Aromatoleum bremense] |
| 09839 | 6 | glycoside hydrolase family 5 protein [Azoarcus olearius] |
| 06496 | 6 | hypothetical protein [Aromatoleum toluclasticum] |
| 03997 | 6 | SPOR domain-containing protein [Aromatoleum toluvorans] |
| 08676 | 6 | phage virion morphogenesis protein [Uliginosibacterium gangwonense] |
| 02834 | 6 | Ig-like domain-containing protein, partial [Azoarcus halotolerans] |
| 01108 | 6 | rhombotarget lipoprotein [Niveibacterium microcysteis] |
| 06435 | 6 | GDP-mannose mannosyl hydrolase [Azoarcus communis] |
| 02827 | 6 | MULTISPECIES: hypothetical protein [Thauera] |
| 10337 | 6 | MULTISPECIES: ribbon-helix-helix protein, CopG family [Thauera] |
| 02786 | 6 | SDR family NAD(P)-dependent oxidoreductase [Aromatoleum petrolei] |
| 07815 | 6 | choice-of-anchor A family protein [Azoarcus indigens] |
| 09824 | 6 | hypothetical protein [Aromatoleum toluclasticum] |
| 08710 | 6 | aspartate/glutamate racemase family protein [Propionivibrio dicarboxylicus] |
| 01128 | 6 | hypothetical protein [Azoarcus olearius] |
| 09820 | 6 | type II toxin-antitoxin system VapC family toxin [Azoarcus olearius] |
| 05278 | 6 | hypothetical protein [Niveibacterium microcysteis] |
| 01130 | 6 | hypothetical protein [Denitromonas halophilus] |
| 00812 | 6 | PAS domain-containing protein [Azoarcus indigens] |
| 01133 | 6 | MULTISPECIES: DUF4007 family protein [Rhodocyclales] |
| 09814 | 6 | hypothetical protein, partial [Aromatoleum buckelii] |
| 08316 | 6 | hypothetical protein [Nitrogeniibacter aestuarii] |
| 04042 | 6 | MULTISPECIES: DUF4124 domain-containing protein [Niveibacterium] |
| 08717 | 6 | DUF1653 domain-containing protein [Candidatus Dactylopiibacterium carminicum] |
| 02901 | 6 | hypothetical protein [Thauera lacus] |
| 09862 | 6 | hypothetical protein [Aromatoleum evansii] |
| 06303 | 6 | TIGR02646 family protein [Aromatoleum aromaticum] |
| 08192 | 6 | hypothetical protein [Azoarcus indigens] |
| 06908 | 6 | CDP-glycerol glycerophosphotransferase family protein [Aromatoleum pumilum] |
| 03911 | 6 | rhomboid family intramembrane serine protease [Niveibacterium microcysteis] |
| 03479 | 6 | branched-chain amino acid ABC transporter permease [Azoarcus taiwanensis] |
| 03912 | 6 | spherulation-specific family 4 protein [Denitromonas halophilus] |
| 03916 | 6 | DUF1778 domain-containing protein [Candidatus Dactylopiibacterium carminicum] |
| 06306 | 6 | DUF4962 domain-containing protein, partial [Thauera aminoaromatica] |
| 05863 | 6 | glucuronate isomerase [Propionivibrio dicarboxylicus] |
| 03294 | 6 | class I SAM-dependent methyltransferase [Azoarcus communis] |
| 03925 | 6 | XRE family transcriptional regulator [Denitromonas halophilus] |
| 08610 | 6 | hypothetical protein [Azoarcus olearius] |
| 03933 | 6 | EamA family transporter [Denitromonas halophilus] |
| 00460 | 6 | exodeoxyribonuclease I [Candidatus Dactylopiibacterium carminicum] |
| 03296 | 6 | DUF4400 domain-containing protein [Azoarcus indigens] |
| 06833 | 6 | 7-cyano-7-deazaguanine synthase [Aromatoleum aromaticum] |
| 06604 | 6 | hypothetical protein [Nitrogeniibacter aestuarii] |
| 02925 | 6 | MULTISPECIES: cyclic nucleotide-binding domain-containing protein [Azospira] |
| 09876 | 6 | hypothetical protein [Thauera phenolivorans] |
| 09874 | 6 | MULTISPECIES: pentaheme c-type cytochrome TorC [Thauera] |
| 01074 | 6 | hypothetical protein [Azospira oryzae] |
| 00848 | 6 | diaminopimelate dehydrogenase [Rhodocyclus purpureus] |
| 00027 | 6 | HD domain-containing protein [Aromatoleum pumilum] |
| 10035 | 6 | zinc ribbon domain-containing protein [Azoarcus olearius] |
| 06920 | 6 | MSMEG_0570 family nitrogen starvation response protein [Azoarcus indigens] |
| 08629 | 6 | alpha/beta hydrolase [Candidatus Dactylopiibacterium carminicum] |
| 09868 | 6 | hypothetical protein [Thauera butanivorans] |
| 05327 | 6 | helix-turn-helix domain-containing protein [Aromatoleum pumilum] |
| 03298 | 6 | alpha/beta hydrolase [Denitromonas halophilus] |
| 08904 | 6 | MULTISPECIES: hypothetical protein [Rhodocyclales] |
| 00526 | 6 | GPO family capsid scaffolding protein [Uliginosibacterium gangwonense] |
| 05786 | 6 | DUF4157 domain-containing protein [Denitromonas halophilus] |
| 05698 | 6 | helix-turn-helix domain-containing protein [Azoarcus indigens] |
| 08251 | 6 | NERD domain-containing protein [Azoarcus olearius] |
| 07489 | 6 | hypothetical protein [Azoarcus nasutitermitis] |
| 10552 | 6 | hypothetical protein [Azoarcus indigens] |
| 02039 | 6 | hypothetical protein [Thauera lacus] |
| 04577 | 6 | GGDEF domain-containing protein [Aromatoleum evansii] |
| 03391 | 6 | GlpM family protein [Azospira oryzae] |

| | | |
|---|---|---|
| 00131 | 6 | hypothetical protein [Denitromonas halophilus] |
| 05696 | 6 | PEP-CTERM sorting domain-containing protein [Azoarcus halotolerans] |
| 08068 | 6 | phage portal protein [Azoarcus indigens] |
| 01602 | 6 | DNA primase family protein [Thauera chlorobenzoica] |
| 09550 | 6 | SGNH/GDSL hydrolase family protein [Azoarcus indigens] |
| 04591 | 6 | hypothetical protein, partial [Denitromonas sp. IR12] |
| 06404 | 6 | polysaccharide pyruvyl transferase family protein [Azoarcus communis] |
| 09258 | 6 | hypothetical protein [Azoarcus sp. DD4] |
| 08070 | 6 | DUF3455 domain-containing protein [Aromatoleum aromaticum] |
| 00723 | 6 | hypothetical protein [Azoarcus communis] |
| 07464 | 6 | ISAs1 family transposase, partial [Azoarcus communis] |
| 04913 | 6 | hypothetical protein [Aromatoleum aromaticum] |
| 10132 | 6 | MULTISPECIES: hypothetical protein [Rhodocyclales] |
| 00290 | 6 | MULTISPECIES: ferredoxin [Aromatoleum] |
| 01997 | 6 | hypothetical protein [Aromatoleum aromaticum] |
| 08106 | 6 | PAS domain S-box protein [Candidatus Dactylopiibacterium carminicum] |
| 06724 | 6 | ATP-binding protein [Aromatoleum toluclasticum] |
| 09526 | 6 | heme-binding protein [Oryzomicrobium terrae] |
| 09522 | 6 | MULTISPECIES: hypothetical protein [Rhodocyclales] |
| 08103 | 6 | FAD-binding oxidoreductase [Aromatoleum petrolei] |
| 01990 | 6 | DUF3164 family protein [Aromatoleum evansii] |
| 10227 | 6 | SMP-30/gluconolactonase/LRE family protein [Candidatus Dactylopiibacterium carminicum] |
| 06761 | 6 | MULTISPECIES: hypothetical protein [Rhodocyclales] |
| 09560 | 6 | GNAT family N-acetyltransferase [Azoarcus taiwanensis] |
| 01571 | 6 | phage tail protein [Uliginosibacterium gangwonense] |
| 03402 | 6 | ATP-binding protein [Azoarcus nasutitermitis] |
| 04560 | 6 | PEP-CTERM sorting domain-containing protein [Denitromonas halophilus] |
| 04485 | 6 | MULTISPECIES: site-specific integrase [Proteobacteria] |
| 10413 | 6 | DUF2502 domain-containing protein [Denitromonas halophilus] |
| 00888 | 6 | hypothetical protein [Azoarcus indigens] |
| 09600 | 6 | PadR family transcriptional regulator [Azoarcus indigens] |
| 08119 | 6 | hypothetical protein [Azoarcus indigens] |
| 07165 | 6 | hypothetical protein [Aromatoleum aromaticum] |
| 01516 | 6 | transporter substrate-binding domain-containing protein [Niveibacterium microcysteis] |
| 02111 | 6 | CIA30 family protein [Azoarcus taiwanensis] |
| 02108 | 6 | hypothetical protein [Azoarcus indigens] |
| 07171 | 6 | TniQ family protein [Azoarcus communis] |
| 04507 | 6 | NIPSNAP family protein [Denitromonas halophilus] |
| 09587 | 6 | hypothetical protein [Azoarcus communis] |
| 04974 | 6 | hypothetical protein [Aromatoleum toluclasticum] |
| 04514 | 6 | DUF2799 domain-containing protein [Aromatoleum pumilum] |
| 09586 | 6 | GNAT family N-acetyltransferase [Niveibacterium microcysteis] |
| 00739 | 6 | DUF1538 domain-containing protein [Nitrogeniibacter aestuarii] |
| 02095 | 6 | AbrB/MazE/SpoVT family DNA-binding domain-containing protein [Aromatoleum bremense] |
| 00018 | 6 | hypothetical protein [Thauera humireducens] |
| 01260 | 6 | 30S ribosomal protein S6 [Thauera humireducens] |
| 06144 | 6 | carboxylesterase family protein [Azoarcus indigens] |
| 07198 | 6 | SMI1/KNR4 family protein [Azoarcus olearius] |
| 04534 | 6 | TVP38/TMEM64 family protein [Azoarcus communis] |
| 02080 | 6 | peptide-methionine (R)-S-oxide reductase MsrB [Denitromonas halophilus] |
| 04549 | 6 | DUF2252 domain-containing protein [Oryzomicrobium terrae] |
| 01551 | 6 | J domain-containing protein [Nitrogeniibacter aestuarii] |
| 04953 | 6 | MULTISPECIES: OsmC family protein [Niveibacterium] |
| 04555 | 6 | hypothetical protein [Candidatus Dactylopiibacterium carminicum] |
| 07206 | 6 | lipopolysaccharide assembly protein LapA domain-containing protein [Denitromonas halophilus] |
| 09563 | 6 | Mov34/MPN/PAD-1 family protein [Azoarcus communis] |
| 09510 | 6 | tetratricopeptide repeat protein [Denitromonas halophilus] |
| 07242 | 6 | hypothetical protein [Aromatoleum toluclasticum] |
| 00594 | 6 | hypothetical protein [Denitromonas halophilus] |
| 03586 | 6 | ImmA/IrrE family metallo-endopeptidase [Denitromonas sp. IR12] |
| 07398 | 6 | glycine zipper 2TM domain-containing protein [Aromatoleum toluclasticum] |
| 05675 | 6 | hypothetical protein [Aromatoleum toluclasticum] |
| 04689 | 6 | helix-turn-helix domain-containing protein [Aromatoleum toluolicum] |
| 04826 | 6 | AAA family ATPase [Azoarcus halotolerans] |
| 10151 | 6 | DUF3369 domain-containing protein [Niveibacterium microcysteis] |
| 06119 | 6 | DUF4124 domain-containing protein [Azoarcus communis] |
| 06101 | 6 | LuxR C-terminal-related transcriptional regulator [Niveibacterium sp. COAC-50] |
| 01881 | 6 | hypothetical protein [Aromatoleum evansii] |
| 10186 | 6 | sarcosine oxidase subunit gamma family protein, partial [Aromatoleum evansii] |
| 07296 | 6 | MULTISPECIES: phosphate ABC transporter substrate-binding protein [Niveibacterium] |
| 09381 | 6 | hypothetical protein [Aromatoleum aromaticum] |
| 04810 | 6 | GNAT family N-acetyltransferase [Denitromonas halophilus] |
| 04724 | 6 | CoA synthetase [Aromatoleum diolicum] |
| 04736 | 6 | prepilin-type N-terminal cleavage/methylation domain-containing protein [Aromatoleum aromaticum] |
| 07345 | 6 | cupin domain-containing protein [Aromatoleum aromaticum] |
| 00155 | 6 | ABC transporter permease subunit [Azoarcus sp. KH32C] |
| 09413 | 6 | TnsD family Tn7-like transposition protein [Zoogloea oleivorans] |
| 07308 | 6 | DUF2802 domain-containing protein [Candidatus Dactylopiibacterium carminicum] |
| 00683 | 6 | hypothetical protein [Azoarcus indigens] |
| 01836 | 6 | cytochrome C [Aromatoleum tolulyticum] |
| 04793 | 6 | BrnA antitoxin family protein [Azoarcus communis] |
| 01833 | 6 | DsrE family protein [Azoarcus halotolerans] |
| 00676 | 6 | YSC84-related protein [Azoarcus indigens] |
| 01799 | 6 | hypothetical protein [Thauera humireducens] |
| 00158 | 6 | MULTISPECIES: hypothetical protein [Niveibacterium] |
| 10179 | 6 | tape measure protein [Aromatoleum aromaticum] |
| 01800 | 6 | VCBS domain-containing protein, partial [Denitromonas sp. IR12] |
| 04779 | 6 | MaoC family dehydratase [Nitrogeniibacter aestuarii] |
| 07324 | 6 | FMN-binding protein [Azoarcus olearius] |
| 01742 | 6 | PhoX family protein [Nitrogeniibacter aestuarii] |
| 06278 | 6 | MULTISPECIES: hypothetical protein [Betaproteobacteria] |
| 01978 | 6 | nuclear transport factor 2 family protein [Denitromonas halophilus] |
| 09473 | 6 | MULTISPECIES: hypothetical protein [Azospira] |
| 09295 | 6 | hypothetical protein [Candidatus Dactylopiibacterium carminicum] |

| | | |
|---|---|---|
| 06376 | 6 | hypothetical protein [Azoarcus indigens] |
| 09302 | 6 | PD40 domain-containing protein [Uliginosibacterium aquaticum] |
| 08179 | 6 | N-acetylmuramoyl-L-alanine amidase [Nitrogeniibacter aestuarii] |
| 04882 | 6 | spore coat U domain-containing protein [Azoarcus sp. KH32C] |
| 04881 | 6 | phytanoyl-CoA dioxygenase family protein [Azoarcus indigens] |
| 01958 | 6 | hypothetical protein [Aromatoleum toluclasticum] |
| 01679 | 6 | zinc transporter ZntB [Nitrogeniibacter aestuarii] |
| 04876 | 6 | MULTISPECIES: glycoside hydrolase family 99-like domain-containing protein [Thauera] |
| 07437 | 6 | ectoine synthase [Aromatoleum pumilum] |
| 07436 | 6 | EthD domain-containing protein [Aromatoleum evansii] |
| 03606 | 6 | alpha/beta hydrolase [Niveibacterium microcysteis] |
| 09318 | 6 | hemerythrin domain-containing protein [Nitrogeniibacter aestuarii] |
| 01947 | 6 | hypothetical protein [Denitromonas halophilus] |
| 09326 | 6 | hypothetical protein [Niveibacterium microcysteis] |
| 03605 | 6 | hypothetical protein [Aromatoleum aromaticum] |
| 08235 | 6 | MULTISPECIES: BphX family protein [Rhodocyclales] |
| 06378 | 6 | MULTISPECIES: acyl carrier protein [Azoarcus] |
| 00305 | 6 | helix-turn-helix domain-containing protein [Azoarcus communis] |
| 06379 | 6 | hypothetical protein [Azoarcus indigens] |
| 09479 | 6 | TetR/AcrR family transcriptional regulator [Azoarcus indigens] |
| 01720 | 6 | PEP-CTERM sorting domain-containing protein [Denitromonas halophilus] |
| 08100 | 6 | MULTISPECIES: MerR family transcriptional regulator [Thauera] |
| 04666 | 6 | hypothetical protein [Azospira oryzae] |
| 05677 | 6 | hemerythrin domain-containing protein [Azoarcus indigens] |
| 07405 | 6 | energy-coupling factor ABC transporter substrate-binding protein [Propionivibrio dicarboxylicus] |
| 04844 | 6 | MULTISPECIES: hypothetical protein [Thauera] |
| 00706 | 6 | MULTISPECIES: peptide MFS transporter [Niveibacterium] |
| 01914 | 6 | COG2958 family protein [Candidatus Dactylopiibacterium carminicum] |
| 08270 | 6 | terminase small subunit [Azoarcus indigens] |
| 04602 | 6 | DUF5676 family membrane protein [Aromatoleum toluclasticum] |
| 03433 | 6 | type II toxin-antitoxin system RelE/ParE family toxin [Azoarcus communis] |
| 02287 | 6 | 2OG-Fe(II) oxygenase family protein [Aromatoleum toluclasticum] |
| 04363 | 6 | hypothetical protein [Aromatoleum bremense] |
| 04292 | 6 | MULTISPECIES: hypothetical protein [Proteobacteria] |
| 00357 | 6 | FIST C-terminal domain-containing protein [Azoarcus taiwanensis] |
| 09666 | 6 | hypothetical protein [Aromatoleum evansii] |
| 02415 | 6 | MULTISPECIES: VOC family protein [Niveibacterium] |
| 03351 | 6 | replication initiation protein [Thauera propionica] |
| 09665 | 6 | MULTISPECIES: Dna2/Cas4 domain-containing protein [Betaproteobacteria] |
| 02432 | 6 | TIGR02594 family protein [Aromatoleum evansii] |
| 05132 | 6 | nicotinate phosphoribosyltransferase [Azoarcus indigens] |
| 04389 | 6 | hypothetical protein [Azoarcus indigens] |
| 09084 | 6 | hypothetical protein [Azoarcus communis] |
| 09021 | 6 | nucleoside 2-deoxyribosyltransferase [Aromatoleum aromaticum] |
| 00759 | 6 | DUF3489 domain-containing protein [Aromatoleum bremense] |
| 07574 | 6 | nucleotidyltransferase domain-containing protein [Denitromonas halophilus] |
| 04404 | 6 | Crp/Fnr family transcriptional regulator [Azoarcus olearius] |
| 09101 | 6 | 23S rRNA (adenine(1618)-N(6))-methyltransferase RlmF [Azospira oryzae] |
| 05049 | 6 | YXWGXW repeat-containing protein [Candidatus Dactylopiibacterium carminicum] |
| 09000 | 6 | CHC2 zinc finger domain-containing protein [Rhodocyclus purpureus] |
| 08296 | 6 | MULTISPECIES: hypothetical protein [Rhodocyclales] |
| 10570 | 6 | hypothetical protein [Aromatoleum tolulyticum] |
| 09727 | 6 | hypothetical protein [Denitromonas halophilus] |
| 06803 | 6 | IS66 family transposase [Thauera aminoaromatica] |
| 00546 | 6 | hypothetical protein [Azoarcus communis] |
| 07615 | 6 | hypothetical protein [Azoarcus olearius] |
| 02248 | 6 | fibronectin type III domain-containing protein, partial [Zoogloea oleivorans] |
| 04334 | 6 | hypothetical protein [Thauera terpenica] |
| 04312 | 6 | AraC family transcriptional regulator [Azoarcus halotolerans] |
| 10444 | 6 | DUF5666 domain-containing protein [Azoarcus halotolerans] |
| 02375 | 6 | IclR family transcriptional regulator [Azoarcus communis] |
| 06415 | 6 | bifunctional glycoside hydrolase 114/ polysaccharide deacetylase family protein [Denitromonas halophilus] |
| 00543 | 6 | DUF1272 domain-containing protein [Aromatoleum diolicum] |
| 01317 | 6 | hypothetical protein [Aromatoleum toluolicum] |
| 05110 | 6 | hypothetical protein [Denitromonas halophilus] |
| 00386 | 6 | hypothetical protein, partial [Azoarcus halotolerans] |
| 03646 | 6 | alpha/beta hydrolase-fold protein [Niveibacterium microcysteis] |
| 05095 | 6 | SpoIIE family protein phosphatase [Azoarcus indigens] |
| 02390 | 6 | hypothetical protein [Aromatoleum diolicum] |
| 04346 | 6 | hypothetical protein [Azoarcus indigens] |
| 06690 | 6 | hypothetical protein [Azoarcus indigens] |
| 01309 | 6 | head completion/stabilization protein [Uliginosibacterium gangwonense] |
| 01304 | 6 | F0F1 ATP synthase subunit epsilon [Azoarcus communis] |
| 09708 | 6 | ethylbenzene dehydrogenase [Aromatoleum buckelii] |
| 06691 | 6 | hypothetical protein [Azoarcus communis] |
| 02361 | 6 | ATP-dependent helicase [Aromatoleum evansii] |
| 00535 | 6 | AAA domain-containing protein [Azoarcus halotolerans] |
| 01356 | 6 | ATP-binding protein [Thauera chlorobenzoica] |
| 01358 | 6 | DUF6036 family nucleotidyltransferase [Aromatoleum aromaticum] |
| 01300 | 6 | MULTISPECIES: hypothetical protein [Rhodocyclales] |
| 01426 | 6 | type II toxin-antitoxin system VapC family toxin [Azoarcus indigens] |
| 09102 | 6 | BREX-2 system phosphatase PglZ [Azoarcus indigens] |
| 04419 | 6 | roadblock/LC7 domain-containing protein [Azoarcus communis] |
| 02460 | 6 | MULTISPECIES: heme-degrading domain-containing protein [Niveibacterium] |
| 02203 | 6 | BON domain-containing protein [Azoarcus indigens] |
| 07550 | 6 | hypothetical protein [Azoarcus halotolerans] |
| 05994 | 6 | MULTISPECIES: phage tail protein [Rhodocyclaceae] |
| 07135 | 6 | hypothetical protein [Azoarcus indigens] |
| 07044 | 6 | hypothetical protein [Azoarcus nasutitermitis] |
| 09152 | 6 | hypothetical protein [Denitromonas halophilus] |
| 00378 | 6 | hypothetical protein [Azoarcus indigens] |
| 02475 | 6 | DUF3597 domain-containing protein [Aromatoleum diolicum] |
| 06363 | 6 | PLxRFG domain-containing protein [Thauera humireducens] |
| 09625 | 6 | hypothetical protein [Nitrogeniibacter aestuarii] |

| | | |
|---|---|---|
| **02175** | 6 | hypothetical protein [Aromatoleum evansii] |
| **01266** | 6 | hypothetical protein [Aromatoleum toluclasticum] |
| **04250** | 6 | OmpA family protein [Niveibacterium umoris] |
| **04473** | 6 | MULTISPECIES: hypothetical protein [Rhodocyclales] |
| **01263** | 6 | four helix bundle protein [Aromatoleum aromaticum] |
| **01468** | 6 | DUF1161 domain-containing protein [Candidatus Dactylopiibacterium carminicum] |
| **05591** | 6 | virulence factor family protein [Azoarcus indigens] |
| **05152** | 6 | NAD/NADP-dependent octopine/nopaline dehydrogenase family protein [Azoarcus indigens] |
| **09608** | 6 | AAA family ATPase [Aromatoleum aromaticum] |
| **07692** | 6 | condensation domain-containing protein, partial [Denitromonas sp. IR12] |
| **05156** | 6 | hypothetical protein [Azoarcus indigens] |
| **09607** | 6 | flagellar assembly peptidoglycan hydrolase FlgJ [Azoarcus indigens] |
| **07039** | 6 | hypothetical protein [Azoarcus indigens] |
| **01444** | 6 | MULTISPECIES: lactoylglutathione lyase [Rhodocyclales] |
| **02379** | 6 | MULTISPECIES: hypothetical protein [Aromatoleum] |
| **03560** | 6 | VOC family protein [Denitromonas halophilus] |
| **10557** | 6 | hypothetical protein [Nitrogeniibacter aestuarii] |
| **04436** | 6 | GNAT family N-acetyltransferase [Azoarcus taiwanensis] |
| **07562** | 6 | DUF2191 domain-containing protein [Aromatoleum buckelii] |
| **04437** | 6 | MULTISPECIES: hypothetical protein [Zoogloea] |
| **02452** | 6 | dodecin family protein [Denitromonas halophilus] |
| **06804** | 6 | hypothetical protein [Candidatus Dactylopiibacterium carminicum] |
| **05035** | 6 | S8 family serine peptidase [Aromatoleum aromaticum] |
| **04425** | 6 | hypothetical protein [Denitromonas halophilus] |
| **05978** | 5 | hypothetical protein [Niveibacterium microcysteis] |
| **10327** | 5 | PEP-CTERM sorting domain-containing protein [Denitromonas halophilus] |
| **06492** | 5 | 4Fe-4S dicluster domain-containing protein, partial [Aromatoleum diolicum] |
| **00436** | 5 | SAM-dependent methyltransferase [Niveibacterium umoris] |
| **00340** | 5 | XRE family transcriptional regulator [Azoarcus halotolerans] |
| **10386** | 5 | MarR family transcriptional regulator [Azoarcus indigens] |
| **05896** | 5 | hypothetical protein [Aromatoleum toluclasticum] |
| **06742** | 5 | dehydrogenase [Aromatoleum tolulyticum] |
| **05633** | 5 | hypothetical protein [Denitromonas halophilus] |
| **00884** | 5 | glucosylglycerol 3-phosphatase [Nitrogeniibacter aestuarii] |
| **06842** | 5 | hypothetical protein [Azoarcus rhizosphaerae] |
| **06512** | 5 | NAD(+)/NADH kinase [Nitrogeniibacter aestuarii] |
| **05680** | 5 | hypothetical protein [Azoarcus sp. KH32C] |
| **10406** | 5 | DUF2384 domain-containing protein [Thauera phenylacetica] |
| **05489** | 5 | ATP synthase subunit I [Denitromonas halophilus] |
| **00405** | 5 | hypothetical protein [Thauera humireducens] |
| **06646** | 5 | ThuA domain-containing protein [Aromatoleum petrolei] |
| **10316** | 5 | DNA methyltransferase [Thauera linaloolentis] |
| **10376** | 5 | DUF805 domain-containing protein [Zoogloea dura] |
| **05781** | 5 | DUF2214 family protein [Azoarcus indigens] |
| **06663** | 5 | hypothetical protein [Azoarcus halotolerans] |
| **00426** | 5 | hypothetical protein [Azoarcus communis] |
| **05980** | 5 | beta-ribofuranosylaminobenzene 5'-phosphate synthase [Aromatoleum toluvorans] |
| **06007** | 5 | hypothetical protein [Aromatoleum toluvorans] |
| **05777** | 5 | CDGSH iron-sulfur domain-containing protein [Denitromonas halophilus] |
| **06676** | 5 | type IV secretion system protein [Thauera terpenica] |
| **00529** | 5 | DUF883 family protein [Azoarcus indigens] |
| **10149** | 5 | hypothetical protein [Niveibacterium umoris] |
| **06521** | 5 | MULTISPECIES: Dabb family protein [Niveibacterium] |
| **05901** | 5 | GNAT family N-acetyltransferase [Niveibacterium umoris] |
| **05795** | 5 | HI0074 family nucleotidyltransferase substrate-binding subunit [Niveibacterium microcysteis] |
| **06734** | 5 | DUF262 domain-containing protein [Niveibacterium microcysteis] |
| **05882** | 5 | TMEM175 family protein [Azoarcus sp. KH32C] |
| **05646** | 5 | cytochrome c3 family protein [Nitrogeniibacter aestuarii] |
| **06511** | 5 | M12 family metallopeptidase [Azoarcus halotolerans] |
| **00590** | 5 | hypothetical protein [Azospira oryzae] |
| **10329** | 5 | hypothetical protein [Azoarcus olearius] |
| **10233** | 5 | nuclear transport factor 2 family protein [Azoarcus indigens] |
| **00736** | 5 | hydrogenase maturation peptidase HycI [Niveibacterium microcysteis] |
| **06771** | 5 | D-alanyl-D-alanine carboxypeptidase family protein [Aromatoleum buckelii] |
| **05524** | 5 | DUF2786 domain-containing protein [Azoarcus communis] |
| **10446** | 5 | sulfotransferase [Denitromonas halophilus] |
| **05705** | 5 | nucleotidyltransferase domain-containing protein [Aromatoleum aromaticum] |
| **00811** | 5 | hypothetical protein [Azoarcus olearius] |
| **10277** | 5 | DUF4276 family protein [Aromatoleum aromaticum] |
| **00853** | 5 | MULTISPECIES: P-loop NTPase fold protein [Betaproteobacteria] |
| **06606** | 5 | type VI secretion system Vgr family protein, partial [Nitrogeniibacter aestuarii] |
| **05589** | 5 | DUF2058 domain-containing protein [Oryzomicrobium terrae] |
| **05608** | 5 | hypothetical protein [Nitrogeniibacter aestuarii] |
| **06595** | 5 | MULTISPECIES: ParB/RepB/Spo0J family partition protein [Bacteria] |
| **06698** | 5 | MULTISPECIES: hypothetical protein [Thauera] |
| **00567** | 5 | biliverdin-producing heme oxygenase [Thauera linaloolentis] |
| **10239** | 5 | phage major capsid protein, P2 family [Uliginosibacterium gangwonense] |
| **10271** | 5 | alpha/beta fold hydrolase [Propionivibrio dicarboxylicus] |
| **10346** | 5 | hypothetical protein [Aromatoleum pumilum] |
| **10040** | 5 | LysR family transcriptional regulator [Denitromonas halophilus] |
| **10124** | 5 | hypothetical protein [Aromatoleum pumilum] |
| **00469** | 5 | type II toxin-antitoxin system VapC family toxin [Aromatoleum pumilum] |
| **00280** | 5 | AraC family transcriptional regulator [Uliginosibacterium gangwonense] |
| **06706** | 5 | hypothetical protein [Azoarcus communis] |
| **00585** | 5 | energy transducer TonB [Azoarcus indigens] |
| **05711** | 5 | hypothetical protein [Aromatoleum pumilum] |
| **00597** | 5 | hypothetical protein [Candidatus Dactylopiibacterium carminicum] |
| **10117** | 5 | tetratricopeptide repeat protein, partial [Propionivibrio dicarboxylicus] |
| **10265** | 5 | DHCW motif cupin fold protein [Denitromonas halophilus] |
| **00595** | 5 | S1C family serine protease [Denitromonas halophilus] |
| **06543** | 5 | LysR family transcriptional regulator [Azoarcus halotolerans] |
| **06713** | 5 | SPOR domain-containing protein [Oryzomicrobium terrae] |
| **10458** | 5 | hypothetical protein [Aromatoleum evansii] |
| **10392** | 5 | YncE family protein [Azoarcus nasutitermitis] |

| | | |
|---|---|---|
| 05737 | 5 | substrate-binding domain-containing protein [Azoarcus indigens] |
| 10135 | 5 | tetratricopeptide repeat protein [Aromatoleum diolicum] |
| 06478 | 5 | MULTISPECIES: NifU family protein [Thauera] |
| 05688 | 5 | hypothetical protein [Niveibacterium microcysteis] |
| 10020 | 5 | BamA/TamA family outer membrane protein [Uliginosibacterium aquaticum] |
| 05627 | 5 | DUF3261 domain-containing protein [Denitromonas halophilus] |
| 10019 | 5 | N-acetyl sugar amidotransferase [Denitromonas halophilus] |
| 10365 | 5 | DUF2523 domain-containing protein [Azoarcus indigens] |
| 00498 | 5 | type I-E CRISPR-associated protein Cse2/CasB [Aromatoleum bremense] |
| 00270 | 5 | MULTISPECIES: plasmid replication initiator TrfA [Proteobacteria] |
| 10333 | 5 | hypothetical protein [Azoarcus taiwanensis] |
| 00346 | 5 | DUF4394 domain-containing protein [Azoarcus communis] |
| 06811 | 5 | flippase-like domain-containing protein [Aromatoleum pumilum] |
| 10408 | 5 | Hpt domain-containing protein [Azospira oryzae] |
| 00768 | 5 | peptidylprolyl isomerase [Azoarcus communis] |
| 10220 | 5 | MULTISPECIES: hypothetical protein [Niveibacterium] |
| 10459 | 5 | dihydrofolate reductase family protein [Denitromonas halophilus] |
| 05872 | 5 | hypothetical protein [Aromatoleum aromaticum] |
| 10363 | 5 | choice-of-anchor L domain-containing protein [Azoarcus communis] |
| 00361 | 5 | hypothetical protein [Azoarcus indigens] |
| 05954 | 5 | MULTISPECIES: cupin domain-containing protein [Proteobacteria] |
| 05869 | 5 | hypothetical protein [Azoarcus communis] |
| 00551 | 5 | Ig domain-containing protein [Thauera hydrothermalis] |
| 05512 | 5 | hypothetical protein, partial [Azoarcus halotolerans] |
| 05619 | 5 | hypothetical protein [Aromatoleum evansii] |
| 00720 | 5 | YkgJ family cysteine cluster protein [Candidatus Dactylopiibacterium carminicum] |
| 10295 | 5 | MULTISPECIES: hypothetical protein [Thauera] |
| 10105 | 5 | TIR domain-containing protein [Aromatoleum toluclasticum] |
| 00319 | 5 | N-6 DNA methylase [Thauera propionica] |
| 10066 | 5 | twin-arginine translocation signal domain-containing protein [Denitromonas halophilus] |
| 00559 | 5 | RusA family crossover junction endodeoxyribonuclease [Azoarcus indigens] |
| 05937 | 5 | XRE family transcriptional regulator [Aromatoleum aromaticum] |
| 00446 | 5 | hypothetical protein [Aromatoleum pumilum] |
| 10332 | 5 | DUF3817 domain-containing protein [Thauera lacus] |
| 10125 | 5 | universal stress protein [Denitromonas halophilus] |
| 01862 | 5 | DUF2442 domain-containing protein [Aromatoleum toluclasticum] |
| 00891 | 5 | hypothetical protein [Aromatoleum aromaticum] |
| 08861 | 5 | hypothetical protein [Denitromonas halophilus] |
| 00068 | 5 | ribbon-helix-helix protein, CopG family [Azoarcus indigens] |
| 02651 | 5 | sulfite exporter TauE/SafE family protein [Azoarcus rhizosphaerae] |
| 08817 | 5 | MULTISPECIES: hypothetical protein [Proteobacteria] |
| 07750 | 5 | retention module-containing protein [Azoarcus olearius] |
| 00072 | 5 | DNA-binding protein [Azoarcus sp. DD4] |
| 08834 | 5 | hypothetical protein [Denitromonas halophilus] |
| 04158 | 5 | hypothetical protein [Candidatus Dactylopiibacterium carminicum] |
| 04159 | 5 | transporter substrate-binding domain-containing protein [Uliginosibacterium gangwonense] |
| 08842 | 5 | tyrosine-type recombinase/integrase [Aromatoleum buckelii] |
| 08847 | 5 | Hcp family type VI secretion system effector [Azoarcus indigens] |
| 08855 | 5 | hypothetical protein [Propionivibrio dicarboxylicus] |
| 04178 | 5 | MULTISPECIES: hypothetical protein [Zoogloeaceae] |
| 02564 | 5 | S41 family peptidase [Niveibacterium microcysteis] |
| 02167 | 5 | hypothetical protein [Azoarcus indigens] |
| 04192 | 5 | lysine N(6)-hydroxylase/L-ornithine N(5)-oxygenase family protein [Azoarcus indigens] |
| 02542 | 5 | ATP-binding protein [Azoarcus communis] |
| 04203 | 5 | pilin [Aromatoleum bremense] |
| 02534 | 5 | transposase, partial [Azoarcus indigens] |
| 07712 | 5 | hypothetical protein [Aromatoleum evansii] |
| 04213 | 5 | sigma-70 family RNA polymerase sigma factor [Azoarcus olearius] |
| 08903 | 5 | cache domain-containing protein, partial [Azoarcus sp. TTM-91] |
| 04228 | 5 | toll/interleukin-1 receptor domain-containing protein [Azoarcus communis] |
| 02510 | 5 | hypothetical protein [Aromatoleum petrolei] |
| 00085 | 5 | hypothetical protein [Denitromonas halophilus] |
| 06342 | 5 | DUF6498-containing protein [Aromatoleum pumilum] |
| 08918 | 5 | MULTISPECIES: YgjV family protein [Uliginosibacterium] |
| 02660 | 5 | MULTISPECIES: hypothetical protein [Thauera] |
| 02669 | 5 | hypothetical protein [Denitromonas halophilus] |
| 06318 | 5 | hypothetical protein [Azoarcus indigens] |
| 02701 | 5 | DUF6152 family protein [Aromatoleum aromaticum] |
| 02841 | 5 | MotA/TolQ/ExbB proton channel family protein [Azoarcus indigens] |
| 02838 | 5 | hypothetical protein [Aromatoleum toluclasticum] |
| 04004 | 5 | MULTISPECIES: broad-spectrum mercury transporter MerE [Proteobacteria] |
| 04005 | 5 | MULTISPECIES: glycosyltransferase [Rhodocyclales] |
| 02833 | 5 | IS3 family transposase [Aromatoleum bremense] |
| 02821 | 5 | hypothetical protein [Azoarcus indigens] |
| 08698 | 5 | PKD domain-containing protein, partial [Zoogloea ramigera] |
| 08699 | 5 | DNA methyltransferase [Aromatoleum aromaticum] |
| 02797 | 5 | nuclear transport factor 2 family protein [Aromatoleum toluclasticum] |
| 07819 | 5 | YjhX family toxin [Azoarcus communis] |
| 02794 | 5 | hypothetical protein [Azoarcus olearius] |
| 07817 | 5 | DUF3396 domain-containing protein [Azoarcus indigens] |
| 08707 | 5 | HipA domain-containing protein [Thauera phenylacetica] |
| 07806 | 5 | DUF6471 domain-containing protein [Aromatoleum pumilum] |
| 08719 | 5 | c-type cytochrome [Aromatoleum diolicum] |
| 06208 | 5 | hypothetical protein [Uliginosibacterium aquaticum] |
| 02744 | 5 | tape measure protein [Aromatoleum toluclasticum] |
| 07787 | 5 | hypothetical protein [Azoarcus indigens] |
| 02738 | 5 | flagellar FliJ family protein [Azoarcus indigens] |
| 08747 | 5 | hypothetical protein [Thauera butanivorans] |
| 02732 | 5 | hypothetical protein [Aromatoleum petrolei] |
| 02728 | 5 | Ig-like domain-containing protein [Aromatoleum diolicum] |
| 08754 | 5 | NUDIX domain-containing protein [Propionivibrio dicarboxylicus] |
| 04099 | 5 | HupE/UreJ family protein [Azoarcus communis] |
| 02706 | 5 | hypothetical protein [Aromatoleum evansii] |
| 02484 | 5 | tannase/feruloyl esterase family alpha/beta hydrolase [Denitromonas sp. IR12] |

| | | |
|---|---|---|
| 07699 | 5 | hypothetical protein [Aromatoleum toluclasticum] |
| 02474 | 5 | DUF4166 domain-containing protein [Denitromonas halophilus] |
| 09041 | 5 | MULTISPECIES: alpha/beta fold hydrolase [Proteobacteria] |
| 09054 | 5 | AbrB/MazE/SpoVT family DNA-binding domain-containing protein [Azoarcus rhizosphaerae] |
| 09058 | 5 | GNAT family N-acetyltransferase [Aromatoleum buckelii] |
| 10559 | 5 | MULTISPECIES: hypothetical protein [Niveibacterium] |
| 09063 | 5 | MULTISPECIES: patatin-like protein [Thauera] |
| 09064 | 5 | hypothetical protein [Aromatoleum diolicum] |
| 04386 | 5 | hypothetical protein [Azoarcus indigens] |
| 09076 | 5 | MULTISPECIES: TrbM/KikA/MpfK family conjugal transfer protein [Proteobacteria] |
| 07586 | 5 | hypothetical protein [Aromatoleum diolicum] |
| 09083 | 5 | restriction endonuclease [Aromatoleum toluclasticum] |
| 07579 | 5 | (p)ppGpp synthetase [Aromatoleum toluolicum] |
| 00113 | 5 | DUF2946 family protein [Aromatoleum toluvorans] |
| 09105 | 5 | hypothetical protein [Azoarcus indigens] |
| 09106 | 5 | hypothetical protein [Azoarcus indigens] |
| 02241 | 5 | EF-hand domain-containing protein [Aromatoleum toluvorans] |
| 02239 | 5 | hypothetical protein [Aromatoleum toluclasticum] |
| 07563 | 5 | transcriptional regulator [Aromatoleum bremense] |
| 09114 | 5 | DNA cytosine methyltransferase [Aromatoleum pumilum] |
| 02229 | 5 | MULTISPECIES: VOC family protein [Niveibacterium] |
| 02219 | 5 | cupin domain-containing protein [Propionivibrio dicarboxylicus] |
| 05481 | 5 | hypothetical protein [Denitromonas halophilus] |
| 02199 | 5 | PASTA domain-containing protein [Aromatoleum toluclasticum] |
| 04464 | 5 | O-antigen ligase family protein [Aromatoleum bremense] |
| 06152 | 5 | hypothetical protein [Aromatoleum toluolicum] |
| 09171 | 5 | hypothetical protein [Aromatoleum pumilum] |
| 04474 | 5 | hypothetical protein [Azoarcus communis] |
| 04371 | 5 | LLM class oxidoreductase [Denitromonas halophilus] |
| 04364 | 5 | hypothetical protein [Nitrogeniibacter aestuarii] |
| 04257 | 5 | VPA1267 family protein [Aromatoleum petrolei] |
| 07609 | 5 | CRISPR system precrRNA processing endoribonuclease RAMP protein Cas6 [Niveibacterium sp. COAC-50] |
| 04269 | 5 | hypothetical protein [Niveibacterium sp. COAC-50] |
| 02444 | 5 | META domain-containing protein [Propionivibrio dicarboxylicus] |
| 08948 | 5 | hypothetical protein [Azoarcus halotolerans] |
| 02438 | 5 | recombinase RecT [Thauera humireducens] |
| 08957 | 5 | glycoside hydrolase family 5 protein [Denitromonas halophilus] |
| 00102 | 5 | Hpt domain-containing protein [Nitrogeniibacter mangrovi] |
| 04280 | 5 | MFS transporter [Aromatoleum toluclasticum] |
| 04287 | 5 | hypothetical protein [Aromatoleum pumilum] |
| 08968 | 5 | SDR family oxidoreductase [Aromatoleum petrolei] |
| 07651 | 5 | glycosyl transferase family 1 [Azoarcus olearius] |
| 04294 | 5 | hypothetical protein [Aromatoleum aromaticum] |
| 06173 | 5 | DUF456 domain-containing protein [Aromatoleum buckelii] |
| 04299 | 5 | MULTISPECIES: hypothetical protein [Aromatoleum] |
| 08984 | 5 | putative molybdenum carrier protein [Aromatoleum toluclasticum] |
| 04309 | 5 | CRISPR-associated protein Csx16 [Niveibacterium sp. COAC-50] |
| 02384 | 5 | TlpA family protein disulfide reductase [Aromatoleum diolicum] |
| 04317 | 5 | DUF58 domain-containing protein [Niveibacterium microcysteis] |
| 09002 | 5 | HPF/RaiA family ribosome-associated protein [Aromatoleum aromaticum] |
| 04323 | 5 | N-acetylglucosamine-specific PTS transporter subunit IIBC [Uliginosibacterium gangwonense] |
| 02367 | 5 | metal-dependent hydrolase [Aromatoleum evansii] |
| 04335 | 5 | hypothetical protein [Thauera terpenica] |
| 09007 | 5 | cyclic peptide export ABC transporter [Azoarcus indigens] |
| 02347 | 5 | VCBS domain-containing protein, partial [Azoarcus halotolerans] |
| 04350 | 5 | GpE family phage tail protein [Denitromonas sp. IR12] |
| 07611 | 5 | hypothetical protein [Niveibacterium sp. COAC-50] |
| 02842 | 5 | hypothetical protein [Aromatoleum diolicum] |
| 03994 | 5 | DUF262 domain-containing protein [Aromatoleum toluvorans] |
| 07835 | 5 | MULTISPECIES: Arc family DNA binding domain-containing protein [Azoarcus] |
| 00019 | 5 | helix-turn-helix transcriptional regulator [Propionivibrio dicarboxylicus] |
| 08266 | 5 | hypothetical protein [Nitrogeniibacter aestuarii] |
| 08054 | 5 | hypothetical protein [Aromatoleum bremense] |
| 08053 | 5 | hypothetical protein [Aromatoleum pumilum] |
| 08279 | 5 | MULTISPECIES: hypothetical protein [Rhodocyclales] |
| 08280 | 5 | sterol desaturase family protein [Azoarcus communis] |
| 08287 | 5 | hypothetical protein [Azoarcus olearius] |
| 03648 | 5 | hypothetical protein [Aromatoleum toluclasticum] |
| 08297 | 5 | hypothetical protein [Aromatoleum diolicum] |
| 03656 | 5 | TetR family transcriptional regulator [Azoarcus sp. TTM-91] |
| 00024 | 5 | CRISPR-associated protein Cse1 [Aromatoleum toluclasticum] |
| 03318 | 5 | cupin domain-containing protein [Azoarcus nasutitermitis] |
| 08019 | 5 | DUF1800 domain-containing protein [Azoarcus indigens] |
| 03674 | 5 | TIR domain-containing protein [Denitromonas halophilus] |
| 03676 | 5 | hypothetical protein [Aromatoleum toluvorans] |
| 06248 | 5 | hemerythrin domain-containing protein [Nitrogeniibacter aestuarii] |
| 08009 | 5 | mechanosensitive ion channel [Niveibacterium microcysteis] |
| 03686 | 5 | MULTISPECIES: hypothetical protein [Proteobacteria] |
| 03290 | 5 | hypothetical protein [Azoarcus olearius] |
| 08335 | 5 | hypothetical protein [Candidatus Dactylopiibacterium carminicum] |
| 08337 | 5 | cysteine hydrolase [Candidatus Dactylopiibacterium carminicum] |
| 07979 | 5 | ROK family protein [Niveibacterium microcysteis] |
| 03274 | 5 | YjbH domain-containing protein [Azoarcus sp. KH32C] |
| 06242 | 5 | class I SAM-dependent methyltransferase [Azoarcus communis] |
| 03246 | 5 | hypothetical protein [Zoogloea oleivorans] |
| 06295 | 5 | phage major tail tube protein [Denitromonas sp. IR12] |
| 03373 | 5 | hypothetical protein [Nitrogeniibacter aestuarii] |
| 03616 | 5 | ATP-binding protein [Aromatoleum aromaticum] |
| 03736 | 5 | MULTISPECIES: glycosyltransferase [Aromatoleum] |
| 08072 | 5 | MULTISPECIES: TraX family protein [Gammaproteobacteria] |
| 08182 | 5 | 3'-5' exoribonuclease [Thauera butanivorans] |
| 08183 | 5 | MULTISPECIES: replication protein C [Proteobacteria] |
| 08170 | 5 | hypothetical protein [Azoarcus indigens] |
| 03487 | 5 | DUF1353 domain-containing protein [Aromatoleum pumilum] |

| | | |
|---|---|---|
| 03519 | 5 | ABC transporter substrate-binding protein [Candidatus Dactylopiibacterium carminicum] |
| 03474 | 5 | chemotaxis protein CheW [Aromatoleum toluclasticum] |
| 03470 | 5 | alpha-D-ribose 1-methylphosphonate 5-triphosphate diphosphatase [Azoarcus sp. L1K30] |
| 03538 | 5 | ACT domain-containing protein [Propionivibrio dicarboxylicus] |
| 03456 | 5 | hypothetical protein [Azoarcus communis] |
| 03540 | 5 | nucleotidyl transferase AbiEii/AbiGii toxin family protein [Azoarcus nasutitermitis] |
| 03432 | 5 | zinc-ribbon domain-containing protein, partial [Denitromonas halophilus] |
| 08133 | 5 | CRISPR-associated ring nuclease Csm6 [Thauera butanivorans] |
| 08130 | 5 | hypothetical protein [Azoarcus halotolerans] |
| 10622 | 5 | GTP-binding protein, partial [Aromatoleum tolulyticum] |
| 08124 | 5 | alpha/beta fold hydrolase [Azoarcus indigens] |
| 08122 | 5 | aldehyde dehydrogenase family protein, partial [Thauera butanivorans] |
| 08121 | 5 | HNH endonuclease [Azoarcus indigens] |
| 08111 | 5 | cupin domain-containing protein [Aromatoleum toluclasticum] |
| 08102 | 5 | response regulator [Aromatoleum tolulyticum] |
| 03581 | 5 | hypothetical protein [Azoarcus indigens] |
| 08099 | 5 | type II toxin-antitoxin system Phd/YefM family antitoxin [Aromatoleum pumilum] |
| 08097 | 5 | IS1634 family transposase, partial [Thauera phenylacetica] |
| 08086 | 5 | hypothetical protein [Azoarcus olearius] |
| 08078 | 5 | hypothetical protein [Azoarcus indigens] |
| 03405 | 5 | DUF6491 family protein [Azoarcus indigens] |
| 00038 | 5 | hypothetical protein [Azospira oryzae] |
| 03740 | 5 | J domain-containing protein [Aromatoleum bremense] |
| 02859 | 5 | hypothetical protein [Azoarcus communis] |
| 03093 | 5 | EF-Tu/IF-2/RF-3 family GTPase, partial [Thauera propionica] |
| 03850 | 5 | FliM/FliN family flagellar motor C-terminal domain-containing protein [Azoarcus indigens] |
| 08528 | 5 | DMT family transporter [Oryzomicrobium terrae] |
| 03859 | 5 | TSUP family transporter [Azoarcus indigens] |
| 06302 | 5 | MoaD/ThiS family protein [Aromatoleum toluclasticum] |
| 03870 | 5 | EAL domain-containing protein [Azospira oryzae] |
| 08549 | 5 | MULTISPECIES: TIGR02391 family protein [Proteobacteria] |
| 08550 | 5 | toll/interleukin-1 receptor domain-containing protein [Azoarcus communis] |
| 08578 | 5 | MULTISPECIES: hypothetical protein [Rhodocyclales] |
| 07877 | 5 | site-2 protease family protein [Denitromonas halophilus] |
| 07873 | 5 | cobalamin B12-binding domain-containing protein [Denitromonas halophilus] |
| 10596 | 5 | glutaredoxin [Denitromonas halophilus] |
| 06217 | 5 | ABC transporter substrate-binding protein [Nitrogeniibacter aestuarii] |
| 03930 | 5 | antibiotic biosynthesis monooxygenase [Denitromonas halophilus] |
| 03935 | 5 | PGDYG domain-containing protein [Denitromonas halophilus] |
| 08612 | 5 | Clp protease ClpP [Oryzomicrobium terrae] |
| 02934 | 5 | hypothetical protein [Azoarcus indigens] |
| 02929 | 5 | MarR family transcriptional regulator [Azoarcus indigens] |
| 08618 | 5 | type IV pilin protein [Aromatoleum pumilum] |
| 03944 | 5 | MULTISPECIES: aspartate 1-decarboxylase [Azospira] |
| 02898 | 5 | tetratricopeptide repeat protein [Azoarcus communis] |
| 02884 | 5 | hypothetical protein [Thauera butanivorans] |
| 08657 | 5 | aspartyl/asparaginyl beta-hydroxylase domain-containing protein [Denitromonas halophilus] |
| 02869 | 5 | class I SAM-dependent methyltransferase [Azoarcus indigens] |
| 10588 | 5 | hypothetical protein [Aromatoleum diolicum] |
| 03986 | 5 | ectoine hydroxylase [Azoarcus taiwanensis] |
| 08498 | 5 | flagellar biosynthesis anti-sigma factor FlgM [Azoarcus indigens] |
| 08486 | 5 | hypothetical protein [Aromatoleum aromaticum] |
| 10604 | 5 | radical SAM protein [Nitrogeniibacter aestuarii] |
| 03105 | 5 | OmpA family protein [Azoarcus olearius] |
| 10598 | 5 | hypothetical protein [Azoarcus olearius] |
| 08411 | 5 | hypothetical protein [Aromatoleum toluclasticum] |
| 06236 | 5 | MULTISPECIES: acyl carrier protein [Niveibacterium] |
| 03187 | 5 | hypothetical protein [Aromatoleum evansii] |
| 03180 | 5 | phage tail tape measure protein [Azoarcus sp. DD4] |
| 08432 | 5 | Fic family protein [Azoarcus olearius] |
| 07945 | 5 | hypothetical protein [Azoarcus communis] |
| 03163 | 5 | MULTISPECIES: patatin-like phospholipase family protein [Azospira] |
| 08440 | 5 | type II secretion system protein GspM [Azoarcus indigens] |
| 08443 | 5 | hypothetical protein [Uliginosibacterium gangwonense] |
| 03156 | 5 | alpha/beta hydrolase [Uliginosibacterium gangwonense] |
| 07939 | 5 | sterol desaturase family protein [Aromatoleum pumilum] |
| 03155 | 5 | phage terminase small subunit [Uliginosibacterium gangwonense] |
| 10597 | 5 | PEPxxWA-CTERM sorting domain-containing protein [Denitromonas halophilus] |
| 03147 | 5 | MULTISPECIES: hypothetical protein [Thauera] |
| 08451 | 5 | DUF3179 domain-containing protein [Aromatoleum toluclasticum] |
| 03139 | 5 | hypothetical protein [Azoarcus olearius] |
| 03805 | 5 | baseplate J/gp47 family protein [Denitromonas sp. IR12] |
| 03126 | 5 | EAL domain-containing protein, partial [Aromatoleum toluclasticum] |
| 07924 | 5 | fatty acid desaturase [Azoarcus communis] |
| 08474 | 5 | helix-turn-helix domain-containing protein [Aromatoleum petrolei] |
| 03115 | 5 | HEAT repeat domain-containing protein [Denitromonas halophilus] |
| 08478 | 5 | DUF2235 domain-containing protein [Aromatoleum pumilum] |
| 08481 | 5 | DUF6306 domain-containing protein [Aromatoleum pumilum] |
| 00045 | 5 | sulfotransferase [Azoarcus indigens] |
| 07529 | 5 | DUF1788 domain-containing protein [Azoarcus olearius] |
| 02213 | 5 | TetR/AcrR family transcriptional regulator [Denitromonas halophilus] |
| 04481 | 5 | kynureninase [Azoarcus taiwanensis] |
| 09752 | 5 | lamin tail domain-containing protein [Denitromonas halophilus] |
| 01348 | 5 | SycD/LcrH family type III secretion system chaperone [Azoarcus rhizosphaerae] |
| 09687 | 5 | MULTISPECIES: hypothetical protein [Rhodocyclales] |
| 07075 | 5 | hypothetical protein [Nitrogeniibacter aestuarii] |
| 06074 | 5 | hypothetical protein [Denitromonas halophilus] |
| 09697 | 5 | hypothetical protein [Azoarcus halotolerans] |
| 10520 | 5 | DUF3144 domain-containing protein [Nitrogeniibacter aestuarii] |
| 01308 | 5 | hypothetical protein [Nitrogeniibacter aestuarii] |
| 06420 | 5 | Sbal_3080 family lipoprotein [Denitromonas halophilus] |
| 09724 | 5 | flagellar basal body P-ring formation chaperone FlgA [Azoarcus indigens] |
| 01278 | 5 | hypothetical protein [Azoarcus communis] |
| 10517 | 5 | MULTISPECIES: hypothetical protein [Niveibacterium] |

| | | |
|---|---|---|
| 01256 | 5 | TetR/AcrR family transcriptional regulator [Denitromonas halophilus] |
| 09755 | 5 | hypothetical protein [Azoarcus olearius] |
| 05239 | 5 | nuclear transport factor 2 family protein [Azoarcus communis] |
| 01249 | 5 | MULTISPECIES: hypothetical protein [Niveibacterium] |
| 01242 | 5 | NUDIX domain-containing protein [Azoarcus nasutitermitis] |
| 09764 | 5 | exosortase H [Aromatoleum pumilum] |
| 05181 | 5 | LuxR C-terminal-related transcriptional regulator [Denitromonas halophilus] |
| 09771 | 5 | ABC transporter substrate-binding protein [Nitrogeniibacter aestuarii] |
| 05191 | 5 | sensor histidine kinase [Azoarcus communis] |
| 07012 | 5 | hypothetical protein [Uliginosibacterium aquaticum] |
| 01221 | 5 | hypothetical protein [Uliginosibacterium gangwonense] |
| 01204 | 5 | COX aromatic rich motif-containing protein [Nitrogeniibacter aestuarii] |
| 05217 | 5 | HNH endonuclease [Aromatoleum aromaticum] |
| 01199 | 5 | FxDxF family PEP-CTERM protein [Thauera lacus] |
| 05229 | 5 | DUF3102 domain-containing protein [Thauera aminoaromatica] |
| 07085 | 5 | spermidine synthase [Denitromonas halophilus] |
| 09678 | 5 | hypothetical protein [Azoarcus olearius] |
| 01367 | 5 | proprotein convertase P-domain-containing protein [Denitromonas halophilus] |
| 01371 | 5 | hypothetical protein [Aromatoleum pumilum] |
| 01586 | 5 | MULTISPECIES: hypothetical protein [Betaproteobacteria] |
| 07214 | 5 | hypothetical protein [Denitromonas halophilus] |
| 07213 | 5 | hypothetical protein [Azoarcus olearius] |
| 06084 | 5 | MULTISPECIES: hypothetical protein [Niveibacterium] |
| 09185 | 5 | hypothetical protein [Niveibacterium sp. COAC-50] |
| 00195 | 5 | PEP-CTERM sorting domain-containing protein [Nitrogeniibacter aestuarii] |
| 01533 | 5 | hypothetical protein [Uliginosibacterium gangwonense] |
| 07188 | 5 | type II secretion system F family protein [Aromatoleum petrolei] |
| 04966 | 5 | hypothetical protein [Denitromonas halophilus] |
| 00197 | 5 | VOC family protein [Nitrogeniibacter aestuarii] |
| 07177 | 5 | DUF3987 domain-containing protein [Denitromonas halophilus] |
| 09593 | 5 | glycosyltransferase [Niveibacterium sp. COAC-50] |
| 01506 | 5 | hypothetical protein [Aromatoleum aromaticum] |
| 09601 | 5 | flagellar export chaperone FlgN [Azoarcus indigens] |
| 01489 | 5 | DUF2628 domain-containing protein [Azoarcus indigens] |
| 09606 | 5 | MULTISPECIES: phospholipid:lipid A palmitoyltransferase [Niveibacterium] |
| 09611 | 5 | DISARM system helicase DrmA [Aromatoleum toluclasticum] |
| 09612 | 5 | OsmC family protein [Candidatus Dactylopiibacterium carminicum] |
| 07130 | 5 | ISAs1 family transposase, partial [Azoarcus communis] |
| 10521 | 5 | hypothetical protein [Azoarcus indigens] |
| 01415 | 5 | MULTISPECIES: SIR2 family protein [Thauera] |
| 09659 | 5 | PRC-barrel domain-containing protein [Azoarcus indigens] |
| 07106 | 5 | O-antigen ligase family protein [Azospira oryzae] |
| 01402 | 5 | hypothetical protein [Denitromonas halophilus] |
| 01384 | 5 | MULTISPECIES: DUF4124 domain-containing protein [Aromatoleum] |
| 09790 | 5 | response regulator [Nitrogeniibacter aestuarii] |
| 01188 | 5 | type II toxin-antitoxin system PemK/MazF family toxin [Aromatoleum toluclasticum] |
| 01597 | 5 | class I SAM-dependent methyltransferase [Azoarcus olearius] |
| 01001 | 5 | NAD(P)/FAD-dependent oxidoreductase [Azoarcus rhizosphaerae] |
| 01054 | 5 | hypothetical protein [Azoarcus olearius] |
| 09883 | 5 | hypothetical protein [Aromatoleum toluclasticum] |
| 01049 | 5 | hypothetical protein [Aromatoleum pumilum] |
| 01038 | 5 | PIN domain-containing protein [Aromatoleum bremense] |
| 01034 | 5 | hypothetical protein [Aromatoleum buckelii] |
| 05377 | 5 | hypothetical protein [Azoarcus indigens] |
| 01016 | 5 | DUF6462 family protein [Azoarcus olearius] |
| 10488 | 5 | flagellar basal body protein [Azoarcus indigens] |
| 09908 | 5 | hypothetical protein [Azoarcus indigens] |
| 05390 | 5 | MULTISPECIES: hypothetical protein [Azospira] |
| 09909 | 5 | type I polyketide synthase [Azoarcus indigens] |
| 09915 | 5 | hypothetical protein [Aromatoleum aromaticum] |
| 05406 | 5 | ATP-binding protein [Aromatoleum toluclasticum] |
| 01187 | 5 | RES family NAD+ phosphorylase [Aromatoleum aromaticum] |
| 05409 | 5 | DUF554 domain-containing protein [Oryzomicrobium terrae] |
| 00993 | 5 | DUF4398 domain-containing protein [Niveibacterium microcysteis] |
| 10487 | 5 | glycoside hydrolase [Aromatoleum diolicum] |
| 06881 | 5 | hypothetical protein [Aromatoleum aromaticum] |
| 06877 | 5 | MULTISPECIES: type II toxin-antitoxin system PrlF family antitoxin [Aromatoleum] |
| 05432 | 5 | PhnD/SsuA/transferrin family substrate-binding protein [Thauera lacus] |
| 09952 | 5 | S8 family peptidase [Zoogloea oleivorans] |
| 00953 | 5 | hypothetical protein [Azoarcus indigens] |
| 09966 | 5 | DUF3732 domain-containing protein [Azoarcus halotolerans] |
| 09976 | 5 | hypothetical protein [Azoarcus communis] |
| 09989 | 5 | helix-turn-helix transcriptional regulator [Azoarcus nasutitermitis] |
| 00900 | 5 | hypothetical protein [Azoarcus indigens] |
| 09882 | 5 | hypothetical protein [Azoarcus indigens] |
| 06917 | 5 | hypothetical protein [Azoarcus indigens] |
| 09877 | 5 | saccharopine dehydrogenase family protein [Aromatoleum pumilum] |
| 01069 | 5 | cobalt-precorrin-6A reductase [Azoarcus communis] |
| 05243 | 5 | ferritin [Azoarcus communis] |
| 09795 | 5 | zf-HC2 domain-containing protein [Aromatoleum toluclasticum] |
| 01173 | 5 | hypothetical protein [Denitromonas halophilus] |
| 01148 | 5 | DUF1232 domain-containing protein [Azoarcus halotolerans] |
| 01142 | 5 | TonB-dependent receptor [Azoarcus sp. TTM-91] |
| 01131 | 5 | HEAT repeat domain-containing protein [Denitromonas halophilus] |
| 06956 | 5 | cupin domain-containing protein [Denitromonas halophilus] |
| 09829 | 5 | DUF115 domain-containing protein [Niveibacterium microcysteis] |
| 01116 | 5 | hypothetical protein [Denitromonas halophilus] |
| 06432 | 5 | endonuclease [Zoogloea oleivorans] |
| 06944 | 5 | hypothetical protein [Aromatoleum petrolei] |
| 06942 | 5 | hypothetical protein [Zoogloea dura] |
| 05302 | 5 | spirocyclase AveC family protein [Denitromonas halophilus] |
| 06049 | 5 | ParB/RepB/Spo0J family partition protein [Azoarcus indigens] |
| 09836 | 5 | NAD-dependent malic enzyme, partial [Uliginosibacterium gangwonense] |
| 01102 | 5 | 4Fe-4S dicluster domain-containing protein [Propionivibrio dicarboxylicus] |

| | | |
|---|---|---|
| 10502 | 5 | heparinase II/III family protein [Candidatus Dactylopiibacterium carminicum] |
| 05310 | 5 | antibiotic biosynthesis monooxygenase [Propionivibrio dicarboxylicus] |
| 01097 | 5 | type III secretion system chaperone [Azoarcus rhizosphaerae] |
| 09855 | 5 | DUF3861 domain-containing protein [Thauera linaloolentis] |
| 05317 | 5 | HAMP domain-containing histidine kinase [Denitromonas halophilus] |
| 05323 | 5 | DNA/RNA non-specific endonuclease [Aromatoleum aromaticum] |
| 05329 | 5 | hypothetical protein [Propionivibrio dicarboxylicus] |
| 09870 | 5 | type II toxin-antitoxin system VapC family toxin [Thauera butanivorans] |
| 05335 | 5 | hypothetical protein [Aromatoleum toluclasticum] |
| 09552 | 5 | transposase [Azoarcus communis] |
| 09575 | 5 | hypothetical protein [Zoogloea dura] |
| 01598 | 5 | M23 family metallopeptidase [Azoarcus communis] |
| 01845 | 5 | MULTISPECIES: SIR2 family protein [Rhodocyclales] |
| 09212 | 5 | EAL domain-containing protein [Nitrogeniibacter aestuarii] |
| 02089 | 5 | hypothetical protein [Azoarcus indigens] |
| 00144 | 5 | hypothetical protein [Azoarcus indigens] |
| 07393 | 5 | MULTISPECIES: hypothetical protein [Azospira] |
| 07391 | 5 | hypothetical protein [Aromatoleum evansii] |
| 04525 | 5 | hypothetical protein [Niveibacterium umoris] |
| 07387 | 5 | type I restriction endonuclease, partial [Zoogloea ramigera] |
| 01903 | 5 | hypothetical protein [Niveibacterium microcysteis] |
| 01898 | 5 | transporter substrate-binding domain-containing protein [Niveibacterium umoris] |
| 09359 | 5 | hypothetical protein, partial [Azoarcus sp. Aa7] |
| 09368 | 5 | DUF4384 domain-containing protein [Nitrogeniibacter aestuarii] |
| 09543 | 5 | HK97 gp10 family phage protein [Aromatoleum evansii] |
| 07369 | 5 | PAS domain-containing protein [Propionivibrio dicarboxylicus] |
| 04711 | 5 | hypothetical protein [Azoarcus indigens] |
| 06386 | 5 | hypothetical protein [Thauera linaloolentis] |
| 01882 | 5 | NF038104 family lipoprotein [Azoarcus halotolerans] |
| 01877 | 5 | DUF4880 domain-containing protein [Azoarcus indigens] |
| 01873 | 5 | radical SAM protein [Aromatoleum diolicum] |
| 07510 | 5 | hypothetical protein [Thauera butanivorans] |
| 09397 | 5 | AAA family ATPase [Azoarcus sp. KH32C] |
| 09205 | 5 | MULTISPECIES: hypothetical protein [Betaproteobacteria] |
| 10430 | 5 | AAA family ATPase [Azoarcus sp. KH32C] |
| 00153 | 5 | phosphatase PAP2 family protein [Candidatus Dactylopiibacterium carminicum] |
| 04679 | 5 | type II toxin-antitoxin system MqsR family toxin [Aromatoleum evansii] |
| 07409 | 5 | hypothetical protein [Azoarcus halotolerans] |
| 00141 | 5 | flagellar hook-length control protein FliK [Azoarcus indigens] |
| 09273 | 5 | RpiB/LacA/LacB family sugar-phosphate isomerase [Propionivibrio dicarboxylicus] |
| 06141 | 5 | hypothetical protein [Azoarcus communis] |
| 02059 | 5 | MULTISPECIES: hypothetical protein [Proteobacteria] |
| 07496 | 5 | PIG-L family deacetylase [Azoarcus indigens] |
| 00128 | 5 | Ti-type conjugative transfer relaxase TraA [Thauera chlorobenzoica] |
| 07492 | 5 | haloacid dehalogenase-like hydrolase [Azoarcus nasutitermitis] |
| 04580 | 5 | hypothetical protein [Nitrogeniibacter aestuarii] |
| 02025 | 5 | MULTISPECIES: hypothetical protein [Niveibacterium] |
| 02000 | 5 | hypothetical protein [Azoarcus indigens] |
| 01998 | 5 | hypothetical protein [Aromatoleum aromaticum] |
| 09268 | 5 | nickel transporter [Aromatoleum toluclasticum] |
| 09291 | 5 | N-acetylmuramoyl-L-alanine amidase [Azoarcus olearius] |
| 01920 | 5 | transposase, partial [Candidatus Dactylopiibacterium carminicum] |
| 04622 | 5 | endonuclease/exonuclease/phosphatase family protein [Azoarcus olearius] |
| 07442 | 5 | hypothetical protein [Azoarcus olearius] |
| 02060 | 5 | c-type cytochrome [Denitromonas halophilus] |
| 01962 | 5 | type I-E CRISPR-associated endoribonuclease Cas2e [Aromatoleum bremense] |
| 09310 | 5 | YbjN domain-containing protein [Niveibacterium microcysteis] |
| 06143 | 5 | FliM/FliN family flagellar motor switch protein [Azoarcus indigens] |
| 01939 | 5 | cyanophycinase [Niveibacterium microcysteis] |
| 01932 | 5 | hypothetical protein [Azoarcus communis] |
| 07422 | 5 | DUF2442 domain-containing protein [Thauera chlorobenzoica] |
| 00140 | 5 | Fis family transcriptional regulator [Azoarcus indigens] |
| 07499 | 5 | hypothetical protein [Azoarcus communis] |
| 09403 | 5 | threonine/serine exporter family protein [Candidatus Dactylopiibacterium carminicum] |
| 02113 | 5 | hypothetical protein [Azoarcus communis] |
| 09508 | 5 | hypothetical protein [Denitromonas halophilus] |
| 02135 | 5 | GtrA family protein [Azoarcus olearius] |
| 09483 | 5 | hypothetical protein [Aromatoleum diolicum] |
| 01706 | 5 | 2,5-dihydroxypyridine 5,6-dioxygenase [Azoarcus indigens] |
| 09489 | 5 | hypothetical protein [Azoarcus indigens] |
| 09497 | 5 | hypothetical protein [Azospira oryzae] |
| 04871 | 5 | DUF3368 domain-containing protein [Aromatoleum diolicum] |
| 04880 | 5 | hypothetical protein [Azoarcus indigens] |
| 07248 | 5 | hypothetical protein [Niveibacterium microcysteis] |
| 04884 | 5 | DUF5911 domain-containing protein [Azoarcus indigens] |
| 01673 | 5 | PD-(D/E)XK nuclease family protein [Thauera humireducens] |
| 01670 | 5 | MULTISPECIES: TrfB-related DNA-binding protein [Bacteria] |
| 01725 | 5 | hypothetical protein [Aromatoleum toluclasticum] |
| 09515 | 5 | hypothetical protein [Azoarcus communis] |
| 01650 | 5 | DUF4438 domain-containing protein [Azoarcus olearius] |
| 04904 | 5 | pectinesterase family protein [Uliginosibacterium aquaticum] |
| 09528 | 5 | BREX-3 system P-loop-containing protein BrxF [Thauera aromatica] |
| 09529 | 5 | DUF4743 domain-containing protein [Azoarcus halotolerans] |
| 00177 | 5 | NifB/NifX family molybdenum-iron cluster-binding protein [Rhodocyclus purpureus] |
| 01627 | 5 | hypothetical protein [Azoarcus indigens] |
| 02142 | 5 | rhodanese-like domain-containing protein [Azoarcus olearius] |
| 09537 | 5 | glycosyl hydrolase [Aromatoleum pumilum] |
| 01618 | 5 | phage portal protein [Uliginosibacterium gangwonense] |
| 09420 | 5 | EthD domain-containing protein [Aromatoleum toluclasticum] |
| 07268 | 5 | MULTISPECIES: glycosyltransferase family 25 protein [Thauera] |
| 09474 | 5 | MULTISPECIES: SPFH domain-containing protein [Azoarcus] |
| 09195 | 5 | hypothetical protein [Nitrogeniibacter aestuarii] |
| 07518 | 5 | MarR family winged helix-turn-helix transcriptional regulator [Denitromonas halophilus] |
| 02121 | 5 | hypothetical protein [Denitromonas halophilus] |

| | | |
|---|---|---|
| **09426** | 5 | MULTISPECIES: hypothetical protein [Proteobacteria] |
| **07332** | 5 | hypothetical protein [Nitrogeniibacter aestuarii] |
| **07323** | 5 | MULTISPECIES: xylose isomerase [Niveibacterium] |
| **09432** | 5 | DNA-binding protein [Thauera propionica] |
| **09437** | 5 | DUF2304 domain-containing protein [Aromatoleum aromaticum] |
| **04789** | 5 | Rha family transcriptional regulator [Azoarcus nasutitermitis] |
| **04792** | 5 | DUF3325 domain-containing protein [Azoarcus rhizosphaerae] |
| **01738** | 5 | polysaccharide deacetylase family protein [Propionivibrio dicarboxylicus] |
| **07313** | 5 | DUF1835 domain-containing protein [Aromatoleum aromaticum] |
| **07306** | 5 | 5-dehydro-4-deoxy-D-glucuronate isomerase [Candidatus Dactylopiibacterium carminicum] |
| **04801** | 5 | glycoside hydrolase family 127 protein [Zoogloea dura] |
| **10538** | 5 | hypothetical protein [Rhodocyclus purpureus] |
| **02127** | 5 | hypothetical protein [Denitromonas halophilus] |
| **09453** | 5 | hypothetical protein [Denitromonas halophilus] |
| **09461** | 5 | hypothetical protein [Azoarcus communis] |
| **04825** | 5 | putative DNA-binding domain-containing protein [Azoarcus olearius] |
| **07501** | 5 | hypothetical protein [Aromatoleum diolicum] |
| **06361** | 4 | DNA cytosine methyltransferase [Azoarcus communis] |
| **10483** | 4 | hypothetical protein [Aromatoleum toluclasticum] |
| **06499** | 4 | DUF2442 domain-containing protein [Aromatoleum toluclasticum] |
| **10435** | 4 | MULTISPECIES: hypothetical protein [Azospira] |
| **06450** | 4 | hypothetical protein [Azoarcus halotolerans] |
| **10625** | 4 | hypothetical protein [Rhodocyclus purpureus] |
| **10441** | 4 | hypothetical protein [Aromatoleum evansii] |
| **06277** | 4 | hypothetical protein [Azoarcus halotolerans] |
| **06394** | 4 | carbon-phosphorus lyase complex subunit PhnI [Azoarcus nasutitermitis] |
| **10522** | 4 | capsular biosynthesis protein [Azoarcus halotolerans] |
| **06319** | 4 | MULTISPECIES: hypothetical protein [Betaproteobacteria] |
| **06308** | 4 | DUF4242 domain-containing protein [Azoarcus sp. DD4] |
| **06398** | 4 | hypothetical protein [Aromatoleum pumilum] |
| **06300** | 4 | helix-turn-helix domain-containing protein [Niveibacterium microcysteis] |
| **06469** | 4 | glycosyltransferase [Azoarcus communis] |
| **06422** | 4 | hypothetical protein [Denitromonas halophilus] |
| **06337** | 4 | hypothetical protein [Aromatoleum aromaticum] |
| **06471** | 4 | MULTISPECIES: hypothetical protein [Betaproteobacteria] |
| **10574** | 4 | hypothetical protein [Niveibacterium microcysteis] |
| **10473** | 4 | phosphonoacetaldehyde hydrolase [Azoarcus halotolerans] |
| **10572** | 4 | amino acid adenylation domain-containing protein [Niveibacterium sp. COAC-50] |
| **06345** | 4 | hypothetical protein [Azoarcus olearius] |
| **06431** | 4 | hypothetical protein [Niveibacterium microcysteis] |
| **06488** | 4 | hypothetical protein [Aromatoleum toluclasticum] |
| **06433** | 4 | hypothetical protein [Azoarcus indigens] |
| **06377** | 4 | phage tail sheath subtilisin-like domain-containing protein [Azoarcus indigens] |
| **10545** | 4 | hypothetical protein [Uliginosibacterium aquaticum] |
| **10560** | 4 | hypothetical protein [Nitrogeniibacter aestuarii] |
| **06356** | 4 | molecular chaperone [Azoarcus communis] |
| **08178** | 4 | hypothetical protein [Azoarcus communis] |
| **09505** | 4 | hypothetical protein [Denitromonas halophilus] |
| **10424** | 4 | hypothetical protein [Zoogloea oleivorans] |
| **08958** | 4 | MULTISPECIES: hypothetical protein [Niveibacterium] |
| **08971** | 4 | holin family protein [Aromatoleum toluclasticum] |
| **07650** | 4 | DUF1214 domain-containing protein [Niveibacterium microcysteis] |
| **07649** | 4 | MULTISPECIES: hypothetical protein [Niveibacterium] |
| **07634** | 4 | glycosyltransferase [Aromatoleum evansii] |
| **07628** | 4 | hypothetical protein [Azoarcus olearius] |
| **07619** | 4 | MASE1 domain-containing protein [Aromatoleum diolicum] |
| **09019** | 4 | mucoidy inhibitor MuiA family protein [Azoarcus rhizosphaerae] |
| **09030** | 4 | MBL fold metallo-hydrolase [Azoarcus nasutitermitis] |
| **09042** | 4 | poly-beta-1,6-N-acetyl-D-glucosamine biosynthesis protein PgaD [Nitrogeniibacter aestuarii] |
| **09060** | 4 | AAA domain-containing protein [Aromatoleum toluvorans] |
| **07592** | 4 | M48 family metallopeptidase [Azoarcus indigens] |
| **09086** | 4 | hypothetical protein [Azoarcus nasutitermitis] |
| **09096** | 4 | MULTISPECIES: hypothetical protein [Proteobacteria] |
| **09104** | 4 | recombination protein NinB [Azoarcus indigens] |
| **07564** | 4 | hypothetical protein [Azoarcus nasutitermitis] |
| **07662** | 4 | TetR/AcrR family transcriptional regulator [Nitrogeniibacter aestuarii] |
| **07666** | 4 | HAMP domain-containing histidine kinase [Aromatoleum toluclasticum] |
| **07557** | 4 | AAA family ATPase [Aromatoleum bremense] |
| **08947** | 4 | S53 family peptidase [Niveibacterium microcysteis] |
| **08837** | 4 | hypothetical protein [Denitromonas halophilus] |
| **08838** | 4 | DUF1194 domain-containing protein [Denitromonas halophilus] |
| **08850** | 4 | glycosyltransferase family 2 protein [Oryzomicrobium terrae] |
| **08870** | 4 | SRPBCC family protein [Aromatoleum aromaticum] |
| **07717** | 4 | DUF126 domain-containing protein [Aromatoleum diolicum] |
| **07716** | 4 | hypothetical protein [Aromatoleum tolulyticum] |
| **08891** | 4 | DUF1302 family protein, partial [Aromatoleum diolicum] |
| **07705** | 4 | hydroxyethylthiazole kinase [Propionivibrio dicarboxylicus] |
| **08906** | 4 | PEP-CTERM sorting domain-containing protein [Azoarcus sp. DD4] |
| **08911** | 4 | flagellar basal body rod protein FlgB [Zoogloea dura] |
| **08923** | 4 | methylglyoxal synthase [Niveibacterium microcysteis] |
| **08927** | 4 | hypothetical protein [Azoarcus indigens] |
| **07689** | 4 | hypothetical protein [Nitrogeniibacter aestuarii] |
| **08939** | 4 | type I-U CRISPR-associated RAMP protein Csb1/Cas7u [Aromatoleum toluolicum] |
| **08942** | 4 | hypothetical protein [Azospira oryzae] |
| **09107** | 4 | hypothetical protein [Azoarcus indigens] |
| **07554** | 4 | hypothetical protein [Azoarcus olearius] |
| **10419** | 4 | DUF2313 domain-containing protein [Azoarcus indigens] |
| **09244** | 4 | MULTISPECIES: methenyltetrahydromethanopterin cyclohydrolase [Thauera] |
| **07467** | 4 | Bro-N domain-containing protein [Candidatus Dactylopiibacterium carminicum] |
| **07452** | 4 | hypothetical protein [Aromatoleum toluolicum] |
| **07447** | 4 | hypothetical protein [Thauera chlorobenzoica] |
| **07444** | 4 | hypothetical protein [Thauera aminoaromatica] |
| **09299** | 4 | hypothetical protein [Azospira oryzae] |
| **09301** | 4 | GNAT family N-acetyltransferase [Candidatus Dactylopiibacterium carminicum] |

| | | |
|---|---|---|
| 09313 | 4 | cytochrome-c oxidase [Aromatoleum pumilum] |
| 07425 | 4 | EutN/CcmL family microcompartment protein [Azoarcus nasutitermitis] |
| 07424 | 4 | hypothetical protein [Azoarcus communis] |
| 09339 | 4 | gamma-mobile-trio recombinase GmtY [Aromatoleum petrolei] |
| 07419 | 4 | hypothetical protein [Aromatoleum toluclasticum] |
| 09343 | 4 | hypothetical protein [Aromatoleum toluolicum] |
| 07412 | 4 | histidine kinase [Niveibacterium microcysteis] |
| 07411 | 4 | GNAT family N-acetyltransferase [Niveibacterium microcysteis] |
| 07401 | 4 | hypothetical protein [Azoarcus sp. KH32C] |
| 07477 | 4 | Ig-like domain-containing protein [Propionivibrio dicarboxylicus] |
| 07484 | 4 | PepSY domain-containing protein [Azoarcus indigens] |
| 09139 | 4 | hypothetical protein [Denitromonas halophilus] |
| 09235 | 4 | DUF1611 domain-containing protein [Denitromonas halophilus] |
| 07543 | 4 | MULTISPECIES: hypothetical protein [Azospira] |
| 07542 | 4 | HDOD domain-containing protein [Nitrogeniibacter aestuarii] |
| 09157 | 4 | ATP-binding protein [Thauera aromatica] |
| 07534 | 4 | hypothetical protein [Azoarcus halotolerans] |
| 09173 | 4 | hypothetical protein [Denitromonas halophilus] |
| 09188 | 4 | DNA cytosine methyltransferase [Azoarcus nasutitermitis] |
| 09189 | 4 | type VI secretion system protein TssA [Azoarcus indigens] |
| 07522 | 4 | hypothetical protein [Denitromonas halophilus] |
| 07514 | 4 | DUF3037 domain-containing protein [Azoarcus communis] |
| 09207 | 4 | hypothetical protein [Nitrogeniibacter aestuarii] |
| 07508 | 4 | transposase [Azoarcus sp. DD4] |
| 09216 | 4 | universal stress protein [Aromatoleum toluclasticum] |
| 09222 | 4 | PAS domain-containing protein [Aromatoleum toluolicum] |
| 07497 | 4 | type II secretion system minor pseudopilin GspH [Azoarcus indigens] |
| 09229 | 4 | AraC family transcriptional regulator [Denitromonas sp. IR12] |
| 08829 | 4 | hypothetical protein [Azoarcus halotolerans] |
| 07743 | 4 | RNA ligase family protein [Thauera phenylacetica] |
| 07748 | 4 | MFS transporter, partial [Thauera phenylacetica] |
| 08022 | 4 | thioredoxin family protein [Niveibacterium microcysteis] |
| 08013 | 4 | DUF1851 domain-containing protein [Nitrogeniibacter aestuarii] |
| 08005 | 4 | S9 family peptidase [Niveibacterium microcysteis] |
| 08001 | 4 | MULTISPECIES: hypothetical protein [Proteobacteria] |
| 08340 | 4 | HlyD family efflux transporter periplasmic adaptor subunit, partial [Aromatoleum toluolicum] |
| 08342 | 4 | hypothetical protein [Azoarcus communis] |
| 08346 | 4 | hypothetical protein [Aromatoleum toluclasticum] |
| 08362 | 4 | GIY-YIG nuclease family protein [Azospira oryzae] |
| 08367 | 4 | nucleotidyltransferase domain-containing protein [Aromatoleum pumilum] |
| 08372 | 4 | hypothetical protein [Azoarcus communis] |
| 07963 | 4 | hypothetical protein [Azoarcus sp. Aa7] |
| 08398 | 4 | DUF6527 family protein [Aromatoleum diolicum] |
| 07958 | 4 | OFA family MFS transporter [Azoarcus indigens] |
| 08412 | 4 | hypothetical protein [Thauera lacus] |
| 07949 | 4 | ABC transporter permease [Azoarcus halotolerans] |
| 07936 | 4 | MbcA/ParS/Xre antitoxin family protein [Aromatoleum diolicum] |
| 08014 | 4 | RNA 2'-phosphotransferase [Azoarcus communis] |
| 08023 | 4 | hypothetical protein [Aromatoleum aromaticum] |
| 08818 | 4 | Mu-like prophage major head subunit gpT family protein [Thauera humireducens] |
| 08026 | 4 | DUF6398 domain-containing protein [Aromatoleum bremense] |
| 08181 | 4 | IS1634 family transposase, partial [Azoarcus communis] |
| 08154 | 4 | hypothetical protein [Azoarcus nasutitermitis] |
| 08202 | 4 | type II toxin-antitoxin system prevent-host-death family antitoxin [Aromatoleum pumilum] |
| 08140 | 4 | peptidoglycan-binding protein [Azoarcus indigens] |
| 08129 | 4 | DUF1566 domain-containing protein [Thauera butanivorans] |
| 08126 | 4 | hypothetical protein [Azoarcus communis] |
| 08117 | 4 | poly-beta-1,6 N-acetyl-D-glucosamine export porin PgaA [Nitrogeniibacter aestuarii] |
| 08230 | 4 | mechanosensitive ion channel [Nitrogeniibacter aestuarii] |
| 08101 | 4 | septal ring lytic transglycosylase RlpA family protein [Aromatoleum aromaticum] |
| 08075 | 4 | SAVED domain-containing protein [Aromatoleum aromaticum] |
| 08260 | 4 | hypothetical protein [Denitromonas halophilus] |
| 08058 | 4 | hypothetical protein [Aromatoleum aromaticum] |
| 08052 | 4 | phosphatase PAP2 family protein [Aromatoleum toluclasticum] |
| 08302 | 4 | PEP-CTERM sorting domain-containing protein [Azoarcus sp. TTM-91] |
| 08305 | 4 | hypothetical protein [Thauera humireducens] |
| 07933 | 4 | DUF4276 family protein [Aromatoleum toluvorans] |
| 06269 | 4 | DUF2795 domain-containing protein [Azoarcus indigens] |
| 07923 | 4 | hypothetical protein [Denitromonas halophilus] |
| 07914 | 4 | hypothetical protein [Aromatoleum evansii] |
| 08678 | 4 | DUF1636 domain-containing protein [Azoarcus indigens] |
| 07803 | 4 | phage major capsid protein [Aromatoleum pumilum] |
| 08723 | 4 | hypothetical protein [Aromatoleum toluclasticum] |
| 07795 | 4 | hypothetical protein [Thauera propionica] |
| 07786 | 4 | TonB-dependent receptor plug domain-containing protein [Thauera phenolivorans] |
| 07773 | 4 | hypothetical protein [Thauera butanivorans] |
| 08750 | 4 | SHOCT domain-containing protein [Azoarcus olearius] |
| 07767 | 4 | PA14 domain-containing protein [Nitrogeniibacter aestuarii] |
| 08772 | 4 | DUF6079 family protein [Thauera aromatica] |
| 07760 | 4 | IPT/TIG domain-containing protein, partial [Uliginosibacterium gangwonense] |
| 08795 | 4 | ABC transporter substrate-binding protein [Azoarcus indigens] |
| 08798 | 4 | helix-turn-helix domain-containing protein [Azoarcus indigens] |
| 08800 | 4 | hypothetical protein [Azoarcus indigens] |
| 08803 | 4 | hypothetical protein [Azoarcus indigens] |
| 08816 | 4 | catalase family peroxidase [Denitromonas halophilus] |
| 08662 | 4 | MULTISPECIES: hypothetical protein [Niveibacterium] |
| 07851 | 4 | cytochrome c [Azoarcus indigens] |
| 08627 | 4 | hypothetical protein [Aromatoleum toluvorans] |
| 08597 | 4 | H4MPT-linked C1 transfer pathway protein [Aromatoleum toluvorans] |
| 08511 | 4 | BMC domain-containing protein [Azoarcus rhizosphaerae] |
| 08515 | 4 | MULTISPECIES: hypothetical protein [Betaproteobacteria] |
| 08520 | 4 | hypothetical protein [Aromatoleum toluvorans] |
| 07904 | 4 | cytochrome c [Nitrogeniibacter aestuarii] |
| 07903 | 4 | DEAD/DEAH box helicase [Azoarcus halotolerans] |

| | | |
|---|---|---|
| 08586 | 4 | MULTISPECIES: helix-turn-helix domain-containing protein [Rhodocyclales] |
| 08600 | 4 | ATP-grasp domain-containing protein [Aromatoleum toluvorans] |
| 08625 | 4 | formate hydrogenlyase maturation HycH family protein [Niveibacterium microcysteis] |
| 08603 | 4 | DUF4214 domain-containing protein [Aromatoleum bremense] |
| 08604 | 4 | putative sulfate exporter family transporter, partial [Aromatoleum diolicum] |
| 08607 | 4 | N-acetyltransferase [Aromatoleum toluolicum] |
| 07865 | 4 | hypothetical protein [Aromatoleum aromaticum] |
| 08608 | 4 | hypothetical protein [Aromatoleum petrolei] |
| 07861 | 4 | hypothetical protein [Azoarcus olearius] |
| 09350 | 4 | TAXI family TRAP transporter solute-binding subunit, partial [Denitromonas halophilus] |
| 09363 | 4 | SLC13 family permease [Propionivibrio dicarboxylicus] |
| 07382 | 4 | PEP-CTERM sorting domain-containing protein [Denitromonas sp. IR12] |
| 10024 | 4 | MULTISPECIES: hypothetical protein [Niveibacterium] |
| 10029 | 4 | hypothetical protein [Niveibacterium microcysteis] |
| 10033 | 4 | hypothetical protein [Candidatus Dactylopiibacterium carminicum] |
| 06827 | 4 | Cro/CI family transcriptional regulator [Azoarcus rhizosphaerae] |
| 10045 | 4 | class I SAM-dependent methyltransferase [Azoarcus olearius] |
| 06819 | 4 | mannonate dehydratase [Candidatus Dactylopiibacterium carminicum] |
| 10072 | 4 | MULTISPECIES: fructose-specific PTS transporter subunit EIIC [Niveibacterium] |
| 10085 | 4 | hypothetical protein [Azoarcus indigens] |
| 10088 | 4 | hypothetical protein [Aromatoleum evansii] |
| 06797 | 4 | MULTISPECIES: conjugal transfer protein TraM [Aromatoleum] |
| 06788 | 4 | DUF1015 domain-containing protein [Candidatus Dactylopiibacterium carminicum] |
| 06787 | 4 | hypothetical protein [Azoarcus rhizosphaerae] |
| 06784 | 4 | heme utilization protein [Propionivibrio dicarboxylicus] |
| 10131 | 4 | MULTISPECIES: formylmethanofuran--tetrahydromethanopterin N-formyltransferase [Thauera] |
| 06764 | 4 | SPOR domain-containing protein [Azoarcus communis] |
| 10138 | 4 | DUF302 domain-containing protein [Aromatoleum diolicum] |
| 10027 | 4 | hypothetical protein [Niveibacterium sp. COAC-50] |
| 10018 | 4 | MULTISPECIES: helix-turn-helix domain-containing protein [Zoogloea] |
| 06911 | 4 | TrfB-related DNA-binding protein [Aromatoleum aromaticum] |
| 10012 | 4 | cyclin-dependent kinase inhibitor 3 family protein [Zoogloea dura] |
| 09892 | 4 | hypothetical protein [Thauera aminoaromatica] |
| 09904 | 4 | DUF3567 family protein [Oryzomicrobium terrae] |
| 06901 | 4 | hypothetical protein [Azospira oryzae] |
| 09928 | 4 | DNA-binding protein [Azoarcus communis] |
| 06879 | 4 | hypothetical protein [Azoarcus nasutitermitis] |
| 06874 | 4 | DUF2971 domain-containing protein [Aromatoleum pumilum] |
| 09941 | 4 | hypothetical protein [Denitromonas halophilus] |
| 06867 | 4 | phage tail assembly protein [Denitromonas sp. IR12] |
| 06865 | 4 | MULTISPECIES: hypothetical protein [Niveibacterium] |
| 09964 | 4 | lipoprotein [Azoarcus olearius] |
| 06860 | 4 | 5-methylcytosine-specific restriction endonuclease system specificity protein McrC [Aromatoleum aromaticum] |
| 09980 | 4 | hypothetical protein [Aromatoleum bremense] |
| 09985 | 4 | ABC transporter permease subunit [Aromatoleum pumilum] |
| 06847 | 4 | hypothetical protein [Aromatoleum toluolicum] |
| 10006 | 4 | hypothetical protein [Niveibacterium umoris] |
| 10156 | 4 | Ig-like domain-containing protein, partial [Azoarcus halotolerans] |
| 10174 | 4 | rod-binding protein [Azoarcus indigens] |
| 10176 | 4 | DUF805 domain-containing protein [Candidatus Dactylopiibacterium carminicum] |
| 10180 | 4 | hypothetical protein [Aromatoleum toluclasticum] |
| 06654 | 4 | MULTISPECIES: D-ribose pyranase [Niveibacterium] |
| 06643 | 4 | hypothetical protein [Thauera phenylacetica] |
| 06633 | 4 | hypothetical protein [Azoarcus olearius] |
| 06628 | 4 | arylamine N-acetyltransferase [Azoarcus olearius] |
| 06619 | 4 | hypothetical protein [Aromatoleum toluclasticum] |
| 06574 | 4 | hypothetical protein [Azoarcus communis] |
| 10382 | 4 | hypothetical protein [Azospira oryzae] |
| 06560 | 4 | PEP-CTERM sorting domain-containing protein [Nitrogeniibacter aestuarii] |
| 10385 | 4 | siderophore-iron reductase FhuF [Azoarcus indigens] |
| 06547 | 4 | hypothetical protein [Aromatoleum toluclasticum] |
| 06540 | 4 | aminopeptidase [Candidatus Dactylopiibacterium carminicum] |
| 10393 | 4 | hypothetical protein [Aromatoleum aromaticum] |
| 06524 | 4 | MULTISPECIES: hypothetical protein [Niveibacterium] |
| 10412 | 4 | sulfurtransferase [Denitromonas halophilus] |
| 06513 | 4 | GNAT family N-acetyltransferase [Niveibacterium microcysteis] |
| 10323 | 4 | alginate export family protein [Thauera aminoaromatica] |
| 06671 | 4 | hypothetical protein [Aromatoleum aromaticum] |
| 10312 | 4 | MULTISPECIES: hypothetical protein [Thauera] |
| 10224 | 4 | MULTISPECIES: zinc ribbon domain-containing protein [Niveibacterium] |
| 06749 | 4 | galactokinase [Niveibacterium microcysteis] |
| 06741 | 4 | L-arabinose isomerase [Niveibacterium microcysteis] |
| 06740 | 4 | UDP-glucose--hexose-1-phosphate uridylyltransferase [Uliginosibacterium gangwonense] |
| 06733 | 4 | hypothetical protein [Niveibacterium microcysteis] |
| 06729 | 4 | hypothetical protein [Azoarcus halotolerans] |
| 10216 | 4 | hypothetical protein [Denitromonas halophilus] |
| 06721 | 4 | MaoC family dehydratase N-terminal domain-containing protein [Aromatoleum toluclasticum] |
| 06684 | 4 | hypothetical protein [Thauera terpenica] |
| 06704 | 4 | UvrD-helicase domain-containing protein [Aromatoleum evansii] |
| 06700 | 4 | MULTISPECIES: TraK family protein [Aromatoleum] |
| 10291 | 4 | MULTISPECIES: hypothetical protein [Thauera] |
| 06695 | 4 | hypothetical protein [Aromatoleum toluvorans] |
| 06693 | 4 | hypothetical protein [Aromatoleum toluclasticum] |
| 06689 | 4 | ethanolamine utilization cob(I)yrinic acid a,c-diamide adenosyltransferase EutT [Azoarcus nasutitermitis] |
| 06904 | 4 | exonuclease domain-containing protein [Azoarcus indigens] |
| 06912 | 4 | hypothetical protein [Azoarcus rhizosphaerae] |
| 09367 | 4 | hypothetical protein [Aromatoleum pumilum] |
| 07276 | 4 | site-specific integrase [Aromatoleum evansii] |
| 09485 | 4 | phosphonate C-P lyase system protein PhnH [Azoarcus nasutitermitis] |
| 07259 | 4 | methyltransferase domain-containing protein [Azoarcus indigens] |
| 09503 | 4 | hypothetical protein [Denitromonas halophilus] |
| 07250 | 4 | YihY/virulence factor BrkB family protein [Aromatoleum diolicum] |
| 07245 | 4 | YccF domain-containing protein [Niveibacterium microcysteis] |
| 07235 | 4 | ATP-binding protein [Azoarcus nasutitermitis] |

| | | |
|---|---|---|
| 09545 | 4 | DUF932 domain-containing protein [Niveibacterium microcysteis] |
| 07232 | 4 | hypothetical protein [Azoarcus communis] |
| 07227 | 4 | hypothetical protein [Azoarcus communis] |
| 09557 | 4 | DUF1538 domain-containing protein [Aromatoleum pumilum] |
| 09567 | 4 | tyrosine-type recombinase/integrase [Zoogloea dura] |
| 07202 | 4 | hypothetical protein, partial [Azoarcus communis] |
| 07201 | 4 | ankyrin repeat domain-containing protein, partial [Azoarcus communis] |
| 09573 | 4 | hypothetical protein [Azoarcus communis] |
| 07192 | 4 | hypothetical protein [Azoarcus indigens] |
| 09476 | 4 | DNA circularization N-terminal domain-containing protein [Azoarcus indigens] |
| 09468 | 4 | hypothetical protein [Azoarcus nasutitermitis] |
| 09879 | 4 | diguanylate cyclase [Azoarcus communis] |
| 09462 | 4 | PEP-CTERM sorting domain-containing protein [Azoarcus communis] |
| 09370 | 4 | phosphoglycerate mutase family protein [Azoarcus indigens] |
| 09375 | 4 | DUF2345 domain-containing protein, partial [Aromatoleum tolulyticum] |
| 07362 | 4 | hypothetical protein [Aromatoleum toluclasticum] |
| 07356 | 4 | DUF4405 domain-containing protein [Nitrogeniibacter aestuarii] |
| 07355 | 4 | CHASE2 domain-containing protein [Nitrogeniibacter aestuarii] |
| 07352 | 4 | hypothetical protein [Nitrogeniibacter aestuarii] |
| 09405 | 4 | NADAR family protein [Uliginosibacterium gangwonense] |
| 07337 | 4 | M23 family metallopeptidase [Nitrogeniibacter aestuarii] |
| 09421 | 4 | hypothetical protein [Aromatoleum aromaticum] |
| 07333 | 4 | hypothetical protein [Nitrogeniibacter aestuarii] |
| 07322 | 4 | hypothetical protein [Aromatoleum pumilum] |
| 07315 | 4 | hypothetical protein [Aromatoleum toluclasticum] |
| 09444 | 4 | DUF2442 domain-containing protein [Aromatoleum pumilum] |
| 09450 | 4 | hypothetical protein [Aromatoleum aromaticum] |
| 07301 | 4 | alpha/beta fold hydrolase [Candidatus Dactylopiibacterium carminicum] |
| 07182 | 4 | AAA family ATPase [Azoarcus halotolerans] |
| 07176 | 4 | DUF4236 domain-containing protein [Denitromonas halophilus] |
| 07124 | 4 | glycosyltransferase family 2 protein [Azoarcus olearius] |
| 09651 | 4 | hypothetical protein [Azoarcus halotolerans] |
| 09782 | 4 | hypothetical protein [Aromatoleum tolulyticum] |
| 09786 | 4 | DNA-binding protein [Candidatus Dactylopiibacterium carminicum] |
| 07000 | 4 | DUF3015 domain-containing protein [Zoogloea ramigera] |
| 09791 | 4 | ferredoxin [Niveibacterium microcysteis] |
| 06983 | 4 | MULTISPECIES: hypothetical protein [Aromatoleum] |
| 09803 | 4 | asparagine synthase-related protein, partial [Thauera propionica] |
| 09807 | 4 | hypothetical protein [Aromatoleum aromaticum] |
| 09821 | 4 | DUF4902 domain-containing protein [Nitrogeniibacter aestuarii] |
| 06948 | 4 | helix-hairpin-helix domain-containing protein [Azoarcus communis] |
| 09830 | 4 | RebB family R body protein [Uliginosibacterium gangwonense] |
| 09832 | 4 | hypothetical protein [Azoarcus communis] |
| 06941 | 4 | TniQ family protein [Aromatoleum pumilum] |
| 06939 | 4 | hypothetical protein [Azospira oryzae] |
| 09853 | 4 | BMC domain-containing protein [Azoarcus nasutitermitis] |
| 06927 | 4 | dsDNA nuclease domain-containing protein [Denitromonas halophilus] |
| 07013 | 4 | hypothetical protein [Denitromonas halophilus] |
| 07022 | 4 | CRISPR-associated protein Csx16 [Niveibacterium sp. COAC-50] |
| 09751 | 4 | NAD(P)H-dependent oxidoreductase [Niveibacterium microcysteis] |
| 09704 | 4 | hypothetical protein [Niveibacterium microcysteis] |
| 09655 | 4 | hypothetical protein [Nitrogeniibacter aestuarii] |
| 09656 | 4 | PEP-CTERM sorting domain-containing protein [Nitrogeniibacter mangrovi] |
| 09669 | 4 | hypothetical protein [Aromatoleum toluclasticum] |
| 07092 | 4 | filamentous hemagglutinin N-terminal domain-containing protein [Nitrogeniibacter aestuarii] |
| 09693 | 4 | hypothetical protein [Thauera humireducens] |
| 09698 | 4 | hypothetical protein [Zoogloea dura] |
| 09709 | 4 | IS5 family transposase [Aromatoleum evansii] |
| 09750 | 4 | PEP-CTERM sorting domain-containing protein [Nitrogeniibacter aestuarii] |
| 09720 | 4 | NUDIX domain-containing protein [Uliginosibacterium aquaticum] |
| 09723 | 4 | MULTISPECIES: hypothetical protein [Niveibacterium] |
| 07051 | 4 | beta-galactosidase [Niveibacterium microcysteis] |
| 09736 | 4 | hypothetical protein [Niveibacterium microcysteis] |
| 09743 | 4 | hypothetical protein [Nitrogeniibacter aestuarii] |
| 07045 | 4 | hypothetical protein [Azoarcus halotolerans] |
| 05314 | 4 | hypothetical protein [Denitromonas halophilus] |
| 10627 | 4 | LLM class flavin-dependent oxidoreductase [Aromatoleum toluclasticum] |
| 03328 | 4 | hypothetical protein [Aromatoleum toluclasticum] |
| 05536 | 4 | hypothetical protein [Aromatoleum diolicum] |
| 00561 | 4 | DUF202 domain-containing protein [Azoarcus indigens] |
| 00563 | 4 | transposase [Azoarcus indigens] |
| 01684 | 4 | glycosyltransferase [Azoarcus indigens] |
| 01680 | 4 | TnsA endonuclease N-terminal domain-containing protein [Azoarcus communis] |
| 03292 | 4 | DUF2190 family protein [Azoarcus communis] |
| 05402 | 4 | AcaB family transcriptional regulator [Azoarcus halotolerans] |
| 05388 | 4 | DUF1566 domain-containing protein [Azoarcus sp. L1K30] |
| 03310 | 4 | ATP-grasp domain-containing protein [Aromatoleum toluvorans] |
| 00580 | 4 | hypothetical protein [Azoarcus indigens] |
| 05365 | 4 | formylmethanofuran dehydrogenase subunit A [Aromatoleum toluvorans] |
| 05360 | 4 | MULTISPECIES: hypothetical protein [Thauera] |
| 03321 | 4 | hypothetical protein [Aromatoleum petrolei] |
| 00584 | 4 | substrate-binding domain-containing protein [Azoarcus indigens] |
| 04434 | 4 | hypothetical protein [Candidatus Dactylopiibacterium carminicum] |
| 00589 | 4 | hypothetical protein [Nitrogeniibacter aestuarii] |
| 00592 | 4 | DUF4105 domain-containing protein [Propionivibrio dicarboxylicus] |
| 05330 | 4 | aldehyde dehydrogenase family protein [Azospira oryzae] |
| 03341 | 4 | hypothetical protein [Azoarcus olearius] |
| 05328 | 4 | dihydroxyacetone kinase subunit DhaK [Propionivibrio dicarboxylicus] |
| 03281 | 4 | DUF937 domain-containing protein [Aromatoleum aromaticum] |
| 01690 | 4 | general secretion pathway protein GspC [Azoarcus indigens] |
| 03270 | 4 | EAL domain-containing protein, partial [Aromatoleum toluclasticum] |
| 00521 | 4 | pilin [Azoarcus sp. L1K30] |
| 01729 | 4 | DUF4157 domain-containing protein [Denitromonas halophilus] |
| 00510 | 4 | (5-formylfuran-3-yl)methyl phosphate synthase [Aromatoleum toluvorans] |

| | | |
|---|---|---|
| 01724 | 4 | hypothetical protein [Denitromonas halophilus] |
| 05509 | 4 | hypothetical protein [Azoarcus halotolerans] |
| 03240 | 4 | helix-turn-helix domain-containing protein [Denitromonas halophilus] |
| 05493 | 4 | DUF4157 domain-containing protein [Denitromonas halophilus] |
| 05490 | 4 | hypothetical protein [Azoarcus olearius] |
| 01709 | 4 | hypothetical protein [Azoarcus communis] |
| 05469 | 4 | type II toxin-antitoxin system VapC family toxin [Azoarcus indigens] |
| 01697 | 4 | MULTISPECIES: TetR/AcrR family transcriptional regulator [Betaproteobacteria] |
| 01708 | 4 | leucine-rich repeat-containing serine/threonine-protein kinase [Azoarcus sp. DD4] |
| 01704 | 4 | prepilin-type N-terminal cleavage/methylation domain-containing protein [Azoarcus indigens] |
| 03256 | 4 | GNAT family N-acetyltransferase [Azoarcus nasutitermitis] |
| 00533 | 4 | hypothetical protein [Azoarcus indigens] |
| 01700 | 4 | MULTISPECIES: hypothetical protein [Thauera] |
| 03258 | 4 | glycosyltransferase [Uliginosibacterium gangwonense] |
| 00548 | 4 | RNA-directed DNA polymerase [Zoogloea sp. G-4-1-14] |
| 03260 | 4 | NERD domain-containing protein [Aromatoleum pumilum] |
| 00593 | 4 | PhnD/SsuA/transferrin family substrate-binding protein [Zoogloea oleivorans] |
| 01628 | 4 | hypothetical protein [Azoarcus indigens] |
| 03356 | 4 | isocitrate lyase/PEP mutase family protein [Denitromonas sp. IR12] |
| 05168 | 4 | cyanophycinase [Niveibacterium microcysteis] |
| 05221 | 4 | hypothetical protein [Aromatoleum evansii] |
| 00636 | 4 | hypothetical protein [Aromatoleum buckelii] |
| 03465 | 4 | SDR family oxidoreductase [Aromatoleum petrolei] |
| 00642 | 4 | MULTISPECIES: hypothetical protein [Thauera] |
| 01532 | 4 | MAPEG family protein [Niveibacterium microcysteis] |
| 05196 | 4 | methyltransferase domain-containing protein [Azoarcus olearius] |
| 05180 | 4 | hypothetical protein [Denitromonas halophilus] |
| 03510 | 4 | hypothetical protein [Candidatus Dactylopiibacterium carminicum] |
| 05166 | 4 | carboxypeptidase regulatory-like domain-containing protein, partial [Thauera phenylacetica] |
| 01562 | 4 | GFA family protein [Nitrogeniibacter aestuarii] |
| 05165 | 4 | response regulator [Aromatoleum evansii] |
| 03517 | 4 | SPFH domain-containing protein [Niveibacterium microcysteis] |
| 03526 | 4 | MBL fold metallo-hydrolase [Zoogloea ramigera] |
| 05145 | 4 | glycoside hydrolase family 108 protein [Azoarcus nasutitermitis] |
| 03543 | 4 | GNAT family N-acetyltransferase [Niveibacterium microcysteis] |
| 05140 | 4 | glycosyltransferase family 2 protein [Azoarcus nasutitermitis] |
| 03551 | 4 | antibiotic biosynthesis monooxygenase [Niveibacterium microcysteis] |
| 03552 | 4 | DUF3302 domain-containing protein [Niveibacterium microcysteis] |
| 03461 | 4 | cobalamin biosynthesis protein CobG [Azoarcus communis] |
| 00629 | 4 | homogentisate 1,2-dioxygenase [Nitrogeniibacter mangrovi] |
| 03359 | 4 | DUF2157 domain-containing protein [Thauera aromatica] |
| 01594 | 4 | DUF4926 domain-containing protein [Aromatoleum aromaticum] |
| 03360 | 4 | hypothetical protein [Thauera aromatica] |
| 03365 | 4 | hypothetical protein [Candidatus Dactylopiibacterium carminicum] |
| 01620 | 4 | beta-galactosidase [Niveibacterium microcysteis] |
| 03367 | 4 | hypothetical protein [Azoarcus sp. DD4] |
| 05301 | 4 | hypothetical protein [Aromatoleum petrolei] |
| 03382 | 4 | type I-E CRISPR-associated protein Cse1/CasA [Aromatoleum toluvorans] |
| 00600 | 4 | HI1506-related protein [Azoarcus rhizosphaerae] |
| 01606 | 4 | helix-turn-helix domain-containing protein [Nitrogeniibacter aestuarii] |
| 00601 | 4 | ATP-binding protein [Aromatoleum toluclasticum] |
| 05237 | 4 | phage tail sheath family protein [Azoarcus communis] |
| 03400 | 4 | TonB family protein [Azoarcus rhizosphaerae] |
| 03416 | 4 | hypothetical protein [Zoogloea ramigera] |
| 05254 | 4 | hypothetical protein [Aromatoleum tolulyticum] |
| 03429 | 4 | C45 family autoproteolytic acyltransferase/hydolase [Denitromonas halophilus] |
| 05248 | 4 | DUF1345 domain-containing protein [Niveibacterium sp. COAC-50] |
| 00610 | 4 | hypothetical protein [Niveibacterium microcysteis] |
| 00612 | 4 | hypothetical protein [Zoogloea dura] |
| 05238 | 4 | MULTISPECIES: Fic family protein [Betaproteobacteria] |
| 05531 | 4 | phosphonate C-P lyase system protein PhnG [Azoarcus nasutitermitis] |
| 01766 | 4 | hypothetical protein [Aromatoleum evansii] |
| 05125 | 4 | hypothetical protein [Azoarcus indigens] |
| 03198 | 4 | MULTISPECIES: ribbon-helix-helix protein, CopG family [Thauera] |
| 05778 | 4 | VOC family protein [Aromatoleum toluolicum] |
| 05774 | 4 | acyltransferase [Azoarcus halotolerans] |
| 00394 | 4 | cytoplasmic protein [Denitromonas halophilus] |
| 02986 | 4 | RNA-directed DNA polymerase [Aromatoleum tolulyticum] |
| 02990 | 4 | Ig-like domain-containing protein, partial [Azoarcus halotolerans] |
| 01889 | 4 | glycosyltransferase [Uliginosibacterium aquaticum] |
| 03001 | 4 | MULTISPECIES: hypothetical protein [Thauera] |
| 05758 | 4 | biotin carboxylase [Denitromonas halophilus] |
| 05757 | 4 | hypothetical protein [Denitromonas halophilus] |
| 05753 | 4 | ferrous iron transport protein A [Denitromonas halophilus] |
| 00417 | 4 | tetratricopeptide repeat protein [Azoarcus indigens] |
| 03018 | 4 | phage head closure protein [Aromatoleum aromaticum] |
| 05725 | 4 | helix-turn-helix domain-containing protein [Thauera phenolivorans] |
| 05723 | 4 | cobyrinic acid a,c-diamide synthase [Azoarcus taiwanensis] |
| 03036 | 4 | MULTISPECIES: MFS transporter [Proteobacteria] |
| 03037 | 4 | DUF2218 domain-containing protein [Azoarcus rhizosphaerae] |
| 00423 | 4 | hypothetical protein [Azoarcus communis] |
| 03040 | 4 | MULTISPECIES: RNA methyltransferase [Niveibacterium] |
| 05712 | 4 | MULTISPECIES: hypothetical protein [Proteobacteria] |
| 01897 | 4 | TIGR00366 family protein [Azoarcus halotolerans] |
| 05784 | 4 | hypothetical protein [Azoarcus sp. KH32C] |
| 01906 | 4 | MULTISPECIES: ion channel [Niveibacterium] |
| 05805 | 4 | DUF3800 domain-containing protein [Azoarcus sp. Aa7] |
| 05827 | 4 | hypothetical protein [Azoarcus indigens] |
| 02921 | 4 | MULTISPECIES: hypothetical protein [Azospira] |
| 02924 | 4 | dihydrodipicolinate reductase [Azoarcus sp. L1K30] |
| 02926 | 4 | FecR domain-containing protein [Propionivibrio dicarboxylicus] |
| 02930 | 4 | CHASE2 domain-containing protein [Azoarcus indigens] |
| 02936 | 4 | ethanolamine utilization protein EutP [Azoarcus nasutitermitis] |
| 02947 | 4 | MFS transporter [Azoarcus rhizosphaerae] |

| | | |
|---|---|---|
| 01921 | 4 | S9 family peptidase [Candidatus Dactylopiibacterium carminicum] |
| 00364 | 4 | phage major capsid protein [Azoarcus indigens] |
| 02969 | 4 | DUF4393 domain-containing protein [Denitromonas halophilus] |
| 00366 | 4 | ATP-grasp domain-containing protein [Azoarcus indigens] |
| 00367 | 4 | ABC transporter permease [Azoarcus indigens] |
| 02955 | 4 | DUF4336 domain-containing protein [Azoarcus olearius] |
| 01918 | 4 | hypothetical protein [Candidatus Dactylopiibacterium carminicum] |
| 01916 | 4 | hypothetical protein [Azoarcus indigens] |
| 01911 | 4 | hypothetical protein [Niveibacterium microcysteis] |
| 01907 | 4 | MULTISPECIES: hypothetical protein [Niveibacterium] |
| 02965 | 4 | hypothetical protein [Denitromonas halophilus] |
| 03050 | 4 | gamma-glutamylcyclotransferase [Azoarcus sp. TTM-91] |
| 01872 | 4 | DDE-type integrase/transposase/recombinase [Aromatoleum pumilum] |
| 01871 | 4 | hypothetical protein [Aromatoleum evansii] |
| 05577 | 4 | HAD family hydrolase [Candidatus Dactylopiibacterium carminicum] |
| 00463 | 4 | hypothetical protein [Aromatoleum aromaticum] |
| 00465 | 4 | GspH/FimT family protein [Aromatoleum aromaticum] |
| 01810 | 4 | hypothetical protein [Azospira oryzae] |
| 03144 | 4 | N(5)-hydroxyornithine transformylase PvdF [Azoarcus indigens] |
| 03145 | 4 | hypothetical protein [Thauera humireducens] |
| 01796 | 4 | GNAT family N-acetyltransferase [Thauera aminoaromatica] |
| 03162 | 4 | PilZ domain-containing protein [Aromatoleum buckelii] |
| 05586 | 4 | BrnT family toxin [Azoarcus sp. Aa7] |
| 00488 | 4 | Fic family protein [Uliginosibacterium gangwonense] |
| 01816 | 4 | DUF4157 domain-containing protein [Thauera humireducens] |
| 03173 | 4 | MULTISPECIES: hypothetical protein [Aromatoleum] |
| 00490 | 4 | transposase, partial [Uliginosibacterium gangwonense] |
| 05570 | 4 | hypothetical protein [Niveibacterium microcysteis] |
| 00491 | 4 | hypothetical protein [Uliginosibacterium gangwonense] |
| 03175 | 4 | hypothetical protein [Thauera aminoaromatica] |
| 01777 | 4 | hypothetical protein [Aromatoleum toluclasticum] |
| 03189 | 4 | ATP-dependent DNA ligase [Azoarcus indigens] |
| 03190 | 4 | HPr-rel-A system PqqD family peptide chaperone [Azoarcus communis] |
| 03141 | 4 | transposase [Denitromonas halophilus] |
| 00455 | 4 | hypothetical protein [Nitrogeniibacter aestuarii] |
| 03059 | 4 | hypothetical protein [Nitrogeniibacter aestuarii] |
| 05661 | 4 | hypothetical protein [Aromatoleum aromaticum] |
| 03060 | 4 | hypothetical protein [Azoarcus sp. Aa7] |
| 05686 | 4 | hypothetical protein [Niveibacterium microcysteis] |
| 05679 | 4 | formylmethanofuran dehydrogenase [Aromatoleum toluvorans] |
| 05672 | 4 | PHB depolymerase family esterase [Thauera butanivorans] |
| 00434 | 4 | metallophosphoesterase [Aromatoleum diolicum] |
| 05669 | 4 | hypothetical protein [Thauera aminoaromatica] |
| 03068 | 4 | baseplate J/gp47 family protein [Azoarcus indigens] |
| 05666 | 4 | hypothetical protein [Azoarcus halotolerans] |
| 01864 | 4 | FkbM family methyltransferase [Aromatoleum diolicum] |
| 05609 | 4 | MULTISPECIES: hypothetical protein [Niveibacterium] |
| 01860 | 4 | DUF6513 domain-containing protein [Aromatoleum toluvorans] |
| 01853 | 4 | helix-turn-helix transcriptional regulator [Azoarcus indigens] |
| 01850 | 4 | MULTISPECIES: hypothetical protein, partial [Proteobacteria] |
| 03098 | 4 | RHS repeat protein [Niveibacterium sp. COAC-50] |
| 00448 | 4 | H-NS histone family protein [Thauera phenolivorans] |
| 01839 | 4 | hypothetical protein [Aromatoleum buckelii] |
| 05617 | 4 | hypothetical protein [Aromatoleum petrolei] |
| 05613 | 4 | hypothetical protein [Aromatoleum petrolei] |
| 03555 | 4 | hypothetical protein [Nitrogeniibacter aestuarii] |
| 01488 | 4 | MULTISPECIES: hypothetical protein [Uliginosibacterium] |
| 02904 | 4 | Arc family DNA-binding protein [Zoogloea dura] |
| 01213 | 4 | carboxymuconolactone decarboxylase family protein [Azoarcus olearius] |
| 00896 | 4 | DUF3106 domain-containing protein [Azospira oryzae] |
| 01121 | 4 | hypothetical protein [Denitromonas halophilus] |
| 01119 | 4 | hypothetical protein [Denitromonas halophilus] |
| 01118 | 4 | helix-turn-helix domain-containing protein [Denitromonas halophilus] |
| 04191 | 4 | hypothetical protein [Azoarcus olearius] |
| 04659 | 4 | hypothetical protein [Aromatoleum aromaticum] |
| 01111 | 4 | hypothetical protein [Zoogloea oleivorans] |
| 04654 | 4 | hypothetical protein [Aromatoleum toluclasticum] |
| 04651 | 4 | hypothetical protein [Niveibacterium microcysteis] |
| 04645 | 4 | phospholipase D-like domain-containing protein [Uliginosibacterium gangwonense] |
| 04242 | 4 | MULTISPECIES: hypothetical protein [Betaproteobacteria] |
| 00920 | 4 | hypothetical protein [Azoarcus communis] |
| 04640 | 4 | DUF447 family protein [Aromatoleum toluvorans] |
| 00922 | 4 | hypothetical protein [Azoarcus sp. DD4] |
| 01099 | 4 | hypothetical protein [Azoarcus indigens] |
| 04628 | 4 | hypothetical protein [Aromatoleum petrolei] |
| 00929 | 4 | ATP-binding protein [Nitrogeniibacter aestuarii] |
| 04263 | 4 | ABC transporter permease [Aromatoleum evansii] |
| 00938 | 4 | 3D domain-containing protein [Denitromonas halophilus] |
| 04187 | 4 | DEAD/DEAH box helicase family protein [Aromatoleum aromaticum] |
| 04671 | 4 | MarR family transcriptional regulator [Azoarcus nasutitermitis] |
| 04144 | 4 | hypothetical protein [Azoarcus halotolerans] |
| 04083 | 4 | CHAT domain-containing protein [Niveibacterium umoris] |
| 00846 | 4 | PEP-CTERM sorting domain-containing protein [Thauera phenylacetica] |
| 04056 | 4 | type I-E CRISPR-associated protein Cse2/CasB [Aromatoleum toluvorans] |
| 04729 | 4 | helix-turn-helix domain-containing protein [Azoarcus indigens] |
| 01184 | 4 | hypothetical protein [Aromatoleum toluclasticum] |
| 00859 | 4 | BREX-1 system phosphatase PglZ type B [Azoarcus olearius] |
| 04717 | 4 | MULTISPECIES: hypothetical protein [Azospira] |
| 04079 | 4 | glycoside hydrolase [Niveibacterium sp. COAC-50] |
| 04713 | 4 | MULTISPECIES: SRPBCC domain-containing protein [Niveibacterium] |
| 01170 | 4 | MULTISPECIES: protein-disulfide reductase DsbD N-terminal domain-containing protein [Betaproteobacteria] |
| 04143 | 4 | MULTISPECIES: complement resistance protein TraT [Thauera] |
| 04089 | 4 | MULTISPECIES: type II toxin-antitoxin system HicB family antitoxin [Aromatoleum] |
| 00868 | 4 | restriction endonuclease subunit S [Azoarcus sp. L1K30] |

| | | |
|---|---|---|
| 04708 | 4 | VTT domain-containing protein [Azoarcus communis] |
| 04102 | 4 | hypothetical protein [Denitromonas halophilus] |
| 01169 | 4 | MULTISPECIES: hypothetical protein [Rhodocyclales] |
| 04114 | 4 | hypothetical protein [Azoarcus rhizosphaerae] |
| 00880 | 4 | hypothetical protein [Azoarcus communis] |
| 00881 | 4 | hypothetical protein [Azoarcus olearius] |
| 02299 | 4 | ethanolamine utilization protein EutJ [Azoarcus nasutitermitis] |
| 04605 | 4 | hypothetical protein, partial [Aromatoleum toluclasticum] |
| 01090 | 4 | FTR1 family iron permease [Azoarcus nasutitermitis] |
| 00986 | 4 | hypothetical protein [Azoarcus communis] |
| 04357 | 4 | hypothetical protein [Thauera propionica] |
| 04369 | 4 | hypothetical protein [Aromatoleum pumilum] |
| 04512 | 4 | AMP-binding protein, partial [Aromatoleum evansii] |
| 04503 | 4 | Ig-like domain-containing protein, partial [Aromatoleum diolicum] |
| 00969 | 4 | hypothetical protein [Azoarcus nasutitermitis] |
| 04388 | 4 | phage baseplate assembly protein V [Azoarcus indigens] |
| 01019 | 4 | HP1 family phage holin [Oryzomicrobium terrae] |
| 00983 | 4 | MULTISPECIES: methylenetetrahydromethanopterin dehydrogenase [Thauera] |
| 04472 | 4 | hypothetical protein [Azoarcus communis] |
| 00961 | 4 | MULTISPECIES: macro domain-containing protein [Thauera] |
| 04398 | 4 | alpha-D-ribose 1-methylphosphonate 5-phosphate C-P-lyase PhnJ [Azoarcus nasutitermitis] |
| 04399 | 4 | hypothetical protein [Thauera butanivorans] |
| 04468 | 4 | hypothetical protein [Thauera butanivorans] |
| 04401 | 4 | hypothetical protein [Azoarcus nasutitermitis] |
| 04466 | 4 | hypothetical protein [Aromatoleum buckelii] |
| 04465 | 4 | hypothetical protein [Aromatoleum aromaticum] |
| 04403 | 4 | GGDEF domain-containing protein [Aromatoleum buckelii] |
| 04458 | 4 | PEP-CTERM sorting domain-containing protein [Denitromonas halophilus] |
| 00963 | 4 | DEAD/DEAH box helicase [Thauera aminoaromatica] |
| 04526 | 4 | PAS domain-containing protein [Candidatus Dactylopiibacterium carminicum] |
| 04597 | 4 | alpha/beta fold hydrolase [Nitrogeniibacter aestuarii] |
| 01072 | 4 | MULTISPECIES: PilZ domain-containing protein [Azospira] |
| 04291 | 4 | amidohydrolase family protein [Aromatoleum evansii] |
| 00949 | 4 | MULTISPECIES: helix-turn-helix domain-containing protein [Rhodocyclales] |
| 01081 | 4 | hypothetical protein [Azoarcus indigens] |
| 04590 | 4 | VOC family protein [Azoarcus olearius] |
| 04296 | 4 | ligase-associated DNA damage response exonuclease [Azoarcus indigens] |
| 04582 | 4 | peptidyl-prolyl cis-trans isomerase [Aromatoleum toluclasticum] |
| 00952 | 4 | hypothetical protein [Aromatoleum aromaticum] |
| 04308 | 4 | DUF4433 domain-containing protein [Aromatoleum evansii] |
| 04316 | 4 | cupin domain-containing protein, partial [Uliginosibacterium gangwonense] |
| 00959 | 4 | invasion associated locus B family protein [Nitrogeniibacter aestuarii] |
| 04318 | 4 | MipA/OmpV family protein [Uliginosibacterium gangwonense] |
| 01059 | 4 | DUF642 domain-containing protein [Niveibacterium microcysteis] |
| 04326 | 4 | energy transducer TonB [Azoarcus indigens] |
| 04330 | 4 | MFS transporter [Thauera terpenica] |
| 01055 | 4 | DUF3427 domain-containing protein [Aromatoleum diolicum] |
| 01047 | 4 | HicA protein [Zoogloea ramigera] |
| 04344 | 4 | hypothetical protein [Azoarcus communis] |
| 04552 | 4 | VCBS domain-containing protein, partial [Niveibacterium sp. COAC-50] |
| 04744 | 4 | phage tail tape measure protein [Aromatoleum pumilum] |
| 04759 | 4 | hypothetical protein [Azoarcus communis] |
| 01487 | 4 | SPFH domain-containing protein [Niveibacterium microcysteis] |
| 04031 | 4 | hypothetical protein [Niveibacterium microcysteis] |
| 01433 | 4 | MULTISPECIES: PadR family transcriptional regulator [Thauera] |
| 03634 | 4 | hypothetical protein [Zoogloea oleivorans] |
| 05010 | 4 | DNA-binding protein [Aromatoleum tolulyticum] |
| 05009 | 4 | hypothetical protein [Aromatoleum evansii] |
| 05008 | 4 | GNAT family N-acetyltransferase [Nitrogeniibacter aestuarii] |
| 05002 | 4 | CoA-binding protein [Propionivibrio dicarboxylicus] |
| 01421 | 4 | ThiF family adenylyltransferase [Aromatoleum diolicum] |
| 04994 | 4 | MULTISPECIES: TniQ family protein [Thauera] |
| 00722 | 4 | MULTISPECIES: hypothetical protein [Aromatoleum] |
| 03669 | 4 | DUF4236 domain-containing protein [Aromatoleum toluclasticum] |
| 03673 | 4 | hypothetical protein [Thauera aromatica] |
| 04982 | 4 | DUF2326 domain-containing protein [Aromatoleum evansii] |
| 04980 | 4 | hypothetical protein [Nitrogeniibacter aestuarii] |
| 03685 | 4 | hypothetical protein [Aromatoleum bremense] |
| 00751 | 4 | hypothetical protein [Uliginosibacterium gangwonense] |
| 04970 | 4 | metalloregulator ArsR/SmtB family transcription factor [Azoarcus nasutitermitis] |
| 03695 | 4 | MULTISPECIES: recombinase family protein [Bacteria] |
| 04964 | 4 | hypothetical protein [Aromatoleum bremense] |
| 01339 | 4 | hydrolase or metal-binding protein [Niveibacterium microcysteis] |
| 05030 | 4 | hypothetical protein [Aromatoleum buckelii] |
| 00707 | 4 | hypothetical protein [Niveibacterium microcysteis] |
| 05038 | 4 | YjbF family lipoprotein [Azoarcus sp. KH32C] |
| 05090 | 4 | AAA family ATPase [Aromatoleum evansii] |
| 01484 | 4 | DUF3313 family protein [Uliginosibacterium gangwonense] |
| 03563 | 4 | MULTISPECIES: hypothetical protein [Niveibacterium] |
| 05111 | 4 | hypothetical protein [Azoarcus communis] |
| 01483 | 4 | DUF2950 domain-containing protein [Niveibacterium sp. COAC-50] |
| 01480 | 4 | glycoside hydrolase [Niveibacterium sp. COAC-50] |
| 00665 | 4 | TssQ family T6SS-associated lipoprotein [Aromatoleum diolicum] |
| 05098 | 4 | MULTISPECIES: phosphate-starvation-inducible PsiE family protein [Niveibacterium] |
| 05097 | 4 | ATP-binding protein [Propionivibrio dicarboxylicus] |
| 00672 | 4 | hypothetical protein [Niveibacterium microcysteis] |
| 00702 | 4 | MULTISPECIES: hypothetical protein [Niveibacterium] |
| 00673 | 4 | hypothetical protein [Azoarcus indigens] |
| 05077 | 4 | major capsid protein [Aromatoleum aromaticum] |
| 00679 | 4 | MAPEG family protein [Azoarcus indigens] |
| 05071 | 4 | MULTISPECIES: hypothetical protein [Zoogloea] |
| 01463 | 4 | MarR family transcriptional regulator [Aromatoleum evansii] |
| 03594 | 4 | class I SAM-dependent methyltransferase [Aromatoleum toluclasticum] |
| 05052 | 4 | Rieske (2Fe-2S) protein [Azoarcus communis] |

| | | |
|---|---|---|
| 01461 | 4 | hypothetical protein [Zoogloea ramigera] |
| 03709 | 4 | hypothetical protein, partial [Azoarcus communis] |
| 03723 | 4 | DUF2848 domain-containing protein [Azoarcus indigens] |
| 01325 | 4 | LuxR C-terminal-related transcriptional regulator [Aromatoleum toluclasticum] |
| 03924 | 4 | glycosyltransferase [Denitromonas halophilus] |
| 04849 | 4 | GIY-YIG nuclease family protein [Aromatoleum evansii] |
| 03899 | 4 | hypothetical protein [Thauera humireducens] |
| 04847 | 4 | EAL domain-containing protein [Aromatoleum evansii] |
| 00805 | 4 | Tm-1-like ATP-binding domain-containing protein [Aromatoleum aromaticum] |
| 04835 | 4 | hypothetical protein [Nitrogeniibacter aestuarii] |
| 04830 | 4 | UDP-N-acetylglucosamine--LPS N-acetylglucosamine transferase [Aromatoleum diolicum] |
| 01259 | 4 | filamentous hemagglutinin N-terminal domain-containing protein [Azoarcus indigens] |
| 03921 | 4 | IS3 family transposase, partial [Denitromonas halophilus] |
| 03928 | 4 | FAD-dependent oxidoreductase, partial [Denitromonas halophilus] |
| 03887 | 4 | class I SAM-dependent methyltransferase [Aromatoleum bremense] |
| 04814 | 4 | hypothetical protein [Denitromonas halophilus] |
| 03931 | 4 | phage GP46 family protein [Azoarcus indigens] |
| 00822 | 4 | hypothetical protein [Azoarcus halotolerans] |
| 04806 | 4 | hypothetical protein [Azoarcus halotolerans] |
| 03959 | 4 | YfdX family protein [Nitrogeniibacter aestuarii] |
| 00830 | 4 | transposase, partial [Uliginosibacterium gangwonense] |
| 03989 | 4 | alpha-ketoacid dehydrogenase subunit alpha/beta [Azoarcus nasutitermitis] |
| 04778 | 4 | hypothetical protein [Azoarcus indigens] |
| 01267 | 4 | DUF3138 family protein [Niveibacterium umoris] |
| 03883 | 4 | alpha/beta-hydrolase family protein [Azoarcus nasutitermitis] |
| 03761 | 4 | AAA family ATPase [Azoarcus halotolerans] |
| 03827 | 4 | DEAD/DEAH box helicase family protein [Aromatoleum aromaticum] |
| 04934 | 4 | hypothetical protein [Rhodocyclus purpureus] |
| 01305 | 4 | hypothetical protein [Azoarcus communis] |
| 00765 | 4 | hypothetical protein [Aromatoleum toluclasticum] |
| 01296 | 4 | tetratricopeptide repeat protein [Niveibacterium umoris] |
| 04923 | 4 | recombinase family protein [Aromatoleum toluolicum] |
| 03796 | 4 | hypothetical protein [Niveibacterium microcysteis] |
| 04906 | 4 | hypothetical protein [Niveibacterium microcysteis] |
| 03808 | 4 | MULTISPECIES: hypothetical protein [Betaproteobacteria] |
| 01279 | 4 | hypothetical protein [Azoarcus communis] |
| 03882 | 4 | ethanolamine utilization microcompartment protein EutL [Azoarcus nasutitermitis] |
| 01272 | 4 | alginate export family protein [Nitrogeniibacter aestuarii] |
| 04888 | 4 | alpha/beta fold hydrolase [Zoogloea dura] |
| 04879 | 4 | DVUA0089 family protein [Azoarcus indigens] |
| 03848 | 4 | cobalamin biosynthesis protein [Azoarcus communis] |
| 03854 | 4 | type II toxin-antitoxin system PemK/MazF family toxin [Azoarcus communis] |
| 04869 | 4 | formylmethanofuran dehydrogenase subunit C [Aromatoleum toluvorans] |
| 00798 | 4 | TIGR00366 family protein [Propionivibrio dicarboxylicus] |
| 03873 | 4 | DUF817 domain-containing protein [Nitrogeniibacter aestuarii] |
| 02910 | 4 | 2-succinyl-5-enolpyruvyl-6-hydroxy-3-cyclohexene-1-carboxylic-acid synthase [Azoarcus communis] |
| 04430 | 4 | hypothetical protein [Denitromonas halophilus] |
| 02684 | 4 | hypothetical protein [Zoogloea dura] |
| 05900 | 4 | type VI secretion system Vgr family protein, partial [Denitromonas halophilus] |
| 02782 | 4 | peptidoglycan recognition protein family protein [Aromatoleum toluclasticum] |
| 02253 | 4 | DUF4112 domain-containing protein [Nitrogeniibacter aestuarii] |
| 02249 | 4 | FixH family protein [Niveibacterium umoris] |
| 05898 | 4 | MULTISPECIES: DUF350 domain-containing protein [Niveibacterium] |
| 02043 | 4 | tetratricopeptide repeat protein [Thauera humireducens] |
| 02587 | 4 | oligosaccharide flippase family protein [Thauera aminoaromatica] |
| 02345 | 4 | GIY-YIG nuclease family protein [Azoarcus halotolerans] |
| 06196 | 4 | hypothetical protein [Nitrogeniibacter aestuarii] |
| 02768 | 4 | MULTISPECIES: hypothetical protein [Niveibacterium] |
| 02487 | 4 | hypothetical protein [Aromatoleum toluolicum] |
| 02351 | 4 | hypothetical protein [Azoarcus indigens] |
| 00048 | 4 | autotransporter outer membrane beta-barrel domain-containing protein [Azoarcus indigens] |
| 02358 | 4 | LuxR C-terminal-related transcriptional regulator [Denitromonas sp. IR12] |
| 00279 | 4 | hypothetical protein [Uliginosibacterium gangwonense] |
| 01986 | 4 | hypothetical protein [Azospira oryzae] |
| 02760 | 4 | OmpA family protein [Niveibacterium microcysteis] |
| 02365 | 4 | hypothetical protein [Aromatoleum evansii] |
| 06111 | 4 | hypothetical protein [Nitrogeniibacter aestuarii] |
| 02785 | 4 | hypothetical protein [Azoarcus halotolerans] |
| 06202 | 4 | hypothetical protein [Nitrogeniibacter aestuarii] |
| 02570 | 4 | MULTISPECIES: hypothetical protein [Rhodocyclales] |
| 01970 | 4 | MFS transporter [Azoarcus communis] |
| 02097 | 4 | phage tail sheath family protein [Azoarcus nasutitermitis] |
| 02576 | 4 | P-type DNA transfer protein VirB5 [Thauera terpenica] |
| 02329 | 4 | hypothetical protein [Zoogloea oleivorans] |
| 06214 | 4 | cytochrome oxidase small assembly protein [Nitrogeniibacter aestuarii] |
| 02819 | 4 | MULTISPECIES: hypothetical protein [Bacteria] |
| 06028 | 4 | DUF5368 domain-containing protein [Azoarcus rhizosphaerae] |
| 02093 | 4 | hypothetical protein [Azoarcus indigens] |
| 02811 | 4 | hypothetical protein [Denitromonas halophilus] |
| 00129 | 4 | tetratricopeptide repeat protein [Azoarcus indigens] |
| 05897 | 4 | hypothetical protein [Niveibacterium microcysteis] |
| 06205 | 4 | AAA family ATPase [Thauera linaloolentis] |
| 01973 | 4 | hypothetical protein [Nitrogeniibacter aestuarii] |
| 06203 | 4 | hypothetical protein [Nitrogeniibacter aestuarii] |
| 00295 | 4 | hypothetical protein [Niveibacterium microcysteis] |
| 00292 | 4 | ArsR family transcriptional regulator [Rhodocyclus purpureus] |
| 02583 | 4 | hypothetical protein [Thauera humireducens] |
| 02793 | 4 | sensor domain-containing diguanylate cyclase [Nitrogeniibacter aestuarii] |
| 00127 | 4 | phosphate ABC transporter permease PstA [Aromatoleum pumilum] |
| 02078 | 4 | hypothetical protein [Azoarcus communis] |
| 02374 | 4 | UvrD-helicase domain-containing protein [Denitromonas halophilus] |
| 00204 | 4 | class I SAM-dependent methyltransferase [Niveibacterium umoris] |
| 02441 | 4 | cysteine-rich CWC family protein [Aromatoleum pumilum] |
| 02023 | 4 | gamma-glutamyl phosphate reductase [Niveibacterium sp. COAC-50] |

| | | |
|---|---|---|
| 02703 | 4 | EAL domain-containing protein [Aromatoleum bremense] |
| 06130 | 4 | MULTISPECIES: oxidoreductase [Azospira] |
| 02202 | 4 | MbtH family NRPS accessory protein [Azoarcus indigens] |
| 02648 | 4 | class I SAM-dependent methyltransferase [Aromatoleum bremense] |
| 00078 | 4 | hypothetical protein [Denitromonas halophilus] |
| 00079 | 4 | type VI secretion system tip protein VgrG, partial [Zoogloea ramigera] |
| 06136 | 4 | hypothetical protein [Aromatoleum diolicum] |
| 02679 | 4 | transposase domain-containing protein, partial [Zoogloea oleivorans] |
| 00107 | 4 | phage integrase N-terminal SAM-like domain-containing protein [Aromatoleum bremense] |
| 00095 | 4 | DNA repair exonuclease [Thauera butanivorans] |
| 00091 | 4 | site-specific integrase [Azoarcus sp. KH32C] |
| 05989 | 4 | MULTISPECIES: hypothetical protein [Thauera] |
| 02435 | 4 | hypothetical protein [Aromatoleum evansii] |
| 02653 | 4 | ethanolamine utilization protein EutH [Azoarcus nasutitermitis] |
| 05985 | 4 | phage/plasmid replication protein, II/X family [Thauera butanivorans] |
| 05981 | 4 | MULTISPECIES: hypothetical protein [Thauera] |
| 05964 | 4 | MarR family EPS-associated transcriptional regulator [Rhodocyclus purpureus] |
| 02191 | 4 | dihydromethanopterin reductase [Aromatoleum toluvorans] |
| 00251 | 4 | antitoxin [Azoarcus nasutitermitis] |
| 02713 | 4 | AAA family ATPase [Denitromonas halophilus] |
| 02715 | 4 | class I SAM-dependent methyltransferase [Azoarcus halotolerans] |
| 06114 | 4 | hypothetical protein [Aromatoleum evansii] |
| 02464 | 4 | pilin [Niveibacterium umoris] |
| 06116 | 4 | ATP-binding protein [Nitrogeniibacter aestuarii] |
| 00054 | 4 | glycosyltransferase [Azoarcus indigens] |
| 00267 | 4 | hypothetical protein [Azoarcus halotolerans] |
| 02619 | 4 | DUF4402 domain-containing protein [Azoarcus communis] |
| 02064 | 4 | HEPN domain-containing protein [Azoarcus sp. Aa7] |
| 02386 | 4 | hypothetical protein [Aromatoleum toluolicum] |
| 00209 | 4 | Na+/H+ antiporter NhaC [Aromatoleum evansii] |
| 02391 | 4 | MULTISPECIES: conjugal transfer protein TraL [Aromatoleum] |
| 01992 | 4 | hypothetical protein [Aromatoleum aromaticum] |
| 02633 | 4 | cobalt ECF transporter T component CbiQ [Azospira oryzae] |
| 02730 | 4 | MULTISPECIES: hypothetical protein [Aromatoleum] |
| 02639 | 4 | ferritin-like domain-containing protein [Azoarcus indigens] |
| 00257 | 4 | AraC family transcriptional regulator [Aromatoleum aromaticum] |
| 05997 | 4 | hypothetical protein [Azoarcus communis] |
| 02207 | 4 | hypothetical protein [Zoogloea dura] |
| 00307 | 4 | DUF4150 domain-containing protein [Thauera butanivorans] |
| 02665 | 4 | DUF4194 domain-containing protein [Azoarcus indigens] |
| 06050 | 4 | DUF1045 domain-containing protein [Azoarcus nasutitermitis] |
| 00341 | 4 | hypothetical protein [Azoarcus communis] |
| 02896 | 4 | hypothetical protein [Azoarcus nasutitermitis] |
| 00139 | 4 | hypothetical protein [Azoarcus indigens] |
| 02119 | 4 | hypothetical protein [Thauera chlorobenzoica] |
| 02169 | 4 | glycosyltransferase [Uliginosibacterium aquaticum] |
| 06233 | 4 | VOC family protein [Aromatoleum petrolei] |
| 02893 | 4 | amidoligase family protein [Aromatoleum pumilum] |
| 05864 | 4 | hypothetical protein [Candidatus Dactylopiibacterium carminicum] |
| 00020 | 4 | glycosyltransferase [Rhodocyclus purpureus] |
| 00004 | 4 | MULTISPECIES: hypothetical protein [Proteobacteria] |
| 02860 | 4 | MarR family transcriptional regulator [Thauera butanivorans] |
| 01938 | 4 | 2OG-Fe dioxygenase family protein [Azoarcus rhizosphaerae] |
| 02294 | 4 | acyl-CoA thioesterase [Azospira oryzae] |
| 02892 | 4 | hypothetical protein [Aromatoleum aromaticum] |
| 02114 | 4 | DEAD/DEAH box helicase family protein [Aromatoleum bremense] |
| 05860 | 4 | MULTISPECIES: hypothetical protein [Azospira] |
| 02559 | 4 | hypothetical protein [Nitrogeniibacter aestuarii] |
| 06243 | 4 | hypothetical protein [Denitromonas halophilus] |
| 02500 | 4 | replication initiation protein [Aromatoleum evansii] |
| 00012 | 4 | aconitase X catalytic domain-containing protein [Aromatoleum diolicum] |
| 02508 | 4 | MULTISPECIES: hypothetical protein [Azospira] |
| 00009 | 4 | triphosphoribosyl-dephospho-CoA synthase [Aromatoleum toluvorans] |
| 02301 | 4 | hypothetical protein [Azoarcus nasutitermitis] |
| 02168 | 4 | anaerobic nitric oxide reductase flavorubredoxin [Aromatoleum aromaticum] |
| 06069 | 4 | hypothetical protein [Denitromonas halophilus] |
| 06055 | 4 | ethanolamine utilization acetate kinase EutQ [Azoarcus nasutitermitis] |
| 06256 | 4 | response regulator [Aromatoleum toluclasticum] |
| 02513 | 4 | BON domain-containing protein [Aromatoleum aromaticum] |
| 05852 | 4 | hypothetical protein [Niveibacterium microcysteis] |
| 06071 | 4 | hypothetical protein [Denitromonas halophilus] |
| 06058 | 4 | carbohydrate binding family 9 domain-containing protein [Thauera phenylacetica] |
| 02568 | 4 | hypothetical protein [Aromatoleum toluclasticum] |
| 00311 | 4 | glutamate--cysteine ligase [Azoarcus communis] |
| 05846 | 4 | DUF6268 family outer membrane beta-barrel protein [Niveibacterium umoris] |
| 02125 | 4 | hypothetical protein [Denitromonas halophilus] |
| 02851 | 4 | hypothetical protein [Nitrogeniibacter aestuarii] |
| 02900 | 4 | DUF3987 domain-containing protein [Aromatoleum pumilum] |
| 02563 | 4 | MULTISPECIES: porin [Niveibacterium] |
| 06044 | 4 | hypothetical protein [Azoarcus nasutitermitis] |
| 05838 | 4 | ATP-binding protein [Aromatoleum toluclasticum] |
| 02303 | 4 | LacI family transcriptional regulator [Candidatus Dactylopiibacterium carminicum] |
| 07882 | 3 | hypothetical protein [Denitromonas halophilus] |
| 07880 | 3 | HAMP domain-containing histidine kinase [Denitromonas halophilus] |
| 08943 | 3 | tetratricopeptide repeat protein [Azoarcus olearius] |
| 01261 | 3 | phage tail assembly protein [Azoarcus indigens] |
| 08895 | 3 | HK97 family phage prohead protease [Azoarcus indigens] |
| 08047 | 3 | DUF1800 domain-containing protein [Nitrogeniibacter aestuarii] |
| 08913 | 3 | hypothetical protein [Candidatus Dactylopiibacterium carminicum] |
| 07887 | 3 | hypothetical protein [Azoarcus rhizosphaerae] |
| 02449 | 3 | hypothetical protein [Niveibacterium microcysteis] |
| 08900 | 3 | FUSC family protein [Azoarcus indigens] |
| 08044 | 3 | phytanoyl-CoA dioxygenase family protein [Nitrogeniibacter aestuarii] |
| 09162 | 3 | hypothetical protein [Zoogloea dura] |

| | | |
|---|---|---|
| **03872** | 3 | hypothetical protein [Uliginosibacterium aquaticum] |
| **07899** | 3 | hypothetical protein [Azoarcus communis] |
| **02183** | 3 | AraC family transcriptional regulator, partial [Azoarcus indigens] |
| **08045** | 3 | lysozyme inhibitor LprI family protein [Nitrogeniibacter aestuarii] |
| **07896** | 3 | aldolase [Zoogloea dura] |
| **07888** | 3 | hypothetical protein [Zoogloea dura] |
| **03874** | 3 | phage head-tail connector protein [Azoarcus indigens] |
| **03875** | 3 | cytochrome P450 [Azoarcus indigens] |
| **07892** | 3 | hypothetical protein [Uliginosibacterium gangwonense] |
| **09158** | 3 | hypothetical protein [Zoogloea dura] |
| **02188** | 3 | DUF637 domain-containing protein [Azoarcus rhizosphaerae] |
| **01269** | 3 | hypothetical protein [Niveibacterium microcysteis] |
| **08046** | 3 | hypothetical protein [Nitrogeniibacter aestuarii] |
| **09159** | 3 | hypothetical protein [Niveibacterium microcysteis] |
| **03901** | 3 | hypothetical protein [Aromatoleum aromaticum] |
| **02443** | 3 | MalM family protein [Niveibacterium microcysteis] |
| **01429** | 3 | DNA-binding protein [Azoarcus taiwanensis] |
| **03934** | 3 | GGDEF domain-containing protein [Denitromonas halophilus] |
| **09753** | 3 | hypothetical protein [Niveibacterium umoris] |
| **02536** | 3 | hypothetical protein [Azoarcus nasutitermitis] |
| **01440** | 3 | GIY-YIG nuclease family protein [Aromatoleum diolicum] |
| **09144** | 3 | TonB family protein [Zoogloea dura] |
| **09754** | 3 | DUF1566 domain-containing protein [Niveibacterium microcysteis] |
| **01441** | 3 | MULTISPECIES: ParB/RepB/Spo0J family partition protein [Betaproteobacteria] |
| **03947** | 3 | Ig-like domain repeat protein [Oryzomicrobium terrae] |
| **03953** | 3 | DUF4124 domain-containing protein [Aromatoleum toluclasticum] |
| **08962** | 3 | MULTISPECIES: HPr-rel-A system PqqD family peptide chaperone [Niveibacterium] |
| **03968** | 3 | hypothetical protein [Aromatoleum aromaticum] |
| **08059** | 3 | type II secretion system protein N [Candidatus Dactylopiibacterium carminicum] |
| **01446** | 3 | hypothetical protein [Aromatoleum aromaticum] |
| **03969** | 3 | hypothetical protein [Aromatoleum aromaticum] |
| **07846** | 3 | DUF2303 family protein [Aromatoleum pumilum] |
| **01244** | 3 | TolC family protein [Candidatus Dactylopiibacterium carminicum] |
| **09141** | 3 | hypothetical protein, partial [Denitromonas halophilus] |
| **07843** | 3 | LuxR C-terminal-related transcriptional regulator [Azoarcus indigens] |
| **07842** | 3 | DUF1656 domain-containing protein [Azoarcus indigens] |
| **07862** | 3 | Fic family protein [Azoarcus olearius] |
| **09147** | 3 | multiubiquitin domain-containing protein [Azoarcus communis] |
| **08956** | 3 | PspC domain-containing protein [Niveibacterium microcysteis] |
| **09653** | 3 | hypothetical protein, partial [Azoarcus halotolerans] |
| **07878** | 3 | chlorite dismutase family protein [Denitromonas halophilus] |
| **09748** | 3 | hypothetical protein [Niveibacterium microcysteis] |
| **02526** | 3 | hypothetical protein [Azoarcus olearius] |
| **07876** | 3 | ethanolamine utilization protein EutQ [Denitromonas halophilus] |
| **08041** | 3 | hypothetical protein [Azoarcus sp. DD4] |
| **09749** | 3 | hypothetical protein [Niveibacterium microcysteis] |
| **02192** | 3 | hypothetical protein [Nitrogeniibacter aestuarii] |
| **03908** | 3 | hypothetical protein [Propionivibrio dicarboxylicus] |
| **07872** | 3 | hypothetical protein [Zoogloea dura] |
| **03923** | 3 | toll/interleukin-1 receptor domain-containing protein [Denitromonas halophilus] |
| **01430** | 3 | MerR family transcriptional regulator [Denitromonas sp. IR12] |
| **02195** | 3 | diguanylate cyclase, partial [Aromatoleum diolicum] |
| **03638** | 3 | AGE family epimerase/isomerase [Uliginosibacterium gangwonense] |
| **02529** | 3 | MULTISPECIES: hypothetical protein [Proteobacteria] |
| **03915** | 3 | AAA family ATPase [Zoogloea ramigera] |
| **07868** | 3 | hypothetical protein [Azoarcus sp. DD4] |
| **08954** | 3 | hypothetical protein [Niveibacterium microcysteis] |
| **07866** | 3 | amino acid synthesis family protein [Nitrogeniibacter aestuarii] |
| **03869** | 3 | YcxB family protein [Azospira oryzae] |
| **03655** | 3 | hypothetical protein [Aromatoleum tolulyticum] |
| **09739** | 3 | DUF2268 domain-containing putative Zn-dependent protease [Niveibacterium microcysteis] |
| **01326** | 3 | hypothetical protein [Aromatoleum diolicum] |
| **03725** | 3 | hypothetical protein [Azoarcus indigens] |
| **03726** | 3 | DoxX family protein [Azoarcus indigens] |
| **03727** | 3 | hypothetical protein [Azoarcus indigens] |
| **08010** | 3 | hypothetical protein [Niveibacterium microcysteis] |
| **07973** | 3 | DUF4114 domain-containing protein [Niveibacterium microcysteis] |
| **07972** | 3 | SDR family oxidoreductase [Niveibacterium microcysteis] |
| **03728** | 3 | STAS domain-containing protein [Azoarcus indigens] |
| **01374** | 3 | DNA damage-inducible protein D [Thauera butanivorans] |
| **07969** | 3 | hypothetical protein [Azoarcus taiwanensis] |
| **09174** | 3 | hypothetical protein [Denitromonas halophilus] |
| **03682** | 3 | toprim domain-containing protein [Denitromonas sp. IR12] |
| **03729** | 3 | AHH domain-containing protein, partial [Azoarcus indigens] |
| **07966** | 3 | hypothetical protein [Candidatus Dactylopiibacterium carminicum] |
| **03735** | 3 | hypothetical protein [Aromatoleum aromaticum] |
| **03748** | 3 | DUF1622 domain-containing protein [Azoarcus olearius] |
| **09700** | 3 | PEP-CTERM sorting domain-containing protein [Azoarcus olearius] |
| **03681** | 3 | hypothetical protein [Thauera humireducens] |
| **03762** | 3 | MULTISPECIES: hypothetical protein [Proteobacteria] |
| **01361** | 3 | hypothetical protein [Aromatoleum pumilum] |
| **02492** | 3 | DUF1269 domain-containing protein [Nitrogeniibacter aestuarii] |
| **08017** | 3 | excisionase [Azoarcus communis] |
| **09683** | 3 | hypothetical protein [Aromatoleum bremense] |
| **03694** | 3 | FAD-dependent oxidoreductase [Aromatoleum pumilum] |
| **02171** | 3 | hypothetical protein [Aromatoleum buckelii] |
| **01347** | 3 | type II toxin-antitoxin system prevent-host-death family antitoxin [Azoarcus nasutitermitis] |
| **09680** | 3 | hypothetical protein [Aromatoleum tolulyticum] |
| **07995** | 3 | hypothetical protein [Propionivibrio dicarboxylicus] |
| **03690** | 3 | hypothetical protein [Aromatoleum aromaticum] |
| **08002** | 3 | hypothetical protein [Niveibacterium microcysteis] |
| **08004** | 3 | pyridoxamine 5'-phosphate oxidase family protein [Niveibacterium microcysteis] |
| **01357** | 3 | DUF1186 domain-containing protein [Zoogloea ramigera] |
| **09692** | 3 | PEP-CTERM sorting domain-containing protein [Azoarcus indigens] |

| | | |
|---|---|---|
| 03700 | 3 | hypothetical protein [Azoarcus nasutitermitis] |
| 08915 | 3 | YiiD C-terminal domain-containing protein [Candidatus Dactylopiibacterium carminicum] |
| 01345 | 3 | BrnA antitoxin family protein [Aromatoleum pumilum] |
| 01343 | 3 | hypothetical protein [Aromatoleum toluvorans] |
| 08916 | 3 | MEKHLA domain-containing protein [Oryzomicrobium terrae] |
| 03707 | 3 | hypothetical protein [Azoarcus indigens] |
| 09172 | 3 | hypothetical protein [Denitromonas halophilus] |
| 09691 | 3 | hypothetical protein [Azoarcus indigens] |
| 08015 | 3 | hypothetical protein [Nitrogeniibacter aestuarii] |
| 02172 | 3 | hypothetical protein [Aromatoleum aromaticum] |
| 03855 | 3 | hypothetical protein, partial [Azoarcus communis] |
| 01404 | 3 | GFA family protein [Nitrogeniibacter aestuarii] |
| 01290 | 3 | polysaccharide pyruvyl transferase family protein [Niveibacterium microcysteis] |
| 07928 | 3 | GNAT family N-acetyltransferase [Azoarcus taiwanensis] |
| 02471 | 3 | hypothetical protein [Aromatoleum aromaticum] |
| 03664 | 3 | TlpA family protein disulfide reductase [Thauera butanivorans] |
| 09725 | 3 | translocation/assembly module TamB domain-containing protein [Candidatus Dactylopiibacterium carminicum] |
| 03812 | 3 | hypothetical protein [Thauera butanivorans] |
| 03816 | 3 | hypothetical protein [Azoarcus olearius] |
| 01282 | 3 | hypothetical protein [Nitrogeniibacter aestuarii] |
| 09164 | 3 | hypothetical protein [Denitromonas halophilus] |
| 01293 | 3 | MULTISPECIES: hypothetical protein [Niveibacterium] |
| 08034 | 3 | DUF2195 family protein [Nitrogeniibacter aestuarii] |
| 01276 | 3 | hypothetical protein [Nitrogeniibacter aestuarii] |
| 08933 | 3 | caspase family protein [Aromatoleum evansii] |
| 07910 | 3 | hypothetical protein [Aromatoleum buckelii] |
| 09737 | 3 | hypothetical protein [Niveibacterium microcysteis] |
| 03653 | 3 | MULTISPECIES: metalloregulator ArsR/SmtB family transcription factor [Thauera] |
| 02463 | 3 | hypothetical protein [Niveibacterium microcysteis] |
| 07905 | 3 | hypothetical protein [Nitrogeniibacter aestuarii] |
| 08029 | 3 | autotransporter domain-containing protein [Azoarcus nasutitermitis] |
| 03800 | 3 | hypothetical protein [Azoarcus taiwanensis] |
| 08020 | 3 | MULTISPECIES: Nif11 family protein [Niveibacterium] |
| 09712 | 3 | LanC-like protein [Niveibacterium microcysteis] |
| 01380 | 3 | hypothetical protein [Azoarcus communis] |
| 02174 | 3 | MULTISPECIES: PRTRC system protein F [Betaproteobacteria] |
| 02485 | 3 | hypothetical protein [Azoarcus sp. DD4] |
| 01382 | 3 | hypothetical protein [Zoogloea dura] |
| 01302 | 3 | hypothetical protein [Denitromonas halophilus] |
| 07950 | 3 | hypothetical protein [Zoogloea dura] |
| 07948 | 3 | phasin family protein [Zoogloea dura] |
| 01297 | 3 | hypothetical protein [Aromatoleum bremense] |
| 08924 | 3 | HAMP domain-containing protein [Niveibacterium microcysteis] |
| 08926 | 3 | major capsid protein [Aromatoleum toluvorans] |
| 03781 | 3 | 2-hydroxycarboxylate transporter family protein [Uliginosibacterium gangwonense] |
| 03782 | 3 | cell division protein FtsI [Candidatus Dactylopiibacterium carminicum] |
| 03666 | 3 | hypothetical protein [Thauera aminoaromatica] |
| 02481 | 3 | hypothetical protein [Niveibacterium sp. COAC-50] |
| 09715 | 3 | MULTISPECIES: hypothetical protein [Niveibacterium] |
| 07938 | 3 | site-specific DNA-methyltransferase [Aromatoleum toluolicum] |
| 03665 | 3 | type III-B CRISPR module-associated protein Cmr5 [Thauera butanivorans] |
| 01294 | 3 | hypothetical protein [Niveibacterium microcysteis] |
| 01311 | 3 | macro domain-containing protein [Aromatoleum bremense] |
| 02903 | 3 | hypothetical protein [Thauera lacus] |
| 03977 | 3 | flagellar hook-length control protein FliK [Aromatoleum evansii] |
| 04278 | 3 | hypothetical protein [Nitrogeniibacter aestuarii] |
| 07659 | 3 | site-specific integrase [Azoarcus sp. L1K30] |
| 04284 | 3 | SRPBCC domain-containing protein [Uliginosibacterium aquaticum] |
| 07656 | 3 | hypothetical protein [Aromatoleum bremense] |
| 09865 | 3 | DUF600 family protein [Azoarcus sp. L1K30] |
| 09032 | 3 | hypothetical protein [Aromatoleum pumilum] |
| 09033 | 3 | LuxR C-terminal-related transcriptional regulator [Aromatoleum toluclasticum] |
| 02259 | 3 | hypothetical protein [Nitrogeniibacter aestuarii] |
| 02262 | 3 | hypothetical protein [Niveibacterium microcysteis] |
| 04304 | 3 | type I-E CRISPR-associated protein Cse1/CasA [Aromatoleum diolicum] |
| 01079 | 3 | DUF1566 domain-containing protein [Azoarcus indigens] |
| 09037 | 3 | hypothetical protein [Aromatoleum aromaticum] |
| 01078 | 3 | hypothetical protein [Azoarcus indigens] |
| 01076 | 3 | hypothetical protein [Azoarcus indigens] |
| 02326 | 3 | hypothetical protein [Aromatoleum buckelii] |
| 09871 | 3 | glycosyltransferase [Azoarcus olearius] |
| 07641 | 3 | PEP-CTERM sorting domain-containing protein [Nitrogeniibacter aestuarii] |
| 02324 | 3 | hypothetical protein [Azoarcus rhizosphaerae] |
| 02266 | 3 | SEL1-like repeat protein [Niveibacterium umoris] |
| 02322 | 3 | cupin domain-containing protein [Denitromonas halophilus] |
| 04322 | 3 | hypothetical protein [Propionivibrio dicarboxylicus] |
| 09090 | 3 | DUF2569 family protein [Azoarcus communis] |
| 04325 | 3 | zinc-ribbon domain-containing protein [Nitrogeniibacter aestuarii] |
| 07630 | 3 | hypothetical protein [Niveibacterium microcysteis] |
| 09099 | 3 | class I SAM-dependent methyltransferase [Azospira oryzae] |
| 07667 | 3 | hypothetical protein [Aromatoleum aromaticum] |
| 01058 | 3 | hypothetical protein [Aromatoleum pumilum] |
| 09856 | 3 | host-nuclease inhibitor Gam family protein [Aromatoleum pumilum] |
| 07697 | 3 | GyrI-like domain-containing protein [Nitrogeniibacter aestuarii] |
| 09022 | 3 | cupin domain-containing protein [Aromatoleum pumilum] |
| 01100 | 3 | hypothetical protein [Azoarcus olearius] |
| 04244 | 3 | hypothetical protein [Azoarcus nasutitermitis] |
| 09843 | 3 | AAA family ATPase [Aromatoleum toluclasticum] |
| 04245 | 3 | hypothetical protein [Nitrogeniibacter aestuarii] |
| 07691 | 3 | hypothetical protein [Azoarcus halotolerans] |
| 04246 | 3 | hypothetical protein [Niveibacterium microcysteis] |
| 04247 | 3 | MULTISPECIES: hypothetical protein [Niveibacterium] |
| 04249 | 3 | hypothetical protein [Niveibacterium microcysteis] |
| 04251 | 3 | DUF4124 domain-containing protein [Niveibacterium microcysteis] |

| | | |
|---|---|---|
| 09845 | 3 | DUF3604 domain-containing protein [Aromatoleum toluclasticum] |
| 09846 | 3 | GIY-YIG nuclease family protein [Aromatoleum toluclasticum] |
| 01095 | 3 | hypothetical protein [Aromatoleum evansii] |
| 07680 | 3 | DUF2183 domain-containing protein [Aromatoleum buckelii] |
| 07679 | 3 | tetratricopeptide repeat protein [Azospira oryzae] |
| 01092 | 3 | helix-turn-helix domain-containing protein [Azoarcus nasutitermitis] |
| 01091 | 3 | GIY-YIG nuclease family protein [Azoarcus nasutitermitis] |
| 04265 | 3 | phage tail protein [Azoarcus sp. L1K30] |
| 02256 | 3 | acetoacetate decarboxylase [Nitrogeniibacter aestuarii] |
| 09029 | 3 | Ig-like domain-containing protein [Niveibacterium sp. COAC-50] |
| 04271 | 3 | cupin domain-containing protein [Aromatoleum toluolicum] |
| 02258 | 3 | site-specific DNA-methyltransferase [Aromatoleum evansii] |
| 02274 | 3 | hypothetical protein [Aromatoleum bremense] |
| 04329 | 3 | aminopeptidase P family N-terminal domain-containing protein, partial [Aromatoleum toluclasticum] |
| 01240 | 3 | radical SAM protein [Candidatus Dactylopiibacterium carminicum] |
| 07589 | 3 | MULTISPECIES: stable inheritance protein KleA [Bacteria] |
| 09903 | 3 | MULTISPECIES: hypothetical protein [Zoogloeaceae] |
| 04392 | 3 | anti-sigma regulatory factor [Azoarcus indigens] |
| 01021 | 3 | DUF5455 family protein [Azoarcus communis] |
| 09065 | 3 | efflux RND transporter permease subunit, partial [Aromatoleum evansii] |
| 09066 | 3 | SUMF1/EgtB/PvdO family nonheme iron enzyme [Aromatoleum buckelii] |
| 09067 | 3 | AAA family ATPase [Azoarcus nasutitermitis] |
| 07580 | 3 | tRNA lysidine(34) synthetase TilS, partial [Aromatoleum diolicum] |
| 09079 | 3 | MULTISPECIES: hypothetical protein [Betaproteobacteria] |
| 01014 | 3 | TIR domain-containing protein [Thauera humireducens] |
| 09070 | 3 | hypothetical protein [Aromatoleum evansii] |
| 09078 | 3 | DUF2357 domain-containing protein [Thauera aromatica] |
| 04406 | 3 | hypothetical protein [Azoarcus olearius] |
| 09077 | 3 | DUF3577 domain-containing protein [Azoarcus taiwanensis] |
| 07571 | 3 | ABC transporter ATP-binding protein [Azoarcus sp. KH32C] |
| 04408 | 3 | serine protease [Azoarcus communis] |
| 04411 | 3 | hypothetical protein [Azoarcus olearius] |
| 04412 | 3 | alpha/beta hydrolase [Aromatoleum tolulyticum] |
| 09910 | 3 | hypothetical protein [Azoarcus indigens] |
| 09911 | 3 | CdiI family contact-dependent growth inhibition immunity protein [Azoarcus indigens] |
| 04414 | 3 | radical SAM protein [Uliginosibacterium gangwonense] |
| 04417 | 3 | hypothetical protein [Zoogloea dura] |
| 01009 | 3 | DNA phosphorothioation system sulfurtransferase DndC [Thauera linaloolentis] |
| 04429 | 3 | hypothetical protein [Denitromonas halophilus] |
| 04387 | 3 | hypothetical protein [Azoarcus indigens] |
| 04384 | 3 | hypothetical protein [Azoarcus sp. KH32C] |
| 04333 | 3 | hypothetical protein [Denitromonas halophilus] |
| 02304 | 3 | DUF2523 domain-containing protein [Aromatoleum toluvorans] |
| 02277 | 3 | hypothetical protein [Aromatoleum bremense] |
| 09051 | 3 | DUF3168 domain-containing protein [Aromatoleum aromaticum] |
| 02278 | 3 | gamma-mobile-trio protein GmtX [Aromatoleum petrolei] |
| 01040 | 3 | major capsid protein [Aromatoleum pumilum] |
| 04348 | 3 | hypothetical protein, partial [Thauera butanivorans] |
| 07614 | 3 | hypothetical protein [Niveibacterium umoris] |
| 02283 | 3 | SH3 domain-containing protein [Azoarcus communis] |
| 04349 | 3 | hypothetical protein [Aromatoleum toluclasticum] |
| 01036 | 3 | AAA family ATPase [Aromatoleum buckelii] |
| 09894 | 3 | hypothetical protein [Oryzomicrobium terrae] |
| 02285 | 3 | DUF3396 domain-containing protein [Azoarcus communis] |
| 09055 | 3 | serine protease inhibitor ecotin [Azoarcus nasutitermitis] |
| 09056 | 3 | hypothetical protein [Azoarcus sp. DD4] |
| 07606 | 3 | GDSL-type esterase/lipase family protein [Niveibacterium microcysteis] |
| 01026 | 3 | efflux RND transporter permease subunit [Azoarcus taiwanensis] |
| 02311 | 3 | hypothetical protein [Denitromonas halophilus] |
| 02310 | 3 | DUF4124 domain-containing protein [Zoogloea dura] |
| 04375 | 3 | type VI secretion system-associated protein TagF [Azoarcus communis] |
| 04377 | 3 | transposase, partial [Azoarcus communis] |
| 07598 | 3 | TolC family protein, partial [Aromatoleum evansii] |
| 07596 | 3 | chemotaxis protein CheW [Azoarcus sp. TTM-91] |
| 07595 | 3 | hypothetical protein [Azoarcus indigens] |
| 04380 | 3 | hypothetical protein [Thauera hydrothermalis] |
| 09109 | 3 | hypothetical protein [Azoarcus indigens] |
| 04241 | 3 | hypothetical protein [Azoarcus indigens] |
| 04240 | 3 | type VI secretion system lipoprotein TssJ [Azoarcus indigens] |
| 01214 | 3 | IS1 family transposase [Zoogloea oleivorans] |
| 04041 | 3 | DUF4442 domain-containing protein [Niveibacterium sp. COAC-50] |
| 02413 | 3 | hypothetical protein [Niveibacterium microcysteis] |
| 09785 | 3 | hypothetical protein [Aromatoleum toluclasticum] |
| 08062 | 3 | MULTISPECIES: hypothetical protein [Azospira] |
| 01209 | 3 | LTA synthase family protein [Azoarcus sp. KH32C] |
| 01202 | 3 | DUF3592 domain-containing protein [Azoarcus indigens] |
| 07797 | 3 | hypothetical protein [Aromatoleum evansii] |
| 01197 | 3 | universal stress protein [Aromatoleum toluolicum] |
| 07794 | 3 | hypothetical protein [Aromatoleum aromaticum] |
| 01191 | 3 | GTP-binding protein [Aromatoleum aromaticum] |
| 04058 | 3 | STAS domain-containing protein [Azoarcus rhizosphaerae] |
| 02204 | 3 | CstA-like transporter-associated (seleno)protein [Azoarcus indigens] |
| 02410 | 3 | PDZ domain-containing protein [Candidatus Dactylopiibacterium carminicum] |
| 09794 | 3 | hypothetical protein [Azospira oryzae] |
| 04066 | 3 | carbohydrate kinase family protein [Niveibacterium microcysteis] |
| 04067 | 3 | beta-xylosidase [Niveibacterium sp. COAC-50] |
| 09797 | 3 | restriction endonuclease [Nitrogeniibacter aestuarii] |
| 07781 | 3 | hypothetical protein [Aromatoleum pumilum] |
| 08975 | 3 | winged helix DNA-binding domain-containing protein [Denitromonas halophilus] |
| 04073 | 3 | LPXTG cell wall anchor domain-containing protein [Nitrogeniibacter aestuarii] |
| 08977 | 3 | hypothetical protein [Denitromonas halophilus] |
| 02408 | 3 | DUF1887 family protein [Zoogloea ramigera] |
| 04081 | 3 | MULTISPECIES: hypothetical protein [Niveibacterium] |
| 04040 | 3 | hypothetical protein [Niveibacterium microcysteis] |

| | | |
|---|---|---|
| **04037** | 3 | DEAD/DEAH box helicase family protein [Niveibacterium sp. COAC-50] |
| **01104** | 3 | malonate transporter subunit MadM [Aromatoleum petrolei] |
| **01217** | 3 | glycosyltransferase [Aromatoleum tolulyticum] |
| **07838** | 3 | YbjN domain-containing protein [Azoarcus indigens] |
| **03987** | 3 | class I SAM-dependent methyltransferase [Azoarcus taiwanensis] |
| **02201** | 3 | MarR family transcriptional regulator [Azoarcus indigens] |
| **01237** | 3 | MULTISPECIES: helix-turn-helix domain-containing protein [Niveibacterium] |
| **04002** | 3 | ATP-binding protein [Zoogloea oleivorans] |
| **07832** | 3 | MULTISPECIES: hypothetical protein [Azospira] |
| **09766** | 3 | nucleotidyltransferase [Aromatoleum pumilum] |
| **01236** | 3 | Imm50 family immunity protein [Niveibacterium microcysteis] |
| **02419** | 3 | hypothetical protein [Azoarcus olearius] |
| **04017** | 3 | methyl-accepting chemotaxis protein [Azoarcus indigens] |
| **01234** | 3 | GNAT family N-acetyltransferase [Niveibacterium microcysteis] |
| **01229** | 3 | hypothetical protein [Niveibacterium microcysteis] |
| **07822** | 3 | hypothetical protein [Azoarcus communis] |
| **09774** | 3 | hypothetical protein, partial [Thauera lacus] |
| **07820** | 3 | hypothetical protein [Azoarcus communis] |
| **04028** | 3 | hypothetical protein [Candidatus Dactylopiibacterium carminicum] |
| **01219** | 3 | hypothetical protein [Candidatus Dactylopiibacterium carminicum] |
| **02418** | 3 | MULTISPECIES: cytochrome c [Niveibacterium] |
| **07816** | 3 | PepSY domain-containing protein [Azoarcus nasutitermitis] |
| **04034** | 3 | MULTISPECIES: hypothetical protein [Niveibacterium] |
| **09778** | 3 | hypothetical protein [Nitrogeniibacter aestuarii] |
| **02414** | 3 | MULTISPECIES: phasin family protein [Niveibacterium] |
| **04036** | 3 | MULTISPECIES: hypothetical protein [Niveibacterium] |
| **02211** | 3 | hypothetical protein [Aromatoleum buckelii] |
| **07775** | 3 | BREX-3 system phosphatase PglZ [Thauera butanivorans] |
| **07774** | 3 | DUF6521 family protein [Aromatoleum evansii] |
| **04084** | 3 | hypothetical protein [Niveibacterium umoris] |
| **04183** | 3 | hypothetical protein [Thauera humireducens] |
| **02228** | 3 | DUF885 domain-containing protein [Niveibacterium sp. COAC-50] |
| **02230** | 3 | DUF2147 domain-containing protein [Niveibacterium microcysteis] |
| **07727** | 3 | PAS domain S-box protein [Thauera terpenica] |
| **07726** | 3 | MULTISPECIES: hypothetical protein [Aromatoleum] |
| **07725** | 3 | hypothetical protein [Azoarcus indigens] |
| **02372** | 3 | hypothetical protein [Azoarcus communis] |
| **07723** | 3 | Ku protein [Aromatoleum bremense] |
| **09008** | 3 | hypothetical protein [Aromatoleum aromaticum] |
| **07720** | 3 | zinc ribbon domain-containing protein [Aromatoleum aromaticum] |
| **01115** | 3 | hypothetical protein [Denitromonas halophilus] |
| **07718** | 3 | hypothetical protein [Aromatoleum aromaticum] |
| **02231** | 3 | MULTISPECIES: hypothetical protein [Niveibacterium] |
| **09010** | 3 | MULTISPECIES: hypothetical protein [Bacteria] |
| **01112** | 3 | hypothetical protein [Azoarcus communis] |
| **07715** | 3 | hypothetical protein [Aromatoleum diolicum] |
| **04205** | 3 | hypothetical protein [Thauera propionica] |
| **04206** | 3 | DUF1902 domain-containing protein [Aromatoleum evansii] |
| **02352** | 3 | MULTISPECIES: hypothetical protein [Bacteria] |
| **04226** | 3 | hypothetical protein [Azoarcus indigens] |
| **07709** | 3 | MoaD/ThiS family protein [Propionivibrio dicarboxylicus] |
| **04227** | 3 | hypothetical protein [Aromatoleum petrolei] |
| **01109** | 3 | c-type cytochrome [Thauera phenylacetica] |
| **01127** | 3 | S24 family peptidase [Aromatoleum aromaticum] |
| **01129** | 3 | hypothetical protein [Denitromonas halophilus] |
| **09819** | 3 | ISAs1 family transposase [Aromatoleum toluclasticum] |
| **04105** | 3 | hypothetical protein, partial [Nitrogeniibacter aestuarii] |
| **09132** | 3 | hypothetical protein [Zoogloea dura] |
| **02399** | 3 | extracellular solute-binding protein [Azoarcus indigens] |
| **04095** | 3 | hypothetical protein [Aromatoleum petrolei] |
| **02397** | 3 | hypothetical protein [Azoarcus indigens] |
| **02216** | 3 | hypothetical protein [Uliginosibacterium aquaticum] |
| **07768** | 3 | DNA (cytosine-5-)-methyltransferase [Azospira oryzae] |
| **02220** | 3 | GNAT family N-acetyltransferase [Propionivibrio dicarboxylicus] |
| **01168** | 3 | exodeoxyribonuclease VII small subunit [Aromatoleum evansii] |
| **07763** | 3 | PAAR domain-containing protein [Aromatoleum toluclasticum] |
| **07762** | 3 | FHA domain-containing protein [Uliginosibacterium aquaticum] |
| **01161** | 3 | hypothetical protein [Aromatoleum aromaticum] |
| **04153** | 3 | putative DNA modification/repair radical SAM protein [Nitrogeniibacter aestuarii] |
| **04119** | 3 | [citrate (pro-3S)-lyase] ligase [Propionivibrio dicarboxylicus] |
| **02224** | 3 | hypothetical protein [Niveibacterium microcysteis] |
| **01159** | 3 | hypothetical protein [Aromatoleum aromaticum] |
| **07754** | 3 | PD-(D/E)XK nuclease family protein [Azoarcus sp. KH32C] |
| **04139** | 3 | M48 family metallopeptidase [Azoarcus halotolerans] |
| **02380** | 3 | hypothetical protein [Azoarcus taiwanensis] |
| **09120** | 3 | DUF4365 domain-containing protein [Aromatoleum tolulyticum] |
| **07746** | 3 | ABC transporter ATP-binding protein [Uliginosibacterium aquaticum] |
| **09815** | 3 | acetate--CoA ligase family protein [Thauera aminoaromatica] |
| **07740** | 3 | BRCT domain-containing protein [Thauera chlorobenzoica] |
| **01448** | 3 | AAA family ATPase, partial [Aromatoleum buckelii] |
| **02116** | 3 | DUF4365 domain-containing protein [Denitromonas halophilus] |
| **03620** | 3 | MULTISPECIES: hypothetical protein [Aromatoleum] |
| **08447** | 3 | hypothetical protein [Candidatus Dactylopiibacterium carminicum] |
| **03157** | 3 | hypothetical protein [Oryzomicrobium terrae] |
| **01794** | 3 | hypothetical protein [Azoarcus indigens] |
| **08725** | 3 | hypothetical protein [Aromatoleum pumilum] |
| **01791** | 3 | restriction endonuclease subunit S [Azoarcus sp. TTM-91] |
| **01790** | 3 | YCF48-related protein, partial [Aromatoleum toluolicum] |
| **08434** | 3 | VPA1269 family protein [Aromatoleum petrolei] |
| **08726** | 3 | hypothetical protein [Aromatoleum toluclasticum] |
| **09443** | 3 | hypothetical protein [Aromatoleum evansii] |
| **08430** | 3 | molybdenum cofactor biosynthesis F family protein [Azoarcus sp. KH32C] |
| **08427** | 3 | hypothetical protein [Candidatus Dactylopiibacterium carminicum] |
| **08729** | 3 | NF038130 family PEP-CTERM protein [Zoogloea ramigera] |

| | | |
|---|---|---|
| 02755 | 3 | GtrA family protein [Aromatoleum pumilum] |
| 01783 | 3 | DNA-binding protein [Candidatus Dactylopiibacterium carminicum] |
| 02754 | 3 | hypothetical protein [Candidatus Dactylopiibacterium carminicum] |
| 01988 | 3 | MULTISPECIES: elongation factor Tu, partial [Proteobacteria] |
| 03181 | 3 | head-tail connector protein [Aromatoleum aromaticum] |
| 01991 | 3 | DNA-binding protein [Thauera propionica] |
| 01776 | 3 | GGDEF domain-containing protein [Azoarcus communis] |
| 09272 | 3 | cellulase family glycosylhydrolase [Uliginosibacterium aquaticum] |
| 08416 | 3 | PRTRC system protein A [Aromatoleum aromaticum] |
| 09271 | 3 | MULTISPECIES: ferrous iron transport protein A [Uliginosibacterium] |
| 03197 | 3 | hypothetical protein [Aromatoleum evansii] |
| 08739 | 3 | hypothetical protein [Thauera humireducens] |
| 03153 | 3 | DUF2934 domain-containing protein [Azoarcus taiwanensis] |
| 08448 | 3 | family 43 glycosylhydrolase [Uliginosibacterium aquaticum] |
| 02741 | 3 | DUF1493 family protein [Azoarcus indigens] |
| 08724 | 3 | hypothetical protein [Aromatoleum toluolicum] |
| 01846 | 3 | hypothetical protein [Azoarcus communis] |
| 09289 | 3 | hypothetical protein [Niveibacterium microcysteis] |
| 03099 | 3 | DUF4440 domain-containing protein [Niveibacterium microcysteis] |
| 08483 | 3 | MULTISPECIES: hypothetical protein [Bacteria] |
| 08482 | 3 | MULTISPECIES: lipoprotein [Proteobacteria] |
| 03113 | 3 | transposase, partial [Denitromonas sp. IR12] |
| 09287 | 3 | glycosyltransferase [Niveibacterium sp. COAC-50] |
| 03118 | 3 | hypothetical protein [Nitrogeniibacter aestuarii] |
| 03121 | 3 | hypothetical protein [Aromatoleum toluclasticum] |
| 09416 | 3 | NAD(P)/FAD-dependent oxidoreductase, partial [Thauera terpenica] |
| 03124 | 3 | ATP-grasp domain-containing protein [Aromatoleum toluclasticum] |
| 08466 | 3 | site-specific integrase [Azoarcus taiwanensis] |
| 09284 | 3 | DUF885 domain-containing protein [Niveibacterium sp. COAC-50] |
| 08462 | 3 | hypothetical protein [Thauera butanivorans] |
| 09283 | 3 | hypothetical protein [Niveibacterium microcysteis] |
| 09281 | 3 | hypothetical protein [Niveibacterium microcysteis] |
| 02771 | 3 | class I SAM-dependent methyltransferase [Azoarcus taiwanensis] |
| 01815 | 3 | hypothetical protein [Azoarcus sp. DD4] |
| 09428 | 3 | hypothetical protein [Azoarcus olearius] |
| 02766 | 3 | DUF1996 domain-containing protein [Niveibacterium microcysteis] |
| 02765 | 3 | AraC family transcriptional regulator ligand-binding domain-containing protein [Niveibacterium microcysteis] |
| 08452 | 3 | hypothetical protein [Aromatoleum toluclasticum] |
| 01804 | 3 | hypothetical protein [Candidatus Dactylopiibacterium carminicum] |
| 01767 | 3 | MULTISPECIES: SIR2 family protein [Rhodocyclales] |
| 09459 | 3 | MFS transporter [Azoarcus communis] |
| 02538 | 3 | hypothetical protein [Aromatoleum toluvorans] |
| 08755 | 3 | hypothetical protein [Thauera butanivorans] |
| 01711 | 3 | hypothetical protein [Azoarcus communis] |
| 02722 | 3 | DUF2007 domain-containing protein [Aromatoleum pumilum] |
| 03245 | 3 | DUF4062 domain-containing protein [Zoogloea ramigera] |
| 08366 | 3 | glycosyl hydrolase family 17 protein [Niveibacterium umoris] |
| 03249 | 3 | hypothetical protein [Denitromonas halophilus] |
| 08363 | 3 | hypothetical protein [Niveibacterium microcysteis] |
| 03255 | 3 | hypothetical protein [Azoarcus sp. DD4] |
| 02711 | 3 | HAMP domain-containing protein, partial [Azoarcus communis] |
| 02710 | 3 | hypothetical protein [Azoarcus communis] |
| 01702 | 3 | diversity-generating retroelement protein Avd [Azoarcus nasutitermitis] |
| 02020 | 3 | nucleotidyltransferase domain-containing protein [Denitromonas halophilus] |
| 09249 | 3 | hypothetical protein [Aromatoleum diolicum] |
| 08353 | 3 | MULTISPECIES: type II toxin-antitoxin system prevent-host-death family antitoxin [Thauera] |
| 09492 | 3 | DUF3565 domain-containing protein [Azoarcus olearius] |
| 02699 | 3 | PEP-CTERM sorting domain-containing protein [Nitrogeniibacter aestuarii] |
| 09494 | 3 | Bro-N domain-containing protein [Oryzomicrobium terrae] |
| 03266 | 3 | cadherin domain-containing protein, partial [Denitromonas sp. IR12] |
| 03267 | 3 | hypothetical protein, partial [Denitromonas sp. IR12] |
| 01695 | 3 | hypothetical protein [Azoarcus indigens] |
| 08344 | 3 | hypothetical protein [Aromatoleum buckelii] |
| 02687 | 3 | DUF6475 domain-containing protein [Oryzomicrobium terrae] |
| 03278 | 3 | 2-iminoacetate synthase ThiH [Propionivibrio dicarboxylicus] |
| 02028 | 3 | hypothetical protein [Aromatoleum toluclasticum] |
| 09481 | 3 | hypothetical protein [Azoarcus indigens] |
| 09480 | 3 | hypothetical protein [Azoarcus indigens] |
| 08405 | 3 | DUF108 domain-containing protein [Candidatus Dactylopiibacterium carminicum] |
| 09260 | 3 | hypothetical protein [Azoarcus indigens] |
| 01759 | 3 | hypothetical protein [Aromatoleum evansii] |
| 03213 | 3 | DinB family protein [Azoarcus rhizosphaerae] |
| 09465 | 3 | hypothetical protein [Azoarcus communis] |
| 02739 | 3 | hypothetical protein [Azoarcus indigens] |
| 01749 | 3 | hypothetical protein [Azoarcus sp. L1K30] |
| 01747 | 3 | helix-turn-helix domain-containing protein, partial [Azoarcus halotolerans] |
| 03217 | 3 | MULTISPECIES: hypothetical protein [Betaproteobacteria] |
| 03218 | 3 | hypothetical protein [Azoarcus indigens] |
| 02737 | 3 | hypothetical protein [Azoarcus indigens] |
| 03226 | 3 | glycosyltransferase [Azospira oryzae] |
| 08386 | 3 | hypothetical protein [Azoarcus communis] |
| 08385 | 3 | DMT family transporter [Azoarcus communis] |
| 03228 | 3 | genomic island protein [Azoarcus indigens] |
| 03230 | 3 | hypothetical protein [Azoarcus indigens] |
| 03231 | 3 | hypothetical protein [Azoarcus indigens] |
| 03233 | 3 | type VI secretion system-associated FHA domain protein TagH [Azoarcus indigens] |
| 03234 | 3 | hypothetical protein [Azoarcus indigens] |
| 08745 | 3 | hypothetical protein [Azoarcus nasutitermitis] |
| 01721 | 3 | hypothetical protein [Denitromonas halophilus] |
| 09477 | 3 | hypothetical protein [Azoarcus indigens] |
| 08375 | 3 | hypothetical protein [Niveibacterium umoris] |
| 01717 | 3 | Hpt domain-containing protein, partial [Zoogloea ramigera] |
| 09262 | 3 | AAC(3) family N-acetyltransferase [Zoogloea ramigera] |
| 03089 | 3 | PEP-CTERM sorting domain-containing protein [Nitrogeniibacter aestuarii] |

| | | |
|---|---|---|
| 03085 | 3 | peptidoglycan-binding protein [Nitrogeniibacter aestuarii] |
| 02802 | 3 | hypothetical protein [Azospira oryzae] |
| 08649 | 3 | glycoside hydrolase [Niveibacterium sp. COAC-50] |
| 08650 | 3 | MULTISPECIES: phospholipase D-like domain-containing protein [Betaproteobacteria] |
| 08651 | 3 | hypothetical protein [Niveibacterium microcysteis] |
| 08652 | 3 | response regulator [Nitrogeniibacter aestuarii] |
| 01904 | 3 | nuclear transport factor 2 family protein [Niveibacterium microcysteis] |
| 01900 | 3 | hypothetical protein [Niveibacterium sp. COAC-50] |
| 08585 | 3 | hypothetical protein [Thauera butanivorans] |
| 02977 | 3 | DNA-binding protein [Zoogloea oleivorans] |
| 09356 | 3 | hypothetical protein [Nitrogeniibacter aestuarii] |
| 02883 | 3 | hypothetical protein [Thauera phenolivorans] |
| 08582 | 3 | hypothetical protein [Aromatoleum evansii] |
| 08581 | 3 | DUF3268 family zinc-finger domain-containing protein [Thauera propionica] |
| 08654 | 3 | GAF domain-containing protein [Nitrogeniibacter aestuarii] |
| 08577 | 3 | MULTISPECIES: TrfB-related DNA-binding protein [Proteobacteria] |
| 09324 | 3 | MULTISPECIES: hypothetical protein [Niveibacterium] |
| 02984 | 3 | hypothetical protein [Azoarcus indigens] |
| 09323 | 3 | hypothetical protein [Niveibacterium microcysteis] |
| 01943 | 3 | tetratricopeptide repeat protein [Niveibacterium microcysteis] |
| 08573 | 3 | LysR substrate-binding domain-containing protein [Aromatoleum toluclasticum] |
| 02870 | 3 | helix-destabilizing protein [Azoarcus communis] |
| 02989 | 3 | hypothetical protein [Azoarcus halotolerans] |
| 08570 | 3 | hypothetical protein [Nitrogeniibacter aestuarii] |
| 08661 | 3 | S8 family serine peptidase [Niveibacterium microcysteis] |
| 01892 | 3 | SoxR reducing system RseC family protein [Uliginosibacterium aquaticum] |
| 09352 | 3 | DUF695 domain-containing protein [Denitromonas halophilus] |
| 08594 | 3 | hypothetical protein [Aromatoleum evansii] |
| 09400 | 3 | DUF459 domain-containing protein [Candidatus Dactylopiibacterium carminicum] |
| 09325 | 3 | MarR family transcriptional regulator [Niveibacterium microcysteis] |
| 02899 | 3 | alpha/beta hydrolase [Nitrogeniibacter aestuarii] |
| 08624 | 3 | DUF4304 domain-containing protein [Niveibacterium microcysteis] |
| 01927 | 3 | hypothetical protein [Nitrogeniibacter aestuarii] |
| 09332 | 3 | glutathione S-transferase [Aromatoleum aromaticum] |
| 08620 | 3 | cytochrome c3 family protein [Niveibacterium microcysteis] |
| 02931 | 3 | hypothetical protein [Azoarcus indigens] |
| 02932 | 3 | hypothetical protein [Azoarcus indigens] |
| 01924 | 3 | hypothetical protein [Azoarcus sp. Aa7] |
| 02933 | 3 | hypothetical protein [Azoarcus indigens] |
| 02943 | 3 | SH3 domain-containing protein [Azoarcus nasutitermitis] |
| 08606 | 3 | MULTISPECIES: hypothetical protein [Betaproteobacteria] |
| 08605 | 3 | hypothetical protein [Aromatoleum diolicum] |
| 01941 | 3 | questin oxidase family protein [Niveibacterium microcysteis] |
| 02950 | 3 | Lar family restriction alleviation protein [Azoarcus communis] |
| 09327 | 3 | alpha amylase C-terminal domain-containing protein [Niveibacterium sp. COAC-50] |
| 08644 | 3 | hypothetical protein [Niveibacterium microcysteis] |
| 01942 | 3 | DUF3313 family protein [Niveibacterium microcysteis] |
| 02953 | 3 | hypothetical protein [Azoarcus olearius] |
| 08646 | 3 | hypothetical protein [Niveibacterium microcysteis] |
| 01919 | 3 | tripartite tricarboxylate transporter TctB family protein [Azoarcus taiwanensis] |
| 08647 | 3 | glutaminase A [Niveibacterium microcysteis] |
| 02958 | 3 | restriction endonuclease subunit S [Aromatoleum bremense] |
| 09348 | 3 | hypothetical protein [Nitrogeniibacter aestuarii] |
| 02995 | 3 | tetratricopeptide repeat protein [Nitrogeniibacter aestuarii] |
| 08562 | 3 | rhomboid family intramembrane serine protease [Azoarcus halotolerans] |
| 08665 | 3 | DUF2442 domain-containing protein [Thauera chlorobenzoica] |
| 02856 | 3 | META domain-containing protein [Azoarcus rhizosphaerae] |
| 08527 | 3 | hypothetical protein [Zoogloea dura] |
| 01959 | 3 | class I SAM-dependent methyltransferase [Thauera aromatica] |
| 09385 | 3 | OmpA family protein [Azoarcus olearius] |
| 01960 | 3 | hypothetical protein [Aromatoleum bremense] |
| 02826 | 3 | acyltransferase [Thauera lacus] |
| 02818 | 3 | PRTRC system ThiF family protein [Aromatoleum aromaticum] |
| 02817 | 3 | hypothetical protein [Aromatoleum aromaticum] |
| 09391 | 3 | outer membrane beta-barrel protein [Uliginosibacterium aquaticum] |
| 03062 | 3 | hypothetical protein [Aromatoleum toluclasticum] |
| 01968 | 3 | VWA domain-containing protein [Azoarcus communis] |
| 01969 | 3 | class I SAM-dependent methyltransferase [Azoarcus communis] |
| 01867 | 3 | hypothetical protein [Aromatoleum diolicum] |
| 03066 | 3 | DUF2220 family protein [Rhodocyclus purpureus] |
| 03067 | 3 | GNAT family N-acetyltransferase [Rhodocyclus purpureus] |
| 08510 | 3 | MULTISPECIES: ATPase [Proteobacteria] |
| 01972 | 3 | RHS repeat protein [Azoarcus communis] |
| 08507 | 3 | hypothetical protein [Azoarcus indigens] |
| 03075 | 3 | hypothetical protein [Niveibacterium microcysteis] |
| 08505 | 3 | hypothetical protein [Azospira oryzae] |
| 03078 | 3 | hypothetical protein [Niveibacterium microcysteis] |
| 01863 | 3 | SAM-dependent methyltransferase [Aromatoleum diolicum] |
| 03080 | 3 | hypothetical protein [Nitrogeniibacter aestuarii] |
| 08499 | 3 | chromosome partitioning protein ParA [Thauera hydrothermalis] |
| 02836 | 3 | capsule biosynthesis GfcC family protein [Azoarcus sp. KH32C] |
| 03047 | 3 | GNAT family N-acetyltransferase [Niveibacterium umoris] |
| 08531 | 3 | LysR substrate-binding domain-containing protein [Thauera butanivorans] |
| 01880 | 3 | hypothetical protein [Azoarcus indigens] |
| 01888 | 3 | hypothetical protein [Aromatoleum toluclasticum] |
| 09369 | 3 | hypothetical protein [Propionivibrio dicarboxylicus] |
| 09315 | 3 | caspase family protein [Niveibacterium microcysteis] |
| 01883 | 3 | hypothetical protein [Zoogloea dura] |
| 03009 | 3 | MULTISPECIES: TniQ family protein [Proteobacteria] |
| 09374 | 3 | hypothetical protein [Aromatoleum evansii] |
| 08551 | 3 | MarR family transcriptional regulator [Aromatoleum diolicum] |
| 03016 | 3 | beta/gamma crystallin-related protein [Aromatoleum diolicum] |
| 02853 | 3 | YHS domain-containing protein [Aromatoleum toluclasticum] |
| 03022 | 3 | hypothetical protein [Azoarcus indigens] |

| | | |
|---|---|---|
| **08546** | 3 | hypothetical protein [Nitrogeniibacter aestuarii] |
| **03044** | 3 | hypothetical protein [Niveibacterium microcysteis] |
| **01878** | 3 | DUF892 family protein [Azoarcus indigens] |
| **09379** | 3 | MULTISPECIES: hypothetical protein [Proteobacteria] |
| **01876** | 3 | hypothetical protein [Azoarcus indigens] |
| **08674** | 3 | hypothetical protein [Azoarcus indigens] |
| **01955** | 3 | hypothetical protein [Niveibacterium microcysteis] |
| **09311** | 3 | hypothetical protein [Niveibacterium microcysteis] |
| **08536** | 3 | cytochrome c family protein [Denitromonas halophilus] |
| **03041** | 3 | MULTISPECIES: hypothetical protein [Niveibacterium] |
| **03043** | 3 | IPT/TIG domain-containing protein [Niveibacterium microcysteis] |
| **01956** | 3 | hypothetical protein [Niveibacterium microcysteis] |
| **03284** | 3 | FAD-binding protein [Aromatoleum buckelii] |
| **03288** | 3 | FAD-binding protein, partial [Aromatoleum toluolicum] |
| **08334** | 3 | sugar phosphate isomerase/epimerase [Niveibacterium microcysteis] |
| **03516** | 3 | nucleotidyltransferase domain-containing protein [Niveibacterium microcysteis] |
| **02584** | 3 | DUF3596 domain-containing protein [Thauera aminoaromatica] |
| **01517** | 3 | hypothetical protein [Niveibacterium umoris] |
| **08156** | 3 | SIR2 family protein [Nitrogeniibacter mangrovi] |
| **03518** | 3 | carbon-nitrogen hydrolase family protein [Niveibacterium microcysteis] |
| **09594** | 3 | MULTISPECIES: hypothetical protein [Niveibacterium] |
| **03525** | 3 | NADAR family protein [Candidatus Dactylopiibacterium carminicum] |
| **02582** | 3 | hypothetical protein [Aromatoleum toluclasticum] |
| **09596** | 3 | MULTISPECIES: YbaY family lipoprotein [Niveibacterium] |
| **01513** | 3 | succinylglutamate desuccinylase [Niveibacterium microcysteis] |
| **03536** | 3 | 4Fe-4S dicluster domain-containing protein [Rhodocyclus purpureus] |
| **02092** | 3 | amidohydrolase family protein [Azoarcus indigens] |
| **08854** | 3 | hypothetical protein [Azoarcus halotolerans] |
| **03546** | 3 | hypothetical protein [Niveibacterium microcysteis] |
| **01502** | 3 | F420-dependent NADP oxidoreductase [Propionivibrio dicarboxylicus] |
| **08856** | 3 | FxDxF family PEP-CTERM protein [Nitrogeniibacter aestuarii] |
| **08138** | 3 | hypothetical protein [Azoarcus nasutitermitis] |
| **08137** | 3 | GspMb/PilO family protein [Nitrogeniibacter mangrovi] |
| **08136** | 3 | hypothetical protein [Aromatoleum aromaticum] |
| **08135** | 3 | hypothetical protein [Azoarcus nasutitermitis] |
| **01495** | 3 | hypothetical protein [Candidatus Dactylopiibacterium carminicum] |
| **08131** | 3 | DUF5716 family protein [Azoarcus indigens] |
| **02571** | 3 | MULTISPECIES: hypothetical protein [Rhodocyclales] |
| **02098** | 3 | DUF2283 domain-containing protein [Azoarcus nasutitermitis] |
| **02091** | 3 | hypothetical protein [Azoarcus indigens] |
| **02585** | 3 | AAA family ATPase [Thauera aromatica] |
| **08193** | 3 | hypothetical protein [Azoarcus indigens] |
| **08161** | 3 | c-type cytochrome [Niveibacterium microcysteis] |
| **03480** | 3 | MULTISPECIES: hypothetical protein [Betaproteobacteria] |
| **03483** | 3 | hypothetical protein [Aromatoleum toluclasticum] |
| **09576** | 3 | hypothetical protein [Azoarcus halotolerans] |
| **01541** | 3 | lipid-A-disaccharide synthase N-terminal domain-containing protein [Aromatoleum aromaticum] |
| **02086** | 3 | DoxX family protein [Azoarcus indigens] |
| **02087** | 3 | hypothetical protein [Azoarcus indigens] |
| **08185** | 3 | type I-E CRISPR-associated protein Cse2/CasB [Aromatoleum diolicum] |
| **03489** | 3 | hypothetical protein [Azoarcus indigens] |
| **03491** | 3 | zinc-binding dehydrogenase [Azoarcus indigens] |
| **03492** | 3 | cytochrome o ubiquinol oxidase subunit IV [Azoarcus indigens] |
| **03495** | 3 | DUF1003 domain-containing protein [Azoarcus indigens] |
| **08180** | 3 | hypothetical protein [Azoarcus communis] |
| **03497** | 3 | hypothetical protein [Azoarcus indigens] |
| **02088** | 3 | hypothetical protein [Azoarcus indigens] |
| **01528** | 3 | hypothetical protein [Azoarcus communis] |
| **01527** | 3 | hypothetical protein [Niveibacterium microcysteis] |
| **03504** | 3 | MULTISPECIES: membrane protease subunit, stomatin/prohibitin [Uliginosibacterium] |
| **01526** | 3 | hypothetical protein [Niveibacterium microcysteis] |
| **08169** | 3 | hypothetical protein [Azoarcus indigens] |
| **01518** | 3 | hypothetical protein [Niveibacterium microcysteis] |
| **09591** | 3 | MULTISPECIES: PEP-CTERM sorting domain-containing protein [Niveibacterium] |
| **03513** | 3 | thioesterase family protein [Niveibacterium microcysteis] |
| **03514** | 3 | MULTISPECIES: DUF962 domain-containing protein [Niveibacterium] |
| **01486** | 3 | DUF6445 family protein [Niveibacterium microcysteis] |
| **03561** | 3 | tetratricopeptide repeat protein [Niveibacterium microcysteis] |
| **03562** | 3 | hypothetical protein [Niveibacterium microcysteis] |
| **03564** | 3 | MULTISPECIES: hypothetical protein [Niveibacterium] |
| **08877** | 3 | hypothetical protein [Azoarcus communis] |
| **03593** | 3 | hypothetical protein [Nitrogeniibacter aestuarii] |
| **08093** | 3 | type I-U CRISPR-associated protein Csb2 [Rhodocyclus purpureus] |
| **08090** | 3 | type III-B CRISPR module RAMP protein Cmr6 [Thauera butanivorans] |
| **03598** | 3 | hypothetical protein [Aromatoleum evansii] |
| **08087** | 3 | hypothetical protein [Azoarcus communis] |
| **08082** | 3 | hypothetical protein [Azoarcus olearius] |
| **08081** | 3 | hypothetical protein [Azoarcus olearius] |
| **01456** | 3 | hypothetical protein [Azoarcus indigens] |
| **09634** | 3 | MULTISPECIES: hypothetical protein [Niveibacterium] |
| **03603** | 3 | hypothetical protein [Azoarcus rhizosphaerae] |
| **01454** | 3 | hypothetical protein [Azoarcus indigens] |
| **09638** | 3 | geranylgeranyl reductase family protein [Rhodocyclus purpureus] |
| **01453** | 3 | hypothetical protein [Azoarcus indigens] |
| **01451** | 3 | hypothetical protein [Aromatoleum petrolei] |
| **08880** | 3 | hypothetical protein [Azoarcus communis] |
| **02543** | 3 | MULTISPECIES: hypothetical protein [Betaproteobacteria] |
| **09641** | 3 | NEW3 domain-containing protein [Aromatoleum aromaticum] |
| **01449** | 3 | acyl-CoA dehydrogenase family protein, partial [Aromatoleum toluolicum] |
| **03612** | 3 | hypothetical protein [Aromatoleum tolulyticum] |
| **03617** | 3 | ATP-binding protein [Aromatoleum evansii] |
| **03618** | 3 | hypothetical protein [Aromatoleum aromaticum] |
| **02539** | 3 | hypothetical protein [Rhodocyclus purpureus] |
| **03591** | 3 | hypothetical protein [Nitrogeniibacter aestuarii] |

| | | |
|---|---|---|
| 03590 | 3 | phage baseplate assembly protein V, partial [Denitromonas sp. IR12] |
| 03589 | 3 | DUF2345 domain-containing protein, partial [Denitromonas halophilus] |
| 08110 | 3 | hypothetical protein [Aromatoleum toluclasticum] |
| 02100 | 3 | TonB-dependent receptor [Azoarcus rhizosphaerae] |
| 03565 | 3 | hypothetical protein [Aromatoleum aromaticum] |
| 03566 | 3 | class I SAM-dependent methyltransferase [Aromatoleum aromaticum] |
| 01478 | 3 | DUF4397 domain-containing protein [Niveibacterium microcysteis] |
| 01476 | 3 | hypothetical protein [Azoarcus sp. KH32C] |
| 02106 | 3 | DUF2946 family protein [Azoarcus rhizosphaerae] |
| 02107 | 3 | LamG domain-containing protein [Thauera propionica] |
| 09615 | 3 | GNAT family N-acetyltransferase [Niveibacterium umoris] |
| 03571 | 3 | TRAP transporter substrate-binding protein DctP [Aromatoleum tolulyticum] |
| 09617 | 3 | nickel/cobalt efflux transporter RcnA [Niveibacterium sp. COAC-50] |
| 09618 | 3 | hypothetical protein [Niveibacterium microcysteis] |
| 01465 | 3 | hypothetical protein [Azoarcus halotolerans] |
| 09619 | 3 | hypothetical protein [Niveibacterium microcysteis] |
| 03574 | 3 | DUF4062 domain-containing protein [Azoarcus communis] |
| 01471 | 3 | hypothetical protein [Niveibacterium microcysteis] |
| 01466 | 3 | GNAT family N-acetyltransferase [Zoogloea dura] |
| 08104 | 3 | VOC family protein [Candidatus Dactylopiibacterium carminicum] |
| 09622 | 3 | acetyl-CoA hydrolase [Niveibacterium sp. COAC-50] |
| 03580 | 3 | DUF2163 domain-containing protein [Candidatus Dactylopiibacterium carminicum] |
| 03583 | 3 | RHS domain-containing protein [Azoarcus sp. L1K30] |
| 02551 | 3 | hypothetical protein [Candidatus Dactylopiibacterium carminicum] |
| 02547 | 3 | fibronectin type III domain-containing protein [Aromatoleum bremense] |
| 01543 | 3 | site-specific integrase [Aromatoleum petrolei] |
| 09574 | 3 | capsule polysaccharide biosynthesis [Azoarcus sp. KH32C] |
| 09501 | 3 | hypothetical protein [Denitromonas halophilus] |
| 02670 | 3 | hypothetical protein [Denitromonas halophilus] |
| 01654 | 3 | acyl-CoA dehydrogenase [Aromatoleum buckelii] |
| 01647 | 3 | PEP-CTERM sorting domain-containing protein [Azoarcus halotolerans] |
| 08796 | 3 | 5'-3'-deoxyribonucleotidase [Azoarcus indigens] |
| 03333 | 3 | MULTISPECIES: IS91 family transposase [Bacteria] |
| 03339 | 3 | type III-B CRISPR module RAMP protein Cmr4 [Thauera butanivorans] |
| 01641 | 3 | helix-turn-helix domain-containing protein [Azoarcus communis] |
| 08797 | 3 | hypothetical protein [Azoarcus indigens] |
| 01632 | 3 | MULTISPECIES: hypothetical protein [Bacteria][Archaea] |
| 02046 | 3 | CRISPR-associated Cmr3 family protein [Thauera phenylacetica] |
| 08799 | 3 | hypothetical protein [Azoarcus indigens] |
| 02656 | 3 | ATP-binding protein [Azoarcus rhizosphaerae] |
| 03354 | 3 | ABC transporter permease subunit [Aromatoleum pumilum] |
| 02048 | 3 | hypothetical protein [Denitromonas sp. IR12] |
| 02650 | 3 | ATP-binding protein [Aromatoleum pumilum] |
| 08806 | 3 | MULTISPECIES: hypothetical protein [Azoarcus] |
| 01616 | 3 | UdgX family uracil-DNA binding protein [Nitrogeniibacter aestuarii] |
| 08807 | 3 | DMT family transporter [Azoarcus indigens] |
| 08269 | 3 | hypothetical protein [Aromatoleum aromaticum] |
| 03370 | 3 | hypothetical protein [Aromatoleum pumilum] |
| 08263 | 3 | hypothetical protein [Nitrogeniibacter aestuarii] |
| 01612 | 3 | reverse transcriptase family protein [Azoarcus indigens] |
| 09546 | 3 | class I SAM-dependent methyltransferase [Aromatoleum evansii] |
| 08259 | 3 | hypothetical protein [Azoarcus halotolerans] |
| 09517 | 3 | hypothetical protein [Zoogloea dura] |
| 03319 | 3 | citrate lyase subunit alpha [Propionivibrio dicarboxylicus] |
| 01544 | 3 | M10 family metallopeptidase C-terminal domain-containing protein [Azoarcus sp. Aa7] |
| 08792 | 3 | FIST C-terminal domain-containing protein [Azospira oryzae] |
| 09246 | 3 | RES domain-containing protein [Thauera lacus] |
| 08782 | 3 | P-loop NTPase fold protein [Azoarcus communis] |
| 08330 | 3 | LD-carboxypeptidase [Niveibacterium microcysteis] |
| 08783 | 3 | RHS repeat-associated core domain-containing protein, partial [Azoarcus indigens] |
| 09504 | 3 | Gfo/Idh/MocA family oxidoreductase [Denitromonas halophilus] |
| 02677 | 3 | type II toxin-antitoxin system HipA family toxin YjjJ [Aromatoleum diolicum] |
| 08786 | 3 | hypothetical protein [Azoarcus halotolerans] |
| 03297 | 3 | hypothetical protein [Thauera phenylacetica] |
| 09507 | 3 | hypothetical protein [Denitromonas halophilus] |
| 08321 | 3 | PD-(D/E)XK motif protein [Aromatoleum aromaticum] |
| 01661 | 3 | DUF3883 domain-containing protein [Aromatoleum toluclasticum] |
| 01660 | 3 | helix-turn-helix domain-containing protein [Thauera propionica] |
| 03305 | 3 | MULTISPECIES: integrase domain-containing protein [Proteobacteria] |
| 09512 | 3 | MULTISPECIES: hypothetical protein [Rhodocyclales] |
| 08315 | 3 | DEAD/DEAH box helicase [Uliginosibacterium sp. IMCC34675] |
| 08314 | 3 | MULTISPECIES: nuclear transport factor 2 family protein [Uliginosibacterium] |
| 02033 | 3 | hypothetical protein [Thauera chlorobenzoica] |
| 03307 | 3 | DNA polymerase/3'-5' exonuclease PolX [Aromatoleum bremense] |
| 02674 | 3 | hypothetical protein [Denitromonas halophilus] |
| 03312 | 3 | ASCH domain-containing protein [Niveibacterium microcysteis] |
| 09514 | 3 | hypothetical protein [Zoogloea dura] |
| 03313 | 3 | DUF4238 domain-containing protein [Aromatoleum toluolicum] |
| 02673 | 3 | hypothetical protein [Denitromonas halophilus] |
| 08258 | 3 | hypothetical protein [Thauera terpenica] |
| 08808 | 3 | heme utilization cystosolic carrier protein HutX [Azoarcus nasutitermitis] |
| 08256 | 3 | cupin domain-containing protein [Azoarcus halotolerans] |
| 08809 | 3 | KilA-N domain-containing protein [Azoarcus nasutitermitis] |
| 03425 | 3 | hypothetical protein [Denitromonas halophilus] |
| 09561 | 3 | ATP phosphoribosyltransferase regulatory subunit [Azoarcus communis] |
| 08821 | 3 | MULTISPECIES: hypothetical protein [Proteobacteria] |
| 08822 | 3 | hypothetical protein [Azoarcus olearius] |
| 03434 | 3 | glycosyltransferase family 9 protein [Zoogloea dura] |
| 08215 | 3 | hypothetical protein, partial [Azoarcus indigens] |
| 03435 | 3 | hypothetical protein [Zoogloea dura] |
| 03440 | 3 | rhodanese-related sulfurtransferase [Azoarcus communis] |
| 08212 | 3 | hypothetical protein [Azoarcus indigens] |
| 02072 | 3 | hypothetical protein [Azoarcus communis] |
| 08835 | 3 | hypothetical protein [Denitromonas halophilus] |

| | | |
|---|---|---|
| 01561 | 3 | penicillin-insensitive murein endopeptidase [Nitrogeniibacter aestuarii] |
| 08209 | 3 | extensin family protein [Azoarcus indigens] |
| 01558 | 3 | methyltransferase domain-containing protein [Aromatoleum pumilum] |
| 01557 | 3 | hypothetical protein [Denitromonas halophilus] |
| 09569 | 3 | hypothetical protein [Azoarcus indigens] |
| 03458 | 3 | RES family NAD+ phosphorylase [Aromatoleum bremense] |
| 03460 | 3 | phage tail length tape measure family protein [Aromatoleum toluolicum] |
| 08201 | 3 | hypothetical protein [Azoarcus nasutitermitis] |
| 02602 | 3 | hypothetical protein [Thauera aromatica] |
| 02595 | 3 | hypothetical protein [Azoarcus olearius] |
| 01550 | 3 | hypothetical protein [Azoarcus sp. L1K30] |
| 02077 | 3 | hypothetical protein [Aromatoleum evansii] |
| 03423 | 3 | DUF1275 domain-containing protein [Denitromonas halophilus] |
| 01575 | 3 | hypothetical protein [Propionivibrio dicarboxylicus] |
| 03422 | 3 | hypothetical protein [Denitromonas halophilus] |
| 03398 | 3 | hypothetical protein [Azoarcus indigens] |
| 08253 | 3 | hypothetical protein [Azospira oryzae] |
| 02649 | 3 | hypothetical protein [Azoarcus rhizosphaerae] |
| 03383 | 3 | RHS repeat-associated core domain-containing protein, partial [Azoarcus communis] |
| 01596 | 3 | DUF2285 domain-containing protein [Aromatoleum toluclasticum] |
| 03394 | 3 | hypothetical protein [Azoarcus indigens] |
| 03396 | 3 | hypothetical protein [Azoarcus indigens] |
| 03397 | 3 | STAS domain-containing protein [Azoarcus indigens] |
| 02644 | 3 | hypothetical protein [Azoarcus indigens] |
| 08244 | 3 | hypothetical protein [Zoogloea dura] |
| 08242 | 3 | hypothetical protein [Zoogloea dura] |
| 08814 | 3 | DUF4258 domain-containing protein [Azoarcus nasutitermitis] |
| 08224 | 3 | hypothetical protein [Zoogloea dura] |
| 08239 | 3 | DUF2845 domain-containing protein [Azoarcus sp. TTM-91] |
| 03399 | 3 | hypothetical protein [Azoarcus indigens] |
| 02640 | 3 | 3-hydroxyanthranilate 3,4-dioxygenase [Azoarcus indigens] |
| 03408 | 3 | hypothetical protein [Aromatoleum pumilum] |
| 08234 | 3 | hypothetical protein [Denitromonas halophilus] |
| 08233 | 3 | hypothetical protein [Denitromonas halophilus] |
| 03415 | 3 | ThiF family adenylyltransferase [Aromatoleum aromaticum] |
| 03420 | 3 | hypothetical protein [Aromatoleum tolulyticum] |
| 08226 | 3 | hypothetical protein [Zoogloea dura] |
| 08225 | 3 | hypothetical protein [Propionivibrio dicarboxylicus] |
| 00001 | 3 | hypothetical protein [Aromatoleum toluclasticum] |
| 04267 | 3 | hypothetical protein [Aromatoleum pumilum] |
| 09914 | 3 | hypothetical protein [Azoarcus nasutitermitis] |
| 05455 | 3 | hypothetical protein [Azoarcus indigens] |
| 10306 | 3 | hypothetical protein [Denitromonas sp. IR12] |
| 06855 | 3 | CRISPR-associated endonuclease Cas1 [Thauera butanivorans] |
| 10308 | 3 | S8 family serine peptidase [Candidatus Dactylopiibacterium carminicum] |
| 06850 | 3 | hypothetical protein [Aromatoleum pumilum] |
| 05484 | 3 | hypothetical protein [Denitromonas halophilus] |
| 05491 | 3 | hypothetical protein [Denitromonas halophilus] |
| 05498 | 3 | hypothetical protein [Zoogloea oleivorans] |
| 05507 | 3 | hypothetical protein [Thauera phenolivorans] |
| 06836 | 3 | cupin domain-containing protein [Aromatoleum toluclasticum] |
| 05518 | 3 | hypothetical protein [Aromatoleum bremense] |
| 00508 | 3 | phage tail assembly chaperone family protein, TAC [Aromatoleum aromaticum] |
| 05521 | 3 | histidine phosphatase family protein [Aromatoleum toluclasticum] |
| 10321 | 3 | hypothetical protein [Azoarcus communis] |
| 05528 | 3 | 1-acyl-sn-glycerol-3-phosphate acyltransferase [Aromatoleum aromaticum] |
| 10322 | 3 | DUF5675 family protein [Azoarcus communis] |
| 05534 | 3 | hypothetical protein [Aromatoleum toluvorans] |
| 05540 | 3 | potassium channel family protein [Aromatoleum diolicum] |
| 05549 | 3 | hypothetical protein [Azoarcus communis] |
| 05550 | 3 | metallophosphoesterase [Azoarcus indigens] |
| 05554 | 3 | hypothetical protein [Zoogloea dura] |
| 00501 | 3 | hypothetical protein [Aromatoleum toluclasticum] |
| 06813 | 3 | hypothetical protein [Aromatoleum evansii] |
| 10330 | 3 | LuxR C-terminal-related transcriptional regulator [Azoarcus olearius] |
| 06810 | 3 | SDR family NAD(P)-dependent oxidoreductase [Zoogloea dura] |
| 05560 | 3 | RpiB/LacA/LacB family sugar-phosphate isomerase [Candidatus Dactylopiibacterium carminicum] |
| 05463 | 3 | AAA domain-containing protein [Azoarcus communis] |
| 00542 | 3 | DISARM system phospholipase D-like protein DrmC [Aromatoleum toluclasticum] |
| 06807 | 3 | hypothetical protein [Thauera humireducens] |
| 00554 | 3 | hypothetical protein [Azospira oryzae] |
| 00596 | 3 | aspartate/glutamate racemase family protein [Denitromonas halophilus] |
| 05324 | 3 | Gfo/Idh/MocA family oxidoreductase [Aromatoleum pumilum] |
| 06922 | 3 | hypothetical protein [Zoogloea dura] |
| 00591 | 3 | hypothetical protein [Zoogloea ramigera] |
| 05347 | 3 | MULTISPECIES: flippase [Thauera] |
| 00588 | 3 | cyclophilin-like fold protein [Azoarcus sp. DD4] |
| 10269 | 3 | ankyrin repeat domain-containing protein [Azospira oryzae] |
| 00571 | 3 | agglutinin biogenesis protein MshP [Azospira oryzae] |
| 10283 | 3 | hypothetical protein [Azoarcus olearius] |
| 05401 | 3 | MULTISPECIES: type I toxin-antitoxin system ptaRNA1 family toxin [Proteobacteria] |
| 10287 | 3 | class I SAM-dependent methyltransferase [Azoarcus olearius] |
| 06886 | 3 | polysaccharide pyruvyl transferase family protein [Azoarcus communis] |
| 00562 | 3 | prepilin-type N-terminal cleavage/methylation domain-containing protein [Azoarcus indigens] |
| 06880 | 3 | DUF4123 domain-containing protein [Nitrogeniibacter aestuarii] |
| 10293 | 3 | MULTISPECIES: hypothetical protein [Thauera] |
| 06878 | 3 | PAS domain-containing protein [Thauera lacus] |
| 05427 | 3 | DUF2761 domain-containing protein [Thauera terpenica] |
| 06876 | 3 | hypothetical protein [Zoogloea dura] |
| 06875 | 3 | TonB family protein [Aromatoleum pumilum] |
| 10294 | 3 | CHAD domain-containing protein [Aromatoleum bremense] |
| 05433 | 3 | helix-turn-helix domain-containing protein [Aromatoleum tolulyticum] |
| 00558 | 3 | hypothetical protein [Azoarcus indigens] |
| 10296 | 3 | ABC transporter substrate-binding protein [Thauera chlorobenzoica] |

| | | |
|---|---|---|
| 05441 | 3 | hypothetical protein [Azoarcus nasutitermitis] |
| 06869 | 3 | hypothetical protein [Thauera propionica] |
| 00497 | 3 | hypothetical protein [Aromatoleum toluclasticum] |
| 05569 | 3 | MULTISPECIES: PsiF family protein [Niveibacterium] |
| 05721 | 3 | YeeE/YedE family protein [Azoarcus halotolerans] |
| 10354 | 3 | TIGR03790 family protein [Azoarcus communis] |
| 05637 | 3 | glutamate--tRNA ligase family protein, partial [Aromatoleum toluclasticum] |
| 05644 | 3 | MBL fold metallo-hydrolase [Propionivibrio dicarboxylicus] |
| 10356 | 3 | ArsR family transcriptional regulator [Azoarcus communis] |
| 06751 | 3 | GNAT family N-acetyltransferase [Niveibacterium microcysteis] |
| 06750 | 3 | MULTISPECIES: helix-hairpin-helix domain-containing protein [Niveibacterium] |
| 05654 | 3 | TIGR04255 family protein [Azoarcus indigens] |
| 05658 | 3 | PRTRC system ParB family protein [Aromatoleum aromaticum] |
| 06747 | 3 | MULTISPECIES: hypothetical protein [Niveibacterium] |
| 00443 | 3 | helix-turn-helix domain-containing protein [Azoarcus indigens] |
| 00439 | 3 | recombinase RecT [Aromatoleum toluolicum] |
| 06736 | 3 | DUF2804 domain-containing protein [Niveibacterium sp. COAC-50] |
| 06731 | 3 | glycosyltransferase [Niveibacterium microcysteis] |
| 00431 | 3 | UDP-N-acetylglucosamine 2-epimerase [Azoarcus communis] |
| 05687 | 3 | DUF4349 domain-containing protein [Niveibacterium microcysteis] |
| 05692 | 3 | MULTISPECIES: phage capsid protein [Bacteria][Archaea] |
| 00428 | 3 | MULTISPECIES: hypothetical protein [Bacteria] |
| 10371 | 3 | PKD domain-containing protein [Uliginosibacterium gangwonense] |
| 10372 | 3 | hypothetical protein [Azospira oryzae] |
| 00427 | 3 | hypothetical protein [Azoarcus sp. KH32C] |
| 06719 | 3 | hypothetical protein [Aromatoleum aromaticum] |
| 06718 | 3 | GNAT family N-acetyltransferase [Candidatus Dactylopiibacterium carminicum] |
| 05714 | 3 | hypothetical protein [Denitromonas sp. IR12] |
| 10374 | 3 | hypothetical protein [Azospira oryzae] |
| 10375 | 3 | PD-(D/E)XK motif protein [Azospira oryzae] |
| 00425 | 3 | MULTISPECIES: hypothetical protein [Betaproteobacteria] |
| 00447 | 3 | hypothetical protein [Zoogloea dura] |
| 05631 | 3 | TIGR00341 family protein [Nitrogeniibacter aestuarii] |
| 05571 | 3 | MerC domain-containing protein [Niveibacterium sp. COAC-50] |
| 06759 | 3 | transposase [Aromatoleum bremense] |
| 06800 | 3 | MULTISPECIES: DUF2188 domain-containing protein [Proteobacteria] |
| 05572 | 3 | M48 family metalloprotease [Niveibacterium microcysteis] |
| 10338 | 3 | AbrB/MazE/SpoVT family DNA-binding domain-containing protein [Thauera chlorobenzoica] |
| 06792 | 3 | hypothetical protein [Aromatoleum aromaticum] |
| 00473 | 3 | MULTISPECIES: hypothetical protein [Thauera] |
| 10340 | 3 | transposase [Azoarcus communis] |
| 10341 | 3 | hypothetical protein [Aromatoleum toluclasticum] |
| 05593 | 3 | hydroxylamine reductase [Propionivibrio dicarboxylicus] |
| 05596 | 3 | N-acylglucosamine 2-epimerase [Niveibacterium sp. COAC-50] |
| 05597 | 3 | hypothetical protein [Niveibacterium microcysteis] |
| 05598 | 3 | XRE family transcriptional regulator [Aromatoleum bremense] |
| 05599 | 3 | hypothetical protein [Niveibacterium microcysteis] |
| 05600 | 3 | Dabb family protein [Niveibacterium microcysteis] |
| 05601 | 3 | hypothetical protein [Niveibacterium microcysteis] |
| 00466 | 3 | hypothetical protein [Aromatoleum evansii] |
| 05602 | 3 | hypothetical protein [Niveibacterium microcysteis] |
| 00457 | 3 | polyhydroxyalkanoic acid system family protein [Niveibacterium microcysteis] |
| 06773 | 3 | substrate-binding domain-containing protein [Azoarcus sp. L1K30] |
| 05604 | 3 | hypothetical protein [Niveibacterium microcysteis] |
| 06770 | 3 | hypothetical protein [Azoarcus communis] |
| 05611 | 3 | MULTISPECIES: hypothetical protein [Niveibacterium] |
| 00453 | 3 | hypothetical protein [Azospira oryzae] |
| 05620 | 3 | DUF2007 domain-containing protein [Azoarcus communis] |
| 05622 | 3 | MarR family transcriptional regulator [Thauera phenolivorans] |
| 05626 | 3 | hypothetical protein [Denitromonas sp. IR12] |
| 10256 | 3 | hypothetical protein [Aromatoleum toluclasticum] |
| 06930 | 3 | glycosyltransferase family 2 protein [Denitromonas halophilus] |
| 06932 | 3 | hypothetical protein [Denitromonas halophilus] |
| 10164 | 3 | MULTISPECIES: hypothetical protein [Niveibacterium] |
| 10166 | 3 | MULTISPECIES: hypothetical protein [Niveibacterium] |
| 05076 | 3 | T6SS effector amidase Tae4 family protein [Thauera butanivorans] |
| 00677 | 3 | acetyltransferase [Azoarcus indigens] |
| 10172 | 3 | hypothetical protein [Niveibacterium microcysteis] |
| 10173 | 3 | tryptophan 2,3-dioxygenase [Niveibacterium microcysteis] |
| 07088 | 3 | hypothetical protein [Azoarcus halotolerans] |
| 05086 | 3 | CopD family protein [Thauera terpenica] |
| 05087 | 3 | AbrB family transcriptional regulator [Thauera chlorobenzoica] |
| 07083 | 3 | hypothetical protein [Denitromonas sp. IR12] |
| 10178 | 3 | type II secretion system protein GspM [Propionivibrio dicarboxylicus] |
| 00664 | 3 | hypothetical protein [Aromatoleum diolicum] |
| 07074 | 3 | 4-amino-4-deoxy-L-arabinose-phospho-UDP flippase [Niveibacterium umoris] |
| 07073 | 3 | MULTISPECIES: hypothetical protein [Aromatoleum] |
| 00663 | 3 | hypothetical protein [Aromatoleum bremense] |
| 10182 | 3 | hypothetical protein [Azoarcus sp. Aa7] |
| 05114 | 3 | hypothetical protein [Candidatus Dactylopiibacterium carminicum] |
| 07066 | 3 | AMP-binding protein, partial [Azoarcus indigens] |
| 07065 | 3 | DUF1998 domain-containing protein [Aromatoleum bremense] |
| 05117 | 3 | AAA family ATPase [Rhodocyclus purpureus] |
| 05119 | 3 | HipA domain-containing protein [Rhodocyclus purpureus] |
| 07061 | 3 | MULTISPECIES: hypothetical protein [Niveibacterium] |
| 05123 | 3 | MULTISPECIES: hypothetical protein [Azospira] |
| 07058 | 3 | hypothetical protein [Niveibacterium umoris] |
| 10187 | 3 | hypothetical protein [Aromatoleum toluclasticum] |
| 05131 | 3 | hypothetical protein [Azoarcus indigens] |
| 05066 | 3 | hypothetical protein [Thauera phenylacetica] |
| 10163 | 3 | N-succinylarginine dihydrolase [Niveibacterium microcysteis] |
| 05316 | 3 | DUF2946 domain-containing protein [Nitrogeniibacter aestuarii] |
| 10161 | 3 | energy transducer TonB [Azospira oryzae] |
| 07155 | 3 | type II toxin-antitoxin system CcdA family antitoxin [Thauera aromatica] |

| | | |
|---|---|---|
| 00719 | 3 | beta-3-deoxy-D-manno-oct-2-ulosonic acid transferase [Zoogloea dura] |
| 00714 | 3 | hemin-degrading factor [Azoarcus nasutitermitis] |
| 10139 | 3 | GYD domain-containing protein [Azoarcus sp. Aa7] |
| 05006 | 3 | citrate:proton symporter [Propionivibrio dicarboxylicus] |
| 07140 | 3 | hypothetical protein [Aromatoleum toluolicum] |
| 10141 | 3 | ABC transporter permease subunit [Aromatoleum toluclasticum] |
| 07138 | 3 | SRPBCC family protein [Niveibacterium microcysteis] |
| 00709 | 3 | hypothetical protein [Niveibacterium microcysteis] |
| 05018 | 3 | MULTISPECIES: hypothetical protein [Rhodocyclales] |
| 05019 | 3 | hypothetical protein [Azoarcus olearius] |
| 00708 | 3 | hypothetical protein [Niveibacterium microcysteis] |
| 07132 | 3 | restriction endonuclease subunit S [Aromatoleum aromaticum] |
| 05026 | 3 | hypothetical protein [Aromatoleum bremense] |
| 05027 | 3 | hemerythrin domain-containing protein [Thauera aminoaromatica] |
| 00705 | 3 | hypothetical protein [Niveibacterium microcysteis] |
| 10148 | 3 | hypothetical protein [Azoarcus indigens] |
| 00703 | 3 | hypothetical protein [Niveibacterium microcysteis] |
| 07123 | 3 | hypothetical protein [Azoarcus olearius] |
| 10150 | 3 | MFS transporter [Azoarcus indigens] |
| 07119 | 3 | MULTISPECIES: PRTRC system protein B [Betaproteobacteria] |
| 05042 | 3 | DUF2939 domain-containing protein [Oryzomicrobium terrae] |
| 05046 | 3 | phage terminase large subunit [Uliginosibacterium gangwonense] |
| 00694 | 3 | EAL domain-containing protein [Niveibacterium microcysteis] |
| 05056 | 3 | MBL fold metallo-hydrolase [Aromatoleum toluclasticum] |
| 05134 | 3 | hypothetical protein [Azoarcus indigens] |
| 05135 | 3 | hypothetical protein [Azoarcus sp. TTM-91] |
| 05139 | 3 | DUF6430 domain-containing protein [Azoarcus nasutitermitis] |
| 10189 | 3 | light-harvesting protein [Rhodocyclus purpureus] |
| 10226 | 3 | MULTISPECIES: hypothetical protein [Niveibacterium] |
| 00624 | 3 | hypothetical protein [Azoarcus halotolerans] |
| 05249 | 3 | CAP domain-containing protein [Candidatus Dactylopiibacterium carminicum] |
| 00608 | 3 | anti-sigma factor [Zoogloea dura] |
| 05257 | 3 | DJ-1/PfpI family protein [Candidatus Dactylopiibacterium carminicum] |
| 05259 | 3 | CHAP domain-containing protein [Niveibacterium microcysteis] |
| 10236 | 3 | IPTL-CTERM sorting domain-containing protein [Azoarcus sp. L1K30] |
| 00605 | 3 | hypothetical protein [Nitrogeniibacter aestuarii] |
| 05273 | 3 | tetratricopeptide repeat protein [Rhodocyclus purpureus] |
| 05275 | 3 | Fic family protein [Azoarcus nasutitermitis] |
| 00603 | 3 | hypothetical protein [Denitromonas halophilus] |
| 06960 | 3 | tyrosine-type recombinase/integrase, partial [Aromatoleum buckelii] |
| 06959 | 3 | MULTISPECIES: transcriptional regulator [Proteobacteria] |
| 05281 | 3 | extracellular solute-binding protein [Niveibacterium microcysteis] |
| 05282 | 3 | DUF2087 domain-containing protein [Niveibacterium microcysteis] |
| 05284 | 3 | PDZ domain-containing protein [Niveibacterium sp. COAC-50] |
| 06954 | 3 | M6 family metalloprotease domain-containing protein [Azoarcus halotolerans] |
| 05287 | 3 | DUF3365 domain-containing protein [Nitrogeniibacter aestuarii] |
| 05289 | 3 | 2OG-Fe(II) oxygenase [Nitrogeniibacter aestuarii] |
| 06949 | 3 | phosphocholine cytidylyltransferase family protein [Denitromonas halophilus] |
| 06947 | 3 | pyruvate phosphate dikinase [Denitromonas halophilus] |
| 05294 | 3 | MULTISPECIES: hypothetical protein [Niveibacterium] |
| 05296 | 3 | DnaJ domain-containing protein [Aromatoleum bremense] |
| 05311 | 3 | hypothetical protein [Aromatoleum diolicum] |
| 05315 | 3 | IS4 family transposase, partial [Denitromonas sp. IR12] |
| 00627 | 3 | PliI family lysozyme inhibitor of I-type lysozyme [Nitrogeniibacter aestuarii] |
| 06989 | 3 | DUF1799 domain-containing protein [Aromatoleum pumilum] |
| 10222 | 3 | phage tail sheath family protein [Niveibacterium microcysteis] |
| 00645 | 3 | hypothetical protein [Aromatoleum evansii] |
| 05148 | 3 | class I SAM-dependent methyltransferase [Azoarcus rhizosphaerae] |
| 05153 | 3 | hypothetical protein [Azoarcus olearius] |
| 10195 | 3 | DUF2635 domain-containing protein [Azoarcus indigens] |
| 00648 | 3 | hypothetical protein [Azoarcus communis] |
| 05158 | 3 | hypothetical protein [Zoogloea dura] |
| 05164 | 3 | hypothetical protein [Zoogloea dura] |
| 05169 | 3 | aminoglycoside phosphotransferase family protein [Denitromonas halophilus] |
| 07027 | 3 | hypothetical protein [Candidatus Dactylopiibacterium carminicum] |
| 05171 | 3 | hypothetical protein [Denitromonas halophilus] |
| 05174 | 3 | hypothetical protein [Denitromonas halophilus] |
| 07024 | 3 | glycine zipper 2TM domain-containing protein [Azoarcus halotolerans] |
| 05179 | 3 | hypothetical protein [Denitromonas halophilus] |
| 10221 | 3 | MULTISPECIES: hypothetical protein [Niveibacterium] |
| 05183 | 3 | MULTISPECIES: antirestriction protein [Bacteria] |
| 10207 | 3 | hypothetical protein [Niveibacterium microcysteis] |
| 07014 | 3 | DUF6250 domain-containing protein [Uliginosibacterium gangwonense] |
| 07011 | 3 | hypothetical protein [Candidatus Dactylopiibacterium carminicum] |
| 05197 | 3 | hypothetical protein [Azoarcus sp. TTM-91] |
| 05208 | 3 | hypothetical protein [Nitrogeniibacter aestuarii] |
| 05209 | 3 | three-Cys-motif partner protein TcmP [Azoarcus sp. M9-3-2] |
| 05214 | 3 | hypothetical protein [Aromatoleum aromaticum] |
| 10211 | 3 | cyclic nucleotide-binding domain-containing protein [Nitrogeniibacter aestuarii] |
| 05225 | 3 | DUF2130 domain-containing protein [Aromatoleum evansii] |
| 06994 | 3 | MFS transporter [Azoarcus sp. DD4] |
| 06711 | 3 | sugar nucleotide-binding protein [Niveibacterium microcysteis] |
| 06708 | 3 | MULTISPECIES: hypothetical protein [Proteobacteria] |
| 04991 | 3 | hypothetical protein [Nitrogeniibacter aestuarii] |
| 06434 | 3 | hypothetical protein [Thauera phenylacetica] |
| 06052 | 3 | hypothetical protein [Azoarcus rhizosphaerae] |
| 00157 | 3 | hypothetical protein [Niveibacterium microcysteis] |
| 06424 | 3 | polysaccharide pyruvyl transferase family protein [Aromatoleum aromaticum] |
| 00143 | 3 | anti-sigma regulatory factor [Azoarcus indigens] |
| 06068 | 3 | hypothetical protein [Denitromonas halophilus] |
| 06421 | 3 | cupin domain-containing protein [Denitromonas halophilus] |
| 06419 | 3 | PEP-CTERM sorting domain-containing protein [Denitromonas halophilus] |
| 06417 | 3 | FAD-binding protein [Denitromonas halophilus] |
| 06072 | 3 | hypothetical protein [Denitromonas halophilus] |

| | | |
|---|---|---|
| **06075** | 3 | hypothetical protein [Denitromonas halophilus] |
| **06080** | 3 | DUF3693 domain-containing protein [Azoarcus olearius] |
| **00136** | 3 | hypothetical protein [Aromatoleum toluolicum] |
| **06408** | 3 | MFS transporter [Nitrogeniibacter aestuarii] |
| **10549** | 3 | 2-dehydro-3-deoxygalactonokinase [Niveibacterium microcysteis] |
| **00134** | 3 | DUF2384 domain-containing protein [Aromatoleum aromaticum] |
| **06089** | 3 | DUF262 domain-containing protein [Azoarcus olearius] |
| **10551** | 3 | anhydro-N-acetylmuramic acid kinase [Niveibacterium microcysteis] |
| **06095** | 3 | hypothetical protein [Aromatoleum tolulyticum] |
| **06096** | 3 | hypothetical protein [Azoarcus indigens] |
| **00123** | 3 | DUF1850 domain-containing protein [Aromatoleum pumilum] |
| **06104** | 3 | contact-dependent growth inhibition system immunity protein [Denitromonas sp. IR12] |
| **00121** | 3 | alpha/beta fold hydrolase [Nitrogeniibacter mangrovi] |
| **00115** | 3 | hypothetical protein [Azoarcus communis] |
| **00110** | 3 | hypothetical protein [Aromatoleum evansii] |
| **10562** | 3 | hypothetical protein [Nitrogeniibacter aestuarii] |
| **06051** | 3 | hypothetical protein [Azoarcus rhizosphaerae] |
| **06438** | 3 | hypothetical protein [Aromatoleum diolicum] |
| **06117** | 3 | DUF2730 family protein [Azoarcus communis] |
| **06048** | 3 | Clp protease ClpP [Aromatoleum aromaticum] |
| **00218** | 3 | MULTISPECIES: ogr/Delta-like zinc finger family protein [Rhodocyclaceae] |
| **05995** | 3 | YdaS family helix-turn-helix protein [Azoarcus nasutitermitis] |
| **00213** | 3 | hypothetical protein [Azoarcus communis] |
| **05999** | 3 | PIN domain-containing protein [Azoarcus nasutitermitis] |
| **10515** | 3 | hypothetical protein [Niveibacterium microcysteis] |
| **06472** | 3 | hypothetical protein [Azoarcus communis] |
| **06003** | 3 | hypothetical protein, partial [Thauera butanivorans] |
| **00205** | 3 | MULTISPECIES: hypothetical protein [Niveibacterium] |
| **10518** | 3 | hypothetical protein [Niveibacterium microcysteis] |
| **00200** | 3 | hypothetical protein [Niveibacterium umoris] |
| **06010** | 3 | hypothetical protein [Thauera propionica] |
| **10525** | 3 | MULTISPECIES: hypothetical protein [Uliginosibacterium] |
| **06014** | 3 | type II toxin-antitoxin system VapC family toxin [Aromatoleum pumilum] |
| **00191** | 3 | polysaccharide deacetylase family protein [Niveibacterium microcysteis] |
| **06018** | 3 | hypothetical protein [Azoarcus rhizosphaerae] |
| **06455** | 3 | hypothetical protein [Nitrogeniibacter aestuarii] |
| **06021** | 3 | hypothetical protein [Azoarcus nasutitermitis] |
| **06022** | 3 | hypothetical protein [Aromatoleum tolulyticum] |
| **06024** | 3 | hypothetical protein [Nitrogeniibacter aestuarii] |
| **00185** | 3 | type II and III secretion system protein [Azoarcus indigens] |
| **00179** | 3 | ABC transporter permease subunit [Rhodocyclus purpureus] |
| **10533** | 3 | hypothetical protein [Rhodocyclus purpureus] |
| **06038** | 3 | hypothetical protein [Azoarcus indigens] |
| **06039** | 3 | hypothetical protein [Azoarcus indigens] |
| **06440** | 3 | hypothetical protein [Azoarcus nasutitermitis] |
| **06387** | 3 | hypothetical protein [Azoarcus halotolerans] |
| **06384** | 3 | major capsid protein [Azoarcus indigens] |
| **10379** | 3 | hypothetical protein [Azoarcus indigens] |
| **00051** | 3 | VRR-NUC domain-containing protein [Azoarcus indigens] |
| **06193** | 3 | GNAT family N-acetyltransferase [Niveibacterium microcysteis] |
| **06195** | 3 | hypothetical protein [Niveibacterium microcysteis] |
| **06326** | 3 | DNA-3-methyladenine glycosylase I [Azoarcus nasutitermitis] |
| **00044** | 3 | helix-turn-helix domain-containing protein [Propionivibrio dicarboxylicus] |
| **06198** | 3 | hypothetical protein [Nitrogeniibacter aestuarii] |
| **06200** | 3 | hypothetical protein [Nitrogeniibacter aestuarii] |
| **10599** | 3 | membrane lipoprotein lipid attachment site-containing protein [Nitrogeniibacter aestuarii] |
| **06313** | 3 | MULTISPECIES: DUF2283 domain-containing protein [Thauera] |
| **06207** | 3 | hypothetical protein [Rhodocyclus purpureus] |
| **06209** | 3 | hypothetical protein [Niveibacterium microcysteis] |
| **06218** | 3 | FecR family protein [Nitrogeniibacter aestuarii] |
| **10608** | 3 | hypothetical protein [Aromatoleum diolicum] |
| **06225** | 3 | MULTISPECIES: hypothetical protein [Niveibacterium] |
| **06227** | 3 | MULTISPECIES: hypothetical protein [Niveibacterium] |
| **06228** | 3 | hypothetical protein [Niveibacterium microcysteis] |
| **06229** | 3 | MULTISPECIES: YgdI/YgdR family lipoprotein [Niveibacterium] |
| **00031** | 3 | hypothetical protein [Azoarcus nasutitermitis] |
| **06292** | 3 | type III secretion system outer membrane ring subunit SctC [Azoarcus rhizosphaerae] |
| **00017** | 3 | MULTISPECIES: conjugal transfer protein TrbD [Proteobacteria] |
| **06286** | 3 | hypothetical protein [Aromatoleum bremense] |
| **06284** | 3 | ATP-binding domain-containing protein, partial [Aromatoleum toluvorans] |
| **06280** | 3 | YbfB/YjiJ family MFS transporter, partial [Thauera phenylacetica] |
| **06279** | 3 | hypothetical protein [Aromatoleum toluvorans] |
| **10621** | 3 | hypothetical protein [Aromatoleum pumilum] |
| **06267** | 3 | hypothetical protein [Aromatoleum aromaticum] |
| **06192** | 3 | Nramp family divalent metal transporter [Niveibacterium microcysteis] |
| **10593** | 3 | hypothetical protein [Aromatoleum evansii] |
| **06383** | 3 | SRPBCC domain-containing protein [Azoarcus indigens] |
| **06187** | 3 | hypothetical protein [Rhodocyclus purpureus] |
| **06382** | 3 | hypothetical protein [Azoarcus indigens] |
| **06118** | 3 | 2Fe-2S iron-sulfur cluster binding domain-containing protein [Azoarcus communis] |
| **06122** | 3 | hypothetical protein [Azoarcus taiwanensis] |
| **06125** | 3 | hypothetical protein [Aromatoleum bremense] |
| **06128** | 3 | MULTISPECIES: SPW repeat protein [Rhodocyclales] |
| **10564** | 3 | hypothetical protein [Azoarcus sp. DD4] |
| **00097** | 3 | OadG family protein [Azoarcus communis] |
| **06145** | 3 | hypothetical protein [Azoarcus indigens] |
| **10568** | 3 | MULTISPECIES: hypothetical protein [Niveibacterium] |
| **06149** | 3 | GNAT family N-acetyltransferase [Propionivibrio dicarboxylicus] |
| **06150** | 3 | serine endopeptidase [Uliginosibacterium aquaticum] |
| **10571** | 3 | transporter [Niveibacterium microcysteis] |
| **00088** | 3 | class I SAM-dependent methyltransferase [Propionivibrio dicarboxylicus] |
| **06159** | 3 | SPOR domain-containing protein [Azospira oryzae] |
| **00081** | 3 | hypothetical protein [Denitromonas halophilus] |
| **06164** | 3 | hypothetical protein [Niveibacterium microcysteis] |

| | | |
|---|---|---|
| 06358 | 3 | KAP family NTPase [Azoarcus communis] |
| 06172 | 3 | endo alpha-1,4 polygalactosaminidase [Niveibacterium microcysteis] |
| 06352 | 3 | hypothetical protein [Aromatoleum pumilum] |
| 06351 | 3 | hypothetical protein [Azoarcus rhizosphaerae] |
| 06181 | 3 | DEAD/DEAH box helicase family protein [Azoarcus sp. Aa7] |
| 06183 | 3 | ATP-binding protein [Rhodocyclus purpureus] |
| 10587 | 3 | hypothetical protein [Azoarcus communis] |
| 00052 | 3 | type VI secretion system-associated protein TagO [Azoarcus indigens] |
| 10592 | 3 | hypothetical protein [Azoarcus rhizosphaerae] |
| 05991 | 3 | DUF2726 domain-containing protein [Thauera humireducens] |
| 10509 | 3 | deoxyribose-phosphate aldolase [Aromatoleum bremense] |
| 06485 | 3 | hypothetical protein, partial [Azoarcus taiwanensis] |
| 06658 | 3 | DUF441 domain-containing protein [Candidatus Dactylopiibacterium carminicum] |
| 10414 | 3 | hypothetical protein [Denitromonas halophilus] |
| 10415 | 3 | hypothetical protein [Denitromonas halophilus] |
| 00363 | 3 | DUF2220 family protein [Azoarcus indigens] |
| 06642 | 3 | serine protease [Azoarcus communis] |
| 10421 | 3 | hypothetical protein [Denitromonas halophilus] |
| 00352 | 3 | hypothetical protein, partial [Azoarcus communis] |
| 06636 | 3 | M15 family metallopeptidase [Azoarcus olearius] |
| 05821 | 3 | sel1 repeat family protein [Aromatoleum diolicum] |
| 05823 | 3 | GNAT family N-acetyltransferase [Aromatoleum tolulyticum] |
| 00350 | 3 | hypothetical protein [Azoarcus olearius] |
| 05830 | 3 | ferritin-like domain-containing protein [Azoarcus indigens] |
| 06629 | 3 | hypothetical protein [Azoarcus olearius] |
| 06627 | 3 | DUF3106 domain-containing protein [Azoarcus olearius] |
| 05834 | 3 | hypothetical protein [Aromatoleum toluolicum] |
| 06625 | 3 | DUF6364 family protein [Aromatoleum aromaticum] |
| 06622 | 3 | hypothetical protein [Aromatoleum aromaticum] |
| 10427 | 3 | hypothetical protein [Zoogloea dura] |
| 05845 | 3 | MULTISPECIES: anti-virulence regulator CigR family protein [Niveibacterium] |
| 06616 | 3 | DUF3124 domain-containing protein [Azospira oryzae] |
| 06615 | 3 | hypothetical protein [Azoarcus communis] |
| 05850 | 3 | class II fructose-bisphosphatase [Niveibacterium microcysteis] |
| 05851 | 3 | MULTISPECIES: phenylalanine 4-monooxygenase [Niveibacterium] |
| 06607 | 3 | hypothetical protein [Aromatoleum toluclasticum] |
| 10434 | 3 | MFS transporter, partial [Thauera aminoaromatica] |
| 06605 | 3 | hypothetical protein [Nitrogeniibacter aestuarii] |
| 06647 | 3 | DUF3373 domain-containing protein [Propionivibrio dicarboxylicus] |
| 06661 | 3 | TerD family protein [Aromatoleum aromaticum] |
| 10506 | 3 | pectate lyase [Uliginosibacterium aquaticum] |
| 06662 | 3 | hypothetical protein [Uliginosibacterium gangwonense] |
| 05729 | 3 | HAD family hydrolase [Azoarcus nasutitermitis] |
| 05731 | 3 | hypothetical protein [Thauera butanivorans] |
| 06701 | 3 | transposase, partial [Azoarcus sp. Aa7] |
| 00421 | 3 | hypothetical protein [Aromatoleum bremense] |
| 00419 | 3 | restriction endonuclease [Aromatoleum evansii] |
| 05744 | 3 | hypothetical protein [Zoogloea dura] |
| 10384 | 3 | hypothetical protein [Azoarcus indigens] |
| 00410 | 3 | Ig-like domain-containing protein [Azoarcus halotolerans] |
| 05761 | 3 | TerB N-terminal domain-containing protein [Aromatoleum toluclasticum] |
| 05764 | 3 | DUF2779 domain-containing protein [Aromatoleum evansii] |
| 05766 | 3 | J domain-containing protein [Azospira oryzae] |
| 05767 | 3 | Eco57I restriction-modification methylase domain-containing protein [Thauera phenylacetica] |
| 06683 | 3 | MULTISPECIES: hypothetical protein [Thauera] |
| 06682 | 3 | MULTISPECIES: restriction endonuclease [Thauera] |
| 00395 | 3 | hypothetical protein [Thauera lacus] |
| 06677 | 3 | MULTISPECIES: TIR domain-containing protein [Thauera] |
| 06675 | 3 | transposase, partial [Thauera terpenica] |
| 00393 | 3 | hypothetical protein [Denitromonas halophilus] |
| 00391 | 3 | HK97 gp10 family phage protein [Aromatoleum aromaticum] |
| 00390 | 3 | hypothetical protein [Denitromonas halophilus] |
| 06668 | 3 | GtrA family protein [Aromatoleum toluclasticum] |
| 10398 | 3 | hemagglutinin repeat-containing protein, partial [Azoarcus rhizosphaerae] |
| 10399 | 3 | toll/interleukin-1 receptor domain-containing protein [Azoarcus olearius] |
| 10401 | 3 | malonate transporter subunit MadL [Aromatoleum petrolei] |
| 10402 | 3 | glycosyltransferase family 39 protein [Azoarcus sp. DD4] |
| 05855 | 3 | DUF4440 domain-containing protein [Niveibacterium microcysteis] |
| 00337 | 3 | PEP-CTERM sorting domain-containing protein [Azoarcus communis] |
| 00333 | 3 | protein phosphatase 2C domain-containing protein [Azoarcus halotolerans] |
| 06600 | 3 | aminopeptidase P family protein [Nitrogeniibacter aestuarii] |
| 06530 | 3 | hypothetical protein [Aromatoleum toluolicum] |
| 06528 | 3 | hypothetical protein [Niveibacterium microcysteis] |
| 06526 | 3 | ImpA family metalloprotease [Niveibacterium microcysteis] |
| 06525 | 3 | hypothetical protein [Niveibacterium microcysteis] |
| 00256 | 3 | MULTISPECIES: PRTRC system protein E [Betaproteobacteria] |
| 05944 | 3 | hypothetical protein [Azoarcus rhizosphaerae] |
| 10490 | 3 | hypothetical protein [Azoarcus indigens] |
| 05947 | 3 | hypothetical protein [Aromatoleum tolulyticum] |
| 10491 | 3 | hypothetical protein [Azoarcus indigens] |
| 10492 | 3 | hypothetical protein [Azoarcus indigens] |
| 10493 | 3 | hypothetical protein [Azoarcus indigens] |
| 10494 | 3 | hypothetical protein [Zoogloea dura] |
| 05956 | 3 | MULTISPECIES: tripartite tricarboxylate transporter TctB family protein [Thauera] |
| 10495 | 3 | hypothetical protein [Azoarcus olearius] |
| 00240 | 3 | replication initiation factor [Nitrogeniibacter aestuarii] |
| 06508 | 3 | flavodoxin family protein [Uliginosibacterium gangwonense] |
| 00236 | 3 | hypothetical protein [Aromatoleum toluclasticum] |
| 06505 | 3 | hypothetical protein [Azoarcus indigens] |
| 06503 | 3 | hypothetical protein [Azoarcus indigens] |
| 00233 | 3 | hypothetical protein [Aromatoleum toluclasticum] |
| 05969 | 3 | phosphoserine transaminase [Candidatus Dactylopiibacterium carminicum] |
| 00228 | 3 | M12 family metallo-peptidase [Nitrogeniibacter aestuarii] |
| 06495 | 3 | DUF72 domain-containing protein [Aromatoleum diolicum] |

| | | |
|---|---|---|
| 05983 | 3 | NERD domain-containing protein [Zoogloea oleivorans] |
| 06493 | 3 | DDE-type integrase/transposase/recombinase, partial [Aromatoleum diolicum] |
| 06536 | 3 | glycosyltransferase [Uliginosibacterium gangwonense] |
| 10476 | 3 | TetR family transcriptional regulator [Uliginosibacterium aquaticum] |
| 06545 | 3 | hypothetical protein [Aromatoleum toluolicum] |
| 05887 | 3 | hypothetical protein [Niveibacterium microcysteis] |
| 06599 | 3 | hypothetical protein [Nitrogeniibacter aestuarii] |
| 05856 | 3 | OmcA/MtrC family decaheme c-type cytochrome [Niveibacterium microcysteis] |
| 05857 | 3 | hypothetical protein [Niveibacterium microcysteis] |
| 00318 | 3 | DUF3427 domain-containing protein [Azoarcus nasutitermitis] |
| 00317 | 3 | hypothetical protein [Thauera aromatica] |
| 00312 | 3 | plasmid mobilization relaxosome protein MobC [Denitromonas sp. IR12] |
| 06587 | 3 | hypothetical protein [Azoarcus sp. L1K30] |
| 05881 | 3 | HPF/RaiA family ribosome-associated protein [Aromatoleum bremense] |
| 06580 | 3 | MULTISPECIES: hypothetical protein [Niveibacterium] |
| 05883 | 3 | zf-TFIIB domain-containing protein [Niveibacterium microcysteis] |
| 00300 | 3 | MULTISPECIES: NAD(P)/FAD-dependent oxidoreductase [Thauera] |
| 06576 | 3 | hypothetical protein [Niveibacterium microcysteis] |
| 05911 | 3 | glycosyltransferase [Azoarcus indigens] |
| 05889 | 3 | hypothetical protein [Nitrogeniibacter aestuarii] |
| 05891 | 3 | Bax inhibitor-1 family protein [Nitrogeniibacter aestuarii] |
| 10453 | 3 | metallophosphoesterase [Aromatoleum buckelii] |
| 06570 | 3 | M48 family metalloprotease [Nitrogeniibacter aestuarii] |
| 06564 | 3 | hypothetical protein [Niveibacterium microcysteis] |
| 06563 | 3 | hypothetical protein [Niveibacterium microcysteis] |
| 06561 | 3 | hypothetical protein [Niveibacterium microcysteis] |
| 06559 | 3 | TonB family protein [Niveibacterium sp. COAC-50] |
| 06557 | 3 | tetratricopeptide repeat protein [Niveibacterium microcysteis] |
| 00284 | 3 | hypothetical protein [Uliginosibacterium aquaticum] |
| 00283 | 3 | NPP1 family protein [Uliginosibacterium gangwonense] |
| 00721 | 3 | hypothetical protein [Azoarcus sp. Aa7] |
| 00573 | 3 | hypothetical protein [Aromatoleum aromaticum] |
| 07360 | 3 | hypothetical protein [Aromatoleum toluclasticum] |
| 00784 | 3 | hypothetical protein [Aromatoleum aromaticum] |
| 00779 | 3 | AMP-binding protein, partial [Denitromonas sp. IR12] |
| 04554 | 3 | hypothetical protein [Niveibacterium microcysteis] |
| 04767 | 3 | hypothetical protein [Denitromonas halophilus] |
| 04921 | 3 | ankyrin repeat domain-containing protein [Candidatus Dactylopiibacterium carminicum] |
| 04914 | 3 | DUF481 domain-containing protein [Candidatus Dactylopiibacterium carminicum] |
| 04556 | 3 | M20/M25/M40 family metallo-hydrolase [Niveibacterium sp. COAC-50] |
| 00780 | 3 | relaxase/mobilization nuclease domain-containing protein [Denitromonas sp. IR12] |
| 00837 | 3 | curli production assembly/transport protein CsgG [Aromatoleum toluclasticum] |
| 00957 | 3 | hypothetical protein [Azoarcus olearius] |
| 07328 | 3 | ABC transporter substrate-binding protein [Niveibacterium microcysteis] |
| 04900 | 3 | alginate lyase family protein [Candidatus Dactylopiibacterium carminicum] |
| 04646 | 3 | BON domain-containing protein [Uliginosibacterium gangwonense] |
| 07238 | 3 | MULTISPECIES: type IV secretion system DNA-binding domain-containing protein [Betaproteobacteria] |
| 00916 | 3 | helix-turn-helix domain-containing protein [Zoogloea ramigera] |
| 07327 | 3 | hypothetical protein [Niveibacterium microcysteis] |
| 04940 | 3 | hypothetical protein [Niveibacterium microcysteis] |
| 07488 | 3 | hypothetical protein [Azoarcus indigens] |
| 04892 | 3 | hypothetical protein, partial [Zoogloea dura] |
| 04569 | 3 | MULTISPECIES: acyl-CoA-binding protein [Niveibacterium] |
| 07326 | 3 | hypothetical protein [Niveibacterium microcysteis] |
| 07244 | 3 | N-formylglutamate deformylase [Azoarcus olearius] |
| 07325 | 3 | FRG domain-containing protein [Niveibacterium microcysteis] |
| 04570 | 3 | hypothetical protein [Niveibacterium umoris] |
| 04781 | 3 | hypothetical protein [Azoarcus indigens] |
| 04886 | 3 | hypothetical protein [Thauera aromatica] |
| 07485 | 3 | hypothetical protein [Azoarcus indigens] |
| 04883 | 3 | hypothetical protein [Azoarcus indigens] |
| 04784 | 3 | MULTISPECIES: hypothetical protein [Proteobacteria] |
| 07318 | 3 | YkgJ family cysteine cluster protein [Aromatoleum toluolicum] |
| 09955 | 3 | hypothetical protein [Oryzomicrobium terrae] |
| 04765 | 3 | type I-E CRISPR-associated protein Cas6/Cse3/CasE [Aromatoleum diolicum] |
| 04763 | 3 | hypothetical protein [Azoarcus communis] |
| 10039 | 3 | Ig-like domain-containing protein, partial [Azoarcus halotolerans] |
| 04922 | 3 | hypothetical protein [Aromatoleum toluolicum] |
| 04506 | 3 | MBL fold metallo-hydrolase [Aromatoleum buckelii] |
| 04664 | 3 | AzlD domain-containing protein [Thauera butanivorans] |
| 04936 | 3 | DUF4150 domain-containing protein [Thauera butanivorans] |
| 04513 | 3 | non-homologous end-joining DNA ligase [Azoarcus indigens] |
| 00844 | 3 | transporter substrate-binding domain-containing protein [Thauera terpenica] |
| 04662 | 3 | MULTISPECIES: hypothetical protein [Proteobacteria] |
| 07415 | 3 | hypothetical protein [Aromatoleum toluclasticum] |
| 07513 | 3 | ABC transporter permease, partial [Thauera aminoaromatica] |
| 00903 | 3 | hypothetical protein [Azoarcus indigens] |
| 00766 | 3 | CHAT domain-containing protein [Azoarcus communis] |
| 09940 | 3 | hypothetical protein [Thauera humireducens] |
| 07220 | 3 | circadian clock KaiB family protein [Aromatoleum buckelii] |
| 10037 | 3 | nitroreductase family protein [Candidatus Dactylopiibacterium carminicum] |
| 04532 | 3 | SLATT domain-containing protein [Niveibacterium microcysteis] |
| 04755 | 3 | YbjN domain-containing protein [Thauera lacus] |
| 00904 | 3 | hypothetical protein [Propionivibrio dicarboxylicus] |
| 07341 | 3 | MULTISPECIES: hypothetical protein [Niveibacterium] |
| 10094 | 3 | GNAT family N-acetyltransferase [Denitromonas halophilus] |
| 04930 | 3 | VCBS domain-containing protein [Rhodocyclus purpureus] |
| 04535 | 3 | MULTISPECIES: autonomous glycyl radical cofactor GrcA [Niveibacterium] |
| 10092 | 3 | hypothetical protein [Aromatoleum aromaticum] |
| 04538 | 3 | hypothetical protein [Candidatus Dactylopiibacterium carminicum] |
| 04542 | 3 | high-potential iron-sulfur protein [Candidatus Dactylopiibacterium carminicum] |
| 09944 | 3 | antibiotic biosynthesis monooxygenase [Denitromonas halophilus] |
| 00772 | 3 | DUF922 domain-containing protein [Denitromonas halophilus] |
| 07340 | 3 | MULTISPECIES: cache domain-containing protein [Niveibacterium] |

| | | |
|---|---|---|
| 07500 | 3 | SEL1-like repeat protein [Azoarcus communis] |
| 07338 | 3 | OmpA family protein [Niveibacterium microcysteis] |
| 04616 | 3 | pyrimidine dimer DNA glycosylase/endonuclease V [Aromatoleum aromaticum] |
| 04877 | 3 | hypothetical protein [Azoarcus indigens] |
| 04572 | 3 | hypothetical protein [Aromatoleum bremense] |
| 10047 | 3 | class I SAM-dependent methyltransferase [Oryzomicrobium terrae] |
| 04853 | 3 | hypothetical protein, partial [Aromatoleum buckelii] |
| 07440 | 3 | DUF4260 domain-containing protein [Azoarcus indigens] |
| 04808 | 3 | hypothetical protein [Denitromonas halophilus] |
| 00927 | 3 | hypothetical protein [Azoarcus olearius] |
| 10070 | 3 | TonB C-terminal domain-containing protein [Niveibacterium microcysteis] |
| 09978 | 3 | hypothetical protein [Azoarcus communis] |
| 04601 | 3 | nucleotidyl transferase AbiEii/AbiGii toxin family protein [Nitrogeniibacter aestuarii] |
| 04809 | 3 | toll/interleukin-1 receptor domain-containing protein [Denitromonas halophilus] |
| 10068 | 3 | MULTISPECIES: hypothetical protein [Niveibacterium] |
| 09977 | 3 | DUF3024 domain-containing protein [Azoarcus communis] |
| 04606 | 3 | hypothetical protein, partial [Aromatoleum toluclasticum] |
| 07458 | 3 | hypothetical protein [Thauera lacus] |
| 00935 | 3 | prepilin-type N-terminal cleavage/methylation domain-containing protein [Denitromonas halophilus] |
| 07446 | 3 | ATP-binding protein [Aromatoleum aromaticum] |
| 07448 | 3 | TIGR04348 family glycosyltransferase [Aromatoleum buckelii] |
| 10064 | 3 | hypothetical protein [Nitrogeniibacter aestuarii] |
| 00819 | 3 | META domain-containing protein [Candidatus Dactylopiibacterium carminicum] |
| 04846 | 3 | hypothetical protein [Aromatoleum evansii] |
| 00930 | 3 | MAPEG family protein [Thauera sp. D20] |
| 04843 | 3 | DMT family transporter [Aromatoleum aromaticum] |
| 04837 | 3 | hypothetical protein [Azoarcus halotolerans] |
| 04614 | 3 | hypothetical protein [Denitromonas halophilus] |
| 04817 | 3 | oligosaccharide flippase family protein [Denitromonas halophilus] |
| 10057 | 3 | hypothetical protein [Niveibacterium microcysteis] |
| 07293 | 3 | hypothetical protein [Niveibacterium microcysteis] |
| 07288 | 3 | M14 family metallopeptidase [Niveibacterium umoris] |
| 07453 | 3 | glucose-inhibited division protein B [Azoarcus taiwanensis] |
| 00809 | 3 | hypothetical protein [Aromatoleum aromaticum] |
| 04615 | 3 | hypothetical protein [Aromatoleum aromaticum] |
| 07292 | 3 | hypothetical protein [Niveibacterium microcysteis] |
| 04630 | 3 | MULTISPECIES: hypothetical protein [Thauera] |
| 07438 | 3 | hypothetical protein [Thauera butanivorans] |
| 04574 | 3 | acyl carrier protein [Aromatoleum aromaticum] |
| 07266 | 3 | tetratricopeptide repeat-containing sulfotransferase family protein [Azoarcus indigens] |
| 09956 | 3 | phasin family protein [Zoogloea dura] |
| 10079 | 3 | ParB N-terminal domain-containing protein [Denitromonas halophilus] |
| 07430 | 3 | TIR domain-containing protein [Thauera humireducens] |
| 04791 | 3 | endopeptidase [Azoarcus rhizosphaerae] |
| 10078 | 3 | hypothetical protein [Aromatoleum bremense] |
| 07254 | 3 | hypothetical protein [Aromatoleum toluolicum] |
| 00795 | 3 | hypothetical protein [Propionivibrio dicarboxylicus] |
| 00832 | 3 | glutaredoxin family protein [Azospira oryzae] |
| 00951 | 3 | MULTISPECIES: hypothetical protein [Aromatoleum] |
| 04795 | 3 | hypothetical protein [Azoarcus communis] |
| 07474 | 3 | McrC family protein [Azospira oryzae] |
| 04593 | 3 | condensation domain-containing protein, partial [Denitromonas sp. IR12] |
| 10049 | 3 | MULTISPECIES: YunC family protein [Betaproteobacteria] |
| 07257 | 3 | TolC family protein, partial [Zoogloea oleivorans] |
| 04868 | 3 | hypothetical protein [Aromatoleum toluvorans] |
| 07260 | 3 | hypothetical protein [Azoarcus indigens] |
| 04796 | 3 | hypothetical protein [Azoarcus rhizosphaerae] |
| 04642 | 3 | DUF4914 family protein [Candidatus Dactylopiibacterium carminicum] |
| 07307 | 3 | DUF3828 domain-containing protein [Uliginosibacterium gangwonense] |
| 04797 | 3 | hypothetical protein [Aromatoleum evansii] |
| 04802 | 3 | alpha-1,2-fucosyltransferase [Azospira oryzae] |
| 00799 | 3 | sugar phosphate isomerase/epimerase [Propionivibrio dicarboxylicus] |
| 10053 | 3 | AzlD domain-containing protein [Azoarcus taiwanensis] |
| 07435 | 3 | glutaredoxin [Aromatoleum toluclasticum] |
| 04855 | 3 | hypothetical protein [Azoarcus communis] |
| 07303 | 3 | DUF488 domain-containing protein [Azoarcus communis] |
| 09967 | 3 | hypothetical protein [Nitrogeniibacter aestuarii] |
| 07264 | 3 | hypothetical protein [Azoarcus indigens] |
| 04854 | 3 | MULTISPECIES: hypothetical protein [Rhodocyclales] |
| 04939 | 3 | thioredoxin family protein [Niveibacterium microcysteis] |
| 04761 | 3 | delta(1)-pyrroline-2-carboxylate reductase family protein [Azoarcus communis] |
| 04682 | 3 | TetR family transcriptional regulator [Azoarcus olearius] |
| 10025 | 3 | MULTISPECIES: RNA-binding protein [Niveibacterium] |
| 04683 | 3 | hypothetical protein [Azoarcus olearius] |
| 04701 | 3 | class I SAM-dependent methyltransferase [Propionivibrio dicarboxylicus] |
| 00997 | 3 | hypothetical protein [Thauera butanivorans] |
| 04965 | 3 | biotin-independent malonate decarboxylase subunit gamma [Aromatoleum petrolei] |
| 04714 | 3 | type III-B CRISPR-associated protein Cas10/Cmr2 [Thauera butanivorans] |
| 04728 | 3 | baseplate protein [Azoarcus indigens] |
| 07548 | 3 | hypothetical protein [Nitrogeniibacter mangrovi] |
| 00746 | 3 | DUF1287 domain-containing protein [Nitrogeniibacter aestuarii] |
| 07519 | 3 | hypothetical protein [Denitromonas halophilus] |
| 01000 | 3 | SIR2 family protein [Denitromonas halophilus] |
| 00898 | 3 | hypothetical protein [Azoarcus indigens] |
| 00761 | 3 | GIY-YIG nuclease family protein [Aromatoleum bremense] |
| 07200 | 3 | RyR domain-containing protein [Azoarcus communis] |
| 04454 | 3 | AAA family ATPase [Nitrogeniibacter aestuarii] |
| 04956 | 3 | P-loop NTPase fold protein [Niveibacterium microcysteis] |
| 07184 | 3 | MULTISPECIES: hypothetical protein [Bacteria] |
| 04955 | 3 | hypothetical protein [Niveibacterium microcysteis] |
| 07373 | 3 | DUF5710 domain-containing protein [Aromatoleum buckelii] |
| 00760 | 3 | hypothetical protein [Aromatoleum bremense] |
| 00752 | 3 | hypothetical protein [Niveibacterium microcysteis] |
| 07180 | 3 | hypothetical protein [Aromatoleum aromaticum] |

| | | |
|---|---|---|
| 00756 | 3 | NAD-dependent epimerase/dehydratase family protein [Aromatoleum aromaticum] |
| 07530 | 3 | thioredoxin family protein [Propionivibrio dicarboxylicus] |
| 10114 | 3 | MULTISPECIES: hypothetical protein [Azospira] |
| 07193 | 3 | hypothetical protein [Azoarcus rhizosphaerae] |
| 07194 | 3 | YdhR family protein [Azoarcus communis] |
| 04478 | 3 | MULTISPECIES: hypothetical protein [Proteobacteria] |
| 10110 | 3 | energy transducer TonB [Azospira oryzae] |
| 07396 | 3 | hypothetical protein [Aromatoleum diolicum] |
| 09922 | 3 | DUF6519 domain-containing protein [Azoarcus communis] |
| 07189 | 3 | prepilin-type N-terminal cleavage/methylation domain-containing protein [Thauera lacus] |
| 07178 | 3 | hypothetical protein [Thauera propionica] |
| 00758 | 3 | hypothetical protein [Aromatoleum aromaticum] |
| 04973 | 3 | hypothetical protein [Azoarcus rhizosphaerae] |
| 04707 | 3 | lactoylglutathione lyase family protein [Denitromonas halophilus] |
| 04960 | 3 | DUF4124 domain-containing protein [Niveibacterium microcysteis] |
| 07181 | 3 | DUF2806 domain-containing protein [Aromatoleum petrolei] |
| 10030 | 3 | alpha/beta fold hydrolase [Niveibacterium sp. COAC-50] |
| 04677 | 3 | hypothetical protein [Thauera butanivorans] |
| 07357 | 3 | hypothetical protein [Aromatoleum aromaticum] |
| 04439 | 3 | hypothetical protein [Zoogloea oleivorans] |
| 04442 | 3 | endonuclease [Azoarcus sp. DD4] |
| 04950 | 3 | META domain-containing protein [Nitrogeniibacter aestuarii] |
| 04737 | 3 | ABC transporter substrate-binding protein [Azoarcus sp. KH32C] |
| 00968 | 3 | type II toxin-antitoxin system VapC family toxin [Azoarcus nasutitermitis] |
| 04700 | 3 | AAA family ATPase [Azoarcus halotolerans] |
| 07351 | 3 | DUF4115 domain-containing protein, partial [Azoarcus sp. L1K30] |
| 00901 | 3 | PepSY domain-containing protein [Azoarcus indigens] |
| 07349 | 3 | PEP-CTERM sorting domain-containing protein [Azoarcus sp. L1K30] |
| 00902 | 3 | GYF domain-containing protein [Azoarcus indigens] |
| 00861 | 3 | DNA methylase [Thauera butanivorans] |
| 09937 | 3 | hypothetical protein [Denitromonas halophilus] |
| 00966 | 3 | MULTISPECIES: hypothetical protein [Thauera] |
| 07187 | 3 | DUF4160 domain-containing protein [Aromatoleum toluvorans] |
| 00845 | 3 | MULTISPECIES: hypothetical protein [Betaproteobacteria] |
| 07372 | 3 | hypothetical protein [Aromatoleum aromaticum] |
| 04469 | 3 | prepilin peptidase [Azoarcus nasutitermitis] |
| 00753 | 3 | DUF2807 domain-containing protein [Niveibacterium microcysteis] |
| 04501 | 3 | diguanylate cyclase [Azoarcus communis] |
| 07370 | 3 | GspH/FimT family protein [Propionivibrio dicarboxylicus] |
| 10023 | 3 | J domain-containing protein [Candidatus Dactylopiibacterium carminicum] |
| 00882 | 3 | hypothetical protein [Azoarcus communis] |
| 04944 | 3 | Spy/CpxP family protein refolding chaperone [Nitrogeniibacter aestuarii] |
| 00899 | 3 | phage antirepressor KilAC domain-containing protein [Azoarcus indigens] |
| 04730 | 3 | hypothetical protein [Azoarcus indigens] |
| 10122 | 3 | hypothetical protein [Denitromonas halophilus] |
| 07205 | 3 | hypothetical protein [Nitrogeniibacter aestuarii] |
| 07408 | 3 | anaerobic ribonucleoside-triphosphate reductase activating protein [Candidatus Dactylopiibacterium carminicum] |
| 04452 | 3 | DDE-type integrase/transposase/recombinase [Aromatoleum aromaticum] |
| 04731 | 3 | hypothetical protein [Azoarcus indigens] |
| 00990 | 3 | Gfo/Idh/MocA family oxidoreductase [Candidatus Dactylopiibacterium carminicum] |
| 04942 | 3 | hypothetical protein [Niveibacterium microcysteis] |
| 10481 | 2 | hypothetical protein [Aromatoleum tolulyticum] |
| 10480 | 2 | MULTISPECIES: hypothetical protein [Betaproteobacteria] |
| 00265 | 2 | hypothetical protein [Denitromonas halophilus] |
| 09330 | 2 | hypothetical protein [Niveibacterium umoris] |
| 02007 | 2 | SAM-dependent methyltransferase [Azoarcus rhizosphaerae] |
| 02011 | 2 | FkbM family methyltransferase [Azospira oryzae] |
| 02012 | 2 | translation initiation factor IF-2 associated domain-containing protein, partial [Thauera aminoaromatica] |
| 02009 | 2 | hypothetical protein [Thauera humireducens] |
| 02005 | 2 | deoxyribodipyrimidine photo-lyase, partial [Thauera aminoaromatica] |
| 09263 | 2 | hypothetical protein [Zoogloea dura] |
| 00261 | 2 | DUF4031 domain-containing protein [Azoarcus olearius] |
| 09817 | 2 | MULTISPECIES: hypothetical protein [Thauera] |
| 00342 | 2 | zinc-binding dehydrogenase [Azoarcus communis] |
| 09983 | 2 | zinc metallopeptidase [Aromatoleum pumilum] |
| 09261 | 2 | hypothetical protein [Thauera aromatica] |
| 01999 | 2 | hypothetical protein [Aromatoleum aromaticum] |
| 01994 | 2 | cysteine protease StiP family protein [Aromatoleum aromaticum] |
| 01101 | 2 | hypothetical protein [Azoarcus olearius] |
| 00343 | 2 | pilin [Azoarcus communis] |
| 09308 | 2 | retention module-containing protein [Rhodocyclus purpureus] |
| 02013 | 2 | PsiF family protein, partial [Thauera aminoaromatica] |
| 09336 | 2 | hypothetical protein [Aromatoleum petrolei] |
| 09337 | 2 | hypothetical protein [Aromatoleum petrolei] |
| 02032 | 2 | DEAD/DEAH box helicase [Thauera phenylacetica] |
| 09812 | 2 | hypothetical protein, partial [Thauera aminoaromatica] |
| 09338 | 2 | hypothetical protein [Aromatoleum petrolei] |
| 00885 | 2 | hypothetical protein [Azoarcus communis] |
| 09975 | 2 | TolC family outer membrane protein [Azoarcus communis] |
| 09243 | 2 | DMT family transporter [Aromatoleum toluvorans] |
| 00351 | 2 | hypothetical protein, partial [Azoarcus communis] |
| 01147 | 2 | hypothetical protein [Zoogloea dura] |
| 10009 | 2 | ATP-binding protein [Oryzomicrobium terrae] |
| 01149 | 2 | substrate-binding domain-containing protein [Azoarcus halotolerans] |
| 01153 | 2 | hypothetical protein [Azoarcus halotolerans] |
| 10010 | 2 | hypothetical protein [Aromatoleum toluclasticum] |
| 10498 | 2 | hypothetical protein [Zoogloea dura] |
| 10420 | 2 | polysaccharide biosynthesis PFTS motif protein [Denitromonas halophilus] |
| 00354 | 2 | DUF4340 domain-containing protein [Nitrogeniibacter mangrovi] |
| 02040 | 2 | hypothetical protein [Azoarcus nasutitermitis] |
| 02029 | 2 | SHOCT domain-containing protein [Aromatoleum bremense] |
| 01929 | 2 | mobile mystery protein A [Azoarcus sp. Aa7] |
| 09840 | 2 | hypothetical protein [Aromatoleum petrolei] |
| 00349 | 2 | hypothetical protein [Azoarcus communis] |

| | | |
|---|---|---|
| 02014 | 2 | PD-(D/E)XK nuclease family protein, partial [Thauera aminoaromatica] |
| 01934 | 2 | restriction endonuclease subunit S [Rhodocyclus purpureus] |
| 02015 | 2 | hypothetical protein [Azoarcus sp. L1K30] |
| 01140 | 2 | helix-turn-helix domain-containing protein [Zoogloea oleivorans] |
| 00249 | 2 | hypothetical protein [Aromatoleum toluolicum] |
| 00248 | 2 | hypothetical protein [Aromatoleum tolulyticum] |
| 02016 | 2 | RsmD family RNA methyltransferase [Aromatoleum evansii] |
| 09252 | 2 | MULTISPECIES: hypothetical protein [Betaproteobacteria] |
| 10426 | 2 | hypothetical protein [Zoogloea oleivorans] |
| 00247 | 2 | hypothetical protein [Propionivibrio dicarboxylicus] |
| 00245 | 2 | glutamate mutase L [Aromatoleum toluvorans] |
| 01141 | 2 | DUF5908 family protein [Zoogloea dura] |
| 09248 | 2 | transposase, partial [Aromatoleum diolicum] |
| 00923 | 2 | hypothetical protein [Azoarcus sp. DD4] |
| 09333 | 2 | hypothetical protein [Aromatoleum bremense] |
| 09844 | 2 | hypothetical protein [Aromatoleum petrolei] |
| 09334 | 2 | addiction module protein [Aromatoleum bremense] |
| 01146 | 2 | glycosyltransferase family 4 protein [Oryzomicrobium terrae] |
| 00273 | 2 | hypothetical protein [Aromatoleum buckelii] |
| 00886 | 2 | DUF3047 domain-containing protein [Azoarcus communis] |
| 01980 | 2 | hypothetical protein [Aromatoleum toluclasticum] |
| 01975 | 2 | hypothetical protein [Nitrogeniibacter aestuarii] |
| 01979 | 2 | hypothetical protein [Nitrogeniibacter aestuarii] |
| 09293 | 2 | DUF58 domain-containing protein [Candidatus Dactylopiibacterium carminicum] |
| 01949 | 2 | hypothetical protein [Nitrogeniibacter aestuarii] |
| 10439 | 2 | DEAD/DEAH box helicase family protein [Oryzomicrobium terrae] |
| 00288 | 2 | hypothetical protein [Aromatoleum toluclasticum] |
| 01948 | 2 | sulfotransferase [Nitrogeniibacter aestuarii] |
| 01945 | 2 | potassium-transporting ATPase subunit F [Niveibacterium microcysteis] |
| 01981 | 2 | hypothetical protein [Aromatoleum toluclasticum] |
| 10456 | 2 | MULTISPECIES: hypothetical protein [Rhodocyclales] |
| 01124 | 2 | DUF2924 domain-containing protein [Aromatoleum toluolicum] |
| 09290 | 2 | hypothetical protein [Niveibacterium sp. COAC-50] |
| 00910 | 2 | hypothetical protein [Azoarcus nasutitermitis] |
| 01983 | 2 | DSD1 family PLP-dependent enzyme [Propionivibrio dicarboxylicus] |
| 09320 | 2 | hypothetical protein [Nitrogeniibacter aestuarii] |
| 01944 | 2 | hypothetical protein [Niveibacterium microcysteis] |
| 09833 | 2 | DUF2892 domain-containing protein [Propionivibrio dicarboxylicus] |
| 00325 | 2 | PilX N-terminal domain-containing pilus assembly protein [Azoarcus rhizosphaerae] |
| 01950 | 2 | hypothetical protein [Nitrogeniibacter aestuarii] |
| 09998 | 2 | hypothetical protein [Aromatoleum toluolicum] |
| 10431 | 2 | PAS domain S-box protein [Zoogloea oleivorans] |
| 01952 | 2 | hypothetical protein [Nitrogeniibacter aestuarii] |
| 00301 | 2 | hypothetical protein [Thauera humireducens] |
| 01957 | 2 | hypothetical protein [Azoarcus sp. KH32C] |
| 00303 | 2 | DUF2523 domain-containing protein [Azoarcus communis] |
| 09994 | 2 | translocation/assembly module TamB domain-containing protein, partial [Aromatoleum diolicum] |
| 09305 | 2 | hypothetical protein [Aromatoleum toluclasticum] |
| 01961 | 2 | DUF4351 domain-containing protein [Aromatoleum bremense] |
| 00308 | 2 | hypothetical protein [Thauera butanivorans] |
| 00309 | 2 | glycosyltransferase [Azoarcus olearius] |
| 01951 | 2 | DUF1444 domain-containing protein [Nitrogeniibacter aestuarii] |
| 00907 | 2 | hypothetical protein [Azoarcus nasutitermitis] |
| 01966 | 2 | class I SAM-dependent methyltransferase [Azoarcus sp. L1K30] |
| 10451 | 2 | hypothetical protein [Aromatoleum bremense] |
| 09992 | 2 | hypothetical protein [Aromatoleum diolicum] |
| 00298 | 2 | hypothetical protein [Thauera linaloolentis] |
| 01971 | 2 | hypothetical protein [Azoarcus sp. L1K30] |
| 10443 | 2 | hypothetical protein, partial [Azoarcus halotolerans] |
| 09298 | 2 | glycosyltransferase [Candidatus Dactylopiibacterium carminicum] |
| 00906 | 2 | succinate dehydrogenase [Azoarcus communis] |
| 09825 | 2 | uracil-DNA glycosylase [Azoarcus sp. KH32C] |
| 09286 | 2 | metallophosphoesterase [Azoarcus sp. KH32C] |
| 00287 | 2 | FlxA-like family protein [Propionivibrio dicarboxylicus] |
| 00271 | 2 | GTPase domain-containing protein [Niveibacterium sp. COAC-50] |
| 10469 | 2 | 50S ribosome-binding GTPase [Aromatoleum pumilum] |
| 09822 | 2 | hypothetical protein [Azoarcus sp. KH32C] |
| 10470 | 2 | DUF3800 domain-containing protein [Aromatoleum evansii] |
| 01987 | 2 | helix-turn-helix transcriptional regulator [Propionivibrio dicarboxylicus] |
| 09275 | 2 | Hpt domain-containing protein [Azoarcus sp. KH32C] |
| 00890 | 2 | hypothetical protein [Candidatus Dactylopiibacterium carminicum] |
| 10472 | 2 | hypothetical protein [Azoarcus communis] |
| 00272 | 2 | DUF4344 domain-containing metallopeptidase [Aromatoleum toluolicum] |
| 09270 | 2 | hypothetical protein [Zoogloea dura] |
| 01984 | 2 | hypothetical protein [Azospira oryzae] |
| 00269 | 2 | tRNA (5-methylaminomethyl-2-thiouridine)(34)-methyltransferase MnmD [Rhodocyclus purpureus] |
| 00889 | 2 | hypothetical protein [Thauera hydrothermalis] |
| 10433 | 2 | MULTISPECIES: hypothetical protein [Proteobacteria] |
| 09838 | 2 | hypothetical protein [Thauera propionica] |
| 01135 | 2 | hypothetical protein [Zoogloea dura] |
| 09269 | 2 | hypothetical protein [Zoogloea dura] |
| 10432 | 2 | baseplate J/gp47 family protein [Zoogloea dura] |
| 10478 | 2 | hypothetical protein [Aromatoleum diolicum] |
| 01103 | 2 | XapX domain-containing protein [Azoarcus rhizosphaerae] |
| 00275 | 2 | sulfotransferase [Azoarcus sp. KH32C] |
| 00276 | 2 | phosphatase PAP2 family protein [Aromatoleum toluclasticum] |
| 09823 | 2 | hypothetical protein [Azoarcus sp. KH32C] |
| 00895 | 2 | hypothetical protein [Azospira oryzae] |
| 00329 | 2 | site-specific integrase [Thauera aromatica] |
| 00285 | 2 | hypothetical protein [Azospira oryzae] |
| 09282 | 2 | hypothetical protein [Niveibacterium microcysteis] |
| 01125 | 2 | hypothetical protein [Azoarcus rhizosphaerae] |
| 00917 | 2 | glycosyltransferase family protein [Azoarcus rhizosphaerae] |
| 09280 | 2 | hypothetical protein [Denitromonas sp. IR12] |

| | | |
|---|---|---|
| 10460 | 2 | efflux RND transporter periplasmic adaptor subunit, partial [Aromatoleum evansii] |
| 09279 | 2 | hypothetical protein [Uliginosibacterium aquaticum] |
| 00918 | 2 | hypothetical protein, partial [Azoarcus rhizosphaerae] |
| 01985 | 2 | hypothetical protein [Propionivibrio dicarboxylicus] |
| 10462 | 2 | hypothetical protein [Azoarcus communis] |
| 10463 | 2 | hypothetical protein [Azoarcus communis] |
| 09835 | 2 | N-acetylglucosamine-6-phosphate deacetylase [Niveibacterium umoris] |
| 00281 | 2 | cellulase family glycosylhydrolase [Uliginosibacterium aquaticum] |
| 09984 | 2 | hypothetical protein [Aromatoleum pumilum] |
| 00894 | 2 | hypothetical protein [Azospira oryzae] |
| 02041 | 2 | fibronectin type III domain-containing protein [Thauera humireducens] |
| 02076 | 2 | hypothetical protein [Azoarcus communis] |
| 09847 | 2 | transposase, partial [Thauera terpenica] |
| 00056 | 2 | hypothetical protein [Azoarcus nasutitermitis] |
| 09898 | 2 | hypothetical protein [Azoarcus communis] |
| 00066 | 2 | DDE-type integrase/transposase/recombinase [Aromatoleum toluclasticum] |
| 00064 | 2 | DUF1442 domain-containing protein [Azoarcus rhizosphaerae] |
| 00062 | 2 | hypothetical protein [Azoarcus rhizosphaerae] |
| 00061 | 2 | helix-turn-helix domain-containing protein [Azoarcus rhizosphaerae] |
| 09134 | 2 | terminase small subunit [Uliginosibacterium gangwonense] |
| 02215 | 2 | hypothetical protein [Aromatoleum toluclasticum] |
| 10582 | 2 | hypothetical protein [Aromatoleum petrolei] |
| 10583 | 2 | SDR family NAD(P)-dependent oxidoreductase [Aromatoleum petrolei] |
| 00059 | 2 | insulinase family protein [Azoarcus nasutitermitis] |
| 09130 | 2 | hypothetical protein [Azoarcus olearius] |
| 02218 | 2 | transporter [Uliginosibacterium aquaticum] |
| 01024 | 2 | hypothetical protein [Thauera butanivorans] |
| 09901 | 2 | glycosyltransferase family 4 protein [Aromatoleum pumilum] |
| 09127 | 2 | hypothetical protein [Azoarcus rhizosphaerae] |
| 02221 | 2 | lipoprotein [Candidatus Dactylopiibacterium carminicum] |
| 02223 | 2 | hypothetical protein [Nitrogeniibacter aestuarii] |
| 10586 | 2 | hypothetical protein [Azoarcus communis] |
| 09124 | 2 | Flp family type IVb pilin [Azoarcus rhizosphaerae] |
| 09123 | 2 | RelA/SpoT domain-containing protein [Azoarcus rhizosphaerae] |
| 01023 | 2 | DUF11 domain-containing protein [Aromatoleum pumilum] |
| 09121 | 2 | HPr-rel-A system PqqD family peptide chaperone [Azoarcus nasutitermitis] |
| 10589 | 2 | RHS repeat-associated core domain-containing protein, partial [Azoarcus sp. L1K30] |
| 09934 | 2 | hypothetical protein [Zoogloea ramigera] |
| 02225 | 2 | C39 family peptidase [Niveibacterium sp. COAC-50] |
| 02227 | 2 | hypothetical protein [Niveibacterium microcysteis] |
| 10577 | 2 | DUF4433 domain-containing protein [Aromatoleum bremense] |
| 09897 | 2 | hypothetical protein, partial [Denitromonas sp. IR12] |
| 01025 | 2 | hypothetical protein [Azoarcus communis] |
| 02200 | 2 | reverse transcriptase domain-containing protein [Aromatoleum toluolicum] |
| 09163 | 2 | DUF945 family protein [Azoarcus communis] |
| 00105 | 2 | hypothetical protein [Aromatoleum aromaticum] |
| 01045 | 2 | hypothetical protein [Thauera humireducens] |
| 01044 | 2 | hypothetical protein [Aromatoleum aromaticum] |
| 09945 | 2 | filamentous hemagglutinin N-terminal domain-containing protein [Zoogloea dura] |
| 02185 | 2 | DUF637 domain-containing protein, partial [Azoarcus indigens] |
| 10563 | 2 | hypothetical protein [Nitrogeniibacter aestuarii] |
| 02190 | 2 | GMC oxidoreductase [Azoarcus taiwanensis] |
| 09155 | 2 | DUF99 family protein [Aromatoleum diolicum] |
| 09154 | 2 | hypothetical protein [Azoarcus sp. DD4] |
| 01030 | 2 | sel1 repeat family protein [Azoarcus olearius] |
| 01029 | 2 | hypothetical protein [Azoarcus olearius] |
| 01028 | 2 | MULTISPECIES: PD-(D/E)XK nuclease domain-containing protein [Thauera] |
| 09151 | 2 | hypothetical protein [Azoarcus communis] |
| 10565 | 2 | hypothetical protein [Nitrogeniibacter aestuarii] |
| 01027 | 2 | hypothetical protein, partial [Thauera butanivorans] |
| 09149 | 2 | hypothetical protein [Azoarcus nasutitermitis] |
| 09148 | 2 | hypothetical protein [Azoarcus nasutitermitis] |
| 00093 | 2 | sulfite exporter TauE/SafE family protein [Azospira oryzae] |
| 09146 | 2 | hypothetical protein [Uliginosibacterium gangwonense] |
| 02196 | 2 | MULTISPECIES: hypothetical protein [Betaproteobacteria] |
| 00965 | 2 | GNAT family N-acetyltransferase [Thauera butanivorans] |
| 02198 | 2 | hypothetical protein [Aromatoleum diolicum] |
| 09142 | 2 | site-specific DNA-methyltransferase [Aromatoleum buckelii] |
| 00089 | 2 | hypothetical protein [Zoogloea dura] |
| 10591 | 2 | hypothetical protein [Azoarcus sp. L1K30] |
| 09116 | 2 | hypothetical protein [Aromatoleum toluvorans] |
| 10499 | 2 | hypothetical protein [Aromatoleum bremense] |
| 00022 | 2 | GNAT family N-acetyltransferase [Thauera lacus] |
| 09095 | 2 | DUF45 domain-containing protein [Azoarcus communis] |
| 10607 | 2 | hypothetical protein [Aromatoleum toluclasticum] |
| 02265 | 2 | hypothetical protein [Azoarcus halotolerans] |
| 01010 | 2 | MULTISPECIES: tetracycline resistance transcriptional repressor TetR(A) [Bacteria] |
| 09921 | 2 | hypothetical protein [Aromatoleum toluolicum] |
| 02267 | 2 | hypothetical protein [Azospira oryzae] |
| 02269 | 2 | type II secretion system protein GspK [Niveibacterium microcysteis] |
| 02270 | 2 | M48 family metallopeptidase [Nitrogeniibacter mangrovi] |
| 00029 | 2 | hypothetical protein [Azoarcus communis] |
| 00028 | 2 | ATP-binding cassette domain-containing protein, partial [Azoarcus communis] |
| 00026 | 2 | hypothetical protein [Aromatoleum toluolicum] |
| 10613 | 2 | ABC transporter permease subunit, partial [Aromatoleum toluclasticum] |
| 02286 | 2 | hypothetical protein [Azoarcus sp. L1K30] |
| 00973 | 2 | DUF6314 family protein [Thauera phenylacetica] |
| 10615 | 2 | pyridoxamine 5'-phosphate oxidase family protein [Azoarcus nasutitermitis] |
| 09082 | 2 | hypothetical protein [Thauera aromatica] |
| 09918 | 2 | DUF2528 family protein [Azoarcus rhizosphaerae] |
| 09080 | 2 | hypothetical protein [Rhodocyclus purpureus] |
| 10618 | 2 | YbaK/prolyl-tRNA synthetase associated domain-containing protein [Propionivibrio dicarboxylicus] |
| 00015 | 2 | DUF2459 domain-containing protein [Aromatoleum pumilum] |
| 10620 | 2 | L-ribulose-5-phosphate 3-epimerase [Propionivibrio dicarboxylicus] |

| | | |
|---|---|---|
| 01004 | 2 | hypothetical protein [Thauera butanivorans] |
| 09916 | 2 | hypothetical protein [Azoarcus nasutitermitis] |
| 10623 | 2 | glycosyltransferase [Azospira oryzae] |
| 00005 | 2 | MULTISPECIES: hypothetical protein [Proteobacteria] |
| 01008 | 2 | hypothetical protein [Thauera butanivorans] |
| 02263 | 2 | hypothetical protein [Niveibacterium umoris] |
| 10606 | 2 | hypothetical protein [Nitrogeniibacter aestuarii] |
| 10605 | 2 | hypothetical protein [Nitrogeniibacter aestuarii] |
| 00039 | 2 | hypothetical protein [Azospira oryzae] |
| 00974 | 2 | MULTISPECIES: hypothetical protein [Thauera] |
| 01022 | 2 | hypothetical protein [Thauera butanivorans] |
| 10595 | 2 | hypothetical protein [Denitromonas sp. IR12] |
| 00975 | 2 | hypothetical protein [Thauera aminoaromatica] |
| 02232 | 2 | hypothetical protein [Azoarcus sp. DD4] |
| 02234 | 2 | hypothetical protein [Candidatus Dactylopiibacterium carminicum] |
| 09111 | 2 | TonB family protein, partial [Azoarcus sp. TTM-91] |
| 02236 | 2 | MULTISPECIES: hypothetical protein [Uliginosibacterium] |
| 00977 | 2 | glycosyltransferase family 4 protein [Azoarcus sp. KH32C] |
| 01020 | 2 | EthD family reductase [Azoarcus sp. DD4] |
| 00980 | 2 | hypothetical protein [Azoarcus sp. KH32C] |
| 02238 | 2 | hypothetical protein [Azoarcus communis] |
| 02247 | 2 | DUF2281 domain-containing protein [Rhodocyclus purpureus] |
| 00043 | 2 | alpha/beta fold hydrolase [Azospira oryzae] |
| 01017 | 2 | hypothetical protein [Thauera humireducens] |
| 02250 | 2 | alpha/beta hydrolase [Niveibacterium sp. COAC-50] |
| 10600 | 2 | hypothetical protein [Nitrogeniibacter aestuarii] |
| 02255 | 2 | hypothetical protein [Nitrogeniibacter aestuarii] |
| 10601 | 2 | sulfotransferase [Nitrogeniibacter aestuarii] |
| 02257 | 2 | hypothetical protein [Azoarcus halotolerans] |
| 10602 | 2 | cache domain-containing protein [Nitrogeniibacter aestuarii] |
| 00992 | 2 | metal-dependent hydrolase [Aromatoleum tolulyticum] |
| 00994 | 2 | hypothetical protein [Uliginosibacterium gangwonense] |
| 10603 | 2 | ATP-binding cassette domain-containing protein [Nitrogeniibacter aestuarii] |
| 00996 | 2 | hypothetical protein [Aromatoleum toluclasticum] |
| 09885 | 2 | hypothetical protein [Thauera phenolivorans] |
| 00111 | 2 | hypothetical protein [Aromatoleum evansii] |
| 02182 | 2 | helix-turn-helix domain-containing protein, partial [Azoarcus indigens] |
| 00193 | 2 | hypothetical protein [Niveibacterium microcysteis] |
| 02066 | 2 | hypothetical protein [Thauera aromatica] |
| 02071 | 2 | SOS response-associated peptidase [Thauera lacus] |
| 01089 | 2 | hypothetical protein [Azoarcus nasutitermitis] |
| 09221 | 2 | hypothetical protein [Aromatoleum bremense] |
| 09220 | 2 | hypothetical protein [Aromatoleum evansii] |
| 00940 | 2 | transposase, partial [Azoarcus halotolerans] |
| 09341 | 2 | hypothetical protein [Azoarcus communis] |
| 09859 | 2 | sulfotransferase [Azoarcus sp. L1K30] |
| 09218 | 2 | hypothetical protein [Aromatoleum evansii] |
| 01087 | 2 | hypothetical protein [Azoarcus nasutitermitis] |
| 00198 | 2 | hypothetical protein [Nitrogeniibacter aestuarii] |
| 00194 | 2 | hypothetical protein [Nitrogeniibacter aestuarii] |
| 09214 | 2 | hypothetical protein [Uliginosibacterium gangwonense] |
| 02180 | 2 | Mur ligase family protein [Azoarcus communis] |
| 09213 | 2 | PRTRC system protein D, partial [Aromatoleum buckelii] |
| 10524 | 2 | hypothetical protein [Candidatus Dactylopiibacterium carminicum] |
| 01086 | 2 | L-2-amino-thiazoline-4-carboxylic acid hydrolase [Azoarcus nasutitermitis] |
| 00192 | 2 | hypothetical protein [Nitrogeniibacter mangrovi] |
| 01085 | 2 | hypothetical protein [Azoarcus nasutitermitis] |
| 10527 | 2 | hypothetical protein [Candidatus Dactylopiibacterium carminicum] |
| 00941 | 2 | hypothetical protein [Aromatoleum evansii] |
| 00944 | 2 | hypothetical protein, partial [Aromatoleum tolulyticum] |
| 01084 | 2 | phosphopantetheine-binding protein [Azoarcus indigens] |
| 00190 | 2 | GNAT family N-acetyltransferase [Niveibacterium microcysteis] |
| 00945 | 2 | hypothetical protein [Aromatoleum tolulyticum] |
| 09864 | 2 | helix-turn-helix domain-containing protein [Thauera butanivorans] |
| 09857 | 2 | hypothetical protein [Thauera humireducens] |
| 09225 | 2 | DUF3077 domain-containing protein [Azoarcus communis] |
| 02065 | 2 | hypothetical protein [Thauera aromatica] |
| 02061 | 2 | hypothetical protein [Aromatoleum pumilum] |
| 10500 | 2 | class I SAM-dependent methyltransferase [Thauera hydrothermalis] |
| 09849 | 2 | hypothetical protein [Rhodocyclus purpureus] |
| 10503 | 2 | hypothetical protein [Candidatus Dactylopiibacterium carminicum] |
| 09973 | 2 | hypothetical protein [Aromatoleum evansii] |
| 01093 | 2 | PTS sugar transporter subunit IIA [Azoarcus nasutitermitis] |
| 02047 | 2 | hypothetical protein [Thauera butanivorans] |
| 00226 | 2 | hypothetical protein [Azoarcus sp. Aa7] |
| 00223 | 2 | NYN domain-containing protein [Denitromonas sp. IR12] |
| 09852 | 2 | hypothetical protein [Azoarcus halotolerans] |
| 02050 | 2 | DUF2357 domain-containing protein [Azoarcus olearius] |
| 10507 | 2 | hypothetical protein [Uliginosibacterium aquaticum] |
| 00222 | 2 | helix-turn-helix transcriptional regulator [Azospira oryzae] |
| 00937 | 2 | hypothetical protein [Denitromonas halophilus] |
| 09854 | 2 | DUF2220 family protein [Azoarcus communis] |
| 02052 | 2 | hypothetical protein [Azoarcus olearius] |
| 09233 | 2 | hypothetical protein, partial [Azoarcus indigens] |
| 00211 | 2 | AAA family ATPase [Aromatoleum diolicum] |
| 09969 | 2 | c-type cytochrome [Aromatoleum aromaticum] |
| 10513 | 2 | M4 family metallopeptidase [Niveibacterium microcysteis] |
| 09232 | 2 | VENN motif pre-toxin domain-containing protein, partial [Azoarcus communis] |
| 09231 | 2 | hypothetical protein [Azoarcus communis] |
| 10514 | 2 | periplasmic heavy metal sensor [Niveibacterium microcysteis] |
| 00939 | 2 | hypothetical protein [Denitromonas halophilus] |
| 02053 | 2 | hypothetical protein [Azoarcus olearius] |
| 02054 | 2 | hypothetical protein [Azoarcus olearius] |
| 00946 | 2 | bacteriophage abortive infection AbiH family protein [Aromatoleum aromaticum] |

| | | |
|---|---|---|
| 10530 | 2 | DUF3883 domain-containing protein [Azoarcus sp. Aa7] |
| 01083 | 2 | Arc family DNA-binding protein [Azoarcus indigens] |
| 02158 | 2 | hypothetical protein [Azoarcus halotolerans] |
| 09183 | 2 | transposase [Thauera lacus] |
| 09182 | 2 | ParB/RepB/Spo0J family partition protein [Candidatus Dactylopiibacterium carminicum] |
| 09954 | 2 | hypothetical protein [Zoogloea oleivorans] |
| 01067 | 2 | oligosaccharide flippase family protein [Azoarcus communis] |
| 09953 | 2 | hypothetical protein [Azoarcus taiwanensis] |
| 09179 | 2 | metallo-mystery pair system four-Cys motif protein [Zoogloea dura] |
| 10546 | 2 | hypothetical protein [Niveibacterium microcysteis] |
| 09178 | 2 | ATP-binding domain-containing protein [Denitromonas halophilus] |
| 02170 | 2 | hypothetical protein [Aromatoleum aromaticum] |
| 09176 | 2 | transposase [Aromatoleum toluolicum] |
| 09175 | 2 | GDP-mannose 4,6-dehydratase, partial [Zoogloea dura] |
| 10547 | 2 | hypothetical protein [Niveibacterium microcysteis] |
| 10548 | 2 | chorismate mutase [Niveibacterium microcysteis] |
| 00954 | 2 | hypothetical protein [Aromatoleum aromaticum] |
| 00955 | 2 | hypothetical protein [Aromatoleum aromaticum] |
| 00956 | 2 | hypothetical protein [Azoarcus olearius] |
| 09948 | 2 | hypothetical protein [Zoogloea oleivorans] |
| 00125 | 2 | ABC transporter ATP-binding protein, partial [Azoarcus taiwanensis] |
| 09947 | 2 | hypothetical protein [Zoogloea oleivorans] |
| 02173 | 2 | hypothetical protein [Aromatoleum bremense] |
| 09881 | 2 | hypothetical protein [Thauera chlorobenzoica] |
| 02177 | 2 | GNAT family N-acetyltransferase [Azoarcus communis] |
| 09946 | 2 | hypothetical protein [Thauera butanivorans] |
| 01052 | 2 | glycosyltransferase [Rhodocyclus purpureus] |
| 02179 | 2 | hypothetical protein [Aromatoleum diolicum] |
| 09184 | 2 | hypothetical protein [Thauera linaloolentis] |
| 01068 | 2 | hypothetical protein, partial [Azoarcus communis] |
| 00950 | 2 | winged helix-turn-helix transcriptional regulator [Aromatoleum aromaticum] |
| 02146 | 2 | nuclear transport factor 2 family protein [Aromatoleum toluclasticum] |
| 00178 | 2 | ATP-binding protein [Azoarcus sp. Aa7] |
| 00175 | 2 | ribonuclease T2 [Rhodocyclus purpureus] |
| 09204 | 2 | hypothetical protein [Azoarcus halotolerans] |
| 02105 | 2 | hypothetical protein [Aromatoleum pumilum] |
| 09958 | 2 | hypothetical protein, partial [Thauera phenylacetica] |
| 00173 | 2 | DUF1441 family protein [Azoarcus sp. Aa7] |
| 00167 | 2 | O-antigen ligase family protein [Candidatus Dactylopiibacterium carminicum] |
| 02117 | 2 | hypothetical protein, partial [Zoogloea ramigera] |
| 09872 | 2 | hypothetical protein [Azoarcus olearius] |
| 09196 | 2 | MULTISPECIES: hypothetical protein [Thauera] |
| 00160 | 2 | hypothetical protein [Niveibacterium microcysteis] |
| 09873 | 2 | hypothetical protein [Aromatoleum petrolei] |
| 09194 | 2 | MULTISPECIES: hypothetical protein [Zoogloeaceae] |
| 10540 | 2 | putative photosynthetic complex assembly protein PuhC [Rhodocyclus purpureus] |
| 10541 | 2 | DUF1851 domain-containing protein [Niveibacterium microcysteis] |
| 02126 | 2 | hypothetical protein [Denitromonas halophilus] |
| 10542 | 2 | hypothetical protein [Niveibacterium microcysteis] |
| 02128 | 2 | hypothetical protein [Azoarcus nasutitermitis] |
| 09191 | 2 | patatin-like phospholipase family protein [Oryzomicrobium terrae] |
| 01073 | 2 | hypothetical protein [Azospira oryzae] |
| 02131 | 2 | HNH endonuclease [Aromatoleum pumilum] |
| 02136 | 2 | DUF2868 domain-containing protein [Zoogloea dura] |
| 09875 | 2 | hypothetical protein [Denitromonas sp. IR12] |
| 02138 | 2 | hypothetical protein [Aromatoleum evansii] |
| 02143 | 2 | Fic family protein [Thauera aromatica] |
| 00355 | 2 | cellulose synthase complex periplasmic endoglucanase BcsZ [Aromatoleum bremense] |
| 00737 | 2 | hypothetical protein [Niveibacterium microcysteis] |
| 10011 | 2 | NAD-binding protein [Propionivibrio dicarboxylicus] |
| 00634 | 2 | HEPN domain-containing protein [Aromatoleum buckelii] |
| 01560 | 2 | hypothetical protein, partial [Denitromonas sp. IR12] |
| 09565 | 2 | ABC transporter substrate-binding protein [Aromatoleum toluolicum] |
| 09718 | 2 | hypothetical protein [Candidatus Dactylopiibacterium carminicum] |
| 00630 | 2 | ThiF family adenylyltransferase [Aromatoleum toluvorans] |
| 09719 | 2 | hypothetical protein [Candidatus Dactylopiibacterium carminicum] |
| 09721 | 2 | DUF5680 domain-containing protein [Candidatus Dactylopiibacterium carminicum] |
| 00615 | 2 | hypothetical protein [Oryzomicrobium terrae] |
| 01567 | 2 | hypothetical protein [Aromatoleum toluclasticum] |
| 10230 | 2 | GNAT family N-acetyltransferase [Aromatoleum bremense] |
| 00609 | 2 | hypothetical protein [Aromatoleum pumilum] |
| 01286 | 2 | hypothetical protein [Azoarcus nasutitermitis] |
| 10086 | 2 | hypothetical protein [Azoarcus sp. L1K30] |
| 10234 | 2 | HNH endonuclease [Azoarcus sp. L1K30] |
| 10235 | 2 | hypothetical protein [Azoarcus sp. L1K30] |
| 01283 | 2 | hypothetical protein [Nitrogeniibacter aestuarii] |
| 00606 | 2 | glycosyltransferase [Denitromonas halophilus] |
| 01576 | 2 | acyclic terpene utilization AtuA family protein [Aromatoleum toluvorans] |
| 09558 | 2 | AAA family ATPase [Aromatoleum bremense] |
| 00781 | 2 | PEP-CTERM sorting domain-containing protein [Nitrogeniibacter aestuarii] |
| 01579 | 2 | OadG family protein [Propionivibrio dicarboxylicus] |
| 00785 | 2 | CopK family periplasmic copper-binding protein [Aromatoleum toluclasticum] |
| 10240 | 2 | cupin domain-containing protein [Denitromonas halophilus] |
| 01585 | 2 | PIN domain nuclease [Azoarcus sp. Aa7] |
| 10241 | 2 | hypothetical protein [Azoarcus taiwanensis] |
| 01281 | 2 | hypothetical protein [Nitrogeniibacter aestuarii] |
| 00633 | 2 | transposase, partial [Aromatoleum buckelii] |
| 10089 | 2 | hypothetical protein [Azoarcus sp. L1K30] |
| 09729 | 2 | hypothetical protein [Rhodocyclus purpureus] |
| 10218 | 2 | methyltransferase domain-containing protein [Nitrogeniibacter aestuarii] |
| 01525 | 2 | MerR family transcriptional regulator [Niveibacterium umoris] |
| 01301 | 2 | NAD-dependent epimerase/dehydratase family protein [Aromatoleum petrolei] |
| 10095 | 2 | hypothetical protein [Denitromonas sp. IR12] |
| 09583 | 2 | DUF3261 domain-containing protein [Niveibacterium microcysteis] |

| | | |
|---|---|---|
| 09582 | 2 | hypothetical protein [Niveibacterium sp. COAC-50] |
| 00644 | 2 | hypothetical protein [Aromatoleum aromaticum] |
| 09710 | 2 | hypothetical protein [Niveibacterium sp. COAC-50] |
| 09580 | 2 | hypothetical protein [Uliginosibacterium aquaticum] |
| 00643 | 2 | ATP-binding protein [Aromatoleum diolicum] |
| 10205 | 2 | T6SS immunity protein Tli4 family protein [Niveibacterium microcysteis] |
| 10206 | 2 | hypothetical protein [Niveibacterium microcysteis] |
| 10208 | 2 | DUF2946 family protein [Niveibacterium microcysteis] |
| 00769 | 2 | hypothetical protein [Azoarcus communis] |
| 01537 | 2 | hypothetical protein [Aromatoleum bremense] |
| 10209 | 2 | hypothetical protein [Aromatoleum tolulyticum] |
| 00770 | 2 | hypothetical protein [Azoarcus rhizosphaerae] |
| 10210 | 2 | hypothetical protein [Niveibacterium microcysteis] |
| 09714 | 2 | hypothetical protein [Denitromonas halophilus] |
| 01295 | 2 | hypothetical protein [Niveibacterium sp. COAC-50] |
| 00640 | 2 | hypothetical protein [Aromatoleum evansii] |
| 09572 | 2 | SIR2 family protein [Aromatoleum buckelii] |
| 00638 | 2 | transposase [Candidatus Dactylopiibacterium carminicum] |
| 01546 | 2 | LysE family transporter [Azoarcus communis] |
| 00637 | 2 | MULTISPECIES: hypothetical protein [Proteobacteria] |
| 10217 | 2 | hypothetical protein [Nitrogeniibacter aestuarii] |
| 01280 | 2 | hypothetical protein [Nitrogeniibacter aestuarii] |
| 01587 | 2 | MULTISPECIES: nucleoid-associated protein [Proteobacteria] |
| 01523 | 2 | DUF3421 domain-containing protein [Nitrogeniibacter aestuarii] |
| 00794 | 2 | hypothetical protein [Propionivibrio dicarboxylicus] |
| 10254 | 2 | hypothetical protein [Aromatoleum toluclasticum] |
| 01621 | 2 | ABC transporter transmembrane domain-containing protein [Uliginosibacterium gangwonense] |
| 10255 | 2 | nucleotidyl transferase AbiEii/AbiGii toxin family protein [Aromatoleum toluclasticum] |
| 01622 | 2 | hypothetical protein [Uliginosibacterium aquaticum] |
| 01626 | 2 | endonuclease NucS [Aromatoleum toluclasticum] |
| 09531 | 2 | hypothetical protein [Azoarcus halotolerans] |
| 01629 | 2 | hypothetical protein [Aromatoleum toluclasticum] |
| 10259 | 2 | hypothetical protein [Propionivibrio dicarboxylicus] |
| 09740 | 2 | TonB-dependent receptor plug domain-containing protein [Niveibacterium microcysteis] |
| 01633 | 2 | glycosyltransferase family 4 protein [Thauera aminoaromatica] |
| 09527 | 2 | hypothetical protein [Azoarcus sp. KH32C] |
| 01638 | 2 | MULTISPECIES: hypothetical protein [Thauera] |
| 10261 | 2 | [FeFe] hydrogenase H-cluster maturation GTPase HydF [Propionivibrio dicarboxylicus] |
| 01642 | 2 | DUF4124 domain-containing protein [Azoarcus communis] |
| 01644 | 2 | hypothetical protein [Azoarcus sp. DD4] |
| 10263 | 2 | DUF4392 domain-containing protein [Azoarcus sp. TTM-91] |
| 01645 | 2 | hypothetical protein [Azoarcus communis] |
| 01646 | 2 | hypothetical protein [Azoarcus sp. Aa7] |
| 09519 | 2 | MULTISPECIES: hypothetical protein [Rhodocyclales] |
| 01270 | 2 | protoglobin domain-containing protein [Niveibacterium microcysteis] |
| 00581 | 2 | hypothetical protein [Azoarcus rhizosphaerae] |
| 10268 | 2 | hypothetical protein [Propionivibrio dicarboxylicus] |
| 01656 | 2 | sulfotransferase [Azoarcus communis] |
| 01658 | 2 | ArsR family transcriptional regulator [Aromatoleum aromaticum] |
| 10270 | 2 | TIM barrel protein [Propionivibrio dicarboxylicus] |
| 09536 | 2 | GNAT family N-acetyltransferase [Nitrogeniibacter mangrovi] |
| 10253 | 2 | hypothetical protein [Aromatoleum toluclasticum] |
| 09730 | 2 | transcription/translation regulatory transformer protein RfaH [Propionivibrio dicarboxylicus] |
| 01615 | 2 | hypothetical protein [Azoarcus sp. TTM-91] |
| 01588 | 2 | CAP domain-containing protein [Aromatoleum aromaticum] |
| 09731 | 2 | hypothetical protein [Azoarcus nasutitermitis] |
| 10243 | 2 | transposase domain-containing protein, partial [Denitromonas sp. IR12] |
| 09732 | 2 | pentapeptide repeat-containing protein [Rhodocyclus purpureus] |
| 10244 | 2 | hypothetical protein [Aromatoleum pumilum] |
| 00788 | 2 | hypothetical protein [Denitromonas sp. IR12] |
| 09551 | 2 | hypothetical protein [Azoarcus communis] |
| 10245 | 2 | hypothetical protein [Denitromonas sp. IR12] |
| 01273 | 2 | hypothetical protein [Niveibacterium microcysteis] |
| 00792 | 2 | DEAD/DEAH box helicase [Propionivibrio dicarboxylicus] |
| 10247 | 2 | transposase, partial [Candidatus Dactylopiibacterium carminicum] |
| 09549 | 2 | hypothetical protein [Azoarcus communis] |
| 01611 | 2 | lipopolysaccharide biosynthesis protein [Azoarcus sp. KH32C] |
| 10248 | 2 | hypothetical protein [Azoarcus communis] |
| 10249 | 2 | hypothetical protein [Nitrogeniibacter aestuarii] |
| 09547 | 2 | efflux RND transporter permease subunit, partial [Aromatoleum evansii] |
| 10250 | 2 | hypothetical protein [Nitrogeniibacter aestuarii] |
| 10251 | 2 | hypothetical protein [Aromatoleum toluclasticum] |
| 09544 | 2 | hypothetical protein [Aromatoleum evansii] |
| 01613 | 2 | hypothetical protein [Niveibacterium umoris] |
| 10252 | 2 | class I SAM-dependent DNA methyltransferase [Azoarcus toluclasticus] |
| 09542 | 2 | hypothetical protein [Aromatoleum evansii] |
| 09541 | 2 | hypothetical protein [Aromatoleum buckelii] |
| 01271 | 2 | PEP-CTERM sorting domain-containing protein [Niveibacterium microcysteis] |
| 01614 | 2 | hypothetical protein [Uliginosibacterium aquaticum] |
| 10098 | 2 | hypothetical protein [Azoarcus olearius] |
| 01522 | 2 | tryptophan-rich sensory protein [Nitrogeniibacter aestuarii] |
| 01923 | 2 | hypothetical protein [Azoarcus sp. Aa7] |
| 01370 | 2 | hypothetical protein [Denitromonas halophilus] |
| 00748 | 2 | hypothetical protein [Nitrogeniibacter aestuarii] |
| 10143 | 2 | NUDIX domain-containing protein [Aromatoleum toluolicum] |
| 01439 | 2 | Nif3-like dinuclear metal center hexameric protein, partial [Aromatoleum diolicum] |
| 00750 | 2 | DUF1206 domain-containing protein [Nitrogeniibacter aestuarii] |
| 09676 | 2 | MULTISPECIES: hypothetical protein [Proteobacteria] |
| 09677 | 2 | arginase family protein, partial [Azoarcus indigens] |
| 09644 | 2 | FumA C-terminus/TtdB family hydratase beta subunit, partial [Aromatoleum evansii] |
| 10145 | 2 | RidA family protein [Azoarcus sp. TTM-91] |
| 01354 | 2 | hypothetical protein [Azoarcus olearius] |
| 09643 | 2 | diguanylate cyclase [Zoogloea dura] |
| 10112 | 2 | inovirus-type Gp2 protein [Niveibacterium microcysteis] |

| | | |
|---|---|---|
| 10152 | 2 | protease inhibitor I42 family protein [Zoogloea dura] |
| 00695 | 2 | tetratricopeptide repeat protein [Uliginosibacterium aquaticum] |
| 10154 | 2 | hypothetical protein [Zoogloea dura] |
| 09636 | 2 | MULTISPECIES: hypothetical protein [Aromatoleum] |
| 01455 | 2 | hypothetical protein, partial [Azoarcus indigens] |
| 09681 | 2 | methylaspartate mutase subunit E [Aromatoleum toluvorans] |
| 01457 | 2 | hypothetical protein, partial [Azoarcus sp. TTM-91] |
| 00692 | 2 | hypothetical protein [Uliginosibacterium aquaticum] |
| 00687 | 2 | class I SAM-dependent methyltransferase [Azoarcus sp. TTM-91] |
| 09631 | 2 | hypothetical protein [Niveibacterium sp. COAC-50] |
| 09630 | 2 | hypothetical protein [Nitrogeniibacter aestuarii] |
| 09629 | 2 | CDP-glycerol glycerophosphotransferase family protein [Nitrogeniibacter aestuarii] |
| 10158 | 2 | hypothetical protein [Uliginosibacterium aquaticum] |
| 00686 | 2 | hypothetical protein [Zoogloea dura] |
| 01431 | 2 | hypothetical protein [Azoarcus communis] |
| 10118 | 2 | chemotaxis protein CheX [Propionivibrio dicarboxylicus] |
| 09682 | 2 | DUF4242 domain-containing protein [Aromatoleum diolicum] |
| 10119 | 2 | alanine-tRNA synthetase second additional domain-containing protein [Aromatoleum toluvorans] |
| 10126 | 2 | AAA domain-containing protein [Azoarcus rhizosphaerae] |
| 10127 | 2 | UUP1 family membrane protein [Aromatoleum pumilum] |
| 01390 | 2 | LUD domain-containing protein [Oryzomicrobium terrae] |
| 01394 | 2 | PEP-CTERM sorting domain-containing protein [Azoarcus halotolerans] |
| 10128 | 2 | MFS transporter [Uliginosibacterium gangwonense] |
| 00742 | 2 | hypothetical protein [Nitrogeniibacter aestuarii] |
| 00732 | 2 | hypothetical protein [Rhodocyclus purpureus] |
| 01398 | 2 | hypothetical protein [Azoarcus halotolerans] |
| 00726 | 2 | photosynthetic reaction center subunit L [Rhodocyclus purpureus] |
| 00744 | 2 | hypothetical protein [Nitrogeniibacter aestuarii] |
| 01408 | 2 | ParB/Srx family N-terminal domain-containing protein [Zoogloea ramigera] |
| 01379 | 2 | autotransporter outer membrane beta-barrel domain-containing protein [Azoarcus communis] |
| 01409 | 2 | hypothetical protein [Aromatoleum evansii] |
| 01410 | 2 | TRAP transporter small permease [Aromatoleum evansii] |
| 01413 | 2 | HK97 gp10 family phage protein [Aromatoleum evansii] |
| 01418 | 2 | tyrosine-type recombinase/integrase [Azoarcus communis] |
| 00718 | 2 | hypothetical protein [Candidatus Dactylopiibacterium carminicum] |
| 09671 | 2 | PEP-CTERM sorting domain-containing protein [Azoarcus communis] |
| 00713 | 2 | DUF2239 family protein [Azoarcus halotolerans] |
| 01422 | 2 | DUF637 domain-containing protein, partial [Azoarcus communis] |
| 00712 | 2 | hypothetical protein [Candidatus Dactylopiibacterium carminicum] |
| 00710 | 2 | DUF4124 domain-containing protein [Candidatus Dactylopiibacterium carminicum] |
| 00745 | 2 | hypothetical protein [Nitrogeniibacter aestuarii] |
| 01424 | 2 | hypothetical protein [Azoarcus communis] |
| 10140 | 2 | MULTISPECIES: antitoxin VbhA family protein [Betaproteobacteria] |
| 09627 | 2 | hypothetical protein [Nitrogeniibacter aestuarii] |
| 00685 | 2 | IS30 family transposase, partial [Zoogloea dura] |
| 01521 | 2 | class I SAM-dependent methyltransferase [Azoarcus communis] |
| 00762 | 2 | cobyrinic acid ac-diamide synthase [Zoogloea oleivorans] |
| 10100 | 2 | hypothetical protein [Denitromonas halophilus] |
| 10188 | 2 | GNAT family N-acetyltransferase [Rhodocyclus purpureus] |
| 09699 | 2 | DUF1302 family protein, partial [Zoogloea dura] |
| 01499 | 2 | ABC transporter substrate-binding protein [Azoarcus nasutitermitis] |
| 00764 | 2 | NIPSNAP family protein [Aromatoleum petrolei] |
| 00655 | 2 | Arm DNA-binding domain-containing protein [Azoarcus taiwanensis] |
| 00653 | 2 | hypothetical protein [Azoarcus taiwanensis] |
| 00649 | 2 | ESPR domain-containing protein, partial [Azoarcus indigens] |
| 01505 | 2 | DUF5681 domain-containing protein [Rhodocyclus purpureus] |
| 01510 | 2 | OmpA family protein [Propionivibrio dicarboxylicus] |
| 10193 | 2 | BCD family MFS transporter [Rhodocyclus purpureus] |
| 01515 | 2 | hypothetical protein [Niveibacterium microcysteis] |
| 01323 | 2 | hypothetical protein [Nitrogeniibacter aestuarii] |
| 10194 | 2 | caspase family protein [Rhodocyclus purpureus] |
| 09595 | 2 | hypothetical protein [Niveibacterium microcysteis] |
| 01322 | 2 | YidC/Oxa1 family membrane protein insertase [Nitrogeniibacter aestuarii] |
| 01320 | 2 | serine/threonine-protein phosphatase [Nitrogeniibacter aestuarii] |
| 00647 | 2 | DNA-binding protein [Azoarcus communis] |
| 01312 | 2 | hypothetical protein [Denitromonas halophilus] |
| 00646 | 2 | hypothetical protein [Azoarcus communis] |
| 09592 | 2 | hypothetical protein [Niveibacterium sp. COAC-50] |
| 10199 | 2 | hypothetical protein [Rhodocyclus purpureus] |
| 10200 | 2 | DUF3656 domain-containing protein [Rhodocyclus purpureus] |
| 10201 | 2 | polymorphic toxin type 15 domain-containing protein [Niveibacterium microcysteis] |
| 01306 | 2 | radical SAM protein [Azoarcus sp. L1K30] |
| 01324 | 2 | hypothetical protein [Nitrogeniibacter aestuarii] |
| 00656 | 2 | hypothetical protein [Aromatoleum toluvorans] |
| 09626 | 2 | hypothetical protein [Nitrogeniibacter aestuarii] |
| 09604 | 2 | DUF45 domain-containing protein, partial [Thauera phenylacetica] |
| 00684 | 2 | transporter substrate-binding domain-containing protein, partial [Zoogloea dura] |
| 00682 | 2 | acyl carrier protein [Azoarcus sp. TTM-91] |
| 01464 | 2 | OmpA family protein [Zoogloea oleivorans] |
| 09686 | 2 | DUF1579 domain-containing protein [Aromatoleum diolicum] |
| 10171 | 2 | hypothetical protein [Niveibacterium microcysteis] |
| 01337 | 2 | hypothetical protein [Propionivibrio dicarboxylicus] |
| 01333 | 2 | ATP-binding protein [Propionivibrio dicarboxylicus] |
| 01332 | 2 | hypothetical protein [Propionivibrio dicarboxylicus] |
| 00669 | 2 | hypothetical protein [Aromatoleum diolicum] |
| 00668 | 2 | CopK family periplasmic copper-binding protein [Aromatoleum evansii] |
| 00667 | 2 | hypothetical protein [Aromatoleum petrolei] |
| 01472 | 2 | K(+)-transporting ATPase subunit F [Uliginosibacterium aquaticum] |
| 00666 | 2 | hypothetical protein [Aromatoleum diolicum] |
| 01330 | 2 | [FeFe] hydrogenase, group A [Propionivibrio dicarboxylicus] |
| 01328 | 2 | phosphotransferase, partial [Aromatoleum toluclasticum] |
| 01327 | 2 | hypothetical protein [Aromatoleum toluclasticum] |
| 10106 | 2 | hypothetical protein [Aromatoleum toluclasticum] |
| 10183 | 2 | hypothetical protein [Aromatoleum petrolei] |

| | | |
|---|---|---|
| 00662 | 2 | hypothetical protein [Aromatoleum diolicum] |
| 10104 | 2 | ABZJ_00895 family protein [Nitrogeniibacter aestuarii] |
| 09609 | 2 | type II toxin-antitoxin system CcdA family antitoxin [Aromatoleum toluolicum] |
| 00660 | 2 | hypothetical protein [Aromatoleum diolicum] |
| 10103 | 2 | hypothetical protein [Nitrogeniibacter aestuarii] |
| 01491 | 2 | hypothetical protein [Niveibacterium microcysteis] |
| 10102 | 2 | hypothetical protein [Azospira oryzae] |
| 10272 | 2 | peptide-methionine (R)-S-oxide reductase MsrB [Propionivibrio dicarboxylicus] |
| 00579 | 2 | heme ABC transporter permease [Azoarcus rhizosphaerae] |
| 10274 | 2 | hypothetical protein [Aromatoleum tolulyticum] |
| 00842 | 2 | hypothetical protein [Thauera chlorobenzoica] |
| 00445 | 2 | glycosyltransferase family 2 protein [Azoarcus communis] |
| 01866 | 2 | hypothetical protein [Aromatoleum diolicum] |
| 00442 | 2 | class I SAM-dependent methyltransferase [Azoarcus sp. TTM-91] |
| 10361 | 2 | hypothetical protein [Azoarcus communis] |
| 09395 | 2 | hypothetical protein [Rhodocyclus purpureus] |
| 09394 | 2 | Hpt domain-containing protein [Zoogloea ramigera] |
| 10362 | 2 | hypothetical protein [Azoarcus communis] |
| 00440 | 2 | hypothetical protein [Aromatoleum toluclasticum] |
| 00438 | 2 | hypothetical protein [Aromatoleum toluolicum] |
| 09392 | 2 | hypothetical protein [Azoarcus halotolerans] |
| 09781 | 2 | hypothetical protein [Nitrogeniibacter aestuarii] |
| 09390 | 2 | helix-turn-helix domain-containing protein [Niveibacterium umoris] |
| 01868 | 2 | MULTISPECIES: hypothetical protein [Betaproteobacteria] |
| 01869 | 2 | hypothetical protein [Aromatoleum evansii] |
| 01212 | 2 | hypothetical protein [Denitromonas halophilus] |
| 00430 | 2 | SAM-dependent methyltransferase [Niveibacterium umoris] |
| 10369 | 2 | hypothetical protein, partial [Aromatoleum evansii] |
| 09386 | 2 | BON domain-containing protein [Azoarcus olearius] |
| 09784 | 2 | hypothetical protein [Aromatoleum toluclasticum] |
| 00850 | 2 | hypothetical protein [Azoarcus sp. KH32C] |
| 01206 | 2 | contractile injection system tape measure protein, partial [Zoogloea oleivorans] |
| 09383 | 2 | DUF262 domain-containing protein [Aromatoleum aromaticum] |
| 00852 | 2 | cobalamin (vitamin B12) biosynthesis CbiX protein [Thauera linaloolentis] |
| 01195 | 2 | SURF1 family protein [Azoarcus sp. DD4] |
| 10031 | 2 | DUF2917 domain-containing protein [Candidatus Dactylopiibacterium carminicum] |
| 10357 | 2 | hypothetical protein [Thauera lacus] |
| 09777 | 2 | OmpA family protein [Nitrogeniibacter aestuarii] |
| 10378 | 2 | HAD-IIIC family phosphatase [Azospira oryzae] |
| 01855 | 2 | hypothetical protein [Azoarcus taiwanensis] |
| 01818 | 2 | DUF4255 domain-containing protein [Zoogloea dura] |
| 01819 | 2 | hypothetical protein [Aromatoleum aromaticum] |
| 01232 | 2 | DUF4391 domain-containing protein [Azoarcus sp. DD4] |
| 01827 | 2 | DUF2169 domain-containing protein [Azoarcus nasutitermitis] |
| 01828 | 2 | hypothetical protein [Azoarcus halotolerans] |
| 09418 | 2 | hypothetical protein [Aromatoleum bremense] |
| 01830 | 2 | nitrogen fixation protein [Azoarcus halotolerans] |
| 01231 | 2 | lysozyme inhibitor LprI family protein [Niveibacterium microcysteis] |
| 10350 | 2 | hypothetical protein [Thauera lacus] |
| 09415 | 2 | VirB3 family type IV secretion system protein [Denitromonas sp. IR12] |
| 01228 | 2 | outer membrane beta-barrel protein [Niveibacterium microcysteis] |
| 01835 | 2 | OmpA family protein [Azoarcus halotolerans] |
| 01226 | 2 | class I SAM-dependent methyltransferase [Niveibacterium sp. COAC-50] |
| 00450 | 2 | hypothetical protein [Azoarcus sp. TTM-91] |
| 09412 | 2 | HAMP domain-containing protein [Aromatoleum toluclasticum] |
| 01840 | 2 | ABC transporter ATP-binding protein [Denitromonas sp. IR12] |
| 09410 | 2 | RDD family protein [Uliginosibacterium gangwonense] |
| 10042 | 2 | hypothetical protein [Aromatoleum evansii] |
| 01842 | 2 | DEAD/DEAH box helicase [Aromatoleum evansii] |
| 00839 | 2 | hypothetical protein [Aromatoleum pumilum] |
| 09408 | 2 | GTP-binding protein [Azoarcus sp. KH32C] |
| 09407 | 2 | DUF2974 domain-containing protein [Uliginosibacterium gangwonense] |
| 09406 | 2 | HAMP domain-containing histidine kinase [Uliginosibacterium aquaticum] |
| 09404 | 2 | hypothetical protein [Uliginosibacterium gangwonense] |
| 09776 | 2 | transposase, partial [Aromatoleum toluclasticum] |
| 09382 | 2 | hypothetical protein [Aromatoleum aromaticum] |
| 01186 | 2 | hypothetical protein [Aromatoleum toluvorans] |
| 01659 | 2 | DUF1643 domain-containing protein [Aromatoleum evansii] |
| 09358 | 2 | aminotransferase class V-fold PLP-dependent enzyme [Nitrogeniibacter aestuarii] |
| 00866 | 2 | hypothetical protein [Thauera chlorobenzoica] |
| 00869 | 2 | hypothetical protein [Thauera butanivorans] |
| 00384 | 2 | hypothetical protein [Aromatoleum evansii] |
| 00381 | 2 | 2Fe-2S iron-sulfur cluster-binding protein [Zoogloea oleivorans] |
| 09802 | 2 | hypothetical protein [Aromatoleum tolulyticum] |
| 00870 | 2 | hypothetical protein [Thauera phenolivorans] |
| 00380 | 2 | hypothetical protein [Zoogloea oleivorans] |
| 10017 | 2 | alpha/beta hydrolase [Uliginosibacterium aquaticum] |
| 00872 | 2 | hypothetical protein [Aromatoleum pumilum] |
| 00873 | 2 | peroxiredoxin [Aromatoleum pumilum] |
| 10403 | 2 | glycosyltransferase [Aromatoleum toluclasticum] |
| 00379 | 2 | SAVED domain-containing protein [Azoarcus sp. DD4] |
| 09805 | 2 | DUF3150 domain-containing protein [Aromatoleum aromaticum] |
| 00377 | 2 | hypothetical protein [Azoarcus communis] |
| 00372 | 2 | DUF4153 domain-containing protein [Azoarcus nasutitermitis] |
| 01917 | 2 | hypothetical protein [Thauera linaloolentis] |
| 00370 | 2 | S-layer family protein [Azoarcus nasutitermitis] |
| 00369 | 2 | hypothetical protein [Aromatoleum toluolicum] |
| 10409 | 2 | DUF3467 domain-containing protein [Azoarcus communis] |
| 10410 | 2 | Rha family transcriptional regulator [Azoarcus rhizosphaerae] |
| 01163 | 2 | hypothetical protein [Aromatoleum buckelii] |
| 00874 | 2 | outer membrane beta-barrel protein [Aromatoleum pumilum] |
| 00876 | 2 | hypothetical protein [Aromatoleum pumilum] |
| 01922 | 2 | MFS transporter [Candidatus Dactylopiibacterium carminicum] |
| 00360 | 2 | hypothetical protein [Azospira oryzae] |

| | | |
|---|---|---|
| 09357 | 2 | thiol-disulfide oxidoreductase DCC family protein [Nitrogeniibacter aestuarii] |
| 00864 | 2 | hypothetical protein [Thauera chlorobenzoica] |
| 09380 | 2 | DUF2304 domain-containing protein [Aromatoleum aromaticum] |
| 09360 | 2 | STAS domain-containing protein [Aromatoleum toluclasticum] |
| 09377 | 2 | DUF3147 family protein [Aromatoleum aromaticum] |
| 09376 | 2 | IS630 family transposase [Aromatoleum bremense] |
| 00855 | 2 | GNAT family N-acetyltransferase [Azoarcus halotolerans] |
| 09668 | 2 | Ig-like domain-containing protein [Aromatoleum evansii] |
| 10380 | 2 | MULTISPECIES: hypothetical protein [Azospira] |
| 01182 | 2 | hypothetical protein [Aromatoleum toluvorans] |
| 01177 | 2 | DUF3305 domain-containing protein [Aromatoleum pumilum] |
| 01175 | 2 | hypothetical protein [Aromatoleum aromaticum] |
| 09373 | 2 | hypothetical protein [Aromatoleum tolulyticum] |
| 09372 | 2 | hypothetical protein [Aromatoleum toluclasticum] |
| 00857 | 2 | hypothetical protein [Aromatoleum toluvorans] |
| 00415 | 2 | hypothetical protein [Aromatoleum buckelii] |
| 01884 | 2 | hypothetical protein [Azoarcus halotolerans] |
| 10026 | 2 | PEP-CTERM sorting domain-containing protein [Niveibacterium microcysteis] |
| 01887 | 2 | HEPN domain-containing protein [Azoarcus sp. DD4] |
| 00401 | 2 | hypothetical protein [Zoogloea dura] |
| 09366 | 2 | hypothetical protein [Candidatus Dactylopiibacterium carminicum] |
| 01172 | 2 | hypothetical protein [Azoarcus communis] |
| 00400 | 2 | hypothetical protein [Zoogloea dura] |
| 00398 | 2 | hypothetical protein [Rhodocyclus purpureus] |
| 01896 | 2 | alpha/beta fold hydrolase [Uliginosibacterium aquaticum] |
| 09362 | 2 | UxaA family hydrolase [Azoarcus communis] |
| 00397 | 2 | hypothetical protein [Zoogloea dura] |
| 01171 | 2 | hypothetical protein [Azoarcus communis] |
| 09361 | 2 | lycopene cyclase domain-containing protein [Aromatoleum toluclasticum] |
| 09423 | 2 | hypothetical protein [Thauera humireducens] |
| 09425 | 2 | MmoB/DmpM family protein [Azoarcus sp. DD4] |
| 00461 | 2 | hypothetical protein [Niveibacterium sp. COAC-50] |
| 10069 | 2 | SPFH domain-containing protein [Niveibacterium umoris] |
| 00803 | 2 | TRAP transporter small permease subunit, partial [Propionivibrio dicarboxylicus] |
| 10297 | 2 | HEPN domain-containing protein [Azoarcus communis] |
| 09491 | 2 | hypothetical protein [Azoarcus olearius] |
| 09490 | 2 | NifU family protein [Azoarcus nasutitermitis] |
| 10065 | 2 | PEP-CTERM sorting domain-containing protein [Nitrogeniibacter aestuarii] |
| 01699 | 2 | CcdB family protein [Thauera linaloolentis] |
| 00550 | 2 | ATP-binding protein [Candidatus Dactylopiibacterium carminicum] |
| 00540 | 2 | DUF5677 domain-containing protein [Azoarcus rhizosphaerae] |
| 00536 | 2 | hypothetical protein [Azoarcus communis] |
| 10063 | 2 | hypothetical protein [Nitrogeniibacter aestuarii] |
| 00528 | 2 | MULTISPECIES: hypothetical protein [Proteobacteria] |
| 00523 | 2 | hypothetical protein [Thauera aromatica] |
| 09484 | 2 | hypothetical protein [Aromatoleum petrolei] |
| 00806 | 2 | hypothetical protein [Aromatoleum aromaticum] |
| 10309 | 2 | hypothetical protein [Aromatoleum toluclasticum] |
| 00518 | 2 | hypothetical protein [Azoarcus olearius] |
| 01255 | 2 | hypothetical protein [Aromatoleum toluclasticum] |
| 00517 | 2 | hypothetical protein [Thauera butanivorans] |
| 01716 | 2 | hypothetical protein [Zoogloea dura] |
| 00516 | 2 | DUF523 domain-containing protein [Azoarcus communis] |
| 00817 | 2 | hypothetical protein [Azoarcus rhizosphaerae] |
| 00515 | 2 | ATP-binding protein [Thauera butanivorans] |
| 10058 | 2 | GNAT family N-acetyltransferase [Niveibacterium microcysteis] |
| 00513 | 2 | MULTISPECIES: type II toxin-antitoxin system HicB family antitoxin [Aromatoleum] |
| 00509 | 2 | DUF6521 family protein [Azoarcus halotolerans] |
| 10067 | 2 | hypothetical protein [Nitrogeniibacter aestuarii] |
| 01265 | 2 | helix-turn-helix domain-containing protein [Aromatoleum buckelii] |
| 10043 | 2 | Fic family protein [Thauera phenylacetica] |
| 09745 | 2 | hypothetical protein [Nitrogeniibacter aestuarii] |
| 00578 | 2 | DUF4160 domain-containing protein [Azoarcus rhizosphaerae] |
| 00577 | 2 | hypothetical protein [Azoarcus nasutitermitis] |
| 00575 | 2 | YdeI/OmpD-associated family protein [Azoarcus nasutitermitis] |
| 00574 | 2 | DUF4857 domain-containing protein [Azoarcus nasutitermitis] |
| 09913 | 2 | MULTISPECIES: hypothetical protein [Proteobacteria] |
| 10280 | 2 | nuclear transport factor 2 family protein [Aromatoleum aromaticum] |
| 10282 | 2 | transposase [Aromatoleum aromaticum] |
| 01669 | 2 | hypothetical protein [Aromatoleum pumilum] |
| 10074 | 2 | hypothetical protein [Aromatoleum aromaticum] |
| 01672 | 2 | hypothetical protein [Thauera aromatica] |
| 00570 | 2 | thioredoxin family protein [Azoarcus nasutitermitis] |
| 09506 | 2 | hypothetical protein [Aromatoleum toluclasticum] |
| 00569 | 2 | DUF4852 domain-containing protein [Azoarcus nasutitermitis] |
| 01674 | 2 | serine protease [Rhodocyclus purpureus] |
| 10286 | 2 | hypothetical protein [Azoarcus olearius] |
| 10288 | 2 | MULTISPECIES: hypothetical protein [Thauera] |
| 01685 | 2 | RHS domain-containing protein [Azoarcus communis] |
| 10292 | 2 | MULTISPECIES: hypothetical protein [Proteobacteria] |
| 09499 | 2 | hypothetical protein, partial [Zoogloea ramigera] |
| 09498 | 2 | PAAR domain-containing protein, partial [Zoogloea ramigera] |
| 10073 | 2 | hypothetical protein [Aromatoleum bremense] |
| 00800 | 2 | ABC transporter permease, partial [Propionivibrio dicarboxylicus] |
| 00802 | 2 | hypothetical protein [Propionivibrio dicarboxylicus] |
| 01268 | 2 | hypothetical protein [Azoarcus sp. KH32C] |
| 01694 | 2 | NAD(P)(+) transhydrogenase (Re/Si-specific) subunit beta, partial [Azoarcus indigens] |
| 01728 | 2 | hypothetical protein, partial [Denitromonas halophilus] |
| 10319 | 2 | N-6 DNA methylase [Thauera humireducens] |
| 10320 | 2 | hypothetical protein [Thauera humireducens] |
| 00820 | 2 | hypothetical protein, partial [Thauera aminoaromatica] |
| 09447 | 2 | hypothetical protein [Aromatoleum pumilum] |
| 01245 | 2 | Hsp20 family protein [Candidatus Dactylopiibacterium carminicum] |
| 10052 | 2 | AcaB family transcriptional regulator [Rhodocyclus purpureus] |

| | | |
|---|---|---|
| **01785** | 2 | MULTISPECIES: toprim domain-containing protein [Betaproteobacteria] |
| **01788** | 2 | RloB family protein [Aromatoleum toluolicum] |
| **00823** | 2 | DUF1343 domain-containing protein [Azoarcus sp. KH32C] |
| **09761** | 2 | methyltransferase domain-containing protein [Azoarcus communis] |
| **00480** | 2 | MULTISPECIES: hypothetical protein [Proteobacteria] |
| **09442** | 2 | hypothetical protein [Thauera propionica] |
| **00478** | 2 | MULTISPECIES: hypothetical protein [Proteobacteria] |
| **09440** | 2 | hypothetical protein [Thauera chlorobenzoica] |
| **09438** | 2 | hypothetical protein [Azoarcus taiwanensis] |
| **01792** | 2 | hypothetical protein [Azoarcus nasutitermitis] |
| **09435** | 2 | hypothetical protein [Thauera aromatica] |
| **01239** | 2 | rhomboid family intramembrane serine protease [Candidatus Dactylopiibacterium carminicum] |
| **09434** | 2 | RES family NAD+ phosphorylase [Thauera phenolivorans] |
| **09433** | 2 | hypothetical protein [Azoarcus sp. KH32C] |
| **00472** | 2 | MULTISPECIES: hypothetical protein [Thauera] |
| **00834** | 2 | S-layer protein [Uliginosibacterium gangwonense] |
| **01238** | 2 | MULTISPECIES: hypothetical protein [Uliginosibacterium] |
| **01803** | 2 | hypothetical protein [Uliginosibacterium aquaticum] |
| **00470** | 2 | FAD/NAD(P)-binding protein [Thauera aminoaromatica] |
| **00835** | 2 | ATP-binding protein [Aromatoleum toluclasticum] |
| **01808** | 2 | ACT domain-containing protein [Uliginosibacterium aquaticum] |
| **10044** | 2 | glycoside hydrolase family 104 protein [Azoarcus sp. Aa7] |
| **00494** | 2 | GDSL-type esterase/lipase family protein [Uliginosibacterium gangwonense] |
| **01779** | 2 | SCO family protein [Aromatoleum toluvorans] |
| **09449** | 2 | hypothetical protein [Aromatoleum pumilum] |
| **10326** | 2 | hypothetical protein [Azoarcus olearius] |
| **01730** | 2 | hypothetical protein [Thauera aromatica] |
| **01733** | 2 | hypothetical protein [Zoogloea dura] |
| **01736** | 2 | PQQ-dependent sugar dehydrogenase [Zoogloea dura] |
| **09469** | 2 | GyrI-like domain-containing protein [Thauera hydrothermalis] |
| **01741** | 2 | DUF3944 domain-containing protein [Oryzomicrobium terrae] |
| **01253** | 2 | hypothetical protein [Azoarcus sp. KH32C] |
| **00507** | 2 | hypothetical protein [Azoarcus communis] |
| **00506** | 2 | surface lipoprotein assembly modifier, partial [Thauera linaloolentis] |
| **10325** | 2 | hypothetical protein, partial [Thauera butanivorans] |
| **01754** | 2 | tyrosine-type recombinase/integrase [Aromatoleum buckelii] |
| **01755** | 2 | winged helix-turn-helix domain-containing protein, partial [Aromatoleum buckelii] |
| **01762** | 2 | hypothetical protein [Aromatoleum evansii] |
| **01246** | 2 | hypothetical protein [Candidatus Dactylopiibacterium carminicum] |
| **01764** | 2 | PEP-CTERM sorting domain-containing protein [Azoarcus communis] |
| **00503** | 2 | hypothetical protein [Aromatoleum toluclasticum] |
| **09458** | 2 | hypothetical protein [Aromatoleum tolulyticum] |
| **01250** | 2 | RluA family pseudouridine synthase, partial [Aromatoleum buckelii] |
| **01248** | 2 | hypothetical protein [Candidatus Dactylopiibacterium carminicum] |
| **01768** | 2 | hypothetical protein, partial [Aromatoleum toluclasticum] |
| **01247** | 2 | phospholipid carrier-dependent glycosyltransferase [Candidatus Dactylopiibacterium carminicum] |
| **09454** | 2 | type II toxin-antitoxin system VapC family toxin [Thauera lacus] |
| **01769** | 2 | HAMP domain-containing histidine kinase, partial [Aromatoleum buckelii] |
| **00499** | 2 | metallophosphoesterase [Aromatoleum toluclasticum] |
| **01772** | 2 | twin-arginine translocation signal domain-containing protein [Aromatoleum pumilum] |
| **02298** | 2 | hypothetical protein [Aromatoleum petrolei] |
| **06270** | 2 | hypothetical protein [Azoarcus olearius] |
| **02300** | 2 | hypothetical protein [Azoarcus nasutitermitis] |
| **07221** | 2 | RES family NAD+ phosphorylase [Thauera linaloolentis] |
| **04978** | 2 | DUF2345 domain-containing protein, partial [Denitromonas sp. IR12] |
| **04977** | 2 | hypothetical protein [Denitromonas halophilus] |
| **04972** | 2 | RHS repeat-associated core domain-containing protein, partial [Azoarcus sp. L1K30] |
| **04971** | 2 | hypothetical protein [Azoarcus sp. L1K30] |
| **04967** | 2 | DUF4189 domain-containing protein [Aromatoleum petrolei] |
| **07185** | 2 | hypothetical protein [Thauera aromatica] |
| **07186** | 2 | hypothetical protein [Azoarcus communis] |
| **04963** | 2 | acetophenone carboxylase epsilon subunit [Aromatoleum aromaticum] |
| **04962** | 2 | hypothetical protein [Aromatoleum bremense] |
| **04961** | 2 | hypothetical protein [Aromatoleum bremense] |
| **07197** | 2 | hypothetical protein [Azoarcus communis] |
| **04959** | 2 | VCBS domain-containing protein, partial [Niveibacterium umoris] |
| **04958** | 2 | cytochrome C [Nitrogeniibacter mangrovi] |
| **07203** | 2 | ring-opening amidohydrolase [Azoarcus communis] |
| **04948** | 2 | hypothetical protein [Nitrogeniibacter aestuarii] |
| **04946** | 2 | hypothetical protein [Nitrogeniibacter aestuarii] |
| **04943** | 2 | hypothetical protein [Nitrogeniibacter aestuarii] |
| **04941** | 2 | hypothetical protein [Niveibacterium microcysteis] |
| **07212** | 2 | TIR domain-containing protein [Zoogloea oleivorans] |
| **04933** | 2 | hypothetical protein [Rhodocyclus purpureus] |
| **07218** | 2 | hypothetical protein [Aromatoleum bremense] |
| **04979** | 2 | HEPN family nuclease [Denitromonas sp. IR12] |
| **07172** | 2 | hypothetical protein [Niveibacterium sp. COAC-50] |
| **07169** | 2 | hypothetical protein [Thauera butanivorans] |
| **07156** | 2 | hypothetical protein [Azoarcus nasutitermitis] |
| **05013** | 2 | T6SS immunity protein Tli4 family protein [Aromatoleum tolulyticum] |
| **07142** | 2 | hypothetical protein [Uliginosibacterium aquaticum] |
| **05005** | 2 | TraB/GumN family protein [Propionivibrio dicarboxylicus] |
| **05001** | 2 | glycosyltransferase [Azospira oryzae] |
| **04999** | 2 | hypothetical protein [Aromatoleum pumilum] |
| **07150** | 2 | hypothetical protein [Aromatoleum toluclasticum] |
| **07151** | 2 | cupin domain-containing protein [Aromatoleum evansii] |
| **04996** | 2 | hypothetical protein [Propionivibrio dicarboxylicus] |
| **07154** | 2 | DUF1090 domain-containing protein [Thauera butanivorans] |
| **04993** | 2 | hypothetical protein [Aromatoleum toluclasticum] |
| **04983** | 2 | hypothetical protein [Nitrogeniibacter aestuarii] |
| **07158** | 2 | hypothetical protein [Thauera humireducens] |
| **07159** | 2 | glycosyltransferase [Azoarcus sp. L1K30] |
| **07161** | 2 | hypothetical protein [Azoarcus olearius] |
| **04990** | 2 | DUF695 domain-containing protein [Nitrogeniibacter aestuarii] |

| | | |
|---|---|---|
| 04989 | 2 | hypothetical protein [Nitrogeniibacter aestuarii] |
| 04988 | 2 | SRPBCC family protein [Nitrogeniibacter aestuarii] |
| 04986 | 2 | hypothetical protein [Nitrogeniibacter aestuarii] |
| 07166 | 2 | Rpn family recombination-promoting nuclease/putative transposase [Azoarcus nasutitermitis] |
| 04984 | 2 | hypothetical protein [Nitrogeniibacter aestuarii] |
| 04931 | 2 | alcohol dehydrogenase catalytic domain-containing protein, partial [Azoarcus communis] |
| 07222 | 2 | DNA/RNA non-specific endonuclease, partial [Aromatoleum buckelii] |
| 07676 | 2 | hypothetical protein [Propionivibrio dicarboxylicus] |
| 07224 | 2 | class I SAM-dependent methyltransferase [Aromatoleum evansii] |
| 04850 | 2 | GIY-YIG nuclease family protein [Aromatoleum evansii] |
| 07274 | 2 | hypothetical protein [Uliginosibacterium aquaticum] |
| 04848 | 2 | sulfotransferase [Aromatoleum evansii] |
| 07277 | 2 | hypothetical protein [Uliginosibacterium aquaticum] |
| 07279 | 2 | 5'-methylthioadenosine/S-adenosylhomocysteine nucleosidase [Uliginosibacterium gangwonense] |
| 04845 | 2 | SoxR reducing system RseC family protein, partial [Aromatoleum evansii] |
| 04842 | 2 | hypothetical protein [Aromatoleum buckelii] |
| 07285 | 2 | DUF2835 domain-containing protein [Candidatus Dactylopiibacterium carminicum] |
| 04824 | 2 | hypothetical protein [Zoogloea oleivorans] |
| 04821 | 2 | GIY-YIG nuclease family protein [Aromatoleum bremense] |
| 04819 | 2 | hypothetical protein [Azospira oryzae] |
| 07294 | 2 | hypothetical protein [Niveibacterium microcysteis] |
| 04812 | 2 | hypothetical protein [Denitromonas halophilus] |
| 07298 | 2 | hypothetical protein [Niveibacterium microcysteis] |
| 04807 | 2 | hypothetical protein [Thauera aromatica] |
| 07304 | 2 | DUF5818 domain-containing protein [Thauera hydrothermalis] |
| 07309 | 2 | hypothetical protein [Azoarcus sp. DD4] |
| 07310 | 2 | DUF917 domain-containing protein [Azoarcus rhizosphaerae] |
| 07311 | 2 | hypothetical protein [Azoarcus olearius] |
| 07312 | 2 | HNH endonuclease [Aromatoleum evansii] |
| 04794 | 2 | type III secretion inner membrane ring lipoprotein SctJ [Azoarcus rhizosphaerae] |
| 07272 | 2 | NifU family protein [Aromatoleum toluolicum] |
| 07271 | 2 | CDGSH iron-sulfur domain-containing protein [Zoogloea dura] |
| 04851 | 2 | hypothetical protein [Aromatoleum evansii] |
| 04898 | 2 | hypothetical protein [Candidatus Dactylopiibacterium carminicum] |
| 04928 | 2 | hypothetical protein [Rhodocyclus purpureus] |
| 04925 | 2 | hypothetical protein [Aromatoleum evansii] |
| 07228 | 2 | hypothetical protein [Azoarcus communis] |
| 07230 | 2 | hypothetical protein [Azoarcus communis] |
| 07231 | 2 | DUF935 family protein [Azoarcus communis] |
| 04912 | 2 | hypothetical protein [Azoarcus taiwanensis] |
| 04911 | 2 | heparinase II/III family protein [Candidatus Dactylopiibacterium carminicum] |
| 04899 | 2 | hypothetical protein [Azoarcus sp. TTM-91] |
| 07240 | 2 | SDR family NAD(P)-dependent oxidoreductase [Azoarcus nasutitermitis] |
| 07243 | 2 | hypothetical protein [Aromatoleum toluvorans] |
| 07269 | 2 | tetratricopeptide repeat protein [Zoogloea dura] |
| 04889 | 2 | class II aldolase/adducin family protein [Thauera humireducens] |
| 07251 | 2 | Hpt domain-containing protein [Aromatoleum diolicum] |
| 07252 | 2 | MULTISPECIES: hypothetical protein [Betaproteobacteria] |
| 07253 | 2 | hypothetical protein [Aromatoleum tolulyticum] |
| 04874 | 2 | hypothetical protein [Thauera aromatica] |
| 04866 | 2 | ATP-dependent zinc protease [Aromatoleum pumilum] |
| 04857 | 2 | hypothetical protein [Azoarcus communis] |
| 04856 | 2 | hypothetical protein [Azoarcus communis] |
| 04852 | 2 | glycine--tRNA ligase subunit alpha, partial [Aromatoleum evansii] |
| 07137 | 2 | hypothetical protein [Uliginosibacterium gangwonense] |
| 05014 | 2 | hypothetical protein [Aromatoleum tolulyticum] |
| 05024 | 2 | MULTISPECIES: hypothetical protein [Azospira] |
| 05029 | 2 | RnfABCDGE type electron transport complex subunit D, partial [Thauera aminoaromatica] |
| 05219 | 2 | hypothetical protein [Aromatoleum evansii] |
| 05216 | 2 | hypothetical protein [Aromatoleum buckelii] |
| 05215 | 2 | hypothetical protein [Aromatoleum bremense] |
| 07004 | 2 | hypothetical protein [Denitromonas halophilus] |
| 07007 | 2 | hypothetical protein [Candidatus Dactylopiibacterium carminicum] |
| 05201 | 2 | PcfJ domain-containing protein [Azoarcus halotolerans] |
| 05193 | 2 | peroxiredoxin [Azoarcus halotolerans] |
| 05188 | 2 | hypothetical protein [Zoogloea oleivorans] |
| 07016 | 2 | transglutaminase-like cysteine peptidase [Niveibacterium microcysteis] |
| 05186 | 2 | PIN domain-containing protein [Aromatoleum bremense] |
| 05185 | 2 | hypothetical protein [Zoogloea dura] |
| 07021 | 2 | hypothetical protein [Niveibacterium sp. COAC-50] |
| 07023 | 2 | hypothetical protein [Niveibacterium microcysteis] |
| 07028 | 2 | hypothetical protein [Candidatus Dactylopiibacterium carminicum] |
| 07029 | 2 | hypothetical protein [Candidatus Dactylopiibacterium carminicum] |
| 07030 | 2 | hypothetical protein [Candidatus Dactylopiibacterium carminicum] |
| 05167 | 2 | hypothetical protein [Aromatoleum toluclasticum] |
| 05159 | 2 | metallophosphoesterase [Rhodocyclus purpureus] |
| 05157 | 2 | filamentous hemagglutinin N-terminal domain-containing protein [Azoarcus communis] |
| 07040 | 2 | hypothetical protein [Aromatoleum aromaticum] |
| 05149 | 2 | hypothetical protein [Azoarcus rhizosphaerae] |
| 05220 | 2 | nucleotidyltransferase domain-containing protein [Aromatoleum evansii] |
| 06999 | 2 | hypothetical protein [Aromatoleum pumilum] |
| 05223 | 2 | glycosyltransferase [Aromatoleum evansii] |
| 05244 | 2 | SDR family oxidoreductase [Niveibacterium microcysteis] |
| 05270 | 2 | hypothetical protein [Rhodocyclus purpureus] |
| 05263 | 2 | hypothetical protein [Azoarcus sp. Aa7] |
| 06968 | 2 | hypothetical protein, partial [Azoarcus indigens] |
| 05260 | 2 | MULTISPECIES: ribbon-helix-helix protein, CopG family [unclassified Azoarcus] |
| 05258 | 2 | phage holin family protein [Candidatus Dactylopiibacterium carminicum] |
| 06973 | 2 | TonB C-terminal domain-containing protein, partial [Thauera phenylacetica] |
| 05253 | 2 | hypothetical protein [Propionivibrio dicarboxylicus] |
| 05247 | 2 | Hint domain-containing protein [Nitrogeniibacter aestuarii] |
| 06979 | 2 | hypothetical protein [Aromatoleum toluvorans] |
| 05241 | 2 | hypothetical protein [Niveibacterium sp. COAC-50] |
| 05224 | 2 | MULTISPECIES: zeta toxin family protein [Betaproteobacteria] |

| | | |
|---|---|---|
| 05234 | 2 | TIGR04255 family protein [Azoarcus communis] |
| 06985 | 2 | hypothetical protein [Aromatoleum toluolicum] |
| 05233 | 2 | hypothetical protein [Thauera phenylacetica] |
| 05231 | 2 | hypothetical protein [Azoarcus communis] |
| 05228 | 2 | nitronate monooxygenase, partial [Aromatoleum evansii] |
| 06990 | 2 | KTSC domain-containing protein [Azoarcus nasutitermitis] |
| 06991 | 2 | hypothetical protein [Aromatoleum toluolicum] |
| 05227 | 2 | hypothetical protein [Azoarcus sp. DD4] |
| 05226 | 2 | hypothetical protein [Aromatoleum evansii] |
| 05147 | 2 | hypothetical protein [Azoarcus nasutitermitis] |
| 05146 | 2 | hypothetical protein [Azoarcus rhizosphaerae] |
| 05144 | 2 | hypothetical protein [Azoarcus nasutitermitis] |
| 07112 | 2 | autotransporter outer membrane beta-barrel domain-containing protein [Propionivibrio dicarboxylicus] |
| 05064 | 2 | hypothetical protein [Azoarcus taiwanensis] |
| 07102 | 2 | hypothetical protein [Propionivibrio dicarboxylicus] |
| 05063 | 2 | hypothetical protein [Thauera humireducens] |
| 05062 | 2 | DUF4145 domain-containing protein [Thauera humireducens] |
| 05060 | 2 | SAVED domain-containing protein [Thauera linaloolentis] |
| 05057 | 2 | hypothetical protein [Aromatoleum aromaticum] |
| 05053 | 2 | hypothetical protein [Aromatoleum toluclasticum] |
| 05048 | 2 | MULTISPECIES: large ATP-binding protein [Rhodocyclales] |
| 07111 | 2 | SLBB domain-containing protein [Propionivibrio dicarboxylicus] |
| 07113 | 2 | hypothetical protein [Niveibacterium umoris] |
| 07096 | 2 | caspase family protein [Propionivibrio dicarboxylicus] |
| 05045 | 2 | DUF4870 domain-containing protein [Candidatus Dactylopiibacterium carminicum] |
| 05044 | 2 | DUF4034 domain-containing protein [Uliginosibacterium gangwonense] |
| 07118 | 2 | phage holin, lambda family [Aromatoleum aromaticum] |
| 05041 | 2 | hypothetical protein [Uliginosibacterium gangwonense] |
| 07122 | 2 | glycosyltransferase [Thauera aminoaromatica] |
| 05037 | 2 | chemotaxis protein CheW [Aromatoleum diolicum] |
| 07125 | 2 | MULTISPECIES: helix-turn-helix domain-containing protein [Proteobacteria] |
| 05033 | 2 | hypothetical protein, partial [Thauera aminoaromatica] |
| 05032 | 2 | DUF2868 domain-containing protein, partial [Aromatoleum buckelii] |
| 05068 | 2 | HNH endonuclease [Thauera butanivorans] |
| 07094 | 2 | ThiF family adenylyltransferase [Nitrogeniibacter aestuarii] |
| 05143 | 2 | Fe-S-containing protein [Azoarcus nasutitermitis] |
| 05101 | 2 | acetamidase/formamidase family protein [Aromatoleum pumilum] |
| 07049 | 2 | hypothetical protein [Azoarcus communis] |
| 05138 | 2 | hypothetical protein [Azoarcus nasutitermitis] |
| 05126 | 2 | GNAT family N-acetyltransferase [Azospira oryzae] |
| 07057 | 2 | hypothetical protein [Niveibacterium umoris] |
| 05115 | 2 | hypothetical protein [Azoarcus communis] |
| 05112 | 2 | hypothetical protein [Azoarcus communis] |
| 05109 | 2 | NAD(P)H-binding protein [Azoarcus communis] |
| 05107 | 2 | DUF6396 domain-containing protein [Thauera lacus] |
| 05106 | 2 | hypothetical protein [Azoarcus taiwanensis] |
| 05100 | 2 | amidohydrolase family protein [Aromatoleum pumilum] |
| 05078 | 2 | LysR substrate-binding domain-containing protein [Thauera butanivorans] |
| 07078 | 2 | glycosyltransferase family 39 protein [Azoarcus sp. L1K30] |
| 05096 | 2 | Com family DNA-binding transcriptional regulator [Aromatoleum pumilum] |
| 05094 | 2 | hypothetical protein [Oryzomicrobium terrae] |
| 05092 | 2 | hypothetical protein [Thauera humireducens] |
| 05088 | 2 | DUF4041 domain-containing protein [Rhodocyclus purpureus] |
| 05083 | 2 | DUF2795 domain-containing protein [Azoarcus olearius] |
| 05082 | 2 | ABC transporter permease [Azoarcus olearius] |
| 07091 | 2 | hypothetical protein [Nitrogeniibacter aestuarii] |
| 05080 | 2 | glycosyltransferase family 39 protein [Azoarcus olearius] |
| 04788 | 2 | 4'-phosphopantetheinyl transferase superfamily protein [Azoarcus nasutitermitis] |
| 04786 | 2 | GntP family permease [Azoarcus nasutitermitis] |
| 07320 | 2 | DUF2628 domain-containing protein [Niveibacterium umoris] |
| 04563 | 2 | hypothetical protein [Nitrogeniibacter aestuarii] |
| 04462 | 2 | hypothetical protein [Aromatoleum aromaticum] |
| 04461 | 2 | PilN domain-containing protein [Thauera butanivorans] |
| 04457 | 2 | hypothetical protein [Nitrogeniibacter mangrovi] |
| 04455 | 2 | hypothetical protein [Nitrogeniibacter aestuarii] |
| 04449 | 2 | hypothetical protein [Azoarcus halotolerans] |
| 07555 | 2 | VCBS repeat-containing protein [Aromatoleum buckelii] |
| 07556 | 2 | hypothetical protein [Aromatoleum buckelii] |
| 04435 | 2 | MBL fold metallo-hydrolase [Zoogloea dura] |
| 07558 | 2 | SDR family NAD(P)-dependent oxidoreductase [Azoarcus sp. L1K30] |
| 04431 | 2 | hypothetical protein [Denitromonas halophilus] |
| 04428 | 2 | hypothetical protein [Denitromonas halophilus] |
| 07566 | 2 | hypothetical protein [Azoarcus communis] |
| 04413 | 2 | sterol desaturase family protein [Zoogloea oleivorans] |
| 04405 | 2 | cell division protein FtsK [Zoogloea dura] |
| 04400 | 2 | hypothetical protein [Azoarcus rhizosphaerae] |
| 04397 | 2 | DUF3540 domain-containing protein [Azoarcus nasutitermitis] |
| 04396 | 2 | DEAD/DEAH box helicase [Azoarcus nasutitermitis] |
| 07582 | 2 | methylaspartate mutase subunit S [Aromatoleum toluvorans] |
| 04394 | 2 | tetratricopeptide repeat protein [Azoarcus nasutitermitis] |
| 07585 | 2 | hypothetical protein [Aromatoleum petrolei] |
| 07587 | 2 | hypothetical protein, partial [Aromatoleum diolicum] |
| 04467 | 2 | VWA domain-containing protein [Aromatoleum aromaticum] |
| 04470 | 2 | hypothetical protein [Aromatoleum evansii] |
| 04471 | 2 | hypothetical protein [Aromatoleum evansii] |
| 04500 | 2 | PriCT-2 domain-containing protein [Aromatoleum diolicum] |
| 07498 | 2 | hypothetical protein [Azoarcus sp. DD4] |
| 04548 | 2 | hypothetical protein [Aromatoleum toluclasticum] |
| 04543 | 2 | DNA mismatch repair protein MutS [Candidatus Dactylopiibacterium carminicum] |
| 07505 | 2 | lasso peptide biosynthesis protein [Aromatoleum evansii] |
| 04524 | 2 | TIGR01777 family oxidoreductase [Candidatus Dactylopiibacterium carminicum] |
| 04523 | 2 | hypothetical protein [Candidatus Dactylopiibacterium carminicum] |
| 04518 | 2 | MFS transporter [Azoarcus nasutitermitis] |
| 04515 | 2 | TIR domain-containing protein [Zoogloea dura] |

| | | |
|---|---|---|
| 07521 | 2 | hypothetical protein [Denitromonas halophilus] |
| 04499 | 2 | DUF1329 domain-containing protein, partial [Aromatoleum diolicum] |
| 07536 | 2 | type VI secretion system tip protein VgrG [Zoogloea dura] |
| 04495 | 2 | hypothetical protein [Aromatoleum diolicum] |
| 04493 | 2 | NAD(P)-binding domain-containing protein [Aromatoleum toluvorans] |
| 04491 | 2 | hypothetical protein [Aromatoleum toluvorans] |
| 04489 | 2 | ATP-binding protein [Azoarcus taiwanensis] |
| 07528 | 2 | hypothetical protein [Zoogloea oleivorans] |
| 04477 | 2 | hypothetical protein [Azoarcus indigens] |
| 04476 | 2 | chromate resistance protein [Zoogloea oleivorans] |
| 07533 | 2 | hypothetical protein [Zoogloea dura] |
| 04475 | 2 | filamentous hemagglutinin N-terminal domain-containing protein [Zoogloea ramigera] |
| 07590 | 2 | DUF5666 domain-containing protein [Aromatoleum tolulyticum] |
| 04383 | 2 | DUF2975 domain-containing protein [Azospira oryzae] |
| 04381 | 2 | hypothetical protein [Dechlorosoma suillum] |
| 07655 | 2 | hypothetical protein [Aromatoleum aromaticum] |
| 07642 | 2 | hypothetical protein [Nitrogeniibacter aestuarii] |
| 07643 | 2 | hypothetical protein [Nitrogeniibacter aestuarii] |
| 04313 | 2 | metallophosphoesterase [Azoarcus sp. KH32C] |
| 04307 | 2 | hypothetical protein [Thauera phenylacetica] |
| 04305 | 2 | AAA family ATPase, partial [Thauera aminoaromatica] |
| 07648 | 2 | hypothetical protein [Niveibacterium microcysteis] |
| 04293 | 2 | DUF4258 domain-containing protein [Aromatoleum aromaticum] |
| 07653 | 2 | hypothetical protein [Aromatoleum bremense] |
| 04289 | 2 | class II fructose-bisphosphate aldolase, partial [Propionivibrio dicarboxylicus] |
| 07657 | 2 | PadR family transcriptional regulator [Aromatoleum petrolei] |
| 07639 | 2 | HEAT repeat domain-containing protein [Nitrogeniibacter aestuarii] |
| 04283 | 2 | hypothetical protein [Denitromonas sp. IR12] |
| 04281 | 2 | hypothetical protein [Aromatoleum toluclasticum] |
| 04279 | 2 | hypothetical protein [Nitrogeniibacter aestuarii] |
| 07663 | 2 | DUF4240 domain-containing protein [Nitrogeniibacter aestuarii] |
| 07665 | 2 | GNAT family N-acetyltransferase [Nitrogeniibacter aestuarii] |
| 04276 | 2 | PEP-CTERM sorting domain-containing protein [Nitrogeniibacter aestuarii] |
| 04275 | 2 | hypothetical protein [Nitrogeniibacter aestuarii] |
| 04274 | 2 | hypothetical protein [Nitrogeniibacter aestuarii] |
| 07671 | 2 | hypothetical protein [Propionivibrio dicarboxylicus] |
| 07640 | 2 | hypothetical protein [Nitrogeniibacter aestuarii] |
| 07638 | 2 | DUF695 domain-containing protein [Nitrogeniibacter aestuarii] |
| 04378 | 2 | TIGR02221 family CRISPR-associated protein [Thauera hydrothermalis] |
| 04345 | 2 | hypothetical protein [Azoarcus nasutitermitis] |
| 04376 | 2 | helix-turn-helix domain-containing protein [Azoarcus communis] |
| 07601 | 2 | hypothetical protein [Uliginosibacterium aquaticum] |
| 07605 | 2 | metallophosphoesterase [Azoarcus sp. L1K30] |
| 07607 | 2 | hypothetical protein [Niveibacterium microcysteis] |
| 07608 | 2 | hypothetical protein [Niveibacterium sp. COAC-50] |
| 04356 | 2 | hypothetical protein [Thauera butanivorans] |
| 07610 | 2 | hypothetical protein [Niveibacterium microcysteis] |
| 04355 | 2 | LysR substrate-binding domain-containing protein [Thauera linaloolentis] |
| 04352 | 2 | hypothetical protein, partial [Thauera butanivorans] |
| 04342 | 2 | hypothetical protein [Denitromonas halophilus] |
| 04315 | 2 | protein phosphatase 2C domain-containing protein [Aromatoleum aromaticum] |
| 04341 | 2 | MULTISPECIES: hypothetical protein [Thauera] |
| 04340 | 2 | hypothetical protein [Thauera butanivorans] |
| 07621 | 2 | CHASE domain-containing protein [Rhodocyclus purpureus] |
| 04339 | 2 | hypothetical protein [Thauera butanivorans] |
| 04337 | 2 | hypothetical protein [Thauera terpenica] |
| 04331 | 2 | chitosanase [Aromatoleum toluclasticum] |
| 04327 | 2 | hypothetical protein [Uliginosibacterium gangwonense] |
| 07632 | 2 | hypothetical protein [Azoarcus nasutitermitis] |
| 07633 | 2 | hypothetical protein [Niveibacterium microcysteis] |
| 04558 | 2 | DUF2845 domain-containing protein [Niveibacterium microcysteis] |
| 07494 | 2 | hypothetical protein [Azoarcus rhizosphaerae] |
| 04783 | 2 | NYN domain-containing protein [Azoarcus nasutitermitis] |
| 04564 | 2 | hypothetical protein [Nitrogeniibacter aestuarii] |
| 07378 | 2 | hypothetical protein [Azoarcus olearius] |
| 04704 | 2 | hypothetical protein [Thauera lacus] |
| 04702 | 2 | hypothetical protein [Candidatus Dactylopiibacterium carminicum] |
| 07383 | 2 | MULTISPECIES: hypothetical protein [Thauera] |
| 07384 | 2 | tRNA (adenosine(37)-N6)-dimethylallyltransferase MiaA, partial [Thauera aminoaromatica] |
| 07386 | 2 | hypothetical protein [Thauera phenylacetica] |
| 04693 | 2 | hypothetical protein [Azoarcus taiwanensis] |
| 07388 | 2 | MULTISPECIES: hypothetical protein [Thauera] |
| 04691 | 2 | hypothetical protein [Thauera chlorobenzoica] |
| 07390 | 2 | hypothetical protein [Aromatoleum toluclasticum] |
| 04690 | 2 | MULTISPECIES: hypothetical protein [Proteobacteria] |
| 04687 | 2 | YjfB family protein [Thauera aromatica] |
| 04686 | 2 | hypothetical protein [Thauera phenolivorans] |
| 04685 | 2 | MFS transporter [Azoarcus sp. KH32C] |
| 04681 | 2 | ankyrin repeat domain-containing protein [Thauera butanivorans] |
| 04678 | 2 | DUF4279 domain-containing protein [Azoarcus communis] |
| 04676 | 2 | hypothetical protein [Thauera butanivorans] |
| 04673 | 2 | hypothetical protein [Thauera butanivorans] |
| 07404 | 2 | DUF4145 domain-containing protein [Uliginosibacterium gangwonense] |
| 04670 | 2 | hypothetical protein [Aromatoleum tolulyticum] |
| 04669 | 2 | ABC transporter substrate-binding protein [Azoarcus sp. TTM-91] |
| 07376 | 2 | hypothetical protein [Aromatoleum aromaticum] |
| 07375 | 2 | cell wall hydrolase [Aromatoleum aromaticum] |
| 07374 | 2 | hypothetical protein [Aromatoleum aromaticum] |
| 04738 | 2 | 50S ribosome-binding GTPase [Aromatoleum pumilum] |
| 04782 | 2 | MULTISPECIES: hypothetical protein [Betaproteobacteria] |
| 04776 | 2 | VOC family protein [Azospira oryzae] |
| 04775 | 2 | hypothetical protein [Azospira oryzae] |
| 04773 | 2 | deoxyribonuclease V [Dechlorosoma suillum] |
| 07335 | 2 | hypothetical protein [Nitrogeniibacter aestuarii] |

| | | |
|---|---|---|
| 04757 | 2 | HPF/RaiA family ribosome-associated protein [Azoarcus communis] |
| 04754 | 2 | type II toxin-antitoxin system VapC family toxin [Denitromonas sp. IR12] |
| 04752 | 2 | transposase [Aromatoleum toluolicum] |
| 04746 | 2 | class I SAM-dependent methyltransferase [Aromatoleum pumilum] |
| 07348 | 2 | hypothetical protein [Aromatoleum bremense] |
| 07371 | 2 | hypothetical protein [Aromatoleum tolulyticum] |
| 07350 | 2 | DUF1697 domain-containing protein [Azoarcus sp. L1K30] |
| 04734 | 2 | hypothetical protein [Azoarcus nasutitermitis] |
| 07358 | 2 | recombinase family protein [Aromatoleum buckelii] |
| 07359 | 2 | ATP synthase subunit C [Aromatoleum toluclasticum] |
| 04719 | 2 | DUF1684 domain-containing protein [Azospira oryzae] |
| 04718 | 2 | SCO family protein [Azospira oryzae] |
| 04715 | 2 | DUF5710 domain-containing protein [Thauera hydrothermalis] |
| 04712 | 2 | RHS repeat-associated core domain-containing protein, partial [Azoarcus communis] |
| 04710 | 2 | hypothetical protein [Azoarcus communis] |
| 04668 | 2 | serine protease [Thauera humireducens] |
| 04667 | 2 | hypothetical protein [Thauera humireducens] |
| 07410 | 2 | HP1 family phage holin [Uliginosibacterium gangwonense] |
| 04592 | 2 | AMP-binding protein, partial [Denitromonas sp. IR12] |
| 04609 | 2 | hypothetical protein [Aromatoleum buckelii] |
| 07463 | 2 | TlpA family protein disulfide reductase [Azoarcus communis] |
| 04600 | 2 | hypothetical protein [Nitrogeniibacter aestuarii] |
| 07465 | 2 | cellulose biosynthesis cyclic di-GMP-binding regulatory protein BcsB [Aromatoleum bremense] |
| 04599 | 2 | hypothetical protein [Nitrogeniibacter aestuarii] |
| 07468 | 2 | GNAT family N-acetyltransferase [Thauera hydrothermalis] |
| 04596 | 2 | TPM domain-containing protein [Nitrogeniibacter aestuarii] |
| 04595 | 2 | roadblock/LC7 domain-containing protein [Nitrogeniibacter aestuarii] |
| 04594 | 2 | fatty acid desaturase [Nitrogeniibacter aestuarii] |
| 04587 | 2 | type IV secretion system protein VirB10 [Denitromonas sp. IR12] |
| 04612 | 2 | tripartite tricarboxylate transporter TctB family protein [Aromatoleum pumilum] |
| 04579 | 2 | site-specific integrase [Uliginosibacterium gangwonense] |
| 07479 | 2 | hypothetical protein [Azospira oryzae] |
| 07480 | 2 | Gfo/Idh/MocA family oxidoreductase [Azospira oryzae] |
| 04578 | 2 | ATP-binding protein [Aromatoleum evansii] |
| 04571 | 2 | cation transporting ATPase C-terminal domain-containing protein [Azoarcus sp. TTM-91] |
| 04568 | 2 | hypothetical protein [Niveibacterium sp. COAC-50] |
| 07490 | 2 | type II toxin-antitoxin system prevent-host-death family antitoxin [Azoarcus nasutitermitis] |
| 04567 | 2 | 2OG-Fe(II) oxygenase [Denitromonas sp. IR12] |
| 04566 | 2 | DUF4157 domain-containing protein [Nitrogeniibacter aestuarii] |
| 04610 | 2 | citrate lyase holo-[acyl-carrier protein] synthase [Propionivibrio dicarboxylicus] |
| 07454 | 2 | class I SAM-dependent methyltransferase [Aromatoleum pumilum] |
| 04665 | 2 | endoribonuclease L-PSP [Thauera humireducens] |
| 07428 | 2 | transposase, partial [Denitromonas halophilus] |
| 04663 | 2 | toxin-activating lysine-acyltransferase [Propionivibrio dicarboxylicus] |
| 04660 | 2 | hypothetical protein [Azoarcus halotolerans] |
| 04658 | 2 | IS5/IS1182 family transposase [Uliginosibacterium gangwonense] |
| 04657 | 2 | hypothetical protein, partial [Aromatoleum toluclasticum] |
| 07420 | 2 | gamma-glutamylcyclotransferase [Aromatoleum toluclasticum] |
| 04656 | 2 | hypothetical protein [Aromatoleum toluclasticum] |
| 04655 | 2 | SDR family NAD(P)-dependent oxidoreductase [Aromatoleum toluclasticum] |
| 07426 | 2 | TIGR03084 family metal-binding protein [Azoarcus sp. KH32C] |
| 07427 | 2 | hypothetical protein [Thauera linaloolentis] |
| 07429 | 2 | hypothetical protein [Thauera humireducens] |
| 07450 | 2 | hypothetical protein [Denitromonas sp. IR12] |
| 07432 | 2 | ImmA/IrrE family metallo-endopeptidase [Azoarcus nasutitermitis] |
| 07434 | 2 | MULTISPECIES: type II toxin-antitoxin system HicA family toxin [Aromatoleum] |
| 04635 | 2 | DUF1837 domain-containing protein [Thauera terpenica] |
| 04627 | 2 | hypothetical protein [Niveibacterium umoris] |
| 04626 | 2 | bifunctional protein-serine/threonine kinase/phosphatase [Thauera phenolivorans] |
| 04621 | 2 | hypothetical protein [Azoarcus olearius] |
| 04620 | 2 | hypothetical protein [Aromatoleum aromaticum] |
| 04618 | 2 | integrase [Aromatoleum aromaticum] |
| 07449 | 2 | hypothetical protein [Aromatoleum pumilum] |
| 05279 | 2 | C1 family peptidase [Niveibacterium microcysteis] |
| 06962 | 2 | DEAD/DEAH box helicase family protein [Aromatoleum evansii] |
| 05280 | 2 | hypothetical protein [Niveibacterium microcysteis] |
| 06033 | 2 | MULTISPECIES: hypothetical protein [Azospira] |
| 06490 | 2 | hypothetical protein [Aromatoleum diolicum] |
| 06491 | 2 | DUF2917 domain-containing protein [Aromatoleum diolicum] |
| 05975 | 2 | L-ascorbate 6-phosphate lactonase [Propionivibrio dicarboxylicus] |
| 05971 | 2 | kinase [Aromatoleum aromaticum] |
| 05968 | 2 | hypothetical protein [Azoarcus communis] |
| 05966 | 2 | hypothetical protein [Aromatoleum diolicum] |
| 06502 | 2 | fimbria/pilus outer membrane usher protein [Azoarcus sp. KH32C] |
| 05961 | 2 | AAA family ATPase [Azoarcus rhizosphaerae] |
| 06507 | 2 | terminase small subunit [Azoarcus nasutitermitis] |
| 05960 | 2 | AMP-binding protein, partial [Thauera phenylacetica] |
| 05959 | 2 | GTP-binding protein, partial [Aromatoleum buckelii] |
| 05955 | 2 | MULTISPECIES: hypothetical protein [Thauera] |
| 05950 | 2 | hypothetical protein [Aromatoleum aromaticum] |
| 05949 | 2 | type II toxin-antitoxin system MqsA family antitoxin [Aromatoleum aromaticum] |
| 06515 | 2 | MULTISPECIES: hypothetical protein [Uliginosibacterium] |
| 05946 | 2 | efflux transporter outer membrane subunit [Propionivibrio dicarboxylicus] |
| 06519 | 2 | hypothetical protein [Uliginosibacterium aquaticum] |
| 05941 | 2 | LysR family transcriptional regulator, partial [Aromatoleum toluclasticum] |
| 05940 | 2 | 2-oxoacid:acceptor oxidoreductase family protein, partial [Aromatoleum toluclasticum] |
| 05939 | 2 | LuxR C-terminal-related transcriptional regulator, partial [Aromatoleum toluclasticum] |
| 05935 | 2 | DNA (cytosine-5-)-methyltransferase [Aromatoleum toluclasticum] |
| 05987 | 2 | PEP-CTERM sorting domain-containing protein [Azoarcus sp. L1K30] |
| 05990 | 2 | hypothetical protein [Denitromonas halophilus] |
| 06486 | 2 | ester cyclase [Aromatoleum toluvorans] |
| 06008 | 2 | methyl-accepting chemotaxis protein [Thauera aromatica] |
| 06019 | 2 | hypothetical protein [Thauera aromatica] |
| 06017 | 2 | hypothetical protein [Aromatoleum pumilum] |

| | | |
|---|---|---|
| 06015 | 2 | diguanylate cyclase [Uliginosibacterium aquaticum] |
| 06460 | 2 | hypothetical protein [Aromatoleum aromaticum] |
| 06461 | 2 | HTH domain-containing protein, partial [Aromatoleum buckelii] |
| 06462 | 2 | MULTISPECIES: hypothetical protein, partial [Zoogloeaceae] |
| 06012 | 2 | NERD domain-containing protein [Thauera phenolivorans] |
| 06464 | 2 | hypothetical protein [Aromatoleum evansii] |
| 06011 | 2 | hypothetical protein [Azoarcus communis] |
| 06468 | 2 | cation transporter, partial [Thauera propionica] |
| 06484 | 2 | hypothetical protein, partial [Azoarcus taiwanensis] |
| 06006 | 2 | glycosyltransferase [Thauera phenolivorans] |
| 06470 | 2 | hypothetical protein [Azoarcus communis] |
| 06473 | 2 | hypothetical protein [Azoarcus communis] |
| 06001 | 2 | hypothetical protein, partial [Thauera butanivorans] |
| 06475 | 2 | aldehyde dehydrogenase family protein, partial [Azoarcus indigens] |
| 06000 | 2 | hypothetical protein [Aromatoleum tolulyticum] |
| 05996 | 2 | hypothetical protein [Azoarcus sp. DD4] |
| 06480 | 2 | Hpt domain-containing protein [Azoarcus taiwanensis] |
| 05992 | 2 | hypothetical protein [Thauera butanivorans] |
| 05933 | 2 | hypothetical protein [Nitrogeniibacter aestuarii] |
| 05932 | 2 | DUF3421 domain-containing protein [Nitrogeniibacter aestuarii] |
| 05930 | 2 | DUF1127 domain-containing protein [Nitrogeniibacter aestuarii] |
| 05876 | 2 | hypothetical protein [Niveibacterium umoris] |
| 05894 | 2 | hypothetical protein [Nitrogeniibacter aestuarii] |
| 06568 | 2 | CGNR zinc finger domain-containing protein [Nitrogeniibacter aestuarii] |
| 05892 | 2 | AHH domain-containing protein [Denitromonas sp. IR12] |
| 06571 | 2 | hypothetical protein [Nitrogeniibacter aestuarii] |
| 06573 | 2 | PEP-CTERM sorting domain-containing protein [Nitrogeniibacter aestuarii] |
| 06575 | 2 | GNAT family N-acetyltransferase [Niveibacterium microcysteis] |
| 05885 | 2 | chitobiase/beta-hexosaminidase C-terminal domain-containing protein [Niveibacterium microcysteis] |
| 06581 | 2 | hypothetical protein [Niveibacterium umoris] |
| 06582 | 2 | PEP-CTERM sorting domain-containing protein [Niveibacterium microcysteis] |
| 06585 | 2 | hypothetical protein [Aromatoleum bremense] |
| 06562 | 2 | hypothetical protein [Niveibacterium microcysteis] |
| 05875 | 2 | apolipoprotein A1/A4/E domain-containing protein [Azospira oryzae] |
| 06588 | 2 | hypothetical protein [Azoarcus sp. L1K30] |
| 06589 | 2 | glycosyltransferase [Aromatoleum pumilum] |
| 05870 | 2 | hypothetical protein [Azoarcus sp. Aa7] |
| 06592 | 2 | hypothetical protein [Azoarcus rhizosphaerae] |
| 05861 | 2 | hypothetical protein [Candidatus Dactylopiibacterium carminicum] |
| 06594 | 2 | zeta toxin family protein [Denitromonas sp. IR12] |
| 06598 | 2 | hypothetical protein [Nitrogeniibacter aestuarii] |
| 06601 | 2 | hypothetical protein [Nitrogeniibacter aestuarii] |
| 06566 | 2 | DUF3857 domain-containing protein [Niveibacterium sp. COAC-50] |
| 05899 | 2 | hypothetical protein [Denitromonas sp. IR12] |
| 05929 | 2 | hypothetical protein [Nitrogeniibacter aestuarii] |
| 06542 | 2 | MULTISPECIES: hypothetical protein [Uliginosibacterium] |
| 06532 | 2 | HAD family hydrolase [Aromatoleum evansii] |
| 05927 | 2 | type VI secretion system Vgr family protein, partial [Aromatoleum toluclasticum] |
| 05926 | 2 | integrase core domain-containing protein, partial [Aromatoleum evansii] |
| 06535 | 2 | hypothetical protein [Niveibacterium microcysteis] |
| 06537 | 2 | DUF1003 domain-containing protein [Niveibacterium microcysteis] |
| 06538 | 2 | adenylate synthase [Candidatus Dactylopiibacterium carminicum] |
| 05923 | 2 | hypothetical protein [Denitromonas sp. IR12] |
| 05921 | 2 | PEP-CTERM sorting domain-containing protein [Denitromonas halophilus] |
| 06541 | 2 | hypothetical protein [Candidatus Dactylopiibacterium carminicum] |
| 05920 | 2 | helix-turn-helix domain-containing protein, partial [Denitromonas sp. IR12] |
| 06556 | 2 | hypothetical protein [Niveibacterium microcysteis] |
| 05918 | 2 | hypothetical protein [Denitromonas sp. IR12] |
| 05916 | 2 | hypothetical protein [Azoarcus communis] |
| 05914 | 2 | hypothetical protein [Azoarcus communis] |
| 05910 | 2 | hypothetical protein [Aromatoleum petrolei] |
| 05909 | 2 | hypothetical protein [Aromatoleum petrolei] |
| 05908 | 2 | hypothetical protein [Aromatoleum petrolei] |
| 05905 | 2 | hypothetical protein [Aromatoleum bremense] |
| 06553 | 2 | hypothetical protein [Rhodocyclus purpureus] |
| 06554 | 2 | hypothetical protein [Rhodocyclus purpureus] |
| 06449 | 2 | DNA primase [Azoarcus halotolerans] |
| 06447 | 2 | hypothetical protein [Azoarcus halotolerans] |
| 06603 | 2 | hypothetical protein [Nitrogeniibacter aestuarii] |
| 06037 | 2 | antA/AntB antirepressor family protein [Azoarcus olearius] |
| 06204 | 2 | hypothetical protein [Nitrogeniibacter aestuarii] |
| 06316 | 2 | hypothetical protein [Thauera aromatica] |
| 06201 | 2 | SEL1-like repeat protein [Nitrogeniibacter aestuarii] |
| 06320 | 2 | type B 50S ribosomal protein L36 [Thauera butanivorans] |
| 06322 | 2 | nuclear transport factor 2 family protein [Thauera butanivorans] |
| 06325 | 2 | ankyrin repeat domain-containing protein [Thauera butanivorans] |
| 06327 | 2 | IS110 family transposase, partial [Thauera butanivorans] |
| 06191 | 2 | hypothetical protein [Niveibacterium microcysteis] |
| 06190 | 2 | hypothetical protein [Niveibacterium microcysteis] |
| 06333 | 2 | sulfatase-like hydrolase/transferase [Aromatoleum bremense] |
| 06335 | 2 | DUF58 domain-containing protein [Niveibacterium umoris] |
| 06336 | 2 | XdhC family protein [Aromatoleum toluclasticum] |
| 06186 | 2 | heme-binding protein [Rhodocyclus purpureus] |
| 06338 | 2 | 3-hydroxyacyl-CoA dehydrogenase family protein [Thauera chlorobenzoica] |
| 06339 | 2 | MULTISPECIES: toll/interleukin-1 receptor domain-containing protein [Thauera] |
| 06340 | 2 | hypothetical protein [Denitromonas halophilus] |
| 06185 | 2 | UbiA family prenyltransferase, partial [Rhodocyclus purpureus] |
| 06182 | 2 | allantoin permease [Azoarcus sp. Aa7] |
| 06344 | 2 | hypothetical protein [Aromatoleum pumilum] |
| 06179 | 2 | hypothetical protein [Azoarcus sp. Aa7] |
| 06177 | 2 | hypothetical protein [Candidatus Dactylopiibacterium carminicum] |
| 06211 | 2 | hypothetical protein [Niveibacterium microcysteis] |
| 06307 | 2 | Ig-like domain-containing protein, partial [Thauera terpenica] |
| 06221 | 2 | hypothetical protein [Nitrogeniibacter aestuarii] |

| | | |
|---|---|---|
| 06252 | 2 | hypothetical protein [Aromatoleum toluclasticum] |
| 06273 | 2 | hypothetical protein [Azoarcus olearius] |
| 06262 | 2 | hypothetical protein [Aromatoleum tolulyticum] |
| 06275 | 2 | hypothetical protein [Candidatus Dactylopiibacterium carminicum] |
| 06276 | 2 | DUF3486 family protein [Thauera phenylacetica] |
| 06259 | 2 | dihydroxyacetone kinase subunit DhaL [Propionivibrio dicarboxylicus] |
| 06258 | 2 | cupin domain-containing protein [Propionivibrio dicarboxylicus] |
| 06281 | 2 | prepilin-type N-terminal cleavage/methylation domain-containing protein, partial [Thauera aminoaromatica] |
| 06283 | 2 | MULTISPECIES: hypothetical protein [Proteobacteria] |
| 06254 | 2 | GAF domain-containing protein [Propionivibrio dicarboxylicus] |
| 06249 | 2 | ankyrin repeat domain-containing protein [Nitrogeniibacter aestuarii] |
| 06305 | 2 | O-antigen ligase family protein [Thauera phenylacetica] |
| 06289 | 2 | HNH endonuclease [Azoarcus sp. KH32C] |
| 06247 | 2 | hypothetical protein [Nitrogeniibacter aestuarii] |
| 06246 | 2 | hypothetical protein [Nitrogeniibacter aestuarii] |
| 06244 | 2 | hypothetical protein [Nitrogeniibacter aestuarii] |
| 06241 | 2 | sigma-70 family RNA polymerase sigma factor [Aromatoleum buckelii] |
| 06238 | 2 | glutaredoxin family protein [Thauera humireducens] |
| 06297 | 2 | dienelactone hydrolase family protein [Uliginosibacterium gangwonense] |
| 06223 | 2 | barstar family protein [Nitrogeniibacter aestuarii] |
| 06222 | 2 | hypothetical protein [Nitrogeniibacter aestuarii] |
| 06176 | 2 | hypothetical protein [Aromatoleum toluvorans] |
| 06349 | 2 | hypothetical protein [Thauera aromatica] |
| 06175 | 2 | LapA family protein [Zoogloea ramigera] |
| 06073 | 2 | hypothetical protein [Aromatoleum evansii] |
| 06092 | 2 | DUF2244 domain-containing protein [Zoogloea dura] |
| 06091 | 2 | hypothetical protein [Thauera propionica] |
| 06090 | 2 | hypothetical protein [Zoogloea oleivorans] |
| 06088 | 2 | MAE_28990/MAE_18760 family HEPN-like nuclease [Zoogloea oleivorans] |
| 06086 | 2 | polysaccharide deacetylase family protein [Zoogloea oleivorans] |
| 06083 | 2 | helix-turn-helix domain-containing protein [Azoarcus nasutitermitis] |
| 06410 | 2 | hypothetical protein [Azoarcus communis] |
| 06081 | 2 | PEP-CTERM sorting domain-containing protein [Thauera aminoaromatica] |
| 06412 | 2 | hypothetical protein [Zoogloea ramigera] |
| 06423 | 2 | acyloxyacyl hydrolase [Denitromonas halophilus] |
| 06102 | 2 | FHA domain-containing protein [Uliginosibacterium gangwonense] |
| 06067 | 2 | linear amide C-N hydrolase [Thauera butanivorans] |
| 06066 | 2 | ankyrin repeat domain-containing protein [Azoarcus sp. Aa7] |
| 06064 | 2 | PIN domain-containing protein [Zoogloea oleivorans] |
| 06059 | 2 | GtrA family protein [Aromatoleum petrolei] |
| 06053 | 2 | hypothetical protein [Azoarcus rhizosphaerae] |
| 06436 | 2 | hypothetical protein [Azoarcus taiwanensis] |
| 06437 | 2 | MULTISPECIES: hypothetical protein [Azospira] |
| 06043 | 2 | hypothetical protein [Azoarcus nasutitermitis] |
| 06042 | 2 | DUF637 domain-containing protein, partial [Azoarcus indigens] |
| 06401 | 2 | AAA family ATPase [Azoarcus olearius] |
| 06396 | 2 | hypothetical protein [Azoarcus rhizosphaerae] |
| 06174 | 2 | STAS domain-containing protein [Candidatus Dactylopiibacterium carminicum] |
| 06369 | 2 | GNAT family N-acetyltransferase [Oryzomicrobium terrae] |
| 06171 | 2 | fused MFS/spermidine synthase [Azoarcus sp. L1K30] |
| 06167 | 2 | DUF4268 domain-containing protein [Niveibacterium sp. COAC-50] |
| 06166 | 2 | hypothetical protein [Niveibacterium sp. COAC-50] |
| 06161 | 2 | hypothetical protein [Niveibacterium microcysteis] |
| 06362 | 2 | hypothetical protein [Azoarcus communis] |
| 06153 | 2 | DUF2158 domain-containing protein [Azoarcus rhizosphaerae] |
| 06364 | 2 | TetR/AcrR family transcriptional regulator [Azoarcus nasutitermitis] |
| 06367 | 2 | hypothetical protein [Thauera hydrothermalis] |
| 06147 | 2 | hypothetical protein [Zoogloea dura] |
| 06146 | 2 | chemotaxis protein CheW [Thauera terpenica] |
| 06108 | 2 | hypothetical protein [Azoarcus halotolerans] |
| 06139 | 2 | hypothetical protein [Aromatoleum tolulyticum] |
| 06135 | 2 | NAD(P)-binding protein, partial [Aromatoleum diolicum] |
| 06132 | 2 | PEP-CTERM sorting domain-containing protein [Aromatoleum diolicum] |
| 06121 | 2 | TolC family protein, partial [Azoarcus indigens] |
| 06388 | 2 | hypothetical protein [Azoarcus nasutitermitis] |
| 06389 | 2 | hypothetical protein [Azoarcus communis] |
| 06112 | 2 | hypothetical protein [Aromatoleum buckelii] |
| 06392 | 2 | hypothetical protein [Azoarcus communis] |
| 06109 | 2 | outer membrane beta-barrel protein [Azoarcus sp. DD4] |
| 06602 | 2 | DUF2314 domain-containing protein [Nitrogeniibacter aestuarii] |
| 05854 | 2 | hypothetical protein [Niveibacterium microcysteis] |
| 06957 | 2 | helix-turn-helix transcriptional regulator [Aromatoleum diolicum] |
| 05581 | 2 | hypothetical protein [Candidatus Dactylopiibacterium carminicum] |
| 05479 | 2 | hypothetical protein [Zoogloea ramigera] |
| 05477 | 2 | ankyrin repeat domain-containing protein [Thauera lacus] |
| 05476 | 2 | class I SAM-dependent methyltransferase [Zoogloea dura] |
| 05475 | 2 | hypothetical protein [Azoarcus sp. KH32C] |
| 05472 | 2 | hypothetical protein [Azoarcus olearius] |
| 05468 | 2 | hypothetical protein [Azoarcus rhizosphaerae] |
| 05467 | 2 | NYN domain-containing protein [Azoarcus rhizosphaerae] |
| 06861 | 2 | hypothetical protein [Thauera phenolivorans] |
| 05454 | 2 | TonB-dependent receptor [Azoarcus indigens] |
| 05451 | 2 | DUF4129 domain-containing protein [Azospira oryzae] |
| 06864 | 2 | hypothetical protein [Thauera aromatica] |
| 05446 | 2 | hypothetical protein [Azospira oryzae] |
| 06866 | 2 | hypothetical protein [Thauera propionica] |
| 05444 | 2 | hypothetical protein [Thauera hydrothermalis] |
| 06868 | 2 | dynamin family protein [Azoarcus halotolerans] |
| 05440 | 2 | transposase, partial [Azoarcus communis] |
| 05438 | 2 | FAD-dependent oxidoreductase, partial [Azoarcus communis] |
| 05428 | 2 | MULTISPECIES: hypothetical protein [Bacteria] |
| 05426 | 2 | hypothetical protein [Aromatoleum pumilum] |
| 05419 | 2 | MULTISPECIES: hypothetical protein [Proteobacteria] |
| 06882 | 2 | hypothetical protein [Azoarcus nasutitermitis] |

| | | |
|---|---|---|
| 05495 | 2 | DUF4347 domain-containing protein, partial [Denitromonas sp. IR12] |
| 05496 | 2 | carboxymuconolactone decarboxylase family protein [Uliginosibacterium aquaticum] |
| 05500 | 2 | tetratricopeptide repeat protein [Zoogloea ramigera] |
| 05542 | 2 | helix-turn-helix domain-containing protein [Aromatoleum diolicum] |
| 05575 | 2 | hypothetical protein [Niveibacterium sp. COAC-50] |
| 06802 | 2 | hypothetical protein [Thauera aminoaromatica] |
| 05568 | 2 | LysM peptidoglycan-binding domain-containing protein [Niveibacterium sp. COAC-50] |
| 05562 | 2 | BRCT domain-containing protein [Denitromonas sp. IR12] |
| 06808 | 2 | hypothetical protein [Thauera aminoaromatica] |
| 05556 | 2 | peptidoglycan DD-metalloendopeptidase family protein [Zoogloea dura] |
| 06812 | 2 | PilZ domain-containing protein [Aromatoleum toluolicum] |
| 05555 | 2 | transposase, partial [Zoogloea dura] |
| 05545 | 2 | hypothetical protein [Aromatoleum toluolicum] |
| 05541 | 2 | HNH endonuclease [Aromatoleum diolicum] |
| 05515 | 2 | hypothetical protein [Azoarcus communis] |
| 05535 | 2 | helix-turn-helix domain containing protein, partial [Aromatoleum toluvorans] |
| 06824 | 2 | ribonuclease E inhibitor RraB [Niveibacterium umoris] |
| 05530 | 2 | MULTISPECIES: hypothetical protein [Proteobacteria] |
| 06829 | 2 | hypothetical protein [Uliginosibacterium aquaticum] |
| 05527 | 2 | hypothetical protein [Azoarcus indigens] |
| 05522 | 2 | thioredoxin family protein [Aromatoleum evansii] |
| 05520 | 2 | hypothetical protein [Aromatoleum evansii] |
| 05517 | 2 | phosphoadenosine phosphosulfate reductase family protein [Aromatoleum aromaticum] |
| 06837 | 2 | hypothetical protein [Aromatoleum toluclasticum] |
| 05415 | 2 | hypothetical protein [Azoarcus sp. Aa7] |
| 05413 | 2 | hypothetical protein [Thauera hydrothermalis] |
| 05410 | 2 | hypothetical protein [Thauera phenolivorans] |
| 06933 | 2 | hypothetical protein [Aromatoleum evansii] |
| 06918 | 2 | hypothetical protein [Aromatoleum diolicum] |
| 05338 | 2 | glycosyl transferase [Niveibacterium umoris] |
| 05331 | 2 | hypothetical protein [Propionivibrio dicarboxylicus] |
| 06923 | 2 | LPP20 family lipoprotein [Zoogloea oleivorans] |
| 05326 | 2 | methylaspartate ammonia-lyase [Aromatoleum toluvorans] |
| 05321 | 2 | biotin/lipoyl-binding protein, partial [Aromatoleum toluclasticum] |
| 05320 | 2 | hypothetical protein [Aromatoleum toluclasticum] |
| 05318 | 2 | hypothetical protein [Aromatoleum toluclasticum] |
| 06931 | 2 | hypothetical protein [Denitromonas halophilus] |
| 06936 | 2 | hypothetical protein [Zoogloea dura] |
| 06916 | 2 | hypothetical protein [Azoarcus olearius] |
| 06937 | 2 | hypothetical protein [Aromatoleum toluvorans] |
| 05309 | 2 | hypothetical protein [Denitromonas sp. IR12] |
| 06940 | 2 | hypothetical protein [Zoogloea oleivorans] |
| 05303 | 2 | DMT family transporter [Aromatoleum petrolei] |
| 05295 | 2 | hypothetical protein [Aromatoleum bremense] |
| 06946 | 2 | ParB N-terminal domain-containing protein [Candidatus Dactylopiibacterium carminicum] |
| 05292 | 2 | GspMb/PilO family protein [Niveibacterium umoris] |
| 06951 | 2 | hypothetical protein [Azoarcus halotolerans] |
| 05285 | 2 | HNH endonuclease [Niveibacterium microcysteis] |
| 05346 | 2 | MULTISPECIES: hypothetical protein [Thauera] |
| 05348 | 2 | S8 family serine peptidase [Thauera aminoaromatica] |
| 05405 | 2 | right-handed parallel beta-helix repeat-containing protein, partial [Thauera butanivorans] |
| 05392 | 2 | hypothetical protein [Azoarcus communis] |
| 06888 | 2 | hypothetical protein [Azoarcus nasutitermitis] |
| 06889 | 2 | hypothetical protein [Aromatoleum toluolicum] |
| 06892 | 2 | hypothetical protein [Azoarcus communis] |
| 05397 | 2 | hypothetical protein [Aromatoleum pumilum] |
| 06894 | 2 | type I restriction-modification system subunit M N-terminal domain-containing protein [Azoarcus communis] |
| 05395 | 2 | hypothetical protein [Azoarcus sp. DD4] |
| 05393 | 2 | hypothetical protein [Thauera linaloolentis] |
| 06897 | 2 | Gldg family protein [Nitrogeniibacter mangrovi] |
| 06898 | 2 | DUF6511 domain-containing protein [Candidatus Dactylopiibacterium carminicum] |
| 05387 | 2 | hypothetical protein [Aromatoleum bremense] |
| 06913 | 2 | hypothetical protein [Azoarcus rhizosphaerae] |
| 05372 | 2 | DUF1269 domain-containing protein [Azoarcus halotolerans] |
| 05371 | 2 | hypothetical protein [Thauera propionica] |
| 05367 | 2 | hypothetical protein [Uliginosibacterium aquaticum] |
| 06905 | 2 | NUMOD4 motif-containing HNH endonuclease [Azoarcus nasutitermitis] |
| 05366 | 2 | hypothetical protein [Aromatoleum buckelii] |
| 05356 | 2 | DUF4276 family protein [Thauera phenolivorans] |
| 06910 | 2 | pyridoxamine 5'-phosphate oxidase family protein [Azoarcus nasutitermitis] |
| 05351 | 2 | response regulator, partial [Thauera aminoaromatica] |
| 05350 | 2 | hypothetical protein, partial [Thauera aminoaromatica] |
| 05576 | 2 | flippase-like domain-containing protein [Zoogloea dura] |
| 05585 | 2 | type II toxin-antitoxin system VapC family toxin [Aromatoleum pumilum] |
| 06610 | 2 | hypothetical protein [Propionivibrio dicarboxylicus] |
| 05587 | 2 | DUF5801 domain-containing protein, partial [Azoarcus sp. Aa7] |
| 05800 | 2 | DUF2513 domain-containing protein [Azoarcus communis] |
| 06655 | 2 | MBL fold metallo-hydrolase [Candidatus Dactylopiibacterium carminicum] |
| 06656 | 2 | class IV adenylate cyclase [Propionivibrio dicarboxylicus] |
| 06657 | 2 | hypothetical protein [Uliginosibacterium gangwonense] |
| 05798 | 2 | LuxR C-terminal-related transcriptional regulator [Aromatoleum evansii] |
| 06660 | 2 | hypothetical protein [Denitromonas sp. IR12] |
| 05796 | 2 | hypothetical protein [Aromatoleum petrolei] |
| 06664 | 2 | hypothetical protein [Oryzomicrobium terrae] |
| 06665 | 2 | hypothetical protein [Azoarcus communis] |
| 06666 | 2 | phosphoribosyltransferase domain-containing protein [Aromatoleum aromaticum] |
| 05793 | 2 | DNA-binding protein [Aromatoleum evansii] |
| 05789 | 2 | hypothetical protein [Aromatoleum aromaticum] |
| 06670 | 2 | hypothetical protein [Aromatoleum petrolei] |
| 06672 | 2 | hypothetical protein [Azoarcus rhizosphaerae] |
| 05783 | 2 | thiamine pyrophosphate-dependent enzyme [Thauera terpenica] |
| 05779 | 2 | hypothetical protein [Azoarcus halotolerans] |
| 05771 | 2 | hypothetical protein [Oryzomicrobium terrae] |
| 05770 | 2 | VOC family protein [Oryzomicrobium terrae] |

| | | |
|---|---|---|
| 06681 | 2 | phosphatidylglycerol lysyltransferase domain-containing protein [Azoarcus sp. KH32C] |
| 06686 | 2 | hypothetical protein [Azoarcus sp. L1K30] |
| 05765 | 2 | hypothetical protein [Zoogloea oleivorans] |
| 06653 | 2 | SRPBCC domain-containing protein [Aromatoleum tolulyticum] |
| 05801 | 2 | hypothetical protein [Azoarcus communis] |
| 06651 | 2 | hypothetical protein [Aromatoleum bremense] |
| 05824 | 2 | hypothetical protein [Aromatoleum toluolicum] |
| 05848 | 2 | Ig-like domain-containing protein, partial [Uliginosibacterium aquaticum] |
| 05847 | 2 | DMT family transporter [Candidatus Dactylopiibacterium carminicum] |
| 06614 | 2 | hypothetical protein [Aromatoleum toluclasticum] |
| 05843 | 2 | hypothetical protein [Uliginosibacterium aquaticum] |
| 05842 | 2 | FKBP-type peptidyl-prolyl cis-trans isomerase [Candidatus Dactylopiibacterium carminicum] |
| 05840 | 2 | ester cyclase [Aromatoleum toluclasticum] |
| 05835 | 2 | midcut-by-XrtH protein [Zoogloea dura] |
| 05832 | 2 | pyridoxamine 5'-phosphate oxidase family protein [Zoogloea dura] |
| 05825 | 2 | class I SAM-dependent methyltransferase [Azoarcus sp. TTM-91] |
| 06637 | 2 | type I restriction-modification system subunit M N-terminal domain-containing protein [Thauera sp. SWB20] |
| 05802 | 2 | hypothetical protein [Azoarcus indigens] |
| 06638 | 2 | MULTISPECIES: hypothetical protein [Thauera] |
| 05820 | 2 | hypothetical protein [Aromatoleum diolicum] |
| 06640 | 2 | hypothetical protein [Thauera phenolivorans] |
| 06641 | 2 | DUF3102 domain-containing protein [Thauera aminoaromatica] |
| 05816 | 2 | hypothetical protein, partial [Aromatoleum toluolicum] |
| 05813 | 2 | FHIPEP family type III secretion protein, partial [Azoarcus taiwanensis] |
| 05810 | 2 | FAD-binding oxidoreductase [Aromatoleum diolicum] |
| 05807 | 2 | hypothetical protein [Azoarcus sp. Aa7] |
| 05803 | 2 | aminotransferase class III-fold pyridoxal phosphate-dependent enzyme, partial [Azoarcus indigens] |
| 05755 | 2 | DUF2190 family protein [Denitromonas halophilus] |
| 05754 | 2 | hypothetical protein [Azoarcus sp. Aa7] |
| 05747 | 2 | NACHT domain-containing protein [Denitromonas sp. IR12] |
| 06768 | 2 | glycosyltransferase [Azoarcus sp. L1K30] |
| 06744 | 2 | hypothetical protein [Nitrogeniibacter aestuarii] |
| 05660 | 2 | hypothetical protein [Aromatoleum aromaticum] |
| 05659 | 2 | hypothetical protein [Aromatoleum aromaticum] |
| 05650 | 2 | hypothetical protein [Aromatoleum tolulyticum] |
| 06755 | 2 | cytidine deaminase [Aromatoleum bremense] |
| 06756 | 2 | hypothetical protein [Aromatoleum bremense] |
| 06758 | 2 | hypothetical protein [Aromatoleum bremense] |
| 06762 | 2 | hypothetical protein [Aromatoleum petrolei] |
| 06766 | 2 | hypothetical protein [Azoarcus communis] |
| 05603 | 2 | toprim domain-containing protein [Azoarcus communis] |
| 05665 | 2 | hypothetical protein [Azoarcus olearius] |
| 06774 | 2 | hypothetical protein [Nitrogeniibacter aestuarii] |
| 06775 | 2 | DUF4157 domain-containing protein [Nitrogeniibacter aestuarii] |
| 06776 | 2 | DUF1566 domain-containing protein [Nitrogeniibacter aestuarii] |
| 06777 | 2 | hypothetical protein [Nitrogeniibacter aestuarii] |
| 06779 | 2 | DUF1508 domain-containing protein [Aromatoleum toluclasticum] |
| 06782 | 2 | hypothetical protein [Denitromonas sp. IR12] |
| 05595 | 2 | DUF3426 domain-containing protein [Azoarcus nasutitermitis] |
| 05592 | 2 | outer membrane beta-barrel protein [Aromatoleum pumilum] |
| 05588 | 2 | prepilin-type N-terminal cleavage/methylation domain-containing protein [Rhodocyclus purpureus] |
| 05662 | 2 | STY4851/ECs_5259 family protein [Azoarcus olearius] |
| 05667 | 2 | aminotransferase class IV, partial [Azoarcus taiwanensis] |
| 05746 | 2 | hypothetical protein [Zoogloea oleivorans] |
| 05717 | 2 | hypothetical protein [Azoarcus communis] |
| 05735 | 2 | transglycosylase SLT domain-containing protein [Azoarcus rhizosphaerae] |
| 06699 | 2 | hypothetical protein [Thauera phenolivorans] |
| 05733 | 2 | Fe-only nitrogenase subunit delta [Azoarcus nasutitermitis] |
| 06702 | 2 | hypothetical protein [Thauera phenolivorans] |
| 05722 | 2 | DUF4194 domain-containing protein [Thauera phenolivorans] |
| 05720 | 2 | hypothetical protein [Thauera butanivorans] |
| 05719 | 2 | hypothetical protein [Candidatus Dactylopiibacterium carminicum] |
| 05718 | 2 | prepilin-type N-terminal cleavage/methylation domain-containing protein [Nitrogeniibacter mangrovi] |
| 06714 | 2 | hypothetical protein [Candidatus Dactylopiibacterium carminicum] |
| 05710 | 2 | radical SAM protein [Azoarcus sp. DD4] |
| 05668 | 2 | hypothetical protein [Thauera aminoaromatica] |
| 05707 | 2 | hypothetical protein [Thauera butanivorans] |
| 05701 | 2 | IS1634 family transposase, partial [Thauera terpenica] |
| 05700 | 2 | hexameric tyrosine-coordinated heme protein [Thauera terpenica] |
| 06725 | 2 | hypothetical protein [Azoarcus sp. Aa7] |
| 05690 | 2 | type IV pili methyl-accepting chemotaxis transducer N-terminal domain-containing protein [Aromatoleum toluclasticum] |
| 05673 | 2 | hypothetical protein, partial [Thauera aminoaromatica] |
| 05671 | 2 | DEAD/DEAH box helicase family protein [Thauera aromatica] |
| 06735 | 2 | DUF2489 domain-containing protein [Niveibacterium microcysteis] |
| 06737 | 2 | GNAT family N-acetyltransferase [Niveibacterium microcysteis] |
| 04261 | 2 | hypothetical protein [Azoarcus communis] |
| 07677 | 2 | IS21 family transposase [Azoarcus sp. Aa7] |
| 09073 | 2 | hypothetical protein [Aromatoleum diolicum] |
| 08664 | 2 | MULTISPECIES: hypothetical protein [Proteobacteria] |
| 08614 | 2 | helix-turn-helix domain-containing protein, partial [Zoogloea dura] |
| 02920 | 2 | type II toxin-antitoxin system RelE/ParE family toxin [Aromatoleum aromaticum] |
| 08623 | 2 | SagB/ThcOx family dehydrogenase [Niveibacterium sp. COAC-50] |
| 02918 | 2 | hypothetical protein, partial [Azoarcus communis] |
| 08626 | 2 | hypothetical protein [Candidatus Dactylopiibacterium carminicum] |
| 02916 | 2 | hypothetical protein [Azoarcus communis] |
| 02909 | 2 | anaerobic C4-dicarboxylate transporter [Azoarcus communis] |
| 02905 | 2 | hypothetical protein [Azoarcus taiwanensis] |
| 08635 | 2 | hypothetical protein [Aromatoleum toluolicum] |
| 02894 | 2 | HD domain-containing protein [Aromatoleum toluvorans] |
| 08638 | 2 | OsmC family protein [Aromatoleum petrolei] |
| 08641 | 2 | DUF3313 domain-containing protein [Rhodocyclus purpureus] |
| 02890 | 2 | hypothetical protein [Denitromonas sp. IR12] |
| 02889 | 2 | RNA-guided endonuclease TnpB family protein [Aromatoleum bremense] |
| 02888 | 2 | hypothetical protein [Azoarcus halotolerans] |

| | | |
|---|---|---|
| 02887 | 2 | hypothetical protein [Thauera chlorobenzoica] |
| 02880 | 2 | hypothetical protein [Azoarcus olearius] |
| 02879 | 2 | hypothetical protein [Azoarcus olearius] |
| 02875 | 2 | hypothetical protein [Aromatoleum aromaticum] |
| 02868 | 2 | ATP-dependent Clp protease proteolytic subunit [Thauera chlorobenzoica] |
| 02863 | 2 | hypothetical protein [Thauera butanivorans] |
| 02937 | 2 | TonB-dependent receptor [Candidatus Dactylopiibacterium carminicum] |
| 02945 | 2 | hypothetical protein [Azoarcus nasutitermitis] |
| 02948 | 2 | hypothetical protein [Aromatoleum toluolicum] |
| 02975 | 2 | PEP-CTERM sorting domain-containing protein [Denitromonas halophilus] |
| 02994 | 2 | hypothetical protein [Denitromonas halophilus] |
| 02993 | 2 | MBL fold metallo-hydrolase [Azoarcus sp. KH32C] |
| 02991 | 2 | cytosine permease [Azoarcus halotolerans] |
| 08575 | 2 | hypothetical protein [Aromatoleum aromaticum] |
| 08576 | 2 | hypothetical protein [Azoarcus nasutitermitis] |
| 02982 | 2 | SH3 domain-containing protein [Zoogloea dura] |
| 02981 | 2 | ParA family protein [Zoogloea ramigera] |
| 08580 | 2 | hypothetical protein [Aromatoleum evansii] |
| 02980 | 2 | hypothetical protein [Zoogloea dura] |
| 08587 | 2 | hypothetical protein [Azoarcus communis] |
| 02949 | 2 | ABC transporter substrate-binding protein [Azoarcus rhizosphaerae] |
| 02974 | 2 | hypothetical protein [Denitromonas halophilus] |
| 02964 | 2 | DEAD/DEAH box helicase family protein, partial [Zoogloea ramigera] |
| 08592 | 2 | hypothetical protein [Azoarcus taiwanensis] |
| 02963 | 2 | hypothetical protein, partial [Zoogloea ramigera] |
| 08595 | 2 | hypothetical protein [Aromatoleum toluvorans] |
| 02961 | 2 | hypothetical protein [Thauera phenylacetica] |
| 08598 | 2 | hypothetical protein [Azoarcus sp. L1K30] |
| 08599 | 2 | thioesterase family protein [Aromatoleum petrolei] |
| 02957 | 2 | helix-turn-helix domain-containing protein [Thauera lacus] |
| 02861 | 2 | aromatic amino acid DMT transporter YddG [Thauera butanivorans] |
| 02858 | 2 | DUF4258 domain-containing protein [Azoarcus halotolerans] |
| 04258 | 2 | hypothetical protein [Aromatoleum evansii] |
| 08667 | 2 | 2-oxoacid:acceptor oxidoreductase subunit alpha [Aromatoleum bremense] |
| 02791 | 2 | hypothetical protein [Denitromonas sp. IR12] |
| 08705 | 2 | lysylphosphatidylglycerol synthase domain-containing protein [Thauera hydrothermalis] |
| 08706 | 2 | hypothetical protein [Azoarcus sp. DD4] |
| 02789 | 2 | hypothetical protein [Rhodocyclus purpureus] |
| 08708 | 2 | hypothetical protein [Azoarcus sp. KH32C] |
| 02788 | 2 | glycosyltransferase family 2 protein [Rhodocyclus purpureus] |
| 02784 | 2 | hypothetical protein [Propionivibrio dicarboxylicus] |
| 08713 | 2 | hypothetical protein [Aromatoleum toluvorans] |
| 02779 | 2 | hypothetical protein [Azoarcus sp. Aa7] |
| 02777 | 2 | hypothetical protein [Candidatus Dactylopiibacterium carminicum] |
| 02775 | 2 | YciI family protein [Candidatus Dactylopiibacterium carminicum] |
| 02772 | 2 | polysaccharide lyase family 7 protein [Candidatus Dactylopiibacterium carminicum] |
| 02769 | 2 | YciI family protein [Niveibacterium microcysteis] |
| 02767 | 2 | hypothetical protein [Niveibacterium sp. COAC-50] |
| 02763 | 2 | hypothetical protein [Niveibacterium microcysteis] |
| 08732 | 2 | hypothetical protein [Azoarcus sp. L1K30] |
| 02753 | 2 | MULTISPECIES: hypothetical protein [Uliginosibacterium] |
| 08734 | 2 | hypothetical protein [Azoarcus nasutitermitis] |
| 02751 | 2 | hypothetical protein, partial [Zoogloea dura] |
| 08736 | 2 | hypothetical protein [Azoarcus communis] |
| 02746 | 2 | zinc dependent phospholipase C family protein [Zoogloea dura] |
| 08703 | 2 | DUF4332 domain-containing protein [Thauera phenylacetica] |
| 02795 | 2 | hypothetical protein [Nitrogeniibacter aestuarii] |
| 02796 | 2 | hypothetical protein [Nitrogeniibacter aestuarii] |
| 02828 | 2 | hypothetical protein [Thauera phenylacetica] |
| 02855 | 2 | type II toxin-antitoxin system RelE/ParE family toxin [Azoarcus sp. Aa7] |
| 08672 | 2 | hypothetical protein [Azoarcus communis] |
| 08673 | 2 | hypothetical protein [Azoarcus sp. L1K30] |
| 02844 | 2 | hypothetical protein [Propionivibrio dicarboxylicus] |
| 02840 | 2 | hypothetical protein [Aromatoleum toluolicum] |
| 02839 | 2 | hypothetical protein [Aromatoleum diolicum] |
| 02832 | 2 | VC_2705 family sodium/solute symporter, partial [Thauera aminoaromatica] |
| 08680 | 2 | hypothetical protein [Aromatoleum aromaticum] |
| 02831 | 2 | tRNA lysidine(34) synthetase TilS, partial [Thauera aminoaromatica] |
| 02823 | 2 | hypothetical protein [Aromatoleum evansii] |
| 02798 | 2 | ATPase [Azoarcus sp. Aa7] |
| 08685 | 2 | hypothetical protein [Aromatoleum aromaticum] |
| 08689 | 2 | hypothetical protein [Azoarcus olearius] |
| 02813 | 2 | DUF4124 domain-containing protein [Aromatoleum tolulyticum] |
| 02809 | 2 | UvrD-helicase domain-containing protein [Aromatoleum toluolicum] |
| 02808 | 2 | hypothetical protein [Propionivibrio dicarboxylicus] |
| 08694 | 2 | CHASE3 domain-containing protein [Thauera phenylacetica] |
| 02807 | 2 | hypothetical protein [Propionivibrio dicarboxylicus] |
| 02806 | 2 | hypothetical protein [Propionivibrio dicarboxylicus] |
| 02804 | 2 | TonB family protein [Propionivibrio dicarboxylicus] |
| 03000 | 2 | hypothetical protein [Aromatoleum aromaticum] |
| 03002 | 2 | hypothetical protein, partial [Aromatoleum evansii] |
| 08560 | 2 | metallophosphoesterase [Azoarcus halotolerans] |
| 08559 | 2 | hypothetical protein [Azoarcus halotolerans] |
| 03159 | 2 | reverse transcriptase family protein [Uliginosibacterium gangwonense] |
| 08441 | 2 | DNA polymerase I [Propionivibrio dicarboxylicus] |
| 03158 | 2 | 6-phosphogluconolactonase [Propionivibrio dicarboxylicus] |
| 03151 | 2 | hypothetical protein [Azoarcus sp. KH32C] |
| 08446 | 2 | hypothetical protein [Uliginosibacterium aquaticum] |
| 06271 | 2 | hypothetical protein [Azoarcus olearius] |
| 03150 | 2 | DUF1810 domain-containing protein [Thauera phenylacetica] |
| 03146 | 2 | MULTISPECIES: hypothetical protein [Thauera] |
| 08455 | 2 | YkgJ family cysteine cluster protein [Thauera humireducens] |
| 03142 | 2 | hypothetical protein, partial [Thauera aminoaromatica] |
| 08457 | 2 | hypothetical protein [Azoarcus nasutitermitis] |

| | | |
|---|---|---|
| 08458 | 2 | hypothetical protein [Azoarcus halotolerans] |
| 08460 | 2 | antibiotic biosynthesis monooxygenase [Azoarcus taiwanensis] |
| 03137 | 2 | hypothetical protein [Aromatoleum aromaticum] |
| 03136 | 2 | hypothetical protein [Aromatoleum aromaticum] |
| 03129 | 2 | hypothetical protein [Propionivibrio dicarboxylicus] |
| 03128 | 2 | ABC transporter substrate-binding protein [Propionivibrio dicarboxylicus] |
| 08467 | 2 | PilZ domain-containing protein [Thauera aromatica] |
| 03125 | 2 | hypothetical protein, partial [Aromatoleum toluclasticum] |
| 08469 | 2 | ParB N-terminal domain-containing protein [Thauera butanivorans] |
| 08470 | 2 | hypothetical protein [Azoarcus sp. Aa7] |
| 03160 | 2 | hypothetical protein [Uliginosibacterium aquaticum] |
| 08437 | 2 | DUF202 domain-containing protein [Uliginosibacterium aquaticum] |
| 08436 | 2 | phosphatase PAP2 family protein [Niveibacterium umoris] |
| 08418 | 2 | hypothetical protein [Aromatoleum aromaticum] |
| 03204 | 2 | hypothetical protein [Aromatoleum pumilum] |
| 03203 | 2 | hypothetical protein [Thauera aromatica] |
| 08406 | 2 | hypothetical protein [Propionivibrio dicarboxylicus] |
| 03202 | 2 | hypothetical protein [Thauera humireducens] |
| 03201 | 2 | MULTISPECIES: pyrroloquinoline quinone precursor peptide PqqA [Thauera] |
| 08409 | 2 | hypothetical protein [Aromatoleum tolulyticum] |
| 03200 | 2 | DUF3280 domain-containing protein [Thauera propionica] |
| 03196 | 2 | hypothetical protein [Thauera chlorobenzoica] |
| 03194 | 2 | GIY-YIG nuclease family protein [Thauera terpenica] |
| 08419 | 2 | hypothetical protein [Aromatoleum aromaticum] |
| 03166 | 2 | NINE protein [Candidatus Dactylopiibacterium carminicum] |
| 08420 | 2 | pilin [Aromatoleum aromaticum] |
| 03182 | 2 | flagellar basal body-associated FliL family protein [Thauera humireducens] |
| 08422 | 2 | hypothetical protein [Azoarcus olearius] |
| 08425 | 2 | DUF305 domain-containing protein [Azoarcus taiwanensis] |
| 08426 | 2 | MFS transporter, partial [Thauera aminoaromatica] |
| 03174 | 2 | hypothetical protein [Aromatoleum toluclasticum] |
| 08429 | 2 | CHAT domain-containing protein [Zoogloea ramigera] |
| 03171 | 2 | phospholipase D family protein [Aromatoleum toluclasticum] |
| 03167 | 2 | hypothetical protein [Denitromonas sp. IR12] |
| 03123 | 2 | molecular chaperone TorD family protein [Aromatoleum toluclasticum] |
| 08473 | 2 | hypothetical protein [Thauera humireducens] |
| 08476 | 2 | mechanosensitive ion channel [Aromatoleum pumilum] |
| 03033 | 2 | DUF2628 domain-containing protein [Uliginosibacterium aquaticum] |
| 03061 | 2 | hypothetical protein [Aromatoleum toluclasticum] |
| 08523 | 2 | DUF1566 domain-containing protein [Azoarcus rhizosphaerae] |
| 08524 | 2 | hypothetical protein [Azoarcus rhizosphaerae] |
| 08525 | 2 | alpha/beta hydrolase [Azoarcus rhizosphaerae] |
| 03054 | 2 | NAD(P)H-binding protein [Azoarcus nasutitermitis] |
| 08529 | 2 | hypothetical protein [Zoogloea oleivorans] |
| 03039 | 2 | carbohydrate binding domain-containing protein [Candidatus Dactylopiibacterium carminicum] |
| 08540 | 2 | hypothetical protein [Denitromonas halophilus] |
| 08541 | 2 | hypothetical protein [Aromatoleum aromaticum] |
| 03031 | 2 | major capsid protein [Azoarcus communis] |
| 08518 | 2 | pilin [Azoarcus nasutitermitis] |
| 03029 | 2 | MULTISPECIES: hypothetical protein [Uliginosibacterium] |
| 03026 | 2 | hypothetical protein [Aromatoleum diolicum] |
| 03025 | 2 | hypothetical protein [Zoogloea dura] |
| 03014 | 2 | 4'-phosphopantetheinyl transferase superfamily protein [Azoarcus rhizosphaerae] |
| 03010 | 2 | hypothetical protein [Aromatoleum pumilum] |
| 08554 | 2 | hypothetical protein [Azospira oryzae] |
| 03007 | 2 | response regulator [Azoarcus indigens] |
| 03006 | 2 | CoA transferase, partial [Aromatoleum toluolicum] |
| 03004 | 2 | hypothetical protein [Azoarcus communis] |
| 08519 | 2 | hypothetical protein [Azoarcus rhizosphaerae] |
| 08517 | 2 | STN domain-containing protein [Azoarcus nasutitermitis] |
| 03116 | 2 | hypothetical protein [Nitrogeniibacter aestuarii] |
| 08493 | 2 | hypothetical protein [Aromatoleum toluclasticum] |
| 03111 | 2 | hypothetical protein [Azoarcus taiwanensis] |
| 08480 | 2 | TraB/GumN family protein [Aromatoleum pumilum] |
| 03106 | 2 | hypothetical protein [Azoarcus rhizosphaerae] |
| 08485 | 2 | DUF4214 domain-containing protein [Thauera aromatica] |
| 03095 | 2 | GSCFA domain-containing protein [Azoarcus halotolerans] |
| 03094 | 2 | DUF2976 domain-containing protein [Denitromonas sp. IR12] |
| 03092 | 2 | hypothetical protein [Azoarcus communis] |
| 08490 | 2 | ATP-binding domain-containing protein [Azoarcus halotolerans] |
| 03090 | 2 | hypothetical protein [Nitrogeniibacter aestuarii] |
| 08495 | 2 | hypothetical protein [Azoarcus communis] |
| 03063 | 2 | putative photosynthetic complex assembly protein PuhE [Rhodocyclus purpureus] |
| 03084 | 2 | flavodoxin family protein [Nitrogeniibacter aestuarii] |
| 08497 | 2 | hypothetical protein [Zoogloea dura] |
| 03081 | 2 | hypothetical protein [Nitrogeniibacter aestuarii] |
| 08501 | 2 | hypothetical protein [Nitrogeniibacter aestuarii] |
| 08502 | 2 | integrase core domain-containing protein [Azoarcus sp. DD4] |
| 03077 | 2 | hypothetical protein [Azoarcus communis] |
| 03074 | 2 | hypothetical protein [Niveibacterium microcysteis] |
| 03071 | 2 | hypothetical protein [Propionivibrio dicarboxylicus] |
| 08514 | 2 | MULTISPECIES: YqaE/Pmp3 family membrane protein [Azospira] |
| 02743 | 2 | hypothetical protein [Zoogloea dura] |
| 08742 | 2 | helix-turn-helix domain-containing protein [Thauera butanivorans] |
| 08744 | 2 | PEP-CTERM sorting domain-containing protein [Thauera butanivorans] |
| 02486 | 2 | condensation domain-containing protein, partial [Denitromonas sp. IR12] |
| 02434 | 2 | hypothetical protein [Azoarcus sp. Aa7] |
| 02433 | 2 | PhoD-like phosphatase N-terminal domain-containing protein, partial [Azoarcus sp. Aa7] |
| 08963 | 2 | 3'-5' exonuclease [Aromatoleum bremense] |
| 02431 | 2 | hypothetical protein [Azoarcus olearius] |
| 02428 | 2 | hypothetical protein [Azoarcus sp. DD4] |
| 02426 | 2 | hypothetical protein [Candidatus Dactylopiibacterium carminicum] |
| 02422 | 2 | PilZ domain-containing protein [Candidatus Dactylopiibacterium carminicum] |
| 02412 | 2 | hypothetical protein [Uliginosibacterium aquaticum] |

| | | |
|---|---|---|
| 08973 | 2 | replication endonuclease [Denitromonas sp. IR12] |
| 02409 | 2 | helix-turn-helix domain-containing protein, partial [Zoogloea dura] |
| 08979 | 2 | AI-2E family transporter [Denitromonas sp. IR12] |
| 02404 | 2 | hypothetical protein [Zoogloea dura] |
| 08981 | 2 | hypothetical protein [Aromatoleum aromaticum] |
| 02402 | 2 | hypothetical protein [Azoarcus rhizosphaerae] |
| 02396 | 2 | hypothetical protein [Azoarcus sp. DD4] |
| 08986 | 2 | V-type ATP synthase subunit F [Aromatoleum toluclasticum] |
| 08987 | 2 | hypothetical protein [Aromatoleum toluclasticum] |
| 02393 | 2 | DUF1566 domain-containing protein, partial [Azoarcus indigens] |
| 02389 | 2 | ThuA domain-containing protein [Aromatoleum petrolei] |
| 02388 | 2 | hypothetical protein [Aromatoleum toluolicum] |
| 08992 | 2 | amidohydrolase family protein [Propionivibrio dicarboxylicus] |
| 08959 | 2 | hypothetical protein [Niveibacterium umoris] |
| 02436 | 2 | S8 family serine peptidase [Rhodocyclus purpureus] |
| 08955 | 2 | hypothetical protein [Nitrogeniibacter mangrovi] |
| 08938 | 2 | toll/interleukin-1 receptor domain-containing protein [Rhodocyclus purpureus] |
| 08921 | 2 | hypothetical protein [Niveibacterium microcysteis] |
| 02483 | 2 | phosphotransferase [Aromatoleum toluolicum] |
| 02480 | 2 | gluconokinase [Azoarcus sp. KH32C] |
| 02476 | 2 | hypothetical protein [Aromatoleum petrolei] |
| 02472 | 2 | hypothetical protein [Aromatoleum bremense] |
| 02469 | 2 | 7,8-didemethyl-8-hydroxy-5-deazariboflavin synthase CofG [Aromatoleum toluclasticum] |
| 02468 | 2 | OsmC family protein [Aromatoleum bremense] |
| 02459 | 2 | hypothetical protein [Nitrogeniibacter aestuarii] |
| 02456 | 2 | hypothetical protein [Nitrogeniibacter aestuarii] |
| 08940 | 2 | DUF4376 domain-containing protein [Aromatoleum pumilum] |
| 02440 | 2 | PAS domain-containing protein [Rhodocyclus purpureus] |
| 02448 | 2 | hypothetical protein [Azoarcus rhizosphaerae] |
| 02447 | 2 | hypothetical protein [Niveibacterium microcysteis] |
| 02446 | 2 | hypothetical protein [Niveibacterium microcysteis] |
| 08945 | 2 | PKD domain-containing protein [Niveibacterium microcysteis] |
| 08946 | 2 | hypothetical protein [Nitrogeniibacter aestuarii] |
| 02445 | 2 | alpha-amylase [Niveibacterium sp. COAC-50] |
| 08949 | 2 | oleate hydratase [Niveibacterium microcysteis] |
| 08950 | 2 | hypothetical protein [Nitrogeniibacter aestuarii] |
| 08951 | 2 | MFS transporter [Nitrogeniibacter aestuarii] |
| 08994 | 2 | C-GCAxxG-C-C family protein [Propionivibrio dicarboxylicus] |
| 08995 | 2 | YecH family protein [Candidatus Dactylopiibacterium carminicum] |
| 08996 | 2 | hypothetical protein [Propionivibrio dicarboxylicus] |
| 09050 | 2 | hypothetical protein [Thauera humireducens] |
| 09038 | 2 | 5-amino-6-(D-ribitylamino)uracil--L-tyrosine 4-hydroxyphenyl transferase CofH [Aromatoleum toluclasticum] |
| 02328 | 2 | META domain-containing protein [Oryzomicrobium terrae] |
| 02325 | 2 | hypothetical protein [Zoogloea oleivorans] |
| 02323 | 2 | caspase family protein [Zoogloea oleivorans] |
| 02320 | 2 | M24 family metallopeptidase, partial [Azoarcus sp. L1K30] |
| 09046 | 2 | Flp family type IVb pilin [Azoarcus nasutitermitis] |
| 09047 | 2 | hypothetical protein [Azoarcus communis] |
| 02319 | 2 | hypothetical protein [Aromatoleum bremense] |
| 09049 | 2 | PEP-CTERM sorting domain-containing protein [Thauera butanivorans] |
| 09052 | 2 | hypothetical protein [Aromatoleum toluvorans] |
| 02330 | 2 | DUF6491 family protein [Zoogloea dura] |
| 02318 | 2 | hypothetical protein [Denitromonas halophilus] |
| 02317 | 2 | O-antigen ligase family protein [Denitromonas halophilus] |
| 02316 | 2 | hypothetical protein [Denitromonas halophilus] |
| 09059 | 2 | hypothetical protein [Aromatoleum petrolei] |
| 02309 | 2 | glycosyltransferase [Aromatoleum evansii] |
| 09062 | 2 | hypothetical protein [Thauera phenolivorans] |
| 02302 | 2 | FkbM family methyltransferase [Azoarcus rhizosphaerae] |
| 09071 | 2 | DDE-type integrase/transposase/recombinase [Aromatoleum diolicum] |
| 09072 | 2 | AbrB/MazE/SpoVT family DNA-binding domain-containing protein [Aromatoleum pumilum] |
| 09036 | 2 | hypothetical protein [Aromatoleum toluclasticum] |
| 09031 | 2 | hypothetical protein [Aromatoleum toluclasticum] |
| 02381 | 2 | DUF6505 family protein [Aromatoleum pumilum] |
| 02357 | 2 | DVUA0089 family protein [Zoogloea ramigera] |
| 08999 | 2 | vitamin K epoxide reductase family protein [Aromatoleum aromaticum] |
| 02378 | 2 | VTT domain-containing protein [Aromatoleum pumilum] |
| 09001 | 2 | hypothetical protein [Aromatoleum aromaticum] |
| 02377 | 2 | hypothetical protein [Proteobacteria] |
| 09004 | 2 | MULTISPECIES: heat resistance protein YfdX2 [Proteobacteria] |
| 02373 | 2 | hypothetical protein [Azoarcus communis] |
| 02363 | 2 | KTSC domain-containing protein [Aromatoleum aromaticum] |
| 02360 | 2 | MULTISPECIES: hypothetical protein [Betaproteobacteria] |
| 09012 | 2 | hypothetical protein [Azoarcus sp. L1K30] |
| 02356 | 2 | asparaginase [Aromatoleum aromaticum] |
| 02342 | 2 | S-adenosyl-l-methionine hydroxide adenosyltransferase family protein [Uliginosibacterium gangwonense] |
| 02349 | 2 | hypothetical protein [Azoarcus halotolerans] |
| 02346 | 2 | polysaccharide export protein [Azoarcus halotolerans] |
| 09020 | 2 | TROVE domain-containing protein [Azoarcus rhizosphaerae] |
| 02343 | 2 | ester cyclase [Aromatoleum toluclasticum] |
| 09024 | 2 | TRIC cation channel family protein [Uliginosibacterium gangwonense] |
| 09025 | 2 | hypothetical protein [Azoarcus sp. KH32C] |
| 09026 | 2 | S8 family serine peptidase [Aromatoleum pumilum] |
| 09027 | 2 | hypothetical protein [Oryzomicrobium terrae] |
| 09028 | 2 | hypothetical protein [Nitrogeniibacter aestuarii] |
| 08920 | 2 | nucleotidyltransferase family protein [Uliginosibacterium aquaticum] |
| 08917 | 2 | SLC13 family permease [Propionivibrio dicarboxylicus] |
| 08746 | 2 | hypothetical protein [Thauera butanivorans] |
| 02491 | 2 | GIY-YIG nuclease family protein [Nitrogeniibacter aestuarii] |
| 08787 | 2 | HIRAN domain-containing protein [Thauera hydrothermalis] |
| 08788 | 2 | hypothetical protein [Thauera hydrothermalis] |
| 08789 | 2 | hypothetical protein [Thauera hydrothermalis] |
| 02663 | 2 | acyl carrier protein [Thauera phenolivorans] |
| 02662 | 2 | nucleotidyl transferase AbiEii/AbiGii toxin family protein [Azoarcus communis] |

| | | |
|---|---|---|
| **02655** | 2 | HNH endonuclease, partial [Azoarcus rhizosphaerae] |
| **02654** | 2 | HrpE/YscL family type III secretion apparatus protein [Azoarcus rhizosphaerae] |
| **08805** | 2 | hypothetical protein, partial [Azoarcus indigens] |
| **08813** | 2 | hypothetical protein [Azoarcus rhizosphaerae] |
| **08815** | 2 | hypothetical protein [Aromatoleum toluvorans] |
| **02631** | 2 | acyl carrier protein [Azospira oryzae] |
| **02628** | 2 | hypothetical protein [Azoarcus communis] |
| **02627** | 2 | pilin [Aromatoleum aromaticum] |
| **02626** | 2 | DUF2514 family protein [Azoarcus communis] |
| **08825** | 2 | hypothetical protein [Zoogloea oleivorans] |
| **08826** | 2 | hypothetical protein [Zoogloea ramigera] |
| **02625** | 2 | helix-turn-helix domain-containing protein [Azoarcus communis] |
| **02621** | 2 | NERD domain-containing protein [Azoarcus communis] |
| **02618** | 2 | SEL1-like repeat protein, partial [Thauera lacus] |
| **02612** | 2 | flagellar protein FlhE [Thauera aromatica] |
| **02611** | 2 | putative sulfate/molybdate transporter [Aromatoleum pumilum] |
| **02676** | 2 | hypothetical protein [Zoogloea oleivorans] |
| **08781** | 2 | polysaccharide pyruvyl transferase family protein [Azoarcus communis] |
| **08780** | 2 | RES family NAD+ phosphorylase [Azoarcus communis] |
| **08759** | 2 | hypothetical protein [Thauera chlorobenzoica] |
| **02735** | 2 | MULTISPECIES: hypothetical protein [Betaproteobacteria] |
| **02734** | 2 | DUF429 domain-containing protein [Aromatoleum toluolicum] |
| **02726** | 2 | FAD-dependent oxidoreductase [Aromatoleum diolicum] |
| **08751** | 2 | hypothetical protein [Azoarcus olearius] |
| **02725** | 2 | glycosyl transferase [Aromatoleum diolicum] |
| **08753** | 2 | hypothetical protein [Thauera phenolivorans] |
| **02724** | 2 | O-antigen translocase [Aromatoleum toluvorans] |
| **02720** | 2 | MerR family transcriptional regulator [Azoarcus taiwanensis] |
| **08758** | 2 | hypothetical protein [Azoarcus sp. Aa7] |
| **02718** | 2 | DUF3080 domain-containing protein [Aromatoleum pumilum] |
| **02678** | 2 | hypothetical protein [Aromatoleum evansii] |
| **08762** | 2 | hypothetical protein [Thauera propionica] |
| **08763** | 2 | type II secretion system protein GspK [Thauera humireducens] |
| **02709** | 2 | MULTISPECIES: hypothetical protein [Rhodocyclales] |
| **08768** | 2 | hypothetical protein [Thauera aromatica] |
| **02705** | 2 | MULTISPECIES: hypothetical protein [Betaproteobacteria] |
| **08770** | 2 | hypothetical protein [Azoarcus rhizosphaerae] |
| **02702** | 2 | OBAP family protein [Aromatoleum aromaticum] |
| **02688** | 2 | DUF3617 family protein [Oryzomicrobium terrae] |
| **02682** | 2 | hypothetical protein [Zoogloea oleivorans] |
| **02608** | 2 | hypothetical protein [Thauera humireducens] |
| **02606** | 2 | hypothetical protein [Niveibacterium sp. COAC-50] |
| **02603** | 2 | retention module-containing protein [Thauera aromatica] |
| **02520** | 2 | hypothetical protein [Aromatoleum aromaticum] |
| **02541** | 2 | 50S ribosome-binding GTPase, partial [Thauera aminoaromatica] |
| **08884** | 2 | hypothetical protein [Azoarcus taiwanensis] |
| **02537** | 2 | AadA family aminoglycoside 3''-O-nucleotidyltransferase [Azoarcus nasutitermitis] |
| **02535** | 2 | MULTISPECIES: hypothetical protein [Thauera] |
| **02532** | 2 | hypothetical protein, partial [Thauera aminoaromatica] |
| **02525** | 2 | hypothetical protein [Azoarcus olearius] |
| **02523** | 2 | hypothetical protein [Azoarcus olearius] |
| **08894** | 2 | MULTISPECIES: hypothetical protein [Betaproteobacteria] |
| **02521** | 2 | hypothetical protein [Azoarcus olearius] |
| **02518** | 2 | hypothetical protein [Aromatoleum aromaticum] |
| **08878** | 2 | hypothetical protein [Azoarcus communis] |
| **02517** | 2 | hypothetical protein [Aromatoleum aromaticum] |
| **02516** | 2 | hypothetical protein [Aromatoleum aromaticum] |
| **02512** | 2 | DUF2271 domain-containing protein [Aromatoleum aromaticum] |
| **02505** | 2 | TRAP transporter small permease subunit [Propionivibrio dicarboxylicus] |
| **08907** | 2 | PIN domain-containing protein [Nitrogeniibacter aestuarii] |
| **02499** | 2 | hypothetical protein [Propionivibrio dicarboxylicus] |
| **08909** | 2 | hypothetical protein [Zoogloea dura] |
| **02498** | 2 | GNAT family N-acetyltransferase [Azoarcus rhizosphaerae] |
| **02497** | 2 | hypothetical protein [Aromatoleum toluclasticum] |
| **08879** | 2 | Hpt domain-containing protein [Azoarcus communis] |
| **02544** | 2 | hypothetical protein [Aromatoleum toluolicum] |
| **02596** | 2 | hypothetical protein [Aromatoleum evansii] |
| **02578** | 2 | type II toxin-antitoxin system MqsA family antitoxin [Aromatoleum bremense] |
| **08840** | 2 | hypothetical protein [Zoogloea oleivorans] |
| **08841** | 2 | hypothetical protein [Zoogloea oleivorans] |
| **02591** | 2 | DUF4224 domain-containing protein [Aromatoleum toluolicum] |
| **08843** | 2 | methyl-accepting chemotaxis protein, partial [Zoogloea dura] |
| **02590** | 2 | SIR2 family protein [Thauera butanivorans] |
| **02589** | 2 | SprT-like domain-containing protein [Aromatoleum evansii] |
| **02588** | 2 | hypothetical protein [Azoarcus nasutitermitis] |
| **08849** | 2 | Spy/CpxP family protein refolding chaperone [Oryzomicrobium terrae] |
| **02579** | 2 | N-acetyltransferase [Thauera linaloolentis] |
| **02574** | 2 | DUF6161 domain-containing protein [Thauera terpenica] |
| **08875** | 2 | ATP-binding protein [Azoarcus rhizosphaerae] |
| **08860** | 2 | hypothetical protein [Azoarcus halotolerans] |
| **02567** | 2 | DUF4258 domain-containing protein [Aromatoleum toluclasticum] |
| **08865** | 2 | PepSY domain-containing protein [Azoarcus halotolerans] |
| **02562** | 2 | MULTISPECIES: hypothetical protein [Uliginosibacterium] |
| **02561** | 2 | phage tail assembly chaperone [Uliginosibacterium gangwonense] |
| **08868** | 2 | hypothetical protein [Aromatoleum aromaticum] |
| **08869** | 2 | acyloxyacyl hydrolase [Aromatoleum aromaticum] |
| **02555** | 2 | zinc ribbon domain-containing protein [Niveibacterium umoris] |
| **08873** | 2 | DUF3560 domain-containing protein [Aromatoleum evansii] |
| **03205** | 2 | porin [Aromatoleum pumilum] |
| **08401** | 2 | hypothetical protein [Aromatoleum toluclasticum] |
| **03206** | 2 | hypothetical protein [Aromatoleum pumilum] |
| **03940** | 2 | hypothetical protein [Zoogloea dura] |
| **07920** | 2 | alkaline phosphatase D family protein [Azoarcus communis] |
| **07921** | 2 | hypothetical protein [Azoarcus communis] |

| | | |
|---|---|---|
| 03826 | 2 | hypothetical protein [Aromatoleum aromaticum] |
| 07925 | 2 | transposase, partial [Azoarcus indigens] |
| 07926 | 2 | hypothetical protein [Aromatoleum bremense] |
| 07929 | 2 | bifunctional ornithine acetyltransferase/N-acetylglutamate synthase, partial [Aromatoleum toluclasticum] |
| 03806 | 2 | hypothetical protein [Azoarcus nasutitermitis] |
| 07931 | 2 | anaerobic ribonucleoside-triphosphate reductase [Azoarcus taiwanensis] |
| 03803 | 2 | HNH endonuclease [Thauera chlorobenzoica] |
| 03802 | 2 | hypothetical protein [Azoarcus communis] |
| 03798 | 2 | hypothetical protein [Rhodocyclus purpureus] |
| 03794 | 2 | hypothetical protein [Azoarcus communis] |
| 03791 | 2 | hypothetical protein [Thauera phenylacetica] |
| 03788 | 2 | hypothetical protein [Denitromonas sp. IR12] |
| 03783 | 2 | GNAT family N-acetyltransferase [Uliginosibacterium gangwonense] |
| 03780 | 2 | PIN domain-containing protein [Azoarcus communis] |
| 03779 | 2 | hypothetical protein [Uliginosibacterium gangwonense] |
| 03776 | 2 | serine protease [Azoarcus halotolerans] |
| 03775 | 2 | sensor histidine kinase [Propionivibrio dicarboxylicus] |
| 03766 | 2 | hypothetical protein [Azoarcus sp. KH32C] |
| 07951 | 2 | hypothetical protein [Zoogloea dura] |
| 07919 | 2 | hypothetical protein [Azoarcus communis] |
| 07917 | 2 | MULTISPECIES: hypothetical protein [Betaproteobacteria] |
| 07916 | 2 | hypothetical protein [Aromatoleum bremense] |
| 07886 | 2 | DUF4384 domain-containing protein [Zoogloea oleivorans] |
| 03938 | 2 | hypothetical protein [Zoogloea oleivorans] |
| 03917 | 2 | hypothetical protein [Azoarcus communis] |
| 07871 | 2 | VCBS domain-containing protein [Zoogloea ramigera] |
| 07874 | 2 | PriCT-2 domain-containing protein [Zoogloea ramigera] |
| 03900 | 2 | hypothetical protein [Azoarcus rhizosphaerae] |
| 03897 | 2 | hypothetical protein [Azoarcus rhizosphaerae] |
| 07883 | 2 | hypothetical protein [Denitromonas halophilus] |
| 03896 | 2 | tape measure protein [Azoarcus communis] |
| 07885 | 2 | hypothetical protein [Azoarcus taiwanensis] |
| 03893 | 2 | hypothetical protein [Azoarcus rhizosphaerae] |
| 07915 | 2 | hypothetical protein [Aromatoleum diolicum] |
| 07898 | 2 | hypothetical protein [Azoarcus halotolerans] |
| 07901 | 2 | 3-hydroxyacyl-CoA dehydrogenase family protein [Aromatoleum toluolicum] |
| 03868 | 2 | TIGR04141 family sporadically distributed protein [Azospira oryzae] |
| 07908 | 2 | DUF1772 domain-containing protein [Aromatoleum buckelii] |
| 07909 | 2 | hypothetical protein [Aromatoleum buckelii] |
| 03846 | 2 | DUF4113 domain-containing protein [Azoarcus communis] |
| 03841 | 2 | roadblock/LC7 domain-containing protein [Azoarcus taiwanensis] |
| 07913 | 2 | hypothetical protein [Aromatoleum evansii] |
| 03828 | 2 | hypothetical protein [Aromatoleum pumilum] |
| 07952 | 2 | hypothetical protein [Zoogloea dura] |
| 03765 | 2 | hypothetical protein [Azoarcus sp. KH32C] |
| 07954 | 2 | MarR family transcriptional regulator [Thauera lacus] |
| 03679 | 2 | hypothetical protein [Thauera chlorobenzoica] |
| 07998 | 2 | PD-(D/E)XK nuclease-like domain-containing protein [Aromatoleum evansii] |
| 08003 | 2 | hypothetical protein [Niveibacterium microcysteis] |
| 03689 | 2 | hypothetical protein [Aromatoleum pumilum] |
| 08006 | 2 | glycoside hydrolase family 16 protein [Niveibacterium microcysteis] |
| 08007 | 2 | hypothetical protein [Niveibacterium microcysteis] |
| 03684 | 2 | hypothetical protein [Aromatoleum toluolicum] |
| 08011 | 2 | hypothetical protein [Niveibacterium microcysteis] |
| 03683 | 2 | hypothetical protein [Denitromonas sp. IR12] |
| 08016 | 2 | nucleotidyltransferase family protein [Nitrogeniibacter aestuarii] |
| 03671 | 2 | hypothetical protein [Azoarcus communis] |
| 03699 | 2 | ectonucleotide pyrophosphatase/phosphodiesterase [Oryzomicrobium terrae] |
| 03670 | 2 | hypothetical protein [Aromatoleum tolulyticum] |
| 03668 | 2 | 2OG-Fe(II) oxygenase [Thauera butanivorans] |
| 08025 | 2 | hypothetical protein [Aromatoleum bremense] |
| 03667 | 2 | hypothetical protein [Thauera butanivorans] |
| 08030 | 2 | hypothetical protein [Azoarcus sp. L1K30] |
| 08032 | 2 | hypothetical protein [Azoarcus sp. L1K30] |
| 03663 | 2 | DUF6364 family protein [Azoarcus nasutitermitis] |
| 03662 | 2 | glycosyltransferase [Aromatoleum pumilum] |
| 03659 | 2 | MULTISPECIES: EreA family erythromycin esterase [Bacteria] |
| 03697 | 2 | hypothetical protein [Thauera aromatica] |
| 07994 | 2 | hypothetical protein [Azoarcus sp. Aa7] |
| 03764 | 2 | hypothetical protein [Thauera phenylacetica] |
| 03712 | 2 | LytTR family transcriptional regulator DNA-binding domain-containing protein [Azospira oryzae] |
| 03760 | 2 | hypothetical protein [Thauera aminoaromatica] |
| 03757 | 2 | PEP-utilizing enzyme, partial [Azoarcus taiwanensis] |
| 07960 | 2 | PilX N-terminal domain-containing pilus assembly protein [Uliginosibacterium gangwonense] |
| 07961 | 2 | hypothetical protein [Candidatus Dactylopiibacterium carminicum] |
| 03754 | 2 | MULTISPECIES: hypothetical protein [Thauera] |
| 03747 | 2 | hypothetical protein [Aromatoleum aromaticum] |
| 03732 | 2 | MFS transporter [Aromatoleum petrolei] |
| 07970 | 2 | zinc-dependent metalloprotease [Uliginosibacterium aquaticum] |
| 03718 | 2 | hypothetical protein [Azospira oryzae] |
| 03711 | 2 | hypothetical protein [Thauera hydrothermalis] |
| 07993 | 2 | general secretion pathway protein GspB, partial [Aromatoleum toluvorans] |
| 03710 | 2 | helix-turn-helix domain-containing protein [Thauera hydrothermalis] |
| 07982 | 2 | pectate lyase [Candidatus Dactylopiibacterium carminicum] |
| 07983 | 2 | glycosyltransferase [Candidatus Dactylopiibacterium carminicum] |
| 07984 | 2 | MliC family protein [Candidatus Dactylopiibacterium carminicum] |
| 03708 | 2 | hypothetical protein [Azoarcus communis] |
| 03705 | 2 | class I SAM-dependent methyltransferase [Azoarcus communis] |
| 07989 | 2 | hypothetical protein [Candidatus Dactylopiibacterium carminicum] |
| 03702 | 2 | hypothetical protein [Aromatoleum petrolei] |
| 07991 | 2 | cell wall hydrolase [Aromatoleum evansii] |
| 07858 | 2 | hypothetical protein [Azoarcus rhizosphaerae] |
| 07856 | 2 | hypothetical protein [Azoarcus nasutitermitis] |
| 08039 | 2 | hypothetical protein [Denitromonas sp. IR12] |

| | | |
|---|---|---|
| 03946 | 2 | hypothetical protein [Oryzomicrobium terrae] |
| 07729 | 2 | ligase-associated DNA damage response endonuclease PdeM [Azoarcus olearius] |
| 04185 | 2 | DUF1428 domain-containing protein [Azoarcus communis] |
| 04176 | 2 | hypothetical protein [Thauera humireducens] |
| 07734 | 2 | Hpt domain-containing protein, partial [Thauera aminoaromatica] |
| 07735 | 2 | hypothetical protein, partial [Thauera aminoaromatica] |
| 07736 | 2 | MULTISPECIES: hypothetical protein [Thauera] |
| 04161 | 2 | beta-propeller fold lactonase family protein [Candidatus Dactylopiibacterium carminicum] |
| 04160 | 2 | hypothetical protein [Uliginosibacterium gangwonense] |
| 07741 | 2 | HAD-IC family P-type ATPase, partial [Thauera aminoaromatica] |
| 07742 | 2 | DUF3375 domain-containing protein [Thauera phenolivorans] |
| 04149 | 2 | alpha/beta fold hydrolase [Azoarcus rhizosphaerae] |
| 04147 | 2 | hypothetical protein [Thauera chlorobenzoica] |
| 04145 | 2 | TauD/TfdA family dioxygenase [Aromatoleum pumilum] |
| 07747 | 2 | hypothetical protein [Aromatoleum buckelii] |
| 04140 | 2 | hypothetical protein [Thauera phenylacetica] |
| 04126 | 2 | helix-turn-helix domain-containing protein [Zoogloea oleivorans] |
| 04125 | 2 | hypothetical protein [Thauera aminoaromatica] |
| 04124 | 2 | hypothetical protein [Aromatoleum aromaticum] |
| 04118 | 2 | TIM barrel protein [Propionivibrio dicarboxylicus] |
| 07759 | 2 | MULTISPECIES: hypothetical protein [Proteobacteria] |
| 04107 | 2 | hypothetical protein [Aromatoleum toluclasticum] |
| 07728 | 2 | FAD-dependent oxidoreductase [Aromatoleum aromaticum] |
| 07724 | 2 | ATP-binding protein [Aromatoleum aromaticum] |
| 07722 | 2 | TIGR04255 family protein [Aromatoleum aromaticum] |
| 07698 | 2 | hypothetical protein [Aromatoleum toluclasticum] |
| 04256 | 2 | hypothetical protein [Aromatoleum petrolei] |
| 04255 | 2 | hypothetical protein [Aromatoleum evansii] |
| 04254 | 2 | protein kinase family protein [Aromatoleum bremense] |
| 04253 | 2 | hypothetical protein [Aromatoleum bremense] |
| 07685 | 2 | hypothetical protein [Azoarcus communis] |
| 07690 | 2 | hypothetical protein [Denitromonas halophilus] |
| 07693 | 2 | hypothetical protein [Nitrogeniibacter aestuarii] |
| 07694 | 2 | toll/interleukin-1 receptor domain-containing protein [Nitrogeniibacter aestuarii] |
| 07696 | 2 | hypothetical protein, partial [Nitrogeniibacter aestuarii] |
| 07700 | 2 | DUF91 domain-containing protein [Aromatoleum toluolicum] |
| 07721 | 2 | carboxymuconolactone decarboxylase family protein [Aromatoleum aromaticum] |
| 04234 | 2 | hypothetical protein [Azospira oryzae] |
| 04233 | 2 | MULTISPECIES: hypothetical protein [Azospira] |
| 04231 | 2 | hypothetical protein [Thauera hydrothermalis] |
| 04229 | 2 | phosphoribosyl-AMP cyclohydrolase [Azoarcus communis] |
| 07707 | 2 | GtrA family protein [Aromatoleum pumilum] |
| 04217 | 2 | STAS domain-containing protein [Azospira oryzae] |
| 04211 | 2 | dihydroxy-acid dehydratase [Aromatoleum pumilum] |
| 04194 | 2 | helix-turn-helix domain-containing protein [Azoarcus olearius] |
| 04190 | 2 | Arc family DNA-binding protein [Thauera butanivorans] |
| 04103 | 2 | hypothetical protein [Nitrogeniibacter aestuarii] |
| 07765 | 2 | elements of external origin [Candidatus Dactylopiibacterium carminicum] |
| 04100 | 2 | glycosyltransferase [Denitromonas sp. IR12] |
| 04012 | 2 | DEAD/DEAH box helicase [Azoarcus sp. DD4] |
| 07814 | 2 | lytic murein transglycosylase [Thauera aromatica] |
| 07818 | 2 | type II toxin-antitoxin system Phd/YefM family antitoxin [Candidatus Dactylopiibacterium carminicum] |
| 04025 | 2 | RusA family crossover junction endodeoxyribonuclease [Azoarcus sp. L1K30] |
| 07823 | 2 | hypothetical protein [Aromatoleum toluclasticum] |
| 04024 | 2 | hypothetical protein [Aromatoleum bremense] |
| 07825 | 2 | hypothetical protein [Azoarcus communis] |
| 04023 | 2 | ATP-binding protein, partial [Thauera linaloolentis] |
| 04021 | 2 | phage holin family protein [Zoogloea dura] |
| 04020 | 2 | hypothetical protein [Zoogloea dura] |
| 04008 | 2 | DUF3883 domain-containing protein [Oryzomicrobium terrae] |
| 07812 | 2 | hypothetical protein [Thauera aromatica] |
| 03991 | 2 | DUF4400 domain-containing protein, partial [Azoarcus taiwanensis] |
| 03988 | 2 | hypothetical protein [Aromatoleum pumilum] |
| 03982 | 2 | HPr-rel-A system PqqD family peptide chaperone [Azoarcus indigens] |
| 03980 | 2 | hypothetical protein [Azoarcus communis] |
| 03976 | 2 | hypothetical protein [Aromatoleum evansii] |
| 03975 | 2 | hypothetical protein, partial [Aromatoleum evansii] |
| 03971 | 2 | AIPR family protein [Aromatoleum aromaticum] |
| 03965 | 2 | HAD hydrolase-like protein [Denitromonas halophilus] |
| 03949 | 2 | DUF535 family protein [Oryzomicrobium terrae] |
| 07813 | 2 | hypothetical protein [Thauera aromatica] |
| 07810 | 2 | DUF4325 domain-containing protein [Thauera aromatica] |
| 04085 | 2 | APC family permease [Niveibacterium umoris] |
| 07793 | 2 | hypothetical protein [Thauera chlorobenzoica] |
| 07772 | 2 | AAA family ATPase, partial [Thauera phenylacetica] |
| 04076 | 2 | thiol-activated cytolysin family protein [Nitrogeniibacter aestuarii] |
| 04069 | 2 | hypothetical protein [Azoarcus nasutitermitis] |
| 07783 | 2 | HPr-rel-A system PqqD family peptide chaperone [Azoarcus olearius] |
| 07784 | 2 | hypothetical protein [Azoarcus olearius] |
| 07785 | 2 | hypothetical protein [Azoarcus olearius] |
| 04063 | 2 | SPOR domain-containing protein [Oryzomicrobium terrae] |
| 07789 | 2 | hypothetical protein [Thauera phenolivorans] |
| 04062 | 2 | hypothetical protein, partial [Rhodocyclus purpureus] |
| 04055 | 2 | hypothetical protein [Propionivibrio dicarboxylicus] |
| 07808 | 2 | three-Cys-motif partner protein TcmP [Aromatoleum pumilum] |
| 07796 | 2 | metallophosphoesterase [Azoarcus communis] |
| 07798 | 2 | hypothetical protein [Thauera humireducens] |
| 07799 | 2 | hypothetical protein [Thauera humireducens] |
| 07800 | 2 | class I SAM-dependent methyltransferase [Aromatoleum aromaticum] |
| 04053 | 2 | hypothetical protein [Rhodocyclus purpureus] |
| 07802 | 2 | cupin domain-containing protein [Aromatoleum pumilum] |
| 04052 | 2 | cold shock domain-containing protein [Azoarcus olearius] |
| 04051 | 2 | hypothetical protein [Aromatoleum toluclasticum] |
| 04046 | 2 | helix-turn-helix domain-containing protein [Candidatus Dactylopiibacterium carminicum] |

| | | |
|---|---|---|
| 03657 | 2 | transcriptional repressor [Thauera butanivorans] |
| 03650 | 2 | toll/interleukin-1 receptor domain-containing protein [Aromatoleum toluclasticum] |
| 03214 | 2 | hypothetical protein [Azoarcus communis] |
| 08222 | 2 | hypothetical protein [Aromatoleum diolicum] |
| 03357 | 2 | IS1595 family transposase [Aromatoleum pumilum] |
| 08281 | 2 | AAA family ATPase, partial [Aromatoleum buckelii] |
| 03352 | 2 | hypothetical protein [Thauera humireducens] |
| 03347 | 2 | hypothetical protein [Nitrogeniibacter aestuarii] |
| 03342 | 2 | hypothetical protein [Azoarcus olearius] |
| 08288 | 2 | DUF4387 domain-containing protein [Aromatoleum toluvorans] |
| 08289 | 2 | molybdopterin-dependent oxidoreductase, partial [Aromatoleum toluvorans] |
| 08290 | 2 | hypothetical protein [Aromatoleum tolulyticum] |
| 08291 | 2 | peroxidase [Aromatoleum diolicum] |
| 03340 | 2 | hypothetical protein [Azoarcus olearius] |
| 08293 | 2 | MULTISPECIES: hypothetical protein [Betaproteobacteria] |
| 03338 | 2 | hypothetical protein [Azoarcus nasutitermitis] |
| 03332 | 2 | aldehyde dehydrogenase family protein, partial [Aromatoleum evansii] |
| 08298 | 2 | GMC family oxidoreductase N-terminal domain-containing protein [Aromatoleum toluolicum] |
| 03330 | 2 | HPr-rel-A system PqqD family peptide chaperone [Azoarcus taiwanensis] |
| 03329 | 2 | hypothetical protein [Aromatoleum toluclasticum] |
| 03327 | 2 | MULTISPECIES: hypothetical protein [Azospira] |
| 08303 | 2 | ATP-binding cassette domain-containing protein, partial [Azoarcus sp. TTM-91] |
| 03322 | 2 | hypothetical protein [Aromatoleum toluclasticum] |
| 03320 | 2 | hypothetical protein [Nitrogeniibacter aestuarii] |
| 08309 | 2 | IPTL-CTERM sorting domain-containing protein [Zoogloea dura] |
| 03364 | 2 | DUF4145 domain-containing protein [Thauera aromatica] |
| 03366 | 2 | hypothetical protein [Azoarcus rhizosphaerae] |
| 08274 | 2 | hypothetical protein [Aromatoleum evansii] |
| 08255 | 2 | hypothetical protein [Azoarcus halotolerans] |
| 03419 | 2 | sulfatase-like hydrolase/transferase [Zoogloea dura] |
| 03412 | 2 | recombinase family protein [Aromatoleum aromaticum] |
| 03409 | 2 | hypothetical protein [Azoarcus rhizosphaerae] |
| 08241 | 2 | hypothetical protein [Rhodocyclus purpureus] |
| 08243 | 2 | hypothetical protein [Propionivibrio dicarboxylicus] |
| 08245 | 2 | hypothetical protein [Zoogloea dura] |
| 03392 | 2 | hypothetical protein [Aromatoleum diolicum] |
| 08250 | 2 | MULTISPECIES: hypothetical protein [Thauera] |
| 03389 | 2 | hypothetical protein [Azospira oryzae] |
| 08257 | 2 | MULTISPECIES: hypothetical protein [Rhodocyclales] |
| 08273 | 2 | type IV secretion system protein [Aromatoleum evansii] |
| 03378 | 2 | hypothetical protein [Azoarcus communis] |
| 08261 | 2 | hypothetical protein [Azoarcus halotolerans] |
| 03374 | 2 | hypothetical protein [Aromatoleum toluclasticum] |
| 08264 | 2 | hypothetical protein [Azoarcus sp. L1K30] |
| 08265 | 2 | GNAT family N-acetyltransferase [Aromatoleum buckelii] |
| 03369 | 2 | hypothetical protein [Thauera lacus] |
| 08268 | 2 | erythromycin esterase family protein [Aromatoleum aromaticum] |
| 03368 | 2 | CcdB family protein [Thauera aromatica] |
| 08271 | 2 | hypothetical protein, partial [Aromatoleum buckelii] |
| 03311 | 2 | fimbria/pilus periplasmic chaperone [Azoarcus sp. TTM-91] |
| 03306 | 2 | hypothetical protein [Thauera terpenica] |
| 03304 | 2 | DUF4131 domain-containing protein [Azoarcus sp. TTM-91] |
| 03235 | 2 | pilin [Azoarcus indigens] |
| 03253 | 2 | hypothetical protein [Aromatoleum petrolei] |
| 08365 | 2 | hypothetical protein [Niveibacterium microcysteis] |
| 03243 | 2 | glycosyltransferase [Azoarcus sp. DD4] |
| 08369 | 2 | beta/gamma crystallin-related protein [Niveibacterium microcysteis] |
| 03241 | 2 | hypothetical protein [Azoarcus rhizosphaerae] |
| 08371 | 2 | hypothetical protein [Nitrogeniibacter aestuarii] |
| 03239 | 2 | glycosyltransferase family 1 protein [Azoarcus nasutitermitis] |
| 08374 | 2 | glycosyltransferase family 1 protein [Niveibacterium umoris] |
| 03237 | 2 | AAA family ATPase [Azoarcus nasutitermitis] |
| 08378 | 2 | hypothetical protein [Azoarcus taiwanensis] |
| 08358 | 2 | hypothetical protein [Propionivibrio dicarboxylicus] |
| 08384 | 2 | hypothetical protein [Aromatoleum bremense] |
| 03223 | 2 | UDP-forming cellulose synthase catalytic subunit [Aromatoleum bremense] |
| 03222 | 2 | hypothetical protein [Thauera hydrothermalis] |
| 08390 | 2 | tripartite tricarboxylate transporter TctB family protein [Azoarcus sp. L1K30] |
| 03221 | 2 | hypothetical protein [Candidatus Dactylopiibacterium carminicum] |
| 03220 | 2 | hypothetical protein [Azoarcus communis] |
| 08394 | 2 | ABC transporter permease [Denitromonas sp. IR12] |
| 03215 | 2 | DUF3465 domain-containing protein [Aromatoleum diolicum] |
| 08397 | 2 | hypothetical protein [Nitrogeniibacter aestuarii] |
| 08360 | 2 | TonB family protein [Rhodocyclus purpureus] |
| 08357 | 2 | DDE-type integrase/transposase/recombinase [Rhodocyclus purpureus] |
| 08319 | 2 | hypothetical protein [Candidatus Dactylopiibacterium carminicum] |
| 08333 | 2 | hypothetical protein [Niveibacterium sp. COAC-50] |
| 08320 | 2 | FkbM family methyltransferase [Aromatoleum pumilum] |
| 03300 | 2 | helix-turn-helix domain-containing protein, partial [Thauera phenylacetica] |
| 08323 | 2 | metallophosphoesterase [Niveibacterium umoris] |
| 03295 | 2 | hypothetical protein [Thauera aminoaromatica] |
| 03291 | 2 | lipase family protein [Azoarcus olearius] |
| 03289 | 2 | hypothetical protein [Azoarcus olearius] |
| 08329 | 2 | hypothetical protein [Niveibacterium sp. COAC-50] |
| 08331 | 2 | hypothetical protein [Niveibacterium sp. COAC-50] |
| 08332 | 2 | phospholipase D-like domain-containing protein [Nitrogeniibacter aestuarii] |
| 03287 | 2 | hypothetical protein [Aromatoleum aromaticum] |
| 03261 | 2 | hypothetical protein [Azoarcus sp. L1K30] |
| 03286 | 2 | hypothetical protein [Aromatoleum aromaticum] |
| 03279 | 2 | hypothetical protein [Aromatoleum diolicum] |
| 08343 | 2 | hypothetical protein [Candidatus Dactylopiibacterium carminicum] |
| 08345 | 2 | hypothetical protein [Azoarcus sp. Aa7] |
| 03265 | 2 | hypothetical protein [Zoogloea dura] |
| 08349 | 2 | hypothetical protein [Aromatoleum toluolicum] |

| | | |
|---|---|---|
| 08350 | 2 | hypothetical protein [Aromatoleum diolicum] |
| 08351 | 2 | sarcosine oxidase subunit delta [Azoarcus sp. Aa7] |
| 03263 | 2 | helix-turn-helix domain-containing protein [Zoogloea oleivorans] |
| 03421 | 2 | copper-binding protein, partial [Zoogloea ramigera] |
| 03428 | 2 | hypothetical protein [Aromatoleum aromaticum] |
| 08043 | 2 | hypothetical protein [Nitrogeniibacter aestuarii] |
| 08216 | 2 | Gfo/Idh/MocA family oxidoreductase [Azoarcus nasutitermitis] |
| 03600 | 2 | hypothetical protein [Azospira oryzae] |
| 08084 | 2 | MULTISPECIES: PEP-CTERM sorting domain-containing protein [Thauera] |
| 08085 | 2 | Hpt domain-containing protein [Thauera aminoaromatica] |
| 03599 | 2 | enoyl-CoA hydratase-related protein, partial [Aromatoleum toluclasticum] |
| 08088 | 2 | MULTISPECIES: type II toxin-antitoxin system VapC family toxin [Thauera] |
| 08092 | 2 | hypothetical protein [Thauera lacus] |
| 08095 | 2 | hypothetical protein [Thauera phenylacetica] |
| 08098 | 2 | DotA/TraY family protein [Thauera chlorobenzoica] |
| 03584 | 2 | hypothetical protein [Denitromonas sp. IR12] |
| 03579 | 2 | hypothetical protein [Azoarcus sp. L1K30] |
| 08105 | 2 | helix-turn-helix domain-containing GNAT family N-acetyltransferase [Azospira oryzae] |
| 03577 | 2 | hypothetical protein [Azoarcus communis] |
| 08108 | 2 | hypothetical protein [Uliginosibacterium gangwonense] |
| 08109 | 2 | response regulator [Propionivibrio dicarboxylicus] |
| 03573 | 2 | MULTISPECIES: NIPSNAP family protein [Thauera] |
| 03572 | 2 | cation:proton antiporter [Aromatoleum petrolei] |
| 03570 | 2 | hypothetical protein [Aromatoleum petrolei] |
| 08115 | 2 | NTP transferase domain-containing protein [Thauera terpenica] |
| 03569 | 2 | hypothetical protein [Aromatoleum petrolei] |
| 03568 | 2 | hypothetical protein [Aromatoleum aromaticum] |
| 08118 | 2 | NAD(P)-binding protein, partial [Thauera terpenica] |
| 08080 | 2 | phosphonopyruvate decarboxylase [Azoarcus olearius] |
| 03601 | 2 | hypothetical protein [Propionivibrio dicarboxylicus] |
| 03602 | 2 | hypothetical protein [Propionivibrio dicarboxylicus] |
| 03622 | 2 | hypothetical protein [Azoarcus olearius] |
| 08048 | 2 | DUF4265 domain-containing protein [Nitrogeniibacter aestuarii] |
| 08049 | 2 | hypothetical protein [Nitrogeniibacter aestuarii] |
| 03649 | 2 | helix-turn-helix domain-containing protein [Aromatoleum evansii] |
| 08051 | 2 | radical SAM protein [Aromatoleum toluclasticum] |
| 03635 | 2 | PepSY domain-containing protein [Aromatoleum toluolicum] |
| 03630 | 2 | lipopolysaccharide biosynthesis protein [Thauera phenolivorans] |
| 03629 | 2 | M23 family metallopeptidase [Thauera phenylacetica] |
| 08057 | 2 | hypothetical protein [Aromatoleum toluclasticum] |
| 03623 | 2 | NB-ARC domain-containing protein [Azoarcus sp. DD4] |
| 08061 | 2 | citrate lyase acyl carrier protein [Propionivibrio dicarboxylicus] |
| 08077 | 2 | hypothetical protein [Aromatoleum aromaticum] |
| 03621 | 2 | hypothetical protein [Azoarcus olearius] |
| 03615 | 2 | hypothetical protein [Aromatoleum aromaticum] |
| 08069 | 2 | hypothetical protein [Aromatoleum aromaticum] |
| 03611 | 2 | HipA domain-containing protein [Aromatoleum tolulyticum] |
| 08071 | 2 | hypothetical protein [Aromatoleum aromaticum] |
| 03610 | 2 | DUF5625 family protein [Propionivibrio dicarboxylicus] |
| 08073 | 2 | hypothetical protein [Aromatoleum aromaticum] |
| 08074 | 2 | hypothetical protein [Aromatoleum aromaticum] |
| 08076 | 2 | hypothetical protein [Aromatoleum aromaticum] |
| 08123 | 2 | hypothetical protein [Thauera linaloolentis] |
| 08125 | 2 | hypothetical protein [Azoarcus communis] |
| 08127 | 2 | hypothetical protein [Thauera humireducens] |
| 08195 | 2 | hypothetical protein [Azoarcus indigens] |
| 08176 | 2 | hypothetical protein [Azoarcus nasutitermitis] |
| 08177 | 2 | 4Fe-4S dicluster domain-containing protein [Azoarcus communis] |
| 03496 | 2 | class II glutamine amidotransferase [Azoarcus indigens] |
| 08186 | 2 | molecular chaperone DnaK [Aromatoleum pumilum] |
| 03488 | 2 | hypothetical protein [Aromatoleum toluolicum] |
| 03486 | 2 | HD domain-containing protein [Aromatoleum tolulyticum] |
| 03485 | 2 | hypothetical protein [Aromatoleum toluolicum] |
| 03475 | 2 | hypothetical protein [Aromatoleum toluvorans] |
| 03467 | 2 | FAD-dependent oxidoreductase, partial [Azoarcus taiwanensis] |
| 08196 | 2 | CpaD family pilus assembly lipoprotein [Azoarcus nasutitermitis] |
| 08174 | 2 | rhomboid family intramembrane serine protease [Oryzomicrobium terrae] |
| 08198 | 2 | phosphopantetheine-binding protein [Azoarcus nasutitermitis] |
| 03464 | 2 | HipA domain-containing protein [Azoarcus communis] |
| 03454 | 2 | PRTRC system protein C [Aromatoleum aromaticum] |
| 03453 | 2 | DUF559 domain-containing protein [Aromatoleum evansii] |
| 08206 | 2 | MULTISPECIES: hypothetical protein [Azospira] |
| 03451 | 2 | hypothetical protein [Aromatoleum buckelii] |
| 03448 | 2 | HNH endonuclease [Aromatoleum buckelii] |
| 03445 | 2 | thiamine pyrophosphate-dependent enzyme, partial [Azoarcus halotolerans] |
| 03442 | 2 | hypothetical protein [Aromatoleum bremense] |
| 08175 | 2 | PIN domain nuclease [Candidatus Dactylopiibacterium carminicum] |
| 03506 | 2 | MULTISPECIES: hypothetical protein [Uliginosibacterium] |
| 08128 | 2 | hypothetical protein [Thauera butanivorans] |
| 08149 | 2 | excisionase family DNA-binding protein [Rhodocyclus purpureus] |
| 03559 | 2 | mechanosensitive ion channel family protein [Nitrogeniibacter aestuarii] |
| 03558 | 2 | hypothetical protein [Denitromonas sp. IR12] |
| 03556 | 2 | PQQ-binding-like beta-propeller repeat protein [Nitrogeniibacter aestuarii] |
| 03553 | 2 | DMT family transporter [Niveibacterium microcysteis] |
| 08141 | 2 | hypothetical protein [Azoarcus olearius] |
| 03549 | 2 | hypothetical protein [Niveibacterium microcysteis] |
| 03533 | 2 | hypothetical protein [Niveibacterium umoris] |
| 03531 | 2 | ATP-binding protein [Aromatoleum toluolicum] |
| 08148 | 2 | MULTISPECIES: glycosyltransferase family 39 protein [Thauera] |
| 03527 | 2 | hypothetical protein [Azoarcus sp. Aa7] |
| 03509 | 2 | PilZ domain-containing protein [Uliginosibacterium aquaticum] |
| 08152 | 2 | MULTISPECIES: conjugal transfer protein TraD [Proteobacteria] |
| 03524 | 2 | AI-2E family transporter [Candidatus Dactylopiibacterium carminicum] |
| 08158 | 2 | hypothetical protein [Thauera humireducens] |

| | | |
|---|---|---|
| **03515** | 2 | hypothetical protein [Niveibacterium microcysteis] |
| **08164** | 2 | hypothetical protein [Aromatoleum pumilum] |
| **08165** | 2 | hypothetical protein [Thauera aromatica] |
| **03512** | 2 | SMI1/KNR4 family protein [Niveibacterium microcysteis] |
| **08167** | 2 | sensor domain-containing diguanylate cyclase [Thauera aromatica] |
| **03511** | 2 | acyl carrier protein [Uliginosibacterium gangwonense] |
| **08449** | 2 | hypothetical protein [Aromatoleum toluclasticum] |

Table S3: Protein subfamilies unique to Denitromonas spp.

| Gene Product Name | Cluster | Number of genomes with gene presence | Percent of genomes with gene presence |
|---|---|---|---|
| hypothetical protein [Denitromonas halophilus] | 04102 | 4 | 100% |
| ferrous iron transport protein A [Denitromonas halophilus] | 05753 | 4 | 100% |
| 3D domain-containing protein [Denitromonas halophilus] | 00938 | 4 | 100% |
| hypothetical protein [Denitromonas halophilus] | 02125 | 4 | 100% |
| hypothetical protein [Denitromonas halophilus] | 05757 | 4 | 100% |
| rhodanese family chromate resistance protein ChrE [Denitromonas halophilus] | 10412 | 4 | 100% |
| hypothetical protein [Denitromonas halophilus] | 08837 | 4 | 100% |
| hypothetical protein [Denitromonas halophilus] | 05314 | 4 | 100% |
| C45 family autoproteolytic acyltransferase/hydolase [Denitromonas halophilus] | 03429 | 4 | 100% |
| biotin carboxylase [Denitromonas halophilus] | 05758 | 4 | 100% |
| hypothetical protein [Denitromonas halophilus] | 09941 | 4 | 100% |
| hypothetical protein [Denitromonas halophilus] | 09503 | 4 | 100% |
| hypothetical protein [Denitromonas halophilus] | 01724 | 4 | 100% |
| hypothetical protein [Denitromonas halophilus] | 09173 | 4 | 100% |
| adenosylcobinamide amidohydrolase [Propionivibrio dicarboxylicus] | 01122 | 4 | 100% |
| hypothetical protein [Denitromonas halophilus] | 06243 | 4 | 100% |
| hypothetical protein [Denitromonas halophilus] | 09505 | 4 | 100% |
| hypothetical protein [Denitromonas halophilus] | 06071 | 4 | 100% |
| lamin tail domain-containing protein [Denitromonas halophilus] | 09752 | 4 | 100% |
| PEPxxWA-CTERM sorting domain-containing protein [Denitromonas halophilus] | 10597 | 4 | 100% |
| DUF4157 domain-containing protein [Denitromonas halophilus] | 05493 | 4 | 100% |
| DUF4157 domain-containing protein [Denitromonas halophilus] | 01729 | 4 | 100% |
| hypothetical protein [Denitromonas halophilus] | 01121 | 4 | 100% |
| glycosyltransferase [Denitromonas halophilus] | 03924 | 4 | 100% |
| DHCW motif cupin fold protein [Propionivibrio dicarboxylicus] | 10265 | 4 | 100% |
| hypothetical protein [Denitromonas halophilus] | 02965 | 4 | 100% |
| DUF2938 domain-containing protein [Denitromonas halophilus] | 06425 | 4 | 100% |
| hypothetical protein [Denitromonas halophilus] | 04430 | 4 | 100% |
| hypothetical protein [Denitromonas halophilus] | 04429 | 3 | 75% |
| hypothetical protein [Denitromonas halophilus] | 04808 | 3 | 75% |
| cytochrome c family protein [Denitromonas halophilus] | 08536 | 3 | 75% |
| hypothetical protein [Denitromonas halophilus] | 05491 | 3 | 75% |
| hypothetical protein [Denitromonas halophilus] | 05174 | 3 | 75% |
| GNAT family N-acetyltransferase [Denitromonas halophilus] | 10094 | 3 | 75% |
| hypothetical protein [Denitromonas halophilus] | 02674 | 3 | 75% |
| hypothetical protein [Denitromonas halophilus] | 08977 | 3 | 75% |
| hypothetical protein [Denitromonas halophilus] | 07519 | 3 | 75% |
| antibiotic biosynthesis monooxygenase [Denitromonas halophilus] | 09944 | 3 | 75% |
| hypothetical protein [Denitromonas halophilus] | 08233 | 3 | 75% |
| hypothetical protein [Denitromonas halophilus] | 00603 | 3 | 75% |
| hypothetical protein [Denitromonas halophilus] | 10414 | 3 | 75% |
| HAMP domain-containing histidine kinase [Denitromonas halophilus] | 07880 | 3 | 75% |
| GGDEF domain-containing protein [Denitromonas halophilus] | 03934 | 3 | 75% |
| type VI secretion system Vgr family protein, partial [Denitromonas halophilus] | 05900 | 3 | 75% |
| hypothetical protein [Denitromonas halophilus] | 10421 | 3 | 75% |
| prepilin-type N-terminal cleavage/methylation domain-containing protein [Denitromonas halophilus] | 00935 | 3 | 75% |
| cupin domain-containing protein [Denitromonas halophilus] | 02322 | 3 | 75% |
| hypothetical protein [Denitromonas halophilus] | 08234 | 3 | 75% |
| chlorite dismutase family protein [Denitromonas halophilus] | 07878 | 3 | 75% |
| PEP-CTERM sorting domain-containing protein [Denitromonas halophilus] | 06419 | 3 | 75% |
| cupin domain-containing protein [Denitromonas halophilus] | 06421 | 3 | 75% |
| hypothetical protein [Denitromonas halophilus] | 05179 | 3 | 75% |
| hypothetical protein [Denitromonas halophilus] | 00393 | 3 | 75% |
| hypothetical protein [Denitromonas halophilus] | 06068 | 3 | 75% |
| hypothetical protein [Denitromonas halophilus] | 01115 | 3 | 75% |
| hypothetical protein [Denitromonas halophilus] | 01721 | 3 | 75% |
| hypothetical protein [Denitromonas halophilus] | 09174 | 3 | 75% |
| hypothetical protein [Denitromonas halophilus] | 00390 | 3 | 75% |
| hypothetical protein [Denitromonas halophilus] | 01129 | 3 | 75% |
| hypothetical protein [Denitromonas halophilus] | 06075 | 3 | 75% |
| hypothetical protein [Denitromonas halophilus] | 02670 | 3 | 75% |
| hypothetical protein [Denitromonas halophilus] | 09507 | 3 | 75% |
| toll/interleukin-1 receptor domain-containing protein [Denitromonas halophilus] | 04809 | 3 | 75% |
| hypothetical protein [Denitromonas halophilus] | 05171 | 3 | 75% |
| hypothetical protein [Denitromonas halophilus] | 06932 | 3 | 75% |
| hypothetical protein [Denitromonas halophilus] | 03249 | 3 | 75% |
| hypothetical protein [Denitromonas halophilus] | 06072 | 3 | 75% |
| GNAT family N-acetyltransferase [Denitromonas halophilus] | 10048 | 3 | 75% |
| hypothetical protein [Denitromonas halophilus] | 09164 | 3 | 75% |
| winged helix DNA-binding domain-containing protein [Denitromonas halophilus] | 08975 | 3 | 75% |
| DUF1275 domain-containing protein [Denitromonas halophilus] | 03423 | 3 | 75% |
| hypothetical protein [Denitromonas halophilus] | 01557 | 3 | 75% |
| hypothetical protein [Denitromonas halophilus] | 09501 | 3 | 75% |
| FAD-binding protein [Denitromonas halophilus] | 06417 | 3 | 75% |
| acyloxyacyl hydrolase [Denitromonas sp. IR12] | 06423 | 2 | 50% |
| PEP-CTERM sorting domain-containing protein [Denitromonas sp. IR12] | 05921 | 2 | 50% |
| O-antigen ligase family protein [Denitromonas sp. IR12] | 02317 | 2 | 50% |
| hypothetical protein [Denitromonas sp. IR12] | 07690 | 2 | 50% |
| FAD-dependent oxidoreductase, partial [Denitromonas halophilus] | 03928 | 2 | 50% |
| hypothetical protein [Denitromonas sp. IR12] | 02974 | 2 | 50% |
| hypothetical protein [Denitromonas halophilus] | 00939 | 2 | 50% |
| cupin domain-containing protein [Denitromonas sp. IR12] | 10240 | 2 | 50% |
| TAXI family TRAP transporter solute-binding subunit, partial [Denitromonas sp. IR12] | 09350 | 2 | 50% |

| | | | |
|---|---|---|---|
| hypothetical protein [Denitromonas sp. IR12] | 04977 | 2 | 50% |
| hypothetical protein [Denitromonas sp. IR12] | 07883 | 2 | 50% |
| hypothetical protein [Denitromonas sp. IR12] | 02126 | 2 | 50% |
| DUF2345 domain-containing protein, partial [Denitromonas sp. IR12] | 03589 | 2 | 50% |
| hypothetical protein [Denitromonas sp. IR12] | 10100 | 2 | 50% |
| hypothetical protein [Denitromonas sp. IR12] | 01312 | 2 | 50% |
| hypothetical protein [Denitromonas halophilus] | 00081 | 2 | 50% |
| hypothetical protein [Denitromonas halophilus] | 04812 | 2 | 50% |
| IS3 family transposase, partial [Denitromonas halophilus] | 03921 | 2 | 50% |
| hypothetical protein [Denitromonas sp. IR12] | 02316 | 2 | 50% |
| phosphocholine cytidylyltransferase family protein [Denitromonas halophilus] | 06949 | 2 | 50% |
| ATP-binding domain-containing protein [Denitromonas halophilus] | 09178 | 2 | 50% |
| DUF695 domain-containing protein [Denitromonas sp. IR12] | 09352 | 2 | 50% |
| hypothetical protein [Denitromonas halophilus] | 08835 | 2 | 50% |
| pyruvate phosphate dikinase [Denitromonas halophilus] | 06947 | 2 | 50% |
| hypothetical protein [Niveibacterium sp. COAC-50] | 02673 | 1 | 25% |
| cadherin domain-containing protein, partial [Denitromonas sp. IR12] | 03266 | 1 | 25% |
| phage late control D family protein, partial [Denitromonas sp. IR12] | 03590 | 1 | 25% |
| hypothetical protein, partial [Denitromonas sp. IR12] | 07692 | 1 | 25% |
| Ig-like domain-containing protein, partial [Denitromonas sp. IR12] | 04591 | 1 | 25% |
| transposase, partial [Denitromonas sp. IR12] | 03113 | 1 | 25% |
| hypothetical protein, partial [Denitromonas sp. IR12] | 03267 | 1 | 25% |
| hypothetical protein [Niveibacterium sp. COAC-50] | 09714 | 1 | 25% |
| hypothetical protein [Denitromonas halophilus] | 07521 | 1 | 25% |
| hypothetical protein [Denitromonas halophilus] | 09141 | 1 | 25% |
| AMP-binding protein, partial [Denitromonas sp. IR12] | 00779 | 1 | 25% |
| AMP-binding protein, partial [Denitromonas sp. IR12] | 04592 | 1 | 25% |
| hypothetical protein [Azospira oryzae] | 04819 | 1 | 25% |
| condensation domain-containing protein, partial [Denitromonas sp. IR12] | 02486 | 1 | 25% |
| hypothetical protein [Niveibacterium sp. COAC-50] | 04431 | 1 | 25% |
| hypothetical protein [Propionivibrio dicarboxylicus] | 06782 | 1 | 25% |
| hypothetical protein [Denitromonas sp. IR12] | 05899 | 1 | 25% |
| DUF2345 domain-containing protein, partial [Denitromonas sp. IR12] | 04978 | 1 | 25% |
| hypothetical protein [Rhodocyclus purpureus] | 08540 | 1 | 25% |
| DUF2190 family protein, partial [Rhodocyclus purpureus] | 05755 | 1 | 25% |
| hypothetical protein [Denitromonas sp. IR12] | 04269 | 1 | 25% |
| hypothetical protein [Denitromonas sp. IR12] | 05309 | 1 | 25% |
| 2OG-Fe(II) oxygenase [Denitromonas sp. IR12] | 04567 | 1 | 25% |
| helix-turn-helix domain-containing protein, partial [Denitromonas sp. IR12] | 05920 | 1 | 25% |
| transposase domain-containing protein, partial [Denitromonas sp. IR12] | 10243 | 1 | 25% |
| phage portal protein [Denitromonas sp. IR12] | 05604 | 1 | 25% |
| ABC transporter permease [Denitromonas sp. IR12] | 08394 | 1 | 25% |
| glycosyltransferase [Denitromonas sp. IR12] | 04100 | 1 | 25% |
| hypothetical protein, partial [Denitromonas sp. IR12] | 01560 | 1 | 25% |
| hypothetical protein [Denitromonas sp. IR12] | 09280 | 1 | 25% |
| hypothetical protein [Denitromonas sp. IR12] | 02229 | 1 | 25% |
| helix-turn-helix transcriptional regulator [Denitromonas sp. IR12] | 00222 | 1 | 25% |
| DUF2975 domain-containing protein [Denitromonas sp. IR12] | 04383 | 1 | 25% |
| hypothetical protein, partial [Denitromonas halophilus] | 01728 | 1 | 25% |

Table S4: Protein subfamilies unique to close relatives of Denitromonas spp.

| Gene Product Name | Cluster | Number of genomes with gene presence | Percent of genomes with gene presence |
|---|---|---|---|
| hypothetical protein [Nitrogeniibacter aestuarii] | 09772 | 7 | 100% |
| hypothetical protein [Nitrogeniibacter aestuarii] | 00413 | 7 | 100% |
| hypothetical protein [Nitrogeniibacter aestuarii] | 06929 | 7 | 100% |
| DUF1289 domain-containing protein [Nitrogeniibacter aestuarii] | 01369 | 7 | 100% |
| hypothetical protein [Nitrogeniibacter aestuarii] | 05205 | 7 | 100% |
| orotidine-5'-phosphate decarboxylase [Nitrogeniibacter aestuarii] | 03110 | 7 | 100% |
| hypothetical protein [Nitrogeniibacter aestuarii] | 05893 | 7 | 100% |
| hypothetical protein [Nitrogeniibacter aestuarii] | 04421 | 7 | 100% |
| hypothetical protein [Nitrogeniibacter aestuarii] | 06413 | 7 | 100% |
| DUF2853 family protein [Nitrogeniibacter aestuarii] | 01275 | 7 | 100% |
| PEP-CTERM sorting domain-containing protein [Nitrogeniibacter aestuarii] | 05513 | 7 | 100% |
| hypothetical protein [Nitrogeniibacter aestuarii] | 06955 | 7 | 100% |
| TetR/AcrR family transcriptional regulator [Nitrogeniibacter aestuarii] | 05623 | 7 | 100% |
| hypothetical protein [Nitrogeniibacter aestuarii] | 01285 | 7 | 100% |
| DUF2007 domain-containing protein [Nitrogeniibacter aestuarii] | 10061 | 7 | 100% |
| hypothetical protein [Nitrogeniibacter aestuarii] | 07330 | 7 | 100% |
| DUF1499 domain-containing protein [Nitrogeniibacter aestuarii] | 10060 | 7 | 100% |
| DUF4040 domain-containing protein [Nitrogeniibacter aestuarii] | 08539 | 7 | 100% |
| EF-hand domain-containing protein [Nitrogeniibacter aestuarii] | 08033 | 7 | 100% |
| hypothetical protein [Nitrogeniibacter aestuarii] | 06070 | 7 | 100% |
| ribosomal protein L7/L12 [Nitrogeniibacter aestuarii] | 02350 | 7 | 100% |
| hypothetical protein [Nitrogeniibacter aestuarii] | 08267 | 7 | 100% |
| hypothetical protein [Nitrogeniibacter aestuarii] | 03079 | 7 | 100% |
| MBL fold metallo-hydrolase [Nitrogeniibacter aestuarii] | 07104 | 7 | 100% |
| hypothetical protein [Nitrogeniibacter aestuarii] | 08978 | 7 | 100% |
| hypothetical protein [Nitrogeniibacter aestuarii] | 07097 | 7 | 100% |
| Na+/H+ antiporter subunit E [Nitrogeniibacter aestuarii] | 09140 | 7 | 100% |
| Na(+)/H(+) antiporter subunit B [Nitrogeniibacter aestuarii] | 09623 | 7 | 100% |
| hypothetical protein [Nitrogeniibacter aestuarii] | 07334 | 7 | 100% |
| GNAT family N-acetyltransferase [Nitrogeniibacter aestuarii] | 04810 | 6 | 86% |
| hypothetical protein [Nitrogeniibacter aestuarii] | 01947 | 6 | 86% |
| VOC family protein [Nitrogeniibacter aestuarii] | 03560 | 6 | 86% |
| dodecin family protein [Nitrogeniibacter aestuarii] | 02452 | 6 | 86% |
| PEP-CTERM sorting domain-containing protein [Nitrogeniibacter aestuarii] | 01720 | 6 | 86% |
| PEP-CTERM sorting domain-containing protein [Nitrogeniibacter aestuarii] | 04560 | 6 | 86% |
| PAS domain-containing sensor histidine kinase [Nitrogeniibacter aestuarii] | 04770 | 6 | 86% |
| nuclear transport factor 2 family protein [Nitrogeniibacter aestuarii] | 01978 | 6 | 86% |
| S1C family serine protease [Denitromonas halophilus] | 00595 | 5 | 71% |
| HAMP domain-containing histidine kinase [Nitrogeniibacter aestuarii] | 05317 | 5 | 71% |
| hypothetical protein [Nitrogeniibacter aestuarii] | 05633 | 5 | 71% |
| hypothetical protein [Denitromonas halophilus] | 08861 | 5 | 71% |
| hypothetical protein [Denitromonas halophilus] | 01402 | 5 | 71% |
| hypothetical protein [Denitromonas halophilus] | 02669 | 5 | 71% |
| HEAT repeat domain-containing protein [Nitrogeniibacter aestuarii] | 03115 | 5 | 71% |
| aspartyl/asparaginyl beta-hydroxylase domain-containing protein [Nitrogeniibacter aestuarii] | 08657 | 5 | 71% |
| hypothetical protein [Nitrogeniibacter aestuarii] | 06074 | 5 | 71% |
| cupin domain-containing protein [Nitrogeniibacter aestuarii] | 06956 | 5 | 71% |
| sulfotransferase [Denitromonas halophilus] | 10446 | 5 | 71% |
| hypothetical protein [Nitrogeniibacter aestuarii] | 10066 | 5 | 71% |
| amidohydrolase family protein [Nitrogeniibacter aestuarii] | 02971 | 5 | 71% |
| PGDYG domain-containing protein [Denitromonas halophilus] | 03935 | 5 | 71% |
| zinc-ribbon domain-containing protein, partial [Nitrogeniibacter aestuarii] | 03432 | 5 | 71% |
| hypothetical protein [Denitromonas halophilus] | 07214 | 5 | 71% |
| DUF4157 domain-containing protein [Denitromonas halophilus] | 05786 | 5 | 71% |
| OPT/YSL family transporter [Nitrogeniibacter aestuarii] | 07631 | 4 | 57% |
| hypothetical protein [Nitrogeniibacter aestuarii] | 07689 | 4 | 57% |
| hypothetical protein [Denitromonas halophilus] | 09503 | 4 | 57% |
| ferrous iron transport protein A [Denitromonas halophilus] | 05753 | 4 | 57% |
| hypothetical protein [Denitromonas halophilus] | 09941 | 4 | 57% |
| hypothetical protein [Denitromonas halophilus] | 01724 | 4 | 57% |
| glycosyltransferase [Denitromonas halophilus] | 03924 | 4 | 57% |
| hypothetical protein [Nitrogeniibacter aestuarii] | 07522 | 4 | 57% |
| hypothetical protein [Nitrogeniibacter aestuarii] | 10216 | 4 | 57% |
| biotin carboxylase [Denitromonas halophilus] | 05758 | 4 | 57% |
| adenosylcobinamide amidohydrolase [Propionivibrio dicarboxylicus] | 01122 | 4 | 57% |
| hypothetical protein [Denitromonas halophilus] | 04430 | 4 | 57% |
| DUF2938 domain-containing protein [Denitromonas halophilus] | 06425 | 4 | 57% |
| DHCW motif cupin fold protein [Propionivibrio dicarboxylicus] | 10265 | 4 | 57% |
| hypothetical protein [Denitromonas halophilus] | 02965 | 4 | 57% |
| C45 family autoproteolytic acyltransferase/hydolase [Denitromonas halophilus] | 03429 | 4 | 57% |
| hypothetical protein [Denitromonas halophilus] | 05314 | 4 | 57% |
| alpha/beta fold hydrolase [Nitrogeniibacter aestuarii] | 04597 | 4 | 57% |
| hypothetical protein [Denitromonas halophilus] | 08837 | 4 | 57% |
| hypothetical protein [Denitromonas halophilus] | 05757 | 4 | 57% |
| hypothetical protein [Denitromonas halophilus] | 09173 | 4 | 57% |
| sulfurtransferase [Denitromonas halophilus] | 10412 | 4 | 57% |
| hypothetical protein [Denitromonas halophilus] | 02125 | 4 | 57% |
| DUF4157 domain-containing protein [Denitromonas halophilus] | 01729 | 4 | 57% |
| hypothetical protein [Nitrogeniibacter aestuarii] | 01973 | 4 | 57% |
| hypothetical protein [Denitromonas halophilus] | 09505 | 4 | 57% |
| hypothetical protein [Denitromonas halophilus] | 06243 | 4 | 57% |
| hypothetical protein [Denitromonas halophilus] | 06071 | 4 | 57% |
| 3D domain-containing protein [Denitromonas halophilus] | 00938 | 4 | 57% |
| lamin tail domain-containing protein [Denitromonas halophilus] | 09752 | 4 | 57% |
| DUF4157 domain-containing protein [Denitromonas halophilus] | 05493 | 4 | 57% |

| | | | |
|---|---|---|---|
| PEPxxWA-CTERM sorting domain-containing protein [Denitromonas halophilus] | 10597 | 4 | 57% |
| hypothetical protein [Denitromonas halophilus] | 01121 | 4 | 57% |
| PEP-CTERM sorting domain-containing protein [Denitromonas halophilus] | 04458 | 4 | 57% |
| hypothetical protein [Denitromonas halophilus] | 04102 | 4 | 57% |
| META domain-containing protein [Nitrogeniibacter aestuarii] | 04950 | 3 | 43% |
| hypothetical protein [Denitromonas halophilus] | 08233 | 3 | 43% |
| hypothetical protein [Denitromonas halophilus] | 07519 | 3 | 43% |
| hypothetical protein [Denitromonas halophilus] | 00393 | 3 | 43% |
| toll/interleukin-1 receptor domain-containing protein [Denitromonas halophilus] | 04809 | 3 | 43% |
| hypothetical protein [Denitromonas halophilus] | 04808 | 3 | 43% |
| hypothetical protein [Denitromonas halophilus] | 00603 | 3 | 43% |
| ABC transporter substrate-binding protein [Nitrogeniibacter aestuarii] | 09771 | 3 | 43% |
| winged helix DNA-binding domain-containing protein [Denitromonas halophilus] | 08975 | 3 | 43% |
| hypothetical protein [Denitromonas halophilus] | 05171 | 3 | 43% |
| hypothetical protein [Denitromonas halophilus] | 04429 | 3 | 43% |
| hypothetical protein [Denitromonas halophilus] | 08977 | 3 | 43% |
| hypothetical protein [Denitromonas halophilus] | 05179 | 3 | 43% |
| Spy/CpxP family protein refolding chaperone [Nitrogeniibacter aestuarii] | 04944 | 3 | 43% |
| hypothetical protein [Denitromonas halophilus] | 09174 | 3 | 43% |
| hypothetical protein [Denitromonas halophilus] | 09507 | 3 | 43% |
| hypothetical protein [Denitromonas halophilus] | 02674 | 3 | 43% |
| hypothetical protein [Denitromonas halophilus] | 09501 | 3 | 43% |
| hypothetical protein [Nitrogeniibacter aestuarii] | 06198 | 3 | 43% |
| M48 family metalloprotease [Nitrogeniibacter aestuarii] | 06570 | 3 | 43% |
| 2OG-Fe(II) oxygenase [Nitrogeniibacter aestuarii] | 05289 | 3 | 43% |
| DUF1275 domain-containing protein [Denitromonas halophilus] | 03423 | 3 | 43% |
| hypothetical protein [Denitromonas halophilus] | 01115 | 3 | 43% |
| hypothetical protein [Denitromonas halophilus] | 00390 | 3 | 43% |
| tetratricopeptide repeat protein [Nitrogeniibacter aestuarii] | 02995 | 3 | 43% |
| TIGR00341 family protein [Nitrogeniibacter aestuarii] | 05631 | 3 | 43% |
| hypothetical protein [Denitromonas halophilus] | 09164 | 3 | 43% |
| GNAT family N-acetyltransferase [Denitromonas halophilus] | 10048 | 3 | 43% |
| hypothetical protein [Denitromonas halophilus] | 01721 | 3 | 43% |
| hypothetical protein [Denitromonas halophilus] | 06068 | 3 | 43% |
| hypothetical protein [Denitromonas halophilus] | 01129 | 3 | 43% |
| hypothetical protein [Denitromonas halophilus] | 06075 | 3 | 43% |
| hypothetical protein [Denitromonas halophilus] | 06932 | 3 | 43% |
| DUF2946 family protein [Nitrogeniibacter aestuarii] | 05316 | 3 | 43% |
| cupin domain-containing protein [Denitromonas halophilus] | 06421 | 3 | 43% |
| hypothetical protein [Denitromonas halophilus] | 02670 | 3 | 43% |
| hypothetical protein [Denitromonas halophilus] | 08234 | 3 | 43% |
| hypothetical protein [Nitrogeniibacter aestuarii] | 08570 | 3 | 43% |
| hypothetical protein [Nitrogeniibacter aestuarii] | 09172 | 3 | 43% |
| PliI family lysozyme inhibitor of I-type lysozyme [Nitrogeniibacter aestuarii] | 00627 | 3 | 43% |
| hypothetical protein [Nitrogeniibacter aestuarii] | 01308 | 3 | 43% |
| DUF1269 domain-containing protein [Nitrogeniibacter aestuarii] | 02492 | 3 | 43% |
| hypothetical protein [Nitrogeniibacter aestuarii] | 09778 | 3 | 43% |
| HAMP domain-containing histidine kinase [Denitromonas halophilus] | 07880 | 3 | 43% |
| hypothetical protein [Denitromonas halophilus] | 03249 | 3 | 43% |
| hypothetical protein [Nitrogeniibacter aestuarii] | 06605 | 3 | 43% |
| type VI secretion system Vgr family protein, partial [Denitromonas halophilus] | 05900 | 3 | 43% |
| hypothetical protein [Denitromonas halophilus] | 10421 | 3 | 43% |
| membrane lipoprotein lipid attachment site-containing protein [Nitrogeniibacter aestuarii] | 10599 | 3 | 43% |
| prepilin-type N-terminal cleavage/methylation domain-containing protein [Denitromonas halophilus] | 00935 | 3 | 43% |
| cupin domain-containing protein [Denitromonas halophilus] | 02322 | 3 | 43% |
| hypothetical protein [Nitrogeniibacter aestuarii] | 06455 | 3 | 43% |
| hypothetical protein [Nitrogeniibacter aestuarii] | 04245 | 3 | 43% |
| alginate export family protein [Nitrogeniibacter aestuarii] | 01272 | 3 | 43% |
| DUF2195 family protein [Nitrogeniibacter aestuarii] | 08034 | 3 | 43% |
| hypothetical protein [Nitrogeniibacter aestuarii] | 00605 | 3 | 43% |
| GAF domain-containing protein [Nitrogeniibacter aestuarii] | 08654 | 3 | 43% |
| DUF1538 domain-containing protein [Nitrogeniibacter aestuarii] | 00739 | 3 | 43% |
| hypothetical protein [Nitrogeniibacter aestuarii] | 08263 | 3 | 43% |
| penicillin-insensitive murein endopeptidase [Nitrogeniibacter aestuarii] | 01561 | 3 | 43% |
| hypothetical protein [Nitrogeniibacter aestuarii] | 09207 | 3 | 43% |
| PEP-CTERM sorting domain-containing protein [Denitromonas halophilus] | 06419 | 3 | 43% |
| GGDEF domain-containing protein [Denitromonas halophilus] | 03934 | 3 | 43% |
| hypothetical protein [Nitrogeniibacter aestuarii] | 03080 | 3 | 43% |
| hypothetical protein [Denitromonas halophilus] | 10414 | 3 | 43% |
| FAD-binding protein [Denitromonas halophilus] | 06417 | 3 | 43% |
| hypothetical protein [Denitromonas halophilus] | 06072 | 3 | 43% |
| hypothetical protein [Denitromonas halophilus] | 01557 | 3 | 43% |
| FecR family protein [Nitrogeniibacter aestuarii] | 06218 | 3 | 43% |
| DUF3365 domain-containing protein [Nitrogeniibacter aestuarii] | 05287 | 3 | 43% |
| PEP-CTERM sorting domain-containing protein [Nitrogeniibacter aestuarii] | 02699 | 3 | 43% |
| hypothetical protein [Nitrogeniibacter aestuarii] | 09318 | 3 | 43% |
| hypothetical protein [Nitrogeniibacter aestuarii] | 05208 | 3 | 43% |
| hypothetical protein [Nitrogeniibacter aestuarii] | 09356 | 3 | 43% |
| hypothetical protein [Nitrogeniibacter aestuarii] | 06200 | 3 | 43% |
| hypothetical protein [Nitrogeniibacter aestuarii] | 07905 | 3 | 43% |
| chlorite dismutase family protein [Denitromonas halophilus] | 07878 | 3 | 43% |
| GFA family protein [Nitrogeniibacter aestuarii] | 01404 | 3 | 43% |
| antibiotic biosynthesis monooxygenase [Denitromonas halophilus] | 09944 | 3 | 43% |
| PEP-CTERM sorting domain-containing protein [Nitrogeniibacter aestuarii] | 07641 | 3 | 43% |
| hypothetical protein [Nitrogeniibacter aestuarii] | 09967 | 3 | 43% |
| GNAT family N-acetyltransferase [Denitromonas halophilus] | 10094 | 3 | 43% |
| hypothetical protein [Denitromonas halophilus] | 05174 | 3 | 43% |

| | | | |
|---|---|---|---|
| hypothetical protein [Nitrogeniibacter aestuarii] | 10064 | 3 | 43% |
| hypothetical protein [Denitromonas halophilus] | 05491 | 3 | 43% |
| hypothetical protein [Nitrogeniibacter aestuarii] | 09348 | 3 | 43% |
| cytochrome c family protein [Denitromonas halophilus] | 08536 | 3 | 43% |
| hypothetical protein [Nitrogeniibacter aestuarii] | 03558 | 2 | 29% |
| hypothetical protein [Nitrogeniibacter aestuarii] | 01949 | 2 | 29% |
| hypothetical protein [Nitrogeniibacter aestuarii] | 08049 | 2 | 29% |
| hypothetical protein [Nitrogeniibacter aestuarii] | 10067 | 2 | 29% |
| hypothetical protein [Nitrogeniibacter aestuarii] | 03584 | 2 | 29% |
| hypothetical protein [Nitrogeniibacter aestuarii] | 00745 | 2 | 29% |
| hypothetical protein [Nitrogeniibacter aestuarii] | 10095 | 2 | 29% |
| hypothetical protein [Nitrogeniibacter aestuarii] | 04991 | 2 | 29% |
| hypothetical protein [Nitrogeniibacter aestuarii] | 02255 | 2 | 29% |
| AHH domain-containing protein [Nitrogeniibacter aestuarii] | 05892 | 2 | 29% |
| hypothetical protein [Nitrogeniibacter aestuarii] | 04455 | 2 | 29% |
| hypothetical protein [Nitrogeniibacter aestuarii] | 08950 | 2 | 29% |
| hypothetical protein [Nitrogeniibacter aestuarii] | 08957 | 2 | 29% |
| DUF2976 domain-containing protein [Nitrogeniibacter aestuarii] | 03094 | 2 | 29% |
| hypothetical protein [Nitrogeniibacter aestuarii] | 06202 | 2 | 29% |
| AAA family ATPase [Nitrogeniibacter aestuarii] | 04454 | 2 | 29% |
| hypothetical protein [Nitrogeniibacter aestuarii] | 06244 | 2 | 29% |
| hypothetical protein [Denitromonas halophilus] | 00081 | 2 | 29% |
| acyloxyacyl hydrolase [Denitromonas sp. IR12] | 06423 | 2 | 29% |
| hypothetical protein [Denitromonas sp. IR12] | 10100 | 2 | 29% |
| hypothetical protein [Denitromonas sp. IR12] | 01312 | 2 | 29% |
| DUF695 domain-containing protein [Denitromonas sp. IR12] | 09352 | 2 | 29% |
| DUF2345 domain-containing protein, partial [Denitromonas sp. IR12] | 03589 | 2 | 29% |
| hypothetical protein [Denitromonas sp. IR12] | 02126 | 2 | 29% |
| hypothetical protein [Denitromonas sp. IR12] | 07883 | 2 | 29% |
| hypothetical protein [Denitromonas sp. IR12] | 04977 | 2 | 29% |
| TAXI family TRAP transporter solute-binding subunit, partial [Denitromonas sp. IR12] | 09350 | 2 | 29% |
| cupin domain-containing protein [Denitromonas sp. IR12] | 10240 | 2 | 29% |
| hypothetical protein [Denitromonas sp. IR12] | 02974 | 2 | 29% |
| hypothetical protein [Denitromonas sp. IR12] | 07690 | 2 | 29% |
| O-antigen ligase family protein [Denitromonas sp. IR12] | 02317 | 2 | 29% |
| PEP-CTERM sorting domain-containing protein [Denitromonas sp. IR12] | 05921 | 2 | 29% |
| hypothetical protein [Denitromonas sp. IR12] | 02316 | 2 | 29% |
| IS3 family transposase, partial [Denitromonas halophilus] | 03921 | 2 | 29% |
| HAD hydrolase-like protein [Nitrogeniibacter mangrovi] | 03965 | 2 | 29% |
| polysaccharide biosynthesis PFTS motif protein [Nitrogeniibacter mangrovi] | 10420 | 2 | 29% |
| hypothetical protein [Nitrogeniibacter mangrovi] | 06931 | 2 | 29% |
| glycosyltransferase family 2 protein [Nitrogeniibacter mangrovi] | 00606 | 2 | 29% |
| hypothetical protein [Nitrogeniibacter mangrovi] | 02994 | 2 | 29% |
| ATP-binding domain-containing protein [Denitromonas halophilus] | 09178 | 2 | 29% |
| hypothetical protein [Denitromonas halophilus] | 08835 | 2 | 29% |
| hypothetical protein [Denitromonas halophilus] | 04812 | 2 | 29% |
| FAD-dependent oxidoreductase, partial [Denitromonas halophilus] | 03928 | 2 | 29% |
| hypothetical protein [Denitromonas halophilus] | 00939 | 2 | 29% |
| phosphocholine cytidylyltransferase family protein [Denitromonas halophilus] | 06949 | 2 | 29% |
| pyruvate phosphate dikinase [Denitromonas halophilus] | 06947 | 2 | 29% |
| hypothetical protein [Nitrogeniibacter aestuarii] | 07335 | 2 | 29% |
| hypothetical protein [Nitrogeniibacter aestuarii] | 08015 | 2 | 29% |
| DUF695 domain-containing protein [Nitrogeniibacter aestuarii] | 04990 | 2 | 29% |
| hypothetical protein [Nitrogeniibacter aestuarii] | 10250 | 2 | 29% |
| hypothetical protein [Nitrogeniibacter aestuarii] | 00748 | 2 | 29% |
| sulfotransferase domain-containing protein [Nitrogeniibacter aestuarii] | 10601 | 2 | 29% |
| hypothetical protein [Nitrogeniibacter aestuarii] | 07642 | 2 | 29% |
| hypothetical protein [Nitrogeniibacter aestuarii] | 04564 | 2 | 29% |
| replication initiation factor [Nitrogeniibacter aestuarii] | 00240 | 2 | 29% |
| hypothetical protein [Nitrogeniibacter aestuarii] | 03090 | 2 | 29% |
| S1 family peptidase [Nitrogeniibacter aestuarii] | 07629 | 2 | 29% |
| PEP-CTERM sorting domain-containing protein [Nitrogeniibacter aestuarii] | 03089 | 2 | 29% |
| hypothetical protein [Nitrogeniibacter aestuarii] | 06601 | 2 | 29% |
| PEP-CTERM sorting domain-containing protein [Nitrogeniibacter aestuarii] | 06560 | 2 | 29% |
| hypothetical protein [Nitrogeniibacter aestuarii] | 04563 | 2 | 29% |
| hypothetical protein [Nitrogeniibacter aestuarii] | 10600 | 2 | 29% |
| hypothetical protein [Nitrogeniibacter aestuarii] | 07640 | 2 | 29% |
| hypothetical protein [Nitrogeniibacter aestuarii] | 01952 | 2 | 29% |
| hypothetical protein [Nitrogeniibacter aestuarii] | 06246 | 2 | 29% |
| hypothetical protein [Nitrogeniibacter aestuarii] | 03116 | 2 | 29% |
| hypothetical protein [Nitrogeniibacter aestuarii] | 09781 | 2 | 29% |
| cache domain-containing protein [Nitrogeniibacter aestuarii] | 10602 | 2 | 29% |
| hypothetical protein [Nitrogeniibacter aestuarii] | 05929 | 2 | 29% |
| hypothetical protein [Nitrogeniibacter aestuarii] | 06247 | 2 | 29% |
| DUF4157 domain-containing protein [Nitrogeniibacter aestuarii] | 06775 | 2 | 29% |
| DUF2314 domain-containing protein [Nitrogeniibacter aestuarii] | 06602 | 2 | 29% |
| response regulator [Nitrogeniibacter aestuarii] | 08652 | 2 | 29% |
| DUF1566 domain-containing protein [Nitrogeniibacter aestuarii] | 06776 | 2 | 29% |
| hypothetical protein [Nitrogeniibacter aestuarii] | 07091 | 2 | 29% |
| DUF1127 domain-containing protein [Nitrogeniibacter aestuarii] | 05930 | 2 | 29% |
| hypothetical protein [Nitrogeniibacter aestuarii] | 06777 | 2 | 29% |
| CGNR zinc finger domain-containing protein [Nitrogeniibacter aestuarii] | 06568 | 2 | 29% |
| hypothetical protein [Nitrogeniibacter aestuarii] | 04983 | 2 | 29% |
| ATP-binding cassette domain-containing protein [Nitrogeniibacter aestuarii] | 10603 | 2 | 29% |
| DUF4240 domain-containing protein [Nitrogeniibacter aestuarii] | 07663 | 2 | 29% |
| hypothetical protein [Nitrogeniibacter aestuarii] | 07333 | 2 | 29% |
| hypothetical protein [Nitrogeniibacter aestuarii] | 09320 | 2 | 29% |

| | | | |
|---|---|---|---|
| hypothetical protein [Nitrogeniibacter aestuarii] | 04103 | 2 | 29% |
| hypothetical protein [Nitrogeniibacter aestuarii] | 09627 | 2 | 29% |
| roadblock/LC7 domain-containing protein [Nitrogeniibacter aestuarii] | 04595 | 2 | 29% |
| DUF1206 domain-containing protein [Nitrogeniibacter aestuarii] | 00750 | 2 | 29% |
| hypothetical protein [Nitrogeniibacter aestuarii] | 09630 | 2 | 29% |
| hypothetical protein [Nitrogeniibacter aestuarii] | 02459 | 2 | 29% |
| hypothetical protein [Nitrogeniibacter aestuarii] | 10245 | 2 | 29% |
| hypothetical protein [Nitrogeniibacter aestuarii] | 05894 | 2 | 29% |
| DUF4157 domain-containing protein [Nitrogeniibacter aestuarii] | 04566 | 2 | 29% |
| hypothetical protein [Nitrogeniibacter aestuarii] | 04274 | 2 | 29% |
| hypothetical protein [Nitrogeniibacter aestuarii] | 08043 | 2 | 29% |
| hypothetical protein [Nitrogeniibacter aestuarii] | 00937 | 2 | 29% |
| hypothetical protein [Nitrogeniibacter aestuarii] | 01280 | 2 | 29% |
| OmpA family protein [Nitrogeniibacter aestuarii] | 09777 | 2 | 29% |
| hypothetical protein [Nitrogeniibacter aestuarii] | 08397 | 2 | 29% |
| hypothetical protein [Nitrogeniibacter aestuarii] | 03081 | 2 | 29% |
| fatty acid desaturase [Nitrogeniibacter aestuarii] | 04594 | 2 | 29% |
| PEP-CTERM sorting domain-containing protein [Nitrogeniibacter aestuarii] | 00781 | 2 | 29% |
| hypothetical protein [Nitrogeniibacter aestuarii] | 07693 | 2 | 29% |
| hypothetical protein [Nitrogeniibacter aestuarii] | 00198 | 2 | 29% |
| hypothetical protein [Nitrogeniibacter aestuarii] | 07643 | 2 | 29% |
| hypothetical protein [Nitrogeniibacter aestuarii] | 01975 | 2 | 29% |
| hypothetical protein [Nitrogeniibacter aestuarii] | 04980 | 2 | 29% |
| toll/interleukin-1 receptor domain-containing protein [Nitrogeniibacter aestuarii] | 07694 | 2 | 29% |
| thiol-activated cytolysin family protein [Nitrogeniibacter aestuarii] | 04076 | 2 | 29% |
| methyltransferase domain-containing protein [Nitrogeniibacter aestuarii] | 10218 | 2 | 29% |
| hypothetical protein [Nitrogeniibacter aestuarii] | 04275 | 2 | 29% |
| TPM domain-containing protein [Nitrogeniibacter aestuarii] | 04596 | 2 | 29% |
| nucleotidyltransferase family protein [Nitrogeniibacter aestuarii] | 08016 | 2 | 29% |
| M23 family metallopeptidase [Nitrogeniibacter aestuarii] | 07337 | 2 | 29% |
| PEP-CTERM sorting domain-containing protein [Nitrogeniibacter aestuarii] | 10065 | 2 | 29% |
| HEAT repeat domain-containing protein [Nitrogeniibacter aestuarii] | 07639 | 2 | 29% |
| DUF1444 domain-containing protein [Nitrogeniibacter aestuarii] | 01951 | 2 | 29% |
| mechanosensitive ion channel family protein [Nitrogeniibacter aestuarii] | 03559 | 2 | 29% |
| hypothetical protein [Nitrogeniibacter aestuarii] | 10063 | 2 | 29% |
| hypothetical protein [Nitrogeniibacter aestuarii] | 06221 | 2 | 29% |
| hypothetical protein [Nitrogeniibacter aestuarii] | 00744 | 2 | 29% |
| hypothetical protein [Nitrogeniibacter aestuarii] | 00742 | 2 | 29% |
| CDP-glycerol glycerophosphotransferase family protein [Nitrogeniibacter aestuarii] | 09629 | 2 | 29% |
| hypothetical protein [Nitrogeniibacter aestuarii] | 01950 | 2 | 29% |
| MFS transporter [Nitrogeniibacter aestuarii] | 08951 | 2 | 29% |
| aminotransferase class V-fold PLP-dependent enzyme [Nitrogeniibacter aestuarii] | 09358 | 2 | 29% |
| DUF3421 domain-containing protein [Nitrogeniibacter aestuarii] | 05932 | 2 | 29% |
| DUF3421 domain-containing protein [Nitrogeniibacter aestuarii] | 01523 | 2 | 29% |
| DUF1800 domain-containing protein [Nitrogeniibacter aestuarii] | 08047 | 2 | 29% |
| hypothetical protein [Nitrogeniibacter aestuarii] | 10565 | 2 | 29% |
| hypothetical protein [Nitrogeniibacter aestuarii] | 04989 | 2 | 29% |
| hypothetical protein [Nitrogeniibacter aestuarii] | 04279 | 2 | 29% |
| hypothetical protein [Nitrogeniibacter aestuarii] | 06598 | 2 | 29% |
| hypothetical protein [Nitrogeniibacter aestuarii] | 10605 | 2 | 29% |
| hypothetical protein [Nitrogeniibacter aestuarii] | 05889 | 2 | 29% |
| barstar family protein [Nitrogeniibacter aestuarii] | 06223 | 2 | 29% |
| SRPBCC family protein [Nitrogeniibacter aestuarii] | 04988 | 2 | 29% |
| hypothetical protein [Nitrogeniibacter aestuarii] | 01323 | 2 | 29% |
| hypothetical protein [Nitrogeniibacter aestuarii] | 00194 | 2 | 29% |
| DUF695 domain-containing protein [Nitrogeniibacter aestuarii] | 07638 | 2 | 29% |
| hypothetical protein [Nitrogeniibacter aestuarii] | 08371 | 2 | 29% |
| hypothetical protein [Nitrogeniibacter aestuarii] | 05933 | 2 | 29% |
| hypothetical protein [Nitrogeniibacter aestuarii] | 10249 | 2 | 29% |
| hypothetical protein [Nitrogeniibacter aestuarii] | 10606 | 2 | 29% |
| hypothetical protein [Nitrogeniibacter aestuarii] | 01324 | 2 | 29% |
| hypothetical protein [Nitrogeniibacter aestuarii] | 06204 | 2 | 29% |
| hypothetical protein [Nitrogeniibacter aestuarii] | 01979 | 2 | 29% |
| hypothetical protein [Nitrogeniibacter aestuarii] | 02456 | 2 | 29% |
| DUF4265 domain-containing protein [Nitrogeniibacter aestuarii] | 08048 | 2 | 29% |
| PQQ-binding-like beta-propeller repeat protein [Nitrogeniibacter aestuarii] | 03556 | 2 | 29% |
| YidC/Oxa1 family membrane protein insertase [Nitrogeniibacter aestuarii] | 01322 | 2 | 29% |
| hypothetical protein [Nitrogeniibacter aestuarii] | 01283 | 2 | 29% |
| hypothetical protein [Nitrogeniibacter aestuarii] | 10563 | 2 | 29% |
| thiol-disulfide oxidoreductase DCC family protein [Nitrogeniibacter aestuarii] | 09357 | 2 | 29% |
| hypothetical protein [Nitrogeniibacter aestuarii] | 04986 | 2 | 29% |
| hypothetical protein [Nitrogeniibacter aestuarii] | 04599 | 2 | 29% |
| PEP-CTERM sorting domain-containing protein [Nitrogeniibacter aestuarii] | 06573 | 2 | 29% |
| hypothetical protein [Nitrogeniibacter aestuarii] | 06222 | 2 | 29% |
| ABZJ_00895 family protein [Nitrogeniibacter aestuarii] | 10104 | 2 | 29% |
| hypothetical protein [Nitrogeniibacter aestuarii] | 10217 | 2 | 29% |
| hypothetical protein [Nitrogeniibacter aestuarii] | 04984 | 2 | 29% |
| hypothetical protein [Nitrogeniibacter aestuarii] | 10560 | 2 | 29% |
| PEP-CTERM sorting domain-containing protein [Nitrogeniibacter aestuarii] | 04276 | 2 | 29% |
| hypothetical protein [Nitrogeniibacter aestuarii] | 06603 | 2 | 29% |
| sulfotransferase [Nitrogeniibacter aestuarii] | 01948 | 2 | 29% |
| hypothetical protein [Nitrogeniibacter aestuarii] | 09745 | 2 | 29% |
| hypothetical protein [Nitrogeniibacter aestuarii] | 04600 | 2 | 29% |
| serine/threonine-protein phosphatase [Nitrogeniibacter aestuarii] | 01320 | 2 | 29% |
| flavodoxin family protein [Nitrogeniibacter aestuarii] | 03084 | 2 | 29% |
| hypothetical protein [Nitrogeniibacter aestuarii] | 01281 | 2 | 29% |
| tryptophan-rich sensory protein [Nitrogeniibacter aestuarii] | 01522 | 2 | 29% |

| | | | |
|---|---|---|---|
| hypothetical protein [Nitrogeniibacter aestuarii] | 09626 | 2 | 29% |
| GNAT family N-acetyltransferase [Nitrogeniibacter aestuarii] | 07665 | 2 | 29% |
| hypothetical protein [Nitrogeniibacter aestuarii] | 10103 | 2 | 29% |
| hypothetical protein [Nitrogeniibacter aestuarii] | 02796 | 2 | 29% |
| hypothetical protein [Nitrogeniibacter aestuarii] | 04948 | 2 | 29% |
| hypothetical protein [Nitrogeniibacter aestuarii] | 06571 | 2 | 29% |
| hypothetical protein [Denitromonas sp. IR12] | 02229 | 1 | 14% |
| helix-turn-helix transcriptional regulator [Denitromonas sp. IR12] | 00222 | 1 | 14% |
| DUF2975 domain-containing protein [Denitromonas sp. IR12] | 04383 | 1 | 14% |
| DUF2345 domain-containing protein, partial [Denitromonas sp. IR12] | 04978 | 1 | 14% |
| hypothetical protein [Denitromonas halophilus] | 07521 | 1 | 14% |
| hypothetical protein [Denitromonas sp. IR12] | 05899 | 1 | 14% |
| hypothetical protein [Nitrogeniibacter aestuarii] | 06599 | 1 | 14% |
| hypothetical protein [Niveibacterium sp. COAC-50] | 04431 | 1 | 14% |
| hypothetical protein [Denitromonas halophilus] | 09141 | 1 | 14% |
| hypothetical protein [Azospira oryzae] | 04819 | 1 | 14% |
| cadherin domain-containing protein, partial [Denitromonas sp. IR12] | 03266 | 1 | 14% |
| phage late control D family protein, partial [Denitromonas sp. IR12] | 03590 | 1 | 14% |
| hypothetical protein, partial [Denitromonas sp. IR12] | 07692 | 1 | 14% |
| Ig-like domain-containing protein, partial [Denitromonas sp. IR12] | 04591 | 1 | 14% |
| hypothetical protein, partial [Denitromonas sp. IR12] | 03267 | 1 | 14% |
| hypothetical protein [Denitromonas sp. IR12] | 09280 | 1 | 14% |
| AMP-binding protein, partial [Denitromonas sp. IR12] | 00779 | 1 | 14% |
| AMP-binding protein, partial [Denitromonas sp. IR12] | 04592 | 1 | 14% |
| hypothetical protein [Niveibacterium sp. COAC-50] | 02673 | 1 | 14% |
| condensation domain-containing protein, partial [Denitromonas sp. IR12] | 02486 | 1 | 14% |
| hypothetical protein [Niveibacterium sp. COAC-50] | 09714 | 1 | 14% |
| transposase, partial [Denitromonas sp. IR12] | 03113 | 1 | 14% |
| SEL1-like repeat protein [Nitrogeniibacter aestuarii] | 06201 | 1 | 14% |
| hypothetical protein, partial [Denitromonas sp. IR12] | 01560 | 1 | 14% |
| hypothetical protein [Nitrogeniibacter mangrovi] | 04457 | 1 | 14% |
| retention module-containing protein, partial [Nitrogeniibacter aestuarii] | 04105 | 1 | 14% |
| hypothetical protein [Propionivibrio dicarboxylicus] | 06782 | 1 | 14% |
| hypothetical protein [Propionivibrio dicarboxylicus] | 01750 | 1 | 14% |
| ThiF family adenylyltransferase [Nitrogeniibacter aestuarii] | 07094 | 1 | 14% |
| phospholipase D-like domain-containing protein [Nitrogeniibacter aestuarii] | 08332 | 1 | 14% |
| hypothetical protein [Rhodocyclus purpureus] | 08540 | 1 | 14% |
| DUF2190 family protein, partial [Rhodocyclus purpureus] | 05755 | 1 | 14% |
| hypothetical protein [Nitrogeniibacter aestuarii] | 02223 | 1 | 14% |
| Hint domain-containing protein [Nitrogeniibacter aestuarii] | 05247 | 1 | 14% |
| hypothetical protein [Nitrogeniibacter aestuarii] | 10102 | 1 | 14% |
| cytochrome C [Nitrogeniibacter mangrovi] | 04958 | 1 | 14% |
| hypothetical protein [Nitrogeniibacter mangrovi] | 08955 | 1 | 14% |
| homogentisate 1,2-dioxygenase [Nitrogeniibacter mangrovi] | 00629 | 1 | 14% |
| hypothetical protein [Nitrogeniibacter aestuarii] | 08946 | 1 | 14% |
| M48 family metallopeptidase [Nitrogeniibacter mangrovi] | 02270 | 1 | 14% |
| hypothetical protein [Nitrogeniibacter mangrovi] | 00192 | 1 | 14% |
| hypothetical protein, partial [Nitrogeniibacter aestuarii] | 07696 | 1 | 14% |
| hypothetical protein [Denitromonas sp. IR12] | 04269 | 1 | 14% |
| hypothetical protein [Denitromonas sp. IR12] | 05309 | 1 | 14% |
| 2OG-Fe(II) oxygenase [Denitromonas sp. IR12] | 04567 | 1 | 14% |
| helix-turn-helix domain-containing protein, partial [Denitromonas sp. IR12] | 05920 | 1 | 14% |
| transposase domain-containing protein, partial [Denitromonas sp. IR12] | 10243 | 1 | 14% |
| phage portal protein [Denitromonas sp. IR12] | 05604 | 1 | 14% |
| ABC transporter permease [Denitromonas sp. IR12] | 08394 | 1 | 14% |
| glycosyltransferase [Denitromonas sp. IR12] | 04100 | 1 | 14% |
| hypothetical protein [Nitrogeniibacter aestuarii] | 06744 | 1 | 14% |
| hypothetical protein, partial [Denitromonas halophilus] | 01728 | 1 | 14% |

Table S5: Protein subfamilies missing from Denitromonas spp. and close relatives

| Gene Product Name | Cluster | Genome Cou |
|---|---|---|
| hypothetical protein [Aromatoleum aromaticum] | 07015 | |
| U32 family peptidase [Thauera linaloolentis] | 05636 | |
| bifunctional DNA-binding transcriptional regulator/O6-methylguanine-DNA methyltransferase Ada [Thauera linaloolentis] | 10160 | |
| membrane protein [Aromatoleum aromaticum] | 02757 | |
| site-specific recombinase [Aromatoleum aromaticum] | 08438 | |
| hypothetical protein [Aromatoleum aromaticum] | 01648 | |
| DUF2946 domain-containing protein [Aromatoleum aromaticum] | 04652 | |
| glycosyltransferase [Aromatoleum aromaticum] | 01406 | |
| beta-1,6-glucan synthase [Aromatoleum aromaticum] | 03755 | |
| AI-2E family transporter [Thauera linaloolentis] | 07403 | |
| hypothetical protein [Aromatoleum aromaticum] | 07353 | |
| glucokinase [Aromatoleum aromaticum] | 00067 | |
| DNA-3-methyladenine glycosylase 2 [Aromatoleum aromaticum] | 08643 | |
| hypothetical protein [Thauera linaloolentis] | 05734 | |
| hypothetical protein [Thauera linaloolentis] | 00617 | |
| glycosyltransferase family 2 protein [Thauera linaloolentis] | 04504 | |
| hypothetical protein [Aromatoleum aromaticum] | 05245 | |
| Na+/H+ antiporter subunit E [Aromatoleum aromaticum] | 02470 | |
| MULTISPECIES: monovalent cation/H(+) antiporter subunit G [Aromatoleum] | 06212 | |
| hypothetical protein [Thauera linaloolentis] | 03005 | |
| PilZ domain-containing protein [Azoarcus indigens] | 09800 | |
| hypothetical protein [Aromatoleum aromaticum] | 05267 | |
| nucleoid-associated protein [Thauera linaloolentis] | 09496 | |
| hypothetical protein [Aromatoleum aromaticum] | 00338 | |
| hypothetical protein [Aromatoleum aromaticum] | 09959 | |
| metal-dependent hydrolase [Aromatoleum aromaticum] | 05456 | |
| hypothetical protein [Aromatoleum aromaticum] | 08637 | |
| EcsC family protein [Aromatoleum aromaticum] | 07377 | |
| LTA synthase family protein [Aromatoleum aromaticum] | 04232 | |
| lipopolysaccharide core heptose(I) kinase RfaP [Aromatoleum aromaticum] | 05462 | |
| ABC transporter permease [Aromatoleum aromaticum] | 03661 | |
| FHA domain-containing protein [Aromatoleum aromaticum] | 10550 | |
| GtrA family protein [Thauera linaloolentis] | 09254 | |
| phosphotransferase [Aromatoleum aromaticum] | 08284 | |
| NAD(P)/FAD-dependent oxidoreductase [Aromatoleum aromaticum] | 01257 | |
| HAD family hydrolase [Aromatoleum aromaticum] | 01758 | |
| hypothetical protein [Thauera linaloolentis] | 10355 | |
| LysR family transcriptional regulator [Azoarcus indigens] | 01291 | |
| 2,3-bisphosphoglycerate-independent phosphoglycerate mutase [Thauera linaloolentis] | 00030 | |
| hypothetical protein [Thauera linaloolentis] | 05828 | |
| superoxide dismutase family protein [Thauera linaloolentis] | 09431 | |
| L-lactate permease [Thauera linaloolentis] | 09017 | |
| GNAT family N-acetyltransferase [Aromatoleum aromaticum] | 02376 | |
| lipocalin family protein [Azoarcus indigens] | 07836 | |
| hypothetical protein [Thauera linaloolentis] | 03654 | |
| MULTISPECIES: hypothetical protein [Aromatoleum] | 06407 | |
| glycoside hydrolase family 5 protein [Azoarcus indigens] | 03120 | |
| tetratricopeptide repeat protein [Thauera linaloolentis] | 09398 | |
| EAL domain-containing protein [Thauera linaloolentis] | 07594 | |
| type II toxin-antitoxin system VapC family toxin [Aromatoleum aromaticum] | 00525 | |
| fused MFS/spermidine synthase [Aromatoleum aromaticum] | 08615 | |
| acetyl-CoA hydrolase/transferase family protein [Thauera linaloolentis] | 08187 | |
| hypothetical protein [Thauera linaloolentis] | 04480 | |
| ABC transporter substrate-binding protein [Thauera linaloolentis] | 08929 | |
| DUF4124 domain-containing protein [Aromatoleum aromaticum] | 05141 | |
| DUF423 domain-containing protein [Thauera linaloolentis] | 02600 | |
| transglutaminase [Aromatoleum aromaticum] | 05773 | |

| | |
|---|---|
| InaA protein [Thauera linaloolentis] | 01773 |
| putative addiction module antidote protein [Aromatoleum aromaticum] | 04748 |
| ChbG/HpnK family deacetylase [Azoarcus indigens] | 00208 |
| glycosyltransferase family 39 protein [Azoarcus indigens] | 01624 |
| outer membrane protein assembly factor BamE [Aromatoleum aromaticum] | 01225 |
| 4-hydroxybenzoyl-CoA reductase subunit beta [Aromatoleum aromaticum] | 04749 |
| phenylacetic acid degradation operon negative regulatory protein PaaX [Aromatoleum aromaticum] | 02017 |
| VOC family protein [Thauera linaloolentis] | 00299 |
| hypothetical protein [Aromatoleum aromaticum] | 07674 |
| dimethyl sulfoxide reductase anchor subunit [Aromatoleum aromaticum] | 00651 |
| DUF3334 family protein [Thauera linaloolentis] | 01770 |
| hypothetical protein [Aromatoleum tolulyticum] | 01181 |
| ABC-type transport auxiliary lipoprotein family protein [Thauera linaloolentis] | 05738 |
| hypothetical protein [Aromatoleum aromaticum] | 08887 |
| cysteine-rich CWC family protein [Aromatoleum aromaticum] | 00356 |
| polysaccharide pyruvyl transferase family protein [Aromatoleum aromaticum] | 00851 |
| hypothetical protein [Thauera linaloolentis] | 05116 |
| TolC family protein [Azoarcus indigens] | 02176 |
| PAS domain-containing protein [Azoarcus indigens] | 06696 |
| MFS transporter [Aromatoleum aromaticum] | 01965 |
| TetR/AcrR family transcriptional regulator [Aromatoleum aromaticum] | 02642 |
| nicotinate phosphoribosyltransferase [Thauera chlorobenzoica] | 08159 |
| YrbL family protein [Aromatoleum aromaticum] | 02681 |
| O-antigen ligase family protein [Aromatoleum aromaticum] | 01299 |
| aminoacyl-histidine dipeptidase [Azoarcus indigens] | 08221 |
| hypothetical protein [Aromatoleum aromaticum] | 03134 |
| DNA oxidative demethylase AlkB [Azoarcus indigens] | 04885 |
| YbdK family carboxylate-amine ligase [Azoarcus indigens] | 10437 |
| amino acid ABC transporter substrate-binding protein [Thauera linaloolentis] | 07443 |
| acyloxyacyl hydrolase [Aromatoleum aromaticum] | 00678 |
| DMT family transporter [Aromatoleum tolulyticum] | 07421 |
| CynX/NimT family MFS transporter [Aromatoleum aromaticum] | 07770 |
| type II toxin-antitoxin system PrlF family antitoxin [Aromatoleum aromaticum] | 04214 |
| hypothetical protein [Aromatoleum aromaticum] | 07219 |
| polysaccharide pyruvyl transferase family protein [Aromatoleum aromaticum] | 04750 |
| type II toxin-antitoxin system Phd/YefM family antitoxin [Aromatoleum aromaticum] | 08300 |
| class I SAM-dependent methyltransferase [Thauera linaloolentis] | 00534 |
| ParD-like family protein [Thauera linaloolentis] | 04516 |
| SdiA-regulated domain-containing protein [Azoarcus indigens] | 07826 |
| hypothetical protein [Thauera linaloolentis] | 05567 |
| hypothetical protein [Aromatoleum aromaticum] | 01088 |
| cytochrome c [Aromatoleum aromaticum] | 06393 |
| MULTISPECIES: hypothetical protein [Aromatoleum] | 07569 |
| hypothetical protein [Aromatoleum aromaticum] | 00914 |
| hypothetical protein [Aromatoleum aromaticum] | 04823 |
| 2-oxoacid:acceptor oxidoreductase family protein [Aromatoleum aromaticum] | 01778 |
| 2'-5' RNA ligase family protein [Thauera linaloolentis] | 10050 |
| phosphatase PAP2 family protein [Azoarcus indigens] | 01479 |
| hypothetical protein [Thauera linaloolentis] | 04390 |
| GatB/YqeY domain-containing protein [Thauera linaloolentis] | 03876 |
| glycogen synthase GlgA [Aromatoleum aromaticum] | 09559 |
| hypothetical protein [Aromatoleum aromaticum] | 08771 |
| 1,4-alpha-glucan branching protein GlgB [Aromatoleum aromaticum] | 01649 |
| Dyp-type peroxidase [Azoarcus indigens] | 00482 |
| methyltransferase domain-containing protein [Azoarcus indigens] | 05748 |
| glycosyltransferase family 2 protein [Aromatoleum aromaticum] | 09396 |
| DUF599 domain-containing protein [Aromatoleum aromaticum] | 10429 |
| type II toxin-antitoxin system prevent-host-death family antitoxin [Aromatoleum aromaticum] | 09850 |
| FAD-binding oxidoreductase [Aromatoleum aromaticum] | 07457 |

| | |
|---|---|
| class I SAM-dependent methyltransferase [Thauera linaloolentis] | 08157 |
| biofilm regulation protein phosphatase SiaA [Aromatoleum aromaticum] | 04696 |
| Txe/YoeB family addiction module toxin [Aromatoleum aromaticum] | 10351 |
| hypothetical protein [Aromatoleum aromaticum] | 09867 |
| glycosyltransferase [Azoarcus indigens] | 08777 |
| mechanosensitive ion channel family protein [Aromatoleum aromaticum] | 04919 |
| hydrogenase maturation protein HypF [Thauera chlorobenzoica] | 08740 |
| flagellar brake protein [Thauera linaloolentis] | 00168 |
| c-type cytochrome [Thauera linaloolentis] | 05142 |
| DNA cytosine methyltransferase [Aromatoleum aromaticum] | 03395 |
| TetR/AcrR family transcriptional regulator [Aromatoleum aromaticum] | 10223 |
| ABC transporter permease [Thauera linaloolentis] | 04769 |
| LysR family transcriptional regulator [Aromatoleum aromaticum] | 06754 |
| U32 family peptidase [Aromatoleum aromaticum] | 03995 |
| DmsE family decaheme c-type cytochrome [Thauera linaloolentis] | 05913 |
| Flp pilus assembly protein CpaB [Aromatoleum aromaticum] | 06720 |
| type II and III secretion system protein family protein [Aromatoleum aromaticum] | 02645 |
| MetQ/NlpA family ABC transporter substrate-binding protein [Thauera linaloolentis] | 05708 |
| hypothetical protein [Thauera linaloolentis] | 00919 |
| alpha-2-macroglobulin family protein [Azoarcus indigens] | 05373 |
| EAL domain-containing response regulator [Aromatoleum aromaticum] | 01542 |
| PDDEXK nuclease domain-containing protein [Thauera linaloolentis] | 07576 |
| malate synthase A [Azoarcus indigens] | 00186 |
| LLM class flavin-dependent oxidoreductase [Thauera linaloolentis] | 07502 |
| hypothetical protein [Azoarcus indigens] | 02055 |
| carbon starvation protein A [Azoarcus indigens] | 07026 |
| rhodanese homology domain-containing protein [Thauera linaloolentis] | 04416 |
| MtrB/PioB family decaheme-associated outer membrane protein [Thauera linaloolentis] | 03547 |
| GNAT family N-acetyltransferase [Aromatoleum aromaticum] | 03973 |
| MFS transporter [Azoarcus indigens] | 04541 |
| PhnD/SsuA/transferrin family substrate-binding protein [Aromatoleum aromaticum] | 04753 |
| helix-turn-helix domain-containing protein [Thauera linaloolentis] | 00847 |
| benzoyl-CoA reductase, bzd-type, subunit Q [Aromatoleum aromaticum] | 01582 |
| Uma2 family endonuclease [Thauera linaloolentis] | 08785 |
| 6-oxocyclohex-1-ene-1-carbonyl-CoA hydratase [Aromatoleum aromaticum] | 02279 |
| MULTISPECIES: 4Fe-4S binding protein [Aromatoleum] | 01604 |
| CoA-binding protein [Aromatoleum aromaticum] | 02276 |
| nicotinamidase [Thauera chlorobenzoica] | 08000 |
| hypothetical protein [Aromatoleum tolulyticum] | 10121 |
| TonB-dependent receptor [Thauera linaloolentis] | 07941 |
| ABC transporter substrate-binding protein [Aromatoleum tolulyticum] | 09097 |
| hypothetical protein [Aromatoleum aromaticum] | 08761 |
| benzoyl-CoA reductase, bzd-type, subunit O [Aromatoleum aromaticum] | 10452 |
| diguanylate cyclase [Azoarcus indigens] | 05799 |
| helix-turn-helix transcriptional regulator [Thauera linaloolentis] | 07927 |
| type II toxin-antitoxin system VapB family antitoxin [Aromatoleum aromaticum] | 03830 |
| carbohydrate ABC transporter permease [Aromatoleum tolulyticum] | 03804 |
| hypothetical protein [Aromatoleum aromaticum] | 01592 |
| benzoyl-CoA reductase, bzd-type, subunit N [Aromatoleum aromaticum] | 07658 |
| hypothetical protein [Azoarcus indigens] | 05069 |
| sugar ABC transporter permease [Aromatoleum tolulyticum] | 05681 |
| type II toxin-antitoxin system CcdA family antitoxin [Aromatoleum aromaticum] | 05797 |
| creatininase [Aromatoleum tolulyticum] | 03350 |
| Zn-ribbon domain-containing OB-fold protein [Aromatoleum aromaticum] | 00836 |
| heavy-metal-associated domain-containing protein [Aromatoleum aromaticum] | 02812 |
| YtxH domain-containing protein [Aromatoleum aromaticum] | 08574 |
| TIGR01212 family radical SAM protein [Thauera chlorobenzoica] | 00981 |
| helix-turn-helix domain-containing protein [Aromatoleum aromaticum] | 09495 |
| thioredoxin domain-containing protein [Aromatoleum tolulyticum] | 05074 |

| | |
|---|---|
| diguanylate cyclase [Thauera linaloolentis] | 00879 |
| flagellar hook-length control protein FliK [Thauera linaloolentis] | 00042 |
| hypothetical protein [Aromatoleum aromaticum] | 01581 |
| prepilin peptidase [Aromatoleum aromaticum] | 00524 |
| aromatic ring-hydroxylating dioxygenase subunit alpha [Azoarcus indigens] | 08027 |
| NAD(P)-dependent oxidoreductase [Azoarcus indigens] | 05246 |
| nitrite reductase small subunit NirD [Thauera linaloolentis] | 03336 |
| pilus assembly protein [Aromatoleum aromaticum] | 02353 |
| histidine kinase [Aromatoleum aromaticum] | 04112 |
| hypothetical protein [Aromatoleum tolulyticum] | 03843 |
| PAS domain-containing protein [Aromatoleum aromaticum] | 04061 |
| ABC transporter permease [Thauera linaloolentis] | 01701 |
| MULTISPECIES: acyl-CoA thioesterase [Aromatoleum] | 09562 |
| VOC family protein [Azoarcus indigens] | 07577 |
| RNA helicase [Aromatoleum aromaticum] | 00076 |
| disulfide bond formation protein B [Aromatoleum tolulyticum] | 05055 |
| DUF3592 domain-containing protein [Aromatoleum aromaticum] | 09556 |
| DEAD/DEAH box helicase [Aromatoleum aromaticum] | 06818 |
| alpha/beta hydrolase [Azoarcus indigens] | 07164 |
| tetratricopeptide repeat protein [Aromatoleum tolulyticum] | 04137 |
| MarR family transcriptional regulator [Aromatoleum aromaticum] | 06966 |
| hypothetical protein [Aromatoleum aromaticum] | 06355 |
| ferritin [Aromatoleum aromaticum] | 03172 |
| dihydrolipoamide acyltransferase [Aromatoleum aromaticum] | 05543 |
| thioredoxin fold domain-containing protein [Thauera linaloolentis] | 03183 |
| glycoside hydrolase family 94 protein [Aromatoleum tolulyticum] | 05283 |
| PilZ domain-containing protein [Azoarcus indigens] | 06359 |
| hypothetical protein [Aromatoleum tolulyticum] | 06883 |
| sel1 repeat family protein [Aromatoleum aromaticum] | 09896 |
| TRAP transporter small permease [Azoarcus indigens] | 03493 |
| META domain-containing protein [Thauera linaloolentis] | 03293 |
| cytochrome oxidase small assembly protein [Aromatoleum aromaticum] | 08671 |
| P-type conjugative transfer protein TrbJ [Aromatoleum aromaticum] | 03471 |
| DUF5610 domain-containing protein [Thauera linaloolentis] | 01895 |
| glycogen debranching protein GlgX [Aromatoleum aromaticum] | 07368 |
| type II toxin-antitoxin system RelB/DinJ family antitoxin [Aromatoleum aromaticum] | 01753 |
| IS21-like element ISAzo4 family transposase [Aromatoleum aromaticum] | 09181 |
| 5-carboxymethyl-2-hydroxymuconate Delta-isomerase [Thauera linaloolentis] | 01064 |
| biofilm regulation phosphoprotein SiaC [Aromatoleum aromaticum] | 09524 |
| biofilm regulation protein kinase SiaB [Aromatoleum aromaticum] | 06366 |
| exo-alpha-sialidase [Azoarcus indigens] | 06445 |
| Flp family type IVb pilin [Aromatoleum aromaticum] | 06487 |
| AAA family ATPase [Thauera linaloolentis] | 01215 |
| HD-GYP domain-containing protein [Thauera linaloolentis] | 04833 |
| formate dehydrogenase accessory protein FdhE [Thauera linaloolentis] | 00060 |
| hypothetical protein [Aromatoleum aromaticum] | 05615 |
| bacterioferritin [Aromatoleum aromaticum] | 03790 |
| DUF3141 domain-containing protein [Aromatoleum aromaticum] | 07602 |
| twitching motility protein PilT [Aromatoleum aromaticum] | 04049 |
| hypothetical protein [Aromatoleum aromaticum] | 04639 |
| methylamine utilization protein [Aromatoleum tolulyticum] | 01848 |
| AlpA family phage regulatory protein [Aromatoleum aromaticum] | 01251 |
| MULTISPECIES: type II toxin-antitoxin system RelE/ParE family toxin [Aromatoleum] | 04550 |
| alpha/beta hydrolase [Aromatoleum tolulyticum] | 01925 |
| DUF2157 domain-containing protein [Thauera linaloolentis] | 07441 |
| DUF4401 domain-containing protein [Thauera linaloolentis] | 08686 |
| type II toxin-antitoxin system YhaV family toxin [Aromatoleum aromaticum] | 00808 |
| MULTISPECIES: molybdopterin-binding protein [Thauera] | 02862 |
| hypothetical protein [Azoarcus indigens] | 07196 |

| | |
|---|---|
| tryptophanase [Aromatoleum aromaticum] | 02731 |
| GspH/FimT family protein [Aromatoleum aromaticum] | 01549 |
| urocanate hydratase [Aromatoleum aromaticum] | 02569 |
| 4a-hydroxytetrahydrobiopterin dehydratase [Aromatoleum aromaticum] | 04336 |
| type II secretion system protein GspK [Aromatoleum aromaticum] | 09758 |
| hypothetical protein [Azoarcus indigens] | 05833 |
| hypothetical protein [Aromatoleum aromaticum] | 01106 |
| hypothetical protein [Thauera linaloolentis] | 09646 |
| YXWGXW repeat-containing protein [Thauera linaloolentis] | 05028 |
| TetR/AcrR family transcriptional regulator [Thauera linaloolentis] | 06311 |
| MULTISPECIES: ion transporter [Proteobacteria] | 00970 |
| ATPase [Aromatoleum aromaticum] | 08683 |
| hypothetical protein [Aromatoleum aromaticum] | 05453 |
| hypothetical protein [Thauera linaloolentis] | 02951 |
| ABC transporter substrate-binding protein [Azoarcus indigens] | 06925 |
| type II secretion system protein GspM [Aromatoleum aromaticum] | 07265 |
| Na+/H+ antiporter subunit E [Thauera chlorobenzoica] | 00371 |
| NF038104 family lipoprotein [Aromatoleum aromaticum] | 04198 |
| AsmA family protein [Aromatoleum aromaticum] | 07117 |
| pyridoxamine 5'-phosphate oxidase family protein [Aromatoleum aromaticum] | 02280 |
| NADH-quinone oxidoreductase subunit B family protein [Aromatoleum aromaticum] | 07613 |
| ABC transporter substrate-binding protein [Aromatoleum tolulyticum] | 01341 |
| MULTISPECIES: formate hydrogenlyase [Aromatoleum] | 09633 |
| DUF2141 domain-containing protein [Thauera linaloolentis] | 02850 |
| PilN domain-containing protein [Aromatoleum aromaticum] | 08459 |
| glycosyltransferase family 39 protein [Azoarcus indigens] | 07591 |
| type II toxin-antitoxin system MqsA family antitoxin [Thauera linaloolentis] | 02849 |
| sulfatase-like hydrolase/transferase [Aromatoleum aromaticum] | 05340 |
| prepilin-type N-terminal cleavage/methylation domain-containing protein [Aromatoleum aromaticum] | 01668 |
| glycoside hydrolase family protein [Aromatoleum aromaticum] | 10113 |
| hypothetical protein [Thauera linaloolentis] | 04741 |
| MULTISPECIES: DeoR/GlpR family DNA-binding transcription regulator [Azoarcus] | 07660 |
| TMAO reductase system periplasmic protein TorT [Aromatoleum tolulyticum] | 01715 |
| PepSY-associated TM helix domain-containing protein [Aromatoleum aromaticum] | 05025 |
| bifunctional 4-hydroxy-2-oxoglutarate aldolase/2-dehydro-3-deoxy-phosphogluconate aldolase [Aromatoleum tolulyticum] | 05858 |
| type II toxin-antitoxin system YafQ family toxin [Aromatoleum tolulyticum] | 01132 |
| cytochrome b/b6 domain-containing protein [Thauera chlorobenzoica] | 06649 |
| acyl-CoA/acyl-ACP dehydrogenase [Thauera linaloolentis] | 06380 |
| ROK family transcriptional regulator [Aromatoleum tolulyticum] | 10034 |
| type II toxin-antitoxin system VapB family antitoxin [Thauera chlorobenzoica] | 05363 |
| alpha-glucan family phosphorylase [Aromatoleum aromaticum] | 05874 |
| antibiotic biosynthesis monooxygenase [Thauera linaloolentis] | 02556 |
| glucose-6-phosphate dehydrogenase [Aromatoleum tolulyticum] | 03998 |
| glycine zipper family protein [Aromatoleum tolulyticum] | 05647 |
| LLM class flavin-dependent oxidoreductase [Thauera linaloolentis] | 08688 |
| pyridoxamine 5'-phosphate oxidase [Thauera linaloolentis] | 01946 |
| sulfatase-like hydrolase/transferase [Thauera linaloolentis] | 02580 |
| MULTISPECIES: YbdD/YjiX family protein [Zoogloeaceae] | 06312 |
| alpha/beta hydrolase [Thauera linaloolentis] | 05497 |
| DUF120 domain-containing protein [Aromatoleum aromaticum] | 08634 |
| phenylphosphate carboxylase subunit gamma [Aromatoleum aromaticum] | 07719 |
| phenylphosphate synthase subunit beta [Aromatoleum aromaticum] | 04110 |
| PEP-utilizing enzyme [Aromatoleum aromaticum] | 01568 |
| tellurite resistance TerB family protein [Thauera linaloolentis] | 06862 |
| HD domain-containing protein [Aromatoleum tolulyticum] | 05212 |
| TMAO reductase system sensor histidine kinase/response regulator TorS [Aromatoleum tolulyticum] | 04106 |
| type II toxin-antitoxin system RelE/ParE family toxin [Aromatoleum aromaticum] | 02540 |
| hypothetical protein [Aromatoleum aromaticum] | 09145 |
| helix-hairpin-helix domain-containing protein [Aromatoleum aromaticum] | 06938 |

| | |
|---|---|
| SoxR reducing system RseC family protein [Azoarcus indigens] | 01678 |
| hypothetical protein [Aromatoleum aromaticum] | 02881 |
| DUF3096 domain-containing protein [Aromatoleum aromaticum] | 01677 |
| hypothetical protein [Aromatoleum aromaticum] | 09757 |
| transcriptional regulator HexR [Aromatoleum tolulyticum] | 04136 |
| TonB-dependent receptor [Azoarcus indigens] | 07512 |
| nickel-responsive transcriptional regulator NikR [Azoarcus indigens] | 08197 |
| DUF4148 domain-containing protein [Thauera linaloolentis] | 00118 |
| DUF2889 domain-containing protein [Aromatoleum tolulyticum] | 09564 |
| hypothetical protein [Aromatoleum tolulyticum] | 02810 |
| conjugal transfer protein TrbF [Aromatoleum aromaticum] | 03786 |
| conjugal transfer system pilin TrbC [Aromatoleum aromaticum] | 09664 |
| glutamate-cysteine ligase family protein [Aromatoleum aromaticum] | 04120 |
| DUF6351 family protein [Aromatoleum aromaticum] | 05430 |
| TIGR02281 family clan AA aspartic protease [Thauera linaloolentis] | 04011 |
| acyl-CoA dehydrogenase family protein [Azoarcus communis] | 02712 |
| hypothetical protein [Aromatoleum aromaticum] | 06230 |
| RimK family alpha-L-glutamate ligase [Aromatoleum aromaticum] | 03768 |
| EF-hand domain-containing protein [Aromatoleum aromaticum] | 00928 |
| response regulator [Thauera linaloolentis] | 07234 |
| SRPBCC family protein [Aromatoleum aromaticum] | 08410 |
| hemerythrin domain-containing protein [Aromatoleum aromaticum] | 05404 |
| 3'-5' exonuclease [Thauera linaloolentis] | 05561 |
| DEAD/DEAH box helicase [Aromatoleum aromaticum] | 05525 |
| hypothetical protein [Azoarcus indigens] | 03375 |
| sterol desaturase family protein [Thauera linaloolentis] | 04861 |
| hypothetical protein [Aromatoleum aromaticum] | 03481 |
| TrbI/VirB10 family protein [Aromatoleum aromaticum] | 10404 |
| PIN domain nuclease [Thauera chlorobenzoica] | 03012 |
| class I SAM-dependent methyltransferase [Thauera phenylacetica] | 06429 |
| dihydrofolate reductase family protein [Azoarcus indigens] | 02527 |
| S49 family peptidase [Aromatoleum aromaticum] | 07233 |
| hypothetical protein [Azoarcus indigens] | 01605 |
| hypothetical protein [Aromatoleum aromaticum] | 02613 |
| DNA recombination protein RmuC [Aromatoleum tolulyticum] | 04260 |
| HAD-IA family hydrolase [Aromatoleum aromaticum] | 00451 |
| hypothetical protein [Aromatoleum aromaticum] | 09970 |
| PepSY-associated TM helix domain-containing protein [Thauera linaloolentis] | 02181 |
| GntP family permease [Aromatoleum tolulyticum] | 02501 |
| DUF2271 domain-containing protein [Thauera linaloolentis] | 03717 |
| hypothetical protein [Aromatoleum aromaticum] | 09605 |
| FecR domain-containing protein [Aromatoleum aromaticum] | 06738 |
| nucleotidyltransferase domain-containing protein [Aromatoleum aromaticum] | 03361 |
| pilus assembly protein [Aromatoleum tolulyticum] | 06334 |
| PAS domain-containing sensor histidine kinase [Aromatoleum aromaticum] | 04632 |
| nucleotidyltransferase domain-containing protein [Aromatoleum aromaticum] | 07801 |
| glycyl radical protein [Aromatoleum aromaticum] | 03308 |
| citrate transporter [Aromatoleum tolulyticum] | 03390 |
| hypothetical protein [Aromatoleum aromaticum] | 00786 |
| DUF4125 family protein [Aromatoleum aromaticum] | 03996 |
| DUF4037 domain-containing protein [Aromatoleum aromaticum] | 04649 |
| MULTISPECIES: cytochrome-c oxidase [Aromatoleum] | 08166 |
| PHP domain-containing protein [Aromatoleum aromaticum] | 01445 |
| hypothetical protein [Aromatoleum aromaticum] | 00631 |
| hypothetical protein [Aromatoleum tolulyticum] | 00549 |
| CDGSH iron-sulfur domain-containing protein [Aromatoleum tolulyticum] | 06467 |
| hypothetical protein [Aromatoleum aromaticum] | 07934 |
| hypothetical protein [Aromatoleum aromaticum] | 04496 |
| ribbon-helix-helix domain-containing protein [Aromatoleum aromaticum] | 03851 |

| | |
|---|---|
| hypothetical protein [Aromatoleum aromaticum] | 02008 |
| GDYXXLXY domain-containing protein [Thauera linaloolentis] | 04680 |
| nucleotide 5'-monophosphate nucleosidase PpnN [Aromatoleum tolulyticum] | 05639 |
| polysulfide reductase NrfD [Aromatoleum aromaticum] | 07114 |
| hemolysin III family protein [Aromatoleum tolulyticum] | 02815 |
| hypothetical protein [Azoarcus communis] | 04623 |
| DUF4198 domain-containing protein [Thauera linaloolentis] | 07829 |
| nuclear transport factor 2 family protein [Thauera linaloolentis] | 01006 |
| conjugative transfer signal peptidase TraF [Aromatoleum aromaticum] | 09660 |
| integrase, partial gene, partial [Aromatoleum aromaticum] | 10535 |
| MULTISPECIES: hypothetical protein [Rhodocyclales] | 05584 |
| hypothetical protein [Thauera linaloolentis] | 04684 |
| hypothetical protein [Aromatoleum aromaticum] | 06859 |
| formylglycine-generating enzyme family protein [Aromatoleum aromaticum] | 01508 |
| hypothetical protein [Thauera linaloolentis] | 02605 |
| HD domain-containing protein [Aromatoleum tolulyticum] | 02756 |
| ATP-binding protein [Azoarcus indigens] | 02099 |
| BREX system P-loop protein BrxC [Aromatoleum aromaticum] | 00611 |
| hypothetical protein [Aromatoleum tolulyticum] | 01820 |
| hypothetical protein [Aromatoleum tolulyticum] | 04870 |
| DUF4157 domain-containing protein [Azoarcus communis] | 03770 |
| hypothetical protein [Aromatoleum toluclasticum] | 03918 |
| hypothetical protein [Aromatoleum aromaticum] | 07553 |
| protease Lon-related BREX system protein BrxL [Aromatoleum aromaticum] | 00519 |
| 3-hydroxyacyl-CoA dehydrogenase/enoyl-CoA hydratase family protein [Aromatoleum aromaticum] | 04167 |
| cobalamin B12-binding domain-containing protein [Aromatoleum aromaticum] | 03974 |
| 5-amino-6-(D-ribitylamino)uracil--L-tyrosine 4-hydroxyphenyl transferase CofH [Aromatoleum aromaticum] | 04751 |
| molybdopterin-binding/glycosyltransferase family 2 protein [Aromatoleum aromaticum] | 01570 |
| CopG family transcriptional regulator [Aromatoleum aromaticum] | 09756 |
| LLM class flavin-dependent oxidoreductase [Aromatoleum tolulyticum] | 07035 |
| metal/formaldehyde-sensitive transcriptional repressor [Thauera linaloolentis] | 10315 |
| response regulator [Aromatoleum tolulyticum] | 04321 |
| archease [Aromatoleum tolulyticum] | 05902 |
| type II toxin-antitoxin system Phd/YefM family antitoxin [Aromatoleum aromaticum] | 09005 |
| fumarate reductase subunit FrdD [Aromatoleum tolulyticum] | 07290 |
| TolC family protein [Thauera linaloolentis] | 07129 |
| fumarate reductase subunit C [Aromatoleum tolulyticum] | 07025 |
| GTPase domain-containing protein [Aromatoleum tolulyticum] | 01134 |
| hypothetical protein [Thauera linaloolentis] | 06873 |
| EAL domain-containing protein [Aromatoleum aromaticum] | 04000 |
| Crp/Fnr family transcriptional regulator [Aromatoleum aromaticum] | 10082 |
| TonB-dependent receptor [Thauera linaloolentis] | 02140 |
| 2-phospho-L-lactate transferase [Aromatoleum aromaticum] | 00237 |
| flavin reductase family protein [Aromatoleum toluclasticum] | 04169 |
| 2-phospho-L-lactate guanylyltransferase [Aromatoleum aromaticum] | 06264 |
| hypothetical protein [Aromatoleum aromaticum] | 01593 |
| coenzyme F420-0:L-glutamate ligase [Aromatoleum aromaticum] | 08450 |
| serine acetyltransferase [Azoarcus indigens] | 00408 |
| SRPBCC family protein [Azoarcus indigens] | 10612 |
| cell envelope integrity protein CreD [Azoarcus indigens] | 04243 |
| MULTISPECIES: IS5 family transposase [Aromatoleum] | 05236 |
| two-component system response regulator [Azoarcus indigens] | 07236 |
| peptidoglycan-binding protein [Azoarcus indigens] | 05826 |
| AbrB/MazE/SpoVT family DNA-binding domain-containing protein [Thauera chlorobenzoica] | 10468 |
| MULTISPECIES: type II toxin-antitoxin system Phd/YefM family antitoxin [Aromatoleum] | 06845 |
| BrnA antitoxin family protein [Thauera chlorobenzoica] | 10336 |
| PIN domain-containing protein [Thauera linaloolentis] | 03379 |
| DNA-binding protein [Aromatoleum aromaticum] | 03981 |
| carbohydrate porin [Aromatoleum tolulyticum] | 06168 |

| | |
|---|---|
| hypothetical protein [Aromatoleum aromaticum] | 06282 |
| MFS transporter [Aromatoleum tolulyticum] | 00484 |
| FapA family protein [Propionivibrio dicarboxylicus] | 01573 |
| hypothetical protein [Aromatoleum tolulyticum] | 09161 |
| hypothetical protein [Aromatoleum aromaticum] | 04463 |
| modified peptide precursor CbpA [Thauera chlorobenzoica] | 07753 |
| hypothetical protein [Aromatoleum tolulyticum] | 02327 |
| aldose 1-epimerase [Aromatoleum tolulyticum] | 00203 |
| nitrogen fixation protein NifZ [Aromatoleum tolulyticum] | 07957 |
| GDP-mannose--glycolipid 4-beta-D-mannosyltransferase [Azoarcus indigens] | 05912 |
| type II toxin-antitoxin system ParD family antitoxin [Thauera linaloolentis] | 03789 |
| zinc ribbon domain-containing protein [Aromatoleum tolulyticum] | 05336 |
| ketoacid CoA transferase [Aromatoleum toluclasticum] | 08275 |
| hypothetical protein [Aromatoleum tolulyticum] | 10273 |
| energy-coupling factor transporter transmembrane protein EcfT [Thauera linaloolentis] | 07731 |
| hypothetical protein [Aromatoleum tolulyticum] | 04094 |
| nuclear transport factor 2 family protein [Aromatoleum toluclasticum] | 08116 |
| hypothetical protein [Aromatoleum tolulyticum] | 10584 |
| hypothetical protein [Aromatoleum tolulyticum] | 01501 |
| hypothetical protein [Aromatoleum toluclasticum] | 06630 |
| OB-fold domain-containing protein [Aromatoleum toluclasticum] | 01630 |
| 5-methyltetrahydropteroyltriglutamate--homocysteine S-methyltransferase [Thauera linaloolentis] | 05526 |
| diguanylate cyclase [Aromatoleum tolulyticum] | 05674 |
| type I 3-dehydroquinate dehydratase [Aromatoleum aromaticum] | 03275 |
| type I-E CRISPR-associated endonuclease Cas1e [Thauera chlorobenzoica] | 09439 |
| type I-E CRISPR-associated protein Cas5/CasD [Thauera chlorobenzoica] | 06365 |
| glycoside hydrolase family 19 protein [Aromatoleum aromaticum] | 00731 |
| type I-E CRISPR-associated protein Cas7/Cse4/CasC [Thauera chlorobenzoica] | 07064 |
| methyltransferase domain-containing protein [Propionivibrio dicarboxylicus] | 01847 |
| hypothetical protein [Thauera chlorobenzoica] | 02243 |
| zinc ribbon domain-containing protein [Aromatoleum aromaticum] | 10075 |
| hypothetical protein [Aromatoleum tolulyticum] | 01193 |
| hypothetical protein [Aromatoleum aromaticum] | 06268 |
| hemagglutinin repeat-containing protein [Thauera linaloolentis] | 04027 |
| response regulator [Thauera linaloolentis] | 00999 |
| DUF3016 domain-containing protein [Thauera linaloolentis] | 02952 |
| pyridoxamine 5'-phosphate oxidase family protein [Aromatoleum aromaticum] | 03785 |
| glycerate kinase [Thauera linaloolentis] | 10108 |
| type II toxin-antitoxin system RelE/ParE family toxin [Aromatoleum aromaticum] | 05067 |
| putative hydro-lyase [Azoarcus indigens] | 07225 |
| iron-containing redox enzyme family protein [Azoarcus indigens] | 03903 |
| hemerythrin domain-containing protein [Aromatoleum aromaticum] | 02400 |
| CRISPR-associated helicase/endonuclease Cas3 [Thauera chlorobenzoica] | 02915 |
| ABC transporter substrate-binding protein [Aromatoleum tolulyticum] | 05652 |
| M67 family metallopeptidase [Aromatoleum tolulyticum] | 04090 |
| L-seryl-tRNA(Sec) selenium transferase [Thauera linaloolentis] | 07622 |
| hypothetical protein [Aromatoleum tolulyticum] | 04009 |
| PepSY domain-containing protein [Azoarcus indigens] | 00146 |
| hypothetical protein [Thauera linaloolentis] | 03810 |
| transcriptional regulator [Aromatoleum aromaticum] | 03902 |
| hypothetical protein [Aromatoleum aromaticum] | 03678 |
| DUF72 domain-containing protein [Thauera linaloolentis] | 00978 |
| DUF927 domain-containing protein [Aromatoleum aromaticum] | 09521 |
| hypothetical protein [Azoarcus indigens] | 04014 |
| MCP four helix bundle domain-containing protein [Aromatoleum tolulyticum] | 09023 |
| MULTISPECIES: IS30-like element ISPsp7 family transposase [Proteobacteria] | 08278 |
| OB-fold domain-containing protein [Aromatoleum toluclasticum] | 05702 |
| hypothetical protein [Aromatoleum tolulyticum] | 02197 |
| mechanosensitive ion channel [Aromatoleum aromaticum] | 00302 |

| | |
|---|---|
| GDSL-type esterase/lipase family protein [Azoarcus indigens] | 08276 |
| tyrosine-type recombinase/integrase [Aromatoleum aromaticum] | 01996 |
| alpha/beta hydrolase-fold protein [Thauera linaloolentis] | 00154 |
| H-NS histone family protein [Aromatoleum aromaticum] | 01935 |
| MULTISPECIES: periplasmic nitrate reductase, NapE protein [Azoarcus] | 08079 |
| hypothetical protein [Aromatoleum tolulyticum] | 06921 |
| Zn-dependent hydrolase [Propionivibrio dicarboxylicus] | 03380 |
| helix-turn-helix transcriptional regulator [Thauera linaloolentis] | 03236 |
| polysaccharide biosynthesis/export family protein [Thauera linaloolentis] | 04633 |
| hypothetical protein [Thauera linaloolentis] | 05470 |
| DUF2274 domain-containing protein [Thauera linaloolentis] | 01355 |
| O-antigen ligase family protein [Zoogloea dura] | 10405 |
| antibiotic biosynthesis monooxygenase [Thauera linaloolentis] | 06400 |
| exopolysaccharide biosynthesis protein [Thauera chlorobenzoica] | 01050 |
| type II toxin-antitoxin system RelE/ParE family toxin [Thauera phenylacetica] | 06350 |
| type IV pilin protein [Aromatoleum tolulyticum] | 08690 |
| 2-phosphosulfolactate phosphatase [Aromatoleum tolulyticum] | 06133 |
| high-potential iron-sulfur protein [Aromatoleum tolulyticum] | 07354 |
| DUF5938 domain-containing protein [Aromatoleum toluclasticum] | 10013 |
| alanine--glyoxylate aminotransferase family protein [Azoarcus indigens] | 09933 |
| type II restriction endonuclease [Aromatoleum tolulyticum] | 04653 |
| SemiSWEET transporter [Aromatoleum tolulyticum] | 03073 |
| hypothetical protein [Thauera linaloolentis] | 05529 |
| nuclear transport factor 2 family protein [Azoarcus indigens] | 10585 |
| type II toxin-antitoxin system RelE/ParE family toxin [Thauera chlorobenzoica] | 03534 |
| TetR family transcriptional regulator [Azoarcus indigens] | 05007 |
| MFS transporter [Azoarcus indigens] | 09929 |
| Hpt domain-containing protein [Thauera linaloolentis] | 02479 |
| hypothetical protein [Aromatoleum aromaticum] | 01854 |
| hypothetical protein [Thauera linaloolentis] | 00065 |
| ribbon-helix-helix protein, CopG family [Thauera linaloolentis] | 09912 |
| hypothetical protein [Thauera linaloolentis] | 10153 |
| energy transducer TonB [Thauera linaloolentis] | 05895 |
| hypothetical protein [Aromatoleum aromaticum] | 06272 |
| type II toxin-antitoxin system RelE/ParE family toxin [Aromatoleum aromaticum] | 00504 |
| hypothetical protein [Aromatoleum aromaticum] | 10080 |
| DUF6285 domain-containing protein [Thauera linaloolentis] | 08347 |
| hypothetical protein [Thauera linaloolentis] | 06390 |
| hypothetical protein [Thauera linaloolentis] | 04201 |
| hypothetical protein [Aromatoleum aromaticum] | 06863 |
| hypothetical protein [Thauera linaloolentis] | 03477 |
| hypothetical protein [Thauera linaloolentis] | 04146 |
| mechanosensitive channel MscK [Thauera linaloolentis] | 08431 |
| hypothetical protein [Aromatoleum aromaticum] | 05538 |
| VirB3 family type IV secretion system protein [Thauera linaloolentis] | 01696 |
| lipocalin-like domain-containing protein [Azoarcus sp. Aa7] | 00979 |
| response regulator [Thauera chlorobenzoica] | 08883 |
| MULTISPECIES: TolC family protein [Rhodocyclales] | 07922 |
| endonuclease/exonuclease/phosphatase family protein [Aromatoleum tolulyticum] | 02369 |
| MliC family protein [Aromatoleum tolulyticum] | 06134 |
| NUDIX domain-containing protein [Aromatoleum tolulyticum] | 04528 |
| DnaJ domain-containing protein [Aromatoleum aromaticum] | 02729 |
| adenosine-specific kinase [Aromatoleum tolulyticum] | 06988 |
| CYTH domain-containing protein [Thauera phenylacetica] | 04581 |
| cytochrome c biogenesis protein ResB [Thauera phenylacetica] | 03545 |
| autotransporter domain-containing protein [Aromatoleum tolulyticum] | 08897 |
| DNA ligase D [Aromatoleum aromaticum] | 03815 |
| glycine zipper 2TM domain-containing protein [Aromatoleum tolulyticum] | 09826 |
| carbonic anhydrase family protein [Aromatoleum tolulyticum] | 05361 |

| | |
|---|---|
| hypothetical protein [Aromatoleum tolulyticum] | 07241 |
| plasmid pRiA4b ORF-3 family protein [Aromatoleum tolulyticum] | 01192 |
| 2-oxoacid:acceptor oxidoreductase subunit alpha [Aromatoleum aromaticum] | 07174 |
| CbtA family protein [Aromatoleum tolulyticum] | 06261 |
| CbtB-domain containing protein [Aromatoleum tolulyticum] | 09760 |
| hypothetical protein [Thauera chlorobenzoica] | 04174 |
| ChaB family protein [Aromatoleum aromaticum] | 09387 |
| hypothetical protein [Zoogloea dura] | 02145 |
| four-helix bundle copper-binding protein [Aromatoleum aromaticum] | 08415 |
| AlpA family phage regulatory protein [Zoogloea dura] | 00310 |
| hypothetical protein [Aromatoleum tolulyticum] | 04494 |
| type II toxin-antitoxin system Phd/YefM family antitoxin [Aromatoleum aromaticum] | 00254 |
| FAD-dependent oxidoreductase [Thauera phenylacetica] | 07459 |
| DUF2818 family protein [Thauera phenylacetica] | 04529 |
| arginine N-succinyltransferase [Aromatoleum tolulyticum] | 01535 |
| DUF433 domain-containing protein [Aromatoleum aromaticum] | 02001 |
| pilus assembly protein TadG-related protein [Aromatoleum aromaticum] | 05334 |
| hypothetical protein [Aromatoleum tolulyticum] | 04047 |
| pilus assembly protein [Aromatoleum aromaticum] | 05877 |
| hypothetical protein [Aromatoleum tolulyticum] | 02723 |
| hypothetical protein [Aromatoleum tolulyticum] | 06659 |
| DNA-binding protein [Aromatoleum tolulyticum] | 03385 |
| hypothetical protein [Aromatoleum tolulyticum] | 00130 |
| phasin family protein [Aromatoleum aromaticum] | 08666 |
| DcaP family trimeric outer membrane transporter [Propionivibrio dicarboxylicus] | 08338 |
| hypothetical protein [Aromatoleum tolulyticum] | 08890 |
| hypothetical protein [Aromatoleum tolulyticum] | 05963 |
| hypothetical protein [Aromatoleum tolulyticum] | 06992 |
| benzylsuccinate synthase gamma subunit family protein [Aromatoleum aromaticum] | 08150 |
| sugar phosphate isomerase/epimerase [Aromatoleum tolulyticum] | 09011 |
| hypothetical protein [Aromatoleum tolulyticum] | 00289 |
| hypothetical protein [Aromatoleum tolulyticum] | 07071 |
| hypothetical protein [Aromatoleum tolulyticum] | 07319 |
| benzylsuccinate synthase subunit beta [Aromatoleum aromaticum] | 01780 |
| hypothetical protein [Aromatoleum aromaticum] | 06624 |
| ribbon-helix-helix protein, CopG family [Thauera chlorobenzoica] | 03625 |
| lmo0937 family membrane protein [Aromatoleum tolulyticum] | 01734 |
| DUF262 domain-containing HNH endonuclease family protein [Aromatoleum aromaticum] | 07481 |
| hypothetical protein [Azoarcus indigens] | 09568 |
| type II toxin-antitoxin system Phd/YefM family antitoxin [Azoarcus indigens] | 01420 |
| hypothetical protein [Azoarcus indigens] | 01652 |
| Ku protein [Azoarcus indigens] | 10300 |
| hsp70 family protein [Aromatoleum aromaticum] | 01681 |
| DUF3175 domain-containing protein [Azoarcus indigens] | 06142 |
| DUF3325 domain-containing protein [Azoarcus indigens] | 09919 |
| DUF2760 domain-containing protein [Aromatoleum aromaticum] | 04197 |
| hypothetical protein [Azoarcus indigens] | 00680 |
| ATP-binding protein [Aromatoleum aromaticum] | 05031 |
| type VI secretion system-associated FHA domain protein TagH [Azoarcus communis] | 06552 |
| hypothetical protein [Aromatoleum aromaticum] | 05300 |
| hypothetical protein [Azoarcus communis] | 09639 |
| hypothetical protein [Azoarcus communis] | 10021 |
| hypothetical protein [Aromatoleum aromaticum] | 07535 |
| hypothetical protein [Thauera linaloolentis] | 02244 |
| hypothetical protein [Aromatoleum aromaticum] | 10436 |
| P-type conjugative transfer protein TrbL [Thauera linaloolentis] | 06397 |
| M20/M25/M40 family metallo-hydrolase [Zoogloea dura] | 02214 |
| hypothetical protein [Azoarcus indigens] | 06501 |
| DUF3427 domain-containing protein [Azoarcus communis] | 04758 |

| | |
|---|---|
| PIN domain-containing protein [Thauera chlorobenzoica] | 02914 |
| amidohydrolase family protein [Thauera chlorobenzoica] | 00661 |
| hypothetical protein [Aromatoleum tolulyticum] | 06978 |
| GrpB family protein [Thauera chlorobenzoica] | 01235 |
| hypothetical protein [Aromatoleum aromaticum] | 06463 |
| Hpt domain-containing protein [Uliginosibacterium aquaticum] | 03639 |
| AI-2E family transporter [Aromatoleum tolulyticum] | 07042 |
| quinol oxidase [Thauera chlorobenzoica] | 07681 |
| site-specific DNA-methyltransferase [Aromatoleum aromaticum] | 10167 |
| hypothetical protein [Aromatoleum tolulyticum] | 02545 |
| Atrophin-1 multi-domain protein [Azoarcus indigens] | 05375 |
| YgiT-type zinc finger protein [Aromatoleum aromaticum] | 00416 |
| hypothetical protein [Azoarcus indigens] | 06815 |
| glycosyltransferase family 2 protein [Azoarcus indigens] | 05457 |
| hypothetical protein [Aromatoleum tolulyticum] | 06288 |
| protease HtpX [Azoarcus indigens] | 09663 |
| zonular occludens toxin [Azoarcus indigens] | 08912 |
| class I SAM-dependent DNA methyltransferase [Aromatoleum aromaticum] | 09759 |
| hypothetical protein [Aromatoleum tolulyticum] | 01761 |
| cysteine dioxygenase type I [Thauera linaloolentis] | 02338 |
| hypothetical protein [Aromatoleum tolulyticum] | 07126 |
| NADPH-dependent FMN reductase [Thauera linaloolentis] | 04286 |
| relaxase/mobilization nuclease and DUF3363 domain-containing protein [Thauera linaloolentis] | 00843 |
| hypothetical protein [Thauera linaloolentis] | 07343 |
| 3-aminobutyryl-CoA ammonia lyase [Aromatoleum tolulyticum] | 08721 |
| PilW family protein [Aromatoleum tolulyticum] | 07504 |
| WbqC family protein [Azoarcus communis] | 07379 |
| nucleotidyltransferase [Thauera linaloolentis] | 10314 |
| flippase-like domain-containing protein, partial [Azoarcus communis] | 06443 |
| hypothetical protein [Aromatoleum toluclasticum] | 01655 |
| azurin [Thauera linaloolentis] | 07483 |
| ATP-binding cassette domain-containing protein, partial [Aromatoleum toluclasticum] | 05809 |
| PEP-CTERM sorting domain-containing protein [Azoarcus communis] | 02707 |
| alpha/beta hydrolase [Azoarcus communis] | 01886 |
| DUF475 domain-containing protein [Thauera linaloolentis] | 06687 |
| MULTISPECIES: D-xylulose 5-phosphate/D-fructose 6-phosphate phosphoketolase [Proteobacteria] | 05670 |
| MULTISPECIES: helix-turn-helix domain-containing protein [Rhodocyclales] | 01423 |
| 6-carboxytetrahydropterin synthase [Azoarcus communis] | 00474 |
| glyoxalase [Azoarcus communis] | 00291 |
| Flp pilus assembly protein CpaB [Thauera phenylacetica] | 06274 |
| helix-turn-helix transcriptional regulator [Thauera phenylacetica] | 00967 |
| acetyltransferase [Thauera phenylacetica] | 03468 |
| SDR family oxidoreductase [Thauera phenylacetica] | 00122 |
| 2-dehydro-3-deoxygalactonokinase [Propionivibrio dicarboxylicus] | 08776 |
| MFS transporter [Propionivibrio dicarboxylicus] | 04361 |
| acetyltransferase [Zoogloea dura] | 10347 |
| SRPBCC family protein [Aromatoleum tolulyticum] | 02634 |
| hypothetical protein [Aromatoleum tolulyticum] | 04536 |
| FapA family protein [Aromatoleum tolulyticum] | 09642 |
| hypothetical protein [Aromatoleum tolulyticum] | 07560 |
| ABC transporter substrate-binding protein [Thauera linaloolentis] | 05070 |
| gamma-glutamylcyclotransferase [Aromatoleum tolulyticum] | 03567 |
| hypothetical protein [Thauera linaloolentis] | 06871 |
| hypothetical protein [Aromatoleum tolulyticum] | 00418 |
| class I SAM-dependent methyltransferase [Aromatoleum tolulyticum] | 05934 |
| ATP-dependent helicase [Thauera linaloolentis] | 04553 |
| TonB-dependent receptor [Thauera linaloolentis] | 00849 |
| type II toxin-antitoxin system VapC family toxin [Thauera linaloolentis] | 02646 |
| glutaredoxin family protein [Thauera chlorobenzoica] | 04631 |

| | |
|---|---|
| type II toxin-antitoxin system PemK/MazF family toxin [Thauera chlorobenzoica] | 04306 |
| hypothetical protein [Thauera chlorobenzoica] | 03140 |
| phosphoethanolamine--lipid A transferase [Thauera chlorobenzoica] | 08205 |
| AIPR family protein [Thauera chlorobenzoica] | 05713 |
| chemotaxis protein CheW [Azoarcus indigens] | 08339 |
| gluconate 2-dehydrogenase subunit 3 family protein [Azoarcus indigens] | 03381 |
| phospholipase D family protein [Aromatoleum toluclasticum] | 04148 |
| dialkylrecorsinol condensing enzyme [Azoarcus communis] | 01227 |
| DUF4198 domain-containing protein [Azoarcus indigens] | 03535 |
| 3,4-dihydroxyphenylacetate 2,3-dioxygenase [Azoarcus indigens] | 07744 |
| MBL fold metallo-hydrolase [Azoarcus indigens] | 01164 |
| putative zinc-binding metallopeptidase [Azoarcus indigens] | 00055 |
| NAD(P)/FAD-dependent oxidoreductase [Azoarcus indigens] | 08630 |
| tetratricopeptide repeat protein [Aromatoleum diolicum] | 05741 |
| hypothetical protein [Thauera linaloolentis] | 09057 |
| hypothetical protein [Thauera linaloolentis] | 00053 |
| sulfatase-like hydrolase/transferase [Thauera linaloolentis] | 05459 |
| histidine phosphatase family protein [Aromatoleum tolulyticum] | 05938 |
| MULTISPECIES: hypothetical protein [Aromatoleum] | 00982 |
| MULTISPECIES: DUF2840 domain-containing protein [Proteobacteria] | 01603 |
| sulfatase-like hydrolase/transferase [Aromatoleum tolulyticum] | 07402 |
| MULTISPECIES: hypothetical protein [Proteobacteria] | 06692 |
| DUF2285 domain-containing protein [Thauera linaloolentis] | 00117 |
| DUF736 domain-containing protein [Thauera linaloolentis] | 01419 |
| hypothetical protein [Aromatoleum tolulyticum] | 06790 |
| monooxygenase [Thauera linaloolentis] | 07506 |
| hypothetical protein [Aromatoleum tolulyticum] | 02733 |
| cupredoxin domain-containing protein [Aromatoleum tolulyticum] | 04091 |
| ethylbenzene dehydrogenase-related protein [Aromatoleum tolulyticum] | 00641 |
| c-type cytochrome [Thauera linaloolentis] | 02939 |
| twin-arginine translocation signal domain-containing protein [Aromatoleum tolulyticum] | 09688 |
| hypothetical protein [Aromatoleum tolulyticum] | 04641 |
| hypothetical protein [Aromatoleum tolulyticum] | 06669 |
| hypothetical protein [Thauera linaloolentis] | 05957 |
| KamA family radical SAM protein [Aromatoleum tolulyticum] | 07148 |
| zinc-binding dehydrogenase [Aromatoleum tolulyticum] | 07583 |
| helix-turn-helix domain-containing protein [Thauera linaloolentis] | 08500 |
| hypothetical protein [Aromatoleum tolulyticum] | 09306 |
| MULTISPECIES: DUF2958 domain-containing protein [Proteobacteria] | 03628 |
| lysine 5,6-aminomutase subunit alpha [Aromatoleum tolulyticum] | 03169 |
| cobalamin-dependent protein [Aromatoleum tolulyticum] | 06586 |
| thioesterase family protein [Aromatoleum tolulyticum] | 01416 |
| hypothetical protein [Aromatoleum tolulyticum] | 10311 |
| DUF3427 domain-containing protein [Aromatoleum tolulyticum] | 06301 |
| hypothetical protein [Aromatoleum toluclasticum] | 02727 |
| ABC transporter permease [Azoarcus communis] | 01740 |
| DUF1640 domain-containing protein [Aromatoleum aromaticum] | 04873 |
| hypothetical protein [Aromatoleum aromaticum] | 03742 |
| DUF421 domain-containing protein [Aromatoleum aromaticum] | 05544 |
| type II toxin-antitoxin system RelE/ParE family toxin [Aromatoleum aromaticum] | 03152 |
| TIGR02677 family protein [Aromatoleum aromaticum] | 05399 |
| hypothetical protein [Aromatoleum aromaticum] | 10484 |
| TIGR02678 family protein [Aromatoleum aromaticum] | 07837 |
| TIGR02679 family protein [Aromatoleum aromaticum] | 03102 |
| hypothetical protein [Aromatoleum aromaticum] | 02134 |
| hypothetical protein [Aromatoleum aromaticum] | 03384 |
| hypothetical protein [Aromatoleum aromaticum] | 00948 |
| hypothetical protein [Aromatoleum aromaticum] | 07399 |
| hypothetical protein [Aromatoleum aromaticum] | 04199 |

| | |
|---|---|
| phage holin family protein [Aromatoleum aromaticum] | 09211 |
| PEP-CTERM sorting domain-containing protein [Aromatoleum aromaticum] | 05213 |
| apolipoprotein A/E/C3 [Aromatoleum aromaticum] | 08687 |
| acyltransferase family protein [Aromatoleum aromaticum] | 07287 |
| hypothetical protein [Aromatoleum aromaticum] | 08712 |
| AbrB/MazE/SpoVT family DNA-binding domain-containing protein [Aromatoleum aromaticum] | 06848 |
| hypothetical protein [Aromatoleum aromaticum] | 08602 |
| hypothetical protein [Aromatoleum aromaticum] | 03280 |
| toprim domain-containing protein [Aromatoleum aromaticum] | 01787 |
| zeta toxin family protein [Aromatoleum aromaticum] | 03978 |
| IS110-like element ISAzo22 family transposase [Aromatoleum aromaticum] | 05051 |
| TIGR02680 family protein [Aromatoleum aromaticum] | 02519 |
| iron-sulfur cluster assembly scaffold protein [Aromatoleum aromaticum] | 01757 |
| type II toxin-antitoxin system VapB family antitoxin [Aromatoleum aromaticum] | 10181 |
| serine hydrolase [Thauera linaloolentis] | 06385 |
| EthD domain-containing protein [Aromatoleum toluclasticum] | 03787 |
| ribonucleoside triphosphate reductase [Thauera linaloolentis] | 00444 |
| cytochrome D1 [Aromatoleum tolulyticum] | 10276 |
| hypothetical protein [Aromatoleum aromaticum] | 09475 |
| hypothetical protein [Aromatoleum tolulyticum] | 08898 |
| potassium-transporting ATPase subunit KdpC [Azoarcus indigens] | 01438 |
| hypothetical protein [Azoarcus communis] | 02101 |
| FHA domain-containing protein [Aromatoleum aromaticum] | 02954 |
| nitroreductase family deazaflavin-dependent oxidoreductase [Aromatoleum toluclasticum] | 03323 |
| DEAD/DEAH box helicase [Aromatoleum toluclasticum] | 02132 |
| tripartite tricarboxylate transporter TctB family protein [Aromatoleum tolulyticum] | 02217 |
| hypothetical protein [Aromatoleum aromaticum] | 07730 |
| AAA family ATPase [Azoarcus indigens] | 02616 |
| hypothetical protein [Thauera linaloolentis] | 09424 |
| M10 family metallopeptidase C-terminal domain-containing protein [Aromatoleum tolulyticum] | 06285 |
| phosphoesterase [Aromatoleum tolulyticum] | 01447 |
| hypothetical protein [Thauera linaloolentis] | 05093 |
| molecular chaperone TorD family protein [Aromatoleum aromaticum] | 02514 |
| hypothetical protein [Thauera phenylacetica] | 08028 |
| IS4-like element ISAzo5 family transposase [Aromatoleum aromaticum] | 01043 |
| type II toxin-antitoxin system Phd/YefM family antitoxin [Azoarcus communis] | 01928 |
| hypothetical protein [Azoarcus indigens] | 07486 |
| glycosyltransferase [Aromatoleum aromaticum] | 03745 |
| hypothetical protein [Azoarcus indigens] | 07263 |
| MaoC family dehydratase N-terminal domain-containing protein [Aromatoleum toluclasticum] | 03269 |
| hypothetical protein [Aromatoleum aromaticum] | 08147 |
| DUF2029 domain-containing protein [Aromatoleum tolulyticum] | 08433 |
| maltose alpha-D-glucosyltransferase [Aromatoleum aromaticum] | 10581 |
| SCO family protein [Aromatoleum tolulyticum] | 09255 |
| DUF11 domain-containing protein [Azoarcus indigens] | 06996 |
| hypothetical protein [Azoarcus indigens] | 07996 |
| hypothetical protein [Aromatoleum tolulyticum] | 05264 |
| SurA N-terminal domain-containing protein [Aromatoleum tolulyticum] | 04545 |
| IS66 family transposase zinc-finger binding domain-containing protein, partial [Thauera terpenica] | 04202 |
| CopD family protein [Aromatoleum aromaticum] | 10281 |
| phosphatase PAP2 family protein [Aromatoleum aromaticum] | 05907 |
| hypothetical protein [Aromatoleum toluclasticum] | 02045 |
| hypothetical protein [Aromatoleum toluclasticum] | 10185 |
| molecular chaperone TorD family protein [Aromatoleum tolulyticum] | 01432 |
| hypothetical protein [Aromatoleum tolulyticum] | 06763 |
| hypothetical protein [Aromatoleum tolulyticum] | 06986 |
| hypothetical protein [Aromatoleum aromaticum] | 04130 |
| isopenicillin N synthase family oxygenase [Azoarcus indigens] | 02641 |
| uracil-DNA glycosylase [Azoarcus rhizosphaerae] | 08791 |

| | |
|---|---|
| DUF2231 domain-containing protein [Aromatoleum aromaticum] | 08066 |
| hypothetical protein [Aromatoleum toluclasticum] | 01789 |
| glycosyltransferase family 2 protein [Aromatoleum aromaticum] | 03132 |
| chaperone NapD [Aromatoleum aromaticum] | 05516 |
| Mov34/MPN/PAD-1 family protein [Aromatoleum aromaticum] | 07764 |
| MULTISPECIES: sugar phosphate isomerase/epimerase [Azoarcus] | 02565 |
| DUF350 domain-containing protein [Azoarcus indigens] | 08823 |
| ABC-type transport auxiliary lipoprotein family protein [Aromatoleum tolulyticum] | 04166 |
| hypothetical protein [Aromatoleum aromaticum] | 06402 |
| hypothetical protein [Aromatoleum aromaticum] | 03706 |
| FAD-binding oxidoreductase [Azoarcus indigens] | 07413 |
| acyltransferase [Aromatoleum aromaticum] | 01589 |
| NAD(P)-dependent oxidoreductase [Azoarcus indigens] | 00635 |
| hypothetical protein [Aromatoleum tolulyticum] | 07462 |
| type II toxin-antitoxin system prevent-host-death family antitoxin [Aromatoleum aromaticum] | 04540 |
| MULTISPECIES: ester cyclase [Aromatoleum] | 08294 |
| type I-F CRISPR-associated protein Csy3 [Zoogloea dura] | 05760 |
| TetR/AcrR family transcriptional regulator [Thauera linaloolentis] | 01005 |
| SRPBCC family protein [Thauera linaloolentis] | 08348 |
| type I-F CRISPR-associated endoribonuclease Cas6/Csy4 [Zoogloea dura] | 07854 |
| radical SAM protein [Aromatoleum toluclasticum] | 09013 |
| hypothetical protein [Aromatoleum tolulyticum] | 01200 |
| hypothetical protein [Aromatoleum aromaticum] | 06626 |
| CoA transferase [Aromatoleum toluclasticum] | 05059 |
| hypothetical protein [Azoarcus indigens] | 04909 |
| DNA-binding transcriptional regulator [Thauera chlorobenzoica] | 04332 |
| glycoside hydrolase family 19 protein [Thauera linaloolentis] | 09419 |
| CopG family transcriptional regulator [Aromatoleum aromaticum] | 03967 |
| CDP-alcohol phosphatidyltransferase family protein [Azoarcus communis] | 07128 |
| FecR family protein [Azoarcus indigens] | 00566 |
| IS1380-like element ISAzo40 family transposase [Aromatoleum aromaticum] | 02575 |
| DEAD/DEAH box helicase [Thauera chlorobenzoica] | 08521 |
| type I-E CRISPR-associated protein Cas6/Cse3/CasE [Thauera chlorobenzoica] | 06411 |
| BREX-1 system adenine-specific DNA-methyltransferase PglX [Aromatoleum aromaticum] | 03462 |
| MULTISPECIES: hypothetical protein [Thauera] | 05546 |
| hypothetical protein [Thauera chlorobenzoica] | 00468 |
| hypothetical protein [Thauera phenylacetica] | 01178 |
| hypothetical protein [Thauera linaloolentis] | 00964 |
| HDIG domain-containing protein [Thauera chlorobenzoica] | 05504 |
| BtrH N-terminal domain-containing protein [Azoarcus communis] | 06296 |
| C-GCAxxG-C-C family protein [Aromatoleum tolulyticum] | 07286 |
| DUF3616 domain-containing protein [Aromatoleum aromaticum] | 06791 |
| hypothetical protein [Thauera chlorobenzoica] | 08700 |
| helix-turn-helix domain-containing protein [Thauera chlorobenzoica] | 09917 |
| DUF4142 domain-containing protein [Aromatoleum aromaticum] | 08139 |
| DUF5602 domain-containing protein [Aromatoleum tolulyticum] | 08966 |
| general secretion pathway protein GspB [Aromatoleum tolulyticum] | 09589 |
| hypothetical protein [Azoarcus communis] | 03212 |
| hypothetical protein [Aromatoleum aromaticum] | 05022 |
| helix-turn-helix domain-containing protein [Aromatoleum aromaticum] | 09988 |
| BON domain-containing protein [Aromatoleum aromaticum] | 03008 |
| hypothetical protein [Thauera phenylacetica] | 02056 |
| hypothetical protein [Thauera phenylacetica] | 03672 |
| hypothetical protein [Thauera phenylacetica] | 01012 |
| hypothetical protein [Thauera linaloolentis] | 01905 |
| lipid A deacylase LpxR family protein [Thauera chlorobenzoica] | 02058 |
| organoarsenical effux MFS transporter ArsJ [Azoarcus communis] | 03631 |
| Arc family DNA-binding protein [Aromatoleum aromaticum] | 04739 |
| hypothetical protein [Aromatoleum aromaticum] | 03741 |

| | |
|---|---|
| DUF2442 domain-containing protein [Thauera chlorobenzoica] | 10076 |
| P-type conjugative transfer protein TrbL [Aromatoleum aromaticum] | 08767 |
| cupin domain-containing protein [Azoarcus rhizosphaerae] | 05730 |
| DUF5615 family PIN-like protein [Aromatoleum aromaticum] | 04692 |
| hypothetical protein [Aromatoleum aromaticum] | 06799 |
| alkaline phosphatase family protein [Thauera chlorobenzoica] | 03680 |
| MULTISPECIES: type II toxin-antitoxin system HicB family antitoxin [Aromatoleum] | 10539 |
| acyl-protein synthetase [Aromatoleum diolicum] | 09616 |
| hypothetical protein [Thauera linaloolentis] | 05394 |
| hypothetical protein [Thauera phenylacetica] | 10331 |
| MULTISPECIES: type II toxin-antitoxin system Phd/YefM family antitoxin [Aromatoleum] | 01162 |
| ferritin-like domain-containing protein [Aromatoleum aromaticum] | 02740 |
| integrase core domain-containing protein, partial [Thauera linaloolentis] | 02814 |
| HutD family protein [Aromatoleum toluclasticum] | 06981 |
| hypothetical protein [Thauera linaloolentis] | 09717 |
| hypothetical protein [Aromatoleum tolulyticum] | 04498 |
| HisA/HisF-related TIM barrel protein [Aromatoleum aromaticum] | 04092 |
| dihydroneopterin aldolase [Aromatoleum tolulyticum] | 03484 |
| CapA family protein [Propionivibrio dicarboxylicus] | 06444 |
| DUF309 domain-containing protein [Aromatoleum aromaticum] | 00106 |
| alpha,alpha-trehalase TreF [Aromatoleum tolulyticum] | 01342 |
| OmpA family protein [Azoarcus communis] | 05811 |
| class I SAM-dependent methyltransferase [Azoarcus communis] | 01298 |
| hypothetical protein [Aromatoleum aromaticum] | 06127 |
| hypothetical protein [Aromatoleum tolulyticum] | 05817 |
| hypothetical protein [Aromatoleum aromaticum] | 01752 |
| helix-turn-helix domain-containing protein [Thauera phenylacetica] | 06329 |
| alpha-1,4-glucan--maltose-1-phosphate maltosyltransferase [Aromatoleum aromaticum] | 00757 |
| DUF4124 domain-containing protein [Azoarcus indigens] | 04511 |
| M48 family metallopeptidase [Thauera chlorobenzoica] | 09445 |
| FlgO family outer membrane protein [Thauera phenylacetica] | 03362 |
| plasmid replication initiator TrfA [Aromatoleum tolulyticum] | 04015 |
| ankyrin repeat domain-containing protein [Aromatoleum tolulyticum] | 03842 |
| PAS domain-containing protein [Aromatoleum tolulyticum] | 03528 |
| PIN domain-containing protein [Azoarcus taiwanensis] | 03101 |
| toxic anion resistance protein [Aromatoleum tolulyticum] | 01056 |
| helix-turn-helix transcriptional regulator [Azoarcus indigens] | 10339 |
| hypothetical protein [Aromatoleum aromaticum] | 08417 |
| class I SAM-dependent methyltransferase [Azoarcus indigens] | 08902 |
| pilus assembly protein TadG-related protein [Aromatoleum toluclasticum] | 06993 |
| bifunctional proline dehydrogenase/L-glutamate gamma-semialdehyde dehydrogenase PutA [Azoarcus indigens] | 00962 |
| pilus assembly protein [Aromatoleum toluclasticum] | 03476 |
| hypothetical protein [Azoarcus indigens] | 08794 |
| alpha/beta fold hydrolase [Azoarcus indigens] | 00557 |
| CoA transferase [Aromatoleum tolulyticum] | 08286 |
| hypothetical protein [Aromatoleum tolulyticum] | 09685 |
| DUF4332 domain-containing protein [Aromatoleum aromaticum] | 00971 |
| glycine/sarcosine/betaine reductase selenoprotein B family protein [Azoarcus indigens] | 03020 |
| hypothetical protein [Aromatoleum tolulyticum] | 10142 |
| helix-turn-helix domain-containing protein [Azoarcus indigens] | 09861 |
| urea carboxylase [Azoarcus communis] | 04264 |
| SEL1-like repeat protein [Aromatoleum aromaticum] | 08681 |
| hypothetical protein [Aromatoleum tolulyticum] | 07068 |
| DUF3024 domain-containing protein [Aromatoleum tolulyticum] | 08318 |
| AAA family ATPase [Zoogloea dura] | 00108 |
| alpha/beta fold hydrolase [Azoarcus indigens] | 05081 |
| DUF504 domain-containing protein [Aromatoleum diolicum] | 10046 |
| addiction module protein [Aromatoleum aromaticum] | 00512 |
| hypothetical protein [Aromatoleum diolicum] | 06231 |

| | |
|---|---|
| hypothetical protein [Azoarcus indigens] | 06427 |
| DUF1778 domain-containing protein [Aromatoleum aromaticum] | 03744 |
| DUF2138 family protein [Azoarcus indigens] | 04798 |
| hypothetical protein [Aromatoleum aromaticum] | 06348 |
| hypothetical protein [Aromatoleum aromaticum] | 03414 |
| EthD domain-containing protein [Aromatoleum tolulyticum] | 00234 |
| hypothetical protein [Azoarcus indigens] | 09925 |
| ATP-binding cassette domain-containing protein, partial [Aromatoleum toluclasticum] | 01414 |
| response regulator, partial [Aromatoleum toluclasticum] | 05230 |
| Txe/YoeB family addiction module toxin [Aromatoleum aromaticum] | 02515 |
| hypothetical protein [Azoarcus indigens] | 09429 |
| hypothetical protein [Aromatoleum tolulyticum] | 08722 |
| diguanylate cyclase [Azoarcus indigens] | 02305 |
| thiol reductant ABC exporter subunit CydD [Azoarcus indigens] | 03814 |
| amino acid ABC transporter permease [Candidatus Dactylopiibacterium carminicum] | 09672 |
| hypothetical protein [Candidatus Dactylopiibacterium carminicum] | 07190 |
| hypothetical protein [Aromatoleum aromaticum] | 03692 |
| MaoC family dehydratase [Aromatoleum aromaticum] | 05814 |
| sensor histidine kinase [Azoarcus communis] | 07183 |
| hypothetical protein [Aromatoleum toluclasticum] | 02387 |
| 2-hydroxyacyl-CoA dehydratase family protein [Aromatoleum toluclasticum] | 08402 |
| site-specific recombinase resolvase [Candidatus Dactylopiibacterium carminicum] | 05866 |
| carboxymuconolactone decarboxylase family protein [Aromatoleum tolulyticum] | 05651 |
| transporter [Aromatoleum tolulyticum] | 00238 |
| toll/interleukin-1 receptor domain-containing protein [Aromatoleum tolulyticum] | 01607 |
| heavy-metal-associated domain-containing protein [Aromatoleum tolulyticum] | 07711 |
| DsbA family protein [Thauera chlorobenzoica] | 06304 |
| PepSY domain-containing protein [Thauera chlorobenzoica] | 01595 |
| YjiH family protein [Thauera chlorobenzoica] | 04604 |
| hypothetical protein [Aromatoleum aromaticum] | 00947 |
| type I-E CRISPR-associated endoribonuclease Cas2e [Thauera chlorobenzoica] | 07792 |
| glycosyltransferase family 4 protein [Thauera chlorobenzoica] | 06705 |
| DNA topoisomerase IB [Aromatoleum aromaticum] | 08211 |
| malto-oligosyltrehalose synthase [Aromatoleum aromaticum] | 03731 |
| glycosyltransferase family 4 protein [Aromatoleum aromaticum] | 04617 |
| MULTISPECIES: MoaD/ThiS family protein [Aromatoleum] | 01183 |
| OmpA family protein [Azoarcus communis] | 07380 |
| BrnT family toxin [Thauera chlorobenzoica] | 01037 |
| ATP-grasp domain-containing protein [Aromatoleum aromaticum] | 03774 |
| hypothetical protein [Aromatoleum aromaticum] | 05657 |
| murein L,D-transpeptidase catalytic domain family protein [Azoarcus communis] | 02598 |
| rhomboid family intramembrane serine protease [Aromatoleum aromaticum] | 07986 |
| DUF1175 domain-containing protein [Azoarcus indigens] | 07414 |
| DUF1788 domain-containing protein [Aromatoleum aromaticum] | 09389 |
| cytosine permease [Aromatoleum toluclasticum] | 00500 |
| DUF1819 family protein [Aromatoleum aromaticum] | 05265 |
| hypothetical protein [Aromatoleum tolulyticum] | 10626 |
| type I-F CRISPR-associated protein Csy2 [Zoogloea dura] | 06909 |
| 3'-5' exonuclease [Azoarcus indigens] | 07567 |
| helix-turn-helix transcriptional regulator [Azoarcus indigens] | 06907 |
| DUF4043 family protein [Azoarcus indigens] | 04650 |
| glycosyltransferase family 4 protein [Aromatoleum toluclasticum] | 01811 |
| hypothetical protein [Azoarcus indigens] | 00365 |
| hypothetical protein [Azoarcus indigens] | 04328 |
| EAL domain-containing protein [Thauera terpenica] | 10471 |
| glycerol dehydrogenase [Azoarcus indigens] | 08543 |
| filamentous hemagglutinin N-terminal domain-containing protein [Azoarcus indigens] | 00831 |
| tetratricopeptide repeat protein [Aromatoleum aromaticum] | 10081 |
| methyl-accepting chemotaxis protein [Propionivibrio dicarboxylicus] | 07475 |

| | |
|---|---|
| hypothetical protein [Azoarcus indigens] | 00376 |
| sulfotransferase [Thauera terpenica] | 09684 |
| hypothetical protein [Thauera terpenica] | 00313 |
| helix-turn-helix domain-containing protein [Aromatoleum aromaticum] | 00441 |
| AAA family ATPase [Aromatoleum aromaticum] | 02897 |
| hypothetical protein [Aromatoleum tolulyticum] | 09245 |
| type I-F CRISPR-associated protein Csy1 [Zoogloea dura] | 06020 |
| hypothetical protein [Aromatoleum toluclasticum] | 00170 |
| potassium-transporting ATPase subunit KdpA [Azoarcus indigens] | 04222 |
| MULTISPECIES: hypothetical protein [Azoarcus] | 05154 |
| replication initiation factor domain-containing protein [Azoarcus indigens] | 08403 |
| nuclear transport factor 2 family protein [Zoogloea dura] | 07942 |
| cupin domain-containing protein [Azoarcus indigens] | 05133 |
| hypothetical protein [Azoarcus indigens] | 04338 |
| hypothetical protein [Aromatoleum tolulyticum] | 01865 |
| penicillin acylase family protein [Azoarcus indigens] | 10569 |
| AAA family ATPase [Aromatoleum toluclasticum] | 01784 |
| glycosyltransferase [Azoarcus indigens] | 02610 |
| DUF2135 domain-containing protein [Azoarcus indigens] | 09384 |
| DUF2300 domain-containing protein [Azoarcus indigens] | 05039 |
| AAA family ATPase [Aromatoleum aromaticum] | 07776 |
| DUF1018 domain-containing protein [Aromatoleum tolulyticum] | 09488 |
| hypothetical protein [Aromatoleum tolulyticum] | 07561 |
| hypothetical protein [Aromatoleum tolulyticum] | 10328 |
| hypothetical protein [Azoarcus indigens] | 01105 |
| heparinase II/III family protein [Thauera linaloolentis] | 10136 |
| GNAT family N-acetyltransferase [Aromatoleum tolulyticum] | 05362 |
| hypothetical protein [Aromatoleum aromaticum] | 01407 |
| DUF4395 family protein [Propionivibrio dicarboxylicus] | 08931 |
| type II toxin-antitoxin system MqsA family antitoxin [Thauera phenylacetica] | 07170 |
| sensory rhodopsin transducer [Aromatoleum aromaticum] | 00632 |
| helix-turn-helix domain-containing protein [Thauera linaloolentis] | 09265 |
| universal stress protein [Aromatoleum toluclasticum] | 00169 |
| MFS transporter [Thauera linaloolentis] | 05422 |
| tetratricopeptide repeat protein [Azoarcus rhizosphaerae] | 06694 |
| DUF2924 domain-containing protein [Propionivibrio dicarboxylicus] | 10231 |
| FHA domain-containing protein [Azoarcus rhizosphaerae] | 00069 |
| polysaccharide deacetylase family protein [Aromatoleum aromaticum] | 03636 |
| DUF2892 domain-containing protein [Aromatoleum aromaticum] | 04129 |
| DNA-binding protein [Aromatoleum toluvorans] | 08491 |
| TRAP transporter small permease subunit [Aromatoleum toluclasticum] | 03457 |
| BREX-1 system phosphatase PglZ type A [Aromatoleum aromaticum] | 06428 |
| aldehyde dehydrogenase EutE [Thauera linaloolentis] | 08408 |
| hypothetical protein [Aromatoleum tolulyticum] | 05380 |
| serine/threonine protein kinase [Aromatoleum aromaticum] | 05339 |
| helicase RepA family protein [Thauera phenolivorans] | 07433 |
| WYL domain-containing protein [Thauera phenylacetica] | 08096 |
| helix-turn-helix domain-containing protein [Zoogloea dura] | 10324 |
| hypothetical protein [Zoogloea dura] | 03242 |
| hypothetical protein [Aromatoleum aromaticum] | 09460 |
| AAA family ATPase [Thauera linaloolentis] | 02038 |
| hypothetical protein [Aromatoleum toluclasticum] | 08544 |
| glycosyltransferase [Aromatoleum aromaticum] | 00259 |
| 4-alpha-glucanotransferase [Thauera linaloolentis] | 04482 |
| hypothetical protein [Aromatoleum toluclasticum] | 00231 |
| gamma-glutamylcyclotransferase [Aromatoleum toluclasticum] | 09640 |
| hypothetical protein [Aromatoleum aromaticum] | 09239 |
| FAD-binding protein, partial [Thauera phenylacetica] | 02592 |
| vinylacetyl-CoA Delta-isomerase [Thauera linaloolentis] | 06381 |

| | |
|---|---|
| DsrE family protein [Thauera linaloolentis] | 10196 |
| hypothetical protein [Aromatoleum toluclasticum] | 09997 |
| glycine cleavage system protein H [Thauera linaloolentis] | 08937 |
| D-serine ammonia-lyase [Aromatoleum toluclasticum] | 00658 |
| GtrA family protein [Zoogloea dura] | 05494 |
| SEC-C domain-containing protein [Aromatoleum aromaticum] | 08702 |
| DUF47 family protein [Thauera linaloolentis] | 01857 |
| transporter substrate-binding domain-containing protein [Propionivibrio dicarboxylicus] | 00754 |
| DUF2474 family protein [Aromatoleum aromaticum] | 08684 |
| Wzz/FepE/Etk N-terminal domain-containing protein [Aromatoleum aromaticum] | 08479 |
| AAA family ATPase [Zoogloea dura] | 07456 |
| replication protein [Azoarcus communis] | 07912 |
| NAD(P)H-dependent oxidoreductase [Azoarcus rhizosphaerae] | 04864 |
| DUF5681 domain-containing protein [Aromatoleum aromaticum] | 08067 |
| DUF192 domain-containing protein [Thauera phenylacetica] | 07131 |
| GTP-binding protein [Thauera linaloolentis] | 04001 |
| hypothetical protein [Aromatoleum aromaticum] | 07167 |
| ATP-binding protein [Aromatoleum aromaticum] | 09813 |
| hypothetical protein [Thauera linaloolentis] | 05998 |
| cytochrome C oxidase subunit IV family protein [Aromatoleum toluclasticum] | 04492 |
| toluene-4-monooxygenase system B family protein [Aromatoleum toluclasticum] | 08387 |
| ChaN family lipoprotein [Aromatoleum aromaticum] | 05762 |
| nuclear transport factor 2 family protein [Zoogloea dura] | 09842 |
| DUF4178 domain-containing protein [Zoogloea dura] | 05884 |
| BREX-1 system adenine-specific DNA-methyltransferase PglX [Azoarcus communis] | 00008 |
| ABC transporter permease [Aromatoleum tolulyticum] | 05222 |
| pentaheme c-type cytochrome TorC [Thauera phenylacetica] | 09874 |
| hypothetical protein, partial [Thauera phenylacetica] | 09814 |
| transposase [Thauera linaloolentis] | 10002 |
| DUF2783 domain-containing protein [Azoarcus indigens] | 06323 |
| DUF3164 family protein [Aromatoleum tolulyticum] | 01990 |
| DUF6036 family nucleotidyltransferase [Aromatoleum aromaticum] | 01358 |
| DUF4254 domain-containing protein [Azoarcus indigens] | 07968 |
| hypothetical protein [Thauera chlorobenzoica] | 02332 |
| hypothetical protein [Aromatoleum tolulyticum] | 09522 |
| hypothetical protein [Aromatoleum tolulyticum] | 07242 |
| hypothetical protein [Thauera linaloolentis] | 02827 |
| entry exclusion lipoprotein TrbK [Aromatoleum aromaticum] | 09891 |
| type II toxin-antitoxin system death-on-curing family toxin [Thauera linaloolentis] | 07859 |
| type I secretion C-terminal target domain-containing protein [Aromatoleum tolulyticum] | 03856 |
| glycine zipper 2TM domain-containing protein [Aromatoleum tolulyticum] | 07398 |
| hypothetical protein [Aromatoleum toluclasticum] | 06496 |
| APC family permease [Azoarcus communis] | 03578 |
| DUF2703 domain-containing protein [Thauera phenylacetica] | 06405 |
| relaxase/mobilization nuclease domain-containing protein [Aromatoleum aromaticum] | 07738 |
| DUF3304 domain-containing protein [Aromatoleum tolulyticum] | 01564 |
| hypothetical protein [Thauera phenylacetica] | 10377 |
| FtsX-like permease family protein [Thauera phenylacetica] | 09457 |
| glycosyltransferase [Thauera phenylacetica] | 07780 |
| retention module-containing protein [Azoarcus indigens] | 03021 |
| hypothetical protein [Thauera phenylacetica] | 06373 |
| hypothetical protein [Aromatoleum tolulyticum] | 05822 |
| hemerythrin domain-containing protein [Azoarcus indigens] | 05677 |
| radical SAM protein [Aromatoleum aromaticum] | 02530 |
| hypothetical protein [Azoarcus indigens] | 07135 |
| Z1 domain-containing protein [Aromatoleum aromaticum] | 06679 |
| hypothetical protein [Thauera phenylacetica] | 04038 |
| class I SAM-dependent methyltransferase [Azoarcus communis] | 03294 |
| MULTISPECIES: BphX family protein [Rhodocyclales] | 08235 |

| | |
|---|---|
| PepSY domain-containing protein [Azoarcus indigens] | 07944 |
| transporter substrate-binding domain-containing protein [Thauera chlorobenzoica] | 04368 |
| single-stranded DNA-binding protein [Thauera chlorobenzoica] | 05790 |
| hypothetical protein [Uliginosibacterium aquaticum] | 01802 |
| DUF4160 domain-containing protein [Thauera chlorobenzoica] | 02275 |
| type II toxin-antitoxin system RelE/ParE family toxin [Azoarcus indigens] | 03895 |
| AAA family ATPase [Azoarcus communis] | 00435 |
| MULTISPECIES: ATP-binding protein [Rhodocyclales] | 08083 |
| hypothetical protein [Aromatoleum tolulyticum] | 06971 |
| thioesterase superfamily protein [Thauera linaloolentis] | 04674 |
| MULTISPECIES: MFS transporter [Thauera] | 08727 |
| MULTISPECIES: phosphoadenosine phosphosulfate reductase family protein [Rhodocyclales] | 05184 |
| UvrD-helicase domain-containing protein [Aromatoleum aromaticum] | 02361 |
| hypothetical protein [Azoarcus indigens] | 00378 |
| WD40 repeat domain-containing protein [Azoarcus indigens] | 03749 |
| hypothetical protein [Azoarcus indigens] | 06376 |
| MmoB/DmpM family protein [Azoarcus indigens] | 00047 |
| alkaline phosphatase D family protein [Aromatoleum aromaticum] | 07990 |
| nitrous oxide reductase accessory protein NosL [Thauera phenylacetica] | 04722 |
| hypothetical protein [Aromatoleum aromaticum] | 06870 |
| glycosyltransferase family 29 protein [Azoarcus communis] | 02659 |
| exopolysaccharide biosynthesis protein [Aromatoleum aromaticum] | 04200 |
| hypothetical protein [Propionivibrio dicarboxylicus] | 10425 |
| hypothetical protein [Azoarcus indigens] | 04346 |
| DUF3445 domain-containing protein [Azoarcus communis] | 01137 |
| PAS domain-containing protein [Azoarcus indigens] | 00812 |
| hypothetical protein [Thauera linaloolentis] | 00206 |
| Wzy polymerase domain-containing protein [Uliginosibacterium aquaticum] | 04865 |
| AAA family ATPase [Aromatoleum tolulyticum] | 06255 |
| 2OG-Fe(II) oxygenase family protein [Aromatoleum toluclasticum] | 02287 |
| polysaccharide pyruvyl transferase family protein [Azoarcus communis] | 06404 |
| terminase small subunit [Azoarcus indigens] | 08270 |
| TetR/AcrR family transcriptional regulator [Thauera linaloolentis] | 09479 |
| NUDIX domain-containing protein [Zoogloea dura] | 06435 |
| DUF4254 domain-containing protein [Aromatoleum aromaticum] | 03282 |
| hypothetical protein [Aromatoleum aromaticum] | 04097 |
| metallophosphoesterase [Azoarcus communis] | 00511 |
| hypothetical protein [Azoarcus communis] | 06840 |
| transposase [Zoogloea dura] | 08313 |
| cupin domain-containing protein [Propionivibrio dicarboxylicus] | 00838 |
| VOC family protein [Thauera linaloolentis] | 00214 |
| glycosyltransferase [Aromatoleum toluclasticum] | 02799 |
| EthD domain-containing protein [Zoogloea dura] | 07436 |
| hypothetical protein [Azoarcus indigens] | 02108 |
| hypothetical protein [Azoarcus indigens] | 08192 |
| hypothetical protein [Aromatoleum tolulyticum] | 01584 |
| CIA30 family protein [Thauera phenylacetica] | 02111 |
| hypothetical protein [Azoarcus communis] | 09094 |
| hypothetical protein [Aromatoleum tolulyticum] | 10084 |
| hypothetical protein [Aromatoleum tolulyticum] | 02281 |
| helix-turn-helix domain-containing protein [Aromatoleum tolulyticum] | 06727 |
| antitoxin MazE family protein [Aromatoleum aromaticum] | 01018 |
| hypothetical protein [Aromatoleum tolulyticum] | 07713 |
| cytochrome b/b6 domain-containing protein [Azoarcus indigens] | 04170 |
| pentapeptide MXKDX repeat protein [Azoarcus indigens] | 08191 |
| DUF4124 domain-containing protein [Azoarcus communis] | 06119 |
| conjugal transfer transcriptional regulator TraJ [Aromatoleum aromaticum] | 02820 |
| SpoIIE family protein phosphatase [Azoarcus indigens] | 05095 |
| DUF3455 domain-containing protein [Aromatoleum aromaticum] | 08070 |

| | |
|---|---|
| SPOR domain-containing protein, partial [Thauera phenylacetica] | 03997 |
| RHS repeat protein [Azoarcus indigens] | 04393 |
| hypothetical protein [Azoarcus indigens] | 05156 |
| FUSC family protein [Azoarcus indigens] | 06732 |
| type II toxin-antitoxin system VapC family toxin [Aromatoleum aromaticum] | 01426 |
| helix-turn-helix domain-containing protein [Aromatoleum aromaticum] | 09806 |
| molecular chaperone TorD [Thauera phenylacetica] | 08752 |
| MULTISPECIES: TAXI family TRAP transporter solute-binding subunit [Aromatoleum] | 05416 |
| FdtA/QdtA family cupin domain-containing protein [Aromatoleum tolulyticum] | 00553 |
| TonB-system energizer ExbB [Azoarcus indigens] | 07145 |
| terminase family protein [Azoarcus communis] | 06804 |
| hypothetical protein [Azoarcus indigens] | 08509 |
| helix-turn-helix domain-containing protein [Thauera terpenica] | 08100 |
| glycoside hydrolase family 5 protein [Uliginosibacterium gangwonense] | 09839 |
| zinc ribbon domain-containing protein [Azoarcus sp. L1K30] | 03188 |
| flocculation-associated PEP-CTERM protein PepA [Aromatoleum aromaticum] | 09993 |
| hypothetical protein [Aromatoleum aromaticum] | 08119 |
| PRC-barrel domain-containing protein [Aromatoleum aromaticum] | 07008 |
| prepilin-type N-terminal cleavage/methylation domain-containing protein [Aromatoleum aromaticum] | 04736 |
| hypothetical protein [Aromatoleum aromaticum] | 10578 |
| TnsA endonuclease N-terminal domain-containing protein [Aromatoleum tolulyticum] | 09523 |
| branched-chain amino acid ABC transporter permease, partial [Thauera phenylacetica] | 03479 |
| amidohydrolase [Zoogloea dura] | 03993 |
| L-lactate dehydrogenase [Azoarcus indigens] | 04044 |
| COG2958 family protein [Candidatus Dactylopiibacterium carminicum] | 01914 |
| ferrous iron transport protein A [Thauera linaloolentis] | 07745 |
| four-carbon acid sugar kinase family protein [Propionivibrio dicarboxylicus] | 01609 |
| ferredoxin [Zoogloea dura] | 00290 |
| DUF6152 family protein [Azoarcus indigens] | 06041 |
| SGNH/GDSL hydrolase family protein [Azoarcus indigens] | 09550 |
| flagellar assembly peptidoglycan hydrolase FlgJ [Azoarcus indigens] | 09607 |
| SLC13 family permease [Thauera chlorobenzoica] | 07389 |
| MarR family transcriptional regulator [Aromatoleum aromaticum] | 08720 |
| helix-turn-helix domain-containing protein [Thauera chlorobenzoica] | 10051 |
| type II toxin-antitoxin system RelE/ParE family toxin [Candidatus Dactylopiibacterium carminicum] | 06551 |
| PIN domain-containing protein [Thauera linaloolentis] | 03410 |
| S8 family serine peptidase [Aromatoleum aromaticum] | 05035 |
| nuclear transport factor 2 family protein [Aromatoleum toluclasticum] | 04134 |
| TonB family protein [Propionivibrio dicarboxylicus] | 09878 |
| hypothetical protein [Zoogloea dura] | 10132 |
| hypothetical protein [Thauera chlorobenzoica] | 09862 |
| queuosine precursor transporter [Azoarcus indigens] | 03490 |
| 4-oxalocrotonate tautomerase [Propionivibrio dicarboxylicus] | 05844 |
| hypothetical protein [Thauera linaloolentis] | 01002 |
| DUF3597 domain-containing protein [Aromatoleum tolulyticum] | 02475 |
| PadR family transcriptional regulator [Azoarcus indigens] | 09600 |
| ribbon-helix-helix domain-containing protein [Aromatoleum aromaticum] | 10337 |
| head decoration protein [Aromatoleum aromaticum] | 05727 |
| hypothetical protein [Aromatoleum toluclasticum] | 01011 |
| hypothetical protein [Aromatoleum toluclasticum] | 09824 |
| hypothetical protein [Aromatoleum toluclasticum] | 01190 |
| hypothetical protein [Thauera linaloolentis] | 02379 |
| porin [Uliginosibacterium aquaticum] | 00690 |
| VOC family protein [Azoarcus communis] | 04557 |
| hypothetical protein [Aromatoleum aromaticum] | 04913 |
| VanZ family protein [Aromatoleum tolulyticum] | 06306 |
| hypothetical protein [Thauera chlorobenzoica] | 08757 |
| DNA primase family protein [Thauera chlorobenzoica] | 01602 |
| alpha/beta fold hydrolase [Azoarcus indigens] | 02941 |

| | |
|---|---|
| type II toxin-antitoxin system RelE/ParE family toxin [Azoarcus communis] | 03433 |
| hypothetical protein [Candidatus Dactylopiibacterium carminicum] | 10035 |
| lactoylglutathione lyase [Zoogloea dura] | 01444 |
| GspH/FimT family pseudopilin [Aromatoleum aromaticum] | 09331 |
| tape measure protein [Aromatoleum aromaticum] | 10179 |
| DUF4917 family protein [Azoarcus communis] | 04858 |
| hypothetical protein [Aromatoleum toluclasticum] | 05675 |
| hypothetical protein [Azoarcus communis] | 09084 |
| heavy-metal-associated domain-containing protein [Azoarcus communis] | 01497 |
| choice-of-anchor A family protein [Thauera linaloolentis] | 07815 |
| ribbon-helix-helix domain-containing protein [Aromatoleum aromaticum] | 04661 |
| RNA-guided endonuclease TnpB family protein [Aromatoleum aromaticum] | 05906 |
| BREX system serine/threonine kinase PglW [Azoarcus indigens] | 05745 |
| LysR family transcriptional regulator [Pseudomonas carnis] | 00433 |
| hypothetical protein [Thauera chlorobenzoica] | 05417 |
| hypothetical protein [Azoarcus indigens] | 00368 |
| hypothetical protein [Zoogloea dura] | 08296 |
| hypothetical protein [Azoarcus indigens] | 09810 |
| ATP-binding protein [Aromatoleum tolulyticum] | 01817 |
| hypothetical protein [Propionivibrio dicarboxylicus] | 07127 |
| capsular polysaccharide biosynthesis protein [Aromatoleum aromaticum] | 02848 |
| type II secretion system protein GspL [Azoarcus indigens] | 01218 |
| Wzz/FepE/Etk N-terminal domain-containing protein [Aromatoleum diolicum] | 10289 |
| MULTISPECIES: transposase [Rhodocyclales] | 00133 |
| DUF1376 domain-containing protein [Aromatoleum aromaticum] | 00252 |
| hypothetical protein [Azoarcus communis] | 04096 |
| RNA polymerase sigma factor [Thauera linaloolentis] | 09906 |
| hypothetical protein [Aromatoleum aromaticum] | 08184 |
| AAA family ATPase [Azoarcus communis] | 07617 |
| MATE family efflux transporter [Uliginosibacterium aquaticum] | 06315 |
| bifunctional lysylphosphatidylglycerol flippase/synthetase MprF [Thauera linaloolentis] | 05276 |
| TAXI family TRAP transporter solute-binding subunit [Zoogloea dura] | 00700 |
| hypothetical protein [Thauera aromatica] | 05792 |
| 7-cyano-7-deazaguanine synthase [Aromatoleum aromaticum] | 06833 |
| hypothetical protein [Azoarcus indigens] | 09926 |
| DUF2191 domain-containing protein [Azoarcus taiwanensis] | 07562 |
| diaminobutyrate acetyltransferase [Thauera linaloolentis] | 03344 |
| DUF29 domain-containing protein [Aromatoleum diolicum] | 03309 |
| TonB-dependent receptor [Zoogloea dura] | 05952 |
| ectoine synthase [Thauera linaloolentis] | 07437 |
| type II toxin-antitoxin system prevent-host-death family antitoxin [Thauera linaloolentis] | 02039 |
| LuxR C-terminal-related transcriptional regulator [Azoarcus indigens] | 03299 |
| type IV pilin protein [Aromatoleum aromaticum] | 06232 |
| SPFH domain-containing protein [Zoogloea dura] | 01211 |
| hypothetical protein [Azoarcus communis] | 00546 |
| MTH1187 family thiamine-binding protein [Thauera chlorobenzoica] | 07673 |
| DUF2875 family protein [Azoarcus indigens] | 00385 |
| DUF2971 domain-containing protein [Azoarcus indigens] | 03165 |
| Abi family protein [Aromatoleum aromaticum] | 09735 |
| Mu transposase C-terminal domain-containing protein [Aromatoleum tolulyticum] | 10144 |
| 5'-nucleotidase [Propionivibrio dicarboxylicus] | 07965 |
| MULTISPECIES: hypothetical protein [Azoarcus] | 04726 |
| energy transducer TonB [Azoarcus indigens] | 00585 |
| type II toxin-antitoxin system VapC family toxin [Thauera phenylacetica] | 00469 |
| DUF2059 domain-containing protein [Azoarcus indigens] | 04774 |
| MotA/TolQ/ExbB proton channel family protein [Azoarcus indigens] | 02841 |
| HIRAN domain-containing protein [Propionivibrio dicarboxylicus] | 04385 |
| hypothetical protein [Aromatoleum toluclasticum] | 02239 |
| GtrA family protein [Zoogloea dura] | 02135 |

| | |
|---|---|
| ectoine hydroxylase [Thauera linaloolentis] | 03986 |
| choice-of-anchor L domain-containing protein [Azoarcus indigens] | 10363 |
| ATP-binding protein [Azoarcus communis] | 02542 |
| terminase family protein [Propionivibrio dicarboxylicus] | 07489 |
| hypothetical protein [Aromatoleum aromaticum] | 01032 |
| WbuC family cupin fold metalloprotein [Propionivibrio dicarboxylicus] | 10116 |
| hypothetical protein [Thauera linaloolentis] | 04666 |
| DUF2442 domain-containing protein [Aromatoleum aromaticum] | 06805 |
| YqaJ viral recombinase family protein [Aromatoleum aromaticum] | 07267 |
| DUF3368 domain-containing protein [Candidatus Dactylopiibacterium carminicum] | 04871 |
| methyltransferase domain-containing protein [Candidatus Dactylopiibacterium carminicum] | 01496 |
| tetratricopeptide repeat protein [Aromatoleum tolulyticum] | 10135 |
| hypothetical protein [Thauera chlorobenzoica] | 02708 |
| flagellar basal body P-ring formation chaperone FlgA [Azoarcus indigens] | 09724 |
| hypothetical protein [Zoogloea dura] | 02212 |
| bacteriohemerythrin [Uliginosibacterium aquaticum] | 04577 |
| DUF3396 domain-containing protein [Azoarcus indigens] | 07817 |
| prepilin-type N-terminal cleavage/methylation domain-containing protein [Propionivibrio dicarboxylicus] | 08845 |
| MULTISPECIES: hypothetical protein [Thauera] | 02660 |
| helix-turn-helix domain-containing protein [Azoarcus indigens] | 05698 |
| glycoside hydrolase family 15 protein [Azoarcus indigens] | 04884 |
| PRC-barrel domain-containing protein [Aromatoleum aromaticum] | 09659 |
| hypothetical protein [Candidatus Dactylopiibacterium carminicum] | 02842 |
| type IV pili methyl-accepting chemotaxis transducer N-terminal domain-containing protein [Thauera phenylacetica] | 02830 |
| hypothetical protein [Azoarcus indigens] | 02394 |
| tetratricopeptide repeat protein [Azoarcus communis] | 02898 |
| hypothetical protein [Zoogloea dura] | 07955 |
| O-antigen ligase family protein [Thauera phenylacetica] | 04464 |
| hypothetical protein [Azoarcus rhizosphaerae] | 08199 |
| Fis family transcriptional regulator [Azoarcus indigens] | 00140 |
| hypothetical protein [Thauera phenylacetica] | 07499 |
| TIGR02391 family protein [Thauera chlorobenzoica] | 08549 |
| alpha/beta fold hydrolase [Thauera phenylacetica] | 09041 |
| histidine phosphatase family protein [Thauera chlorobenzoica] | 09924 |
| hypothetical protein [Zoogloea dura] | 01881 |
| hypothetical protein [Thauera chlorobenzoica] | 03825 |
| plasmid replication initiator TrfA [Thauera phenylacetica] | 00270 |
| hypothetical protein [Thauera chlorobenzoica] | 01016 |
| pilin [Thauera chlorobenzoica] | 04203 |
| MULTISPECIES: hypothetical protein [Rhodocyclales] | 08279 |
| ethanolamine ammonia-lyase reactivating factor EutA [Thauera linaloolentis] | 06395 |
| efflux RND transporter periplasmic adaptor subunit [Propionivibrio dicarboxylicus] | 03939 |
| DUF1553 domain-containing protein [Zoogloea dura] | 06945 |
| toll/interleukin-1 receptor domain-containing protein [Azoarcus communis] | 08550 |
| hypothetical protein [Azoarcus communis] | 05431 |
| S8 family peptidase [Azoarcus communis] | 00074 |
| virulence factor family protein [Thauera linaloolentis] | 05591 |
| hypothetical protein [Aromatoleum tolulyticum] | 02732 |
| nucleotidyltransferase domain-containing protein [Azoarcus communis] | 06546 |
| DUF4880 domain-containing protein [Thauera linaloolentis] | 01877 |
| MULTISPECIES: hypothetical protein [Zoogloeaceae] | 05886 |
| PASTA domain-containing protein [Aromatoleum tolulyticum] | 02199 |
| DUF1501 domain-containing protein [Zoogloea dura] | 04891 |
| NAD(P)/FAD-dependent oxidoreductase [Uliginosibacterium aquaticum] | 01001 |
| hypothetical protein [Aromatoleum tolulyticum] | 05335 |
| hypothetical protein [Thauera chlorobenzoica] | 05711 |
| hypothetical protein [Zoogloea dura] | 05979 |
| TonB-dependent receptor [Thauera linaloolentis] | 01142 |
| AAA family ATPase [Zoogloea dura] | 03184 |

| | |
|---|---|
| type II toxin-antitoxin system Phd/YefM family antitoxin [Thauera phenylacetica] | 08099 |
| NifU family protein [Thauera chlorobenzoica] | 06478 |
| ImmA/IrrE family metallo-endopeptidase [Propionivibrio dicarboxylicus] | 05379 |
| hypothetical protein [Thauera chlorobenzoica] | 01054 |
| type II toxin-antitoxin system VapC family toxin [Thauera phenylacetica] | 09820 |
| hypothetical protein [Zoogloea dura] | 05192 |
| hypothetical protein [Thauera chlorobenzoica] | 06706 |
| hypothetical protein [Thauera chlorobenzoica] | 04689 |
| hypothetical protein [Thauera chlorobenzoica] | 03676 |
| ATP-binding protein [Thauera chlorobenzoica] | 01356 |
| DUF6491 family protein [Thauera linaloolentis] | 03405 |
| DUF3817 domain-containing protein [Thauera linaloolentis] | 10332 |
| YkgJ family cysteine cluster protein [Uliginosibacterium aquaticum] | 00720 |
| hypothetical protein [Aromatoleum aromaticum] | 04292 |
| DNA cytosine methyltransferase [Zoogloea dura] | 09114 |
| patatin-like protein [Thauera chlorobenzoica] | 09063 |
| hypothetical protein [Propionivibrio dicarboxylicus] | 00486 |
| hypothetical protein [Zoogloea dura] | 02742 |
| DUF2442 domain-containing protein [Thauera chlorobenzoica] | 01862 |
| type II toxin-antitoxin system PemK/MazF family toxin [Aromatoleum aromaticum] | 00464 |
| hypothetical protein [Aromatoleum aromaticum] | 02511 |
| helix-hairpin-helix domain-containing protein [Uliginosibacterium aquaticum] | 09713 |
| exosortase H [Zoogloea dura] | 09764 |
| hypothetical protein [Zoogloea dura] | 00446 |
| cupin domain-containing protein [Aromatoleum tolulyticum] | 08111 |
| hypothetical protein [Azoarcus communis] | 03187 |
| zf-HC2 domain-containing protein [Aromatoleum tolulyticum] | 09795 |
| hypothetical protein [Thauera phenylacetica] | 06278 |
| hypothetical protein [Azoarcus indigens] | 08078 |
| hypothetical protein [Azoarcus communis] | 09515 |
| NirD/YgiW/YdeI family stress tolerance protein [Azoarcus indigens] | 02424 |
| MFS transporter [Thauera linaloolentis] | 09986 |
| TSUP family transporter [Azoarcus indigens] | 03859 |
| nuclear transport factor 2 family protein [Aromatoleum tolulyticum] | 02797 |
| Ti-type conjugative transfer relaxase TraA [Thauera chlorobenzoica] | 00128 |
| MULTISPECIES: broad-spectrum mercury transporter MerE [Proteobacteria] | 04004 |
| DEAD/DEAH box helicase [Aromatoleum aromaticum] | 07344 |
| sterol desaturase family protein [Azoarcus indigens] | 08280 |
| chemotaxis protein CheW [Azoarcus rhizosphaerae] | 08903 |
| CRISPR-associated endonuclease Cas2 [Thauera phenylacetica] | 08530 |
| nuclear transport factor 2 family protein [Azoarcus communis] | 05239 |
| DNA-binding protein [Aromatoleum aromaticum] | 01160 |
| putative DNA-binding domain-containing protein [Zoogloea dura] | 04825 |
| HAD hydrolase-like protein [Aromatoleum aromaticum] | 01039 |
| hypothetical protein [Zoogloea dura] | 06152 |
| MULTISPECIES: PD-(D/E)XK nuclease family protein [Betaproteobacteria] | 09665 |
| hypothetical protein [Uliginosibacterium aquaticum] | 05563 |
| peptidylprolyl isomerase [Azoarcus communis] | 00768 |
| MULTISPECIES: hypothetical protein [Rhodocyclales] | 04473 |
| DUF2442 domain-containing protein [Thauera chlorobenzoica] | 07422 |
| MULTISPECIES: DUF87 domain-containing protein [Rhodocyclales] | 02675 |
| MULTISPECIES: hypothetical protein [Rhodocyclales] | 08904 |
| MULTISPECIES: hypothetical protein [Rhodocyclales] | 06761 |
| DUF4394 domain-containing protein [Azoarcus communis] | 00346 |
| OmpA family protein [Azoarcus sp. L1K30] | 03105 |
| MULTISPECIES: hypothetical protein [Rhodocyclales] | 01300 |
| MULTISPECIES: DUF4007 family protein [Rhodocyclales] | 01133 |
| PEP-CTERM sorting domain-containing protein [Zoogloea ramigera] | 06635 |
| MULTISPECIES: hypothetical protein [Rhodocyclales] | 09687 |

| | |
|---|---|
| PIG-L family deacetylase [Thauera linaloolentis] | 07496 |
| spore maturation protein [Zoogloea dura] | 07012 |
| hypothetical protein [Zoogloea dura] | 09180 |
| prepilin-type N-terminal cleavage/methylation domain-containing protein [Propionivibrio dicarboxylicus] | 06260 |
| IS200/IS605-like element ISAzo20 family transposase [Aromatoleum aromaticum] | 05656 |
| TetR/AcrR family transcriptional regulator [Thauera linaloolentis] | 03656 |
| hypothetical protein [Azoarcus indigens] | 07945 |
| primase C-terminal domain-containing protein [Thauera phenylacetica] | 03351 |
| PEP-CTERM sorting domain-containing protein [Aromatoleum tolulyticum] | 01199 |
| hypothetical protein [Zoogloea dura] | 01034 |
| hypothetical protein [Azoarcus communis] | 02375 |
| coenzyme A transferase [Thauera linaloolentis] | 04724 |
| hypothetical protein [Aromatoleum aromaticum] | 01998 |
| type II toxin-antitoxin system PemK/MazF family toxin [Aromatoleum toluclasticum] | 01188 |
| retention module-containing protein [Aromatoleum aromaticum] | 07750 |
| Fic family protein [Azoarcus communis] | 08432 |
| transcriptional regulator KorA [Aromatoleum aromaticum] | 01670 |
| hypothetical protein [Aromatoleum tolulyticum] | 07699 |
| GIY-YIG nuclease family protein [Aromatoleum tolulyticum] | 05251 |
| hypothetical protein [Aromatoleum toluclasticum] | 03648 |
| hypothetical protein [Azoarcus indigens] | 02821 |
| hypothetical protein [Aromatoleum tolulyticum] | 10329 |
| hypothetical protein [Aromatoleum aromaticum] | 06466 |
| hypothetical protein [Aromatoleum toluclasticum] | 02838 |
| DUF1214 domain-containing protein [Thauera terpenica] | 02780 |
| response regulator [Aromatoleum tolulyticum] | 08102 |
| hypothetical protein [Aromatoleum aromaticum] | 02167 |
| DUF4276 family protein [Aromatoleum aromaticum] | 10277 |
| hypothetical protein [Aromatoleum aromaticum] | 02000 |
| DUF4438 domain-containing protein [Aromatoleum tolulyticum] | 01650 |
| patatin-like phospholipase family protein [Azoarcus indigens] | 07782 |
| hypothetical protein [Aromatoleum tolulyticum] | 08287 |
| hypothetical protein [Aromatoleum tolulyticum] | 04299 |
| DNA methyltransferase [Aromatoleum aromaticum] | 08699 |
| hypothetical protein [Thauera linaloolentis] | 07510 |
| hypothetical protein [Thauera phenylacetica] | 06698 |
| HPF/RaiA family ribosome-associated protein [Aromatoleum aromaticum] | 09002 |
| DUF4105 domain-containing protein [Azoarcus indigens] | 10344 |
| hypothetical protein [Aromatoleum tolulyticum] | 05619 |
| hypothetical protein [Thauera terpenica] | 07586 |
| hypothetical protein [Propionivibrio dicarboxylicus] | 00727 |
| MULTISPECIES: type II toxin-antitoxin system PrlF family antitoxin [Proteobacteria] | 06877 |
| hypothetical protein [Azoarcus communis] | 03459 |
| substrate-binding domain-containing protein [Azoarcus indigens] | 05737 |
| hypothetical protein [Aromatoleum aromaticum] | 00953 |
| hypothetical protein [Azoarcus rhizosphaerae] | 10124 |
| hypothetical protein [Azoarcus indigens] | 09489 |
| c-type cytochrome [Aromatoleum tolulyticum] | 08719 |
| diaminopimelate dehydrogenase [Thauera linaloolentis] | 00848 |
| alpha/beta fold hydrolase [Thauera linaloolentis] | 08124 |
| hypothetical protein [Aromatoleum diolicum] | 07501 |
| DUF6484 domain-containing protein [Thauera phenylacetica] | 02261 |
| carboxynorspermidine decarboxylase [Thauera linaloolentis] | 08738 |
| AAA family ATPase, partial [Thauera phenylacetica] | 10430 |
| hypothetical protein [Azoarcus indigens] | 09882 |
| hypothetical protein [Azoarcus indigens] | 09908 |
| hypothetical protein [Azoarcus indigens] | 05377 |
| DUF1835 domain-containing protein [Aromatoleum aromaticum] | 07313 |
| chemotaxis protein CheW [Aromatoleum tolulyticum] | 03474 |

| | |
|---|---|
| Hpt domain-containing protein, partial [Thauera phenylacetica] | 08633 |
| hypothetical protein [Azoarcus communis] | 05229 |
| hypothetical protein [Azoarcus indigens] | 04386 |
| hypothetical protein [Aromatoleum toluclasticum] | 09883 |
| sulfotransferase [Azoarcus indigens] | 00045 |
| DNA alkylation repair protein [Azoarcus indigens] | 03045 |
| haloacid dehalogenase-like hydrolase [Thauera linaloolentis] | 07492 |
| nuclear transport factor 2 family protein [Azoarcus indigens] | 10233 |
| PQQ-binding-like beta-propeller repeat protein [Aromatoleum tolulyticum] | 05537 |
| hypothetical protein [Aromatoleum tolulyticum] | 02510 |
| helix-turn-helix domain-containing protein [Aromatoleum tolulyticum] | 00109 |
| FliM/FliN family flagellar motor switch protein [Azoarcus indigens] | 06143 |
| FliM/FliN family flagellar motor C-terminal domain-containing protein [Azoarcus indigens] | 03850 |
| flagellar FliJ family protein [Azoarcus indigens] | 02738 |
| hypothetical protein [Azoarcus indigens] | 10521 |
| flagellar hook-length control protein FliK [Azoarcus indigens] | 00141 |
| flagellar basal body protein [Azoarcus indigens] | 10488 |
| hypothetical protein [Thauera phenolivorans] | 10333 |
| flagellar biosynthesis anti-sigma factor FlgM [Azoarcus indigens] | 08498 |
| flagellar export chaperone FlgN [Azoarcus indigens] | 09601 |
| flagellar basal body protein FlgE [Azoarcus indigens] | 08552 |
| hypothetical protein [Aromatoleum tolulyticum] | 00113 |
| saccharopine dehydrogenase family protein [Thauera linaloolentis] | 09877 |
| helix-turn-helix transcriptional regulator [Azoarcus indigens] | 06328 |
| hypothetical protein [Rhodocyclus purpureus] | 01260 |
| hypothetical protein [Propionivibrio dicarboxylicus] | 09364 |
| hypothetical protein [Azoarcus indigens] | 00361 |
| XRE family transcriptional regulator [Aromatoleum aromaticum] | 05937 |
| hypothetical protein [Aromatoleum tolulyticum] | 09678 |
| SIR2 family protein [Aromatoleum aromaticum] | 01845 |
| hypothetical protein [Thauera terpenica] | 04844 |
| hypothetical protein, partial [Aromatoleum toluclasticum] | 01434 |
| hypothetical protein [Aromatoleum aromaticum] | 09915 |
| hypothetical protein [Aromatoleum aromaticum] | 09868 |
| metal-dependent hydrolase [Aromatoleum tolulyticum] | 02367 |
| hypothetical protein [Aromatoleum tolulyticum] | 09801 |
| MarR family transcriptional regulator [Azoarcus indigens] | 02929 |
| hypothetical protein [Azoarcus indigens] | 04880 |
| type II secretion system protein GspM [Azoarcus indigens] | 08440 |
| Ig-like domain-containing protein [Aromatoleum aromaticum] | 00551 |
| hypothetical protein [Aromatoleum diolicum] | 09064 |
| BREX-2 system phosphatase PglZ [Azoarcus indigens] | 09102 |
| hypothetical protein [Azoarcus indigens] | 07039 |
| hypothetical protein [Azoarcus indigens] | 07787 |
| hypothetical protein [Aromatoleum toluclasticum] | 01958 |
| hypothetical protein [Aromatoleum aromaticum] | 08486 |
| hypothetical protein [Aromatoleum tolulyticum] | 02706 |
| hypothetical protein [Aromatoleum diolicum] | 10588 |
| DUF2304 domain-containing protein [Aromatoleum aromaticum] | 09437 |
| hypothetical protein [Aromatoleum diolicum] | 09483 |
| oxalurate catabolism protein HpxZ [Azoarcus communis] | 09979 |
| urea carboxylase-associated family protein [Azoarcus communis] | 01682 |
| tetratricopeptide repeat protein [Aromatoleum aromaticum] | 06887 |
| transposase, partial [Thauera terpenica] | 06803 |
| hypothetical protein [Azoarcus indigens] | 09105 |
| glycosyltransferase [Azoarcus sp. DD4] | 07651 |
| hypothetical protein [Aromatoleum toluclasticum] | 09432 |
| DDE-type integrase/transposase/recombinase [Azoarcus communis] | 07739 |
| glycoside hydrolase [Aromatoleum tolulyticum] | 10487 |

| | |
|---|---|
| type II toxin-antitoxin system VapC family toxin [Aromatoleum aromaticum] | 09106 |
| DUF1788 domain-containing protein [Thauera aromatica] | 07529 |
| lysine N(6)-hydroxylase/L-ornithine N(5)-oxygenase family protein [Azoarcus indigens] | 04192 |
| hypothetical protein [Aromatoleum toluclasticum] | 00264 |
| hypothetical protein [Propionivibrio dicarboxylicus] | 09972 |
| rhodanese-like domain-containing protein [Zoogloea dura] | 02142 |
| hypothetical protein [Thauera linaloolentis] | 08741 |
| hypothetical protein [Aromatoleum toluclasticum] | 00853 |
| SIR2 family protein [Aromatoleum tolulyticum] | 01415 |
| hypothetical protein [Aromatoleum tolulyticum] | 08053 |
| DUF3486 family protein [Azoarcus communis] | 00071 |
| EthD domain-containing protein [Thauera linaloolentis] | 09420 |
| cadherin-like domain-containing protein [Aromatoleum tolulyticum] | 00432 |
| hypothetical protein [Azoarcus indigens] | 06141 |
| fatty acid desaturase [Azoarcus indigens] | 07924 |
| DUF3179 domain-containing protein [Aromatoleum toluclasticum] | 08451 |
| helix-turn-helix domain-containing protein [Propionivibrio dicarboxylicus] | 05327 |
| HNH endonuclease [Thauera linaloolentis] | 08121 |
| ATP-binding protein [Aromatoleum toluclasticum] | 06724 |
| helix-turn-helix transcriptional regulator [Aromatoleum tolulyticum] | 06558 |
| type II secretion system F family protein, partial [Thauera phenylacetica] | 07188 |
| MULTISPECIES: DUF4124 domain-containing protein [Aromatoleum] | 01384 |
| Ldh family oxidoreductase [Propionivibrio dicarboxylicus] | 04219 |
| DUF4160 domain-containing protein [Aromatoleum aromaticum] | 05478 |
| hypothetical protein [Aromatoleum tolulyticum] | 08054 |
| DUF2384 domain-containing protein [Thauera phenylacetica] | 10406 |
| DNA-binding protein [Aromatoleum toluclasticum] | 06533 |
| hypothetical protein [Propionivibrio dicarboxylicus] | 10219 |
| hypothetical protein [Aromatoleum tolulyticum] | 07712 |
| terpene cyclase/mutase family protein [Aromatoleum tolulyticum] | 04801 |
| hypothetical protein [Azoarcus indigens] | 01627 |
| regulatory protein GemA [Azoarcus communis] | 03979 |
| type I polyketide synthase [Azoarcus indigens] | 09909 |
| hypothetical protein [Thauera linaloolentis] | 06842 |
| replication protein C [Thauera phenolivorans] | 08183 |
| menaquinone-dependent protoporphyrinogen IX dehydrogenase [Propionivibrio dicarboxylicus] | 04629 |
| ATP-binding protein [Aromatoleum toluclasticum] | 05406 |
| helix-turn-helix transcriptional regulator [Thauera aminoaromatica] | 04479 |
| class I SAM-dependent methyltransferase [Thauera phenolivorans] | 01597 |
| DUF2523 domain-containing protein [Azoarcus indigens] | 10365 |
| MULTISPECIES: hypothetical protein [Proteobacteria] | 09426 |
| hypothetical protein [Azoarcus indigens] | 02934 |
| hypothetical protein [Azoarcus communis] | 03456 |
| hypothetical protein [Thauera linaloolentis] | 02884 |
| prolyl oligopeptidase family serine peptidase [Thauera linaloolentis] | 10391 |
| TlpA family protein disulfide reductase [Aromatoleum diolicum] | 02384 |
| aromatic ring-hydroxylating dioxygenase subunit alpha [Thauera linaloolentis] | 06126 |
| ThuA domain-containing protein [Thauera linaloolentis] | 06646 |
| hypothetical protein [Azoarcus indigens] | 06379 |
| hypothetical protein [Azoarcus indigens] | 02089 |
| RusA family crossover junction endodeoxyribonuclease [Azoarcus indigens] | 00559 |
| MULTISPECIES: hypothetical protein [Proteobacteria] | 08817 |
| hypothetical protein [Aromatoleum toluclasticum] | 05980 |
| HPP family protein [Propionivibrio dicarboxylicus] | 04117 |
| hypothetical protein [Thauera linaloolentis] | 03290 |
| hypothetical protein [Thauera aromatica] | 07442 |
| AbrB/MazE/SpoVT family DNA-binding domain-containing protein [Thauera linaloolentis] | 02095 |
| peptidoglycan DD-metalloendopeptidase family protein, partial [Thauera phenylacetica] | 01598 |
| hypothetical protein [Azoarcus indigens] | 06317 |

| | |
|---|---|
| NERD domain-containing protein [Aromatoleum tolulyticum] | 08251 |
| hypothetical protein [Azoarcus indigens] | 03581 |
| MULTISPECIES: glycosyltransferase [Rhodocyclales] | 04005 |
| hypothetical protein [Propionivibrio dicarboxylicus] | 10038 |
| hypothetical protein [Aromatoleum diolicum] | 08297 |
| HK97 gp10 family phage protein [Azoarcus communis] | 09543 |
| Ig-like domain-containing protein [Aromatoleum tolulyticum] | 02728 |
| type II toxin-antitoxin system Phd/YefM family antitoxin [Aromatoleum aromaticum] | 00112 |
| 3-hydroxyacyl-CoA dehydrogenase NAD-binding domain-containing protein [Aromatoleum tolulyticum] | 08591 |
| hypothetical protein [Azoarcus indigens] | 00900 |
| tape measure protein [Aromatoleum tolulyticum] | 02744 |
| hypothetical protein [Azoarcus sp. DD4] | 09755 |
| ABC transporter substrate-binding protein [Aromatoleum aromaticum] | 10301 |
| type II secretion system protein GspM [Propionivibrio dicarboxylicus] | 09470 |
| DUF1778 domain-containing protein [Zoogloea dura] | 03916 |
| endonuclease/exonuclease/phosphatase family protein [Azoarcus sp. L1K30] | 04622 |
| hypothetical protein [Thauera phenylacetica] | 01035 |
| DNA methyltransferase [Thauera linaloolentis] | 10316 |
| uridine kinase [Zoogloea dura] | 02747 |
| DUF2252 family protein [Zoogloea dura] | 04549 |
| HlyD family efflux transporter periplasmic adaptor subunit, partial [Thauera phenylacetica] | 08340 |
| LLM class flavin-dependent oxidoreductase [Aromatoleum tolulyticum] | 10627 |
| MULTISPECIES: hypothetical protein [Zoogloea] | 02331 |
| MULTISPECIES: nuclear transport factor 2 family protein [Rhodocyclales] | 05050 |
| MaoC family dehydratase N-terminal domain-containing protein [Aromatoleum tolulyticum] | 06721 |
| hypothetical protein [Propionivibrio dicarboxylicus] | 06226 |
| DUF2799 domain-containing protein [Propionivibrio dicarboxylicus] | 04514 |
| hypothetical protein [Aromatoleum tolulyticum] | 03321 |
| hypothetical protein [Thauera chlorobenzoica] | 01041 |
| MULTISPECIES: hypothetical protein [Zoogloea] | 05071 |
| lipid-binding SYLF domain-containing protein [Zoogloea dura] | 06346 |
| CRISPR-associated protein Csx16 [Thauera phenylacetica] | 04309 |
| SRPBCC family protein [Aromatoleum aromaticum] | 08870 |
| 3'-5' exoribonuclease [Thauera chlorobenzoica] | 08182 |
| (d)CMP kinase, partial [Thauera phenylacetica] | 05255 |
| hypothetical protein [Zoogloea dura] | 04434 |
| mucoidy inhibitor MuiA family protein [Zoogloea dura] | 09019 |
| CRISPR system precrRNA processing endoribonuclease RAMP protein Cas6 [Thauera phenylacetica] | 07609 |
| ribbon-helix-helix protein, CopG family [Thauera chlorobenzoica] | 03198 |
| ABC transporter substrate-binding protein [Aromatoleum tolulyticum] | 05337 |
| hypothetical protein [Aromatoleum tolulyticum] | 06547 |
| H-NS histone family protein [Zoogloea dura] | 00448 |
| DEAD/DEAH box helicase family protein [Aromatoleum aromaticum] | 03827 |
| class I SAM-dependent methyltransferase [Aromatoleum tolulyticum] | 03594 |
| hypothetical protein [Aromatoleum tolulyticum] | 08411 |
| hypothetical protein [Zoogloea dura] | 08504 |
| hypothetical protein [Thauera chlorobenzoica] | 02119 |
| hypothetical protein [Zoogloea dura] | 09185 |
| hypothetical protein [Zoogloea dura] | 01799 |
| hypothetical protein [Thauera chlorobenzoica] | 08154 |
| ATP-binding protein [Aromatoleum aromaticum] | 03616 |
| hypothetical protein [Zoogloea dura] | 00018 |
| hypothetical protein [Aromatoleum aromaticum] | 06337 |
| YihY/virulence factor BrkB family protein [Aromatoleum tolulyticum] | 07250 |
| hypothetical protein [Aromatoleum tolulyticum] | 02365 |
| cupin domain-containing protein [Aromatoleum aromaticum] | 07345 |
| hypothetical protein [Aromatoleum tolulyticum] | 09666 |
| hypothetical protein [Aromatoleum tolulyticum] | 04628 |
| aconitase X catalytic domain-containing protein [Aromatoleum tolulyticum] | 00012 |

| | |
|---|---|
| glycosyltransferase family 4 protein [Zoogloea dura] | 03258 |
| type II toxin-antitoxin system HicA family toxin [Thauera phenylacetica] | 06809 |
| hypothetical protein [Azoarcus sp. L1K30] | 07124 |
| hypothetical protein [Zoogloea dura] | 06363 |
| arylamine N-acetyltransferase [Azoarcus sp. L1K30] | 06628 |
| hypothetical protein [Thauera lacus] | 00881 |
| phage tail sheath subtilisin-like domain-containing protein [Zoogloea dura] | 02097 |
| hypothetical protein [Zoogloea dura] | 05360 |
| NUDIX hydrolase [Zoogloea dura] | 01700 |
| tyrosine-type recombinase/integrase [Aromatoleum tolulyticum] | 09567 |
| MULTISPECIES: hypothetical protein [Aromatoleum] | 00722 |
| hypothetical protein [Zoogloea dura] | 06345 |
| type II toxin-antitoxin system HipA family toxin [Aromatoleum aromaticum] | 08707 |
| EamA family transporter [Propionivibrio dicarboxylicus] | 08622 |
| HD domain-containing protein [Propionivibrio dicarboxylicus] | 00027 |
| hypothetical protein [Propionivibrio dicarboxylicus] | 01049 |
| TniQ family protein [Aromatoleum tolulyticum] | 06789 |
| glycosyltransferase family 25 protein [Zoogloea dura] | 07268 |
| hypothetical protein [Thauera phenylacetica] | 00969 |
| BON domain-containing protein [Aromatoleum aromaticum] | 02513 |
| FkbM family methyltransferase [Zoogloea dura] | 01864 |
| DUF3696 domain-containing protein [Propionivibrio dicarboxylicus] | 02680 |
| 4Fe-4S dicluster domain-containing protein, partial [Thauera phenylacetica] | 06492 |
| mechanosensitive ion channel [Propionivibrio dicarboxylicus] | 07006 |
| PEP-CTERM sorting domain-containing protein [Thauera phenylacetica] | 08302 |
| hypothetical protein [Thauera phenylacetica] | 01463 |
| carboxypeptidase regulatory-like domain-containing protein, partial [Thauera phenylacetica] | 05166 |
| DUF937 domain-containing protein [Aromatoleum aromaticum] | 03281 |
| nucleotidyltransferase domain-containing protein [Aromatoleum aromaticum] | 05705 |
| transcriptional regulator [Aromatoleum aromaticum] | 07563 |
| type II toxin-antitoxin system HicB family antitoxin [Aromatoleum aromaticum] | 02466 |
| GNAT family N-acetyltransferase [Azoarcus sp. L1K30] | 02955 |
| ATP-binding protein [Aromatoleum aromaticum] | 07557 |
| DUF6306 domain-containing protein [Zoogloea dura] | 08481 |
| 5-methylcytosine-specific restriction endonuclease system specificity protein McrC [Aromatoleum aromaticum] | 06860 |
| OsmC family protein [Zoogloea dura] | 09612 |
| PEP-CTERM sorting domain-containing protein [Zoogloea dura] | 08906 |
| tyrosine-type recombinase/integrase [Thauera phenylacetica] | 07391 |
| hypothetical protein [Aromatoleum tolulyticum] | 08346 |
| restriction endonuclease subunit S [Zoogloea dura] | 00868 |
| nickel transporter [Zoogloea dura] | 09268 |
| hypothetical protein [Thauera phenylacetica] | 08627 |
| hypothetical protein [Aromatoleum tolulyticum] | 06136 |
| MASE1 domain-containing protein [Aromatoleum tolulyticum] | 07619 |
| hypothetical protein [Thauera phenylacetica] | 03175 |
| CRISPR-associated ring nuclease Csm6 [Thauera phenylacetica] | 08133 |
| hypothetical protein [Aromatoleum aromaticum] | 00251 |
| RES family NAD+ phosphorylase [Aromatoleum aromaticum] | 01187 |
| prepilin-type N-terminal cleavage/methylation domain-containing protein [Propionivibrio dicarboxylicus] | 07870 |
| HEPN domain-containing protein [Thauera phenylacetica] | 02064 |
| DEAD/DEAH box helicase family protein [Aromatoleum aromaticum] | 04187 |
| iron transporter [Aromatoleum tolulyticum] | 07714 |
| cyclin-dependent kinase inhibitor 3 family protein [Zoogloea dura] | 10012 |
| MULTISPECIES: hypothetical protein [Aromatoleum] | 02730 |
| FIST C-terminal domain-containing protein [Propionivibrio dicarboxylicus] | 00357 |
| DUF6471 domain-containing protein [Aromatoleum tolulyticum] | 07806 |
| ThiF family adenylyltransferase [Propionivibrio dicarboxylicus] | 00263 |
| DUF6398 domain-containing protein [Thauera chlorobenzoica] | 08026 |
| MULTISPECIES: DUF2442 domain-containing protein [Aromatoleum] | 09444 |

| | |
|---|---|
| hypothetical protein [Aromatoleum aromaticum] | 10393 |
| molecular chaperone DnaJ [Aromatoleum aromaticum] | 03740 |
| hypothetical protein [Aromatoleum tolulyticum] | 02568 |
| hypothetical protein [Aromatoleum tolulyticum] | 04654 |
| protein-disulfide reductase DsbD N-terminal domain-containing protein, partial [Thauera phenylacetica] | 01170 |
| leucine-rich repeat-containing serine/threonine-protein kinase [Zoogloea dura] | 01708 |
| thiamine pyrophosphate-binding protein [Azoarcus nasutitermitis] | 06633 |
| hypothetical protein [Azoarcus communis] | 07452 |
| hypothetical protein [Aromatoleum tolulyticum] | 05009 |
| hypothetical protein [Zoogloea dura] | 02684 |
| DUF4157 domain-containing protein, partial [Thauera phenylacetica] | 01816 |
| hypothetical protein [Aromatoleum aromaticum] | 09421 |
| hypothetical protein [Aromatoleum tolulyticum] | 08608 |
| DUF126 domain-containing protein [Aromatoleum tolulyticum] | 07717 |
| GNAT family N-acetyltransferase [Zoogloea dura] | 01796 |
| hypothetical protein [Thauera phenylacetica] | 07444 |
| H4MPT-linked C1 transfer pathway protein [Aromatoleum toluvorans] | 08597 |
| type II secretion system minor pseudopilin GspH [Azoarcus indigens] | 07497 |
| DUF6079 family protein [Thauera chlorobenzoica] | 08772 |
| hypothetical protein [Azoarcus communis] | 09258 |
| ABC transporter permease [Aromatoleum toluclasticum] | 04263 |
| cobalt-precorrin-6A reductase [Thauera linaloolentis] | 01069 |
| sterol desaturase family protein [Azoarcus taiwanensis] | 07939 |
| DUF4351 domain-containing protein [Thauera linaloolentis] | 05988 |
| ABC transporter permease subunit [Azoarcus taiwanensis] | 09985 |
| TniQ family protein [Azoarcus communis] | 07171 |
| hypothetical protein [Aromatoleum aromaticum] | 07865 |
| nitroreductase family protein [Azoarcus indigens] | 10162 |
| hypothetical protein [Azoarcus communis] | 09832 |
| hypothetical protein [Azoarcus indigens] | 01916 |
| hypothetical protein [Azoarcus communis] | 07232 |
| hypothetical protein [Azoarcus communis] | 00467 |
| DUF302 domain-containing protein [Aromatoleum diolicum] | 10138 |
| M48 family metallopeptidase [Azoarcus indigens] | 07592 |
| hypothetical protein [Aromatoleum toluclasticum] | 07315 |
| CHASE2 domain-containing protein [Azoarcus indigens] | 02930 |
| ABC transporter substrate-binding protein [Azoarcus indigens] | 08795 |
| cytochrome c [Azoarcus indigens] | 07851 |
| DUF2442 domain-containing protein [Aromatoleum toluclasticum] | 06499 |
| hypothetical protein [Azoarcus indigens] | 08927 |
| rod-binding protein [Azoarcus indigens] | 10174 |
| hypothetical protein [Azoarcus indigens] | 00673 |
| amidoligase family protein [Azoarcus taiwanensis] | 02893 |
| Hpt domain-containing protein, partial [Aromatoleum diolicum] | 10408 |
| ribbon-helix-helix protein, CopG family [Azoarcus indigens] | 00068 |
| hypothetical protein [Thauera linaloolentis] | 00423 |
| hypothetical protein [Aromatoleum aromaticum] | 04659 |
| hypothetical protein [Thauera phenolivorans] | 00319 |
| biliverdin-producing heme oxygenase [Thauera linaloolentis] | 00567 |
| hypothetical protein [Thauera phenolivorans] | 05712 |
| HI0074 family nucleotidyltransferase substrate-binding subunit [Thauera aromatica] | 05795 |
| LuxR C-terminal-related transcriptional regulator [Aromatoleum toluclasticum] | 01325 |
| ethanolamine utilization acetate kinase EutQ [Thauera linaloolentis] | 06055 |
| alpha/beta hydrolase-fold protein [Thauera linaloolentis] | 05672 |
| sodium/glutamate symporter [Thauera aromatica] | 04957 |
| GTP-binding protein, partial [Thauera linaloolentis] | 10622 |
| hypothetical protein [Aromatoleum toluclasticum] | 09669 |
| hypothetical protein [Thauera linaloolentis] | 04759 |
| helix-turn-helix domain-containing protein [Aromatoleum toluclasticum] | 09786 |

| | |
|---|---|
| aldehyde dehydrogenase family protein, partial [Thauera linaloolentis] | 08122 |
| cobalamin biosynthesis protein [Thauera linaloolentis] | 03848 |
| metallophosphoesterase [Aromatoleum toluclasticum] | 00434 |
| MULTISPECIES: recombinase family protein [Proteobacteria] | 03695 |
| DUF3427 domain-containing protein [Aromatoleum diolicum] | 01055 |
| hypothetical protein [Aromatoleum aromaticum] | 06881 |
| hypothetical protein [Azoarcus communis] | 02113 |
| hypothetical protein [Azoarcus communis] | 04474 |
| hypothetical protein [Azoarcus communis] | 00880 |
| DUF1538 domain-containing protein [Azoarcus taiwanensis] | 09557 |
| hypothetical protein [Aromatoleum diolicum] | 02390 |
| hypothetical protein [Aromatoleum aromaticum] | 05872 |
| ParB/RepB/Spo0J family partition protein [Azoarcus indigens] | 06049 |
| DUF3489 domain-containing protein [Thauera terpenica] | 00759 |
| RNA ligase family protein [Thauera phenylacetica] | 07743 |
| peptidoglycan recognition protein family protein [Aromatoleum toluclasticum] | 02782 |
| hypothetical protein [Azoarcus indigens] | 07192 |
| helix-turn-helix domain-containing protein [Azoarcus indigens] | 08798 |
| hypothetical protein [Aromatoleum toluclasticum] | 08723 |
| hypothetical protein [Thauera terpenica] | 08126 |
| hypothetical protein [Thauera terpenica] | 06695 |
| phage major capsid protein [Azoarcus indigens] | 00364 |
| hypothetical protein [Azoarcus indigens] | 06318 |
| MULTISPECIES: TraK family protein [Aromatoleum] | 06700 |
| helix-turn-helix transcriptional regulator [Azoarcus indigens] | 01853 |
| 2,5-dihydroxypyridine 5,6-dioxygenase [Azoarcus indigens] | 01706 |
| hypothetical protein [Azoarcus indigens] | 00139 |
| MULTISPECIES: conjugal transfer protein TraL [Aromatoleum] | 02391 |
| MULTISPECIES: conjugal transfer protein TraM [Aromatoleum] | 06797 |
| hypothetical protein [Azoarcus indigens] | 04711 |
| MULTISPECIES: hypothetical protein [Thauera] | 05981 |
| hypothetical protein [Azoarcus indigens] | 00144 |
| restriction endonuclease [Aromatoleum toluclasticum] | 09083 |
| MULTISPECIES: hypothetical protein [Rhodocyclales] | 01169 |
| hypothetical protein [Azoarcus indigens] | 08800 |
| hypothetical protein [Thauera terpenica] | 00642 |
| hypothetical protein [Thauera terpenica] | 05723 |
| AAA family ATPase [Thauera terpenica] | 09060 |
| ligase-associated DNA damage response exonuclease [Azoarcus indigens] | 04296 |
| hypothetical protein [Aromatoleum diolicum] | 07861 |
| filamentous hemagglutinin N-terminal domain-containing protein [Azoarcus indigens] | 06162 |
| hypothetical protein [Aromatoleum aromaticum] | 01506 |
| NAD/NADP-dependent octopine/nopaline dehydrogenase family protein [Azoarcus indigens] | 05152 |
| helix-turn-helix domain-containing protein [Thauera chlorobenzoica] | 00949 |
| DUF4194 domain-containing protein [Azoarcus indigens] | 02665 |
| hypothetical protein [Aromatoleum diolicum] | 05819 |
| hypothetical protein [Aromatoleum toluclasticum] | 01184 |
| general secretion pathway protein GspC [Azoarcus indigens] | 01690 |
| substrate-binding domain-containing protein [Azoarcus indigens] | 00584 |
| glycosyltransferase, partial [Azoarcus taiwanensis] | 05187 |
| glycosyltransferase [Azoarcus indigens] | 01684 |
| lipase chaperone [Thauera terpenica] | 00986 |
| cupin domain-containing protein [Azoarcus indigens] | 00897 |
| phosphate ABC transporter permease PstA [Azoarcus taiwanensis] | 00127 |
| hypothetical protein [Thauera terpenica] | 05989 |
| hypothetical protein [Aromatoleum toluclasticum] | 07419 |
| hypothetical protein [Azoarcus communis] | 08342 |
| hypothetical protein [Azoarcus communis] | 07554 |
| DUF1800 domain-containing protein [Azoarcus indigens] | 08019 |

| | |
|---|---|
| ATP-dependent DNA ligase [Azoarcus indigens] | 03189 |
| alginate export family protein [Thauera linaloolentis] | 10323 |
| tyrosine-type recombinase/integrase [Aromatoleum aromaticum] | 00091 |
| EAL domain-containing protein [Thauera linaloolentis] | 03870 |
| hypothetical protein [Thauera aminoaromatica] | 05030 |
| hypothetical protein [Thauera linaloolentis] | 02904 |
| glutamate--cysteine ligase [Uliginosibacterium gangwonense] | 00311 |
| septal ring lytic transglycosylase RlpA family protein [Aromatoleum aromaticum] | 08101 |
| hypothetical protein [Aromatoleum aromaticum] | 05661 |
| phage portal protein [Uliginosibacterium gangwonense] | 01618 |
| glycoside hydrolase family 108 protein [Azoarcus rhizosphaerae] | 05145 |
| MULTISPECIES: hypothetical protein [Aromatoleum] | 06983 |
| DUF5368 domain-containing protein [Azoarcus rhizosphaerae] | 06028 |
| MarR family transcriptional regulator [Thauera linaloolentis] | 02860 |
| hypothetical protein [Uliginosibacterium gangwonense] | 06874 |
| type II toxin-antitoxin system prevent-host-death family antitoxin [Azoarcus rhizosphaerae] | 08202 |
| hypothetical protein [Azoarcus rhizosphaerae] | 09468 |
| class I SAM-dependent methyltransferase [Azoarcus rhizosphaerae] | 02648 |
| hypothetical protein [Azoarcus rhizosphaerae] | 10129 |
| AbrB/MazE/SpoVT family DNA-binding domain-containing protein [Azoarcus rhizosphaerae] | 09054 |
| hypothetical protein [Aromatoleum aromaticum] | 06471 |
| hypothetical protein [Thauera aminoaromatica] | 08086 |
| oligosaccharide flippase family protein [Thauera aminoaromatica] | 02587 |
| hypothetical protein [Aromatoleum toluclasticum] | 06488 |
| hypothetical protein [Aromatoleum aromaticum] | 00463 |
| hypothetical protein [Aromatoleum toluclasticum] | 00765 |
| AAA family ATPase [Aromatoleum aromaticum] | 09608 |
| hypothetical protein [Azoarcus rhizosphaerae] | 06879 |
| ATP-grasp domain-containing protein [Aromatoleum toluvorans] | 08600 |
| (5-formylfuran-3-yl)methyl phosphate synthase [Aromatoleum toluvorans] | 00510 |
| hypothetical protein [Thauera linaloolentis] | 01099 |
| formylmethanofuran dehydrogenase [Aromatoleum toluvorans] | 05679 |
| formylmethanofuran dehydrogenase subunit A [Aromatoleum toluvorans] | 05365 |
| hypothetical protein [Thauera linaloolentis] | 01349 |
| hypothetical protein [Thauera linaloolentis] | 03407 |
| DUF3325 domain-containing protein [Thauera linaloolentis] | 04792 |
| DUF447 family protein [Aromatoleum toluvorans] | 04640 |
| formylmethanofuran--tetrahydromethanopterin N-formyltransferase [Aromatoleum toluvorans] | 10131 |
| formylmethanofuran dehydrogenase subunit C [Aromatoleum toluvorans] | 04869 |
| DUF6513 domain-containing protein [Aromatoleum toluvorans] | 01860 |
| formaldehyde-activating enzyme [Aromatoleum toluvorans] | 03349 |
| dihydromethanopterin reductase [Aromatoleum toluvorans] | 02191 |
| triphosphoribosyl-dephospho-CoA synthase [Aromatoleum toluvorans] | 00009 |
| ethanolamine utilization cob(I)yrinic acid a,c-diamide adenosyltransferase EutT [Thauera linaloolentis] | 06689 |
| MULTISPECIES: ethanolamine utilization microcompartment protein EutM [Thauera] | 09112 |
| ethanolamine utilization microcompartment protein EutN [Thauera linaloolentis] | 07425 |
| ethanolamine utilization protein EutJ [Thauera linaloolentis] | 02299 |
| ethanolamine utilization protein EutH [Thauera linaloolentis] | 02653 |
| methenyltetrahydromethanopterin cyclohydrolase [Aromatoleum toluvorans] | 09244 |
| methylenetetrahydromethanopterin dehydrogenase [Aromatoleum toluvorans] | 00983 |
| GspH/FimT family protein [Aromatoleum aromaticum] | 00465 |
| hypothetical protein [Aromatoleum toluclasticum] | 10483 |
| TetR/AcrR family transcriptional regulator [Thauera linaloolentis] | 01697 |
| DUF4926 domain-containing protein [Aromatoleum aromaticum] | 01594 |
| PD-(D/E)XK nuclease family protein [Thauera aromatica] | 01673 |
| MFS transporter [Azoarcus communis] | 01970 |
| hypothetical protein [Azoarcus communis] | 00932 |
| hypothetical protein [Aromatoleum aromaticum] | 02892 |
| precorrin-3B synthase [Thauera linaloolentis] | 03461 |

| | |
|---|---|
| phage tail protein [Aromatoleum aromaticum] | 07564 |
| BREX-1 system phosphatase PglZ type B [Thauera phenolivorans] | 00859 |
| hypothetical protein [Rhodocyclus purpureus] | 06944 |
| BMC domain-containing protein [Thauera linaloolentis] | 09853 |
| ethanolamine utilization microcompartment protein EutL [Thauera linaloolentis] | 03882 |
| ethanolamine utilization microcompartment protein EutS [Thauera linaloolentis] | 08511 |
| ethanolamine utilization protein EutP [Thauera linaloolentis] | 02936 |
| hypothetical protein [Azoarcus communis] | 01709 |
| Fic family protein [Azoarcus communis] | 05238 |
| bifunctional GNAT family N-acetyltransferase/nucleoside diphosphate kinase regulator [Rhodocyclus purpureus] | 09058 |
| hypothetical protein [Rhodocyclus purpureus] | 01932 |
| SDR family oxidoreductase [Rhodocyclus purpureus] | 02786 |
| FAD-binding oxidoreductase [Rhodocyclus purpureus] | 08103 |
| flippase-like domain-containing protein [Rhodocyclus purpureus] | 06811 |
| MFS transporter [Thauera linaloolentis] | 04280 |
| DegT/DnrJ/EryC1/StrS family aminotransferase [Rhodocyclus purpureus] | 04790 |
| hypothetical protein [Rhodocyclus purpureus] | 08960 |
| AraC family transcriptional regulator [Aromatoleum aromaticum] | 00257 |
| phage integrase N-terminal SAM-like domain-containing protein [Thauera aromatica] | 00107 |
| hypothetical protein [Azoarcus rhizosphaerae] | 09086 |
| GGDEF domain-containing protein [Azoarcus rhizosphaerae] | 04403 |
| hypothetical protein [Azoarcus rhizosphaerae] | 02301 |
| preprotein translocase subunit YajC [Azoarcus rhizosphaerae] | 04399 |
| DUF1045 domain-containing protein [Azoarcus rhizosphaerae] | 06050 |
| TrfB-related DNA-binding protein [Aromatoleum aromaticum] | 06911 |
| phosphonate C-P lyase system protein PhnG [Azoarcus rhizosphaerae] | 05531 |
| phosphonate C-P lyase system protein PhnH [Azoarcus rhizosphaerae] | 09485 |
| carbon-phosphorus lyase complex subunit PhnI [Azoarcus rhizosphaerae] | 06394 |
| alpha-D-ribose 1-methylphosphonate 5-phosphate C-P-lyase PhnJ [Azoarcus rhizosphaerae] | 04398 |
| alpha-D-ribose 1-methylphosphonate 5-triphosphate diphosphatase [Azoarcus rhizosphaerae] | 03470 |
| SHOCT domain-containing protein [Azoarcus sp. Aa7] | 08750 |
| hypothetical protein [Aromatoleum aromaticum] | 01992 |
| DNA cytosine methyltransferase [Azoarcus rhizosphaerae] | 06361 |
| DUF2795 domain-containing protein [Aromatoleum aromaticum] | 06269 |
| hypothetical protein [Thauera linaloolentis] | 04191 |
| hypothetical protein [Azoarcus sp. Aa7] | 08520 |
| BrnT family toxin [Azoarcus sp. Aa7] | 05586 |
| hypothetical protein [Azoarcus sp. Aa7] | 01871 |
| DUF4276 family protein [Azoarcus sp. Aa7] | 07933 |
| hypothetical protein [Azoarcus sp. Aa7] | 02386 |
| hypothetical protein [Azoarcus communis] | 09980 |
| DUF4236 domain-containing protein [Thauera linaloolentis] | 03669 |
| hypothetical protein [Aromatoleum aromaticum] | 08474 |
| hypothetical protein [Aromatoleum aromaticum] | 00952 |
| DUF1353 domain-containing protein [Aromatoleum toluclasticum] | 03487 |
| hypothetical protein [Thauera phenylacetica] | 05669 |
| RDD family protein [Uliginosibacterium aquaticum] | 02758 |
| molecular chaperone [Azoarcus communis] | 06356 |
| TonB family protein [Azoarcus indigens] | 09128 |
| transposase [Azoarcus indigens] | 00563 |
| ATP-binding protein [Thauera phenylacetica] | 09157 |
| FTR1 family iron permease [Candidatus Dactylopiibacterium carminicum] | 01090 |
| hypothetical protein [Azoarcus communis] | 01305 |
| DUF4402 domain-containing protein [Azoarcus communis] | 02619 |
| DDE-type integrase/transposase/recombinase [Aromatoleum pumilum] | 01872 |
| hypothetical protein [Azoarcus indigens] | 06433 |
| mitochondrial fission ELM1 family protein [Thauera phenylacetica] | 00477 |
| hypothetical protein [Azoarcus communis] | 06764 |
| hypothetical protein [Uliginosibacterium aquaticum] | 08648 |

| | |
|---|---|
| YjhX family toxin [Azoarcus communis] | 07819 |
| hypothetical protein [Azoarcus indigens] | 01628 |
| glycosyltransferase [Azoarcus indigens] | 00054 |
| tetratricopeptide repeat protein [Azoarcus indigens] | 00417 |
| major capsid protein [Aromatoleum aromaticum] | 05077 |
| CDP-glycerol glycerophosphotransferase family protein [Aromatoleum pumilum] | 06908 |
| PilZ domain-containing protein [Uliginosibacterium aquaticum] | 03162 |
| IS1595 family transposase [Candidatus Dactylopiibacterium carminicum] | 07987 |
| EAL domain-containing protein, partial [Aromatoleum toluclasticum] | 03126 |
| hypothetical protein [Azoarcus communis] | 08178 |
| ATP-grasp domain-containing protein [Aromatoleum toluvorans] | 03310 |
| hypothetical protein [Azoarcus indigens] | 05827 |
| competence protein ComEC [Thauera chlorobenzoica] | 03526 |
| tetratricopeptide repeat protein [Thauera phenylacetica] | 02043 |
| patatin-like phospholipase family protein [Uliginosibacterium aquaticum] | 03163 |
| DUF1116 domain-containing protein [Azoarcus indigens] | 01013 |
| hypothetical protein [Azoarcus indigens] | 02093 |
| glycine zipper 2TM domain-containing protein [Uliginosibacterium aquaticum] | 04910 |
| glycosyltransferase [Uliginosibacterium aquaticum] | 01889 |
| Bro-N domain-containing protein [Candidatus Dactylopiibacterium carminicum] | 07467 |
| hypothetical protein [Azoarcus indigens] | 00533 |
| long-chain fatty acid--CoA ligase, partial [Aromatoleum toluclasticum] | 04512 |
| hypothetical protein [Aromatoleum aromaticum] | 08023 |
| DUF2218 domain-containing protein [Uliginosibacterium aquaticum] | 03037 |
| MMPL family transporter, partial [Aromatoleum toluclasticum] | 03271 |
| Mov34/MPN/PAD-1 family protein [Thauera chlorobenzoica] | 09563 |
| DUF2157 domain-containing protein [Thauera chlorobenzoica] | 03359 |
| hypothetical protein [Azoarcus communis] | 00636 |
| YiiX family permuted papain-like enzyme [Uliginosibacterium aquaticum] | 03703 |
| sodium-translocating pyrophosphatase [Propionivibrio dicarboxylicus] | 03866 |
| transposase, partial [Azoarcus indigens] | 02704 |
| MULTISPECIES: hypothetical protein [Thauera] | 10291 |
| heme-binding protein [Uliginosibacterium aquaticum] | 09526 |
| HlyD family secretion protein [Azoarcus indigens] | 09553 |
| MULTISPECIES: hypothetical protein [Thauera] | 02030 |
| PIN domain-containing protein [Thauera chlorobenzoica] | 01038 |
| HNH endonuclease [Aromatoleum aromaticum] | 05217 |
| hypothetical protein [Uliginosibacterium aquaticum] | 07963 |
| exonuclease domain-containing protein [Azoarcus indigens] | 06904 |
| glycosyltransferase [Uliginosibacterium aquaticum] | 02169 |
| complement resistance protein TraT [Aromatoleum aromaticum] | 04143 |
| HPr-rel-A system PqqD family peptide chaperone [Azoarcus communis] | 03190 |
| hypothetical protein [Thauera chlorobenzoica] | 03673 |
| MarR family winged helix-turn-helix transcriptional regulator [Thauera chlorobenzoica] | 00877 |
| hypothetical protein [Azoarcus indigens] | 00580 |
| hypothetical protein [Aromatoleum aromaticum] | 08515 |
| prepilin-type N-terminal cleavage/methylation domain-containing protein [Azoarcus indigens] | 01704 |
| anaerobic nitric oxide reductase flavorubredoxin [Aromatoleum aromaticum] | 02168 |
| methyltransferase [Uliginosibacterium aquaticum] | 00436 |
| PH domain-containing protein [Aromatoleum aromaticum] | 04742 |
| hypothetical protein [Thauera chlorobenzoica] | 02435 |
| MBL fold metallo-hydrolase [Uliginosibacterium aquaticum] | 09030 |
| response regulator [Uliginosibacterium aquaticum] | 00033 |
| alpha-ketoacid dehydrogenase subunit alpha/beta [Thauera chlorobenzoica] | 03989 |
| hypothetical protein [Azoarcus communis] | 09573 |
| cysteine hydrolase [Uliginosibacterium aquaticum] | 08337 |
| DUF4350 domain-containing protein [Uliginosibacterium aquaticum] | 07981 |
| DUF4129 domain-containing protein [Uliginosibacterium aquaticum] | 03317 |
| oxidoreductase [Thauera chlorobenzoica] | 01102 |

| | |
|---|---|
| stage II sporulation protein M [Uliginosibacterium aquaticum] | 04366 |
| helix-turn-helix domain-containing protein [Azoarcus indigens] | 04729 |
| tape measure protein [Thauera butanivorans] | 03180 |
| SDR family oxidoreductase, partial [Aromatoleum toluclasticum] | 03465 |
| DUF4150 domain-containing protein [Thauera butanivorans] | 00307 |
| Na+/H+ antiporter NhaC [Aromatoleum toluclasticum] | 00209 |
| MULTISPECIES: hypothetical protein [Rhodocyclales] | 08578 |
| MULTISPECIES: HAD family phosphatase [Uliginosibacterium] | 05577 |
| hypothetical protein [Thauera phenylacetica] | 00922 |
| hypothetical protein [Azoarcus indigens] | 06917 |
| hypothetical protein [Thauera phenylacetica] | 03685 |
| MULTISPECIES: hypothetical protein [Rhodocyclales] | 02570 |
| alpha/beta fold hydrolase [Aromatoleum toluclasticum] | 02944 |
| PepSY domain-containing protein [Azoarcus indigens] | 07484 |
| hypothetical protein [Azoarcus communis] | 08818 |
| ferritin-like domain-containing protein [Azoarcus indigens] | 02639 |
| DUF2848 domain-containing protein [Azoarcus indigens] | 03723 |
| DUF202 domain-containing protein [Azoarcus indigens] | 00561 |
| MULTISPECIES: K(+)-transporting ATPase subunit F [Azoarcus] | 07841 |
| cyclic peptide export ABC transporter [Azoarcus indigens] | 09007 |
| siderophore-iron reductase FhuF [Azoarcus indigens] | 10385 |
| cytochrome c3 family protein, partial [Propionivibrio dicarboxylicus] | 01467 |
| hypothetical protein [Azoarcus communis] | 06114 |
| methyltransferase domain-containing protein [Azoarcus indigens] | 07259 |
| phage tail tape measure protein [Thauera chlorobenzoica] | 04744 |
| hypothetical protein [Aromatoleum petrolei] | 04466 |
| hypothetical protein [Azoarcus communis] | 07914 |
| hypothetical protein [Propionivibrio dicarboxylicus] | 07044 |
| tetratricopeptide repeat protein [Azoarcus indigens] | 00129 |
| peptidoglycan-binding protein [Azoarcus indigens] | 08140 |
| type II toxin-antitoxin system HicB family antitoxin [Thauera chlorobenzoica] | 04089 |
| hypothetical protein [Azoarcus communis] | 01766 |
| hypothetical protein [Aromatoleum petrolei] | 01839 |
| hypothetical protein [Thauera phenylacetica] | 09892 |
| MarR family transcriptional regulator [Thauera phenylacetica] | 04671 |
| PadR family transcriptional regulator [Thauera phenylacetica] | 01433 |
| TniQ family protein [Aromatoleum pumilum] | 06941 |
| hypothetical protein [Aromatoleum aromaticum] | 02819 |
| type II toxin-antitoxin system VapC family toxin [Azoarcus indigens] | 05469 |
| DUF3300 domain-containing protein [Azoarcus indigens] | 01875 |
| type II toxin-antitoxin system MqsR family toxin [Aromatoleum toluclasticum] | 04679 |
| hypothetical protein [Thauera chlorobenzoica] | 02896 |
| energy transducer TonB [Azoarcus indigens] | 04326 |
| DUF4433 domain-containing protein [Thauera phenylacetica] | 04308 |
| N(5)-hydroxyornithine transformylase PvdF [Azoarcus indigens] | 03144 |
| hypothetical protein [Aromatoleum toluclasticum] | 01777 |
| hypothetical protein [Aromatoleum toluclasticum] | 05613 |
| hypothetical protein [Uliginosibacterium aquaticum] | 05221 |
| ABC transporter permease [Azoarcus indigens] | 00367 |
| TnsD family transposase [Aromatoleum toluclasticum] | 09413 |
| hypothetical protein [Uliginosibacterium aquaticum] | 10088 |
| type I-E CRISPR-associated endoribonuclease Cas2e [Aromatoleum toluclasticum] | 01962 |
| type I-E CRISPR-associated protein Cse2/CasB [Aromatoleum toluclasticum] | 00498 |
| CRISPR-associated protein Cse1 [Aromatoleum toluclasticum] | 00024 |
| hypothetical protein [Candidatus Dactylopiibacterium carminicum] | 02900 |
| hypothetical protein [Uliginosibacterium aquaticum] | 01805 |
| DVUA0089 family protein [Azoarcus indigens] | 04879 |
| hypothetical protein [Thauera phenylacetica] | 03001 |
| hypothetical protein [Thauera chlorobenzoica] | 05869 |

| | |
|---|---|
| hypothetical protein [Candidatus Dactylopiibacterium carminicum] | 04401 |
| metallohydrolase [Thauera chlorobenzoica] | 01266 |
| hypothetical protein [Uliginosibacterium aquaticum] | 03970 |
| Rieske 2Fe-2S domain-containing protein [Aromatoleum toluclasticum] | 05052 |
| MbtH family NRPS accessory protein [Azoarcus indigens] | 02202 |
| hypothetical protein [Azoarcus indigens] | 09107 |
| hypothetical protein [Aromatoleum toluclasticum] | 06398 |
| MAPEG family protein [Azoarcus indigens] | 00679 |
| class I SAM-dependent methyltransferase [Azoarcus indigens] | 02869 |
| hypothetical protein [Azoarcus indigens] | 08803 |
| hypothetical protein [Azoarcus indigens] | 07227 |
| OFA family MFS transporter [Azoarcus indigens] | 07958 |
| macro domain-containing protein [Thauera phenylacetica] | 00961 |
| site-specific integrase [Uliginosibacterium aquaticum] | 07276 |
| hypothetical protein, partial [Azoarcus sp. TTM-91] | 03341 |
| DUF2786 domain-containing protein [Azoarcus communis] | 05524 |
| hypothetical protein [Aromatoleum toluclasticum] | 09787 |
| diguanylate cyclase [Thauera chlorobenzoica] | 09879 |
| type III-B CRISPR-associated protein Cas10/Cmr2 [Thauera phenylacetica] | 04714 |
| DUF5676 family membrane protein [Aromatoleum toluclasticum] | 04602 |
| IS1634 family transposase, partial [Thauera phenylacetica] | 08097 |
| CHAD domain-containing protein, partial [Thauera phenylacetica] | 10294 |
| hypothetical protein [Thauera phenylacetica] | 00966 |
| ISAs1 family transposase, partial [Thauera phenylacetica] | 07464 |
| TolC family protein, partial [Thauera phenylacetica] | 07598 |
| type III-B CRISPR module-associated protein Cmr5 [Thauera phenylacetica] | 03665 |
| hypothetical protein [Aromatoleum toluclasticum] | 00501 |
| hypothetical protein [Thauera phenylacetica] | 02274 |
| MULTISPECIES: hypothetical protein [Betaproteobacteria] | 02543 |
| glycoside hydrolase family 3 protein [Thauera phenylacetica] | 08714 |
| hypothetical protein [Thauera phenylacetica] | 00473 |
| type III-B CRISPR module RAMP protein Cmr6 [Thauera phenylacetica] | 08090 |
| hypothetical protein [Thauera phenylacetica] | 03634 |
| hypothetical protein [Thauera phenylacetica] | 02883 |
| hypothetical protein [Thauera phenylacetica] | 04934 |
| ISAs1 family transposase, partial [Thauera phenylacetica] | 07130 |
| hypothetical protein [Azoarcus communis] | 10180 |
| MULTISPECIES: CRISPR-associated endonuclease Cas1 [Thauera] | 06855 |
| hypothetical protein [Thauera phenylacetica] | 00997 |
| hypothetical protein [Thauera phenylacetica] | 03245 |
| hypothetical protein [Thauera phenylacetica] | 07387 |
| hypothetical protein [Thauera phenylacetica] | 06434 |
| hypothetical protein [Aromatoleum toluclasticum] | 02028 |
| MULTISPECIES: hypothetical protein [Proteobacteria] | 04662 |
| type III-B CRISPR module RAMP protein Cmr4 [Thauera phenylacetica] | 03339 |
| MFS transporter, partial [Thauera phenylacetica] | 10434 |
| hypothetical protein [Aromatoleum toluclasticum] | 03062 |
| hypothetical protein [Thauera phenylacetica] | 08427 |
| hypothetical protein [Azoarcus sp. DD4] | 02595 |
| CRISPR-associated Cmr3 family protein [Thauera phenylacetica] | 02046 |
| ATP-binding domain-containing protein, partial [Thauera phenylacetica] | 06284 |
| DUF1622 domain-containing protein [Azoarcus olearius] | 03748 |
| universal stress protein [Azoarcus sp. KH32C] | 01197 |
| hypothetical protein [Thauera phenylacetica] | 03145 |
| LuxR C-terminal-related transcriptional regulator [Aromatoleum toluclasticum] | 09033 |
| molybdenum cofactor biosynthesis F family protein [Thauera phenylacetica] | 08430 |
| hypothetical protein [Aromatoleum toluclasticum] | 07415 |
| ISAs1 family transposase, partial [Thauera phenylacetica] | 09819 |
| hypothetical protein [Thauera phenylacetica] | 04796 |

| | |
|---|---|
| hypothetical protein [Azoarcus sp. DD4] | 04411 |
| hypothetical protein [Thauera phenylacetica] | 04630 |
| hypothetical protein [Azoarcus sp. DD4] | 08081 |
| hypothetical protein [Azoarcus sp. DD4] | 02953 |
| TniQ family protein [Aromatoleum toluclasticum] | 04994 |
| type II toxin-antitoxin system VapC family toxin [Thauera phenylacetica] | 00968 |
| OmpA family protein [Azoarcus sp. DD4] | 09385 |
| sarcosine oxidase subunit gamma family protein, partial [Aromatoleum toluclasticum] | 10186 |
| type II toxin-antitoxin system prevent-host-death family antitoxin [Thauera phenylacetica] | 01347 |
| hypothetical protein [Aromatoleum toluclasticum] | 00664 |
| ABC transporter permease, partial [Thauera phenylacetica] | 07513 |
| TolC family protein, partial [Thauera phenylacetica] | 01244 |
| glycosyltransferase [Aromatoleum toluclasticum] | 07634 |
| type I-U CRISPR-associated RAMP protein Csb1/Cas7u [Thauera phenylacetica] | 08939 |
| phosphatase PAP2 family protein [Aromatoleum toluclasticum] | 08052 |
| hypothetical protein [Azoarcus sp. DD4] | 00927 |
| EAL domain-containing protein, partial [Aromatoleum toluclasticum] | 03270 |
| hypothetical protein [Aromatoleum toluclasticum] | 00497 |
| type I-U CRISPR-associated helicase/endonuclease Cas3 [Thauera phenylacetica] | 00963 |
| hypothetical protein, partial [Aromatoleum toluclasticum] | 01850 |
| hypothetical protein [Aromatoleum toluclasticum] | 03598 |
| hypothetical protein [Aromatoleum toluclasticum] | 03408 |
| hypothetical protein [Aromatoleum toluclasticum] | 08110 |
| hypothetical protein [Aromatoleum toluclasticum] | 03483 |
| META domain-containing protein [Thauera phenylacetica] | 00819 |
| GtrA family protein [Aromatoleum toluclasticum] | 06668 |
| histidine phosphatase family protein [Aromatoleum toluclasticum] | 05521 |
| hypothetical protein [Aromatoleum toluclasticum] | 01888 |
| curli production assembly/transport protein CsgG [Aromatoleum toluclasticum] | 00837 |
| MULTISPECIES: metalloregulator ArsR/SmtB family transcription factor [Thauera] | 03653 |
| hypothetical protein [Aromatoleum toluclasticum] | 10187 |
| hypothetical protein [Aromatoleum toluclasticum] | 07360 |
| DUF4115 domain-containing protein, partial [Thauera phenylacetica] | 07351 |
| class I SAM-dependent methyltransferase [Azoarcus sp. DD4] | 10287 |
| hypothetical protein [Thauera phenylacetica] | 10293 |
| tRNA lysidine(34) synthetase TilS, partial [Thauera phenylacetica] | 07580 |
| porin [Thauera phenylacetica] | 09391 |
| ATP-binding cassette domain-containing protein, partial [Thauera phenylacetica] | 07746 |
| hypothetical protein [Aromatoleum toluclasticum] | 08726 |
| hypothetical protein [Aromatoleum toluclasticum] | 06607 |
| radical SAM protein [Aromatoleum toluclasticum] | 05573 |
| TIR domain-containing protein [Aromatoleum toluclasticum] | 10105 |
| hypothetical protein [Aromatoleum toluclasticum] | 03121 |
| hypothetical protein [Azoarcus sp. DD4] | 08822 |
| hypothetical protein [Thauera phenylacetica] | 03666 |
| hypothetical protein [Thauera phenylacetica] | 06279 |
| hypothetical protein [Azoarcus sp. KH32C] | 04682 |
| Eco57I restriction-modification methylase domain-containing protein [Thauera phenylacetica] | 05767 |
| SPOR domain-containing protein, partial [Thauera phenylacetica] | 06713 |
| flagellar hook-length control protein FliK, partial [Thauera phenylacetica] | 03977 |
| DUF3565 domain-containing protein [Azoarcus sp. DD4] | 09492 |
| DUF1302 family protein, partial [Aromatoleum toluclasticum] | 08891 |
| hypothetical protein [Thauera phenylacetica] | 02538 |
| hypothetical protein [Aromatoleum toluclasticum] | 10341 |
| DUF2285 domain-containing protein [Aromatoleum toluclasticum] | 01596 |
| hypothetical protein, partial [Thauera phenylacetica] | 06807 |
| hypothetical protein [Aromatoleum toluclasticum] | 00233 |
| hypothetical protein [Thauera phenylacetica] | 07438 |
| translocation/assembly module TamB domain-containing protein, partial [Thauera phenylacetica] | 09725 |

| | |
|---|---|
| nucleotidyltransferase [Thauera phenylacetica] | 09766 |
| tripartite tricarboxylate transporter TctB family protein [Thauera phenylacetica] | 05956 |
| MULTISPECIES: hypothetical protein [Proteobacteria] | 04784 |
| MULTISPECIES: DUF6430 domain-containing protein [Proteobacteria] | 05139 |
| hypothetical protein [Aromatoleum toluclasticum] | 01326 |
| hypothetical protein [Aromatoleum toluclasticum] | 10608 |
| recombinase family protein [Aromatoleum toluclasticum] | 04923 |
| glutamate--tRNA ligase family protein, partial [Aromatoleum toluclasticum] | 05637 |
| MULTISPECIES: GNAT family N-acetyltransferase [Bacteria] | 06170 |
| MULTISPECIES: phage capsid protein, partial [Bacteria] | 05692 |
| MULTISPECIES: hypothetical protein [Bacteria][Archaea] | 01632 |
| MULTISPECIES: TniQ family protein [Proteobacteria] | 03009 |
| hypothetical protein [Thauera phenylacetica] | 08087 |
| peptidyl-prolyl cis-trans isomerase [Aromatoleum toluclasticum] | 04582 |
| hypothetical protein [Thauera phenylacetica] | 02536 |
| hypothetical protein [Aromatoleum toluclasticum] | 09785 |
| radical SAM protein [Thauera phenylacetica] | 01240 |
| hypothetical protein [Thauera phenylacetica] | 05086 |
| hypothetical protein [Thauera phenylacetica] | 01429 |
| PAAR domain-containing protein [Aromatoleum toluclasticum] | 07763 |
| hypothetical protein [Uliginosibacterium aquaticum] | 08443 |
| hypothetical protein [Zoogloea dura] | 08242 |
| ABC transporter permease subunit [Aromatoleum toluclasticum] | 10141 |
| GIY-YIG nuclease family protein [Aromatoleum toluclasticum] | 09846 |
| DISARM system phospholipase D-like protein DrmC [Aromatoleum toluclasticum] | 00542 |
| LysR substrate-binding domain-containing protein [Aromatoleum toluclasticum] | 08573 |
| malonate transporter subunit MadL [Aromatoleum petrolei] | 10401 |
| caspase family protein [Aromatoleum toluclasticum] | 08933 |
| malonate transporter subunit MadM [Aromatoleum petrolei] | 01104 |
| universal stress protein [Aromatoleum toluclasticum] | 09216 |
| MULTISPECIES: SPW repeat protein [Rhodocyclales] | 06128 |
| hypothetical protein [Azoarcus toluclasticus] | 04797 |
| transposase, partial [Aromatoleum toluclasticum] | 04227 |
| hypothetical protein [Aromatoleum petrolei] | 01451 |
| AMP-binding protein, partial [Azoarcus indigens] | 07066 |
| hypothetical protein [Aromatoleum petrolei] | 04095 |
| hypothetical protein [Aromatoleum toluclasticum] | 08344 |
| hypothetical protein [Aromatoleum toluclasticum] | 02175 |
| hypothetical protein [Aromatoleum toluclasticum] | 08452 |
| MFS transporter [Aromatoleum toluclasticum] | 04330 |
| hypothetical protein [Aromatoleum toluclasticum] | 09032 |
| hypothetical protein [Azoarcus toluclasticus] | 08810 |
| hypothetical protein [Aromatoleum toluclasticum] | 04349 |
| DUF3883 domain-containing protein [Aromatoleum toluclasticum] | 01661 |
| hypothetical protein [Aromatoleum petrolei] | 01960 |
| GIY-YIG nuclease family protein [Thauera terpenica] | 04849 |
| hypothetical protein [Thauera terpenica] | 04334 |
| MliC family protein [Thauera terpenica] | 02856 |
| hypothetical protein [Thauera terpenica] | 09876 |
| hypothetical protein [Thauera terpenica] | 05983 |
| alpha/beta-hydrolase family protein, partial [Azoarcus taiwanensis] | 03883 |
| GNAT family N-acetyltransferase [Azoarcus taiwanensis] | 07928 |
| AAA family ATPase, partial [Azoarcus taiwanensis] | 01448 |
| hypothetical protein [Azoarcus taiwanensis] | 00428 |
| DUF262 domain-containing protein [Azoarcus taiwanensis] | 06734 |
| DNA damage-inducible protein D [Thauera terpenica] | 01374 |
| hypothetical protein [Thauera terpenica] | 03800 |
| DUF2761 domain-containing protein [Thauera terpenica] | 05427 |
| class I SAM-dependent methyltransferase [Azoarcus taiwanensis] | 01969 |

| | |
|---|---|
| GGDEF domain-containing protein [Azoarcus taiwanensis] | 01776 |
| EAL domain-containing protein [Thauera terpenica] | 04847 |
| RHS repeat protein [Azoarcus taiwanensis] | 02917 |
| hypothetical protein [Thauera terpenica] | 00845 |
| hypothetical protein [Aromatoleum petrolei] | 04357 |
| DNA polymerase/3'-5' exonuclease PolX [Aromatoleum petrolei] | 03307 |
| biotin-independent malonate decarboxylase subunit gamma [Aromatoleum petrolei] | 04965 |
| class I SAM-dependent methyltransferase [Thauera terpenica] | 01959 |
| transporter substrate-binding domain-containing protein [Thauera terpenica] | 00844 |
| hypothetical protein [Thauera terpenica] | 06684 |
| hypothetical protein [Thauera terpenica] | 04964 |
| recombinase RecT [Azoarcus indigens] | 00439 |
| DUF2188 domain-containing protein [Aromatoleum toluclasticum] | 06800 |
| DUF4398 domain-containing protein [Zoogloea dura] | 00993 |
| hypothetical protein [Uliginosibacterium aquaticum] | 07966 |
| DUF2802 domain-containing protein [Uliginosibacterium aquaticum] | 07308 |
| hypothetical protein [Thauera butanivorans] | 09056 |
| DUF459 domain-containing protein [Uliginosibacterium aquaticum] | 09400 |
| putative motility protein [Uliginosibacterium aquaticum] | 05974 |
| hypothetical protein [Uliginosibacterium aquaticum] | 07969 |
| hypothetical protein [Uliginosibacterium aquaticum] | 00588 |
| FHA domain-containing protein [Uliginosibacterium aquaticum] | 07762 |
| rhamnogalacturonan lyase [Uliginosibacterium aquaticum] | 08335 |
| phosphoserine transaminase [Uliginosibacterium aquaticum] | 05969 |
| hypothetical protein [Thauera butanivorans] | 08138 |
| T6SS effector amidase Tae4 family protein [Uliginosibacterium aquaticum] | 05076 |
| alkane 1-monooxygenase [Aromatoleum toluclasticum] | 06210 |
| hypothetical protein [Uliginosibacterium aquaticum] | 10321 |
| ankyrin repeat domain-containing protein [Uliginosibacterium aquaticum] | 04921 |
| J domain-containing protein [Uliginosibacterium aquaticum] | 10023 |
| DUF481 domain-containing protein [Uliginosibacterium aquaticum] | 04914 |
| type VI secretion system-associated protein TagF [Thauera butanivorans] | 04375 |
| restriction endonuclease subunit S [Zoogloea ramigera] | 02958 |
| glycoside hydrolase family 28 protein [Uliginosibacterium aquaticum] | 07406 |
| type 4a pilus biogenesis protein PilO [Uliginosibacterium aquaticum] | 07027 |
| hypothetical protein [Uliginosibacterium aquaticum] | 04555 |
| CAP domain-containing protein [Uliginosibacterium aquaticum] | 05249 |
| DUF2523 domain-containing protein [Candidatus Dactylopiibacterium carminicum] | 02304 |
| major capsid protein [Candidatus Dactylopiibacterium carminicum] | 08926 |
| DJ-1/PfpI family protein [Candidatus Dactylopiibacterium carminicum] | 02183 |
| DUF2007 domain-containing protein [Candidatus Dactylopiibacterium carminicum] | 05620 |
| spore coat protein U domain-containing protein [Azoarcus indigens] | 04882 |
| MULTISPECIES: IS91 family transposase [Bacteria] | 03333 |
| YdaS family helix-turn-helix protein [Thauera butanivorans] | 05995 |
| DUF1998 domain-containing protein [Aromatoleum toluclasticum] | 07065 |
| DISARM system helicase DrmA [Aromatoleum toluclasticum] | 09611 |
| hypothetical protein [Aromatoleum toluclasticum] | 09561 |
| ABC transporter permease subunit [Candidatus Dactylopiibacterium carminicum] | 00155 |
| NAD(P)/FAD-dependent oxidoreductase [Thauera butanivorans] | 00300 |
| MipA/OmpV family protein [Candidatus Dactylopiibacterium carminicum] | 04318 |
| hypothetical protein [Candidatus Dactylopiibacterium carminicum] | 10371 |
| hypothetical protein [Candidatus Dactylopiibacterium carminicum] | 01924 |
| phage baseplate assembly protein V [Thauera butanivorans] | 00079 |
| DUF5675 family protein [Thauera butanivorans] | 10322 |
| DUF4150 domain-containing protein [Thauera butanivorans] | 06691 |
| LysR substrate-binding domain-containing protein [Thauera butanivorans] | 08531 |
| hypothetical protein [Thauera butanivorans] | 10587 |
| hypothetical protein [Thauera butanivorans] | 02283 |
| hemin-degrading factor [Candidatus Dactylopiibacterium carminicum] | 00714 |

| | |
|---|---|
| BrnA antitoxin family protein [Thauera butanivorans] | 01345 |
| hypothetical protein [Candidatus Dactylopiibacterium carminicum] | 07140 |
| LacI family transcriptional regulator [Candidatus Dactylopiibacterium carminicum] | 02303 |
| type IV secretion system DNA-binding domain-containing protein, partial [Azoarcus taiwanensis] | 07238 |
| hypothetical protein [Azoarcus taiwanensis] | 03370 |
| DUF72 domain-containing protein [Aromatoleum diolicum] | 06495 |
| DUF4258 domain-containing protein [Azoarcus rhizosphaerae] | 08814 |
| hypothetical protein [Azoarcus rhizosphaerae] | 07887 |
| DUF3596 domain-containing protein [Thauera aminoaromatica] | 02584 |
| histidine kinase [Azoarcus rhizosphaerae] | 06878 |
| MULTISPECIES: hypothetical protein [Proteobacteria] | 02529 |
| serine protease [Azoarcus sp. Aa7] | 04408 |
| restriction endonuclease [Azoarcus sp. Aa7] | 00419 |
| DEAD/DEAH box helicase family protein [Azoarcus sp. Aa7] | 06181 |
| hypothetical protein [Azoarcus sp. Aa7] | 03060 |
| GIY-YIG nuclease family protein [Azoarcus sp. Aa7] | 01091 |
| DUF2283 domain-containing protein [Azoarcus rhizosphaerae] | 02098 |
| viperin family antiviral radical SAM protein [Azoarcus sp. Aa7] | 04414 |
| metallophosphoesterase [Aromatoleum diolicum] | 10453 |
| mobile mystery protein B [Azoarcus sp. Aa7] | 00488 |
| helix-turn-helix transcriptional regulator [Azoarcus rhizosphaerae] | 09989 |
| transposase, partial [Azoarcus sp. Aa7] | 06701 |
| hypothetical protein [Azoarcus sp. Aa7] | 06545 |
| hypothetical protein [Azoarcus rhizosphaerae] | 08201 |
| hypothetical protein [Azoarcus sp. Aa7] | 08582 |
| hypothetical protein [Azoarcus sp. Aa7] | 04922 |
| hypothetical protein [Rhodocyclus purpureus] | 07786 |
| DUF2726 domain-containing protein [Thauera phenolivorans] | 05991 |
| MarR family transcriptional regulator [Thauera phenolivorans] | 05622 |
| hypothetical protein [Azoarcus rhizosphaerae] | 02324 |
| ethylbenzene dehydrogenase-related protein [Azoarcus rhizosphaerae] | 09708 |
| hypothetical protein [Azoarcus rhizosphaerae] | 01100 |
| hypothetical protein [Azoarcus rhizosphaerae] | 06351 |
| hypothetical protein [Aromatoleum toluvorans] | 04868 |
| DUF1902 domain-containing protein [Aromatoleum toluvorans] | 04206 |
| N-formylglutamate deformylase [Aromatoleum toluvorans] | 07244 |
| hypothetical protein [Azoarcus rhizosphaerae] | 06051 |
| hypothetical protein [Uliginosibacterium gangwonense] | 08239 |
| hypothetical protein [Uliginosibacterium gangwonense] | 08821 |
| hypothetical protein [Uliginosibacterium gangwonense] | 07073 |
| cupin domain-containing protein [Uliginosibacterium gangwonense] | 03318 |
| gamma-mobile-trio recombinase GmtY [Uliginosibacterium gangwonense] | 09339 |
| hypothetical protein [Uliginosibacterium gangwonense] | 07892 |
| site-specific integrase [Uliginosibacterium gangwonense] | 01543 |
| methyltransferase domain-containing protein [Uliginosibacterium gangwonense] | 05196 |
| DUF637 domain-containing protein [Azoarcus rhizosphaerae] | 02188 |
| hypothetical protein [Azoarcus rhizosphaerae] | 06352 |
| ATP-binding protein [Azoarcus rhizosphaerae] | 02650 |
| DinB family protein [Azoarcus rhizosphaerae] | 03213 |
| PepSY domain-containing protein [Azoarcus rhizosphaerae] | 07816 |
| MULTISPECIES: hypothetical protein [Thauera] | 06683 |
| oligosaccharide flippase family protein [Thauera aminoaromatica] | 09092 |
| type II toxin-antitoxin system VapC family toxin [Azoarcus rhizosphaerae] | 09870 |
| hypothetical protein [Azoarcus rhizosphaerae] | 06021 |
| pyruvate kinase [Rhodocyclus purpureus] | 00527 |
| hypothetical protein [Rhodocyclus purpureus] | 07322 |
| hypothetical protein [Rhodocyclus purpureus] | 09587 |
| hypothetical protein [Aromatoleum diolicum] | 07401 |
| sel1 repeat family protein [Aromatoleum diolicum] | 05821 |

| | |
|---|---|
| beta/gamma crystallin-related protein [Aromatoleum diolicum] | 03016 |
| DUF6498-containing protein [Azoarcus taiwanensis] | 06342 |
| DUF1272 domain-containing protein [Aromatoleum diolicum] | 00543 |
| hypothetical protein [Aromatoleum diolicum] | 08605 |
| metalloregulator ArsR/SmtB family transcription factor [Azoarcus taiwanensis] | 04970 |
| type II toxin-antitoxin system HipA family toxin YjjJ [Aromatoleum diolicum] | 02677 |
| glucose-inhibited division protein B [Azoarcus taiwanensis] | 07453 |
| DUF1850 domain-containing protein [Azoarcus taiwanensis] | 00123 |
| hypothetical protein [Aromatoleum diolicum] | 00663 |
| hypothetical protein [Azoarcus taiwanensis] | 01058 |
| hypothetical protein [Aromatoleum diolicum] | 09249 |
| SAM-dependent methyltransferase [Aromatoleum diolicum] | 01863 |
| class I SAM-dependent methyltransferase [Aromatoleum diolicum] | 03887 |
| fibronectin type III domain-containing protein [Aromatoleum diolicum] | 02547 |
| hypothetical protein [Aromatoleum diolicum] | 06286 |
| hypothetical protein [Aromatoleum diolicum] | 09683 |
| hypothetical protein [Aromatoleum diolicum] | 07396 |
| type I-E CRISPR-associated protein Cas6/Cse3/CasE [Aromatoleum diolicum] | 04765 |
| type I-E CRISPR-associated protein Cse2/CasB [Aromatoleum diolicum] | 08185 |
| type I-E CRISPR-associated protein Cse1/CasA [Aromatoleum diolicum] | 04304 |
| hypothetical protein [Azoarcus taiwanensis] | 00427 |
| hypothetical protein [Aromatoleum diolicum] | 04501 |
| type I-E CRISPR-associated protein Cse1/CasA [Rhodocyclus purpureus] | 03382 |
| MULTISPECIES: hypothetical protein [Proteobacteria] | 02352 |
| type I-E CRISPR-associated protein Cse2/CasB [Rhodocyclus purpureus] | 04056 |
| class I SAM-dependent methyltransferase [Rhodocyclus purpureus] | 10045 |
| UDP-N-acetylglucosamine 2-epimerase [Thauera phenolivorans] | 00431 |
| glycosyltransferase family 2 protein [Thauera aromatica] | 00020 |
| hypothetical protein [Thauera phenolivorans] | 02380 |
| AAA family ATPase [Thauera aromatica] | 02585 |
| PIN domain-containing protein [Thauera aromatica] | 05999 |
| restriction endonuclease [Thauera aromatica] | 06682 |
| MULTISPECIES: DNA cytosine methyltransferase [Betaproteobacteria] | 09188 |
| MULTISPECIES: hypothetical protein [Betaproteobacteria] | 09079 |
| hypothetical protein [Thauera aromatica] | 05026 |
| KilA-N domain-containing protein [Thauera aromatica] | 08809 |
| MULTISPECIES: DUF3577 domain-containing protein [Proteobacteria] | 09077 |
| MULTISPECIES: hypothetical protein [Proteobacteria] | 04478 |
| efflux RND transporter permease subunit, partial [Thauera phenolivorans] | 01026 |
| MULTISPECIES: antirestriction protein [Proteobacteria] | 05183 |
| efflux RND transporter permease subunit, partial [Thauera phenolivorans] | 09065 |
| MULTISPECIES: hypothetical protein [Proteobacteria] | 08483 |
| site-specific integrase [Thauera phenolivorans] | 08466 |
| MULTISPECIES: hypothetical protein [Proteobacteria] | 07184 |
| MULTISPECIES: stable inheritance protein KleA [Proteobacteria] | 07589 |
| cell division protein FtsI [Uliginosibacterium aquaticum] | 03782 |
| hypothetical protein [Uliginosibacterium aquaticum] | 01495 |
| YXWGXW repeat-containing protein [Uliginosibacterium aquaticum] | 05049 |
| phasin family protein [Zoogloea dura] | 09956 |
| hypothetical protein [Zoogloea dura] | 06939 |
| hypothetical protein [Azoarcus sp. DD4] | 02107 |
| hypothetical protein [Azoarcus sp. DD4] | 05019 |
| hypothetical protein [Azoarcus sp. DD4] | 10283 |
| hypothetical protein [Azoarcus sp. DD4] | 00957 |
| M15 family metallopeptidase [Azoarcus sp. DD4] | 06636 |
| hypothetical protein [Zoogloea dura] | 06922 |
| hypothetical protein [Zoogloea dura] | 08224 |
| hypothetical protein [Zoogloea dura] | 08527 |
| hypothetical protein [Zoogloea dura] | 05554 |

| | |
|---|---|
| hypothetical protein [Zoogloea dura] | 09517 |
| TIR domain-containing protein [Thauera humireducens] | 07430 |
| hypothetical protein [Zoogloea dura] | 00612 |
| hypothetical protein [Zoogloea dura] | 02750 |
| hypothetical protein [Zoogloea dura] | 05164 |
| zinc ribbon domain-containing protein [Zoogloea dura] | 06150 |
| hypothetical protein [Zoogloea dura] | 04417 |
| hypothetical protein [Zoogloea dura] | 02921 |
| hypothetical protein [Azoarcus sp. DD4] | 08082 |
| hypothetical protein [Zoogloea dura] | 01382 |
| KAP family NTPase [Zoogloea dura] | 06358 |
| hypothetical protein [Zoogloea dura] | 00590 |
| hypothetical protein [Azoarcus sp. DD4] | 04406 |
| hypothetical protein [Zoogloea dura] | 08244 |
| hypothetical protein, partial [Aromatoleum evansii] | 04853 |
| RES family NAD+ phosphorylase [Aromatoleum evansii] | 03458 |
| ABC transporter substrate-binding protein [Thauera humireducens] | 04737 |
| hypothetical protein [Azoarcus sp. L1K30] | 09922 |
| hypothetical protein [Zoogloea dura] | 01111 |
| dihydrodipicolinate reductase [Zoogloea dura] | 02924 |
| hypothetical protein [Azoarcus sp. L1K30] | 09621 |
| hypothetical protein [Zoogloea dura] | 07888 |
| hypothetical protein [Azoarcus sp. L1K30] | 04244 |
| hypothetical protein [Zoogloea dura] | 00882 |
| hypothetical protein [Zoogloea dura] | 05744 |
| hypothetical protein [Zoogloea dura] | 02432 |
| hypothetical protein [Zoogloea dura] | 05190 |
| DUF1186 domain-containing protein [Zoogloea sp. G-4-1-14] | 01357 |
| PEP-CTERM sorting domain-containing protein [Azoarcus sp. L1K30] | 00337 |
| hypothetical protein [Thauera lacus] | 00395 |
| hypothetical protein [Aromatoleum evansii] | 02171 |
| MAPEG family protein [Zoogloea dura] | 00930 |
| exodeoxyribonuclease VII small subunit [Aromatoleum evansii] | 01168 |
| IS5 family transposase [Aromatoleum evansii] | 09709 |
| hypothetical protein [Aromatoleum evansii] | 00466 |
| DUF2779 domain-containing protein [Aromatoleum evansii] | 05764 |
| hypothetical protein [Aromatoleum evansii] | 06813 |
| hypothetical protein [Zoogloea dura] | 07950 |
| hypothetical protein [Azoarcus sp. DD4] | 09428 |
| hypothetical protein [Zoogloea dura] | 08226 |
| phasin family protein [Zoogloea dura] | 07948 |
| DUF4062 domain-containing protein [Zoogloea dura] | 03574 |
| hypothetical protein [Zoogloea dura] | 07872 |
| hypothetical protein [Azoarcus sp. DD4] | 07123 |
| hypothetical protein [Zoogloea dura] | 00447 |
| hypothetical protein [Zoogloea dura] | 00038 |
| DUF4124 domain-containing protein [Zoogloea dura] | 02310 |
| hypothetical protein [Zoogloea dura] | 09132 |
| PEP-CTERM sorting domain-containing protein [Zoogloea dura] | 09711 |
| hypothetical protein [Zoogloea dura] | 04886 |
| S8 family serine peptidase [Zoogloea dura] | 09952 |
| flavodoxin family protein [Zoogloea dura] | 06508 |
| DUF3369 domain-containing protein [Zoogloea dura] | 10151 |
| transporter substrate-binding domain-containing protein [Zoogloea dura] | 06579 |
| hypothetical protein [Zoogloea dura] | 10494 |
| endonuclease [Zoogloea dura] | 06432 |
| hypothetical protein [Zoogloea dura] | 09693 |
| hypothetical protein [Zoogloea dura] | 00405 |
| hypothetical protein [Zoogloea dura] | 08745 |

| | |
|---|---|
| helix-turn-helix domain-containing protein [Zoogloea dura] | 08305 |
| FMN-binding protein [Zoogloea dura] | 07324 |
| DUF4405 domain-containing protein [Zoogloea dura] | 08610 |
| toll/interleukin-1 receptor domain-containing protein [Zoogloea dura] | 00766 |
| hypothetical protein [Zoogloea dura] | 02333 |
| hypothetical protein [Zoogloea dura] | 09575 |
| rhombosortase [Zoogloea dura] | 03911 |
| hypothetical protein [Zoogloea dura] | 03246 |
| hypothetical protein [Azoarcus sp. DD4] | 05153 |
| hypothetical protein [Zoogloea dura] | 08499 |
| hypothetical protein [Zoogloea dura] | 03435 |
| LTA synthase family protein [Zoogloea sp. G-4-1-14] | 01209 |
| beta-3-deoxy-D-manno-oct-2-ulosonic acid transferase [Zoogloea dura] | 00719 |
| ABC transporter ATP-binding protein [Zoogloea dura] | 07571 |
| toll/interleukin-1 receptor domain-containing protein [Azoarcus sp. DD4] | 10399 |
| hypothetical protein [Azoarcus sp. DD4] | 02526 |
| hypothetical protein [Zoogloea dura] | 09162 |
| hypothetical protein [Zoogloea dura] | 01883 |
| hypothetical protein [Zoogloea dura] | 09158 |
| hypothetical protein [Azoarcus sp. DD4] | 03816 |
| hypothetical protein [Zoogloea dura] | 02248 |
| hypothetical protein [Zoogloea dura] | 09514 |
| hypothetical protein [Zoogloea dura] | 10495 |
| glycosyltransferase family 9 protein [Zoogloea dura] | 03434 |
| hypothetical protein [Zoogloea dura] | 05158 |
| Clp protease ClpP [Zoogloea dura] | 08612 |
| hypothetical protein [Zoogloea dura] | 01461 |
| rhombotarget lipoprotein [Zoogloea dura] | 01108 |
| hypothetical protein [Thauera humireducens] | 00421 |
| transposase domain-containing protein [Azoarcus communis] | 02679 |
| hypothetical protein [Uliginosibacterium aquaticum] | 04154 |
| threonine/serine exporter family protein [Uliginosibacterium aquaticum] | 09403 |
| YiiD C-terminal domain-containing protein [Uliginosibacterium aquaticum] | 08915 |
| PD40 domain-containing protein [Uliginosibacterium aquaticum] | 09302 |
| hypothetical protein [Uliginosibacterium aquaticum] | 05864 |
| hypothetical protein [Uliginosibacterium aquaticum] | 08850 |
| hypothetical protein [Uliginosibacterium aquaticum] | 09299 |
| GtrA family protein [Uliginosibacterium aquaticum] | 02755 |
| hypothetical protein [Uliginosibacterium aquaticum] | 08447 |
| ABC transporter substrate-binding protein [Uliginosibacterium aquaticum] | 03519 |
| hypothetical protein [Uliginosibacterium aquaticum] | 08913 |
| helix-turn-helix domain-containing protein [Uliginosibacterium aquaticum] | 06080 |
| anaerobic ribonucleoside-triphosphate reductase activating protein [Uliginosibacterium aquaticum] | 07408 |
| polysaccharide pyruvyl transferase family protein [Azoarcus communis] | 06886 |
| RpiB/LacA/LacB family sugar-phosphate isomerase [Uliginosibacterium aquaticum] | 05560 |
| rhamnogalacturonan acetylesterase [Uliginosibacterium aquaticum] | 08322 |
| ABC transporter permease subunit [Aromatoleum pumilum] | 03354 |
| hypothetical protein [Aromatoleum pumilum] | 10236 |
| pectate lyase [Uliginosibacterium aquaticum] | 04537 |
| glycosyltransferase [Aromatoleum pumilum] | 09871 |
| Gfo/Idh/MocA family oxidoreductase [Uliginosibacterium aquaticum] | 00990 |
| hypothetical protein [Uliginosibacterium aquaticum] | 01219 |
| hypothetical protein [Uliginosibacterium aquaticum] | 04538 |
| hypothetical protein [Uliginosibacterium aquaticum] | 07307 |
| hypothetical protein [Uliginosibacterium aquaticum] | 01804 |
| Fic family protein [Azoarcus sp. TTM-91] | 07862 |
| GNAT family N-acetyltransferase [Uliginosibacterium aquaticum] | 06718 |
| DUF441 domain-containing protein [Uliginosibacterium aquaticum] | 06658 |
| deoxyribose-phosphate aldolase [Uliginosibacterium aquaticum] | 10509 |

| | |
|---|---|
| alpha/beta fold hydrolase [Uliginosibacterium aquaticum] | 07301 |
| TetR family transcriptional regulator [Uliginosibacterium aquaticum] | 10476 |
| alginate lyase family protein [Uliginosibacterium aquaticum] | 04900 |
| hypothetical protein [Uliginosibacterium aquaticum] | 07868 |
| high-potential iron-sulfur protein [Uliginosibacterium aquaticum] | 04542 |
| aldose 1-epimerase [Uliginosibacterium aquaticum] | 03057 |
| hypothetical protein [Uliginosibacterium aquaticum] | 02551 |
| DEAD/DEAH box helicase family protein [Zoogloea ramigera] | 02114 |
| hypothetical protein [Zoogloea ramigera] | 05111 |
| VOC family protein [Uliginosibacterium aquaticum] | 08104 |
| DUF4914 family protein [Uliginosibacterium aquaticum] | 04642 |
| oligosaccharide flippase family protein [Uliginosibacterium aquaticum] | 05347 |
| NUDIX domain-containing protein [Uliginosibacterium aquaticum] | 09720 |
| NADAR family protein [Uliginosibacterium aquaticum] | 03525 |
| hypothetical protein [Uliginosibacterium aquaticum] | 07011 |
| tyrosine-type recombinase/integrase [Oryzomicrobium terrae] | 08842 |
| PAS domain S-box protein [Uliginosibacterium aquaticum] | 08106 |
| TIGR02285 family protein [Uliginosibacterium aquaticum] | 03314 |
| DNA-binding protein [Azoarcus sp. TTM-91] | 00072 |
| exodeoxyribonuclease I [Uliginosibacterium aquaticum] | 00460 |
| pectinesterase family protein [Uliginosibacterium aquaticum] | 04029 |
| TRAP transporter substrate-binding protein DctP [Uliginosibacterium aquaticum] | 02237 |
| YCF48-related protein [Uliginosibacterium aquaticum] | 08317 |
| hypothetical protein [Zoogloea dura] | 02408 |
| recombinase RecT [Zoogloea dura] | 02438 |
| TonB family protein [Zoogloea dura] | 09144 |
| hypothetical protein [Zoogloea dura] | 03416 |
| hypothetical protein [Zoogloea dura] | 09698 |
| hypothetical protein [Zoogloea dura] | 06942 |
| MULTISPECIES: helix-turn-helix domain-containing protein [Zoogloea] | 10018 |
| GNAT family N-acetyltransferase [Zoogloea dura] | 01466 |
| hypothetical protein [Zoogloea dura] | 01214 |
| hypothetical protein [Thauera hydrothermalis] | 05441 |
| hypothetical protein [Thauera hydrothermalis] | 00031 |
| hypothetical protein [Thauera hydrothermalis] | 03681 |
| hypothetical protein [Thauera hydrothermalis] | 03700 |
| hypothetical protein [Zoogloea dura] | 10014 |
| hypothetical protein [Zoogloea dura] | 10427 |
| chemotaxis protein CheW [Zoogloea dura] | 07596 |
| AAA domain-containing protein [Azoarcus communis] | 05463 |
| hypothetical protein [Azoarcus communis] | 03480 |
| anti-sigma factor [Zoogloea dura] | 00608 |
| hypothetical protein [Azoarcus communis] | 03217 |
| hypothetical protein [Azoarcus communis] | 00350 |
| sensor histidine kinase [Azoarcus communis] | 05191 |
| DUF4157 domain-containing protein [Zoogloea dura] | 00614 |
| VWA domain-containing protein [Zoogloea dura] | 01968 |
| hypothetical protein [Uliginosibacterium aquaticum] | 04028 |
| SDR family NAD(P)-dependent oxidoreductase [Zoogloea dura] | 06810 |
| hypothetical protein [Uliginosibacterium aquaticum] | 09295 |
| hypothetical protein [Uliginosibacterium aquaticum] | 04158 |
| hypothetical protein [Uliginosibacterium aquaticum] | 03053 |
| hypothetical protein [Uliginosibacterium aquaticum] | 02754 |
| site-specific integrase [Uliginosibacterium aquaticum] | 07659 |
| DEAD/DEAH box helicase [Uliginosibacterium sp. IMCC34675] | 08315 |
| GIY-YIG nuclease family protein [Uliginosibacterium aquaticum] | 00761 |
| heparinase II/III family protein [Uliginosibacterium aquaticum] | 10502 |
| MULTISPECIES: pectate lyase [Uliginosibacterium] | 07967 |
| MULTISPECIES: DUF1161 domain-containing protein [Uliginosibacterium] | 01468 |

| | |
|---|---|
| MULTISPECIES: hypothetical protein [Uliginosibacterium] | 10525 |
| MULTISPECIES: DUF1653 domain-containing protein [Uliginosibacterium] | 08717 |
| MULTISPECIES: hypothetical protein [Uliginosibacterium] | 00991 |
| MULTISPECIES: heme-degrading domain-containing protein [Uliginosibacterium] | 02460 |
| MULTISPECIES: hypothetical protein [Uliginosibacterium] | 05860 |
| MULTISPECIES: hypothetical protein [Uliginosibacterium] | 01221 |
| MULTISPECIES: pectate lyase [Uliginosibacterium] | 05566 |
| MULTISPECIES: right-handed parallel beta-helix repeat-containing protein [Uliginosibacterium] | 09578 |
| PEP-CTERM sorting domain-containing protein [Zoogloea dura] | 09700 |
| DUF2058 domain-containing protein [Zoogloea dura] | 05589 |
| glycosyltransferase family 39 protein [Zoogloea dura] | 10402 |
| MULTISPECIES: hypothetical protein [Rhodocyclales] | 02571 |
| MULTISPECIES: hypothetical protein [Betaproteobacteria] | 01586 |
| hypothetical protein [Azoarcus indigens] | 04387 |
| HK97 family phage prohead protease [Azoarcus indigens] | 08895 |
| minor coat protein [Thauera linaloolentis] | 01021 |
| DUF3015 family protein [Propionivibrio dicarboxylicus] | 07000 |
| hypothetical protein [Azoarcus indigens] | 03218 |
| hypothetical protein [Azoarcus indigens] | 00903 |
| DUF1493 family protein [Azoarcus indigens] | 02741 |
| phage/plasmid replication protein, II/X family [Thauera linaloolentis] | 05985 |
| hypothetical protein [Thauera linaloolentis] | 02870 |
| DUF1003 domain-containing protein [Azoarcus indigens] | 03495 |
| DUF1993 domain-containing protein [Propionivibrio dicarboxylicus] | 05559 |
| hypothetical protein [Azoarcus indigens] | 06145 |
| acyl-CoA thioesterase [Propionivibrio dicarboxylicus] | 10266 |
| carboxymuconolactone decarboxylase family protein [Propionivibrio dicarboxylicus] | 01213 |
| type VI secretion system protein TssA [Azoarcus indigens] | 09189 |
| hypothetical protein [Azoarcus indigens] | 10085 |
| hypothetical protein [Azoarcus indigens] | 01880 |
| hypothetical protein [Azoarcus indigens] | 02644 |
| type II toxin-antitoxin system CcdA family antitoxin [Thauera linaloolentis] | 07155 |
| type II secretion system GspH family protein [Propionivibrio dicarboxylicus] | 06293 |
| prepilin-type N-terminal cleavage/methylation domain-containing protein [Propionivibrio dicarboxylicus] | 00201 |
| anti-sigma regulatory factor [Azoarcus indigens] | 04392 |
| anti-sigma regulatory factor [Azoarcus indigens] | 00143 |
| STAS domain-containing protein [Azoarcus indigens] | 03728 |
| DMT family transporter [Azoarcus indigens] | 08807 |
| hypothetical protein [Azoarcus indigens] | 08169 |
| class I SAM-dependent methyltransferase [Aromatoleum aromaticum] | 03566 |
| hypothetical protein [Aromatoleum aromaticum] | 07372 |
| hypothetical protein [Thauera linaloolentis] | 02582 |
| hypothetical protein [Azoarcus indigens] | 01695 |
| zonular occludens toxin [Thauera linaloolentis] | 02710 |
| AHH domain-containing protein, partial [Azoarcus indigens] | 03729 |
| PRTRC system ParB family protein [Aromatoleum aromaticum] | 05658 |
| transposase [Thauera linaloolentis] | 02211 |
| thioredoxin family protein [Thauera linaloolentis] | 07435 |
| DUF134 domain-containing protein [Propionivibrio dicarboxylicus] | 08407 |
| hypothetical protein [Propionivibrio dicarboxylicus] | 09276 |
| DNA methylase [Thauera chlorobenzoica] | 00861 |
| BREX-3 system phosphatase PglZ [Thauera chlorobenzoica] | 07775 |
| PRTRC system ThiF family protein [Aromatoleum aromaticum] | 02818 |
| PRTRC system protein A [Aromatoleum aromaticum] | 08416 |
| PRTRC system protein B [Aromatoleum aromaticum] | 07119 |
| PRTRC system protein F [Aromatoleum aromaticum] | 02174 |
| hypothetical protein [Thauera chlorobenzoica] | 08755 |
| PRTRC system protein E [Aromatoleum aromaticum] | 00256 |
| hypothetical protein [Thauera chlorobenzoica] | 02077 |

| | |
|---|---|
| non-homologous end-joining DNA ligase [Azoarcus indigens] | 04513 |
| DUF4435 domain-containing protein [Thauera chlorobenzoica] | 01036 |
| YjbH domain-containing protein [Propionivibrio dicarboxylicus] | 03274 |
| AzlD domain-containing protein [Thauera linaloolentis] | 04664 |
| UvrD-helicase domain-containing protein [Thauera chlorobenzoica] | 06704 |
| nucleoside 2-deoxyribosyltransferase [Aromatoleum aromaticum] | 09021 |
| DUF2442 domain-containing protein [Thauera chlorobenzoica] | 08665 |
| TIGR04255 family protein [Azoarcus indigens] | 05654 |
| NUDIX domain-containing protein [Propionivibrio dicarboxylicus] | 08754 |
| hypothetical protein [Azoarcus indigens] | 01361 |
| filamentous hemagglutinin N-terminal domain-containing protein [Azoarcus indigens] | 01259 |
| prepilin peptidase [Thauera linaloolentis] | 04469 |
| response regulator [Azoarcus indigens] | 05165 |
| LuxR C-terminal-related transcriptional regulator [Azoarcus indigens] | 07843 |
| hypothetical protein [Thauera linaloolentis] | 05390 |
| type II toxin-antitoxin system PemK/MazF family toxin [Thauera chlorobenzoica] | 03854 |
| hypothetical protein [Aromatoleum aromaticum] | 02817 |
| extracellular solute-binding protein [Azoarcus indigens] | 02399 |
| hypothetical protein [Azoarcus indigens] | 04226 |
| RHS repeat-associated core domain-containing protein, partial [Azoarcus indigens] | 08783 |
| MoaD/ThiS family protein [Aromatoleum aromaticum] | 06302 |
| hypothetical protein [Propionivibrio dicarboxylicus] | 06787 |
| hypothetical protein [Aromatoleum aromaticum] | 03605 |
| contact-dependent growth inhibition system immunity protein [Azoarcus indigens] | 09911 |
| acid-shock protein [Propionivibrio dicarboxylicus] | 06619 |
| hypothetical protein, partial [Azoarcus indigens] | 08215 |
| CstA-like transporter-associated (seleno)protein [Azoarcus indigens] | 02204 |
| phosphate/phosphite/phosphonate ABC transporter substrate-binding protein [Propionivibrio dicarboxylicus] | 05432 |
| hypothetical protein [Propionivibrio dicarboxylicus] | 04525 |
| serine protease inhibitor ecotin [Thauera linaloolentis] | 09055 |
| hypothetical protein [Azoarcus indigens] | 03399 |
| zinc-binding dehydrogenase [Azoarcus indigens] | 03491 |
| ATP-binding protein [Propionivibrio dicarboxylicus] | 07235 |
| DUF4105 domain-containing protein [Propionivibrio dicarboxylicus] | 00592 |
| DUF1015 family protein [Propionivibrio dicarboxylicus] | 06788 |
| phytanoyl-CoA dioxygenase family protein [Azoarcus indigens] | 04881 |
| hypothetical protein [Aromatoleum aromaticum] | 09008 |
| hypothetical protein [Aromatoleum aromaticum] | 03969 |
| GlpM family protein [Propionivibrio dicarboxylicus] | 03391 |
| amidohydrolase family protein [Azoarcus indigens] | 02092 |
| hypothetical protein [Azoarcus indigens] | 10148 |
| hypothetical protein [Azoarcus indigens] | 06505 |
| TonB family protein [Thauera linaloolentis] | 03400 |
| hypothetical protein [Propionivibrio dicarboxylicus] | 01918 |
| class I SAM-dependent methyltransferase [Thauera linaloolentis] | 05148 |
| hypothetical protein [Azoarcus indigens] | 06382 |
| DoxX family protein [Azoarcus indigens] | 03726 |
| major capsid protein [Azoarcus indigens] | 06384 |
| J domain-containing protein [Aromatoleum aromaticum] | 05296 |
| hypothetical protein [Aromatoleum aromaticum] | 01161 |
| hypothetical protein [Azoarcus indigens] | 04241 |
| hypothetical protein [Aromatoleum aromaticum] | 00758 |
| glycosyltransferase [Aromatoleum aromaticum] | 04115 |
| hypothetical protein [Azoarcus indigens] | 03727 |
| hypothetical protein [Aromatoleum aromaticum] | 07723 |
| MULTISPECIES: hypothetical protein [Azoarcus] | 08806 |
| hypothetical protein [Azoarcus indigens] | 02739 |
| pyrimidine dimer DNA glycosylase/endonuclease V [Aromatoleum aromaticum] | 04616 |
| tagaturonate reductase [Propionivibrio dicarboxylicus] | 01912 |

| | |
|---|---|
| type VI secretion system lipoprotein TssJ [Azoarcus indigens] | 04240 |
| mannonate dehydratase [Propionivibrio dicarboxylicus] | 06819 |
| SRPBCC domain-containing protein [Azoarcus indigens] | 06383 |
| DUF3592 domain-containing protein [Azoarcus indigens] | 01202 |
| hypothetical protein [Azoarcus indigens] | 06503 |
| hypothetical protein [Propionivibrio dicarboxylicus] | 09497 |
| hypothetical protein [Azoarcus indigens] | 10490 |
| DNA-3-methyladenine glycosylase I [Thauera linaloolentis] | 06326 |
| DUF6364 family protein [Aromatoleum aromaticum] | 06625 |
| DUF3861 domain-containing protein [Thauera linaloolentis] | 09855 |
| heme utilization cystosolic carrier protein HutX [Thauera linaloolentis] | 08808 |
| ACT domain-containing protein [Propionivibrio dicarboxylicus] | 03538 |
| hypothetical protein [Azoarcus indigens] | 03234 |
| hypothetical protein [Azoarcus indigens] | 05134 |
| 5-deoxy-glucuronate isomerase [Propionivibrio dicarboxylicus] | 07306 |
| DUF3037 domain-containing protein [Thauera chlorobenzoica] | 07514 |
| DUF2283 domain-containing protein [Aromatoleum aromaticum] | 06313 |
| hypothetical protein [Thauera linaloolentis] | 06386 |
| HP1 family phage holin [Thauera chlorobenzoica] | 01019 |
| PD-(D/E)XK motif protein [Aromatoleum aromaticum] | 08321 |
| DUF2183 domain-containing protein [Aromatoleum tolulyticum] | 07680 |
| hypothetical protein [Aromatoleum tolulyticum] | 00236 |
| YdaS family helix-turn-helix protein [Thauera chlorobenzoica] | 06827 |
| hypothetical protein [Aromatoleum tolulyticum] | 01867 |
| hypothetical protein [Thauera chlorobenzoica] | 09067 |
| hypothetical protein [Aromatoleum aromaticum] | 04465 |
| hypothetical protein [Aromatoleum aromaticum] | 07727 |
| hypothetical protein [Aromatoleum tolulyticum] | 01759 |
| outer membrane protein assembly factor BamE [Aromatoleum tolulyticum] | 01343 |
| MULTISPECIES: hypothetical protein [Aromatoleum] | 07726 |
| hypothetical protein [Aromatoleum tolulyticum] | 08594 |
| siderophore biosynthesis protein [Aromatoleum tolulyticum] | 03124 |
| GNAT family N-acetyltransferase [Aromatoleum tolulyticum] | 05823 |
| hypothetical protein [Aromatoleum tolulyticum] | 03612 |
| hypothetical protein [Aromatoleum aromaticum] | 07357 |
| hypothetical protein [Aromatoleum tolulyticum] | 05947 |
| hypothetical protein [Aromatoleum aromaticum] | 03288 |
| hypothetical protein [Thauera chlorobenzoica] | 06850 |
| methyl-accepting chemotaxis protein [Azoarcus indigens] | 04017 |
| hypothetical protein [Aromatoleum aromaticum] | 09807 |
| amidohydrolase family protein [Aromatoleum tolulyticum] | 04291 |
| NEW3 domain-containing protein [Aromatoleum aromaticum] | 09641 |
| major capsid protein [Thauera chlorobenzoica] | 01040 |
| helix-turn-helix domain-containing protein [Aromatoleum tolulyticum] | 05433 |
| hypothetical protein [Thauera chlorobenzoica] | 07781 |
| TRAP transporter substrate-binding protein DctP [Aromatoleum tolulyticum] | 03571 |
| type II toxin-antitoxin system HicA family toxin [Thauera chlorobenzoica] | 01047 |
| hypothetical protein [Aromatoleum tolulyticum] | 04846 |
| hypothetical protein [Aromatoleum tolulyticum] | 05097 |
| hypothetical protein [Thauera chlorobenzoica] | 00317 |
| hypothetical protein [Thauera chlorobenzoica] | 08135 |
| hypothetical protein [Thauera chlorobenzoica] | 03197 |
| hypothetical protein [Thauera chlorobenzoica] | 00318 |
| hypothetical protein [Aromatoleum aromaticum] | 02471 |
| hypothetical protein [Thauera chlorobenzoica] | 02033 |
| DNA/RNA non-specific endonuclease [Aromatoleum aromaticum] | 05323 |
| hypothetical protein [Aromatoleum aromaticum] | 09379 |
| conjugal transfer protein TrbH [Aromatoleum aromaticum] | 08482 |
| Rha family transcriptional regulator [Thauera chlorobenzoica] | 04789 |

| | |
|---|---|
| terminase small subunit [Thauera chlorobenzoica] | 00292 |
| DUF4160 domain-containing protein [Thauera chlorobenzoica] | 07187 |
| MBL fold metallo-hydrolase [Thauera chlorobenzoica] | 05056 |
| hypothetical protein [Thauera chlorobenzoica] | 00425 |
| hypothetical protein [Aromatoleum aromaticum] | 05214 |
| UDP-N-acetylglucosamine--LPS N-acetylglucosamine transferase [Aromatoleum tolulyticum] | 04830 |
| hypothetical protein [Aromatoleum aromaticum] | 07180 |
| hypothetical protein [Thauera chlorobenzoica] | 02326 |
| hypothetical protein [Thauera chlorobenzoica] | 00001 |
| acyl-CoA dehydrogenase [Thauera chlorobenzoica] | 01654 |
| phage major capsid protein [Aromatoleum tolulyticum] | 07803 |
| uracil-DNA glycosylase [Aromatoleum tolulyticum] | 10625 |
| Clp protease ClpP [Aromatoleum aromaticum] | 06048 |
| hypothetical protein [Aromatoleum aromaticum] | 08136 |
| hypothetical protein [Aromatoleum aromaticum] | 00573 |
| MULTISPECIES: hypothetical protein [Aromatoleum] | 03620 |
| hypothetical protein [Aromatoleum aromaticum] | 03565 |
| LuxR C-terminal-related transcriptional regulator [Aromatoleum tolulyticum] | 10330 |
| potassium channel family protein [Aromatoleum tolulyticum] | 05540 |
| hypothetical protein [Propionivibrio dicarboxylicus] | 00177 |
| hypothetical protein [Aromatoleum aromaticum] | 08058 |
| TonB-dependent receptor [Thauera linaloolentis] | 05466 |
| hypothetical protein [Thauera linaloolentis] | 10078 |
| hypothetical protein [Thauera linaloolentis] | 05518 |
| transcriptional regulator [Aromatoleum aromaticum] | 06959 |
| cupin domain-containing protein [Propionivibrio dicarboxylicus] | 02219 |
| hypothetical protein [Thauera linaloolentis] | 03899 |
| alpha/beta hydrolase [Propionivibrio dicarboxylicus] | 08629 |
| glucuronate isomerase [Propionivibrio dicarboxylicus] | 05863 |
| conjugal transfer protein TrbD [Aromatoleum aromaticum] | 00017 |
| hypothetical protein [Thauera chlorobenzoica] | 03367 |
| restriction endonuclease subunit S [Aromatoleum aromaticum] | 07132 |
| DUF4351 domain-containing protein [Thauera linaloolentis] | 07849 |
| sulfite exporter TauE/SafE family protein [Propionivibrio dicarboxylicus] | 02651 |
| hypothetical protein [Aromatoleum aromaticum] | 01997 |
| zincin-like metallopeptidase domain-containing protein [Aromatoleum aromaticum] | 07373 |
| DMT family transporter [Aromatoleum aromaticum] | 04843 |
| TIR domain-containing protein [Thauera linaloolentis] | 06677 |
| hypothetical protein [Aromatoleum aromaticum] | 03617 |
| PfkB family carbohydrate kinase [Propionivibrio dicarboxylicus] | 05973 |
| nucleotidyltransferase domain-containing protein [Thauera linaloolentis] | 08367 |
| LuxR C-terminal-related transcriptional regulator [Thauera linaloolentis] | 00019 |
| CHAT domain-containing protein [Propionivibrio dicarboxylicus] | 07645 |
| hypothetical protein [Aromatoleum aromaticum] | 04294 |
| AbrB family transcriptional regulator [Thauera chlorobenzoica] | 05087 |
| DUF4384 domain-containing protein [Propionivibrio dicarboxylicus] | 07003 |
| phosphate acetyltransferase [Propionivibrio dicarboxylicus] | 00188 |
| zinc-finger-containing protein [Aromatoleum aromaticum] | 08581 |
| type I toxin-antitoxin system ptaRNA1 family toxin [Aromatoleum aromaticum] | 05401 |
| acyltransferase [Thauera linaloolentis] | 02826 |
| hypothetical protein [Thauera chlorobenzoica] | 03365 |
| hypothetical protein [Thauera chlorobenzoica] | 03762 |
| ankyrin repeat domain-containing protein [Thauera chlorobenzoica] | 05090 |
| TIGR04348 family glycosyltransferase [Thauera chlorobenzoica] | 07448 |
| SUMF1/EgtB/PvdO family nonheme iron enzyme [Thauera chlorobenzoica] | 09066 |
| hypothetical protein [Thauera chlorobenzoica] | 00760 |
| hypothetical protein [Aromatoleum tolulyticum] | 09374 |
| hypothetical protein [Aromatoleum aromaticum] | 08269 |
| hypothetical protein [Aromatoleum tolulyticum] | 06022 |

| | |
|---|---|
| hypothetical protein [Aromatoleum tolulyticum] | 09367 |
| hypothetical protein [Aromatoleum aromaticum] | 05018 |
| retention module-containing protein, partial [Aromatoleum tolulyticum] | 04503 |
| transposase [Aromatoleum tolulyticum] | 07508 |
| hypothetical protein [Thauera chlorobenzoica] | 01297 |
| ABC transporter substrate-binding protein [Thauera chlorobenzoica] | 10296 |
| 1-acyl-sn-glycerol-3-phosphate acyltransferase [Aromatoleum aromaticum] | 05528 |
| hypothetical protein [Aromatoleum tolulyticum] | 09680 |
| hemerythrin domain-containing protein [Thauera chlorobenzoica] | 05027 |
| hypothetical protein [Aromatoleum aromaticum] | 03618 |
| hypothetical protein [Thauera chlorobenzoica] | 02583 |
| alpha/beta hydrolase [Aromatoleum tolulyticum] | 04412 |
| hypothetical protein [Aromatoleum tolulyticum] | 01991 |
| hypothetical protein [Aromatoleum aromaticum] | 01446 |
| hypothetical protein [Aromatoleum aromaticum] | 00548 |
| hypothetical protein [Aromatoleum tolulyticum] | 03655 |
| lipid-A-disaccharide synthase N-terminal domain-containing protein [Aromatoleum aromaticum] | 01541 |
| DUF4365 domain-containing protein [Aromatoleum tolulyticum] | 09120 |
| hypothetical protein [Aromatoleum aromaticum] | 07725 |
| acyl carrier protein [Aromatoleum aromaticum] | 04574 |
| hypothetical protein [Azoarcus indigens] | 10493 |
| hypothetical protein [Propionivibrio dicarboxylicus] | 07393 |
| hypothetical protein [Azoarcus indigens] | 07595 |
| hypothetical protein [Aromatoleum aromaticum] | 06792 |
| DUF892 family protein [Azoarcus indigens] | 01878 |
| hypothetical protein [Azoarcus communis] | 01095 |
| glutathione S-transferase [Aromatoleum aromaticum] | 09332 |
| PEP-CTERM sorting domain-containing protein [Azoarcus indigens] | 06902 |
| DUF2190 family protein [Azoarcus communis] | 03292 |
| hypothetical protein [Azoarcus indigens] | 03396 |
| hypothetical protein [Azoarcus communis] | 10312 |
| phage BR0599 family protein [Azoarcus communis] | 03580 |
| hypothetical protein [Azoarcus indigens] | 09477 |
| hypothetical protein [Azoarcus communis] | 08877 |
| hypothetical protein [Azoarcus communis] | 09978 |
| hypothetical protein [Azoarcus indigens] | 05125 |
| hypothetical protein [Azoarcus indigens] | 03230 |
| extensin family protein [Azoarcus indigens] | 08209 |
| ferritin-like domain-containing protein [Azoarcus indigens] | 05830 |
| helix-turn-helix domain-containing protein [Azoarcus communis] | 01641 |
| ATP-binding protein [Aromatoleum aromaticum] | 07446 |
| hypothetical protein [Aromatoleum aromaticum] | 03968 |
| hypothetical protein [Aromatoleum aromaticum] | 09381 |
| DUF2730 family protein [Azoarcus communis] | 06117 |
| hypothetical protein [Aromatoleum aromaticum] | 09037 |
| hypothetical protein [Aromatoleum aromaticum] | 07718 |
| hypothetical protein [Aromatoleum aromaticum] | 04615 |
| ATP-grasp domain-containing protein [Azoarcus indigens] | 00366 |
| YSC84-related protein [Azoarcus indigens] | 00676 |
| Tm-1-like ATP-binding domain-containing protein [Aromatoleum aromaticum] | 00805 |
| DoxX family protein [Azoarcus indigens] | 02086 |
| hypothetical protein [Azoarcus indigens] | 04730 |
| hypothetical protein [Aromatoleum aromaticum] | 07194 |
| twin-arginine translocation signal domain-containing protein [Aromatoleum aromaticum] | 02602 |
| YkgJ family cysteine cluster protein [Azoarcus communis] | 07318 |
| hypothetical protein [Azoarcus indigens] | 03231 |
| glycosyltransferase [Azoarcus indigens] | 05911 |
| prepilin-type N-terminal cleavage/methylation domain-containing protein [Azoarcus indigens] | 00562 |
| hypothetical protein [Azoarcus communis] | 00920 |

| | |
|---|---|
| type VI secretion system-associated FHA domain protein TagH [Azoarcus indigens] | 03233 |
| hypothetical protein [Azoarcus communis] | 04855 |
| hypothetical protein [Azoarcus indigens] | 07485 |
| hypothetical protein [Aromatoleum aromaticum] | 00784 |
| hypothetical protein [Azoarcus communis] | 04316 |
| hypothetical protein [Azoarcus indigens] | 10492 |
| GTP-binding protein [Aromatoleum aromaticum] | 01191 |
| 3-hydroxyanthranilate 3,4-dioxygenase [Azoarcus indigens] | 02640 |
| hypothetical protein [Azoarcus indigens] | 08193 |
| hypothetical protein [Aromatoleum aromaticum] | 01159 |
| hypothetical protein [Azoarcus indigens] | 06038 |
| hypothetical protein [Azoarcus indigens] | 07264 |
| ATPase [Azoarcus communis] | 08510 |
| host-nuclease inhibitor Gam family protein [Azoarcus communis] | 09856 |
| hypothetical protein [Azoarcus communis] | 00115 |
| type VI secretion system-associated protein TagO [Azoarcus indigens] | 00052 |
| hypothetical protein [Aromatoleum aromaticum] | 07667 |
| hypothetical protein [Azoarcus indigens] | 02933 |
| Lar family restriction alleviation protein [Azoarcus communis] | 02950 |
| DUF1566 domain-containing protein [Azoarcus indigens] | 01079 |
| acetyltransferase [Azoarcus indigens] | 00677 |
| hypothetical protein [Azoarcus indigens] | 01453 |
| VRR-NUC domain-containing protein [Azoarcus indigens] | 00051 |
| PEP-CTERM sorting domain-containing protein [Azoarcus indigens] | 09692 |
| hypothetical protein [Aromatoleum aromaticum] | 06719 |
| hypothetical protein [Azoarcus indigens] | 01456 |
| hypothetical protein [Aromatoleum aromaticum] | 07424 |
| DUF2569 family protein [Azoarcus communis] | 09090 |
| hypothetical protein [Azoarcus indigens] | 04731 |
| hypothetical protein [Azoarcus indigens] | 01078 |
| hypothetical protein [Aromatoleum aromaticum] | 06622 |
| hypothetical protein [Azoarcus indigens] | 02737 |
| hypothetical protein [Azoarcus indigens] | 09260 |
| hypothetical protein [Azoarcus indigens] | 09691 |
| hypothetical protein [Azoarcus indigens] | 00898 |
| hypothetical protein [Azoarcus communis] | 01112 |
| phage head-tail connector protein [Azoarcus indigens] | 03874 |
| hypothetical protein [Azoarcus indigens] | 01794 |
| DUF4260 domain-containing protein [Azoarcus indigens] | 07440 |
| hypothetical protein [Azoarcus indigens] | 04883 |
| hypothetical protein [Azoarcus indigens] | 08799 |
| phage antirepressor KilAC domain-containing protein [Azoarcus indigens] | 00899 |
| cytochrome P450 [Azoarcus indigens] | 03875 |
| hypothetical protein [Azoarcus communis] | 08386 |
| hypothetical protein [Azoarcus indigens] | 01454 |
| hypothetical protein [Azoarcus indigens] | 02087 |
| phosphoglycerate mutase family protein [Azoarcus indigens] | 09370 |
| hypothetical protein [Azoarcus communis] | 05549 |
| tetratricopeptide repeat-containing sulfotransferase family protein [Azoarcus indigens] | 07266 |
| hypothetical protein [Azoarcus indigens] | 00558 |
| hypothetical protein [Azoarcus communis] | 04795 |
| hypothetical protein [Azoarcus communis] | 08880 |
| hypothetical protein, partial [Azoarcus communis] | 01846 |
| transposase [Azoarcus communis] | 10340 |
| hypothetical protein [Azoarcus indigens] | 10384 |
| ArsR family transcriptional regulator [Azoarcus communis] | 10356 |
| hypothetical protein [Azoarcus indigens] | 10491 |
| hypothetical protein [Azoarcus indigens] | 08797 |
| hypothetical protein [Azoarcus indigens] | 01876 |

| | |
|---|---|
| hypothetical protein [Azoarcus indigens] | 09569 |
| Hcp family type VI secretion system effector [Azoarcus indigens] | 08847 |
| hypothetical protein [Azoarcus indigens] | 04877 |
| hypothetical protein [Azoarcus communis] | 00648 |
| autotransporter outer membrane beta-barrel domain-containing protein [Azoarcus indigens] | 00048 |
| hypothetical protein [Azoarcus indigens] | 02088 |
| MarR family transcriptional regulator [Azoarcus indigens] | 02201 |
| cytochrome o ubiquinol oxidase subunit IV [Azoarcus indigens] | 03492 |
| hypothetical protein [Azoarcus indigens] | 05131 |
| TnsA endonuclease N-terminal domain-containing protein [Azoarcus communis] | 01680 |
| hypothetical protein [Azoarcus indigens] | 08212 |
| hypothetical protein [Azoarcus indigens] | 02932 |
| genomic island protein [Azoarcus indigens] | 03228 |
| NAD-dependent epimerase/dehydratase family protein [Aromatoleum aromaticum] | 00756 |
| hypothetical protein [Azoarcus indigens] | 03489 |
| recombination protein NinB [Azoarcus indigens] | 09104 |
| 5'-3'-deoxyribonucleotidase [Azoarcus indigens] | 08796 |
| hypothetical protein [Azoarcus indigens] | 03394 |
| delta(1)-pyrroline-2-carboxylate reductase family protein [Azoarcus communis] | 04761 |
| carboxylesterase family protein [Azoarcus indigens] | 06144 |
| hypothetical protein [Azoarcus indigens] | 07260 |
| MFS transporter [Azoarcus indigens] | 10150 |
| MULTISPECIES: Arc family DNA binding domain-containing protein [Azoarcus] | 07835 |
| MULTISPECIES: SPFH domain-containing protein [Azoarcus] | 09474 |
| phage tail sheath family protein [Azoarcus communis] | 05237 |
| hypothetical protein [Azoarcus communis] | 01711 |
| hypothetical protein [Azoarcus indigens] | 03707 |
| hypothetical protein [Azoarcus indigens] | 01076 |
| hypothetical protein [Azoarcus indigens] | 08507 |
| reverse transcriptase family protein [Azoarcus indigens] | 01612 |
| ThiF family adenylyltransferase [Azoarcus communis] | 01421 |
| hypothetical protein [Aromatoleum aromaticum] | 03735 |
| YfdQ family protein [Aromatoleum aromaticum] | 07846 |
| metallophosphoesterase [Azoarcus indigens] | 05550 |
| hypothetical protein [Azoarcus indigens] | 03022 |
| hypothetical protein [Azoarcus indigens] | 06039 |
| DUF2220 family protein [Azoarcus indigens] | 00363 |
| DUF5716 family protein [Azoarcus indigens] | 08131 |
| hypothetical protein [Aromatoleum aromaticum] | 03690 |
| MULTISPECIES: hypothetical protein [Rhodocyclales] | 09512 |
| YbjN domain-containing protein [Azoarcus indigens] | 07838 |
| hypothetical protein [Aromatoleum aromaticum] | 02172 |
| SAVED domain-containing protein [Aromatoleum aromaticum] | 08075 |
| hypothetical protein [Azoarcus indigens] | 03497 |
| MULTISPECIES: hypothetical protein [Rhodocyclales] | 04854 |
| hypothetical protein [Azoarcus communis] | 04472 |
| hypothetical protein [Azoarcus communis] | 09465 |
| hypothetical protein [Azoarcus indigens] | 04781 |
| hypothetical protein [Azoarcus indigens] | 07488 |
| zinc ribbon domain-containing protein [Aromatoleum aromaticum] | 07720 |
| hypothetical protein [Azoarcus indigens] | 02397 |
| TIGR03790 family protein [Azoarcus communis] | 10354 |
| 3'-5' exoribonuclease [Azoarcus communis] | 01901 |
| EAL domain-containing protein [Propionivibrio dicarboxylicus] | 02703 |
| hypothetical protein [Azoarcus communis] | 04468 |
| DNA phosphorothioation system sulfurtransferase DndC [Thauera linaloolentis] | 01009 |
| hypothetical protein [Azoarcus indigens] | 09910 |
| hypothetical protein [Azoarcus communis] | 00213 |
| helix-turn-helix domain-containing protein [Azoarcus indigens] | 00443 |

| | |
|---|---|
| toprim domain-containing protein [Azoarcus communis] | 07254 |
| helix-turn-helix domain-containing protein [Propionivibrio dicarboxylicus] | 08586 |
| DNA-binding protein [Azoarcus communis] | 09928 |
| SEL1-like repeat protein [Azoarcus communis] | 07500 |
| DUF1656 domain-containing protein [Azoarcus indigens] | 07842 |
| hypothetical protein [Azoarcus indigens] | 03398 |
| S1/P1 nuclease [Propionivibrio dicarboxylicus] | 03436 |
| hypothetical protein [Aromatoleum aromaticum] | 03901 |
| hypothetical protein [Aromatoleum aromaticum] | 09803 |
| serine protease [Azoarcus communis] | 06642 |
| hypothetical protein [Azoarcus indigens] | 02091 |
| GYF domain-containing protein [Azoarcus indigens] | 00902 |
| FUSC family protein [Azoarcus indigens] | 08900 |
| glycosyltransferase [Azoarcus communis] | 06469 |
| STAS domain-containing protein [Azoarcus indigens] | 03397 |
| energy transducer TonB [Thauera linaloolentis] | 06875 |
| hypothetical protein [Azoarcus communis] | 02372 |
| helix-hairpin-helix domain-containing protein [Azoarcus communis] | 06948 |
| hypothetical protein [Azoarcus communis] | 07822 |
| peroxidase [Aromatoleum diolicum] | 08291 |
| Hpt domain-containing protein [Aromatoleum diolicum] | 07251 |
| hypothetical protein [Thauera aromatica] | 04807 |
| DUF262 domain-containing protein [Aromatoleum aromaticum] | 09383 |
| RelA/SpoT domain-containing protein [Azoarcus rhizosphaerae] | 09123 |
| hypothetical protein [Thauera aromatica] | 08770 |
| hypothetical protein [Thauera aromatica] | 02419 |
| LuxR C-terminal-related transcriptional regulator [Aromatoleum toluvorans] | 05798 |
| hypothetical protein [Thauera aromatica] | 09894 |
| L-2-amino-thiazoline-4-carboxylic acid hydrolase [Azoarcus rhizosphaerae] | 01086 |
| DUF3465 domain-containing protein [Aromatoleum diolicum] | 03215 |
| hypothetical protein [Thauera aromatica] | 04183 |
| hypothetical protein [Thauera linaloolentis] | 07156 |
| hypothetical protein [Thauera linaloolentis] | 05426 |
| hypothetical protein [Aromatoleum diolicum] | 05966 |
| hypothetical protein [Thauera aromatica] | 06316 |
| hypothetical protein [Thauera aromatica] | 01672 |
| hypothetical protein [Aromatoleum diolicum] | 04495 |
| ATP-binding protein [Aromatoleum diolicum] | 00643 |
| hypothetical protein [Thauera aromatica] | 03366 |
| YdeI/OmpD-associated family protein [Azoarcus rhizosphaerae] | 00575 |
| hypothetical protein [Thauera linaloolentis] | 01008 |
| hypothetical protein [Thauera aromatica] | 06019 |
| PTS sugar transporter subunit IIA [Azoarcus rhizosphaerae] | 01093 |
| hypothetical protein [Thauera aromatica] | 07858 |
| class I SAM-dependent methyltransferase [Azoarcus rhizosphaerae] | 03987 |
| hypothetical protein [Thauera aromatica] | 01730 |
| hypothetical protein [Aromatoleum toluvorans] | 09116 |
| thioesterase family protein, partial [Aromatoleum toluvorans] | 08599 |
| DUF4242 domain-containing protein [Aromatoleum diolicum] | 09682 |
| hypothetical protein [Aromatoleum toluvorans] | 04851 |
| hypothetical protein [Aromatoleum toluvorans] | 05226 |
| hypothetical protein [Thauera linaloolentis] | 00308 |
| hypothetical protein [Thauera aromatica] | 04874 |
| hypothetical protein [Aromatoleum toluvorans] | 06937 |
| hypothetical protein, partial [Aromatoleum toluvorans] | 05816 |
| helix-turn-helix domain-containing protein [Aromatoleum diolicum] | 05542 |
| pilin [Aromatoleum aromaticum] | 02627 |
| PriCT-2 domain-containing protein [Aromatoleum diolicum] | 04500 |
| hypothetical protein [Aromatoleum aromaticum] | 00955 |

| | |
|---|---|
| hypothetical protein [Aromatoleum aromaticum] | 03747 |
| McrC family protein [Thauera aromatica] | 07474 |
| hypothetical protein [Thauera linaloolentis] | 08462 |
| hypothetical protein [Thauera linaloolentis] | 05393 |
| hypothetical protein [Thauera linaloolentis] | 04665 |
| Fic family protein [Thauera aromatica] | 02143 |
| FAD-binding oxidoreductase [Aromatoleum diolicum] | 05810 |
| transposase [Thauera aromatica] | 04752 |
| hypothetical protein [Aromatoleum diolicum] | 02839 |
| PD-(D/E)XK motif protein [Thauera linaloolentis] | 10375 |
| hypothetical protein [Aromatoleum toluvorans] | 03475 |
| hypothetical protein [Thauera linaloolentis] | 00301 |
| ribbon-helix-helix domain-containing protein [Thauera aromatica] | 08180 |
| hypothetical protein [Aromatoleum toluvorans] | 07585 |
| hypothetical protein [Aromatoleum diolicum] | 09073 |
| transposase [Aromatoleum aromaticum] | 10282 |
| hypothetical protein [Aromatoleum toluvorans] | 08815 |
| XRE family transcriptional regulator [Aromatoleum toluvorans] | 05598 |
| type II restriction endonuclease [Aromatoleum toluvorans] | 08598 |
| hypothetical protein [Aromatoleum diolicum] | 06490 |
| molybdopterin-dependent oxidoreductase, partial [Aromatoleum toluvorans] | 08289 |
| hypothetical protein [Aromatoleum toluvorans] | 05824 |
| hypothetical protein [Thauera linaloolentis] | 03239 |
| hypothetical protein [Aromatoleum diolicum] | 05820 |
| hypothetical protein [Aromatoleum toluvorans] | 00857 |
| pilin [Aromatoleum aromaticum] | 08420 |
| DDE-type integrase/transposase/recombinase [Aromatoleum diolicum] | 09071 |
| DNA (cytosine-5-)-methyltransferase [Thauera linaloolentis] | 07768 |
| AAA family ATPase [Aromatoleum diolicum] | 00211 |
| hypothetical protein [Aromatoleum toluvorans] | 09052 |
| hypothetical protein [Thauera aromatica] | 03203 |
| hypothetical protein [Azoarcus rhizosphaerae] | 02945 |
| MULTISPECIES: ribbon-helix-helix protein, CopG family [unclassified Azoarcus] | 05260 |
| ThiF family adenylyltransferase [Aromatoleum aromaticum] | 03415 |
| DUF4214 domain-containing protein [Thauera terpenica] | 08485 |
| ATP-binding protein, partial [Thauera linaloolentis] | 04023 |
| hypothetical protein [Thauera linaloolentis] | 01917 |
| hypothetical protein [Thauera linaloolentis] | 02040 |
| hypothetical protein [Aromatoleum aromaticum] | 08076 |
| hypothetical protein [Thauera aromatica] | 05668 |
| DUF4160 domain-containing protein [Azoarcus rhizosphaerae] | 00578 |
| hypothetical protein [Azoarcus rhizosphaerae] | 05144 |
| PEP-CTERM sorting domain-containing protein [Thauera linaloolentis] | 05987 |
| hypothetical protein [Aromatoleum aromaticum] | 10074 |
| NUMOD4 motif-containing HNH endonuclease [Thauera aromatica] | 06905 |
| XapX domain-containing protein [Azoarcus rhizosphaerae] | 01103 |
| glycosyltransferase family 4 protein [Thauera terpenica] | 01633 |
| hypothetical protein [Azoarcus rhizosphaerae] | 01089 |
| hypothetical protein [Thauera linaloolentis] | 09184 |
| hypothetical protein [Thauera linaloolentis] | 05392 |
| hypothetical protein [Aromatoleum aromaticum] | 07122 |
| GIY-YIG nuclease family protein [Thauera terpenica] | 03194 |
| chemotaxis protein CheW [Thauera terpenica] | 06146 |
| DUF2528 family protein [Azoarcus rhizosphaerae] | 09918 |
| DUF4325 domain-containing protein [Thauera aromatica] | 07810 |
| hypothetical protein [Thauera linaloolentis] | 00666 |
| hypothetical protein [Azoarcus rhizosphaerae] | 05731 |
| PRTRC system protein C [Aromatoleum aromaticum] | 03454 |
| hypothetical protein [Thauera linaloolentis] | 08746 |

| | |
|---|---|
| hypothetical protein [Azoarcus rhizosphaerae] | 02298 |
| ATP-binding protein [Azoarcus rhizosphaerae] | 08875 |
| winged helix-turn-helix domain-containing protein [Azoarcus rhizosphaerae] | 00916 |
| hypothetical protein [Aromatoleum aromaticum] | 08419 |
| hypothetical protein [Aromatoleum aromaticum] | 03137 |
| hypothetical protein [Azoarcus rhizosphaerae] | 07494 |
| hypothetical protein [Azoarcus rhizosphaerae] | 05147 |
| MULTISPECIES: EreA family erythromycin esterase [Proteobacteria] | 03659 |
| IS1634 family transposase, partial [Thauera terpenica] | 08181 |
| hypothetical protein [Azoarcus rhizosphaerae] | 09149 |
| MULTISPECIES: hypothetical protein [Proteobacteria] | 05428 |
| heme ABC transporter permease [Azoarcus rhizosphaerae] | 00579 |
| hypothetical protein [Aromatoleum aromaticum] | 05950 |
| hypothetical protein [Azoarcus rhizosphaerae] | 09148 |
| hypothetical protein [Azoarcus rhizosphaerae] | 06053 |
| hypothetical protein [Aromatoleum aromaticum] | 02170 |
| hypothetical protein [Aromatoleum aromaticum] | 08685 |
| hypothetical protein [Thauera terpenica] | 09418 |
| MULTISPECIES: hypothetical protein [Proteobacteria] | 00528 |
| MULTISPECIES: tetracycline resistance transcriptional repressor TetR(A) [Bacteria] | 01010 |
| type II toxin-antitoxin system MqsA family antitoxin [Thauera terpenica] | 02578 |
| hypothetical protein [Thauera aromatica] | 08768 |
| hypothetical protein [Thauera aromatica] | 07812 |
| PilX N-terminal domain-containing pilus assembly protein [Azoarcus rhizosphaerae] | 00325 |
| hypothetical protein [Azoarcus rhizosphaerae] | 00577 |
| four helix bundle protein [Thauera aromatica] | 01702 |
| hypothetical protein [Azoarcus rhizosphaerae] | 05520 |
| DUF99 family protein [Aromatoleum diolicum] | 09155 |
| YCF48-related protein, partial [Aromatoleum diolicum] | 01790 |
| hypothetical protein [Thauera linaloolentis] | 09914 |
| hypothetical protein [Azoarcus rhizosphaerae] | 03893 |
| hypothetical protein [Azoarcus rhizosphaerae] | 06592 |
| hypothetical protein [Aromatoleum diolicum] | 00660 |
| hypothetical protein [Azoarcus rhizosphaerae] | 04400 |
| hypothetical protein [Aromatoleum diolicum] | 08350 |
| DUF6475 domain-containing protein [Thauera aromatica] | 02687 |
| DMT family transporter [Thauera linaloolentis] | 06097 |
| hypothetical protein, partial [Thauera terpenica] | 10320 |
| hypothetical protein [Aromatoleum aromaticum] | 02520 |
| hypothetical protein [Aromatoleum aromaticum] | 08077 |
| hypothetical protein [Aromatoleum aromaticum] | 05057 |
| Ig-like domain-containing protein, partial [Thauera terpenica] | 06307 |
| hypothetical protein [Thauera terpenica] | 06979 |
| hypothetical protein [Azoarcus rhizosphaerae] | 10589 |
| DUF1442 domain-containing protein [Azoarcus rhizosphaerae] | 00064 |
| hypothetical protein [Azoarcus rhizosphaerae] | 06440 |
| ArsR family transcriptional regulator [Aromatoleum aromaticum] | 01658 |
| hypothetical protein [Thauera terpenica] | 03794 |
| class I SAM-dependent methyltransferase [Aromatoleum aromaticum] | 07800 |
| hypothetical protein [Aromatoleum aromaticum] | 07376 |
| hypothetical protein [Aromatoleum aromaticum] | 09382 |
| hypothetical protein [Thauera terpenica] | 05387 |
| hypothetical protein [Aromatoleum aromaticum] | 03287 |
| hypothetical protein [Aromatoleum aromaticum] | 00105 |
| DUF5677 domain-containing protein [Azoarcus rhizosphaerae] | 00540 |
| type II toxin-antitoxin system prevent-host-death family antitoxin [Thauera aromatica] | 08353 |
| FAD-dependent oxidoreductase [Aromatoleum aromaticum] | 07728 |
| hypothetical protein [Aromatoleum diolicum] | 09992 |
| hypothetical protein [Rhodocyclus purpureus] | 02977 |

| | |
|---|---|
| CcdB family protein [Thauera linaloolentis] | 01699 |
| AAA family ATPase [Rhodocyclus purpureus] | 05117 |
| MULTISPECIES: hypothetical protein [Proteobacteria] | 09913 |
| MULTISPECIES: helix-turn-helix domain-containing protein [Proteobacteria] | 07125 |
| diguanylate cyclase, partial [Aromatoleum diolicum] | 02195 |
| hypothetical protein [Thauera linaloolentis] | 08734 |
| M10 family metallopeptidase C-terminal domain-containing protein [Azoarcus sp. Aa7] | 01544 |
| CpaD family pilus assembly lipoprotein [Thauera linaloolentis] | 08196 |
| mobile mystery protein A [Azoarcus sp. Aa7] | 01929 |
| hypothetical protein [Azoarcus sp. Aa7] | 02190 |
| allantoin permease [Azoarcus sp. Aa7] | 06182 |
| NB-ARC domain-containing protein [Thauera aminoaromatica] | 03623 |
| hypothetical protein [Azoarcus sp. Aa7] | 08635 |
| phosphatidylglycerol lysyltransferase domain-containing protein [Thauera linaloolentis] | 06681 |
| cold shock domain-containing protein [Azoarcus sp. Aa7] | 04052 |
| hypothetical protein [Azoarcus sp. Aa7] | 08349 |
| hypothetical protein [Azoarcus sp. Aa7] | 02388 |
| Flp family type IVb pilin [Thauera linaloolentis] | 09046 |
| hypothetical protein [Azoarcus sp. Aa7] | 08524 |
| Ig-like domain-containing protein [Azoarcus sp. Aa7] | 09668 |
| hypothetical protein [Thauera linaloolentis] | 04339 |
| MULTISPECIES: glycosyltransferase family 39 protein [Thauera] | 08148 |
| hypothetical protein [Thauera linaloolentis] | 09718 |
| DUF1090 domain-containing protein [Thauera linaloolentis] | 07154 |
| AAA family ATPase, partial [Thauera aminoaromatica] | 04305 |
| PEP-utilizing enzyme, partial [Thauera aminoaromatica] | 03757 |
| MULTISPECIES: hypothetical protein [Thauera] | 10288 |
| FAD-dependent oxidoreductase [Aromatoleum diolicum] | 02726 |
| serine/threonine protein kinase [Thauera aminoaromatica] | 04626 |
| hypothetical protein [Rhodocyclus purpureus] | 04572 |
| MULTISPECIES: hypothetical protein [Thauera] | 03754 |
| MULTISPECIES: YbjN domain-containing protein [Thauera] | 04755 |
| MULTISPECIES: DUF305 domain-containing protein [Betaproteobacteria] | 08425 |
| DUF2934 domain-containing protein [Rhodocyclus purpureus] | 03153 |
| hypothetical protein [Aromatoleum diolicum] | 00669 |
| FAD/NAD(P)-binding protein [Thauera aminoaromatica] | 00470 |
| hypothetical protein [Thauera linaloolentis] | 00507 |
| hypothetical protein [Thauera linaloolentis] | 02047 |
| hypothetical protein [Thauera aminoaromatica] | 00876 |
| hypothetical protein [Aromatoleum diolicum] | 02198 |
| MULTISPECIES: hypothetical protein [Thauera] | 05346 |
| outer membrane beta-barrel protein [Thauera aminoaromatica] | 00517 |
| hypothetical protein [Thauera linaloolentis] | 10292 |
| PEP-CTERM sorting domain-containing protein [Thauera linaloolentis] | 08744 |
| aminotransferase class IV, partial [Thauera aminoaromatica] | 05667 |
| DUF4857 domain-containing protein [Thauera linaloolentis] | 00574 |
| DUF2628 domain-containing protein [Azoarcus indigens] | 01489 |
| hypothetical protein [Azoarcus sp. Aa7] | 04253 |
| hypothetical protein [Azoarcus sp. Aa7] | 01646 |
| hypothetical protein [Thauera linaloolentis] | 07169 |
| HNH endonuclease [Azoarcus sp. Aa7] | 07312 |
| hypothetical protein [Azoarcus sp. Aa7] | 10042 |
| hypothetical protein [Aromatoleum diolicum] | 07915 |
| hypothetical protein [Aromatoleum diolicum] | 00662 |
| HK97 gp10 family phage protein [Azoarcus sp. Aa7] | 01413 |
| hypothetical protein [Thauera linaloolentis] | 04340 |
| Fic family protein [Rhodocyclus purpureus] | 05275 |
| hypothetical protein [Aromatoleum diolicum] | 06918 |
| tetratricopeptide repeat protein [Rhodocyclus purpureus] | 08943 |

| | |
|---|---|
| hypothetical protein [Aromatoleum diolicum] | 08222 |
| HPF/RaiA family ribosome-associated protein [Rhodocyclus purpureus] | 05881 |
| TIGR03747 family integrating conjugative element membrane protein [Azoarcus communis] | 03991 |
| hypothetical protein [Thauera linaloolentis] | 00910 |
| chemotaxis protein CheW [Aromatoleum diolicum] | 05037 |
| hypothetical protein [Thauera linaloolentis] | 05263 |
| cell wall hydrolase [Azoarcus sp. Aa7] | 07991 |
| TIGR03084 family metal-binding protein [Thauera linaloolentis] | 07426 |
| tetratricopeptide repeat protein [Thauera linaloolentis] | 04394 |
| hypothetical protein [Azoarcus sp. Aa7] | 05807 |
| ankyrin repeat domain-containing protein [Azoarcus sp. Aa7] | 06066 |
| hypothetical protein [Thauera linaloolentis] | 00298 |
| hypothetical protein [Azoarcus sp. Aa7] | 07243 |
| DUF5801 domain-containing protein, partial [Azoarcus sp. Aa7] | 05587 |
| cobalamin (vitamin B12) biosynthesis CbiX protein [Thauera linaloolentis] | 00852 |
| hypothetical protein [Thauera linaloolentis] | 02078 |
| glycoside hydrolase family 57 protein [Rhodocyclus purpureus] | 01480 |
| general secretion pathway protein GspB, partial [Azoarcus sp. Aa7] | 07993 |
| GYD domain-containing protein [Aromatoleum diolicum] | 10139 |
| hypothetical protein [Azoarcus sp. Aa7] | 04925 |
| hypothetical protein [Azoarcus sp. Aa7] | 10183 |
| type II toxin-antitoxin system CcdA family antitoxin [Azoarcus sp. Aa7] | 09609 |
| DUF1579 domain-containing protein [Aromatoleum diolicum] | 09686 |
| type II toxin-antitoxin system VapC family toxin [Azoarcus sp. Aa7] | 05585 |
| Flp family type IVb pilin [Azoarcus sp. Aa7] | 09124 |
| transporter [Azoarcus sp. Aa7] | 01253 |
| 2-succinyl-5-enolpyruvyl-6-hydroxy-3-cyclohexene-1-carboxylic-acid synthase [Rhodocyclus purpureus] | 02910 |
| aldehyde dehydrogenase family protein, partial [Thauera aminoaromatica] | 06475 |
| DUF262 domain-containing protein [Rhodocyclus purpureus] | 06089 |
| ATP-binding protein [Rhodocyclus purpureus] | 06183 |
| RES family NAD+ phosphorylase [Thauera linaloolentis] | 07221 |
| type II toxin-antitoxin system RelE/ParE family toxin [Azoarcus indigens] | 04477 |
| hypothetical protein [Uliginosibacterium gangwonense] | 00438 |
| hypothetical protein [Uliginosibacterium gangwonense] | 09214 |
| insulinase family protein [Uliginosibacterium gangwonense] | 08445 |
| NYN domain-containing protein [Thauera linaloolentis] | 05467 |
| DUF4124 domain-containing protein [Uliginosibacterium gangwonense] | 00126 |
| HNH endonuclease [Aromatoleum diolicum] | 05541 |
| hypothetical protein [Uliginosibacterium gangwonense] | 05048 |
| DUF1805 domain-containing protein [Rhodocyclus purpureus] | 10049 |
| Ig-like domain-containing protein [Uliginosibacterium gangwonense] | 09029 |
| S8 family serine peptidase [Uliginosibacterium gangwonense] | 09026 |
| ABC transporter transmembrane domain-containing protein [Uliginosibacterium gangwonense] | 01621 |
| GTP-binding protein [Uliginosibacterium gangwonense] | 09408 |
| site-specific integrase [Uliginosibacterium gangwonense] | 04579 |
| hypothetical protein [Uliginosibacterium gangwonense] | 06662 |
| hypothetical protein [Uliginosibacterium gangwonense] | 09025 |
| transcriptional regulator [Uliginosibacterium gangwonense] | 05725 |
| hypothetical protein [Uliginosibacterium gangwonense] | 09838 |
| hypothetical protein [Uliginosibacterium gangwonense] | 06665 |
| DUF2220 family protein [Thauera linaloolentis] | 09854 |
| hypothetical protein [Rhodocyclus purpureus] | 07910 |
| tetratricopeptide repeat protein [Rhodocyclus purpureus] | 02943 |
| hypothetical protein [Thauera linaloolentis] | 09047 |
| hypothetical protein [Thauera linaloolentis] | 08736 |
| SAVED domain-containing protein [Thauera linaloolentis] | 05060 |
| hypothetical protein [Aromatoleum toluvorans] | 08595 |
| HAMP domain-containing histidine kinase [Aromatoleum toluvorans] | 00515 |
| cupin domain-containing protein [Rhodocyclus purpureus] | 09022 |

| | |
|---|---|
| hypothetical protein [Aromatoleum toluvorans] | 08713 |
| hypothetical protein [Aromatoleum diolicum] | 06960 |
| DMT family transporter [Aromatoleum toluvorans] | 09243 |
| MFS transporter [Uliginosibacterium gangwonense] | 10128 |
| hypothetical protein [Aromatoleum toluvorans] | 04491 |
| phosphoribosyl-AMP cyclohydrolase [Thauera linaloolentis] | 04229 |
| hypothetical protein [Aromatoleum toluvorans] | 00764 |
| HD domain-containing protein [Aromatoleum toluvorans] | 02894 |
| hypothetical protein [Thauera linaloolentis] | 08732 |
| GNAT family N-acetyltransferase [Rhodocyclus purpureus] | 04705 |
| S-layer protein [Uliginosibacterium gangwonense] | 00834 |
| hypothetical protein [Uliginosibacterium gangwonense] | 09519 |
| terminase small subunit [Uliginosibacterium gangwonense] | 09134 |
| transcriptional regulator [Uliginosibacterium gangwonense] | 09864 |
| MULTISPECIES: PD-(D/E)XK nuclease domain-containing protein [Thauera] | 01028 |
| PEP-CTERM sorting domain-containing protein [Thauera aminoaromatica] | 06081 |
| MarR family transcriptional regulator [Aromatoleum diolicum] | 08551 |
| DUF4238 domain-containing protein [Uliginosibacterium gangwonense] | 03313 |
| N-acetyltransferase [Thauera linaloolentis] | 02579 |
| DUF4145 domain-containing protein [Uliginosibacterium gangwonense] | 07404 |
| hypothetical protein [Uliginosibacterium gangwonense] | 05968 |
| hypothetical protein [Thauera linaloolentis] | 07427 |
| TROVE domain-containing protein [Uliginosibacterium gangwonense] | 09020 |
| aldehyde dehydrogenase family protein, partial [Thauera linaloolentis] | 03332 |
| LysR substrate-binding domain-containing protein [Thauera linaloolentis] | 04355 |
| hypothetical protein [Uliginosibacterium gangwonense] | 05910 |
| aromatic amino acid DMT transporter YddG [Thauera linaloolentis] | 02861 |
| hypothetical protein [Thauera linaloolentis] | 06043 |
| hypothetical protein [Thauera aminoaromatica] | 03760 |
| hypothetical protein [Thauera linaloolentis] | 07197 |
| hypothetical protein [Thauera linaloolentis] | 01004 |
| putative sulfate exporter family transporter, partial [Aromatoleum diolicum] | 08604 |
| hypothetical protein [Thauera linaloolentis] | 03897 |
| hypothetical protein [Thauera aminoaromatica] | 08884 |
| capsule biosynthesis GfcC family protein [Rhodocyclus purpureus] | 02836 |
| DUF1343 domain-containing protein [Uliginosibacterium gangwonense] | 00823 |
| hypothetical protein [Thauera linaloolentis] | 02863 |
| ABC transporter permease [Uliginosibacterium gangwonense] | 03890 |
| hypothetical protein [Thauera linaloolentis] | 03667 |
| hypothetical protein [Uliginosibacterium gangwonense] | 01409 |
| hypothetical protein [Aromatoleum diolicum] | 01866 |
| HPF/RaiA family ribosome-associated protein [Uliginosibacterium gangwonense] | 04757 |
| NADAR family protein [Uliginosibacterium gangwonense] | 09405 |
| transcriptional repressor [Thauera linaloolentis] | 03657 |
| hypothetical protein [Uliginosibacterium gangwonense] | 05680 |
| hypothetical protein [Uliginosibacterium gangwonense] | 05371 |
| transposase, partial [Uliginosibacterium gangwonense] | 00830 |
| acyl carrier protein [Uliginosibacterium gangwonense] | 06236 |
| hypothetical protein [Uliginosibacterium gangwonense] | 09146 |
| hypothetical protein [Rhodocyclus purpureus] | 06187 |
| hypothetical protein [Uliginosibacterium gangwonense] | 01087 |
| DUF58 domain-containing protein [Uliginosibacterium gangwonense] | 04317 |
| DUF2939 domain-containing protein [Uliginosibacterium gangwonense] | 05042 |
| type III secretion inner membrane ring lipoprotein SctJ [Uliginosibacterium gangwonense] | 04794 |
| hypothetical protein [Uliginosibacterium gangwonense] | 01615 |
| ATP-binding protein [Aromatoleum diolicum] | 04002 |
| type B 50S ribosomal protein L36 [Thauera linaloolentis] | 06320 |
| hypothetical protein, partial [Azoarcus communis] | 09231 |
| hypothetical protein [Aromatoleum aromaticum] | 04126 |

| | |
|---|---|
| hypothetical protein [Azoarcus sp. L1K30] | 02286 |
| transporter [Aromatoleum aromaticum] | 00415 |
| pyridoxamine 5'-phosphate oxidase family protein [Azoarcus sp. L1K30] | 02277 |
| SDR family NAD(P)-dependent oxidoreductase [Azoarcus sp. L1K30] | 07558 |
| hypothetical protein [Azoarcus sp. L1K30] | 06588 |
| HNH endonuclease [Azoarcus sp. L1K30] | 10234 |
| gluconokinase [Azoarcus sp. L1K30] | 02480 |
| hypothetical protein [Azoarcus sp. L1K30] | 03577 |
| OBAP family protein [Aromatoleum aromaticum] | 02702 |
| hypothetical protein [Aromatoleum aromaticum] | 03615 |
| tyrosine-type recombinase/integrase [Azoarcus sp. L1K30] | 01754 |
| hypothetical protein [Azoarcus sp. L1K30] | 05916 |
| hypothetical protein [Azoarcus sp. L1K30] | 03579 |
| hypothetical protein [Azoarcus sp. L1K30] | 00056 |
| asparaginase [Aromatoleum aromaticum] | 02356 |
| hypothetical protein [Azoarcus sp. L1K30] | 09341 |
| LysE family transporter [Azoarcus sp. L1K30] | 01546 |
| 4Fe-4S dicluster domain-containing protein [Azoarcus sp. L1K30] | 08177 |
| hypothetical protein [Azoarcus sp. L1K30] | 04261 |
| hypothetical protein [Azoarcus sp. L1K30] | 05914 |
| hypothetical protein [Azoarcus sp. L1K30] | 10586 |
| methyltransferase domain-containing protein [Azoarcus sp. L1K30] | 09761 |
| ImmA/IrrE family metallo-endopeptidase [Thauera humireducens] | 07432 |
| serine protease [Thauera humireducens] | 04668 |
| hypothetical protein [Thauera humireducens] | 04205 |
| radical SAM protein [Thauera humireducens] | 05710 |
| hypothetical protein [Thauera humireducens] | 01045 |
| hypothetical protein [Thauera humireducens] | 07178 |
| hypothetical protein [Thauera humireducens] | 05092 |
| hypothetical protein [Thauera humireducens] | 06686 |
| glutaredoxin family protein [Thauera humireducens] | 06238 |
| DUF4433 domain-containing protein [Thauera humireducens] | 10577 |
| hypothetical protein [Azoarcus sp. L1K30] | 03261 |
| hypothetical protein [Azoarcus sp. L1K30] | 06766 |
| MULTISPECIES: antitoxin VbhA family protein [Betaproteobacteria] | 10140 |
| KTSC domain-containing protein [Azoarcus nasutitermitis] | 06990 |
| succinate dehydrogenase [Azoarcus nasutitermitis] | 00906 |
| AAA family ATPase [Azoarcus nasutitermitis] | 03237 |
| GntP family permease [Azoarcus nasutitermitis] | 04786 |
| VCBS repeat-containing protein [Azoarcus nasutitermitis] | 07555 |
| DEAD/DEAH box helicase [Azoarcus nasutitermitis] | 04396 |
| hypothetical protein [Azoarcus nasutitermitis] | 09916 |
| hypothetical protein [Azoarcus nasutitermitis] | 06888 |
| vitamin K epoxide reductase family protein [Aromatoleum aromaticum] | 08999 |
| MBL fold metallo-hydrolase [Niveibacterium umoris] | 04506 |
| hypothetical protein [Niveibacterium umoris] | 00811 |
| N-acetylmuramoyl-L-alanine amidase [Niveibacterium umoris] | 09291 |
| MULTISPECIES: hypothetical protein [Betaproteobacteria] | 02735 |
| MULTISPECIES: hypothetical protein [Betaproteobacteria] | 09252 |
| MULTISPECIES: hypothetical protein [Betaproteobacteria] | 02360 |
| CAP domain-containing protein [Aromatoleum aromaticum] | 01588 |
| helix-turn-helix domain-containing protein [Azoarcus nasutitermitis] | 06083 |
| hypothetical protein [Azoarcus nasutitermitis] | 08576 |
| MULTISPECIES: hypothetical protein [Proteobacteria] | 10433 |
| hypothetical protein [Thauera lacus] | 06882 |
| SOS response-associated peptidase [Thauera lacus] | 02071 |
| hypothetical protein [Thauera lacus] | 00554 |
| hypothetical protein [Thauera lacus] | 04471 |
| hypothetical protein [Thauera lacus] | 10357 |

| | |
|---|---|
| hypothetical protein [Thauera lacus] | 08176 |
| transposase [Thauera lacus] | 09183 |
| thioredoxin family protein [Thauera lacus] | 00570 |
| hypothetical protein [Thauera lacus] | 00571 |
| type II toxin-antitoxin system prevent-host-death family antitoxin [Thauera lacus] | 07490 |
| hypothetical protein [Azoarcus sp. L1K30] | 07685 |
| c-type cytochrome [Aromatoleum aromaticum] | 09969 |
| hypothetical protein [Aromatoleum aromaticum] | 08868 |
| type II secretion system protein GspK [Thauera humireducens] | 08763 |
| YkgJ family cysteine cluster protein [Thauera humireducens] | 08455 |
| hypothetical protein [Thauera humireducens] | 07798 |
| hypothetical protein [Azoarcus communis] | 08125 |
| hypothetical protein [Azoarcus communis] | 06010 |
| hypothetical protein [Azoarcus communis] | 00136 |
| DUF3396 domain-containing protein [Azoarcus communis] | 02285 |
| DUF3147 family protein [Aromatoleum aromaticum] | 09377 |
| hypothetical protein [Azoarcus communis] | 04763 |
| metallophosphoesterase [Azoarcus communis] | 07796 |
| hypothetical protein [Azoarcus communis] | 08672 |
| flagellar basal body-associated FliL family protein [Thauera humireducens] | 03182 |
| hypothetical protein [Azoarcus communis] | 02916 |
| hypothetical protein [Azoarcus communis] | 00769 |
| oligosaccharide flippase family protein [Azoarcus communis] | 01067 |
| AIPR family protein [Aromatoleum aromaticum] | 03971 |
| hypothetical protein [Aromatoleum aromaticum] | 08069 |
| hypothetical protein [Azoarcus communis] | 05216 |
| integrase core domain-containing protein [Thauera hydrothermalis] | 08502 |
| glycosyl hydrolase family 8 [Thauera hydrothermalis] | 00355 |
| UDP-forming cellulose synthase catalytic subunit [Thauera hydrothermalis] | 03223 |
| cellulose biosynthesis cyclic di-GMP-binding regulatory protein BcsB [Thauera hydrothermalis] | 07465 |
| TetR/AcrR family transcriptional regulator [Thauera hydrothermalis] | 06364 |
| phage holin, lambda family [Aromatoleum aromaticum] | 07118 |
| dihydroxy-acid dehydratase [Aromatoleum pumilum] | 04211 |
| amidohydrolase family protein [Aromatoleum pumilum] | 05100 |
| class I SAM-dependent methyltransferase [Aromatoleum pumilum] | 07454 |
| hypothetical protein [Aromatoleum pumilum] | 00609 |
| hypothetical protein [Aromatoleum aromaticum] | 02516 |
| glycosyltransferase family 4 protein [Aromatoleum pumilum] | 09901 |
| transcriptional regulator [Aromatoleum pumilum] | 10089 |
| glycosyltransferase [Aromatoleum pumilum] | 06589 |
| hypothetical protein [Aromatoleum aromaticum] | 00644 |
| kinase [Aromatoleum aromaticum] | 05971 |
| hypothetical protein [Aromatoleum pumilum] | 03204 |
| TIGR04255 family protein [Aromatoleum aromaticum] | 07722 |
| hypothetical protein [Azoarcus communis] | 07078 |
| oligosaccharide flippase family protein [Aromatoleum aromaticum] | 04125 |
| hypothetical protein [Thauera humireducens] | 08158 |
| class I SAM-dependent methyltransferase [Thauera humireducens] | 01966 |
| sulfotransferase [Thauera humireducens] | 09859 |
| glycosyltransferase [Thauera humireducens] | 07159 |
| hypothetical protein [Thauera humireducens] | 09423 |
| hypothetical protein [Thauera humireducens] | 03352 |
| IS630-like element ISAzo9 family transposase [Aromatoleum aromaticum] | 09376 |
| hypothetical protein [Thauera humireducens] | 08127 |
| carboxymuconolactone decarboxylase family protein [Aromatoleum aromaticum] | 07721 |
| hypothetical protein [Thauera humireducens] | 06112 |
| hypothetical protein [Thauera humireducens] | 04962 |
| hypothetical protein [Thauera humireducens] | 09050 |
| hypothetical protein [Thauera humireducens] | 09857 |

| | |
|---|---|
| hypothetical protein [Thauera humireducens] | 08473 |
| hypothetical protein [Thauera humireducens] | 07799 |
| hypothetical protein [Thauera humireducens] | 03202 |
| hypothetical protein, partial [Azoarcus communis] | 10369 |
| hypothetical protein [Thauera humireducens] | 01971 |
| hypothetical protein [Thauera humireducens] | 02608 |
| hypothetical protein [Thauera humireducens] | 00850 |
| hypothetical protein [Thauera humireducens] | 07498 |
| hypothetical protein [Thauera humireducens] | 05395 |
| MULTISPECIES: hypothetical protein [Thauera] | 04341 |
| MULTISPECIES: NIPSNAP family protein [Thauera] | 03573 |
| MULTISPECIES: toll/interleukin-1 receptor domain-containing protein [Thauera] | 06339 |
| MULTISPECIES: pyrroloquinoline quinone precursor peptide PqqA [Thauera] | 03201 |
| MULTISPECIES: DUF3280 domain-containing protein [Thauera] | 03200 |
| MULTISPECIES: hypothetical protein [Thauera] | 01638 |
| hypothetical protein [Aromatoleum aromaticum] | 01175 |
| hypothetical protein [Azoarcus communis] | 06389 |
| hypothetical protein, partial [Azoarcus communis] | 00352 |
| MULTISPECIES: hypothetical protein [Betaproteobacteria] | 02705 |
| MULTISPECIES: toprim domain-containing protein [Betaproteobacteria] | 01785 |
| DUF2304 domain-containing protein [Aromatoleum aromaticum] | 09380 |
| hypothetical protein [Azoarcus olearius] | 02521 |
| hypothetical protein [Azoarcus sp. KH32C] | 06991 |
| hypothetical protein [Azoarcus sp. KH32C] | 05366 |
| hypothetical protein [Azoarcus sp. KH32C] | 02840 |
| HNH endonuclease [Azoarcus sp. KH32C] | 06289 |
| hypothetical protein [Azoarcus olearius] | 06985 |
| hypothetical protein [Azoarcus olearius] | 10326 |
| hypothetical protein [Azoarcus olearius] | 00956 |
| hypothetical protein [Azoarcus olearius] | 07378 |
| ligase-associated DNA damage response endonuclease PdeM [Azoarcus olearius] | 07729 |
| hypothetical protein [Azoarcus olearius] | 06270 |
| HPr-rel-A system PqqD family peptide chaperone [Azoarcus olearius] | 07783 |
| hypothetical protein [Azoarcus olearius] | 02523 |
| hypothetical protein [Azoarcus olearius] | 06271 |
| BON domain-containing protein [Azoarcus olearius] | 09386 |
| hypothetical protein [Azoarcus sp. KH32C] | 06651 |
| hypothetical protein [Azoarcus sp. KH32C] | 09541 |
| hypothetical protein [Azoarcus sp. KH32C] | 10073 |
| hypothetical protein [Azoarcus sp. KH32C] | 00273 |
| PepSY domain-containing protein [Azoarcus sp. KH32C] | 03635 |
| PilZ domain-containing protein [Azoarcus sp. KH32C] | 06812 |
| hypothetical protein [Azoarcus sp. KH32C] | 00272 |
| hypothetical protein [Azoarcus sp. KH32C] | 02544 |
| DUF4113 domain-containing protein [Azoarcus communis] | 03846 |
| OsmC family protein [Azoarcus communis] | 02468 |
| hypothetical protein [Azoarcus communis] | 06889 |
| hypothetical protein [Azoarcus communis] | 09998 |
| hypothetical protein [Azoarcus communis] | 00026 |
| autotransporter outer membrane beta-barrel domain-containing protein [Azoarcus communis] | 01379 |
| hypothetical protein [Aromatoleum aromaticum] | 06460 |
| RNA-guided endonuclease TnpB family protein [Thauera propionica] | 02889 |
| MULTISPECIES: hypothetical protein [Proteobacteria] | 03686 |
| hypothetical protein [Azoarcus olearius] | 07784 |
| hypothetical protein [Azoarcus olearius] | 08141 |
| MULTISPECIES: hypothetical protein [Betaproteobacteria] | 08894 |
| acyl-CoA dehydrogenase family protein, partial [Aromatoleum buckelii] | 01449 |
| hypothetical protein [Aromatoleum buckelii] | 10451 |
| hypothetical protein [Aromatoleum buckelii] | 07218 |

| | |
|---|---|
| hypothetical protein [Aromatoleum buckelii] | 06758 |
| hypothetical protein [Aromatoleum buckelii] | 07909 |
| hypothetical protein [Aromatoleum buckelii] | 05215 |
| site-specific DNA-methyltransferase [Aromatoleum buckelii] | 09142 |
| hypothetical protein, partial [Aromatoleum buckelii] | 02173 |
| hypothetical protein [Azoarcus olearius] | 02525 |
| hypothetical protein [Aromatoleum bremense] | 07653 |
| PIN domain-containing protein [Aromatoleum bremense] | 05186 |
| GIY-YIG nuclease family protein [Aromatoleum bremense] | 04821 |
| hypothetical protein [Aromatoleum bremense] | 00762 |
| hypothetical protein [Zoogloea oleivorans] | 03485 |
| TolC family protein, partial [Zoogloea oleivorans] | 07257 |
| hypothetical protein [Azoarcus olearius] | 10286 |
| hypothetical protein [Azoarcus olearius] | 07785 |
| phosphonopyruvate decarboxylase [Azoarcus olearius] | 08080 |
| hypothetical protein [Azoarcus olearius] | 06273 |
| hypothetical protein [Azoarcus olearius] | 03289 |
| DUF2795 domain-containing protein [Azoarcus olearius] | 05083 |
| sel1 repeat family protein [Azoarcus olearius] | 01030 |
| hypothetical protein [Azoarcus olearius] | 03622 |
| lipase family protein [Azoarcus olearius] | 03291 |
| glycosyltransferase [Azoarcus olearius] | 00309 |
| hypothetical protein [Azoarcus olearius] | 05665 |
| hypothetical protein [Azoarcus olearius] | 02052 |
| hypothetical protein [Azoarcus olearius] | 08422 |
| hypothetical protein [Azoarcus olearius] | 03621 |
| hypothetical protein [Azoarcus olearius] | 04621 |
| hypothetical protein [Azoarcus olearius] | 01029 |
| glycosyltransferase family 39 protein [Azoarcus olearius] | 05080 |
| hypothetical protein [Thauera propionica] | 03766 |
| hypothetical protein [Thauera propionica] | 06011 |
| hypothetical protein [Thauera propionica] | 08708 |
| hypothetical protein [Aromatoleum evansii] | 07656 |
| GIY-YIG nuclease family protein [Aromatoleum evansii] | 04850 |
| DEAD/DEAH box helicase family protein [Aromatoleum evansii] | 06962 |
| hypothetical protein [Aromatoleum evansii] | 00941 |
| hypothetical protein [Aromatoleum evansii] | 02138 |
| DUF2271 domain-containing protein [Aromatoleum aromaticum] | 02512 |
| hypothetical protein [Aromatoleum evansii] | 09221 |
| hypothetical protein [Aromatoleum evansii] | 07916 |
| hypothetical protein [Thauera propionica] | 03684 |
| hypothetical protein [Aromatoleum aromaticum] | 03568 |
| hypothetical protein [Aromatoleum evansii] | 07917 |
| glycosyltransferase family 4 protein, partial [Aromatoleum evansii] | 03243 |
| hypothetical protein [Aromatoleum evansii] | 00111 |
| hypothetical protein, partial [Aromatoleum evansii] | 03002 |
| hypothetical protein [Aromatoleum evansii] | 06468 |
| hypothetical protein [Aromatoleum evansii] | 02823 |
| hypothetical protein [Aromatoleum evansii] | 10480 |
| zeta toxin family protein [Aromatoleum evansii] | 05224 |
| winged helix-turn-helix domain-containing protein, partial [Aromatoleum evansii] | 01755 |
| CopK family periplasmic copper-binding protein [Aromatoleum evansii] | 00668 |
| hypothetical protein [Aromatoleum aromaticum] | 04124 |
| lasso peptide biosynthesis protein [Aromatoleum evansii] | 07505 |
| SoxR reducing system RseC family protein, partial [Aromatoleum evansii] | 04845 |
| hypothetical protein [Aromatoleum evansii] | 02678 |
| DUF1643 domain-containing protein [Aromatoleum evansii] | 01659 |
| DNA-binding protein [Aromatoleum evansii] | 05793 |
| DUF559 domain-containing protein [Aromatoleum evansii] | 03453 |

| | |
|---|---|
| hypothetical protein [Aromatoleum evansii] | 00261 |
| DUF3560 domain-containing protein [Aromatoleum evansii] | 08873 |
| MULTISPECIES: hypothetical protein [Betaproteobacteria] | 02196 |
| MULTISPECIES: hypothetical protein [Betaproteobacteria] | 01868 |
| hypothetical protein [Aromatoleum evansii] | 07252 |
| hypothetical protein, partial [Aromatoleum evansii] | 05227 |
| hypothetical protein [Aromatoleum evansii] | 01869 |
| hypothetical protein [Aromatoleum evansii] | 03976 |
| hypothetical protein [Azoarcus sp. DD4] | 02879 |
| hypothetical protein [Thauera propionica] | 09823 |
| hypothetical protein [Thauera propionica] | 06091 |
| hypothetical protein [Thauera propionica] | 06866 |
| sulfotransferase [Thauera propionica] | 01656 |
| DEAD/DEAH box helicase [Azoarcus sp. DD4] | 04012 |
| SAVED domain-containing protein [Azoarcus sp. DD4] | 00379 |
| hypothetical protein [Azoarcus sp. DD4] | 02053 |
| hypothetical protein [Azoarcus sp. DD4] | 09130 |
| hypothetical protein [Aromatoleum aromaticum] | 02875 |
| hypothetical protein [Aromatoleum aromaticum] | 01587 |
| hypothetical protein [Aromatoleum aromaticum] | 08680 |
| SURF1 family protein [Azoarcus sp. DD4] | 01195 |
| hypothetical protein [Aromatoleum aromaticum] | 07655 |
| hypothetical protein [Aromatoleum aromaticum] | 09450 |
| hypothetical protein [Azoarcus sp. DD4] | 05472 |
| efflux RND transporter permease subunit, partial [Aromatoleum evansii] | 09547 |
| hypothetical protein [Azoarcus sp. DD4] | 00518 |
| hypothetical protein [Azoarcus sp. DD4] | 01354 |
| hypothetical protein [Azoarcus sp. DD4] | 01101 |
| hypothetical protein [Azoarcus sp. DD4] | 03340 |
| hypothetical protein [Azoarcus sp. DD4] | 09872 |
| AAA family ATPase [Azoarcus sp. DD4] | 06401 |
| hypothetical protein [Azoarcus sp. DD4] | 09491 |
| acetophenone carboxylase epsilon subunit [Aromatoleum aromaticum] | 04963 |
| hypothetical protein [Azoarcus sp. DD4] | 06916 |
| bacteriophage abortive infection AbiH family protein [Aromatoleum aromaticum] | 00946 |
| acyloxyacyl hydrolase [Aromatoleum aromaticum] | 08869 |
| hypothetical protein [Azoarcus sp. DD4] | 09921 |
| hypothetical protein [Alicycliphilus denitrificans] | 08293 |
| FumA C-terminus/TtdB family hydratase beta subunit, partial [Aromatoleum evansii] | 09644 |
| protein phosphatase 2C domain-containing protein [Aromatoleum aromaticum] | 04315 |
| type II toxin-antitoxin system VapC family toxin [Aromatoleum pumilum] | 06014 |
| FAD-dependent oxidoreductase [Aromatoleum pumilum] | 03694 |
| type IV secretion system protein [Azoarcus taiwanensis] | 08273 |
| Gfo/Idh/MocA family oxidoreductase [Azoarcus taiwanensis] | 08216 |
| MULTISPECIES: type II toxin-antitoxin system HicA family toxin [Aromatoleum] | 07434 |
| DUF4258 domain-containing protein [Aromatoleum aromaticum] | 04293 |
| Hpt domain-containing protein [Azoarcus taiwanensis] | 06480 |
| type II toxin-antitoxin system MqsA family antitoxin [Aromatoleum aromaticum] | 05949 |
| hypothetical protein [Azoarcus taiwanensis] | 08580 |
| pyridoxamine 5'-phosphate oxidase family protein [Azoarcus taiwanensis] | 06910 |
| hypothetical protein [Azoarcus taiwanensis] | 05106 |
| TraB/GumN family protein [Azoarcus taiwanensis] | 08480 |
| hypothetical protein [Azoarcus taiwanensis] | 03828 |
| antibiotic biosynthesis monooxygenase [Azoarcus taiwanensis] | 08460 |
| OmpA family protein [Azoarcus taiwanensis] | 04250 |
| transposase [Aromatoleum aromaticum] | 06759 |
| integrase [Aromatoleum aromaticum] | 04618 |
| 4'-phosphopantetheinyl transferase superfamily protein [Azoarcus taiwanensis] | 04788 |
| DUF3080 family protein [Azoarcus taiwanensis] | 02718 |

| | |
|---|---|
| protein kinase family protein [Azoarcus taiwanensis] | 04254 |
| hypothetical protein [Azoarcus taiwanensis] | 07926 |
| ATP-binding protein [Azoarcus taiwanensis] | 09558 |
| hypothetical protein [Azoarcus taiwanensis] | 00015 |
| hypothetical protein [Azoarcus taiwanensis] | 03988 |
| hypothetical protein [Azoarcus taiwanensis] | 04782 |
| DUF3800 domain-containing protein [Azoarcus taiwanensis] | 10470 |
| hypothetical protein [Azoarcus taiwanensis] | 05064 |
| MULTISPECIES: type II toxin-antitoxin system HicB family antitoxin [Aromatoleum] | 00513 |
| hypothetical protein [Aromatoleum aromaticum] | 08073 |
| hypothetical protein [Azoarcus taiwanensis] | 08164 |
| twin-arginine translocation signal domain-containing protein [Azoarcus taiwanensis] | 01772 |
| DUF3305 domain-containing protein [Azoarcus taiwanensis] | 01177 |
| hypothetical protein [Aromatoleum aromaticum] | 07374 |
| GTP-binding DUF697 domain-containing protein [Azoarcus taiwanensis] | 10469 |
| MerR family transcriptional regulator [Azoarcus taiwanensis] | 02720 |
| hypothetical protein [Azoarcus taiwanensis] | 05219 |
| hypothetical protein [Aromatoleum pumilum] | 05397 |
| hypothetical protein [Aromatoleum petrolei] | 03569 |
| hypothetical protein [Aromatoleum petrolei] | 04258 |
| ATP-binding protein [Aromatoleum petrolei] | 04578 |
| hypothetical protein [Aromatoleum petrolei] | 09873 |
| hypothetical protein [Aromatoleum petrolei] | 09059 |
| hypothetical protein [Aromatoleum petrolei] | 09542 |
| hypothetical protein [Aromatoleum petrolei] | 09484 |
| hypothetical protein [Aromatoleum petrolei] | 03702 |
| UUP1 family membrane protein, partial [Azoarcus taiwanensis] | 10127 |
| hypothetical protein [Aromatoleum aromaticum] | 08575 |
| SDR family NAD(P)-dependent oxidoreductase [Aromatoleum petrolei] | 10583 |
| PadR family transcriptional regulator [Aromatoleum petrolei] | 07657 |
| hypothetical protein [Aromatoleum petrolei] | 09636 |
| hypothetical protein [Aromatoleum aromaticum] | 05659 |
| NAD-dependent epimerase/dehydratase family protein [Aromatoleum petrolei] | 01301 |
| hypothetical protein [Aromatoleum petrolei] | 09973 |
| transposase [Aromatoleum aromaticum] | 00490 |
| hypothetical protein [Aromatoleum petrolei] | 00667 |
| VPA1267 family protein [Aromatoleum petrolei] | 04257 |
| gamma-mobile-trio protein GmtX [Aromatoleum petrolei] | 02278 |
| ThuA domain-containing protein [Aromatoleum petrolei] | 02389 |
| hypothetical protein [Aromatoleum petrolei] | 04256 |
| hypothetical protein [Aromatoleum petrolei] | 04255 |
| Arm DNA-binding domain-containing protein [Azoarcus taiwanensis] | 00655 |
| SDR family NAD(P)-dependent oxidoreductase [Azoarcus taiwanensis] | 07240 |
| ATP-binding protein [Azoarcus taiwanensis] | 04489 |
| peroxiredoxin [Azoarcus taiwanensis] | 00873 |
| arginase family protein, partial [Azoarcus taiwanensis] | 09677 |
| hypothetical protein [Aromatoleum aromaticum] | 03284 |
| RimK/LysX family protein, partial [Azoarcus taiwanensis] | 04866 |
| DUF6505 family protein [Azoarcus taiwanensis] | 02381 |
| hypothetical protein [Azoarcus taiwanensis] | 06436 |
| hypothetical protein [Azoarcus taiwanensis] | 02905 |
| 50S ribosome-binding GTPase [Azoarcus taiwanensis] | 04738 |
| VTT domain-containing protein, partial [Azoarcus taiwanensis] | 02378 |
| HPr-rel-A system PqqD family peptide chaperone [Azoarcus rhizosphaerae] | 09121 |
| hypothetical protein [Azoarcus rhizosphaerae] | 00377 |
| V-containing nitrogenase subunit delta [Azoarcus rhizosphaerae] | 05733 |
| NifU family protein [Azoarcus rhizosphaerae] | 09490 |
| hypothetical protein [Azoarcus rhizosphaerae] | 00770 |
| DUF2946 family protein [Azoarcus rhizosphaerae] | 02106 |

| | |
|---|---|
| DUF2514 family protein [Azoarcus rhizosphaerae] | 02626 |
| AAA domain-containing protein [Azoarcus rhizosphaerae] | 10126 |
| hypothetical protein [Azoarcus taiwanensis] | 04693 |
| hypothetical protein [Aromatoleum aromaticum] | 01044 |
| hypothetical protein [Azoarcus rhizosphaerae] | 02105 |
| cell wall hydrolase [Aromatoleum aromaticum] | 07375 |
| Rha family transcriptional regulator [Azoarcus rhizosphaerae] | 10410 |
| transglycosylase SLT domain-containing protein [Azoarcus rhizosphaerae] | 05735 |
| 4'-phosphopantetheinyl transferase superfamily protein [Azoarcus rhizosphaerae] | 03014 |
| hypothetical protein [Azoarcus rhizosphaerae] | 05468 |
| hypothetical protein [Azoarcus rhizosphaerae] | 03106 |
| hypothetical protein [Azoarcus rhizosphaerae] | 02948 |
| hypothetical protein [Azoarcus rhizosphaerae] | 08739 |
| MULTISPECIES: hypothetical protein [Proteobacteria] | 07759 |
| polysaccharide pyruvyl transferase family protein [Azoarcus rhizosphaerae] | 08781 |
| DUF2384 domain-containing protein [Aromatoleum aromaticum] | 00134 |
| cation:proton antiporter [Azoarcus rhizosphaerae] | 03572 |
| hypothetical protein [Aromatoleum aromaticum] | 03286 |
| hypothetical protein [Azoarcus rhizosphaerae] | 07856 |
| hypothetical protein [Aromatoleum aromaticum] | 08418 |
| hypothetical protein [Aromatoleum aromaticum] | 02518 |
| hypothetical protein [Azoarcus rhizosphaerae] | 02300 |
| hypothetical protein [Aromatoleum aromaticum] | 07794 |
| alpha/beta fold hydrolase [Azoarcus rhizosphaerae] | 04149 |
| phosphoribosyltransferase domain-containing protein [Aromatoleum aromaticum] | 06666 |
| hypothetical protein [Aromatoleum aromaticum] | 04620 |
| DUF4153 domain-containing protein [Azoarcus rhizosphaerae] | 00372 |
| hypothetical protein [Thauera phenolivorans] | 02054 |
| hypothetical protein [Aromatoleum aromaticum] | 04462 |
| hypothetical protein [Aromatoleum aromaticum] | 03136 |
| hypothetical protein [Azoarcus taiwanensis] | 06122 |
| polysaccharide deacetylase family protein, partial [Azoarcus taiwanensis] | 08751 |
| hypothetical protein [Azoarcus taiwanensis] | 09953 |
| roadblock/LC7 domain-containing protein [Azoarcus taiwanensis] | 03841 |
| hypothetical protein [Azoarcus taiwanensis] | 09984 |
| zinc metallopeptidase [Azoarcus taiwanensis] | 09983 |
| hypothetical protein [Azoarcus taiwanensis] | 03010 |
| type II toxin-antitoxin system RelE/ParE family toxin [Aromatoleum aromaticum] | 02920 |
| hypothetical protein [Aromatoleum aromaticum] | 08074 |
| MULTISPECIES: hypothetical protein [Rhodocyclales] | 10456 |
| NERD domain-containing protein [Thauera phenolivorans] | 06012 |
| hypothetical protein [Thauera phenolivorans] | 09885 |
| hypothetical protein [Aromatoleum aromaticum] | 05789 |
| winged helix-turn-helix transcriptional regulator [Aromatoleum aromaticum] | 00950 |
| MFS transporter, partial [Thauera phenolivorans] | 04685 |
| hypothetical protein [Thauera phenolivorans] | 09062 |
| membrane protein [Aromatoleum aromaticum] | 07908 |
| hypothetical protein [Thauera phenolivorans] | 03765 |
| hypothetical protein [Thauera phenolivorans] | 00870 |
| hypothetical protein [Thauera phenolivorans] | 02590 |
| hypothetical protein [Thauera phenolivorans] | 06861 |
| plasmid-related transcriptional repressor protein [Thauera phenolivorans] | 09433 |
| methyltransferase domain-containing protein [Thauera phenolivorans] | 09546 |
| hypothetical protein [Thauera phenolivorans] | 07789 |
| sulfatase-like hydrolase/transferase [Thauera phenolivorans] | 06333 |
| erythromycin esterase family protein [Aromatoleum aromaticum] | 08268 |
| hypothetical protein [Thauera phenolivorans] | 05413 |
| hypothetical protein [Thauera phenolivorans] | 06869 |
| hypothetical protein [Thauera phenolivorans] | 02596 |

| | |
|---|---|
| acyl carrier protein [Thauera phenolivorans] | 02663 |
| GtrA family protein [Aromatoleum petrolei] | 06059 |
| GNAT family N-acetyltransferase [Aromatoleum petrolei] | 10230 |
| hypothetical protein [Aromatoleum petrolei] | 02476 |
| phosphate/phosphite/phosphonate ABC transporter substrate-binding protein [Zoogloea ramigera] | 00593 |
| WS/DGAT domain-containing protein [Zoogloea ramigera] | 10228 |
| hypothetical protein [Aromatoleum aromaticum] | 09001 |
| hypothetical protein [Zoogloea ramigera] | 08825 |
| hypothetical protein [Aromatoleum aromaticum] | 02517 |
| hypothetical protein [Zoogloea ramigera] | 05746 |
| PIN domain-containing protein, partial [Zoogloea ramigera] | 06064 |
| hypothetical protein [Zoogloea ramigera] | 08841 |
| ankyrin repeat domain-containing protein, partial [Thauera butanivorans] | 07201 |
| tetratricopeptide repeat protein [Zoogloea ramigera] | 05500 |
| hypothetical protein [Zoogloea ramigera] | 08529 |
| DUF4384 domain-containing protein [Zoogloea ramigera] | 07886 |
| caspase family protein [Zoogloea ramigera] | 02323 |
| hypothetical protein [Zoogloea ramigera] | 02676 |
| polysaccharide deacetylase family protein [Zoogloea ramigera] | 06086 |
| 2Fe-2S iron-sulfur cluster-binding protein [Zoogloea ramigera] | 00381 |
| hypothetical protein [Zoogloea ramigera] | 00380 |
| hypothetical protein [Zoogloea ramigera] | 05475 |
| LPP20 family lipoprotein [Zoogloea ramigera] | 06923 |
| hypothetical protein [Zoogloea ramigera] | 10424 |
| ParB/Srx family N-terminal domain-containing protein [Zoogloea ramigera] | 01408 |
| helix-turn-helix domain-containing protein [Zoogloea ramigera] | 02957 |
| class I SAM-dependent methyltransferase [Zoogloea ramigera] | 07224 |
| hypothetical protein [Zoogloea ramigera] | 05188 |
| N-6 DNA methylase [Zoogloea ramigera] | 10319 |
| hypothetical protein [Aromatoleum aromaticum] | 01819 |
| hypothetical protein [Aromatoleum aromaticum] | 06267 |
| hypothetical protein [Thauera butanivorans] | 08032 |
| NUDIX hydrolase [Thauera butanivorans] | 10143 |
| linear amide C-N hydrolase [Thauera butanivorans] | 06067 |
| hypothetical protein [Aromatoleum aromaticum] | 03826 |
| KTSC domain-containing protein [Aromatoleum aromaticum] | 02363 |
| sterol desaturase family protein [Zoogloea ramigera] | 04413 |
| ankyrin repeat domain-containing protein [Zoogloea ramigera] | 05477 |
| hypothetical protein [Zoogloea ramigera] | 10426 |
| DUF3106 domain-containing protein [Oryzomicrobium terrae] | 06627 |
| molecular chaperone DnaK [Aromatoleum pumilum] | 08186 |
| SIR2 family protein [Azoarcus sp. TTM-91] | 09572 |
| TerD family protein [Aromatoleum aromaticum] | 00985 |
| hypothetical protein [Aromatoleum aromaticum] | 00806 |
| hypothetical protein [Azoarcus sp. TTM-91] | 01815 |
| hypothetical protein [Azoarcus sp. TTM-91] | 02396 |
| cysteine protease StiP family protein [Aromatoleum aromaticum] | 01994 |
| fimbria/pilus outer membrane usher protein [Azoarcus sp. TTM-91] | 06502 |
| fimbria/pilus periplasmic chaperone [Azoarcus sp. TTM-91] | 03311 |
| hypothetical protein [Azoarcus sp. TTM-91] | 01792 |
| autotransporter domain-containing protein [Oryzomicrobium terrae] | 08029 |
| TerD family protein [Aromatoleum aromaticum] | 06661 |
| MFS transporter [Oryzomicrobium terrae] | 06994 |
| PilZ domain-containing protein [Oryzomicrobium terrae] | 01072 |
| hypothetical protein [Zoogloea ramigera] | 04439 |
| hypothetical protein [Zoogloea ramigera] | 03938 |
| DUF945 family protein [Zoogloea ramigera] | 09163 |
| PriCT-2 domain-containing protein [Zoogloea ramigera] | 07874 |
| glycosyltransferase [Zoogloea ramigera] | 02309 |

| | |
|---|---|
| MAE_28990/MAE_18760 family HEPN-like nuclease [Zoogloea ramigera] | 06088 |
| ParA family protein [Zoogloea ramigera] | 02981 |
| hypothetical protein [Zoogloea ramigera] | 02682 |
| ATP-binding protein [Aromatoleum aromaticum] | 07724 |
| hypothetical protein [Zoogloea ramigera] | 02329 |
| hypothetical protein [Aromatoleum aromaticum] | 08071 |
| hypothetical protein [Zoogloea ramigera] | 00384 |
| DVUA0089 family protein [Zoogloea ramigera] | 02357 |
| TIGR02221 family CRISPR-associated protein [Zoogloea ramigera] | 04378 |
| hypothetical protein [Zoogloea ramigera] | 09948 |
| hypothetical protein [Zoogloea ramigera] | 02325 |
| hypothetical protein [Thauera butanivorans] | 00029 |
| DUF4279 domain-containing protein [Thauera butanivorans] | 04678 |
| integrase family protein [Aromatoleum petrolei] | 08434 |
| hypothetical protein [Thauera butanivorans] | 07161 |
| CoA transferase, partial [Azoarcus indigens] | 03006 |
| response regulator [Azoarcus indigens] | 03007 |
| DUF3150 domain-containing protein [Aromatoleum aromaticum] | 09805 |
| ABC transporter substrate-binding protein [Thauera butanivorans] | 04669 |
| hypothetical protein [Thauera butanivorans] | 08128 |
| PEP-CTERM sorting domain-containing protein [Thauera butanivorans] | 09049 |
| hypothetical protein [Thauera butanivorans] | 05707 |
| VWA domain-containing protein [Aromatoleum aromaticum] | 04467 |
| hypothetical protein [Thauera butanivorans] | 00869 |
| hypothetical protein [Thauera butanivorans] | 04356 |
| hypothetical protein [Thauera butanivorans] | 07773 |
| hypothetical protein [Thauera butanivorans] | 03806 |
| hypothetical protein [Thauera butanivorans] | 03812 |
| hypothetical protein [Thauera butanivorans] | 08457 |
| AAA family ATPase, partial [Azoarcus indigens] | 08281 |
| HAMP domain-containing histidine kinase, partial [Azoarcus indigens] | 01769 |
| TolC family protein, partial [Azoarcus indigens] | 06121 |
| chromate resistance protein, partial [Azoarcus indigens] | 04476 |
| aminotransferase class III-fold pyridoxal phosphate-dependent enzyme, partial [Azoarcus indigens] | 05803 |
| RHS repeat-associated core domain-containing protein, partial [Azoarcus indigens] | 03855 |
| filamentous hemagglutinin N-terminal domain-containing protein, partial [Azoarcus indigens] | 04475 |
| DUF1566 domain-containing protein, partial [Azoarcus indigens] | 02393 |
| hypothetical protein, partial [Azoarcus indigens] | 01455 |
| HPr-rel-A system PqqD family peptide chaperone [Azoarcus indigens] | 03982 |
| nuclear transport factor 2 family protein [Aromatoleum aromaticum] | 10280 |
| class II glutamine amidotransferase [Azoarcus indigens] | 03496 |
| MFS transporter [Aromatoleum petrolei] | 03732 |
| hypothetical protein [Aromatoleum petrolei] | 03263 |
| recombinase family protein [Aromatoleum aromaticum] | 03412 |
| hypothetical protein [Aromatoleum petrolei] | 05617 |
| hypothetical protein [Aromatoleum petrolei] | 05796 |
| phosphoadenosine phosphosulfate reductase family protein [Aromatoleum aromaticum] | 05517 |
| hypothetical protein [Thauera butanivorans] | 04345 |
| glycosyltransferase [Thauera butanivorans] | 05223 |
| hypothetical protein [Thauera butanivorans] | 01022 |
| hypothetical protein [Thauera butanivorans] | 01645 |
| hypothetical protein [Thauera butanivorans] | 09946 |
| hypothetical protein [Thauera butanivorans] | 05082 |
| hypothetical protein [Thauera butanivorans] | 01644 |
| DUF4224 domain-containing protein [Thauera butanivorans] | 02591 |
| hypothetical protein [Thauera butanivorans] | 04676 |
| hypothetical protein [Aromatoleum aromaticum] | 03000 |
| MULTISPECIES: DUF523 domain-containing protein [Betaproteobacteria] | 00516 |
| RHS domain-containing protein [Thauera butanivorans] | 03583 |

| | |
|---|---|
| DUF11 domain-containing protein [Thauera butanivorans] | 01023 |
| hypothetical protein [Thauera butanivorans] | 04380 |
| hypothetical protein [Thauera butanivorans] | 09494 |
| SprT-like domain-containing protein, partial [Thauera butanivorans] | 02589 |
| Rpn family recombination-promoting nuclease/putative transposase [Thauera butanivorans] | 07166 |
| hypothetical protein [Thauera butanivorans] | 09865 |
| hypothetical protein [Thauera butanivorans] | 05996 |
| hypothetical protein [Thauera butanivorans] | 02588 |
| hypothetical protein [Thauera butanivorans] | 03338 |
| AbrB/MazE/SpoVT family DNA-binding domain-containing protein [Thauera butanivorans] | 09072 |
| DUF6364 family protein [Thauera butanivorans] | 03663 |
| hypothetical protein [Thauera butanivorans] | 08747 |
| hypothetical protein [Thauera butanivorans] | 05992 |
| hypothetical protein [Thauera butanivorans] | 06892 |
| hypothetical protein [Thauera butanivorans] | 04734 |
| glycosyltransferase [Thauera butanivorans] | 03662 |
| EamA family transporter [Thauera butanivorans] | 07074 |
| DUF1428 domain-containing protein [Thauera butanivorans] | 04185 |
| EthD family reductase [Thauera butanivorans] | 01020 |
| MmoB/DmpM family protein [Thauera butanivorans] | 09425 |
| hypothetical protein [Thauera butanivorans] | 04673 |
| hypothetical protein [Azoarcus taiwanensis] | 00022 |
| circadian clock KaiB family protein [Aromatoleum diolicum] | 07220 |
| hypothetical protein [Thauera terpenica] | 08025 |
| diguanylate cyclase [Zoogloea dura] | 09643 |
| hypothetical protein [Zoogloea dura] | 02404 |
| DUF5908 family protein [Zoogloea dura] | 01141 |
| tripartite tricarboxylate transporter TctB family protein [Propionivibrio dicarboxylicus] | 01919 |
| class I SAM-dependent methyltransferase [Zoogloea dura] | 00442 |
| acyl carrier protein [Zoogloea dura] | 00682 |
| hypothetical protein [Zoogloea dura] | 06147 |
| filamentous hemagglutinin N-terminal domain-containing protein [Zoogloea dura] | 09945 |
| DUF748 domain-containing protein [Propionivibrio dicarboxylicus] | 00151 |
| hydroxyethylthiazole kinase [Propionivibrio dicarboxylicus] | 07705 |
| polyketide cyclase [Propionivibrio dicarboxylicus] | 04284 |
| hypothetical protein [Zoogloea dura] | 04020 |
| hypothetical protein [Zoogloea dura] | 05185 |
| DUF108 domain-containing protein [Propionivibrio dicarboxylicus] | 08405 |
| DUF6491 family protein [Zoogloea dura] | 02330 |
| DUF932 domain-containing protein [Propionivibrio dicarboxylicus] | 09545 |
| hypothetical protein [Zoogloea dura] | 09082 |
| sulfide/dihydroorotate dehydrogenase-like FAD/NAD-binding protein [Propionivibrio dicarboxylicus] | 05945 |
| hypothetical protein [Zoogloea dura] | 09263 |
| TIR domain-containing protein [Zoogloea dura] | 04515 |
| hypothetical protein [Zoogloea dura] | 02743 |
| DUF3373 domain-containing protein [Propionivibrio dicarboxylicus] | 06647 |
| glycosyltransferase family 2 protein [Zoogloea dura] | 00445 |
| hypothetical protein [Zoogloea dura] | 07200 |
| hypothetical protein [Propionivibrio dicarboxylicus] | 02410 |
| type II secretion system protein GspM [Propionivibrio dicarboxylicus] | 10178 |
| terminase small subunit [Zoogloea dura] | 06507 |
| type II secretion system protein N [Propionivibrio dicarboxylicus] | 08059 |
| hypothetical protein [Zoogloea dura] | 02402 |
| hypothetical protein [Propionivibrio dicarboxylicus] | 06052 |
| baseplate J/gp47 family protein [Zoogloea dura] | 10432 |
| hypothetical protein [Zoogloea dura] | 07533 |
| hypothetical protein [Uliginosibacterium aquaticum] | 02216 |
| hypothetical protein [Zoogloea dura] | 01206 |
| hypothetical protein [Aromatoleum tolulyticum] | 03420 |

| | |
|---|---|
| hypothetical protein [Aromatoleum tolulyticum] | 00945 |
| hypothetical protein [Zoogloea dura] | 02207 |
| hypothetical protein [Aromatoleum tolulyticum] | 01585 |
| hypothetical protein [Zoogloea dura] | 05834 |
| sulfatase-like hydrolase/transferase [Zoogloea dura] | 03419 |
| hypothetical protein [Aromatoleum tolulyticum] | 04993 |
| DUF5666 domain-containing protein [Aromatoleum tolulyticum] | 07590 |
| DUF2244 domain-containing protein [Zoogloea dura] | 06092 |
| methyltransferase type 12 [Zoogloea dura] | 03705 |
| SRPBCC domain-containing protein [Aromatoleum tolulyticum] | 06653 |
| hypothetical protein [Zoogloea dura] | 03940 |
| tetratricopeptide repeat protein [Zoogloea dura] | 07269 |
| zinc dependent phospholipase C family protein [Zoogloea dura] | 02746 |
| pyridoxamine 5'-phosphate oxidase family protein [Zoogloea dura] | 05832 |
| VOC family protein [Propionivibrio dicarboxylicus] | 02415 |
| hypothetical protein [Propionivibrio dicarboxylicus] | 07362 |
| hypothetical protein [Zoogloea dura] | 01716 |
| class I SAM-dependent methyltransferase [Propionivibrio dicarboxylicus] | 04701 |
| hypothetical protein [Propionivibrio dicarboxylicus] | 10346 |
| DUF554 domain-containing protein [Propionivibrio dicarboxylicus] | 05409 |
| protease inhibitor I42 family protein [Zoogloea dura] | 10152 |
| hypothetical protein [Zoogloea dura] | 00397 |
| type I restriction-modification system subunit M N-terminal domain-containing protein [Zoogloea dura] | 06637 |
| type III secretion system outer membrane ring subunit SctC [Propionivibrio dicarboxylicus] | 06292 |
| hypothetical protein [Zoogloea dura] | 06084 |
| hypothetical protein [Zoogloea dura] | 02980 |
| DUF4337 domain-containing protein [Propionivibrio dicarboxylicus] | 03773 |
| helix-turn-helix transcriptional regulator [Propionivibrio dicarboxylicus] | 09222 |
| hypothetical protein [Propionivibrio dicarboxylicus] | 03603 |
| hypothetical protein [Propionivibrio dicarboxylicus] | 05944 |
| cell division protein FtsK [Zoogloea dura] | 04405 |
| hypothetical protein [Propionivibrio dicarboxylicus] | 04973 |
| FHIPEP family type III secretion protein, partial [Thauera phenylacetica] | 05813 |
| ParA family protein [Thauera phenylacetica] | 05961 |
| hypothetical protein [Thauera phenylacetica] | 07390 |
| MULTISPECIES: CcdB family protein [Thauera] | 03368 |
| hypothetical protein [Thauera phenylacetica] | 07158 |
| thioredoxin family protein [Propionivibrio dicarboxylicus] | 07530 |
| hypothetical protein [Thauera phenylacetica] | 05720 |
| hypothetical protein [Thauera phenylacetica] | 06283 |
| DEAD/DEAH box helicase [Thauera phenylacetica] | 02032 |
| TauD/TfdA family dioxygenase [Thauera phenylacetica] | 04145 |
| HAD family hydrolase [Propionivibrio dicarboxylicus] | 05729 |
| CoA-binding protein [Propionivibrio dicarboxylicus] | 05002 |
| peptidoglycan-binding protein, partial [Thauera phenylacetica] | 07814 |
| helix-turn-helix domain-containing protein, partial [Thauera phenylacetica] | 08742 |
| hypothetical protein [Propionivibrio dicarboxylicus] | 06629 |
| FAD-dependent oxidoreductase, partial [Thauera phenylacetica] | 05438 |
| AMP-binding protein, partial [Thauera phenylacetica] | 05960 |
| hypothetical protein, partial [Thauera phenylacetica] | 03306 |
| PD-(D/E)XK nuclease family protein, partial [Thauera phenylacetica] | 02014 |
| hypothetical protein, partial [Thauera phenylacetica] | 09958 |
| DUF2868 domain-containing protein, partial [Thauera phenylacetica] | 05032 |
| prepilin-type N-terminal cleavage/methylation domain-containing protein, partial [Thauera phenylacetica] | 06281 |
| aspartate/glutamate racemase family protein [Propionivibrio dicarboxylicus] | 08710 |
| NAD(P)/FAD-dependent oxidoreductase, partial [Thauera phenylacetica] | 09416 |
| hypothetical protein, partial [Thauera phenylacetica] | 07735 |
| hypothetical protein [Propionivibrio dicarboxylicus] | 06044 |
| Hpt domain-containing protein, partial [Thauera phenylacetica] | 07734 |

| | |
|---|---|
| DUF1810 domain-containing protein [Thauera phenylacetica] | 03150 |
| CHAT domain-containing protein, partial [Thauera phenylacetica] | 08429 |
| hypothetical protein [Thauera phenylacetica] | 05066 |
| AAA family ATPase, partial [Thauera phenylacetica] | 07772 |
| deoxyribodipyrimidine photo-lyase, partial [Thauera phenylacetica] | 02005 |
| hypothetical protein [Thauera phenylacetica] | 00974 |
| hypothetical protein [Propionivibrio dicarboxylicus] | 02649 |
| hypothetical protein [Zoogloea dura] | 06999 |
| hypothetical protein [Propionivibrio dicarboxylicus] | 04791 |
| AAC(3) family N-acetyltransferase [Zoogloea dura] | 09262 |
| site-specific DNA-methyltransferase [Propionivibrio dicarboxylicus] | 07938 |
| portal protein [Propionivibrio dicarboxylicus] | 01550 |
| hypothetical protein [Zoogloea dura] | 09261 |
| DUF2235 domain-containing protein [Thauera phenylacetica] | 08478 |
| hypothetical protein [Thauera phenylacetica] | 00975 |
| hypothetical protein, partial [Thauera phenylacetica] | 05673 |
| class I SAM-dependent methyltransferase [Propionivibrio dicarboxylicus] | 00088 |
| hypothetical protein [Thauera phenylacetica] | 06808 |
| hypothetical protein [Thauera phenylacetica] | 05955 |
| hypothetical protein [Thauera phenylacetica] | 09012 |
| hypothetical protein, partial [Thauera phenylacetica] | 05033 |
| hypothetical protein [Thauera phenylacetica] | 08706 |
| restriction endonuclease subunit S [Thauera phenylacetica] | 01791 |
| hypothetical protein [Thauera phenylacetica] | 10024 |
| hypothetical protein [Thauera phenylacetica] | 10530 |
| hypothetical protein [Thauera phenylacetica] | 05530 |
| YjbF family lipoprotein [Propionivibrio dicarboxylicus] | 05038 |
| SAM-dependent methyltransferase [Thauera phenylacetica] | 02016 |
| lysylphosphatidylglycerol synthase domain-containing protein [Thauera phenylacetica] | 08705 |
| DUF4276 family protein [Thauera phenylacetica] | 05356 |
| DUF6314 family protein [Thauera phenylacetica] | 00973 |
| SAVED domain-containing protein [Thauera phenylacetica] | 04635 |
| response regulator [Propionivibrio dicarboxylicus] | 06256 |
| hypothetical protein [Thauera phenylacetica] | 02015 |
| DUF805 domain-containing protein [Zoogloea dura] | 10376 |
| hypothetical protein [Zoogloea dura] | 00591 |
| PQQ-dependent sugar dehydrogenase [Zoogloea dura] | 01736 |
| 3-hydroxyacyl-CoA dehydrogenase family protein [Thauera chlorobenzoica] | 06338 |
| MULTISPECIES: hypothetical protein [Proteobacteria] | 09676 |
| CDGSH iron-sulfur domain-containing protein [Zoogloea dura] | 07271 |
| class I SAM-dependent methyltransferase [Zoogloea dura] | 05476 |
| hypothetical protein [Zoogloea dura] | 07137 |
| hypothetical protein [Thauera chlorobenzoica] | 04147 |
| hypothetical protein [Thauera chlorobenzoica] | 03196 |
| hypothetical protein [Zoogloea dura] | 03026 |
| hypothetical protein [Zoogloea dura] | 07954 |
| hypothetical protein [Thauera chlorobenzoica] | 02377 |
| hypothetical protein [Zoogloea dura] | 06876 |
| RusA family crossover junction endodeoxyribonuclease [Zoogloea dura] | 04025 |
| hypothetical protein [Thauera chlorobenzoica] | 08759 |
| VCBS domain-containing protein, partial [Zoogloea dura] | 04552 |
| hypothetical protein [Zoogloea dura] | 00686 |
| endonuclease NucS [Zoogloea dura] | 01626 |
| hypothetical protein [Thauera chlorobenzoica] | 03527 |
| hypothetical protein [Zoogloea dura] | 08497 |
| hypothetical protein [Thauera chlorobenzoica] | 02434 |
| MBL fold metallo-hydrolase [Zoogloea dura] | 04435 |
| hypothetical protein [Zoogloea dura] | 04024 |
| hypothetical protein [Zoogloea dura] | 10499 |

| | |
|---|---|
| hypothetical protein [Thauera chlorobenzoica] | 06756 |
| hypothetical protein [Thauera chlorobenzoica] | 06179 |
| hypothetical protein [Thauera chlorobenzoica] | 04691 |
| hypothetical protein [Zoogloea dura] | 08909 |
| hypothetical protein [Thauera chlorobenzoica] | 00866 |
| hypothetical protein [Zoogloea dura] | 01182 |
| YodC family protein [Zoogloea dura] | 06153 |
| DotA/TraY family protein [Thauera chlorobenzoica] | 08098 |
| class I SAM-dependent methyltransferase [Zoogloea dura] | 00687 |
| DUF4255 domain-containing protein [Zoogloea dura] | 01818 |
| type VI secretion system tip protein VgrG [Zoogloea dura] | 07536 |
| MULTISPECIES: hypothetical protein [Uliginosibacterium] | 02753 |
| MULTISPECIES: hypothetical protein [Uliginosibacterium] | 03029 |
| MULTISPECIES: hypothetical protein [Uliginosibacterium] | 06542 |
| MULTISPECIES: hypothetical protein [Uliginosibacterium] | 06515 |
| ParB N-terminal domain-containing protein [Thauera chlorobenzoica] | 08469 |
| MULTISPECIES: nuclear transport factor 2 family protein [Uliginosibacterium] | 08314 |
| MULTISPECIES: hypothetical protein [Uliginosibacterium] | 02562 |
| MULTISPECIES: hypothetical protein [Uliginosibacterium] | 02025 |
| hypothetical protein [Thauera chlorobenzoica] | 00864 |
| hypothetical protein [Thauera chlorobenzoica] | 03679 |
| hypothetical protein [Thauera chlorobenzoica] | 02065 |
| MULTISPECIES: hypothetical protein [Uliginosibacterium] | 02236 |
| hypothetical protein [Thauera chlorobenzoica] | 02066 |
| hypothetical protein [Thauera chlorobenzoica] | 08470 |
| hypothetical protein [Thauera chlorobenzoica] | 05870 |
| class I SAM-dependent methyltransferase [Zoogloea dura] | 10500 |
| hypothetical protein [Thauera chlorobenzoica] | 08758 |
| MULTISPECIES: hypothetical protein [Uliginosibacterium] | 01238 |
| MULTISPECIES: membrane protease subunit, stomatin/prohibitin [Uliginosibacterium] | 03504 |
| hypothetical protein [Thauera chlorobenzoica] | 07449 |
| flagellar basal body rod protein FlgB [Zoogloea dura] | 08911 |
| Hpt domain-containing protein [Zoogloea dura] | 01717 |
| methyl-accepting chemotaxis protein, partial [Zoogloea dura] | 08843 |
| hypothetical protein [Thauera chlorobenzoica] | 00721 |
| NifU family protein [Zoogloea dura] | 07272 |
| PEP-CTERM sorting domain-containing protein [Zoogloea dura] | 06132 |
| methyl-accepting chemotaxis protein [Thauera chlorobenzoica] | 06008 |
| MFS transporter [Zoogloea dura] | 04518 |
| site-specific integrase [Thauera chlorobenzoica] | 00329 |
| hypothetical protein [Zoogloea dura] | 00453 |
| hypothetical protein [Aromatoleum tolulyticum] | 07253 |
| hypothetical protein [Aromatoleum tolulyticum] | 07371 |
| SH3 domain-containing protein [Zoogloea dura] | 02982 |
| PAS domain-containing protein [Zoogloea dura] | 10431 |
| hypothetical protein [Zoogloea dura] | 07951 |
| DUF4124 domain-containing protein [Aromatoleum tolulyticum] | 02813 |
| phosphate ABC transporter substrate-binding protein [Zoogloea dura] | 07296 |
| hypothetical protein [Zoogloea dura] | 00450 |
| hypothetical protein [Zoogloea dura] | 08345 |
| hypothetical protein [Zoogloea dura] | 00005 |
| MULTISPECIES: elongation factor Tu, partial [Proteobacteria] | 01988 |
| hypothetical protein [Zoogloea dura] | 10498 |
| hypothetical protein [Aromatoleum tolulyticum] | 06438 |
| DUF2868 domain-containing protein [Zoogloea dura] | 02136 |
| hypothetical protein [Azoarcus tolulyticus] | 00249 |
| hypothetical protein [Zoogloea dura] | 01733 |
| hypothetical protein [Zoogloea dura] | 05498 |
| hypothetical protein [Zoogloea dura] | 10154 |

| | |
|---|---|
| hypothetical protein [Zoogloea dura] | 08245 |
| hypothetical protein [Aromatoleum tolulyticum] | 09802 |
| phage holin family protein [Zoogloea dura] | 04021 |
| hypothetical protein [Aromatoleum tolulyticum] | 03670 |
| hypothetical protein [Aromatoleum tolulyticum] | 07715 |
| hypothetical protein [Zoogloea dura] | 01464 |
| hypothetical protein [Zoogloea dura] | 00089 |
| cyanophycinase [Zoogloea dura] | 01939 |
| hypothetical protein [Zoogloea dura] | 03265 |
| aldolase [Zoogloea dura] | 07896 |
| hypothetical protein [Aromatoleum tolulyticum] | 10481 |
| hypothetical protein [Aromatoleum tolulyticum] | 06139 |
| hypothetical protein [Zoogloea dura] | 01085 |
| hypothetical protein [Aromatoleum tolulyticum] | 08412 |
| hypothetical protein [Aromatoleum tolulyticum] | 06837 |
| hypothetical protein [Zoogloea dura] | 01135 |
| hypothetical protein [Aromatoleum tolulyticum] | 08290 |
| dehydrogenase [Aromatoleum tolulyticum] | 06742 |
| metal-dependent hydrolase [Aromatoleum tolulyticum] | 00992 |
| hypothetical protein [Aromatoleum tolulyticum] | 06262 |
| hypothetical protein [Aromatoleum tolulyticum] | 05650 |
| hypothetical protein [Zoogloea dura] | 06936 |
| HNH endonuclease [Zoogloea dura] | 03448 |
| hypothetical protein [Aromatoleum tolulyticum] | 03279 |
| hypothetical protein [Aromatoleum tolulyticum] | 09372 |
| hypothetical protein [Aromatoleum tolulyticum] | 01669 |
| kynureninase [Zoogloea sp. G-4-1-14] | 04481 |
| HD domain-containing protein [Aromatoleum tolulyticum] | 03486 |
| hypothetical protein [Zoogloea dura] | 08813 |
| hypothetical protein [Aromatoleum tolulyticum] | 06095 |
| glycosyltransferase [Aromatoleum tolulyticum] | 01217 |
| hypothetical protein [Aromatoleum tolulyticum] | 09458 |
| alkaline phosphatase D family protein [Aromatoleum tolulyticum] | 02433 |
| hypothetical protein [Aromatoleum tolulyticum] | 08409 |
| hypothetical protein [Zoogloea dura] | 00472 |
| hypothetical protein [Zoogloea dura] | 07383 |
| hypothetical protein [Aromatoleum tolulyticum] | 04670 |
| metallo-mystery pair system four-Cys motif protein [Zoogloea dura] | 09179 |
| hypothetical protein [Zoogloea dura] | 00400 |
| hypothetical protein [Zoogloea dura] | 00401 |
| peptidoglycan DD-metalloendopeptidase family protein [Zoogloea dura] | 05556 |
| class II aldolase/adducin family protein [Zoogloea dura] | 04889 |
| hypothetical protein [Aromatoleum tolulyticum] | 06000 |
| hypothetical protein [Aromatoleum tolulyticum] | 10253 |
| GNAT family N-acetyltransferase [Zoogloea dura] | 03067 |
| hypothetical protein [Aromatoleum tolulyticum] | 09373 |
| translocation/assembly module TamB domain-containing protein, partial [Thauera phenylacetica] | 09994 |
| DUF2130 domain-containing protein [Propionivibrio dicarboxylicus] | 05225 |
| Hpt domain-containing protein, partial [Thauera phenylacetica] | 09394 |
| hypothetical protein [Azoarcus communis] | 01528 |
| hypothetical protein [Aromatoleum toluclasticum] | 10254 |
| hypothetical protein [Aromatoleum toluclasticum] | 09036 |
| diguanylate cyclase [Aromatoleum toluclasticum] | 09412 |
| DNA-binding protein [Azoarcus communis] | 00647 |
| hypothetical protein [Aromatoleum toluclasticum] | 10309 |
| hypothetical protein [Aromatoleum toluclasticum] | 01327 |
| hypothetical protein [Aromatoleum toluclasticum] | 08987 |
| DDE-type integrase/transposase/recombinase [Aromatoleum toluclasticum] | 00066 |
| DUF1508 domain-containing protein [Aromatoleum toluclasticum] | 06779 |

| | |
|---|---|
| TIGR04255 family protein [Azoarcus communis] | 05234 |
| type I restriction-modification system subunit M N-terminal domain-containing protein [Azoarcus communis] | 06894 |
| hypothetical protein [Azoarcus communis] | 00349 |
| type IV pili methyl-accepting chemotaxis transducer N-terminal domain-containing protein [Aromatoleum toluclasticum] | 05690 |
| hypothetical protein [Azoarcus communis] | 03917 |
| DUF935 family protein [Azoarcus communis] | 07231 |
| hypothetical protein [Aromatoleum toluclasticum] | 01980 |
| hypothetical protein [Aromatoleum toluclasticum] | 01255 |
| hypothetical protein [Aromatoleum toluclasticum] | 04051 |
| ATP synthase subunit C [Aromatoleum toluclasticum] | 07359 |
| V-type ATP synthase subunit F [Aromatoleum toluclasticum] | 08986 |
| hypothetical protein [Aromatoleum toluclasticum] | 10106 |
| V-type ATP synthase subunit D [Aromatoleum toluclasticum] | 02798 |
| hypothetical protein [Azoarcus communis] | 01380 |
| hypothetical protein [Aromatoleum toluclasticum] | 10252 |
| hypothetical protein [Azoarcus communis] | 01025 |
| molecular chaperone TorD family protein [Aromatoleum toluclasticum] | 03123 |
| hypothetical protein [Azoarcus communis] | 07921 |
| hypothetical protein [Azoarcus communis] | 04856 |
| hypothetical protein [Azoarcus communis] | 06472 |
| hypothetical protein [Azoarcus communis] | 10463 |
| major capsid protein [Azoarcus communis] | 05233 |
| hypothetical protein [Azoarcus communis] | 02238 |
| hypothetical protein [Aromatoleum toluclasticum] | 00503 |
| 2Fe-2S iron-sulfur cluster binding domain-containing protein [Azoarcus communis] | 06118 |
| hypothetical protein [Azoarcus communis] | 08384 |
| endonuclease NucS [Aromatoleum toluclasticum] | 07700 |
| type IV toxin-antitoxin system AbiEi family antitoxin [Aromatoleum toluclasticum] | 07825 |
| nucleotidyl transferase AbiEii/AbiGii toxin family protein [Aromatoleum toluclasticum] | 10255 |
| hypothetical protein [Azoarcus communis] | 02574 |
| hypothetical protein [Azoarcus communis] | 06470 |
| XdhC family protein [Aromatoleum toluclasticum] | 06336 |
| hypothetical protein [Aromatoleum toluclasticum] | 01981 |
| hypothetical protein [Azoarcus communis] | 07228 |
| DUF4124 domain-containing protein [Azoarcus communis] | 01642 |
| hypothetical protein [Azoarcus communis] | 09549 |
| hypothetical protein [Aromatoleum toluclasticum] | 05909 |
| glycosyltransferase family 4 protein, partial [Aromatoleum toluclasticum] | 06768 |
| hypothetical protein [Azoarcus communis] | 00646 |
| hypothetical protein [Azoarcus communis] | 09151 |
| tape measure protein [Azoarcus communis] | 03896 |
| GNAT family N-acetyltransferase [Azoarcus communis] | 02177 |
| hypothetical protein [Aromatoleum toluclasticum] | 02215 |
| hypothetical protein [Azoarcus communis] | 03378 |
| hypothetical protein [Azoarcus communis] | 07919 |
| hypothetical protein [Aromatoleum toluclasticum] | 05053 |
| hypothetical protein [Aromatoleum toluclasticum] | 09031 |
| PEP-CTERM sorting domain-containing protein [Azoarcus communis] | 09462 |
| hypothetical protein [Azoarcus communis] | 10362 |
| hypothetical protein [Azoarcus communis] | 05112 |
| hypothetical protein [Azoarcus communis] | 06362 |
| hypothetical protein [Azoarcus communis] | 07230 |
| nucleotidyltransferase domain-containing protein [Aromatoleum toluclasticum] | 05220 |
| HAMP domain-containing protein, partial [Azoarcus communis] | 02711 |
| DUF3467 domain-containing protein [Azoarcus communis] | 10409 |
| hypothetical protein [Azoarcus communis] | 03004 |
| 7,8-didemethyl-8-hydroxy-5-deazariboflavin synthase CofG [Aromatoleum toluclasticum] | 02469 |
| ester cyclase [Aromatoleum toluclasticum] | 05840 |
| hypothetical protein, partial [Azoarcus communis] | 08879 |

| | |
|---|---|
| hypothetical protein [Azoarcus communis] | 09551 |
| PEP-CTERM sorting domain-containing protein [Azoarcus communis] | 09671 |
| hypothetical protein [Aromatoleum toluclasticum] | 10256 |
| hypothetical protein [Aromatoleum toluclasticum] | 01567 |
| glutamate ligase [Azoarcus communis] | 02180 |
| hypothetical protein [Aromatoleum toluclasticum] | 10607 |
| hypothetical protein [Azoarcus communis] | 04857 |
| hypothetical protein [Azoarcus communis] | 08587 |
| hypothetical protein [Azoarcus communis] | 03297 |
| hypothetical protein [Aromatoleum toluclasticum] | 00996 |
| hypothetical protein [Aromatoleum toluclasticum] | 09784 |
| STAS domain-containing protein [Aromatoleum toluclasticum] | 09360 |
| hypothetical protein [Azoarcus communis] | 00478 |
| hypothetical protein [Azoarcus communis] | 04710 |
| hypothetical protein [Azoarcus communis] | 06392 |
| hypothetical protein [Azoarcus communis] | 08495 |
| PIN domain-containing protein [Azoarcus communis] | 03780 |
| DUF637 domain-containing protein, partial [Azoarcus communis] | 01422 |
| hypothetical protein [Azoarcus communis] | 05801 |
| DUF45 domain-containing protein [Azoarcus communis] | 09095 |
| hypothetical protein [Azoarcus communis] | 07348 |
| ring-opening amidohydrolase [Azoarcus communis] | 07203 |
| HipA domain-containing protein [Azoarcus communis] | 03464 |
| DUF3047 domain-containing protein [Azoarcus communis] | 00886 |
| nucleotidyl transferase AbiEii/AbiGii toxin family protein [Azoarcus communis] | 02662 |
| hypothetical protein [Azoarcus communis] | 06410 |
| 5-amino-6-(D-ribitylamino)uracil--L-tyrosine 4-hydroxyphenyl transferase CofH [Aromatoleum toluclasticum] | 09038 |
| DUF3604 domain-containing protein [Aromatoleum toluclasticum] | 09845 |
| hypothetical protein [Azoarcus communis] | 08878 |
| hypothetical protein [Azoarcus communis] | 07566 |
| radical SAM protein [Aromatoleum toluclasticum] | 08051 |
| hypothetical protein [Aromatoleum toluclasticum] | 07698 |
| glycosyltransferase [Aromatoleum toluclasticum] | 10403 |
| hypothetical protein [Azoarcus communis] | 02373 |
| hypothetical protein [Aromatoleum toluclasticum] | 10010 |
| hypothetical protein [Azoarcus communis] | 06473 |
| hypothetical protein [Azoarcus communis] | 03980 |
| TlpA family protein disulfide reductase [Azoarcus communis] | 07463 |
| hypothetical protein [Aromatoleum toluclasticum] | 04281 |
| multiubiquitin domain-containing protein [Azoarcus communis] | 09147 |
| recombinase family protein [Aromatoleum toluclasticum] | 07358 |
| hypothetical protein [Azoarcus communis] | 01431 |
| sulfite exporter TauE/SafE family protein [Aromatoleum toluclasticum] | 04656 |
| gamma-glutamylcyclotransferase [Aromatoleum toluclasticum] | 07420 |
| hypothetical protein [Aromatoleum toluclasticum] | 03174 |
| hypothetical protein [Aromatoleum toluclasticum] | 08401 |
| hypothetical protein [Aromatoleum toluclasticum] | 08449 |
| hypothetical protein [Aromatoleum toluclasticum] | 05318 |
| hypothetical protein [Aromatoleum toluclasticum] | 04107 |
| hypothetical protein [Azoarcus communis] | 10472 |
| hypothetical protein [Aromatoleum toluclasticum] | 10251 |
| hypothetical protein [Azoarcus communis] | 02179 |
| thioredoxin family protein [Aromatoleum toluclasticum] | 05522 |
| hypothetical protein [Aromatoleum toluclasticum] | 06252 |
| hypothetical protein [Aromatoleum toluclasticum] | 02497 |
| lycopene cyclase domain-containing protein [Aromatoleum toluclasticum] | 09361 |
| hypothetical protein [Aromatoleum toluclasticum] | 00288 |
| CopK family periplasmic copper-binding protein [Aromatoleum toluclasticum] | 00785 |
| filamentous hemagglutinin N-terminal domain-containing protein [Azoarcus communis] | 05157 |

| | |
|---|---|
| pentapeptide repeat-containing protein [Azoarcus communis] | 07871 |
| alpha/beta hydrolase [Propionivibrio dicarboxylicus] | 03156 |
| hypothetical protein [Propionivibrio dicarboxylicus] | 08606 |
| hypothetical protein [Propionivibrio dicarboxylicus] | 08225 |
| hypothetical protein [Thauera phenylacetica] | 02009 |
| hypothetical protein [Thauera phenylacetica] | 07388 |
| hypothetical protein, partial [Thauera phenylacetica] | 03146 |
| anion transporter [Propionivibrio dicarboxylicus] | 02843 |
| hypothetical protein [Propionivibrio dicarboxylicus] | 10621 |
| DUF1799 domain-containing protein [Propionivibrio dicarboxylicus] | 06989 |
| phage tail length tape measure family protein [Propionivibrio dicarboxylicus] | 03460 |
| 50S ribosome-binding GTPase, partial [Thauera phenylacetica] | 02541 |
| citrate lyase subunit alpha [Propionivibrio dicarboxylicus] | 03319 |
| hypothetical protein, partial [Thauera phenylacetica] | 09812 |
| RES family NAD+ phosphorylase [Thauera phenylacetica] | 09434 |
| hypothetical protein [Thauera phenylacetica] | 06699 |
| YbfB/YjiJ family MFS transporter, partial [Thauera phenylacetica] | 06280 |
| hypothetical protein, partial [Thauera phenylacetica] | 02532 |
| Nif3-like dinuclear metal center hexameric protein, partial [Thauera phenylacetica] | 01439 |
| hypothetical protein [Thauera phenylacetica] | 03791 |
| type I-U CRISPR-associated protein Csb2 [Thauera phenylacetica] | 08093 |
| hypothetical protein [Thauera phenylacetica] | 02539 |
| transposase [Thauera phenylacetica] | 06802 |
| hypothetical protein [Thauera phenylacetica] | 08092 |
| hypothetical protein [Thauera phenylacetica] | 10086 |
| hypothetical protein [Propionivibrio dicarboxylicus] | 10592 |
| hypothetical protein [Thauera phenylacetica] | 02828 |
| DUF3486 family protein [Thauera phenylacetica] | 06276 |
| hypothetical protein [Thauera phenylacetica] | 06641 |
| TRAP transporter large permease, partial [Propionivibrio dicarboxylicus] | 01577 |
| hypothetical protein [Thauera phenylacetica] | 01186 |
| RnfABCDGE type electron transport complex subunit D, partial [Thauera phenylacetica] | 05029 |
| F420-dependent NADP oxidoreductase [Propionivibrio dicarboxylicus] | 01502 |
| hypothetical protein, partial [Thauera phenylacetica] | 03147 |
| MFS transporter, partial [Thauera phenylacetica] | 07748 |
| hypothetical protein [Thauera phenylacetica] | 03295 |
| response regulator, partial [Thauera phenylacetica] | 05351 |
| hypothetical protein, partial [Thauera phenylacetica] | 03142 |
| VC_2705 family sodium/solute symporter, partial [Thauera phenylacetica] | 02832 |
| GTP-binding protein, partial [Thauera phenylacetica] | 05959 |
| PsiF family protein, partial [Thauera phenylacetica] | 02013 |
| tRNA lysidine(34) synthetase TilS, partial [Thauera phenylacetica] | 02831 |
| translation initiation factor IF-2 associated domain-containing protein, partial [Thauera phenylacetica] | 02012 |
| MFS transporter, partial [Thauera phenylacetica] | 08426 |
| TonB C-terminal domain-containing protein, partial [Thauera phenylacetica] | 06973 |
| Hpt domain-containing protein, partial [Thauera phenylacetica] | 08085 |
| hypothetical protein [Thauera phenylacetica] | 07747 |
| tRNA (adenosine(37)-N6)-dimethylallyltransferase MiaA, partial [Thauera phenylacetica] | 07384 |
| GNAT family N-acetyltransferase [Propionivibrio dicarboxylicus] | 06149 |
| hypothetical protein, partial [Thauera phenylacetica] | 00820 |
| DUF4131 domain-containing protein, partial [Thauera phenylacetica] | 03304 |
| PEP-CTERM sorting domain-containing protein [Thauera phenylacetica] | 08084 |
| integrase domain-containing protein, partial [Thauera phenylacetica] | 03305 |
| 2OG-Fe(II) oxygenase [Thauera phenylacetica] | 03668 |
| CHASE3 domain-containing protein [Thauera phenylacetica] | 08694 |
| hypothetical protein [Thauera phenylacetica] | 03764 |
| DUF4332 domain-containing protein [Thauera phenylacetica] | 08703 |
| hypothetical protein [Thauera phenylacetica] | 09817 |
| hypothetical protein [Thauera phenylacetica] | 06643 |

| | |
|---|---|
| S8 family serine peptidase, partial [Thauera phenylacetica] | 05348 |
| DEAD/DEAH box helicase family protein [Thauera phenylacetica] | 05671 |
| mechanosensitive ion channel [Thauera phenylacetica] | 08476 |
| type II secretion system protein M, partial [Thauera phenylacetica] | 05350 |
| DUF45 domain-containing protein, partial [Thauera phenylacetica] | 09604 |
| HAMP domain-containing histidine kinase [Propionivibrio dicarboxylicus] | 07666 |
| type II toxin-antitoxin system VapC family toxin [Thauera phenylacetica] | 09454 |
| TolC family outer membrane protein [Azoarcus communis] | 09975 |
| hypothetical protein [Propionivibrio dicarboxylicus] | 01339 |
| hypothetical protein [Azoarcus communis] | 07556 |
| hypothetical protein [Azoarcus communis] | 07186 |
| DUF3077 domain-containing protein [Azoarcus communis] | 09225 |
| hypothetical protein [Azoarcus communis] | 03802 |
| PEP-CTERM sorting domain-containing protein [Azoarcus communis] | 01764 |
| hypothetical protein [Azoarcus communis] | 05115 |
| type II toxin-antitoxin system HipA family toxin [Thauera phenylacetica] | 05119 |
| hypothetical protein [Azoarcus communis] | 03671 |
| hypothetical protein [Azoarcus communis] | 08585 |
| hypothetical protein [Azoarcus communis] | 05717 |
| DUF2523 domain-containing protein [Azoarcus communis] | 00303 |
| LysR family transcriptional regulator, partial [Aromatoleum toluclasticum] | 05941 |
| enoyl-CoA hydratase-related protein, partial [Aromatoleum toluclasticum] | 03599 |
| helix-turn-helix transcriptional regulator [Azoarcus communis] | 02625 |
| biotin/lipoyl-binding protein, partial [Aromatoleum toluclasticum] | 05321 |
| ABC transporter permease subunit, partial [Aromatoleum toluclasticum] | 10613 |
| bifunctional ornithine acetyltransferase/N-acetylglutamate synthase, partial [Aromatoleum toluclasticum] | 07929 |
| hypothetical protein, partial [Aromatoleum toluclasticum] | 04140 |
| HEPN domain-containing protein [Azoarcus communis] | 10297 |
| hypothetical protein [Azoarcus communis] | 01171 |
| LuxR C-terminal-related transcriptional regulator, partial [Aromatoleum toluclasticum] | 05939 |
| hypothetical protein, partial [Aromatoleum toluclasticum] | 03975 |
| hypothetical protein, partial [Aromatoleum toluclasticum] | 01768 |
| hypothetical protein [Aromatoleum toluclasticum] | 05534 |
| hypothetical protein [Aromatoleum toluclasticum] | 04114 |
| hypothetical protein [Aromatoleum toluclasticum] | 08057 |
| hypothetical protein [Azoarcus communis] | 05231 |
| chitosanase [Aromatoleum toluclasticum] | 04331 |
| hypothetical protein [Azoarcus communis] | 02709 |
| hypothetical protein [Azoarcus communis] | 00536 |
| MULTISPECIES: SIR2 family protein [Rhodocyclales] | 01767 |
| MULTISPECIES: ANT(3'')-Ia family aminoglycoside nucleotidyltransferase AadA4 [Proteobacteria] | 02537 |
| hypothetical protein [Thauera phenylacetica] | 02535 |
| hypothetical protein [Thauera phenylacetica] | 10533 |
| Fic family protein [Thauera phenylacetica] | 10043 |
| DUF3375 domain-containing protein [Thauera phenylacetica] | 07742 |
| DUF4194 domain-containing protein [Thauera phenylacetica] | 05722 |
| DUF2220 family protein [Thauera phenylacetica] | 03066 |
| hypothetical protein [Thauera phenylacetica] | 02961 |
| HAD-IC family P-type ATPase, partial [Thauera phenylacetica] | 07741 |
| hypothetical protein [Thauera phenylacetica] | 01903 |
| lipopolysaccharide biosynthesis protein [Thauera phenylacetica] | 03630 |
| YgjV family protein [Propionivibrio dicarboxylicus] | 08918 |
| DUF1887 family CARF protein [Propionivibrio dicarboxylicus] | 07022 |
| hypothetical protein [Thauera phenylacetica] | 02007 |
| hypothetical protein [Thauera phenylacetica] | 00817 |
| prolyl oligopeptidase family serine peptidase [Propionivibrio dicarboxylicus] | 10271 |
| MULTISPECIES: hypothetical protein [Proteobacteria] | 00965 |
| GNAT family N-acetyltransferase [Propionivibrio dicarboxylicus] | 02220 |
| hypothetical protein [Thauera phenylacetica] | 06640 |

| | |
|---|---|
| hypothetical protein [Thauera phenylacetica] | 04307 |
| O-antigen ligase family protein [Thauera phenylacetica] | 06305 |
| ferrous iron transport protein A [Propionivibrio dicarboxylicus] | 09271 |
| hypothetical protein [Thauera phenylacetica] | 06367 |
| hypothetical protein [Thauera phenylacetica] | 02076 |
| GspH/FimT family protein [Propionivibrio dicarboxylicus] | 07370 |
| MULTISPECIES: hypothetical protein [Proteobacteria] | 00637 |
| MULTISPECIES: DUF456 family protein [Betaproteobacteria] | 06173 |
| type II toxin-antitoxin system VapC family toxin [Thauera phenylacetica] | 08088 |
| MULTISPECIES: hypothetical protein, partial [Zoogloeaceae] | 06462 |
| DUF5710 domain-containing protein [Thauera phenylacetica] | 04715 |
| MULTISPECIES: DUF115 domain-containing protein [Uliginosibacterium] | 09829 |
| hypothetical protein [Aromatoleum tolulyticum] | 07150 |
| hypothetical protein [Uliginosibacterium aquaticum] | 01246 |
| Fe-S-containing protein [Candidatus Dactylopiibacterium carminicum] | 05143 |
| sensor domain-containing diguanylate cyclase [Thauera chlorobenzoica] | 08167 |
| lipopolysaccharide biosynthesis protein [Uliginosibacterium aquaticum] | 07983 |
| hypothetical protein [Uliginosibacterium aquaticum] | 07960 |
| NAD(P)(+) transhydrogenase (Re/Si-specific) subunit beta, partial [Azoarcus indigens] | 01694 |
| hypothetical protein [Uliginosibacterium aquaticum] | 04899 |
| hypothetical protein [Uliginosibacterium aquaticum] | 08446 |
| hypothetical protein [Candidatus Dactylopiibacterium carminicum] | 07989 |
| hypothetical protein [Uliginosibacterium aquaticum] | 08319 |
| hypothetical protein [Candidatus Dactylopiibacterium carminicum] | 00718 |
| DUF917 domain-containing protein [Candidatus Dactylopiibacterium carminicum] | 07310 |
| hypothetical protein [Candidatus Dactylopiibacterium carminicum] | 06003 |
| elements of external origin [Candidatus Dactylopiibacterium carminicum] | 07765 |
| hypothetical protein [Candidatus Dactylopiibacterium carminicum] | 07030 |
| DUF637 domain-containing protein, partial [Azoarcus indigens] | 06042 |
| hypothetical protein [Candidatus Dactylopiibacterium carminicum] | 01250 |
| transposase [Candidatus Dactylopiibacterium carminicum] | 00638 |
| hypothetical protein [Candidatus Dactylopiibacterium carminicum] | 07311 |
| type II and III secretion system protein [Azoarcus indigens] | 00185 |
| hypothetical protein [Candidatus Dactylopiibacterium carminicum] | 01249 |
| type II toxin-antitoxin system Phd/YefM family antitoxin [Candidatus Dactylopiibacterium carminicum] | 07818 |
| siderophore ABC transporter substrate-binding protein [Candidatus Dactylopiibacterium carminicum] | 01499 |
| hypothetical protein [Candidatus Dactylopiibacterium carminicum] | 07029 |
| ATP-binding protein [Candidatus Dactylopiibacterium carminicum] | 00550 |
| tetratricopeptide repeat protein [Uliginosibacterium aquaticum] | 00695 |
| hypothetical protein [Uliginosibacterium aquaticum] | 09024 |
| hypothetical protein [Candidatus Dactylopiibacterium carminicum] | 03646 |
| polysaccharide lyase family 7 protein [Uliginosibacterium aquaticum] | 02772 |
| hypothetical protein [Candidatus Dactylopiibacterium carminicum] | 06177 |
| Hpt domain-containing protein [Uliginosibacterium aquaticum] | 09275 |
| hypothetical protein [Candidatus Dactylopiibacterium carminicum] | 02428 |
| hypothetical protein [Candidatus Dactylopiibacterium carminicum] | 07309 |
| YncE family protein [Candidatus Dactylopiibacterium carminicum] | 10392 |
| hypothetical protein [Candidatus Dactylopiibacterium carminicum] | 02431 |
| alpha/beta fold hydrolase [Uliginosibacterium aquaticum] | 01896 |
| zinc-dependent metalloprotease [Uliginosibacterium aquaticum] | 07970 |
| YHS domain-containing protein [Aromatoleum toluclasticum] | 02853 |
| hypothetical protein [Aromatoleum toluclasticum] | 06670 |
| hypothetical protein [Aromatoleum toluclasticum] | 10582 |
| nuclear transport factor 2 family protein [Aromatoleum toluclasticum] | 02146 |
| TerB N-terminal domain-containing protein [Aromatoleum toluclasticum] | 05761 |
| phospholipase D family protein [Aromatoleum toluclasticum] | 03171 |
| AAA family ATPase [Aromatoleum toluclasticum] | 09843 |
| hypothetical protein [Aromatoleum toluclasticum] | 08724 |
| hypothetical protein [Aromatoleum toluclasticum] | 09840 |

| | |
|---|---|
| hypothetical protein [Aromatoleum toluclasticum] | 03061 |
| hypothetical protein [Aromatoleum toluclasticum] | 08493 |
| TMEM175 family protein [Uliginosibacterium aquaticum] | 05882 |
| alpha/beta hydrolase-fold protein [Azoarcus indigens] | 09312 |
| nucleotidyltransferase family protein [Uliginosibacterium aquaticum] | 08920 |
| PilZ domain-containing protein [Thauera chlorobenzoica] | 08467 |
| PilZ domain-containing protein [Uliginosibacterium aquaticum] | 03509 |
| OmpA family protein [Uliginosibacterium aquaticum] | 02760 |
| alpha/beta hydrolase [Uliginosibacterium aquaticum] | 10017 |
| PepSY domain-containing protein [Azoarcus indigens] | 00901 |
| hypothetical protein [Uliginosibacterium aquaticum] | 03036 |
| hypothetical protein [Candidatus Dactylopiibacterium carminicum] | 06275 |
| hypothetical protein [Candidatus Dactylopiibacterium carminicum] | 08343 |
| hypothetical protein [Aromatoleum toluclasticum] | 05167 |
| DUF4258 domain-containing protein [Aromatoleum toluclasticum] | 02567 |
| hypothetical protein [Uliginosibacterium aquaticum] | 05688 |
| hypothetical protein [Aromatoleum toluclasticum] | 03374 |
| TIGR01777 family oxidoreductase [Uliginosibacterium aquaticum] | 04524 |
| hypothetical protein [Aromatoleum toluclasticum] | 05908 |
| hypothetical protein [Azoarcus toluclasticus] | 03488 |
| hypothetical protein [Candidatus Dactylopiibacterium carminicum] | 02232 |
| ATP-binding protein [Aromatoleum toluclasticum] | 00835 |
| hypothetical protein [Uliginosibacterium aquaticum] | 06519 |
| addiction module protein [Thauera chlorobenzoica] | 09334 |
| cytidine deaminase [Uliginosibacterium aquaticum] | 06755 |
| hypothetical protein [Azoarcus indigens] | 09109 |
| adenylate synthase [Candidatus Dactylopiibacterium carminicum] | 06538 |
| SMP-30/gluconolactonase/LRE family protein [Candidatus Dactylopiibacterium carminicum] | 10227 |
| ParB N-terminal domain-containing protein [Candidatus Dactylopiibacterium carminicum] | 06946 |
| ParB/RepB/Spo0J family partition protein [Candidatus Dactylopiibacterium carminicum] | 09182 |
| hypothetical protein [Uliginosibacterium aquaticum] | 10033 |
| hypothetical protein [Azoarcus indigens] | 03725 |
| DUF805 domain-containing protein [Candidatus Dactylopiibacterium carminicum] | 10176 |
| hypothetical protein [Candidatus Dactylopiibacterium carminicum] | 07304 |
| DUF6250 domain-containing protein [Uliginosibacterium aquaticum] | 07014 |
| DUF6511 domain-containing protein [Candidatus Dactylopiibacterium carminicum] | 06898 |
| phospholipid carrier-dependent glycosyltransferase [Uliginosibacterium aquaticum] | 01247 |
| phage holin family protein [Uliginosibacterium aquaticum] | 05258 |
| flagellar protein FlhE [Thauera chlorobenzoica] | 02612 |
| STAS domain-containing protein [Uliginosibacterium aquaticum] | 06174 |
| hypothetical protein [Uliginosibacterium aquaticum] | 07961 |
| MBL fold metallo-hydrolase [Thauera chlorobenzoica] | 05644 |
| hypothetical protein [Azoarcus indigens] | 09481 |
| hypothetical protein [Candidatus Dactylopiibacterium carminicum] | 00890 |
| DUF2917 domain-containing protein [Uliginosibacterium aquaticum] | 10031 |
| DUF4870 domain-containing protein [Candidatus Dactylopiibacterium carminicum] | 05045 |
| phosphatase PAP2 family protein [Uliginosibacterium aquaticum] | 00153 |
| xylose isomerase [Uliginosibacterium aquaticum] | 07323 |
| hypothetical protein [Azoarcus indigens] | 08195 |
| hypothetical protein [Uliginosibacterium aquaticum] | 02777 |
| hypothetical protein [Uliginosibacterium aquaticum] | 10161 |
| diguanylate cyclase [Uliginosibacterium aquaticum] | 06015 |
| S8 family serine peptidase [Candidatus Dactylopiibacterium carminicum] | 10308 |
| pectate lyase [Uliginosibacterium aquaticum] | 06157 |
| hypothetical protein [Thauera chlorobenzoica] | 08165 |
| hypothetical protein [Azoarcus indigens] | 10379 |
| hypothetical protein [Candidatus Dactylopiibacterium carminicum] | 03221 |
| hypothetical protein [Uliginosibacterium aquaticum] | 05581 |
| hypothetical protein, partial [Azoarcus indigens] | 08805 |

| | |
|---|---|
| hypothetical protein [Candidatus Dactylopiibacterium carminicum] | 02234 |
| hypothetical protein [Uliginosibacterium aquaticum] | 06714 |
| hypothetical protein [Uliginosibacterium aquaticum] | 04898 |
| hypothetical protein [Azoarcus indigens] | 05802 |
| carbohydrate binding domain-containing protein [Uliginosibacterium aquaticum] | 03039 |
| NINE protein [Candidatus Dactylopiibacterium carminicum] | 03166 |
| glycosyltransferase [Candidatus Dactylopiibacterium carminicum] | 09298 |
| hypothetical protein [Uliginosibacterium aquaticum] | 09279 |
| hypothetical protein [Uliginosibacterium aquaticum] | 09407 |
| porin [Thauera chlorobenzoica] | 03205 |
| hypothetical protein [Candidatus Dactylopiibacterium carminicum] | 08592 |
| hypothetical protein [Candidatus Dactylopiibacterium carminicum] | 06176 |
| hypothetical protein [Uliginosibacterium aquaticum] | 00284 |
| hypothetical protein [Candidatus Dactylopiibacterium carminicum] | 01248 |
| pectate lyase [Uliginosibacterium aquaticum] | 07982 |
| oxidoreductase-like protein [Uliginosibacterium aquaticum] | 03510 |
| hypothetical protein [Candidatus Dactylopiibacterium carminicum] | 05861 |
| O-antigen ligase family protein [Candidatus Dactylopiibacterium carminicum] | 00167 |
| hypothetical protein [Uliginosibacterium aquaticum] | 07142 |
| hypothetical protein [Uliginosibacterium aquaticum] | 08320 |
| NUDIX domain-containing protein [Candidatus Dactylopiibacterium carminicum] | 01242 |
| DUF2628 domain-containing protein [Uliginosibacterium aquaticum] | 03033 |
| hypothetical protein [Azoarcus indigens] | 00369 |
| NAD(P)H-binding protein [Candidatus Dactylopiibacterium carminicum] | 03054 |
| MliC family protein [Candidatus Dactylopiibacterium carminicum] | 07984 |
| HAMP domain-containing histidine kinase [Uliginosibacterium aquaticum] | 09406 |
| hypothetical protein [Candidatus Dactylopiibacterium carminicum] | 06541 |
| DUF5680 domain-containing protein [Candidatus Dactylopiibacterium carminicum] | 09721 |
| hypothetical protein [Candidatus Dactylopiibacterium carminicum] | 07028 |
| beta-propeller fold lactonase family protein [Candidatus Dactylopiibacterium carminicum] | 04161 |
| hypothetical protein [Candidatus Dactylopiibacterium carminicum] | 05719 |
| hypothetical protein [Candidatus Dactylopiibacterium carminicum] | 04702 |
| DUF202 domain-containing protein [Uliginosibacterium aquaticum] | 08437 |
| hypothetical protein [Candidatus Dactylopiibacterium carminicum] | 00279 |
| SPOR domain-containing protein [Uliginosibacterium aquaticum] | 06159 |
| PIN domain nuclease [Candidatus Dactylopiibacterium carminicum] | 08175 |
| 4Fe-4S dicluster domain-containing protein [Uliginosibacterium aquaticum] | 03536 |
| MBL fold metallo-hydrolase [Candidatus Dactylopiibacterium carminicum] | 06655 |
| hypothetical protein [Uliginosibacterium aquaticum] | 08626 |
| hypothetical protein [Aromatoleum toluclasticum] | 09305 |
| S41 family peptidase [Uliginosibacterium aquaticum] | 02564 |
| phosphopantetheine-binding protein [Candidatus Dactylopiibacterium carminicum] | 08198 |
| energy transducer TonB [Azoarcus indigens] | 09111 |
| hypothetical protein [Azoarcus indigens] | 02984 |
| cellulase family glycosylhydrolase [Uliginosibacterium aquaticum] | 00281 |
| hypothetical protein [Uliginosibacterium aquaticum] | 04523 |
| hypothetical protein [Uliginosibacterium aquaticum] | 00692 |
| Com family DNA-binding transcriptional regulator [Candidatus Dactylopiibacterium carminicum] | 05096 |
| hypothetical protein [Uliginosibacterium aquaticum] | 03038 |
| hypothetical protein [Thauera chlorobenzoica] | 07813 |
| hypothetical protein [Thauera chlorobenzoica] | 06864 |
| methyl-accepting chemotaxis protein, partial [Azoarcus indigens] | 05135 |
| MULTISPECIES: hypothetical protein [Uliginosibacterium] | 03506 |
| hypothetical protein [Uliginosibacterium aquaticum] | 07601 |
| hypothetical protein [Thauera chlorobenzoica] | 09438 |
| hypothetical protein [Uliginosibacterium aquaticum] | 07277 |
| AI-2E family transporter [Uliginosibacterium aquaticum] | 03524 |
| TlpA family protein disulfide reductase [Thauera terpenica] | 03664 |
| SoxR reducing system RseC family protein [Uliginosibacterium aquaticum] | 01892 |

| | |
|---|---|
| hypothetical protein [Thauera terpenica] | 03697 |
| YjfB family protein [Thauera chlorobenzoica] | 04687 |
| PAS domain-containing protein [Uliginosibacterium aquaticum] | 04526 |
| contractile injection system protein, VgrG/Pvc8 family [Azoarcus indigens] | 04728 |
| hypothetical protein [Thauera chlorobenzoica] | 06349 |
| hypothetical protein [Uliginosibacterium aquaticum] | 10503 |
| hypothetical protein [Thauera chlorobenzoica] | 07185 |
| hypothetical protein [Thauera terpenica] | 08789 |
| PD-(D/E)XK nuclease family protein [Thauera chlorobenzoica] | 07754 |
| hypothetical protein [Uliginosibacterium aquaticum] | 10158 |
| hypothetical protein [Uliginosibacterium aquaticum] | 07007 |
| hypothetical protein [Uliginosibacterium aquaticum] | 09955 |
| hypothetical protein [Uliginosibacterium aquaticum] | 01803 |
| cellulase family glycosylhydrolase [Uliginosibacterium aquaticum] | 09272 |
| retention module-containing protein [Thauera chlorobenzoica] | 02603 |
| hypothetical protein [Azoarcus indigens] | 06096 |
| hypothetical protein [Uliginosibacterium aquaticum] | 03160 |
| pilin [Azoarcus indigens] | 03235 |
| hypothetical protein [Thauera chlorobenzoica] | 07793 |
| conjugal transfer protein TraD [Thauera chlorobenzoica] | 08152 |
| hypothetical protein [Thauera chlorobenzoica] | 06708 |
| hypothetical protein [Uliginosibacterium aquaticum] | 00712 |
| hypothetical protein [Uliginosibacterium aquaticum] | 10524 |
| pseudouridine synthase [Uliginosibacterium aquaticum] | 10527 |
| hypothetical protein [Uliginosibacterium aquaticum] | 02412 |
| hypothetical protein [Uliginosibacterium aquaticum] | 05843 |
| FKBP-type peptidyl-prolyl cis-trans isomerase [Uliginosibacterium sp. IMCC34675] | 05842 |
| HEPN domain-containing protein [Uliginosibacterium aquaticum] | 01887 |
| hypothetical protein [Uliginosibacterium aquaticum] | 09723 |
| hypothetical protein [Azoarcus indigens] | 01083 |
| hypothetical protein [Uliginosibacterium aquaticum] | 05367 |
| hypothetical protein [Uliginosibacterium aquaticum] | 10507 |
| hypothetical protein [Thauera chlorobenzoica] | 09440 |
| hypothetical protein [Uliginosibacterium aquaticum] | 01622 |
| pectinesterase family protein [Uliginosibacterium aquaticum] | 04904 |
| carboxymuconolactone decarboxylase family protein [Uliginosibacterium aquaticum] | 05496 |
| DMT family transporter [Uliginosibacterium sp. IMCC34675] | 05847 |
| hypothetical protein [Thauera chlorobenzoica] | 04690 |
| hypothetical protein [Uliginosibacterium aquaticum] | 09753 |
| phosphopantetheine-binding protein [Uliginosibacterium aquaticum] | 03511 |
| hypothetical protein [Azoarcus indigens] | 05527 |
| lipoprotein [Uliginosibacterium aquaticum] | 02221 |
| ATP-dependent Clp protease proteolytic subunit [Thauera chlorobenzoica] | 02868 |
| STY4851/ECs_5259 family protein [Thauera terpenica] | 05662 |
| hypothetical protein [Thauera terpenica] | 03214 |
| hypothetical protein [Thauera terpenica] | 09898 |
| BRCT domain-containing protein [Thauera chlorobenzoica] | 07740 |
| hypothetical protein [Thauera chlorobenzoica] | 00842 |
| lipopolysaccharide biosynthesis protein [Uliginosibacterium aquaticum] | 01611 |
| DNA-binding protein [Candidatus Dactylopiibacterium carminicum] | 01783 |
| hypothetical protein [Thauera chlorobenzoica] | 00523 |
| hypothetical protein [Uliginosibacterium aquaticum] | 01476 |
| hypothetical protein [Uliginosibacterium aquaticum] | 05295 |
| toll/interleukin-1 receptor domain-containing protein [Aromatoleum toluclasticum] | 03650 |
| hypothetical protein [Aromatoleum toluclasticum] | 03329 |
| hypothetical protein [Azoarcus indigens] | 05455 |
| phage tail assembly protein [Azoarcus indigens] | 01261 |
| major capsid protein [Uliginosibacterium aquaticum] | 03031 |
| 5'-methylthioadenosine/adenosylhomocysteine nucleosidase [Uliginosibacterium aquaticum] | 07279 |

| | |
|---|---|
| helix-turn-helix domain-containing protein [Aromatoleum toluclasticum] | 03649 |
| hypothetical protein [Aromatoleum toluclasticum] | 09844 |
| hypothetical protein [Aromatoleum toluclasticum] | 00440 |
| ACT domain-containing protein [Uliginosibacterium aquaticum] | 01808 |
| hypothetical protein [Candidatus Dactylopiibacterium carminicum] | 04912 |
| hypothetical protein [Uliginosibacterium aquaticum] | 09719 |
| DUF4349 domain-containing protein [Uliginosibacterium aquaticum] | 05687 |
| insulinase family protein [Uliginosibacterium aquaticum] | 00059 |
| hypothetical protein [Uliginosibacterium aquaticum] | 01614 |
| hypothetical protein [Aromatoleum toluclasticum] | 01629 |
| hypothetical protein [Aromatoleum toluclasticum] | 00839 |
| DUF58 domain-containing protein [Uliginosibacterium aquaticum] | 09293 |
| hypothetical protein [Aromatoleum toluclasticum] | 03570 |
| Signal transduction histidine kinase [Candidatus Dactylopiibacterium carminicum] | 06175 |
| K(+)-transporting ATPase subunit F [Uliginosibacterium aquaticum] | 01472 |
| Hsp20 family protein [Uliginosibacterium aquaticum] | 01245 |
| phosphatase PAP2 family protein [Aromatoleum toluclasticum] | 00276 |
| SHOCT domain-containing protein [Aromatoleum toluclasticum] | 02029 |
| hypothetical protein [Thauera chlorobenzoica] | 07795 |
| hypothetical protein [Aromatoleum toluclasticum] | 05896 |
| DUF2835 family protein [Uliginosibacterium aquaticum] | 07285 |
| hypothetical protein [Aromatoleum toluclasticum] | 03364 |
| cupin domain-containing protein [Aromatoleum toluclasticum] | 07151 |
| SDR family NAD(P)-dependent oxidoreductase [Aromatoleum toluclasticum] | 04655 |
| NPP1 family protein [Uliginosibacterium aquaticum] | 00283 |
| hypothetical protein [Aromatoleum toluclasticum] | 07823 |
| hypothetical protein [Thauera terpenica] | 03360 |
| hypothetical protein [Thauera terpenica] | 03342 |
| TonB-dependent receptor [Azoarcus indigens] | 05454 |
| nuclear transport factor 2 family protein [Thauera terpenica] | 06322 |
| NTP transferase domain-containing protein [Thauera terpenica] | 08115 |
| hypothetical protein [Uliginosibacterium aquaticum] | 01471 |
| endo-1,4-beta-xylanase [Uliginosibacterium aquaticum] | 02759 |
| type II toxin-antitoxin system RelE/ParE family toxin [Thauera terpenica] | 02855 |
| hypothetical protein [Thauera terpenica] | 03330 |
| hypothetical protein [Thauera chlorobenzoica] | 07447 |
| pectate lyase [Uliginosibacterium aquaticum] | 10506 |
| HNH endonuclease [Thauera chlorobenzoica] | 03803 |
| hypothetical protein [Azoarcus indigens] | 02931 |
| hypothetical protein [Thauera terpenica] | 04176 |
| DUF2635 domain-containing protein [Azoarcus indigens] | 10195 |
| hypothetical protein [Thauera terpenica] | 09333 |
| hypothetical protein [Uliginosibacterium aquaticum] | 06829 |
| hypothetical protein [Thauera chlorobenzoica] | 09435 |
| family 43 glycosylhydrolase [Uliginosibacterium aquaticum] | 08448 |
| TonB-dependent receptor [Candidatus Dactylopiibacterium carminicum] | 02937 |
| IS5/IS1182 family transposase, partial [Thauera terpenica] | 04658 |
| DUF4124 domain-containing protein [Candidatus Dactylopiibacterium carminicum] | 00710 |
| hypothetical protein [Thauera terpenica] | 06672 |
| hypothetical protein [Thauera terpenica] | 08762 |
| DUF637 domain-containing protein, partial [Azoarcus indigens] | 02185 |
| hypothetical protein [Uliginosibacterium aquaticum] | 03151 |
| hypothetical protein [Thauera chlorobenzoica] | 05415 |
| DJ-1/PfpI family protein [Candidatus Dactylopiibacterium carminicum] | 05257 |
| hypothetical protein [Propionivibrio dicarboxylicus] | 03900 |
| RHS repeat-associated core domain-containing protein, partial [Azoarcus sp. L1K30] | 04972 |
| RloB family protein [Rhodocyclus purpureus] | 01788 |
| geranylgeranyl reductase family protein [Rhodocyclus purpureus] | 09638 |
| RES family NAD+ phosphorylase [Rhodocyclus purpureus] | 08780 |

| | |
|---|---|
| hypothetical protein [Propionivibrio dicarboxylicus] | 03908 |
| NAD-binding protein [Propionivibrio dicarboxylicus] | 10011 |
| helix-turn-helix transcriptional regulator [Propionivibrio dicarboxylicus] | 01987 |
| fused MFS/spermidine synthase [Azoarcus sp. L1K30] | 06171 |
| NAD-dependent malic enzyme [Propionivibrio dicarboxylicus] | 09836 |
| hypothetical protein [Azoarcus sp. L1K30] | 08030 |
| helix-turn-helix domain-containing protein [Thauera phenylacetica] | 09390 |
| metallophosphoesterase [Azoarcus sp. L1K30] | 07605 |
| methyltransferase domain-containing protein [Azoarcus sp. L1K30] | 02481 |
| hypothetical protein [Azoarcus sp. L1K30] | 04971 |
| ATP-binding protein [Rhodocyclus purpureus] | 03531 |
| exo-alpha-sialidase [Azoarcus sp. L1K30] | 08375 |
| hypothetical protein [Propionivibrio dicarboxylicus] | 05253 |
| helix-turn-helix domain-containing protein [Propionivibrio dicarboxylicus] | 07677 |
| efflux transporter outer membrane subunit [Propionivibrio dicarboxylicus] | 05946 |
| hypothetical protein, partial [Azoarcus communis] | 02918 |
| transposase, partial [Azoarcus communis] | 04377 |
| DUF3313 domain-containing protein [Rhodocyclus purpureus] | 08641 |
| hypothetical protein [Thauera phenylacetica] | 08095 |
| hypothetical protein, partial [Azoarcus communis] | 07202 |
| DDE-type integrase/transposase/recombinase [Rhodocyclus purpureus] | 08357 |
| RHS domain-containing protein [Azoarcus communis] | 01685 |
| hypothetical protein [Propionivibrio dicarboxylicus] | 09449 |
| hypothetical protein [Azoarcus communis] | 02628 |
| helix-turn-helix domain-containing protein, partial [Thauera phenylacetica] | 03300 |
| hypothetical protein [Rhodocyclus purpureus] | 06553 |
| glycosyltransferase family 2 protein [Rhodocyclus purpureus] | 02788 |
| citrate lyase holo-[acyl-carrier protein] synthase [Propionivibrio dicarboxylicus] | 04610 |
| hypothetical protein [Rhodocyclus purpureus] | 08241 |
| citrate lyase acyl carrier protein [Propionivibrio dicarboxylicus] | 08061 |
| 2-hydroxycarboxylate transporter family protein [Propionivibrio dicarboxylicus] | 03781 |
| fibronectin type III domain-containing protein [Thauera humireducens] | 02041 |
| transposase, partial [Azoarcus communis] | 05440 |
| hypothetical protein, partial [Azoarcus communis] | 03709 |
| serine protease [Rhodocyclus purpureus] | 01674 |
| hypothetical protein, partial [Propionivibrio dicarboxylicus] | 06784 |
| hypothetical protein [Azoarcus communis] | 07049 |
| hypothetical protein [Azoarcus communis] | 03077 |
| hypothetical protein [Azoarcus communis] | 00885 |
| GNAT family acetyltransferase [Propionivibrio dicarboxylicus] | 03783 |
| MoaD/ThiS family protein [Propionivibrio dicarboxylicus] | 07709 |
| TRAP transporter small permease subunit [Propionivibrio dicarboxylicus] | 02505 |
| YhcH/YjgK/YiaL family protein [Propionivibrio dicarboxylicus] | 06298 |
| hypothetical protein [Azoarcus communis] | 03220 |
| hypothetical protein, partial [Azoarcus communis] | 00351 |
| pentapeptide repeat-containing protein [Rhodocyclus purpureus] | 09732 |
| hypothetical protein [Uliginosibacterium aquaticum] | 06208 |
| RHS repeat-associated core domain-containing protein, partial [Azoarcus communis] | 03383 |
| hypothetical protein [Azoarcus communis] | 03708 |
| ATP-binding cassette domain-containing protein, partial [Azoarcus communis] | 00028 |
| RHS repeat-associated core domain-containing protein, partial [Azoarcus communis] | 04712 |
| hypothetical protein, partial [Azoarcus communis] | 01068 |
| hypothetical protein [Rhodocyclus purpureus] | 00398 |
| hypothetical protein [Propionivibrio dicarboxylicus] | 09366 |
| metallophosphoesterase [Rhodocyclus purpureus] | 05159 |
| FlxA-like family protein [Propionivibrio dicarboxylicus] | 00287 |
| class IV adenylate cyclase [Propionivibrio dicarboxylicus] | 06656 |
| tripartite tricarboxylate transporter TctB family protein [Propionivibrio dicarboxylicus] | 04612 |
| SDR family oxidoreductase [Rhodocyclus purpureus] | 05109 |

| | |
|---|---|
| terminase [Propionivibrio dicarboxylicus] | 05046 |
| hypothetical protein [Propionivibrio dicarboxylicus] | 09220 |
| outer membrane beta-barrel protein [Rhodocyclus purpureus] | 05592 |
| outer membrane beta-barrel protein [Rhodocyclus purpureus] | 00874 |
| SLBB domain-containing protein [Propionivibrio dicarboxylicus] | 07111 |
| Ig-like domain-containing protein [Propionivibrio dicarboxylicus] | 07477 |
| UvrD-helicase domain-containing protein [Propionivibrio dicarboxylicus] | 02809 |
| STAS domain-containing protein [Rhodocyclus purpureus] | 04058 |
| restriction endonuclease subunit S [Thauera phenylacetica] | 01934 |
| hypothetical protein [Rhodocyclus purpureus] | 06725 |
| hypothetical protein [Rhodocyclus purpureus] | 00178 |
| chemotaxis protein CheX [Propionivibrio dicarboxylicus] | 10118 |
| heparinase II/III family protein [Candidatus Dactylopiibacterium carminicum] | 04911 |
| hypothetical protein [Propionivibrio dicarboxylicus] | 09473 |
| BON domain-containing protein [Rhodocyclus purpureus] | 04646 |
| hypothetical protein [Propionivibrio dicarboxylicus] | 02844 |
| hypothetical protein [Propionivibrio dicarboxylicus] | 08243 |
| DUF4145 domain-containing protein [Thauera humireducens] | 05062 |
| hydrogenase maturation protease [Rhodocyclus purpureus] | 00736 |
| retention module-containing protein [Rhodocyclus purpureus] | 09308 |
| aspartate 1-decarboxylase [Propionivibrio dicarboxylicus] | 03944 |
| hypothetical protein [Thauera humireducens] | 05063 |
| hypothetical protein [Thauera humireducens] | 04667 |
| hypothetical protein [Thauera humireducens] | 01017 |
| hypothetical protein [Propionivibrio dicarboxylicus] | 10119 |
| hypothetical protein [Rhodocyclus purpureus] | 08689 |
| hypothetical protein [Rhodocyclus purpureus] | 09731 |
| tetratricopeptide repeat protein [Propionivibrio dicarboxylicus] | 01296 |
| filamentous hemagglutinin N-terminal domain-containing protein [Thauera phenylacetica] | 10194 |
| DUF4376 domain-containing protein [Rhodocyclus purpureus] | 08940 |
| Dabb family protein [Uliginosibacterium aquaticum] | 06521 |
| hypothetical protein [Propionivibrio dicarboxylicus] | 05146 |
| PD-(D/E)XK nuclease-like domain-containing protein [Rhodocyclus purpureus] | 07998 |
| class I SAM-dependent methyltransferase [Rhodocyclus purpureus] | 04746 |
| aminopeptidase [Candidatus Dactylopiibacterium carminicum] | 06540 |
| type II secretion system GspH family protein [Thauera lacus] | 07189 |
| hypothetical protein [Aromatoleum toluclasticum] | 03327 |
| DUF2917 domain-containing protein [Aromatoleum diolicum] | 06491 |
| glycosyltransferase family 4 protein [Azoarcus sp. KH32C] | 00977 |
| metallophosphoesterase [Azoarcus sp. KH32C] | 09286 |
| hypothetical protein [Azoarcus sp. KH32C] | 01268 |
| cupin domain-containing protein [Aromatoleum toluclasticum] | 06836 |
| hypothetical protein [Azoarcus sp. KH32C] | 04384 |
| glycosyltransferase [Aromatoleum diolicum] | 02725 |
| hypothetical protein [Aromatoleum toluclasticum] | 04548 |
| DUF2513 domain-containing protein [Azoarcus communis] | 05800 |
| hypothetical protein [Niveibacterium sp. COAC-50] | 06530 |
| hypothetical protein [Niveibacterium sp. COAC-50] | 07172 |
| DUF2357 domain-containing protein [Niveibacterium sp. COAC-50] | 02050 |
| NAD(P)-binding protein, partial [Aromatoleum diolicum] | 06135 |
| hypothetical protein [Aromatoleum toluclasticum] | 06167 |
| hypothetical protein, partial [Aromatoleum toluclasticum] | 04657 |
| MULTISPECIES: hypothetical protein [Thauera] | 08250 |
| hypothetical protein [Aromatoleum diolicum] | 10478 |
| transposase, partial [Aromatoleum diolicum] | 09248 |
| hypothetical protein [Aromatoleum toluclasticum] | 03322 |
| hypothetical protein [Thauera propionica] | 09442 |
| hypothetical protein [Candidatus Dactylopiibacterium carminicum] | 05259 |
| hypothetical protein [Azoarcus sp. DD4] | 09154 |

| | |
|---|---|
| hypothetical protein [Azoarcus sp. DD4] | 00923 |
| hypothetical protein [Aromatoleum aromaticum] | 05338 |
| hypothetical protein, partial [Aromatoleum diolicum] | 06207 |
| hypothetical protein [Aromatoleum diolicum] | 03392 |
| hypothetical protein [Thauera linaloolentis] | 08123 |
| DUF1329 domain-containing protein, partial [Aromatoleum diolicum] | 04499 |
| hypothetical protein, partial [Aromatoleum diolicum] | 07587 |
| SNF2-related protein [Azoarcus sp. DD4] | 04037 |
| hypothetical protein [Thauera linaloolentis] | 03798 |
| MULTISPECIES: hypothetical protein [Proteobacteria] | 05419 |
| helix-turn-helix domain-containing protein [Thauera propionica] | 01660 |
| DNA (cytosine-5-)-methyltransferase [Aromatoleum toluclasticum] | 05935 |
| metallophosphoesterase [Aromatoleum toluclasticum] | 00499 |
| phytochelatin synthase family protein [Aromatoleum diolicum] | 07248 |
| transposase, partial [Thauera terpenica] | 06675 |
| Imm50 family immunity protein [Thauera linaloolentis] | 01236 |
| hypothetical protein [Zoogloea oleivorans] | 06090 |
| hypothetical protein [Aromatoleum bremense] | 05905 |
| 2-oxoacid:acceptor oxidoreductase subunit alpha [Aromatoleum bremense] | 08667 |
| VENN motif pre-toxin domain-containing protein, partial [Azoarcus communis] | 09232 |
| hypothetical protein [Zoogloea oleivorans] | 08840 |
| hypothetical protein [Zoogloea oleivorans] | 07528 |
| hypothetical protein [Zoogloea oleivorans] | 06940 |
| hypothetical protein [Zoogloea oleivorans] | 09947 |
| helix-turn-helix domain-containing protein [Zoogloea oleivorans] | 01140 |
| transposase, partial [Thauera terpenica] | 09847 |
| hypothetical protein [Zoogloea oleivorans] | 04824 |
| IS1634 family transposase, partial [Thauera terpenica] | 05701 |
| hypothetical protein [Zoogloea oleivorans] | 05765 |
| hypothetical protein [Zoogloea oleivorans] | 09954 |
| NAD(P)-binding protein, partial [Thauera terpenica] | 08118 |
| hypothetical protein [Aromatoleum toluolicum] | 05545 |
| ABC transporter substrate-binding protein [Aromatoleum toluolicum] | 09565 |
| DUF4351 domain-containing protein [Aromatoleum bremense] | 01961 |
| MULTISPECIES: hypothetical protein [Proteobacteria] | 08664 |
| hypothetical protein [Azospira oryzae] | 04961 |
| hypothetical protein [Azospira oryzae] | 06585 |
| hypothetical protein [Thauera terpenica] | 09849 |
| hypothetical protein [Azoarcus olearius] | 02880 |
| transposase, partial [Aromatoleum buckelii] | 00633 |
| HEPN domain-containing protein [Aromatoleum buckelii] | 00634 |
| hypothetical protein [Azoarcus communis] | 01424 |
| alcohol dehydrogenase catalytic domain-containing protein, partial [Azoarcus communis] | 04931 |
| hypothetical protein [Aromatoleum buckelii] | 03451 |
| hypothetical protein [Aromatoleum buckelii] | 04842 |
| HTH domain-containing protein, partial [Aromatoleum buckelii] | 06461 |
| PRTRC system protein D, partial [Aromatoleum buckelii] | 09213 |
| hypothetical protein, partial [Aromatoleum buckelii] | 08271 |
| DNA/RNA non-specific endonuclease, partial [Aromatoleum buckelii] | 07222 |
| hypothetical protein [Thauera terpenica] | 04337 |
| MULTISPECIES: hypothetical protein [Azospira] | 05024 |
| antA/AntB antirepressor family protein [Azospira oryzae] | 06037 |
| MFS transporter [Candidatus Dactylopiibacterium carminicum] | 01922 |
| GNAT family N-acetyltransferase [Candidatus Dactylopiibacterium carminicum] | 09301 |
| type II secretion system GspH family protein [Thauera lacus] | 10350 |
| TonB-dependent receptor [Azoarcus nasutitermitis] | 08517 |
| PilZ domain-containing protein [Candidatus Dactylopiibacterium carminicum] | 02422 |
| M23 family metallopeptidase [Thauera phenylacetica] | 03629 |
| DUF4852 domain-containing protein [Azoarcus nasutitermitis] | 00569 |

| | |
|---|---|
| glycosyltransferase family 2 protein [Azoarcus nasutitermitis] | 05140 |
| hypothetical protein [Thauera aromatica] | 06554 |
| hypothetical protein [Azoarcus nasutitermitis] | 06388 |
| YqaE/Pmp3 family membrane protein [Azoarcus nasutitermitis] | 08514 |
| HNH endonuclease [Thauera aromatica] | 05285 |
| ribonuclease [Thauera phenylacetica] | 00175 |
| hypothetical protein [Azoarcus nasutitermitis] | 00907 |
| retention module-containing protein, partial [Azoarcus nasutitermitis] | 01344 |
| hypothetical protein [Thauera phenylacetica] | 07021 |
| hypothetical protein [Niveibacterium umoris] | 00610 |
| MerR family transcriptional regulator [Niveibacterium umoris] | 01525 |
| hypothetical protein [Niveibacterium umoris] | 07628 |
| ferritin-like domain-containing protein [Niveibacterium umoris] | 05243 |
| S-layer family protein [Azoarcus nasutitermitis] | 00370 |
| hypothetical protein [Thauera phenylacetica] | 07386 |
| hypothetical protein [Azoarcus nasutitermitis] | 05138 |
| hypothetical protein [Azoarcus nasutitermitis] | 07632 |
| toll/interleukin-1 receptor domain-containing protein [Rhodocyclus purpureus] | 08938 |
| J domain-containing protein [Rhodocyclus purpureus] | 05766 |
| toxin-activating lysine-acyltransferase [Propionivibrio dicarboxylicus] | 04663 |
| rhomboid family intramembrane serine protease [Candidatus Dactylopiibacterium carminicum] | 01239 |
| hypothetical protein [Thauera lacus] | 02903 |
| hypothetical protein [Thauera lacus] | 07458 |
| hypothetical protein [Propionivibrio dicarboxylicus] | 00802 |
| hypothetical protein [Thauera lacus] | 03369 |
| DUF6396 domain-containing protein [Thauera lacus] | 05107 |
| STAS domain-containing protein [Thauera lacus] | 04217 |
| glycosyltransferase [Rhodocyclus purpureus] | 01052 |
| pilin [Propionivibrio dicarboxylicus] | 00343 |
| OsmC family protein [Rhodocyclus purpureus] | 08638 |
| alpha/beta hydrolase [Rhodocyclus purpureus] | 06163 |
| hypothetical protein [Azoarcus nasutitermitis] | 04069 |
| SPFH domain-containing protein [Niveibacterium umoris] | 10069 |
| metallophosphoesterase [Niveibacterium umoris] | 04313 |
| phospholipid:lipid A palmitoyltransferase [Candidatus Dactylopiibacterium carminicum] | 09606 |
| glycine--tRNA ligase subunit alpha, partial [Aromatoleum evansii] | 04852 |
| autotransporter serine protease [Thauera aromatica] | 02436 |
| helix-turn-helix domain-containing protein [Candidatus Dactylopiibacterium carminicum] | 04046 |
| VOC family protein [Thauera linaloolentis] | 05770 |
| DNA mismatch repair protein MutS [Candidatus Dactylopiibacterium carminicum] | 04543 |
| tetratricopeptide repeat protein [Thauera linaloolentis] | 06557 |
| hypothetical protein [Thauera aromatica] | 04053 |
| hypothetical protein [Candidatus Dactylopiibacterium carminicum] | 02426 |
| surface lipoprotein assembly modifier, partial [Thauera linaloolentis] | 00506 |
| MULTISPECIES: phospholipase D-like domain-containing protein [Betaproteobacteria] | 08650 |
| hypothetical protein [Azoarcus communis] | 10462 |
| S9 family peptidase [Candidatus Dactylopiibacterium carminicum] | 01921 |
| nitronate monooxygenase, partial [Aromatoleum evansii] | 05228 |
| hypothetical protein [Azoarcus communis] | 01172 |
| efflux RND transporter periplasmic adaptor subunit, partial [Aromatoleum evansii] | 10460 |
| DUF4391 domain-containing protein [Azoarcus sp. DD4] | 01232 |
| hypothetical protein [Azoarcus communis] | 07920 |
| hypothetical protein [Aromatoleum toluclasticum] | 05320 |
| hypothetical protein, partial [Aromatoleum toluclasticum] | 03125 |
| hypothetical protein [Aromatoleum evansii] | 09544 |
| hypothetical protein [Aromatoleum evansii] | 09218 |
| hypothetical protein [Niveibacterium umoris] | 06865 |
| hypothetical protein [Niveibacterium umoris] | 06431 |
| pilin [Niveibacterium umoris] | 02464 |

| | |
|---|---|
| DUF1254 domain-containing protein [Niveibacterium umoris] | 07650 |
| hypothetical protein [Niveibacterium umoris] | 01957 |
| YciI family protein [Candidatus Dactylopiibacterium carminicum] | 02775 |
| hypothetical protein [Thauera aromatica] | 01505 |
| phosphotransferase, partial [Aromatoleum toluclasticum] | 01328 |
| HAD family hydrolase [Aromatoleum evansii] | 06532 |
| hypothetical protein [Aromatoleum evansii] | 07913 |
| tetratricopeptide repeat protein [Candidatus Dactylopiibacterium carminicum] | 02225 |
| DUF5801 domain-containing protein, partial [Aromatoleum toluclasticum] | 04606 |
| cupin domain-containing protein [Propionivibrio dicarboxylicus] | 06258 |
| TRAP transporter small permease [Aromatoleum evansii] | 01410 |
| hypothetical protein [Thauera aromatica] | 09080 |
| toprim domain-containing protein [Azoarcus communis] | 05603 |
| type III secretion system chaperone [Azoarcus rhizosphaerae] | 01097 |
| L-ascorbate 6-phosphate lactonase [Propionivibrio dicarboxylicus] | 05975 |
| alpha/beta fold hydrolase [Uliginosibacterium gangwonense] | 00043 |
| hypothetical protein [Uliginosibacterium gangwonense] | 00751 |
| pre-peptidase C-terminal domain-containing protein, partial [Uliginosibacterium gangwonense] | 07760 |
| reverse transcriptase family protein [Uliginosibacterium gangwonense] | 03159 |
| ATP-binding cassette domain-containing protein, partial [Azoarcus taiwanensis] | 00125 |
| ribonuclease E inhibitor RraB [Uliginosibacterium gangwonense] | 06824 |
| hypothetical protein [Uliginosibacterium gangwonense] | 05044 |
| hypothetical protein [Thauera linaloolentis] | 10029 |
| N-acetylglucosamine-6-phosphate deacetylase [Uliginosibacterium gangwonense] | 09835 |
| class I SAM-dependent methyltransferase [Azoarcus taiwanensis] | 02771 |
| class I SAM-dependent methyltransferase [Uliginosibacterium aquaticum] | 00204 |
| arabinogalactan endo-1,4-beta-galactosidase [Uliginosibacterium gangwonense] | 05047 |
| MAPEG family protein [Uliginosibacterium aquaticum] | 01532 |
| phosphatase PAP2 family protein [Uliginosibacterium gangwonense] | 08436 |
| hypothetical protein, partial [Aromatoleum tolulyticum] | 00944 |
| hypothetical protein [Uliginosibacterium gangwonense] | 03779 |
| DUF2950 family protein [Uliginosibacterium aquaticum] | 01483 |
| hypothetical protein [Azoarcus taiwanensis] | 01855 |
| helix-turn-helix domain-containing GNAT family N-acetyltransferase [Uliginosibacterium gangwonense] | 08105 |
| maltose/maltodextrin ABC transporter substrate-binding protein MalE [Uliginosibacterium gangwonense] | 06831 |
| hexameric tyrosine-coordinated heme protein [Thauera terpenica] | 05700 |
| hypothetical protein [Aromatoleum aromaticum] | 05660 |
| formate hydrogenlyase maturation HycH family protein [Uliginosibacterium gangwonense] | 08625 |
| SPFH domain-containing protein [Uliginosibacterium gangwonense] | 03517 |
| glutaredoxin family protein [Uliginosibacterium gangwonense] | 00832 |
| T6SS immunity protein Tli4 family protein [Aromatoleum tolulyticum] | 05013 |
| hypothetical protein [Aromatoleum tolulyticum] | 10209 |
| transposase, partial [Zoogloea dura] | 05555 |
| hypothetical protein [Uliginosibacterium gangwonense] | 01533 |
| hypothetical protein [Uliginosibacterium gangwonense] | 04651 |
| beta-galactosidase [Uliginosibacterium gangwonense] | 07051 |
| DUF1302 family protein, partial [Zoogloea dura] | 09699 |
| midcut-by-XrtH protein [Zoogloea dura] | 05835 |
| beta-galactosidase [Uliginosibacterium gangwonense] | 01620 |
| UDP-glucose--hexose-1-phosphate uridylyltransferase [Uliginosibacterium gangwonense] | 06740 |
| galactokinase [Uliginosibacterium gangwonense] | 06749 |
| hypothetical protein [Uliginosibacterium gangwonense] | 00285 |
| hypothetical protein [Uliginosibacterium gangwonense] | 01619 |
| lactate utilization protein [Azoarcus taiwanensis] | 01390 |
| BamA/TamA family outer membrane protein [Uliginosibacterium aquaticum] | 10020 |
| phage tail assembly chaperone [Uliginosibacterium gangwonense] | 02561 |
| NAD(P)/FAD-dependent oxidoreductase, partial [Azoarcus taiwanensis] | 03467 |
| IPTL-CTERM sorting domain-containing protein [Zoogloea dura] | 08309 |
| helix-turn-helix domain-containing protein, partial [Zoogloea dura] | 08614 |

| | |
|---|---|
| hypothetical protein [Thauera linaloolentis] | 03098 |
| alpha/beta fold hydrolase [Zoogloea dura] | 04888 |
| hypothetical protein [Aromatoleum aromaticum] | 07040 |
| ATP-binding protein [Zoogloea dura] | 10009 |
| hypothetical protein [Azoarcus taiwanensis] | 00653 |
| hypothetical protein [Azoarcus taiwanensis] | 08378 |
| PAS domain-containing transcriptional regulator [Azoarcus taiwanensis] | 03712 |
| hypothetical protein [Propionivibrio dicarboxylicus] | 00581 |
| glycosyltransferase [Uliginosibacterium gangwonense] | 06731 |
| methyltransferase domain-containing protein [Uliginosibacterium gangwonense] | 09287 |
| hypothetical protein [Uliginosibacterium gangwonense] | 02023 |
| DMT family transporter [Propionivibrio dicarboxylicus] | 08528 |
| hypothetical protein [Thauera linaloolentis] | 00295 |
| hypothetical protein [Uliginosibacterium gangwonense] | 05041 |
| hypothetical protein [Uliginosibacterium gangwonense] | 09404 |
| hypothetical protein [Uliginosibacterium gangwonense] | 05575 |
| AraC family transcriptional regulator [Uliginosibacterium gangwonense] | 00280 |
| 23S rRNA (adenine(1618)-N(6))-methyltransferase RlmF [Uliginosibacterium aquaticum] | 09101 |
| hypothetical protein [Uliginosibacterium gangwonense] | 04327 |
| thioredoxin family protein [Uliginosibacterium gangwonense] | 08022 |
| [FeFe] hydrogenase H-cluster maturation GTPase HydF [Propionivibrio dicarboxylicus] | 10261 |
| hypothetical protein [Uliginosibacterium gangwonense] | 01911 |
| Nramp family divalent metal transporter [Uliginosibacterium gangwonense] | 06192 |
| [FeFe] hydrogenase, group A [Propionivibrio dicarboxylicus] | 01330 |
| hypothetical protein [Aromatoleum petrolei] | 06762 |
| 6-phosphogluconolactonase [Propionivibrio dicarboxylicus] | 03158 |
| L-arabinose isomerase [Uliginosibacterium gangwonense] | 06741 |
| M13 family metallopeptidase [Uliginosibacterium gangwonense] | 03076 |
| RebB family R body protein [Uliginosibacterium gangwonense] | 09830 |
| energy-coupling factor ABC transporter substrate-binding protein [Propionivibrio dicarboxylicus] | 07405 |
| energy-coupling factor transporter transmembrane protein EcfT [Propionivibrio dicarboxylicus] | 02633 |
| HP1 family phage holin [Uliginosibacterium gangwonense] | 07410 |
| flippase-like domain-containing protein [Zoogloea dura] | 05576 |
| hypothetical protein [Thauera linaloolentis] | 03796 |
| transporter substrate-binding domain-containing protein [Uliginosibacterium gangwonense] | 04159 |
| zinc ribbon domain-containing protein [Uliginosibacterium gangwonense] | 02555 |
| SHOCT domain-containing protein [Uliginosibacterium gangwonense] | 03315 |
| TonB-dependent receptor plug domain-containing protein, partial [Azoarcus taiwanensis] | 09740 |
| hypothetical protein [Uliginosibacterium gangwonense] | 03548 |
| hypothetical protein [Propionivibrio dicarboxylicus] | 07671 |
| glycoside hydrolase family 16 protein [Uliginosibacterium gangwonense] | 04162 |
| anaerobic C4-dicarboxylate transporter [Propionivibrio dicarboxylicus] | 02909 |
| class I SAM-dependent methyltransferase [Azoarcus taiwanensis] | 00430 |
| hypothetical protein, partial [Azoarcus taiwanensis] | 06484 |
| hypothetical protein, partial [Azoarcus taiwanensis] | 06485 |
| hypothetical protein, partial [Zoogloea dura] | 04892 |
| hypothetical protein [Uliginosibacterium gangwonense] | 02024 |
| hypothetical protein [Uliginosibacterium gangwonense] | 00994 |
| hypothetical protein [Uliginosibacterium gangwonense] | 06657 |
| hypothetical protein [Propionivibrio dicarboxylicus] | 07995 |
| anaerobic ribonucleoside-triphosphate reductase [Azoarcus taiwanensis] | 07931 |
| UxaA family hydrolase [Propionivibrio dicarboxylicus] | 09362 |
| fructose-specific PTS transporter subunit EIIC [Uliginosibacterium gangwonense] | 10072 |
| MULTISPECIES: hypothetical protein [Proteobacteria] | 08001 |
| hydroxylamine reductase [Propionivibrio dicarboxylicus] | 05593 |
| hypothetical protein, partial [Zoogloea dura] | 02751 |
| SCO family protein [Aromatoleum toluvorans] | 01779 |
| hemagglutinin repeat-containing protein [Azoarcus rhizosphaerae] | 10398 |
| HNH endonuclease, partial [Azoarcus rhizosphaerae] | 02655 |

| | |
|---|---|
| hypothetical protein [Thauera chlorobenzoica] | 09881 |
| hypothetical protein [Zoogloea dura] | 10372 |
| hypothetical protein [Zoogloea dura] | 01147 |
| DUF1566 domain-containing protein [Azoarcus rhizosphaerae] | 08523 |
| hypothetical protein [Uliginosibacterium aquaticum] | 04031 |
| ester cyclase [Aromatoleum toluvorans] | 06486 |
| hypothetical protein [Uliginosibacterium gangwonense] | 00707 |
| endo alpha-1,4 polygalactosaminidase [Uliginosibacterium gangwonense] | 06172 |
| YcxB family protein [Zoogloea dura] | 03869 |
| MFS transporter [Azoarcus rhizosphaerae] | 02947 |
| HrpE/YscL family type III secretion apparatus protein [Azoarcus rhizosphaerae] | 02654 |
| tetratricopeptide repeat protein [Azoarcus rhizosphaerae] | 01348 |
| hypothetical protein [Zoogloea dura] | 09270 |
| 2OG-Fe dioxygenase family protein [Azoarcus rhizosphaerae] | 01938 |
| hypothetical protein [Thauera chlorobenzoica] | 02606 |
| fibronectin type III domain-containing protein [Uliginosibacterium gangwonense] | 03947 |
| pullulanase-type alpha-1,6-glucosidase [Uliginosibacterium gangwonense] | 00496 |
| hypothetical protein [Azoarcus rhizosphaerae] | 03409 |
| hypothetical protein [Uliginosibacterium aquaticum] | 03872 |
| helix-turn-helix domain-containing protein, partial [Zoogloea dura] | 02409 |
| HipA domain-containing protein [Aromatoleum tolulyticum] | 03611 |
| DUF4041 domain-containing protein [Thauera chlorobenzoica] | 05088 |
| hypothetical protein [Uliginosibacterium sp. IMCC34675] | 05852 |
| helix-turn-helix domain-containing protein [Azoarcus rhizosphaerae] | 00061 |
| hypothetical protein [Uliginosibacterium aquaticum] | 01487 |
| hypothetical protein [Uliginosibacterium aquaticum] | 07274 |
| hypothetical protein [Uliginosibacterium aquaticum] | 08662 |
| hypothetical protein [Azoarcus rhizosphaerae] | 06396 |
| hypothetical protein [Azoarcus rhizosphaerae] | 00062 |
| hypothetical protein [Thauera chlorobenzoica] | 06610 |
| hypothetical protein [Azoarcus rhizosphaerae] | 07493 |
| hypothetical protein, partial [Aromatoleum toluvorans] | 07245 |
| TIGR04141 family sporadically distributed protein [Uliginosibacterium aquaticum] | 03868 |
| hypothetical protein [Uliginosibacterium aquaticum] | 07293 |
| DUF1800 domain-containing protein [Uliginosibacterium aquaticum] | 04087 |
| MULTISPECIES: hypothetical protein [Uliginosibacterium] | 01488 |
| pyrroline-5-carboxylate reductase [Aromatoleum toluvorans] | 04493 |
| hypothetical protein [Uliginosibacterium aquaticum] | 09580 |
| hypothetical protein [Azoarcus rhizosphaerae] | 08519 |
| helix-turn-helix domain containing protein, partial [Aromatoleum toluvorans] | 05535 |
| GNAT family N-acetyltransferase [Zoogloea dura] | 06513 |
| hypothetical protein [Thauera chlorobenzoica] | 02887 |
| hypothetical protein [Uliginosibacterium aquaticum] | 06733 |
| anti-virulence regulator CigR family protein [Uliginosibacterium aquaticum] | 05845 |
| hypothetical protein [Azoarcus rhizosphaerae] | 05149 |
| ABC transporter substrate-binding protein [Azoarcus rhizosphaerae] | 02949 |
| hypothetical protein [Uliginosibacterium gangwonense] | 06664 |
| hypothetical protein [Uliginosibacterium gangwonense] | 08108 |
| transporter [Uliginosibacterium aquaticum] | 02218 |
| hypothetical protein [Uliginosibacterium aquaticum] | 05241 |
| hypothetical protein [Thauera phenolivorans] | 04686 |
| glycosyltransferase [Thauera phenolivorans] | 06006 |
| hypothetical protein [Uliginosibacterium aquaticum] | 05278 |
| glycoside hydrolase family 31 protein [Uliginosibacterium gangwonense] | 04157 |
| glycosyltransferase [Uliginosibacterium gangwonense] | 06536 |
| hypothetical protein [Thauera phenolivorans] | 06702 |
| patatin-like phospholipase family protein [Zoogloea dura] | 09191 |
| DUF58 domain-containing protein, partial [Thauera phenolivorans] | 06335 |
| hypothetical protein [Zoogloea dura] | 07952 |

| | |
|---|---|
| hypothetical protein [Uliginosibacterium gangwonense] | 07649 |
| IS30 family transposase, partial [Zoogloea dura] | 00685 |
| histidine kinase [Uliginosibacterium gangwonense] | 07412 |
| dienelactone hydrolase family protein [Uliginosibacterium gangwonense] | 06297 |
| GNAT family N-acetyltransferase [Uliginosibacterium gangwonense] | 07411 |
| hypothetical protein [Uliginosibacterium gangwonense] | 00491 |
| hypothetical protein [Uliginosibacterium gangwonense] | 05978 |
| hypothetical protein [Uliginosibacterium gangwonense] | 05897 |
| PTS glucose transporter subunit IIBC [Uliginosibacterium gangwonense] | 04323 |
| hypothetical protein [Thauera phenolivorans] | 05410 |
| hypothetical protein [Thauera phenolivorans] | 05609 |
| hypothetical protein [Uliginosibacterium gangwonense] | 05686 |
| hypothetical protein [Azoarcus rhizosphaerae] | 02448 |
| hypothetical protein [Uliginosibacterium gangwonense] | 03074 |
| hypothetical protein [Uliginosibacterium gangwonense] | 09027 |
| hypothetical protein [Aromatoleum aromaticum] | 01999 |
| porin [Uliginosibacterium gangwonense] | 02563 |
| hypothetical protein [Uliginosibacterium gangwonense] | 09736 |
| glycosyl hydrolase [Uliginosibacterium gangwonense] | 04079 |
| glycosyltransferase [Uliginosibacterium gangwonense] | 09593 |
| hypothetical protein [Zoogloea dura] | 09269 |
| transporter substrate-binding domain-containing protein, partial [Zoogloea dura] | 00684 |
| helix-turn-helix domain-containing protein [Uliginosibacterium gangwonense] | 06300 |
| DUF3617 family protein [Zoogloea dura] | 02688 |
| hypothetical protein, partial [Azoarcus rhizosphaerae] | 00918 |
| hypothetical protein [Uliginosibacterium gangwonense] | 04160 |
| FkbM family methyltransferase [Azoarcus rhizosphaerae] | 02302 |
| hypothetical protein [Zoogloea dura] | 03025 |
| RDD family protein [Uliginosibacterium gangwonense] | 09410 |
| S53 family peptidase [Uliginosibacterium gangwonense] | 08947 |
| hypothetical protein [Aromatoleum tolulyticum] | 00248 |
| hypothetical protein [Thauera phenolivorans] | 08753 |
| GH1 family beta-glucosidase [Uliginosibacterium gangwonense] | 09738 |
| hypothetical protein [Aromatoleum petrolei] | 09338 |
| DUF3313 family protein [Uliginosibacterium gangwonense] | 01484 |
| methylaspartate mutase subunit S [Propionivibrio dicarboxylicus] | 07582 |
| DUF2169 domain-containing protein [Oryzomicrobium terrae] | 01827 |
| hypothetical protein [Oryzomicrobium terrae] | 05197 |
| DUF3944 domain-containing protein [Oryzomicrobium terrae] | 01741 |
| autotransporter domain-containing protein [Oryzomicrobium terrae] | 07349 |
| DUF3540 domain-containing protein [Oryzomicrobium terrae] | 04397 |
| rhomboid family intramembrane serine protease [Oryzomicrobium terrae] | 08174 |
| META domain-containing protein [Oryzomicrobium terrae] | 02328 |
| DUF3883 domain-containing protein [Oryzomicrobium terrae] | 04008 |
| hypothetical protein [Uliginosibacterium aquaticum] | 06166 |
| pilin [Oryzomicrobium terrae] | 08518 |
| cation transporting ATPase C-terminal domain-containing protein [Azoarcus sp. TTM-91] | 04571 |
| gamma-glutamylcyclotransferase [Azoarcus sp. TTM-91] | 03050 |
| hypothetical protein [Azoarcus sp. TTM-91] | 08958 |
| phosphoesterase, partial [Azoarcus sp. TTM-91] | 00672 |
| class I SAM-dependent methyltransferase [Azoarcus sp. TTM-91] | 05825 |
| hypothetical protein [Azoarcus sp. TTM-91] | 07327 |
| potassium channel family protein [Uliginosibacterium aquaticum] | 01906 |
| hypothetical protein [Uliginosibacterium aquaticum] | 04533 |
| tRNA (5-methylaminomethyl-2-thiouridine)(34)-methyltransferase MnmD, partial [Thauera phenylacetica] | 00269 |
| RidA family protein [Azoarcus sp. TTM-91] | 10145 |
| putative sulfate/molybdate transporter [Propionivibrio dicarboxylicus] | 02611 |
| ATP-binding cassette domain-containing protein, partial [Azoarcus sp. TTM-91] | 08303 |
| hypothetical protein [Oryzomicrobium terrae] | 09527 |

| | |
|---|---|
| ectonucleotide pyrophosphatase/phosphodiesterase [Oryzomicrobium terrae] | 03699 |
| YbaK/prolyl-tRNA synthetase associated domain-containing protein [Propionivibrio dicarboxylicus] | 10618 |
| hypothetical protein [Oryzomicrobium terrae] | 05094 |
| MEKHLA domain-containing protein [Uliginosibacterium aquaticum] | 08916 |
| sarcosine oxidase subunit delta, partial [Azoarcus sp. Aa7] | 08351 |
| hypothetical protein [Azoarcus sp. Aa7] | 07994 |
| 2-iminoacetate synthase ThiH [Propionivibrio dicarboxylicus] | 03278 |
| hypothetical protein [Zoogloea ramigera] | 06412 |
| hypothetical protein [Propionivibrio dicarboxylicus] | 03606 |
| DEAD/DEAH box helicase family protein, partial [Zoogloea ramigera] | 02964 |
| glycosyltransferase family 4 protein [Zoogloea ramigera] | 01146 |
| cyanophycinase [Zoogloea ramigera] | 05168 |
| hypothetical protein [Zoogloea ramigera] | 05479 |
| copper-binding protein, partial [Zoogloea ramigera] | 03421 |
| PAAR domain-containing protein, partial [Zoogloea ramigera] | 09498 |
| hypothetical protein, partial [Zoogloea ramigera] | 09499 |
| hypothetical protein, partial [Zoogloea ramigera] | 02963 |
| hypothetical protein [Azoarcus sp. Aa7] | 02779 |
| hypothetical protein [Oryzomicrobium terrae] | 08963 |
| sel1 repeat family protein [Propionivibrio dicarboxylicus] | 02266 |
| hypothetical protein [Oryzomicrobium terrae] | 00980 |
| hypothetical protein [Oryzomicrobium terrae] | 01537 |
| endonuclease [Oryzomicrobium terrae] | 04442 |
| cupin domain-containing protein [Oryzomicrobium terrae] | 07802 |
| hypothetical protein, partial [Azoarcus sp. TTM-91] | 01457 |
| hypothetical protein [Azoarcus sp. TTM-91] | 10149 |
| response regulator [Propionivibrio dicarboxylicus] | 08109 |
| cytochrome-c oxidase [Aromatoleum pumilum] | 09313 |
| hypothetical protein [Thauera hydrothermalis] | 00889 |
| DUF3426 domain-containing protein [Rhodocyclus purpureus] | 05595 |
| hypothetical protein [Rhodocyclus purpureus] | 01265 |
| helix-turn-helix domain-containing protein [Thauera hydrothermalis] | 03710 |
| GNAT family N-acetyltransferase [Thauera hydrothermalis] | 07468 |
| GyrI-like domain-containing protein [Thauera hydrothermalis] | 09469 |
| hypothetical protein [Rhodocyclus purpureus] | 10199 |
| hypothetical protein [Rhodocyclus purpureus] | 09822 |
| hypothetical protein [Thauera hydrothermalis] | 08788 |
| hypothetical protein [Thauera hydrothermalis] | 04231 |
| HIRAN domain-containing protein [Thauera hydrothermalis] | 08787 |
| hypothetical protein [Thauera hydrothermalis] | 03222 |
| hypothetical protein [Thauera hydrothermalis] | 05444 |
| DUF2281 domain-containing protein [Rhodocyclus purpureus] | 02247 |
| ATP-binding protein [Rhodocyclus purpureus] | 02656 |
| glutamate mutase L [Propionivibrio dicarboxylicus] | 00245 |
| hypothetical protein [Thauera hydrothermalis] | 03711 |
| DUF429 domain-containing protein [Rhodocyclus purpureus] | 02734 |
| hypothetical protein [Uliginosibacterium aquaticum] | 04234 |
| DUF3302 domain-containing protein [Uliginosibacterium aquaticum] | 03552 |
| AcaB family transcriptional regulator [Rhodocyclus purpureus] | 10052 |
| methylaspartate mutase subunit E [Propionivibrio dicarboxylicus] | 09681 |
| SRPBCC domain-containing protein [Thauera hydrothermalis] | 04713 |
| Ig-like domain-containing protein, partial [Uliginosibacterium aquaticum] | 05848 |
| DUF4387 domain-containing protein [Propionivibrio dicarboxylicus] | 08288 |
| FkbM family methyltransferase [Thauera phenylacetica] | 02011 |
| cysteine-rich CWC family protein [Aromatoleum pumilum] | 02441 |
| EAL domain-containing protein [Propionivibrio dicarboxylicus] | 00694 |
| sugar phosphate isomerase/epimerase [Propionivibrio dicarboxylicus] | 00799 |
| hypothetical protein [Propionivibrio dicarboxylicus] | 04609 |
| hypothetical protein [Aromatoleum pumilum] | 06344 |

| | |
|---|---|
| cytochrome c [Aromatoleum pumilum] | 08161 |
| L-ribulose-5-phosphate 3-epimerase [Propionivibrio dicarboxylicus] | 10620 |
| hypothetical protein [Aromatoleum pumilum] | 03689 |
| hypothetical protein [Aromatoleum pumilum] | 00872 |
| DMT family transporter [Rhodocyclus purpureus] | 05303 |
| DUF350 domain-containing protein [Azoarcus sp. Aa7] | 05898 |
| phosphate-starvation-inducible PsiE family protein [Aromatoleum pumilum] | 05098 |
| hypothetical protein [Azoarcus sp. Aa7] | 02768 |
| acyclic terpene utilization AtuA family protein [Propionivibrio dicarboxylicus] | 01576 |
| methylaspartate ammonia-lyase [Propionivibrio dicarboxylicus] | 05326 |
| hypothetical protein [Aromatoleum pumilum] | 03206 |
| DUF3656 domain-containing protein [Rhodocyclus purpureus] | 10200 |
| acetamidase/formamidase family protein [Aromatoleum pumilum] | 05101 |
| hypothetical protein [Aromatoleum pumilum] | 06017 |
| metallophosphoesterase [Uliginosibacterium aquaticum] | 08323 |
| hypothetical protein [Propionivibrio dicarboxylicus] | 06615 |
| DUF4189 domain-containing protein [Aromatoleum petrolei] | 04967 |
| hypothetical protein [Uliginosibacterium gangwonense] | 00494 |
| hypothetical protein [Uliginosibacterium gangwonense] | 01613 |
| helix-turn-helix domain-containing protein [Thauera chlorobenzoica] | 08149 |
| hypothetical protein [Thauera aminoaromatica] | 04627 |
| hypothetical protein [Zoogloea dura] | 10006 |
| GNAT family N-acetyltransferase [Propionivibrio dicarboxylicus] | 02498 |
| hypothetical protein [Propionivibrio dicarboxylicus] | 04999 |
| DUF1566 domain-containing protein [Thauera butanivorans] | 08129 |
| HNH endonuclease [Thauera butanivorans] | 05068 |
| Arc family DNA-binding protein [Thauera butanivorans] | 04190 |
| RpiB/LacA/LacB family sugar-phosphate isomerase [Propionivibrio dicarboxylicus] | 09273 |
| TIGR00366 family protein [Propionivibrio dicarboxylicus] | 00798 |
| SLC13 family permease [Propionivibrio dicarboxylicus] | 08917 |
| DEAD/DEAH box helicase [Propionivibrio dicarboxylicus] | 00792 |
| DNA polymerase I [Propionivibrio dicarboxylicus] | 08441 |
| ankyrin repeat domain-containing protein [Thauera butanivorans] | 04348 |
| DUF3293 domain-containing protein [Propionivibrio dicarboxylicus] | 01530 |
| sensor histidine kinase [Propionivibrio dicarboxylicus] | 03775 |
| AzlD domain-containing protein [Propionivibrio dicarboxylicus] | 10053 |
| GTPase domain-containing protein [Thauera phenylacetica] | 00271 |
| DUF4150 domain-containing protein [Thauera butanivorans] | 04936 |
| GtrA family protein [Propionivibrio dicarboxylicus] | 07707 |
| ESPR domain-containing protein, partial [Azoarcus indigens] | 00649 |
| DUF1345 domain-containing protein [Uliginosibacterium gangwonense] | 05248 |
| YecH family protein [Propionivibrio dicarboxylicus] | 08995 |
| hypothetical protein, partial [Azoarcus indigens] | 06968 |
| hypothetical protein [Uliginosibacterium gangwonense] | 07020 |
| hypothetical protein [Uliginosibacterium aquaticum] | 10574 |
| AGE family epimerase/isomerase [Uliginosibacterium gangwonense] | 03638 |
| ASCH domain-containing protein [Uliginosibacterium gangwonense] | 03312 |
| Cof-type HAD-IIB family hydrolase [Propionivibrio dicarboxylicus] | 05610 |
| cellulase family glycosylhydrolase [Uliginosibacterium gangwonense] | 05972 |
| ROK family protein [Propionivibrio dicarboxylicus] | 07979 |
| SRPBCC family protein [Uliginosibacterium gangwonense] | 07138 |
| DUF4392 domain-containing protein [Propionivibrio dicarboxylicus] | 10263 |
| zinc ribbon domain-containing protein [Uliginosibacterium gangwonense] | 10224 |
| helix-turn-helix domain-containing protein, partial [Azoarcus indigens] | 02182 |
| hypothetical protein, partial [Azoarcus indigens] | 09233 |
| pyridoxamine 5'-phosphate oxidase family protein [Propionivibrio dicarboxylicus] | 10615 |
| hypothetical protein [Zoogloea dura] | 10374 |
| flagellar biosynthesis protein FlgA [Propionivibrio dicarboxylicus] | 05324 |
| hypothetical protein [Propionivibrio dicarboxylicus] | 03602 |

| | |
|---|---|
| non-ribosomal peptide synthetase [Uliginosibacterium gangwonense] | 10572 |
| hypothetical protein [Propionivibrio dicarboxylicus] | 04322 |
| PAS domain S-box protein [Propionivibrio dicarboxylicus] | 07369 |
| hypothetical protein [Uliginosibacterium aquaticum] | 01810 |
| transposase, partial [Azoarcus indigens] | 07925 |
| GNAT family N-acetyltransferase [Propionivibrio dicarboxylicus] | 09560 |
| MarR family EPS-associated transcriptional regulator [Propionivibrio dicarboxylicus] | 05964 |
| LysR substrate-binding domain-containing protein [Thauera butanivorans] | 05078 |
| MULTISPECIES: hypothetical protein [Thauera] | 06638 |
| hypothetical protein, partial [Thauera butanivorans] | 01027 |
| transporter substrate-binding domain-containing protein [Propionivibrio dicarboxylicus] | 07971 |
| glycoside hydrolase family 104 protein [Azoarcus sp. Aa7] | 10044 |
| DUF2892 domain-containing protein [Propionivibrio dicarboxylicus] | 09833 |
| hypothetical protein [Uliginosibacterium aquaticum] | 10545 |
| mechanosensitive ion channel [Uliginosibacterium aquaticum] | 08009 |
| DUF4124 domain-containing protein [Uliginosibacterium aquaticum] | 04042 |
| hypothetical protein [Uliginosibacterium aquaticum] | 09704 |
| hypothetical protein [Uliginosibacterium aquaticum] | 04906 |
| DUF1441 family protein [Azoarcus sp. Aa7] | 00173 |
| MULTISPECIES: hypothetical protein [Proteobacteria] | 00480 |
| MULTISPECIES: hypothetical protein [Bacteria] | 09010 |
| PDZ domain-containing protein [Zoogloea ramigera] | 06033 |
| hypothetical protein [Propionivibrio dicarboxylicus] | 00795 |
| [citrate (pro-3S)-lyase] ligase [Propionivibrio dicarboxylicus] | 04119 |
| hypothetical protein [Zoogloea ramigera] | 08826 |
| hypothetical protein, partial [Zoogloea ramigera] | 02117 |
| hypothetical protein [Zoogloea ramigera] | 09159 |
| DEAD/DEAH box helicase family protein [Zoogloea ramigera] | 10439 |
| hypothetical protein [Uliginosibacterium aquaticum] | 00158 |
| hypothetical protein [Propionivibrio dicarboxylicus] | 03157 |
| hypothetical protein [Propionivibrio dicarboxylicus] | 07102 |
| hypothetical protein [Propionivibrio dicarboxylicus] | 02061 |
| right-handed parallel beta-helix repeat-containing protein, partial [Thauera butanivorans] | 05405 |
| hypothetical protein [Propionivibrio dicarboxylicus] | 02499 |
| ankyrin repeat domain-containing protein [Thauera butanivorans] | 06325 |
| hypothetical protein [Thauera phenylacetica] | 09395 |
| glycosyltransferase family protein [Propionivibrio dicarboxylicus] | 00917 |
| DUF1851 domain-containing protein [Thauera butanivorans] | 10541 |
| alpha-1,2-fucosyltransferase [Propionivibrio dicarboxylicus] | 04802 |
| hypothetical protein [Thauera butanivorans] | 01024 |
| MULTISPECIES: hypothetical protein [Thauera] | 07736 |
| RbsD or FucU transport [Propionivibrio dicarboxylicus] | 06654 |
| Cyanophycin synthase [Thauera aminoaromatica] | 09815 |
| DUF6268 family outer membrane beta-barrel protein [Uliginosibacterium aquaticum] | 05846 |
| hypothetical protein [Propionivibrio dicarboxylicus] | 06614 |
| hypothetical protein [Thauera butanivorans] | 04677 |
| hypothetical protein [Azoarcus indigens] | 00904 |
| IS110 family transposase, partial [Thauera butanivorans] | 06327 |
| ankyrin repeat domain-containing protein [Thauera butanivorans] | 04681 |
| polymorphic toxin type 15 domain-containing protein [Thauera butanivorans] | 10201 |
| hypothetical protein, partial [Thauera butanivorans] | 10325 |
| MULTISPECIES: heat resistance protein YfdX2 [Proteobacteria] | 09004 |
| hypothetical protein, partial [Thauera butanivorans] | 04352 |
| hypothetical protein, partial [Thauera butanivorans] | 06001 |
| GMC family oxidoreductase N-terminal domain-containing protein [Aromatoleum toluolicum] | 08298 |

### Table S6: Average nucleotide identities presented as percent pairwise identity

| Query | Zoogloea | Zoogloea | Zoogloea | Cognataz | Nitrogeni | Nitrogeni | Nitrogeni | Denitrom | Denitrom | Denitrom | Denitrom | Pseudoaz | Pseudoaz | Aromatol | Aromatol | Aromatol | Aromatol | Azoarcus | Aromatol | Aromatol | Azoarcus | Aromatol | Aromatol | Aromatol | Aromatol |
|---|---|---|---|---|---|---|---|---|---|---|---|---|---|---|---|---|---|---|---|---|---|---|---|---|---|
| Zoogloea oleivorans strain Buc | 99.997 | 83.6785 | 83.2605 | 78.4389 | 78.7746 | 78.1575 | 78.2023 | 78.9677 | 78.5539 | 78.7723 | 78.823 | 77.9065 | 78.3155 | 79.0706 | 78.4746 | 78.0889 | 78.6937 | 78.6965 | 78.8944 | 79.8182 | 78.7392 | 78.7323 | 78.7741 | 80.371 | 79.138 |
| Zoogloea dura | 83.9281 | 100 | 83.3643 | 78.7278 | 79.085 | 78.3263 | 78.2042 | 79.0797 | 78.4932 | 78.533 | 78.5786 | 78.0656 | 78.4458 | 78.7196 | 78.2838 | 77.959 | 78.5688 | 78.666 | 78.8243 | 79.5955 | 78.8375 | 78.7899 | 78.4652 | 80.3896 | 78.9439 |
| Zoogloea ramigera | 83.27 | 83.4569 | 100 | 78.6242 | 79.3336 | 78.0276 | 77.9277 | 79.4654 | 78.8011 | 78.9408 | 78.9443 | 78.5162 | 78.3262 | 78.6867 | 78.5696 | 78.4133 | 78.5851 | 78.9759 | 79.2723 | 79.3261 | 79.14 | 79.3288 | 79.0795 | 79.3052 | 79.2616 |
| Cognatazoarcus halotolerans | 78.4705 | 78.6529 | 78.5935 | 100 | 78.7193 | 78.1089 | 78.2315 | 79.3298 | 78.8024 | 78.8325 | 78.8236 | 77.8753 | 78.721 | 78.5298 | 78.5496 | 77.9886 | 78.2539 | 78.3477 | 78.6183 | 78.8814 | 78.6572 | 78.4218 | 78.458 | 78.6334 | 78.7302 |
| Nitrogeniibacter mangrovi | 78.7582 | 79.1116 | 79.3437 | 78.8167 | 100 | 81.6308 | 81.6355 | 82.007 | 80.3703 | 80.264 | 80.2679 | 79.139 | 78.4876 | 78.5234 | 78.3168 | 78.2317 | 78.7124 | 78.9717 | 79.0632 | 79.0734 | 78.9301 | 79.1243 | 78.9092 | 78.949 | 78.8678 |
| Nitrogeniibacter aestuarii sp. ZN11-R3-1 | 78.1761 | 78.269 | 77.9943 | 78.1219 | 81.5795 | 100 | 98.2779 | 79.852 | 79.2836 | 79.3112 | 79.3107 | 77.7925 | | | | | 77.8512 | 77.9586 | 77.84 | 78.0393 | 78.0232 | 78.139 | 77.6504 | 78.071 | 77.8912 |
| Nitrogeniibacter aestuarii sp. H1-1-2A | 78.1465 | 78.2422 | 78.0344 | 78.28 | 81.5801 | 98.2229 | 100 | 79.8659 | 79.2581 | 79.3145 | 79.2793 | 77.9784 | | | | | | 78.1604 | 77.7921 | 77.9322 | 77.8366 | 78.0893 | 77.8181 | 77.794 | 77.7334 |
| Denitromonas iodatirespirans IR12 | 79.0492 | 78.8188 | 79.3214 | 79.3069 | 81.6841 | 79.9494 | 79.8697 | 100 | 85.0271 | 85.1743 | 85.1556 | 79.1433 | 78.7796 | 78.9229 | 78.8132 | 78.8077 | 78.8926 | 79.3783 | 79.711 | 79.4637 | 79.2907 | 79.4651 | 79.2666 | 79.2264 | 79.5628 |
| Denitromonas halophila SFB-3 | 78.3873 | 78.3276 | 78.7201 | 78.8621 | 80.3705 | 79.3326 | 79.2678 | 85.1105 | 100 | 94.4583 | 94.4447 | 78.1474 | 78.1888 | 77.9819 | 78.0711 | 77.9597 | 78.2867 | 78.5678 | 78.7703 | 78.5456 | 78.5936 | 78.6234 | 78.3339 | 78.4887 | 78.6092 |
| Denitromonas ohlonensis SFB-1 | 78.8445 | 78.5105 | 78.7245 | 79.0267 | 80.3969 | 79.2206 | 79.2995 | 85.2383 | 94.4713 | 100 | 99.9959 | 78.2858 | 78.6371 | 78.5652 | 78.1592 | 78.3054 | 78.4756 | 78.4517 | 78.7999 | 78.8084 | 78.5011 | 78.4804 | 78.5417 | 78.8048 | 79.0537 |
| Denitromonas ohlonensis SFB-2 | 78.8999 | 78.544 | 78.671 | 79.0904 | 80.3835 | 79.2617 | 79.3219 | 85.2423 | 94.4263 | 99.9939 | 100 | 78.3009 | 78.5495 | 78.5202 | 78.1673 | 78.2245 | 78.4959 | 78.4709 | 78.9066 | 78.848 | 78.5636 | 78.5683 | 78.5237 | 78.8261 | 79.0284 |
| Pseudoazoarcus pumilus | 77.9415 | 77.911 | 78.5785 | 77.9888 | 79.0125 | 77.8812 | 78.0399 | 79.1603 | 78.3339 | 78.3407 | 78.3621 | 100 | 79.0211 | 79.2169 | 79.048 | 79.138 | 79.1161 | 79.1407 | 79.6125 | 79.4655 | 79.3437 | 79.4671 | 79.5053 | 79.6666 | 79.3464 |
| Pseudoazoarcus taiwanensis | 78.3194 | 78.2972 | 78.3689 | 78.6698 | 78.428 | | | 78.6058 | 78.1284 | 78.416 | 78.3948 | 78.9693 | 100 | 79.2387 | 78.6823 | 78.5466 | 78.5947 | 78.4902 | 78.7415 | 78.9031 | 78.6741 | 78.9032 | 78.6745 | 79.0611 | 78.9388 |
| Aromatoleum bremense strain PbN1 | 78.8201 | 78.6804 | 78.6524 | 78.6722 | 78.6572 | | | 78.8841 | 78.1032 | 78.5394 | 78.5461 | 79.094 | 79.3539 | 100 | 92.4623 | 92.2625 | 82.2928 | 81.504 | 82.7645 | 82.3739 | 82.1818 | 82.2699 | 81.9799 | 82.4218 | 82.4968 |
| Aromatoleum aromaticum EbN1 | 78.5551 | 78.2461 | 78.5183 | 78.6534 | 78.3355 | | | 78.7883 | 78.0637 | 78.1162 | 78.1654 | 79.0025 | 78.628 | 92.5272 | 100 | 93.2847 | 81.5702 | 81.334 | 82.4274 | 82.3209 | 82.0039 | 81.847 | 81.9722 | 82.3038 | 82.1406 |
| Aromatoleum buckelii | 77.941 | 77.9483 | 78.3781 | 78.0309 | 78.303 | | | 78.6208 | 77.9097 | 78.0153 | 78.016 | 79.1492 | 78.2746 | 92.3349 | 93.1742 | 100 | 81.6962 | 81.3773 | 82.5134 | 82.4652 | 82.0829 | 82.4674 | 81.8966 | 82.3277 | 82.0125 |
| Aromatoleum diolicum | 78.5239 | 78.533 | 78.5777 | 78.4528 | 78.7803 | | 77.7572 | 79.0912 | 78.2291 | 78.3895 | 78.3821 | 79.0726 | 78.5715 | 81.9723 | 81.5054 | 81.6654 | 100 | 82.4924 | 83.493 | 84.283 | 83.6051 | 83.2029 | 82.8331 | 83.7202 | 83.1732 |
| Azoarcus sp. KH32C | 78.7559 | 78.7086 | 78.9887 | 78.3835 | 78.7372 | 78.1121 | 78.0342 | 79.302 | 78.4757 | 78.4569 | 78.4524 | 78.9955 | 78.3781 | 81.5294 | 81.2428 | 81.3092 | 82.5981 | 100 | 83.7756 | 83.7871 | 83.4757 | 83.4823 | 83.1055 | 83.6944 | 83.2332 |
| Aromatoleum toluvorans | 78.8185 | 78.8946 | 79.4774 | 78.713 | 79.0387 | 77.9311 | 77.8134 | 79.5907 | 78.7996 | 78.5819 | 78.5656 | 79.6767 | 78.8003 | 82.6374 | 82.1965 | 82.4795 | 83.3333 | 83.7628 | 100 | 87.569 | 87.5392 | 87.4278 | 87.2395 | 87.6396 | 88.1856 |
| Aromatoleum toluolicum | 79.7234 | 79.7142 | 79.2755 | 78.8401 | 78.8981 | 77.8141 | 77.9132 | 79.4453 | 78.5646 | 78.6613 | 78.7015 | 79.4155 | 79.14 | 82.3495 | 82.188 | 82.207 | 84.2044 | 83.796 | 87.5515 | 100 | 92.4977 | 89.3202 | 88.8774 | 93.3019 | 89.6973 |
| Azoarcus sp. Aa7 | 78.77 | 78.7432 | 79.3985 | 78.5635 | 78.8253 | 78.0587 | 77.9395 | 79.3698 | 78.4635 | 78.5754 | 78.5554 | 79.362 | 78.6636 | 82.1398 | 82.0146 | 82.1144 | 83.7038 | 83.5682 | 87.5593 | 92.488 | 100 | 89.3263 | 88.775 | 92.731 | 90.4708 |
| Aromatoleum evansii | 78.5667 | 78.7887 | 79.1255 | 78.3439 | 79.0917 | 77.9729 | 78.0023 | 79.3492 | 78.4309 | 78.5956 | 78.6156 | 79.3565 | 78.8406 | 82.0432 | 81.8458 | 82.3427 | 83.2079 | 83.4349 | 87.3125 | 89.3911 | 89.339 | 100 | 91.4249 | 89.5542 | 89.2684 |
| Aromatoleum petrolei | 78.5395 | 78.2941 | 79.1617 | 78.5451 | 78.9137 | 77.7799 | 77.8292 | 79.1193 | 78.3618 | 78.521 | 78.5375 | 79.3767 | 78.6265 | 81.8838 | 81.785 | 81.9945 | 82.8863 | 82.9593 | 87.0902 | 88.9137 | 88.8844 | 91.4475 | 100 | 88.9413 | 88.9719 |
| Aromatoleum tolulyticum | 80.2531 | 80.0398 | 79.4532 | 78.9476 | 78.9477 | 77.9517 | 77.8959 | 79.4748 | 78.6944 | 78.8532 | 78.894 | 79.5832 | 79.1327 | 82.3956 | 82.2573 | 82.4623 | 83.7642 | 83.5802 | 87.6361 | 93.2478 | 92.7191 | 89.4555 | 88.9008 | 100 | 90.2375 |
| Aromatoleum toluclasticum ATCC 700605 | 78.903 | 78.9427 | 79.2482 | 78.6477 | 78.912 | 77.7166 | 77.7281 | 79.5356 | 78.491 | 79.1057 | 79.0798 | 79.2639 | 78.966 | 82.4559 | 82.1789 | 82.0724 | 83.2929 | 83.2235 | 88.0689 | 89.7807 | 90.5142 | 89.192 | 89.1145 | 90.3465 | 100 |
| Aromatoleum toluclasticum TC-10 | 79.2799 | 79.1524 | 79.2159 | 78.8385 | 78.864 | 77.8523 | 77.7546 | 79.5214 | 78.5025 | 78.8924 | 78.9155 | 79.3657 | 78.941 | 82.3005 | 82.1608 | 82.0158 | 83.3185 | 83.1666 | 87.9008 | 89.6335 | 90.2934 | 89.3535 | 88.8617 | 90.3158 | 98.116 |
| Pseudothauera lacus sp. D20 | 79.3151 | 79.5087 | 80.1861 | 78.8856 | 80.0241 | 78.4888 | 78.6101 | 80.2466 | 79.0092 | 79.1228 | 79.133 | 79.9601 | 79.3367 | 79.6907 | 79.5262 | 79.4104 | 80.0869 | 79.9716 | 80.7325 | 80.617 | 80.5094 | 80.7595 | 80.3178 | 80.7108 | 80.5118 |
| Pseudothauera hydrothermalis | 77.73 | 77.6249 | 77.9499 | 78.2141 | 78.2701 | 77.5682 | 77.7351 | 78.3614 | 77.724 | 77.6859 | 77.6775 | 78.2103 | 78.0113 | 77.9549 | 77.9452 | 77.7701 | 78.2145 | 77.9226 | 78.3961 | 78.2383 | 78.0764 | 78.3048 | 78.3613 | 78.1663 | 78.145 |
| Pseudothauera nasutitermitis | 79.2219 | 79.2834 | 79.6718 | 78.5647 | 79.9394 | 78.3223 | 78.4408 | 80.0946 | 78.8917 | 78.8623 | 78.8468 | 80.0071 | 79.2128 | 79.7878 | 79.4788 | 79.69 | 79.4821 | 79.5299 | 80.6837 | 80.3428 | 80.3456 | 80.2226 | 80.3002 | 80.3556 | 80.2364 |
| Pseudothauera rhizosphaerae | 79.1756 | 79.5856 | 80.1481 | 78.558 | 80.319 | 78.899 | 78.9004 | 79.9141 | 79.1538 | 78.9275 | 78.9242 | 79.9354 | 78.7549 | 79.6425 | 79.5236 | 79.48 | 79.7525 | 79.6799 | 80.6019 | 80.4392 | 80.4222 | 80.5159 | 80.2833 | 80.2685 | 80.2589 |
| Parazoarcus communis LMG 5514 | 80.4127 | 79.9111 | 78.6483 | 78.7302 | 78.5786 | 77.8369 | 77.9473 | 78.6925 | 78.3479 | 78.7779 | 78.7842 | 78.1744 | 79.3787 | 79.0392 | 78.8283 | 78.3116 | 79.124 | 78.8267 | 79.1761 | 79.9612 | 79.1387 | 78.9921 | 78.8134 | 79.9894 | 79.0499 |
| Parazoarcus communis SWub3 | 78.6674 | 78.7862 | 78.5208 | 78.6304 | 78.6201 | 77.9746 | 78.0522 | 78.7067 | 78.2826 | 78.2439 | 78.2614 | 78.1887 | 78.6409 | 78.774 | 78.6307 | 78.4039 | 78.9557 | 78.826 | 79.3145 | 78.928 | 79.0818 | 79.1874 | 78.9465 | 78.8704 | 78.8407 |
| Parazoarcus communis | 78.9484 | 78.7349 | 78.8261 | 78.7267 | 78.9491 | 78.2927 | 78.3903 | 79.3164 | 78.7506 | 79.0047 | 78.9829 | 78.6955 | 78.3067 | 79.0451 | 78.828 | 78.9007 | 79.1716 | 79.1639 | 79.3527 | 79.4241 | 79.2309 | 79.4864 | 79.4255 | 79.0886 | 79.4062 |
| Azoarcus sp. L1K30 | 78.4003 | 78.3339 | 78.213 | 78.8527 | 78.5039 | 78.1001 | 78.3167 | 79.0005 | 78.5632 | 78.2423 | 78.2438 | 78.2969 | 78.5253 | 78.6864 | 78.4036 | 78.5545 | 78.7 | 78.8863 | 79.34 | 79.0084 | 78.999 | 79.226 | 78.967 | 79.046 | 79.0948 |
| Azoarcus sp. TTM-91 | 78.9584 | 79.0962 | 79.2456 | 78.4233 | 79.1862 | 78.1154 | 78.1111 | 79.3174 | 78.4873 | 78.6546 | 78.6831 | 78.8473 | 78.8123 | 79.0787 | 78.8878 | 78.916 | 79.133 | 79.1747 | 79.9685 | 79.814 | 79.742 | 79.8049 | 79.362 | 79.5954 | 79.527 |
| Azoarcus indigens VB32 | 78.6514 | 79.0188 | 79.4612 | 78.9676 | 79.3697 | 78.086 | 78.1297 | 79.2175 | 78.5409 | 78.4824 | 78.477 | 78.9338 | 78.9092 | 79.2287 | 79.0843 | 78.8265 | 79.312 | 79.1901 | 80.0865 | 79.8082 | 79.6557 | 79.7925 | 79.5709 | 79.6104 | 79.647 |
| Azoarcus indigens | 78.5937 | 79.1719 | 79.2926 | 78.9816 | 79.2864 | 78.3199 | 78.1484 | 79.1294 | 78.4175 | 78.3643 | 78.336 | 78.7266 | 78.7796 | 79.2591 | 79.1545 | 78.7566 | 79.1782 | 79.1982 | 79.9995 | 79.68 | 79.6352 | 79.6883 | 79.3772 | 79.726 | 79.5213 |
| Azoarcus sp. DD4 | 79.4985 | 79.6398 | 79.6573 | 79.3348 | 79.5048 | 78.2924 | 78.5476 | 79.6375 | 78.9241 | 79.4056 | 79.4165 | 79.3178 | 79.4825 | 80.1916 | 79.885 | 79.6413 | 80.3459 | 80.2666 | 80.8101 | 80.8055 | 80.715 | 80.7824 | 80.5522 | 80.7553 | 80.6575 |
| Azoarcus olearius BH72 | 79.1597 | 79.548 | 79.8157 | 79.1624 | 79.6003 | 78.6979 | 78.6843 | 79.7792 | 79.0662 | 79.2953 | 79.2919 | 79.3669 | 78.8071 | 80.2312 | 79.8318 | 79.8099 | 80.1193 | 80.0955 | 81.0832 | 80.6505 | 80.5957 | 80.697 | 80.6371 | 80.5866 | 80.4079 |
| Azoarcus olearius DQS-4 | 79.3115 | 79.5188 | 79.7076 | 79.2157 | 79.505 | 78.5289 | 78.42 | 79.764 | 78.9773 | 79.1716 | 79.1619 | 79.6313 | 78.7522 | 80.3078 | 79.8514 | 79.9799 | 80.1388 | 80.0567 | 81.1487 | 80.6977 | 80.5827 | 80.8849 | 80.3992 | 80.623 | 80.6425 |
| Thauera phenolivorans | 78.8403 | 79.1182 | 80.0562 | 79.1178 | 79.4538 | 78.2306 | 78.3638 | 79.7637 | 78.7033 | 78.926 | 78.9053 | 79.7343 | 79.4168 | 80.2214 | 80.2561 | 79.8444 | 79.7071 | 80.0904 | 80.9628 | 80.5411 | 80.6817 | 80.6283 | 80.6386 | 80.8199 | 80.4637 |
| Thauera aminoaromatica S2 | 79.2258 | 79.5221 | 80.4163 | 79.3054 | 79.9478 | 78.4831 | 78.3994 | 80.261 | 79.1137 | 79.1043 | 79.1013 | 80.0692 | 79.4559 | 80.1623 | 79.8952 | 79.8752 | 79.8489 | 80.2835 | 81.2552 | 80.8867 | 80.9483 | 81.1938 | 80.8399 | 81.058 | 80.9199 |
| Thauera phenylacetica | 79.3334 | 79.6944 | 80.8752 | 79.4426 | 80.0033 | 78.5867 | 78.6006 | 80.3189 | 79.2581 | 79.2552 | 79.2608 | 80.4695 | 80.0457 | 80.567 | 80.1271 | 80.0649 | 80.0126 | 80.3077 | 81.3495 | 81.1195 | 81.0293 | 81.1493 | 80.7592 | 81.1474 | 81.0545 |
| Thauera aromatica K172 | 79.1471 | 79.271 | 79.9828 | 78.8082 | 79.5614 | 78.218 | 78.2634 | 79.7716 | 78.7803 | 78.7677 | 78.7299 | 80.0714 | 79.5182 | 80.6487 | 80.8206 | 80.4912 | 80.0899 | 80.1801 | 81.1621 | 80.9257 | 81.0019 | 80.9279 | 80.7792 | 80.7901 | 80.881 |
| Thauera chlorobenzoica | 79.1513 | 79.0279 | 79.9478 | 78.8276 | 79.774 | 78.1523 | 78.1515 | 79.9667 | 78.9309 | 78.9992 | 78.9922 | 79.8151 | 79.4551 | 80.9033 | 80.8037 | 80.6437 | 80.1233 | 80.1488 | 81.0896 | 80.9621 | 81.102 | 81.1048 | 80.7753 | 80.9832 | 80.9043 |
| Thauera propionica | 78.9105 | 79.248 | 79.8874 | 78.8434 | 79.4638 | 78.3884 | 78.2677 | 79.8397 | 78.9506 | 78.9751 | 78.9644 | 79.6045 | 79.1298 | 79.8585 | 79.6365 | 79.6555 | 80.0488 | 80.1111 | 81.0208 | 80.6087 | 80.7991 | 81.0836 | 80.4897 | 80.708 | 80.632 |
| Thauera terpenica | 78.4991 | 78.7937 | 78.9681 | 79.4648 | 78.7127 | 77.996 | 78.093 | 79.0039 | 78.2922 | 78.6961 | 78.6741 | 78.9121 | 79.1139 | 79.2906 | 79.0817 | 78.8095 | 79.0068 | 79.0461 | 79.7483 | 79.6432 | 79.5924 | 79.5591 | 79.4195 | 79.5864 | 79.5556 |
| Thauera linaloolentis | 79.0893 | 79.0569 | 79.717 | 78.7657 | 79.1572 | 77.9425 | 78.0803 | 79.691 | 78.8171 | 78.8048 | 78.8323 | 79.5176 | 78.965 | 79.8984 | 79.5144 | 79.5478 | 79.6706 | 80.021 | 80.8333 | 80.5064 | 80.4018 | 80.3919 | 80.0582 | 80.552 | 80.2307 |
| Thauera butanivorans | 79.2987 | 79.6691 | 80.2467 | 79.9121 | 79.6976 | 78.3109 | 78.2504 | 79.9634 | 78.7341 | 79.0933 | 79.1231 | 80.1324 | 79.7702 | 80.554 | 80.5388 | 80.3196 | 80.1523 | 80.2357 | 80.9078 | 80.8886 | 80.8223 | 81.0479 | 80.5148 | 81.0112 | 80.7829 |

| Query | Aromatol | Pseudoth | Pseudoth | Pseudoth | Pseudoth | Parazoarc | Parazoarc | Parazoarc | Azoarcus | Azoarcus | Azoarcus | Azoarcus | Azoarcus | Azoarcus | Azoarcus | Thauera p | Thauera a | Thauera p | Thauera a | Thauera c | Thauera p | Thauera t | Thauera l | Thauera butanivorans |
|---|---|---|---|---|---|---|---|---|---|---|---|---|---|---|---|---|---|---|---|---|---|---|---|---|
| Zoogloea oleivorans strain Buc | 79.3772 | 79.1339 | 77.6749 | 79.0879 | 79.132 | 80.3995 | 78.5494 | 78.6357 | 78.2753 | 78.8392 | 78.9129 | 78.8214 | 79.5471 | 79.3375 | 79.3285 | 78.8057 | 79.3479 | 79.4424 | 79.0229 | 78.9975 | 79.163 | 78.6476 | 79.1587 | 79.3861 |
| Zoogloea dura | 79.128 | 79.3501 | 77.8179 | 79.2945 | 79.689 | 79.9415 | 78.7469 | 78.6964 | 78.3981 | 79.2728 | 79.289 | 79.1935 | 79.6605 | 79.4869 | 79.4779 | 79.2555 | 79.5788 | 79.7501 | 79.1614 | 79.0514 | 79.2294 | 78.7481 | 79.1765 | 79.6404 |
| Zoogloea ramigera | 79.3074 | 79.994 | 78.0568 | 79.6753 | 79.9632 | 78.7126 | 78.4955 | 78.9638 | 78.1518 | 79.0356 | 79.2483 | 79.2719 | 79.5796 | 79.6275 | 79.6953 | 79.9386 | 80.5208 | 81.052 | 79.7413 | 79.7701 | 79.8994 | 78.9947 | 79.8997 | 80.1065 |
| Cognatazoarcus halotolerans | 78.6353 | 78.8855 | 78.1083 | 78.5574 | 78.6166 | 78.7844 | 78.5745 | 78.7914 | 78.8531 | 78.4958 | 79.088 | 79.0305 | 79.2702 | 79.1235 | 79.1848 | 79.174 | 79.427 | 79.5234 | 78.6645 | 78.8813 | 78.8316 | 79.4503 | 78.6978 | 79.8001 |
| Nitrogeniibacter mangrovi | 78.923 | 79.8481 | 78.4043 | 80.0349 | 80.1548 | 78.7455 | 78.6041 | 78.9419 | 78.5741 | 79.0785 | 79.2668 | 79.2924 | 79.6131 | 79.5502 | 79.5201 | 79.3742 | 79.7728 | 80.022 | 79.488 | 79.64 | 79.4341 | 78.5648 | 79.2495 | 79.5699 |
| Nitrogeniibacter aestuarii sp. ZN11-R3-1 | 78.0163 | 78.4276 | 77.577 | 78.3994 | 78.8748 | 77.8909 | 77.9733 | 78.2371 | 78.0256 | 78.1191 | 78.4241 | 78.506 | 78.3876 | 78.5815 | 78.5864 | 78.1921 | 78.342 | 78.4948 | 78.0461 | 77.9953 | 78.6096 | 77.9723 | 78.1661 | 78.345 |
| Nitrogeniibacter aestuarii sp. H1-1-2A | 77.8237 | 78.5429 | 77.5669 | 78.4028 | 78.8924 | 77.9398 | 78.0995 | 78.4512 | 78.1533 | 78.2027 | 78.3089 | 78.3944 | 78.3841 | 78.5291 | 78.5574 | 78.3968 | 78.4524 | 78.585 | 78.1218 | 78.283 | 78.5606 | 77.8507 | 77.9629 | 78.2279 |
| Denitromonas iodatirespirans IR12 | 79.4982 | 80.276 | 78.2571 | 79.7964 | 80.0722 | 78.746 | 78.5769 | 79.3049 | 78.9761 | 79.082 | 79.1329 | 79.0877 | 79.7476 | 79.8475 | 79.9637 | 79.7663 | 80.2909 | 80.5201 | 79.6014 | 79.9459 | 79.6766 | 78.9537 | 79.6924 | 79.9402 |
| Denitromonas halophila SFB-3 | 78.573 | 79.0907 | 77.6357 | 78.8726 | 79.1891 | 78.4074 | 78.359 | 78.7299 | 78.5201 | 78.5452 | 78.4945 | 78.5355 | 78.9039 | 78.9142 | 78.9508 | 78.7771 | 78.965 | 79.1552 | 78.7012 | 78.8151 | 78.9415 | 78.3056 | 78.7543 | 79.0078 |
| Denitromonas ohlonensis SFB-1 | 78.9757 | 78.8779 | 77.9091 | 78.7816 | 78.98 | 78.7491 | 78.3068 | 78.7609 | 78.4147 | 78.5807 | 78.5059 | 78.5754 | 79.3403 | 79.2797 | 79.2619 | 78.6542 | 79.2759 | 79.4467 | 78.7446 | 78.7965 | 78.9654 | 78.5894 | 78.5256 | 79.0879 |
| Denitromonas ohlonensis SFB-2 | 79.0001 | 78.9363 | 77.9219 | 78.8299 | 79.0267 | 78.7546 | 78.3081 | 78.8449 | 78.4303 | 78.4966 | 78.5091 | 78.5407 | 79.3658 | 79.2339 | 79.2749 | 78.6079 | 79.3026 | 79.4156 | 78.7972 | 78.7307 | 78.9833 | 78.5645 | 78.5347 | 79.025 |
| Pseudoazoarcus pumilus | 79.4994 | 80.0147 | 78.3608 | 80.1141 | 79.7735 | 78.2014 | 78.323 | 78.6442 | 78.4727 | 78.9645 | 78.8905 | 78.8853 | 79.3182 | 79.3479 | 79.5017 | 79.7501 | 80.0856 | 80.4902 | 79.8517 | 79.8992 | 79.6268 | 78.8852 | 79.4905 | 80.2434 |
| Pseudoazoarcus taiwanensis | 79.0943 | 79.4376 | 77.9386 | 79.2327 | 79.0361 | 79.2838 | 78.4837 | 78.3794 | 78.2835 | 78.7291 | 78.7606 | 78.6974 | 79.6588 | 78.8501 | 79.0287 | 79.6189 | 79.9811 | 79.9769 | 79.4196 | 79.5257 | 79.3364 | 79.26 | 79.0234 | 79.7399 |
| Aromatoleum bremense strain PbN1 | 82.3106 | 79.6793 | 77.9612 | 79.6821 | 79.6641 | 78.9211 | 78.4733 | 78.9009 | 78.5537 | 79.0471 | 79.219 | 79.1511 | 80.0127 | 80.2946 | 80.2266 | 80.1672 | 80.2323 | 80.506 | 80.696 | 80.7821 | 79.9731 | 79.209 | 79.7518 | 80.6052 |
| Aromatoleum aromaticum EbN1 | 82.2789 | 79.3296 | 77.9623 | 79.6046 | 79.5552 | 78.874 | 78.3086 | 78.6596 | 78.4745 | 78.8556 | 79.046 | 79.0597 | 80.0703 | 80.0369 | 80.009 | 80.3002 | 80.2654 | 80.2826 | 80.5523 | 80.6947 | 79.718 | 78.8359 | 79.5475 | 80.5406 |
| Aromatoleum buckelii | 82.232 | 79.3519 | 77.7706 | 79.614 | 79.467 | 78.4018 | 78.4506 | 78.6961 | 78.5099 | 78.8548 | 78.8398 | 78.8738 | 79.5938 | 79.9254 | 79.8896 | 79.5875 | 79.9015 | 80.1116 | 80.4693 | 80.7168 | 79.8207 | 78.7178 | 79.6888 | 80.01 |
| Aromatoleum diolicum | 83.1912 | 79.7904 | 78.0592 | 79.3638 | 79.7145 | 79.0299 | 78.8732 | 79.1923 | 78.7491 | 79.4578 | 79.1095 | 79.1454 | 80.3151 | 80.227 | 80.186 | 79.7898 | 79.8329 | 79.8224 | 80.1468 | 80.0442 | 79.9329 | 78.9527 | 79.5331 | 80.2569 |
| Azoarcus sp. KH32C | 83.2071 | 80.0422 | 78.003 | 79.4634 | 79.6059 | 78.9551 | 79.0487 | 79.0868 | 78.7462 | 79.2973 | 79.1892 | 79.1931 | 80.3 | 80.0601 | 80.1006 | 80.027 | 80.1701 | 80.2599 | 80.1399 | 80.1331 | 80.1928 | 78.9818 | 79.9136 | 80.0535 |
| Aromatoleum toluvorans | 87.9184 | 80.8242 | 78.4569 | 80.6549 | 80.5085 | 79.2056 | 79.0879 | 79.4388 | 79.1727 | 80.066 | 80.0496 | 80.0132 | 80.7482 | 80.9553 | 80.9917 | 80.9113 | 81.2592 | 81.5808 | 80.8943 | 81.0901 | 80.9232 | 79.6018 | 80.7048 | 81.05 |
| Aromatoleum toluolicum | 89.5767 | 80.5868 | 78.176 | 80.2923 | 80.5768 | 79.9627 | 79.0115 | 79.4089 | 79.1452 | 79.8126 | 79.9175 | 79.8774 | 80.7276 | 80.611 | 80.6976 | 80.7064 | 81.0486 | 81.1127 | 80.7017 | 80.6744 | 80.6608 | 79.6475 | 80.3328 | 80.952 |
| Azoarcus sp. Aa7 | 90.2071 | 80.5492 | 78.1644 | 80.3503 | 80.3134 | 78.9173 | 78.905 | 79.3763 | 79.0319 | 79.7275 | 79.6902 | 79.6999 | 80.7805 | 80.6057 | 80.5843 | 80.5848 | 80.8915 | 81.2026 | 80.9025 | 80.9284 | 80.7279 | 79.3647 | 80.3857 | 80.856 |
| Aromatoleum evansii | 89.4139 | 80.6134 | 77.9908 | 80.3262 | 80.5052 | 79.0377 | 79.1198 | 79.5337 | 79.1273 | 79.6349 | 79.7878 | 79.7664 | 80.6561 | 80.5782 | 80.5698 | 80.6902 | 81.076 | 81.2646 | 80.8871 | 80.9277 | 80.9185 | 79.4805 | 80.2872 | 80.8898 |
| Aromatoleum petrolei | 88.9356 | 80.3749 | 78.2587 | 80.381 | 80.2705 | 78.8563 | 78.7675 | 79.3294 | 78.8649 | 79.4568 | 79.3534 | 79.3092 | 80.4656 | 80.4774 | 80.5296 | 80.5718 | 80.7666 | 80.8561 | 80.511 | 80.709 | 80.5353 | 79.0902 | 80.12 | 80.5598 |
| Aromatoleum tolulyticum | 90.252 | 80.5411 | 78.1021 | 80.4338 | 80.3356 | 80.0066 | 78.8273 | 79.3852 | 79.1032 | 79.725 | 79.8903 | 79.8895 | 80.8305 | 80.6591 | 80.7458 | 80.8157 | 81.1752 | 81.3154 | 80.7649 | 80.9579 | 80.6723 | 79.5427 | 80.4416 | 80.9125 |
| Aromatoleum toluclasticum ATCC 700605 | 98.1284 | 80.5186 | 78.1951 | 80.0564 | 80.44 | 79.0389 | 78.9572 | 79.2325 | 78.9823 | 79.6308 | 79.6843 | 79.7288 | 80.7647 | 80.5279 | 80.546 | 80.5032 | 81.1658 | 81.1514 | 80.8471 | 80.7595 | 80.8516 | 79.5718 | 80.3959 | 80.813 |
| Aromatoleum toluclasticum TC-10 | 100 | 80.5615 | 78.1803 | 80.325 | 80.3483 | 79.1607 | 78.9854 | 79.4232 | 79.1331 | 79.7837 | 79.8041 | 79.6885 | 80.6416 | 80.5332 | 80.5191 | 80.5452 | 81.2324 | 81.1428 | 80.785 | 80.6988 | 80.6101 | 79.3855 | 80.3697 | 80.713 |
| Pseudothauera lacus sp. D20 | 80.505 | 100 | 80.8947 | 83.8638 | 84.0345 | 80.0174 | 79.77 | 80.1329 | 79.5427 | 81.0881 | 80.8954 | 80.8401 | 81.1977 | 81.3839 | 81.3043 | 81.4347 | 82.1366 | 82.4501 | 81.709 | 81.8924 | 81.5152 | 80.5061 | 81.0541 | 81.5991 |
| Pseudothauera hydrothermalis | 78.1612 | 81.093 | 100 | 80.7899 | 81.0103 | 78.5065 | 78.3715 | 78.4924 | 78.355 | 78.7926 | 78.7656 | 78.7851 | 79.07 | 79.3401 | 79.1907 | 78.8033 | 79.0724 | 79.5218 | 79.2324 | 79.3211 | 79.069 | 78.42 | 78.8203 | 79.0773 |
| Pseudothauera nasutitermitis | 80.2376 | 83.9617 | 80.8167 | 100 | 85.3747 | 79.6474 | 79.5463 | 79.6672 | 79.4005 | 81.1916 | 81.2123 | 81.309 | 81.276 | 81.6136 | 81.581 | 80.7921 | 81.753 | 82.0697 | 81.0143 | 81.3655 | 81.1673 | 80.1678 | 82.3975 | 82.216 |
| Pseudothauera rhizosphaerae | 80.4135 | 83.8303 | 81.0905 | 85.3591 | 100 | 79.6387 | 79.5973 | 79.7585 | 79.3307 | 81.5438 | 81.5759 | 81.5606 | 81.6083 | 81.7723 | 81.7728 | 81.1925 | 81.9289 | 82.3123 | 81.2997 | 81.6473 | 81.0355 | 80.0306 | 81.501 | 81.919 |
| Parazoarcus communis LMG 5514 | 79.2608 | 79.8506 | 78.5173 | 79.6583 | 79.3914 | 100 | 98.6462 | 80.4989 | 80.0549 | 79.7664 | 79.7416 | 79.7386 | 80.9866 | 80.2721 | 80.2162 | 79.6819 | 80.3028 | 80.5842 | 79.9149 | 80.263 | 80.0549 | 79.8775 | 79.8174 | 80.0865 |
| Parazoarcus communis SWub3 | 78.9293 | 79.8339 | 78.5994 | 79.5626 | 79.3958 | 98.6708 | 100 | 80.5212 | 80.511 | 79.7211 | 79.6261 | 79.6095 | 80.4345 | 80.5232 | 80.4858 | 79.5927 | 79.8379 | 79.947 | 79.8148 | 79.7935 | 80.2457 | 79.4425 | 79.7729 | 80.2272 |
| Parazoarcus communis | 79.6011 | 79.9802 | 78.3406 | 79.6341 | 79.7894 | 80.5385 | 80.499 | 100 | 83.344 | 79.6238 | 79.6919 | 79.6497 | 80.4989 | 80.6423 | 80.6189 | 80.0822 | 80.1738 | 80.3979 | 80.1509 | 80.2998 | 80.3621 | 79.5425 | 79.9856 | 80.1787 |
| Azoarcus sp. L1K30 | 79.2308 | 79.6372 | 78.2016 | 79.1908 | 79.251 | 80.0968 | 80.3075 | 83.2728 | 100 | 79.3236 | 79.2743 | 79.2598 | 80.1884 | 80.2267 | 80.1984 | 79.7324 | 79.7823 | 80.0936 | 79.843 | 79.9269 | 80.1521 | 79.2132 | 79.575 | 79.6757 |
| Azoarcus sp. TTM-91 | 79.7906 | 80.9952 | 78.768 | 81.1723 | 81.5797 | 79.8495 | 79.9166 | 79.6479 | 79.3717 | 100 | 93.9885 | 93.9476 | 82.7514 | 82.4628 | 82.4413 | 80.4993 | 81.0014 | 80.9779 | 80.6539 | 80.7607 | 80.8541 | 79.679 | 80.6101 | 80.9202 |
| Azoarcus indigens VB32 | 79.6914 | 80.8929 | 78.903 | 81.2052 | 81.4803 | 79.7361 | 79.799 | 79.526 | 79.441 | 93.9075 | 100 | 99.9924 | 82.8405 | 82.4849 | 82.4358 | 80.3187 | 80.9008 | 80.9796 | 80.5381 | 80.6543 | 80.8646 | 79.4893 | 80.7622 | 81.147 |
| Azoarcus indigens | 79.629 | 80.8998 | 78.8007 | 81.218 | 81.6347 | 79.5838 | 79.6761 | 79.6211 | 79.3204 | 93.8945 | 99.9807 | 100 | 82.8771 | 82.5433 | 82.464 | 80.4631 | 80.8034 | 80.9588 | 80.5384 | 80.6249 | 80.9442 | 79.5768 | 80.7458 | 81.1506 |
| Azoarcus sp. DD4 | 80.771 | 81.2771 | 79.1768 | 81.3464 | 81.6277 | 80.9165 | 80.4156 | 80.7091 | 80.22 | 82.8467 | 82.8539 | 82.8606 | 100 | 86.4373 | 86.4099 | 81.482 | 82.0421 | 82.3341 | 81.7346 | 82.0321 | 81.6916 | 80.938 | 81.496 | 82.1426 |
| Azoarcus olearius BH72 | 80.4527 | 81.4394 | 79.3192 | 81.3925 | 81.8339 | 80.2265 | 80.3827 | 80.4699 | 80.1992 | 82.6316 | 82.7318 | 82.5565 | 86.4463 | 100 | 98.7536 | 81.4368 | 81.901 | 82.1664 | 81.5146 | 81.7866 | 81.8618 | 80.1195 | 81.525 | 81.7146 |
| Azoarcus olearius DQS-4 | 80.6746 | 81.3208 | 79.2237 | 81.3931 | 81.6575 | 80.3486 | 80.4862 | 80.5334 | 80.1627 | 82.531 | 82.5945 | 82.5916 | 86.3509 | 98.768 | 100 | 81.5134 | 81.8893 | 82.2296 | 81.5836 | 81.6817 | 81.828 | 80.1062 | 81.4586 | 81.5353 |
| Thauera phenolivorans | 80.4678 | 81.4045 | 78.7831 | 80.9995 | 81.3183 | 79.8693 | 79.6344 | 80.1238 | 79.6825 | 80.5325 | 80.4524 | 80.3521 | 81.5239 | 81.5153 | 81.628 | 100 | 84.0836 | 84.8395 | 83.1929 | 83.5008 | 82.7517 | 81.1813 | 82.1863 | 82.865 |
| Thauera aminoaromatica S2 | 80.814 | 81.9527 | 79.0844 | 81.7077 | 81.7452 | 80.4424 | 79.7804 | 80.164 | 79.7717 | 81.1023 | 81.046 | 80.915 | 82.0257 | 81.8951 | 82.0162 | 83.9407 | 100 | 92.0408 | 84.4276 | 84.6995 | 84.4562 | 82.4682 | 83.4839 | 84.0926 |
| Thauera phenylacetica | 81.0646 | 82.1758 | 79.3002 | 81.9801 | 82.1864 | 80.2582 | 79.9062 | 80.45 | 79.8574 | 80.9203 | 80.8297 | 80.8704 | 82.2646 | 82.0446 | 82.151 | 84.5822 | 91.9008 | 100 | 84.6895 | 85.131 | 84.5304 | 82.9673 | 83.5244 | 84.1439 |
| Thauera aromatica K172 | 80.914 | 81.7364 | 79.119 | 81.1579 | 81.376 | 80.137 | 79.9106 | 80.0064 | 79.8984 | 80.6583 | 80.6935 | 80.7366 | 81.8906 | 81.7062 | 81.5983 | 83.2184 | 84.3439 | 84.9195 | 100 | 93.3578 | 83.3575 | 81.6324 | 82.7846 | 83.2539 |
| Thauera chlorobenzoica | 80.8621 | 81.8885 | 79.3668 | 81.4791 | 81.4398 | 80.1065 | 79.7055 | 80.322 | 79.9069 | 80.8849 | 80.7262 | 80.6938 | 82.0076 | 81.9495 | 81.914 | 83.4295 | 84.7191 | 85.2531 | 93.3297 | 100 | 83.4454 | 82.2156 | 82.912 | 83.685 |
| Thauera propionica | 80.7075 | 81.4059 | 78.9625 | 81.2573 | 81.1455 | 80.1873 | 80.2216 | 80.4256 | 80.0637 | 80.8829 | 80.7992 | 80.875 | 81.7446 | 81.9661 | 81.9314 | 82.6574 | 84.5369 | 84.6506 | 83.2468 | 83.1632 | 100 | 82.6667 | 84.7005 | 84.7644 |
| Thauera terpenica | 79.5542 | 80.4269 | 78.8395 | 79.9879 | 80.0242 | 79.9523 | 79.4732 | 79.735 | 79.307 | 79.5592 | 79.658 | 79.645 | 80.9852 | 80.1457 | 80.2745 | 81.4455 | 82.7198 | 83.3819 | 81.3967 | 82.2352 | 82.9176 | 100 | 82.4934 | 82.8594 |
| Thauera linaloolentis | 80.4098 | 81.0512 | 78.7288 | 82.3889 | 81.5746 | 79.7765 | 79.8822 | 80.2101 | 79.5925 | 80.6997 | 80.9204 | 80.8953 | 81.4313 | 81.4729 | 81.5134 | 82.0909 | 83.5758 | 83.8044 | 82.736 | 82.8082 | 84.7201 | 82.458 | 100 | 88.3879 |
| Thauera butanivorans | 80.8474 | 81.4589 | 79.1101 | 82.1544 | 82.0167 | 80.2632 | 80.4069 | 80.2817 | 79.8242 | 81.0896 | 81.287 | 81.1922 | 81.8791 | 81.8601 | 81.7143 | 82.6452 | 83.9171 | 84.214 | 83.2197 | 83.4761 | 84.7322 | 82.6351 | 88.2937 | 100 |

### Table S7: Jaccard similarity index scores

[illegible]
